## Supplementary Figures for "Loss of TET proteins in regulatory T cells unleashes effector function"

### Supplementary Figure 1

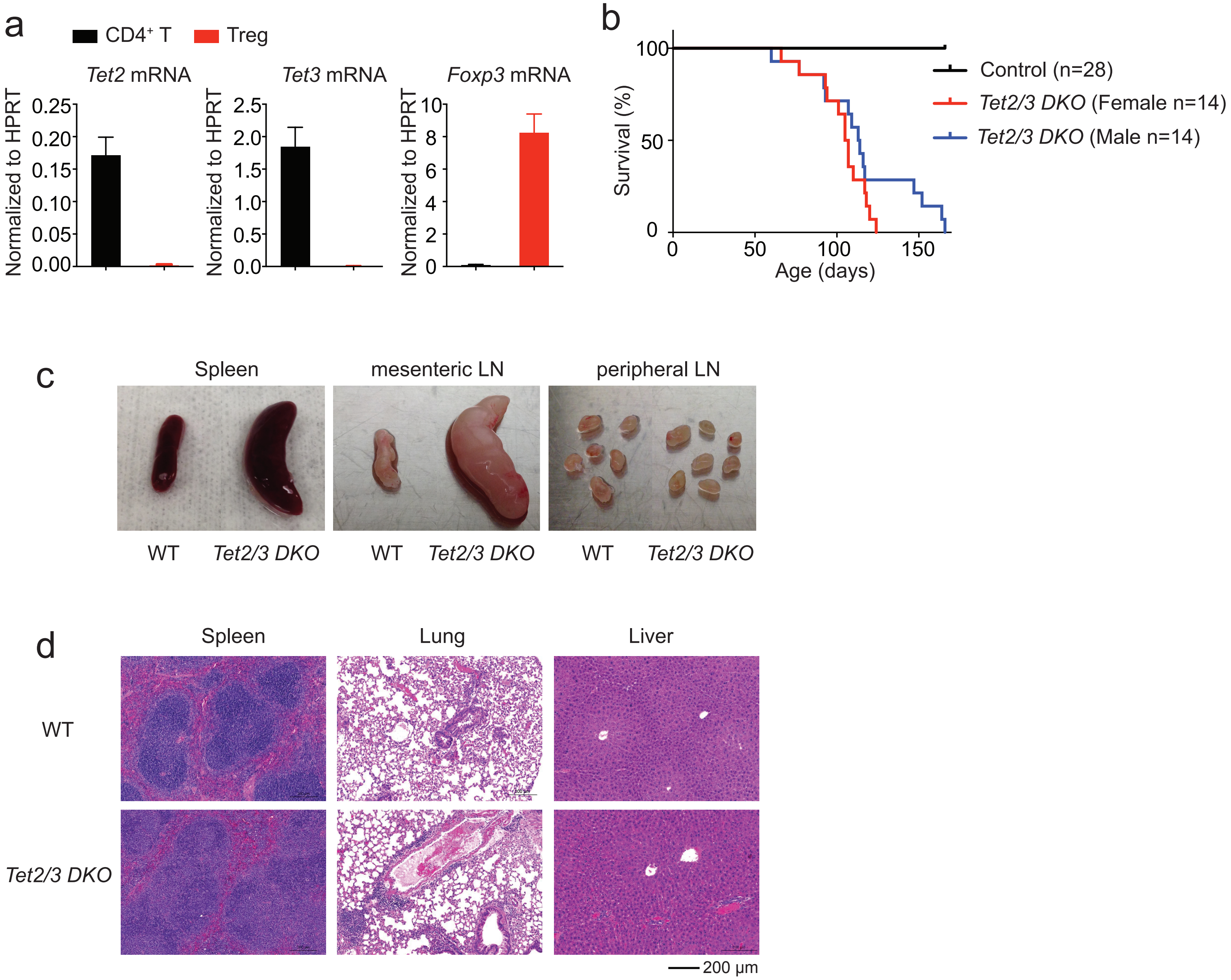


**Supplementary Figure 1. Characterization of mice with Treg-specific deficiency of *Tet2* and *Tet3*.**

**a**. Quantitative real-time PCR analysis of *Tet2*, *Tet3* and *Foxp3* expression level in $CD4^+Foxp3^-$ T cells ($CD4^+$ T) and $CD4^+Foxp3^+$ Treg cells (Treg) isolated from $Tet2/3^{fl/fl}Foxp3^{Cre}$ DKO mice (13-16 weeks old). **b**. Survival curves for control WT (n=28) and $Tet2/3^{fl/fl}Foxp3^{Cre}$ DKO mice separated into female and male groups (n=14 for each group). **c**. Representative pictures of spleen, mesenteric lymph nodes and peripheral lymph nodes (pLN) from WT and $Tet2/3^{fl/fl}Foxp3^{Cre}$ DKO mice (14 weeks old). **d**. H&E staining of spleen, lung and liver of WT and $Tet2/3^{fl/fl}Foxp3^{Cre}$ DKO mice (15-16 weeks old).

Supplementary Figure 2

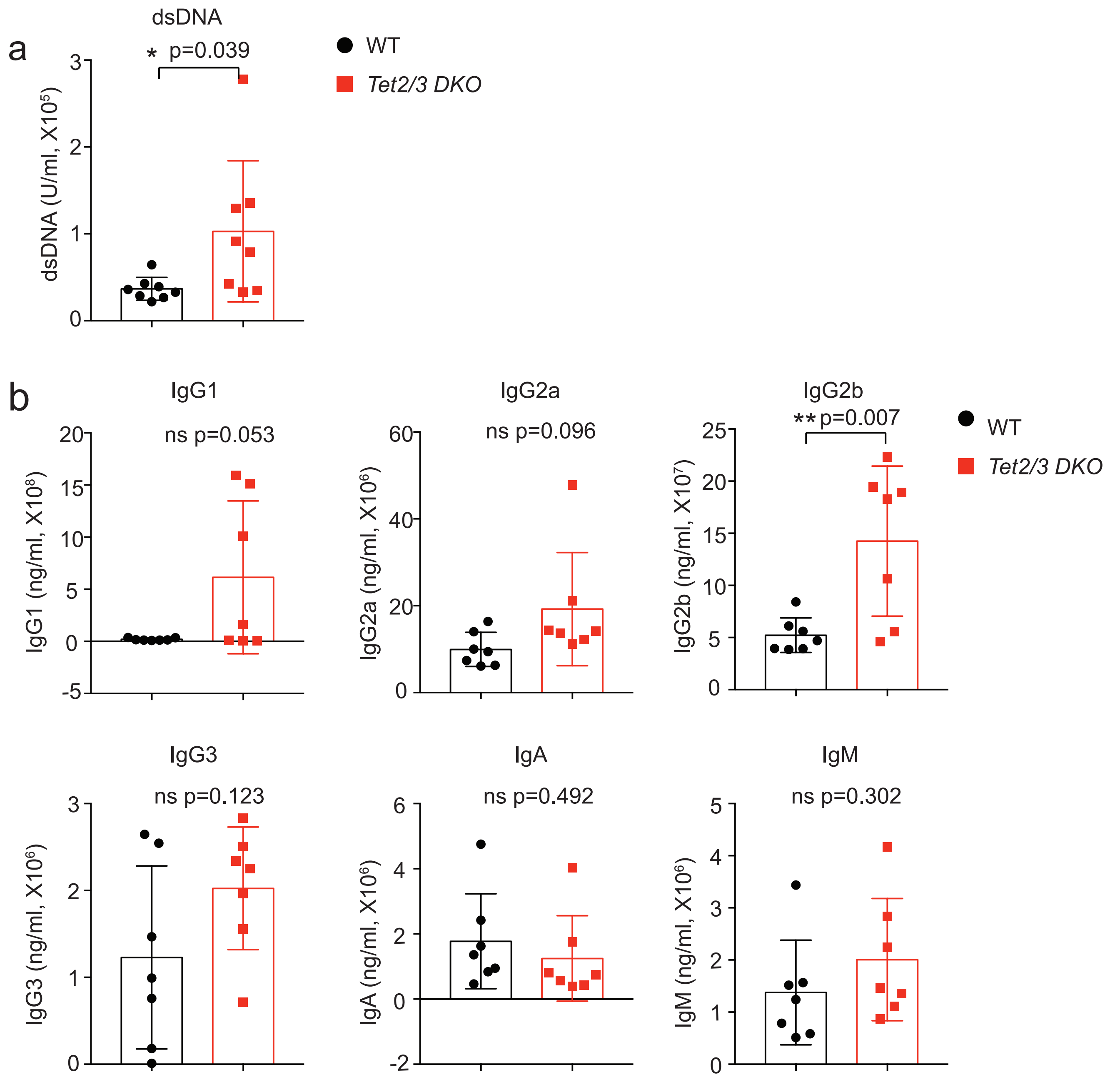


**Supplementary Figure 2. Analysis of anti-dsDNA antibody and immunoglobulin levels in the serum isolated from *Tet2/3$^{fl/fl}$Foxp3$^{Cre}$* DKO mice.**

**a**. Quantification of anti-dsDNA antibody in the serum of WT and *Tet2/3$^{fl/fl}$Foxp3$^{Cre}$* DKO mice (11-16 weeks old, WT n=8, *Tet2/3$^{fl/fl}$Foxp3$^{Cre}$* DKO mice n=7). **b**. Quantification of immunoglobulin IgG1, IgG2a, IgG2b, IgG3, IgA and IgM in the serum of WT and *Tet2/3$^{fl/fl}$Foxp3$^{Cre}$* DKO mice (11-16 weeks old, WT n=7, *Tet2/3$^{fl/fl}$Foxp3$^{Cre}$* DKO mice n=7). Data are mean ± S.D. from two independent experiments. Statistical analysis was performed using two-tailed unpaired student's t test (*P<0.05, **P<0.01).

Supplementary Figure 3

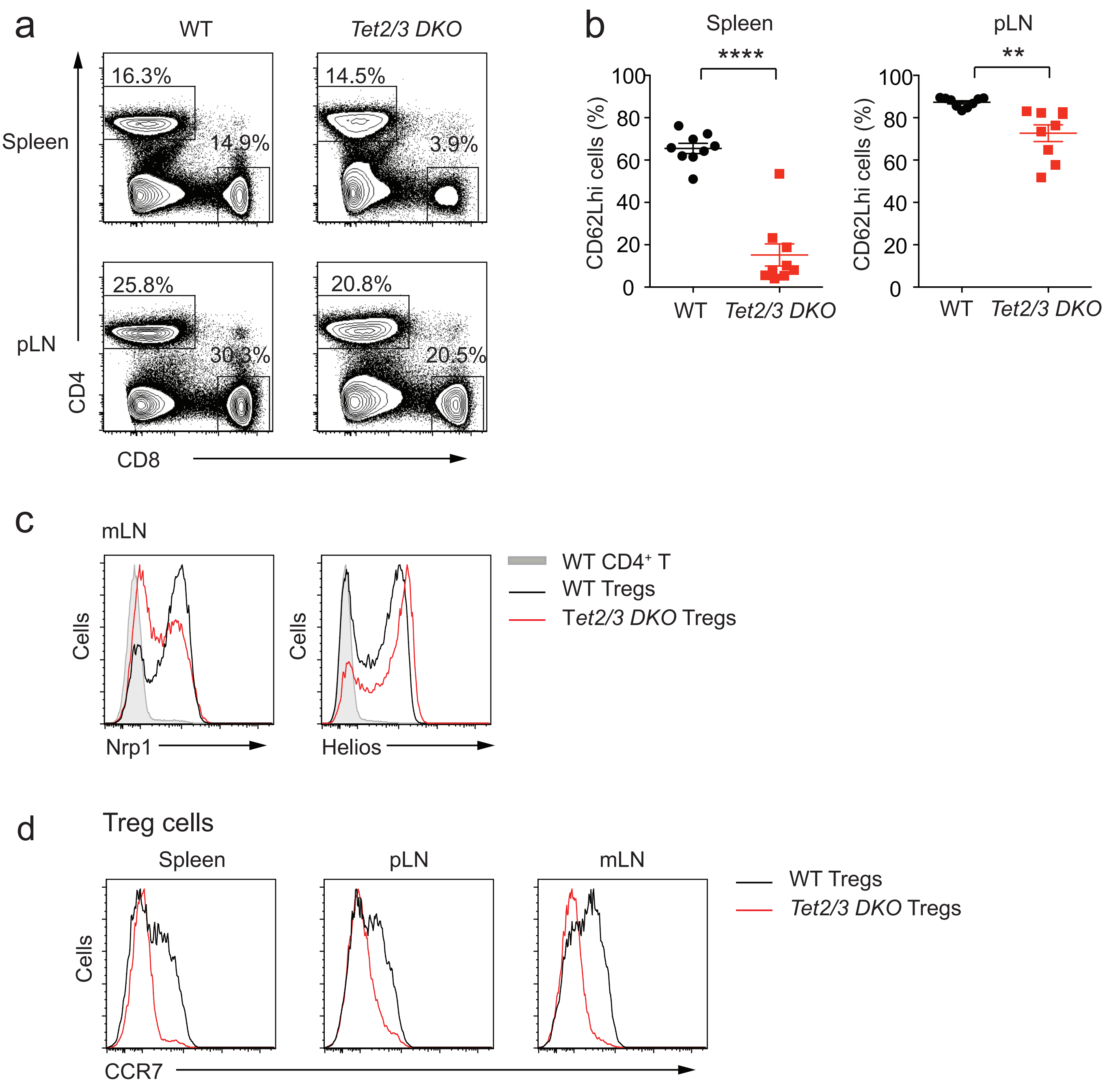


**Supplementary Figure 3. Characterization of $CD4^+$ and $CD8^+$ T cell compartments and Treg cells in *Tet2/3$^{fl/fl}$Foxp3$^{Cre}$* DKO mice.**

**a**. Representative flow cytometry analysis of CD4 and CD8 expression in spleen (*upper panels*) and pLN (*lower panels*) from WT and *Tet2/3$^{fl/fl}$Foxp3$^{Cre}$* DKO mice (13-16 weeks old). **b**. Quantification of the percentage of CD62L$^{high}$ cells in spleen and pLN from WT and *Tet2/3$^{fl/fl}$Foxp3$^{Cre}$* DKO mice (13-16 weeks old, see flow cytometry plots of Figure 1D). Statistical analysis was performed using two-tailed unpaired student's t test (**P<0.01, ***P<0.001). **c**. Flow cytometry analysis of *Tet2/3*-deficient Treg cells (13-16 weeks old) from mLN for the expression of Nrp1 and Helios. Shown are WT $CD4^+$ T cells (shaded grey); WT Tregs (black line); DKO Tregs (red line). **d**. Flow cytometry analysis of *Tet2/3*-deficient Treg cells (13-16 weeks old) from spleen (*left*), pLN (*middle*) and mLN (*right*) for the expression of CCR7. Shown are WT Tregs (black line); DKO Tregs (red line).

Supplementary Figure 4

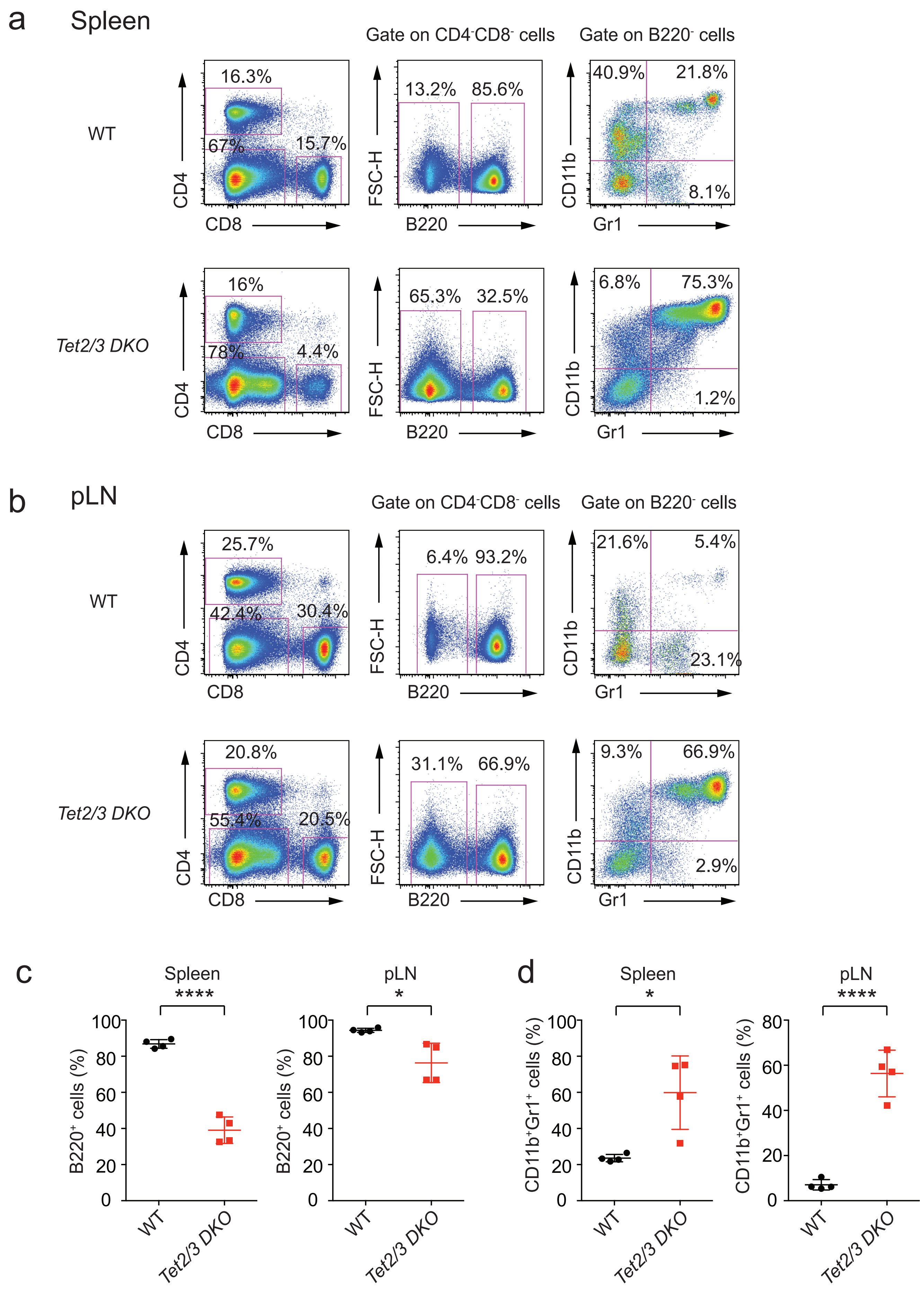

**Supplementary Figure 4. Analysis of B cell and Myeloid cell lineages in *Tet2/3$^{fl/fl}$Foxp3$^{Cre}$ DKO* mice.**

**a-b**. Flow cytometry staining of cells from spleen (**a**) and peripheral lymph nodes (pLN) (**b**) isolated from WT (*upper panels*) and *Tet2/3$^{fl/fl}$Foxp3$^{Cre}$ DKO* mice (*lower panels*) for the cell surface markers of CD4 and CD8 (*left panels*), B220 (*middle panels*, gated on $CD4^-CD8^-$ cells), CD11b and Gr1 (*right panels*, gated on $CD4^-CD8^-B220^-$ cells). **c-d**. Quantification of the percentage of $B220^+$ cells (**c**) and $CD11b^+Gr1^+$ cells (**d**) in spleen (*left panels*) and pLN (*right panels*) isolated from WT and *Tet2/3$^{fl/fl}$Foxp3$^{Cre}$ DKO* mice. Data are representative of four independent experiments. Statistical analysis was performed using two-tailed unpaired student's t test (*P<0.05, ****P<0.0001).

### Supplementary Figure 5

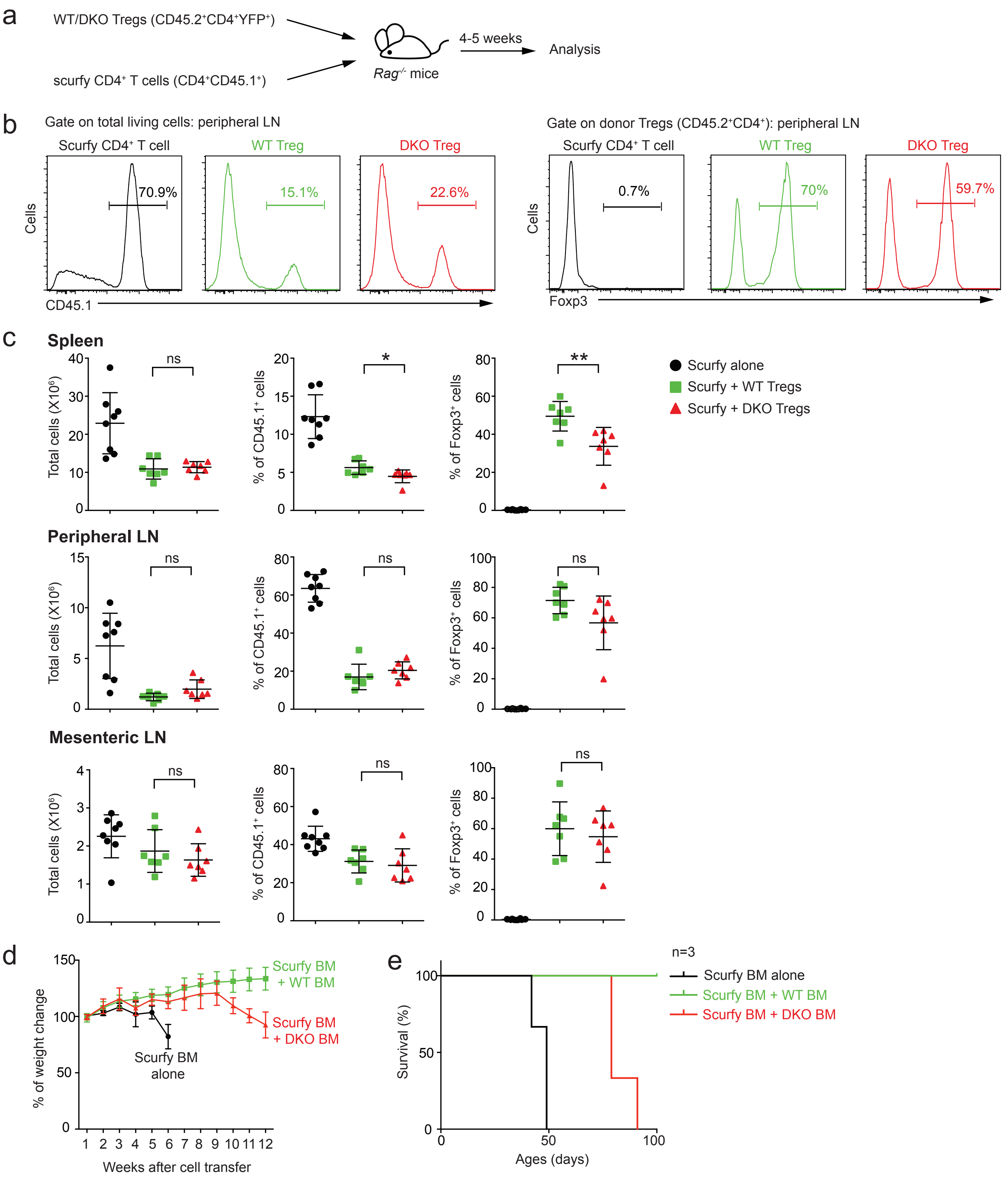

**Supplementary Figure 5. *Tet2/3 DKO* Treg cells can correct the *scurfy* phenotype in short term but not long-term assays.**

**a**. Schematic representation of the adoptive transfer experiment to assess the stability and suppressive function of Treg cells from 11-14 week-old *Tet2/3 DKO* mice *in vivo*. **b**. Representative histograms of CD45.1$^+$ cells (*left panel*) and CD45.2$^+$CD4$^+$Foxp3$^+$ cells (*right panel*) in peripheral lymph nodes from *Rag*-deficient mice 4-5 weeks after adoptive transfer of *scurfy* CD4$^+$ T cells alone or together with WT or DKO Treg cells. **c**. Graphs quantifying the total cellularity, percentage of CD45.1$^+$ cells and percentage of Foxp3$^+$ cells in transferred Treg cells from *Rag*-deficient mice, 4-5 weeks after adoptive transfer of scurfy CD4$^+$ T cells alone or together with WT or DKO Treg cells. WT and DKO Tregs show an equivalent ability to suppress the uncontrolled expansion of scurfy T cells. **d-e**. The percent weight change (**d**) and survival curves (**e**) after adoptive transfer of scurfy bone marrow cells, alone or together with WT or DKO bone marrow cells, into sub-lethally irradiated *Rag*-deficient mice (n=3). Statistical analysis was performed using two-tailed unpaired student's t test (*P<0.05, **P<0.01).

Supplementary Figure 6

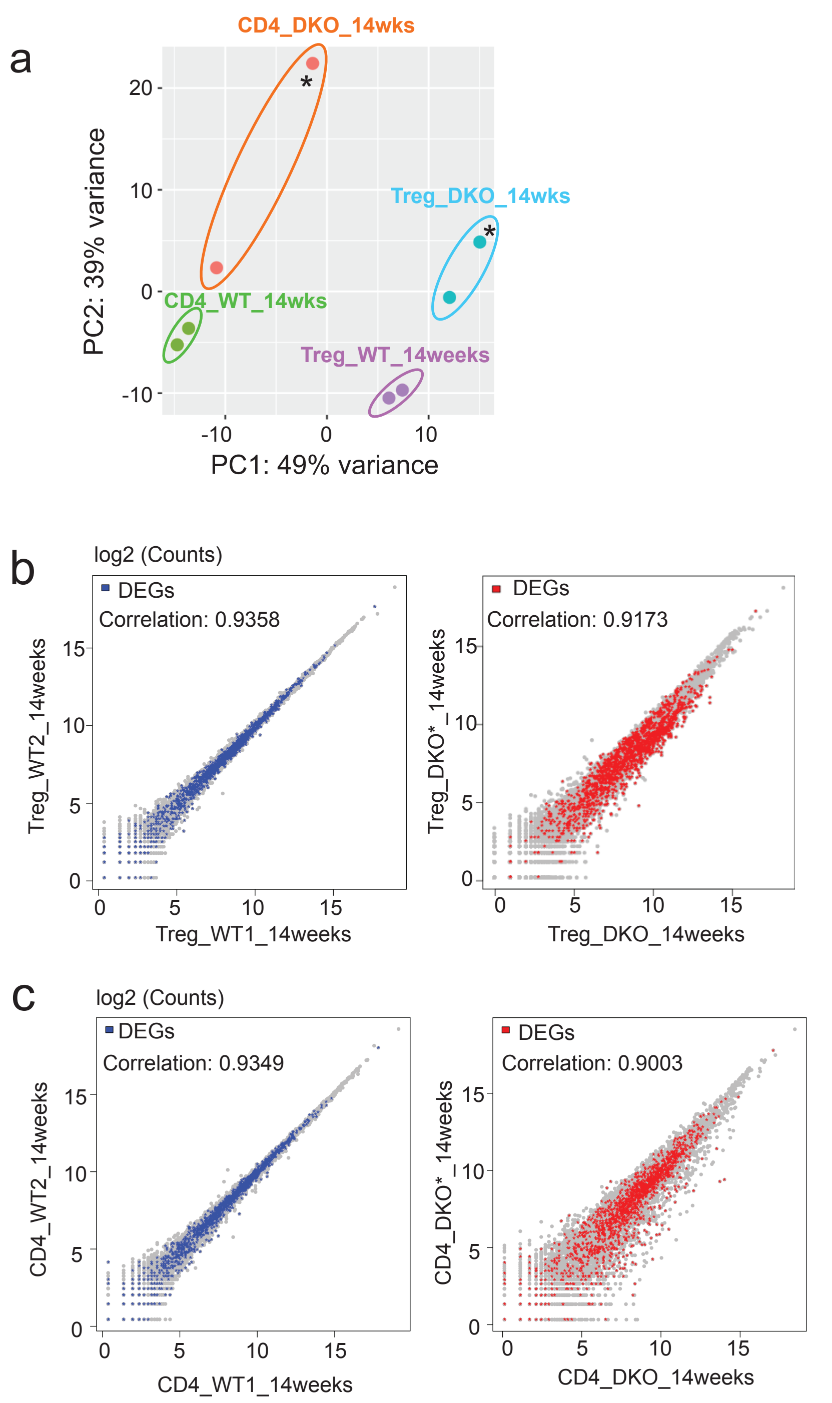


**Supplementary Figure 6. PCA and correlation analysis for RNA-seq replicates.**

**a**. Principal component analysis (PCA) plot for replicate RNA-seq samples: comparison of WT and DKO Tregs and WT and DKO CD4$^+$Foxp3$^-$ T cells from 14-weeks old mice. **b-c**. Scatter plots showing the correlation between replicates for Treg cells (**b**) and CD4$^+$Foxp3$^-$ T cells (**c**) from 14-weeks old WT and *Tet2/3*$^{fl/fl}$*Foxp3*$^{Cre}$ DKO mice.

Supplementary Figure 7

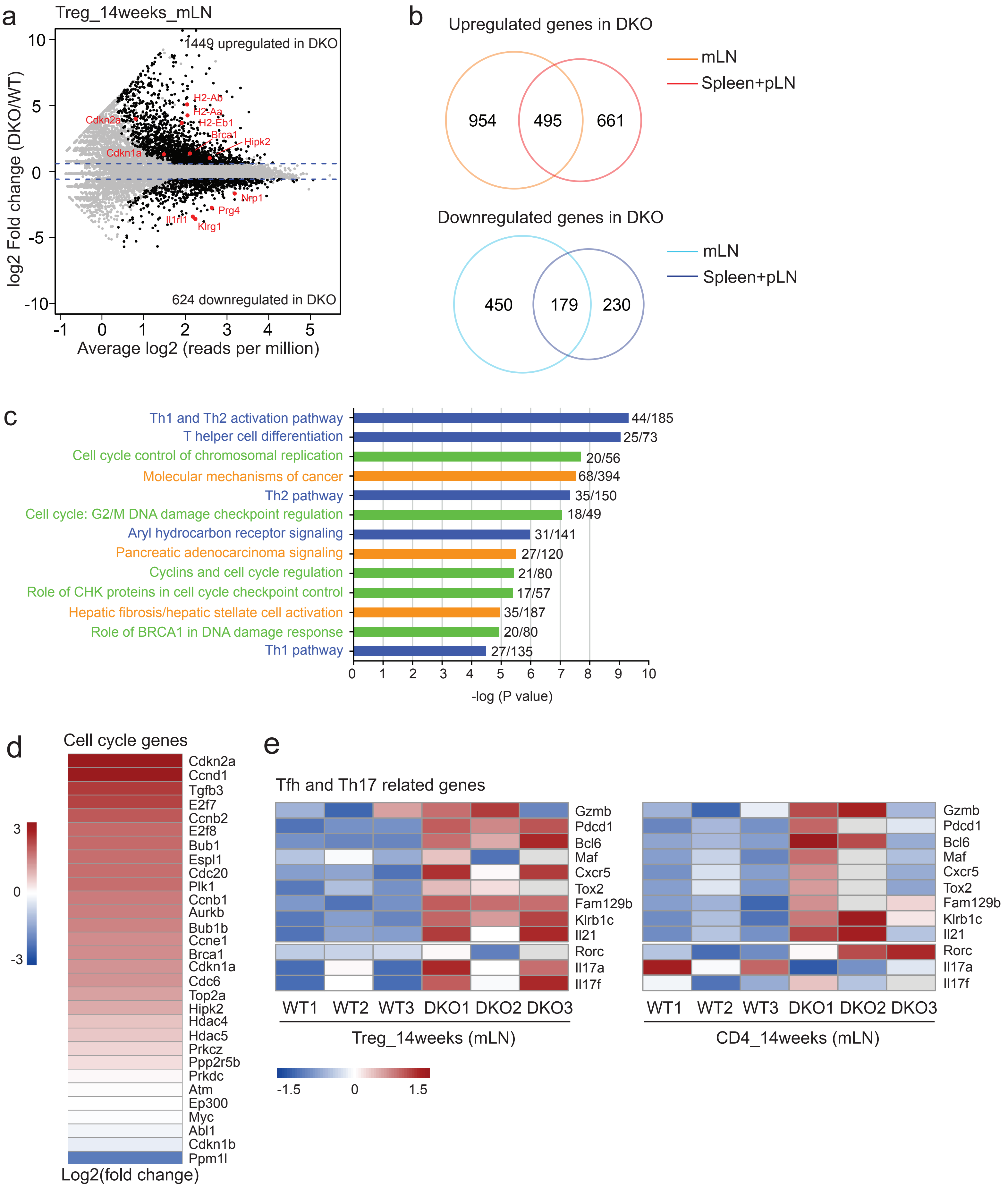

**Supplementary Figure 7. RNA-seq analysis for Treg cells isolated from mLN of WT or *Tet2/3$^{fl/fl}$Foxp3$^{Cre}$* *DKO* mice.**

**a**. Mean average (MA) plot of genes differentially expressed in *Tet2/3 DKO* Tregs (14-weeks old, isolated from mLN) relative to their expression in WT Treg cells. **b**. Overlap of differentially expressed genes that are upregulated (upper panel) or downregulated (lower panel) in *Tet2/3* DKO Treg cells isolated from either pooled spleen and pLNs or from mLN. **c**. IPA analysis of canonical pathways for differentially expressed genes in *Tet2/3 DKO* Treg cells isolated from mLN. Green, categories related to DNA repair, DNA damage and cell cycle; blue, categories related to immune cell function; orange, category related to cancer. **d**. Heatmap for the expression of selected genes encoding cell cycle regulators. The color gradient indicates Log2 Fold Change (DKO/WT). **e**. Heatmaps showing expression (row z score of log2 TPM values) of Tfh and Th17 related genes in WT and DKO Tregs (*left panel*) and WT and DKO $CD4^{+}Foxp3^{-}$ cells (*right panel*).

### Supplementary Figure 8

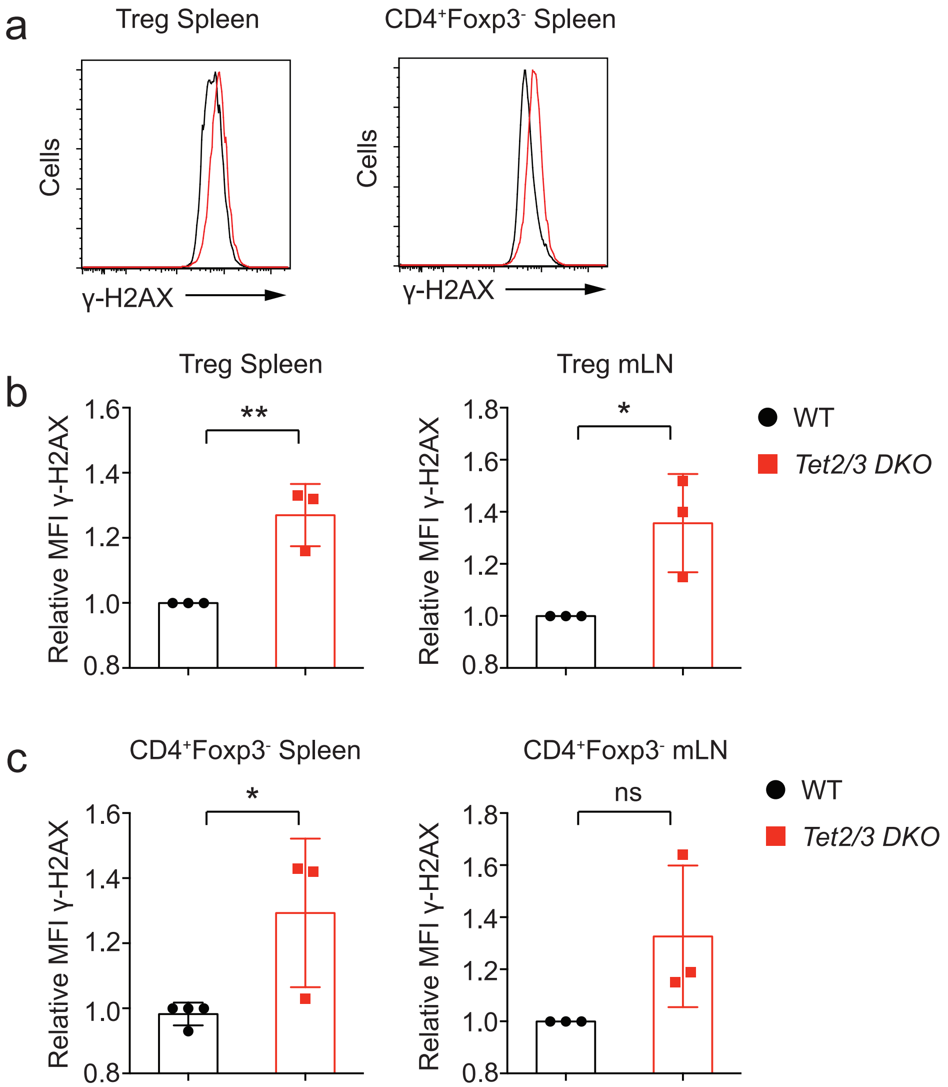


**Supplementary Figure 8. *Tet2/3* DKO cells show increased level of DNA damage.**

**a**. Representative flow cytometry analysis of γ-H2AX in Treg cells (*left panel*) and CD4$^+$Foxp3$^-$ T cells (*right panel*) in spleen isolated from WT and *Tet2/3*$^{fl/fl}$*Foxp3*$^{Cre}$ DKO mice (13-16 weeks old) **b**. Relative MFI (mean fluorescent intensity) of γ-H2AX in Treg cells from WT and *Tet2/3*$^{fl/fl}$*Foxp3*$^{Cre}$ DKO mice (13-16 weeks old) in Spleen (*left panel*) and mLN (*right panel*). **c**. Relative MFI of γ-H2AX in CD4$^+$Foxp3$^-$ cells from WT and *Tet2/3*$^{fl/fl}$*Foxp3*$^{Cre}$ DKO mice (13-16 weeks old) in Spleen (*left panel*) and mLN (*right panel*). Data are mean ± S.D. from three independent experiments. Statistical analysis was performed using two-tailed unpaired student's t test (*P<0.05, **P<0.01, ns: not significant).

Supplementary Figure 9

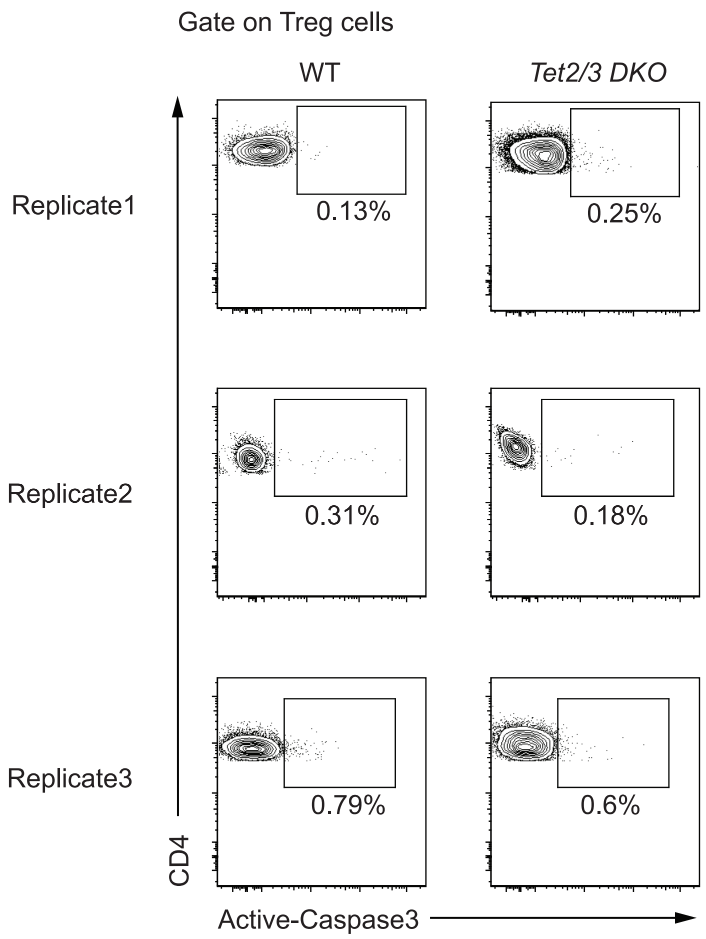


**Supplementary Figure 9. Caspase 3 staining to evaluate the apoptosis in *Tet2/3* DKO Treg cells.**

Flow cytometry staining of active caspase-3 in WT and *Tet2/3* DKO Treg cells using PE-conjugated anti-active Caspase-3 antibody. Data are from three independent experiments.

Supplementary Figure 10

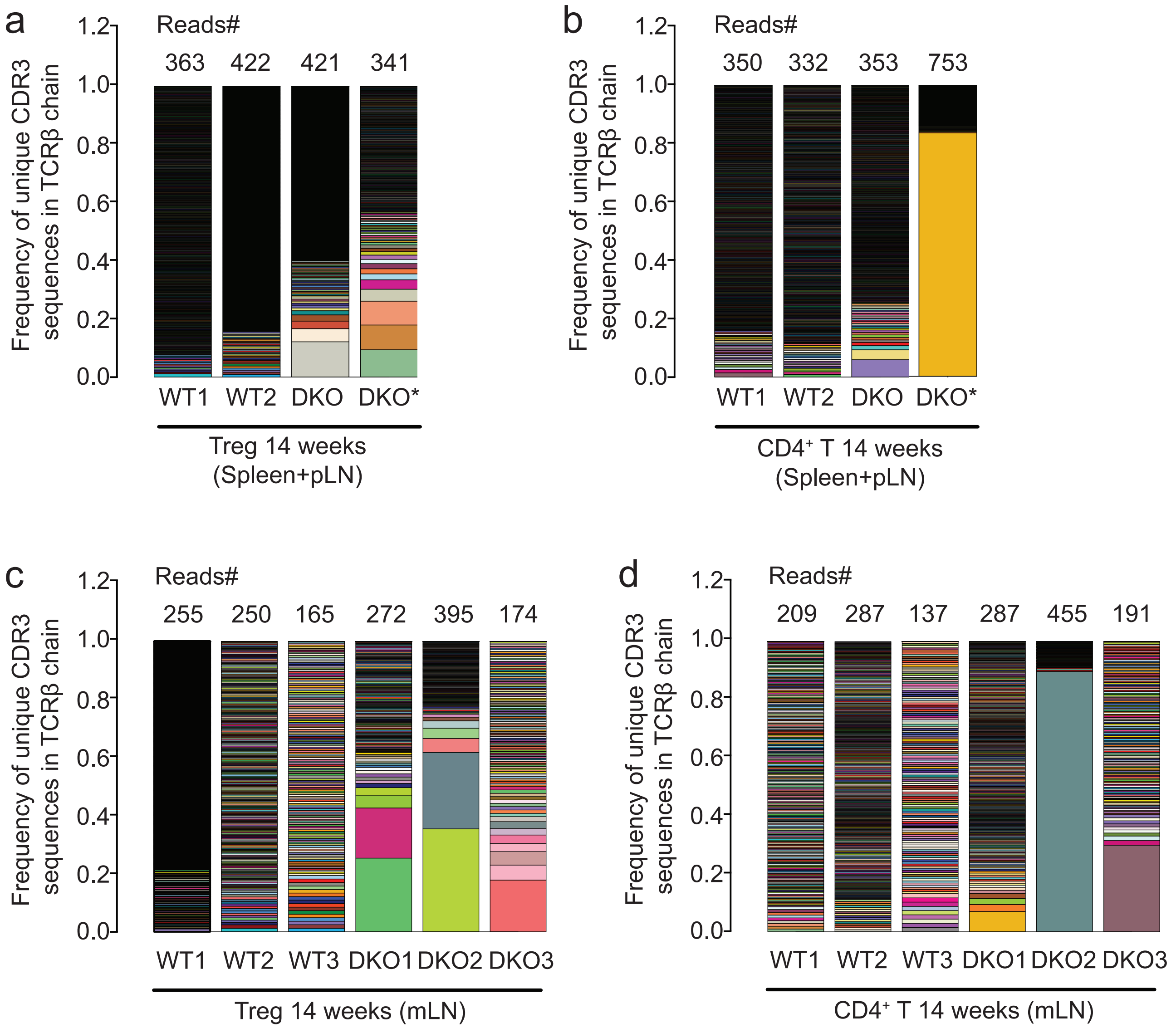


**Supplementary Figure 10. *Tet2/3* DKO cells show increased T cell clonal expansion.**

**a-b**. The frequency of reads (from the RNA-seq data of pooled spleen and pLNs) mapping to unique CDR3 sequences in TCRβ chain in WT and DKO Tregs from 14-week-old WT and *Tet2/3*$^{fl/fl}$*Foxp3*$^{Cre}$ DKO mice (**a**), in WT and DKO CD4$^{+}$Foxp3$^{-}$ T cells from 14-week-old WT and *Tet2/3*$^{fl/fl}$*Foxp3*$^{Cre}$ DKO mice (**b**). **c-d**. The frequency of reads (from the RNA-seq data of mLN) mapping to unique CDR3 sequences in TCRβ chain in WT and DKO Tregs from 14-week-old WT and *Tet2/3*$^{fl/fl}$*Foxp3*$^{Cre}$ DKO mice (**c**), in WT and DKO CD4$^{+}$Foxp3$^{-}$ T cells from 14-week-old WT and *Tet2/3*$^{fl/fl}$*Foxp3*$^{Cre}$ DKO mice (**d**). Each color represents a different TCRβ CDR3 sequence; the number of reads is shown on top of each graph.

Supplementary Figure 11

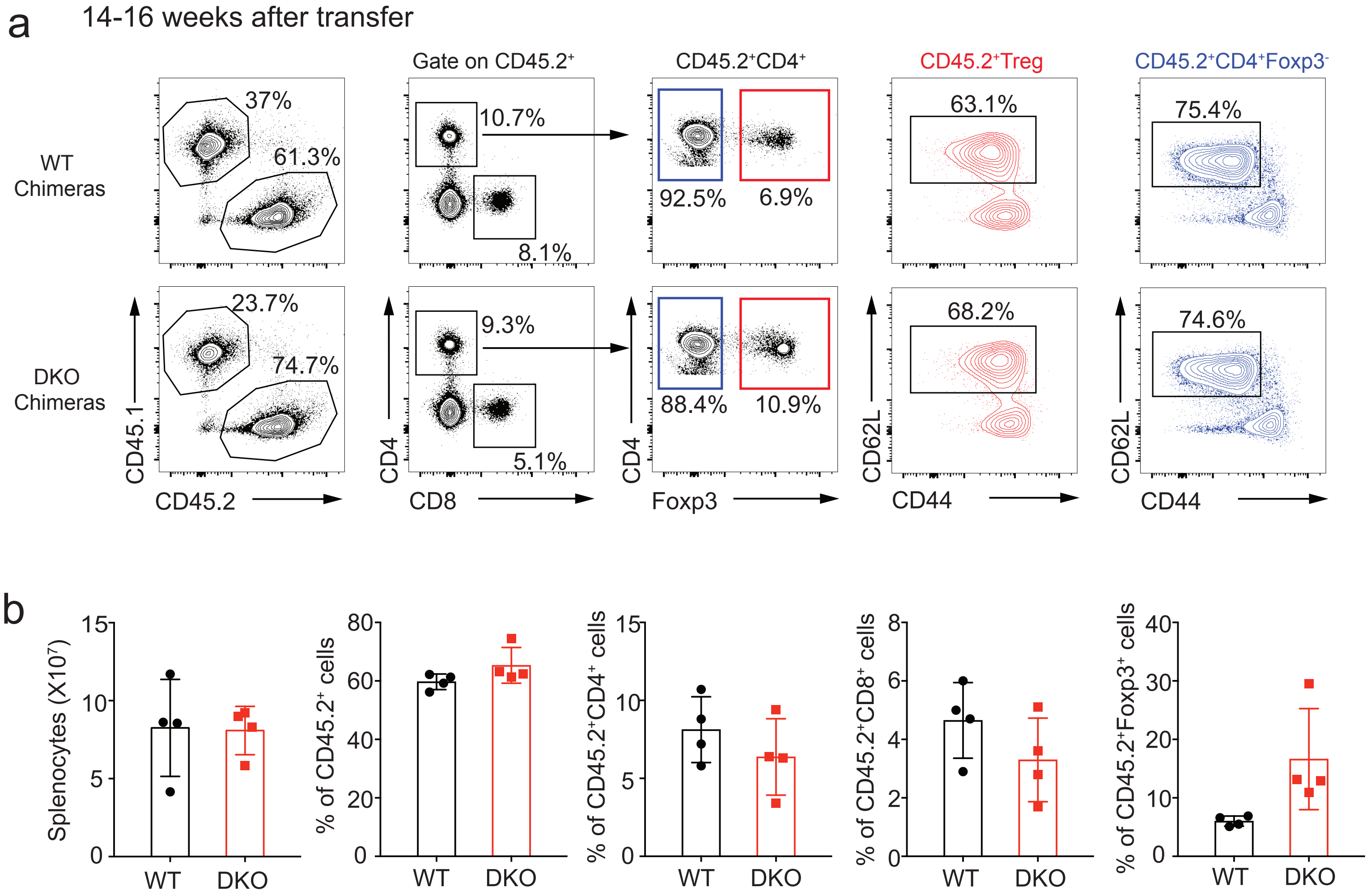


**Supplementary Figure 11. Analyses of bone marrow chimeras at earlier time points, prior to splenomegaly.**

**a**. Representative flow cytometry analysis for WT and DKO mixed bone marrow chimeras 14-16 weeks after transfer, before the development of splenomegaly. **b**. From left to right, the graphs show quantifications for the total number of splenocytes, the percentage of $CD45.2^+$ cells, the percentage of $CD4^+$ and $CD8^+$ cells within the $CD45.2^+$ cells and the percentage of $Foxp3^+$ cells within the $CD45.2^+CD4^+$ cells (n=4).

Supplementary Figure 12

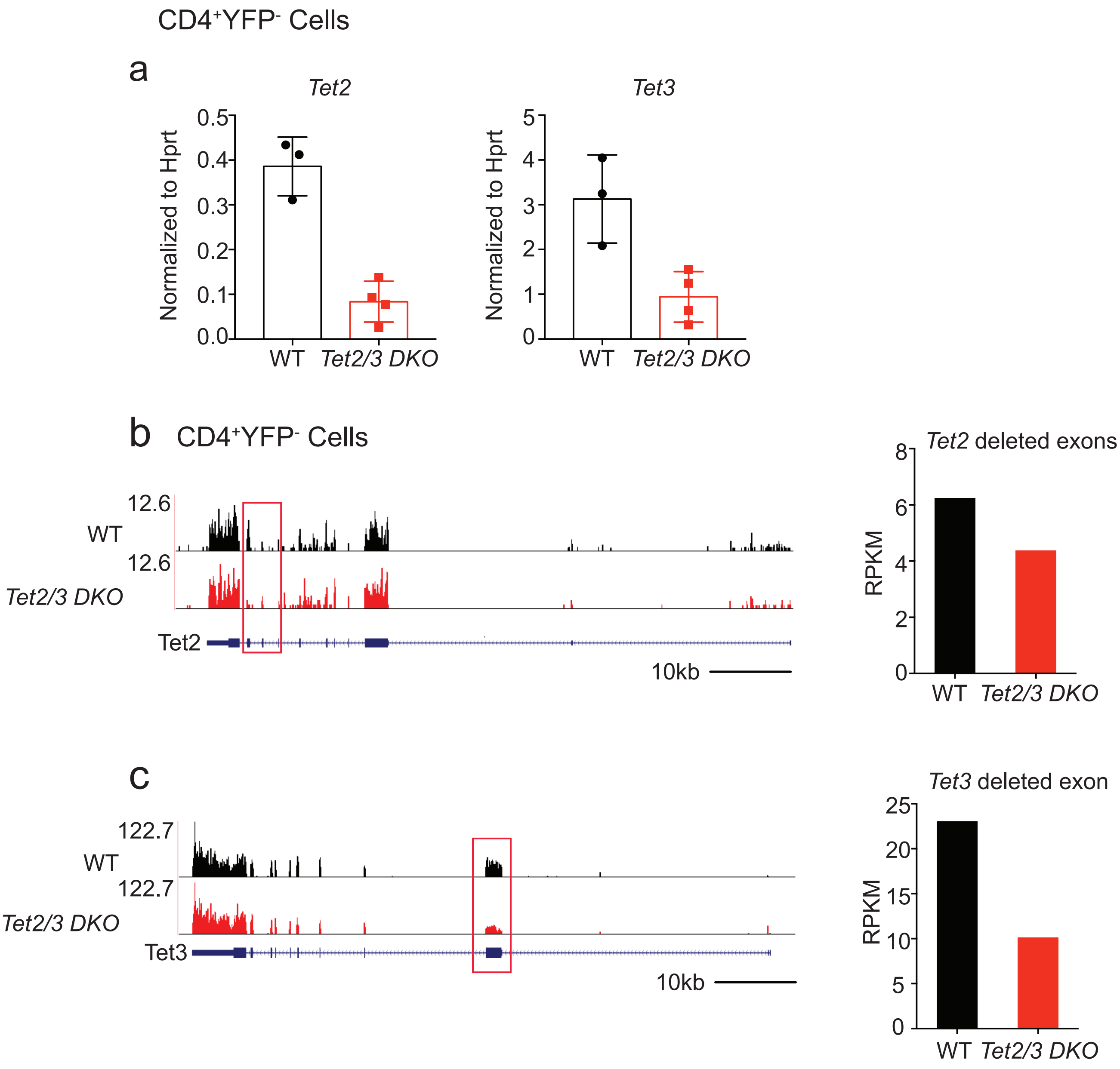


**Supplementary Figure 12. Analyses of *Tet2* and *Tet3* expression at mRNA level in CD4$^+$YFP$^-$ T cells from *Tet2/3$^{fl/fl}$Foxp3$^{Cre}$* DKO mice.**

**a**. Quantitative real-time PCR analysis of *Tet2* and *Tet3* expression in CD4$^+$YFP$^-$ cells from WT and *Tet2/3$^{fl/fl}$Foxp3$^{Cre}$* DKO mice (13-16 weeks old). **b-c**. Genome browser tracks of *Tet2* and *Tet3* with the deleted exons highlighted in rectangles (*left panels*) and the number of normalized reads within the deleted exons (*right panels*).
