## Supplementary Tables for "Loss of TET proteins in regulatory T cells unleashes effector function"

#### Supplementary Table1. Hematological parameters in *Tet2/3* DKO mice

| | WT | DKO | P value |
|---|---|---|---|
| WBC ($10^3$/µl) | 9.66±1.62 | 8.66±0.95 | 0.3298 |
| Neutrophil (103/ul) | 1.91±0.30 | 2.98±0.84 | 0.0525 |
| Lymphocyte (103/ul) | 7.47±1.30 | 5.29±1.04 | *, 0.0401 |
| Monocyte (103/ul) | 0.20±0.06 | 0.26±0.06 | 0.2248 |
| Eosinophil (103/ul) | 0.07±0.02 | 0.10±0.05 | 0.2508 |
| Basophil (103/ul) | 0.02±0.01 | 0.03±0.01 | 0.3202 |
| RBC (106/ul) | 9.75±0.25 | 9.48±0.33 | 0.2332 |
| Hemoglobin (g/dL) | 12.93±0.26 | 12.48±0.81 | 0.3291 |
| Hematocrit (%) | 46.05±0.91 | 44.50±1.82 | 0.1783 |
| Platelet (103/ul) | 1043.75±85.18 | 782±214.82 | 0.0641 |

#### Supplementary Table 2. Altered Treg signature genes in 14 week-old Tet2/3 DKO Tregs

**Pooled spleen and pLNs**

| Gene name | Log2 fold change | P value | Ajusted P value |
|---|---|---|---|
| Myo10 | 2.768 | 0.002 | 0.017 |
| Nrn1 | 2.639 | 0.000 | 0.000 |
| Plcb4 | 2.580 | 0.000 | 0.000 |
| Nr4a2 | 2.498 | 0.000 | 0.002 |
| Ntrk3 | 2.145 | 0.000 | 0.000 |
| Abhd15 | 2.118 | 0.000 | 0.001 |
| Hipk2 | 2.086 | 0.000 | 0.000 |
| Spnb2 | 1.803 | 0.000 | 0.000 |
| Hivep3 | 1.684 | 0.001 | 0.009 |
| Appl2 | 1.660 | 0.000 | 0.000 |
| Larp1b | 1.630 | 0.000 | 0.000 |
| Alcam | 1.545 | 0.001 | 0.013 |
| Ahnak | 1.355 | 0.002 | 0.020 |
| Pde3b | 1.286 | 0.000 | 0.002 |
| E130308A19Rik | 1.246 | 0.007 | 0.040 |
| Naip5 | 1.238 | 0.000 | 0.000 |
| C230081A13Rik | 1.233 | 0.001 | 0.008 |
| Matn2 | 1.223 | 0.002 | 0.018 |
| Clip1 | 1.130 | 0.002 | 0.017 |
| Foxp3 | 1.065 | 0.000 | 0.001 |
| Cdcp1 | 1.059 | 0.004 | 0.029 |
| Bcl9 | 1.043 | 0.004 | 0.027 |
| Itih5 | 1.039 | 0.004 | 0.029 |
| Serpinb6b | 1.021 | 0.000 | 0.000 |
| Lyst | 0.980 | 0.008 | 0.045 |
| Pim2 | 0.946 | 0.000 | 0.000 |
| Cst7 | 0.883 | 0.000 | 0.004 |
| Ppic | 0.882 | 0.002 | 0.020 |
| Slc12a2 | 0.839 | 0.002 | 0.020 |
| Slc35d1 | 0.733 | 0.004 | 0.027 |
| Zfp467 | 0.712 | 0.000 | 0.005 |
| Sema4f | 0.707 | 0.000 | 0.000 |
| Acot7 | 0.669 | 0.001 | 0.011 |
| St3gal2 | 0.652 | 0.004 | 0.027 |
| Gramd1a | 0.623 | 0.000 | 0.005 |
| Gsn | 0.591 | 0.001 | 0.008 |
| Emb | -0.591 | 0.000 | 0.000 |
| Gimap7 | -0.594 | 0.000 | 0.003 |
| Ptger4 | -0.668 | 0.003 | 0.026 |
| Tmem64 | -0.678 | 0.003 | 0.023 |
| Mbnl3 | -0.691 | 0.000 | 0.004 |
| Trim59 | -0.709 | 0.001 | 0.012 |
| Tnfsf11 | -0.710 | 0.001 | 0.012 |
| Lysmd2 | -0.740 | 0.006 | 0.037 |
| Gpr83 | -0.822 | 0.004 | 0.027 |
| Slc14a1 | -0.849 | 0.001 | 0.012 |
| A130040M12Rik | -0.858 | 0.001 | 0.013 |
| 1190002H23Rik | -0.865 | 0.001 | 0.013 |
| Sh3bgrl2 | -0.874 | 0.000 | 0.000 |
| 2010111I01Rik | -0.906 | 0.000 | 0.004 |
| C030044B11Rik | -0.910 | 0.000 | 0.000 |
| Iigp1 | -0.917 | 0.008 | 0.045 |
| F2rl1 | -0.950 | 0.000 | 0.006 |

| | | | |
|---|---|---|---|
| Cish | -0.965 | 0.001 | 0.014 |
| Slamf1 | -0.973 | 0.000 | 0.000 |
| 2510009E07Rik | -0.990 | 0.001 | 0.009 |
| Gata1 | -1.066 | 0.002 | 0.016 |
| She | -1.080 | 0.001 | 0.012 |
| B930041F14Rik | -1.105 | 0.002 | 0.016 |
| Sgms1 | -1.201 | 0.000 | 0.001 |
| Nrp1 | -1.216 | 0.000 | 0.000 |
| Tgfbr3 | -1.247 | 0.000 | 0.000 |
| Ppm1l | -1.292 | 0.002 | 0.016 |
| Socs2 | -1.309 | 0.000 | 0.007 |
| Cpe | -1.385 | 0.000 | 0.003 |
| Ecm1 | -1.390 | 0.000 | 0.000 |
| Tmcc3 | -1.412 | 0.003 | 0.024 |
| Sema4a | -1.415 | 0.000 | 0.007 |
| Xdh | -1.552 | 0.000 | 0.000 |
| Gbp1 | -1.706 | 0.000 | 0.000 |
| March3' | -1.870 | 0.000 | 0.000 |
| Klrg1 | -2.147 | 0.000 | 0.000 |
| Dapl1 | -2.186 | 0.000 | 0.001 |
| Prg4 | -2.451 | 0.000 | 0.000 |
| Dnahc7b | -3.360 | 0.000 | 0.000 |
| Il1rl1 | -4.012 | 0.000 | 0.000 |

**mLNs**

| Gene name | Log2 fold change | P value | Ajusted P value |
|---|---|---|---|
| Nrn1 | 4.498 | 0.000 | 0.000 |
| Ntrk3 | 4.222 | 0.000 | 0.000 |
| Pou2af1 | 3.103 | 0.000 | 0.000 |
| Appl2 | 2.615 | 0.000 | 0.000 |
| Alcam | 2.384 | 0.000 | 0.001 |
| Abhd15 | 2.367 | 0.000 | 0.000 |
| Bhlhe40 | 2.174 | 0.000 | 0.000 |
| Klrd1 | 2.120 | 0.003 | 0.015 |
| Sestd1 | 2.049 | 0.000 | 0.000 |
| Gem | 2.021 | 0.000 | 0.001 |
| Pmaip1 | 2.018 | 0.000 | 0.000 |
| Nr4a2 | 1.862 | 0.000 | 0.000 |
| Eea1 | 1.768 | 0.000 | 0.000 |
| Hmgn3 | 1.710 | 0.000 | 0.000 |
| Cst7 | 1.690 | 0.003 | 0.016 |
| Cd81 | 1.658 | 0.001 | 0.005 |
| Odc1 | 1.517 | 0.000 | 0.000 |
| Tgm2 | 1.456 | 0.001 | 0.006 |
| Sema4f | 1.441 | 0.000 | 0.000 |
| Melk | 1.263 | 0.000 | 0.000 |
| Plcb4 | 1.262 | 0.007 | 0.029 |
| Stx11 | 1.259 | 0.000 | 0.000 |
| Slc12a2 | 1.229 | 0.000 | 0.002 |
| Pim2 | 1.218 | 0.000 | 0.000 |
| Serpinb6b | 1.200 | 0.000 | 0.000 |
| Id2 | 1.197 | 0.013 | 0.046 |
| Myo10 | 1.176 | 0.010 | 0.038 |
| E130308A19Rik | 1.164 | 0.000 | 0.000 |
| Gabarapl1 | 1.141 | 0.000 | 0.003 |
| Larp1b | 1.118 | 0.000 | 0.000 |
| Phlpp1 | 1.111 | 0.001 | 0.008 |

| | | | |
|---|---|---|---|
| Gsn | 1.087 | 0.000 | 0.000 |
| Otud1 | 1.075 | 0.000 | 0.000 |
| Cdk6 | 1.056 | 0.000 | 0.002 |
| Sla | 1.050 | 0.000 | 0.001 |
| Itgb8 | 1.026 | 0.002 | 0.010 |
| Hipk2 | 1.016 | 0.000 | 0.000 |
| 6330403K07Rik | 1.012 | 0.012 | 0.042 |
| Tnfrsf9 | 0.964 | 0.000 | 0.001 |
| Zc3h12d | 0.955 | 0.000 | 0.000 |
| Hivep3 | 0.947 | 0.001 | 0.006 |
| Frmd4b | 0.943 | 0.007 | 0.029 |
| Gpr174 | 0.909 | 0.000 | 0.000 |
| Laptm4b | 0.882 | 0.004 | 0.019 |
| Wwox | 0.839 | 0.000 | 0.001 |
| Apol7b | 0.815 | 0.005 | 0.023 |
| Pole2 | 0.739 | 0.000 | 0.002 |
| Anxa4 | 0.724 | 0.001 | 0.004 |
| Klhdc2 | 0.699 | 0.000 | 0.000 |
| Acot7 | 0.652 | 0.000 | 0.002 |
| Myo1e | 0.650 | 0.002 | 0.009 |
| AW112010 | -0.596 | 0.003 | 0.013 |
| Chchd10 | -0.606 | 0.008 | 0.032 |
| Ahcyl2 | -0.609 | 0.000 | 0.000 |
| Slc22a5 | -0.611 | 0.006 | 0.026 |
| Gfi1 | -0.637 | 0.002 | 0.010 |
| Kcnk6 | -0.644 | 0.007 | 0.027 |
| Lrig1 | -0.655 | 0.000 | 0.000 |
| Slc14a1 | -0.673 | 0.000 | 0.000 |
| Phlda1 | -0.680 | 0.009 | 0.035 |
| Rora | -0.680 | 0.014 | 0.048 |
| Rhoq | -0.709 | 0.010 | 0.036 |
| Ugcg | -0.712 | 0.007 | 0.027 |
| Dennd5a | -0.718 | 0.005 | 0.021 |
| Pik3r5 | -0.761 | 0.002 | 0.009 |
| Tgfbr2 | -0.769 | 0.004 | 0.017 |
| Mbnl3 | -0.776 | 0.000 | 0.000 |
| Scml4 | -0.789 | 0.000 | 0.000 |
| Frmd6 | -0.801 | 0.000 | 0.000 |
| Adam19 | -0.824 | 0.000 | 0.002 |
| Cyth3 | -0.825 | 0.000 | 0.000 |
| Trim59 | -0.845 | 0.000 | 0.001 |
| Swap70 | -0.870 | 0.001 | 0.007 |
| Lef1 | -0.871 | 0.004 | 0.016 |
| Sgk1 | -0.889 | 0.008 | 0.030 |
| 1110032F04Rik | -0.910 | 0.000 | 0.000 |
| Eno3 | -0.937 | 0.000 | 0.000 |
| Mboat1 | -0.957 | 0.003 | 0.013 |
| Add3 | -0.963 | 0.000 | 0.000 |
| Ikzf3 | -0.985 | 0.000 | 0.000 |
| Ift80 | -1.025 | 0.000 | 0.000 |
| Gpr183 | -1.027 | 0.000 | 0.000 |
| Rgs1 | -1.032 | 0.000 | 0.002 |
| B930041F14Rik | -1.048 | 0.000 | 0.003 |
| Igf2r | -1.061 | 0.000 | 0.003 |
| Ptger4 | -1.077 | 0.000 | 0.000 |
| Pros1 | -1.082 | 0.000 | 0.000 |
| Il2ra | -1.112 | 0.000 | 0.000 |

| | | | |
|---|---|---|---|
| 2510009E07Rik | -1.121 | 0.012 | 0.044 |
| St8sia1 | -1.132 | 0.000 | 0.000 |
| Tgfbr1 | -1.146 | 0.000 | 0.000 |
| Maml2 | -1.148 | 0.000 | 0.000 |
| Arrdc4 | -1.149 | 0.001 | 0.006 |
| Arhgap20 | -1.172 | 0.013 | 0.046 |
| Atp6v0a1 | -1.195 | 0.000 | 0.000 |
| Usp27x | -1.226 | 0.003 | 0.015 |
| Capn3 | -1.257 | 0.001 | 0.003 |
| 2010111I01Rik | -1.266 | 0.000 | 0.000 |
| Tgfbr3 | -1.266 | 0.010 | 0.036 |
| Themis | -1.291 | 0.000 | 0.000 |
| Als2cl | -1.307 | 0.003 | 0.014 |
| Ahr | -1.330 | 0.000 | 0.000 |
| Slamf1 | -1.352 | 0.000 | 0.000 |
| Gata1 | -1.396 | 0.001 | 0.004 |
| Gpr83 | -1.406 | 0.000 | 0.000 |
| Ikzf4 | -1.421 | 0.000 | 0.000 |
| Lta | -1.452 | 0.000 | 0.000 |
| Ebi3 | -1.465 | 0.010 | 0.037 |
| Specc1 | -0.260 | 0.056 | 0.140 |
| Plagl1 | -1.593 | 0.000 | 0.000 |
| Entpd1 | -1.621 | 0.000 | 0.002 |
| Rnase4 | -1.638 | 0.000 | 0.000 |
| Rapgef4 | -1.640 | 0.000 | 0.000 |
| Ggt5 | -1.650 | 0.008 | 0.032 |
| Nrp1 | -1.673 | 0.000 | 0.000 |
| Tmie | -1.733 | 0.000 | 0.001 |
| Ecm1 | -1.735 | 0.000 | 0.000 |
| Camk2n1 | -1.762 | 0.000 | 0.000 |
| Igfbp4 | -1.858 | 0.001 | 0.007 |
| Cish | -1.886 | 0.000 | 0.000 |
| She | -1.927 | 0.000 | 0.000 |
| Sema4a | -1.999 | 0.000 | 0.000 |
| Ppm1l | -2.043 | 0.000 | 0.000 |
| Tmcc3 | -2.173 | 0.000 | 0.000 |
| Lypd6b | -2.177 | 0.002 | 0.009 |
| Slc22a2 | -0.205 | 0.628 | 0.760 |
| Ddc | -2.431 | 0.000 | 0.000 |
| Socs2 | -2.579 | 0.000 | 0.000 |
| Prg4 | -2.754 | 0.000 | 0.000 |
| Adh1 | -3.157 | 0.000 | 0.000 |
| Il1rl1 | -3.421 | 0.000 | 0.000 |
| Klrg1 | -3.598 | 0.000 | 0.000 |
| Epas1 | -3.796 | 0.000 | 0.000 |

#### Supplementary Table 3. Differentially expressed genes invovled in different pathways (pooled spleen and pLNs)

##### Category 1: genes invovled in DNA damage, DNA repair and Cell cycle

**DNA Double-Strand Break Repair by Homologous Recombination**

| Gene name | Log2 fold change | P value | Ajusted P value |
|---|---|---|---|
| Lig1 | 1.103 | 0.003 | 0.026 |
| Gen1 | 1.460 | 0.000 | 0.004 |
| Pola1 | 0.779 | 0.003 | 0.021 |
| Atrx | 1.159 | 0.001 | 0.008 |
| Abl1 | 0.668 | 0.003 | 0.024 |
| Brca1 | 1.813 | 0.000 | 0.006 |
| Rad50 | 1.435 | 0.003 | 0.024 |
| Atm | 0.601 | 0.001 | 0.008 |

**Cell Cycle: G2/M DNA Damage Checkpoint Regulation**

| Gene name | Log2 fold change | P value | Ajusted P value |
|---|---|---|---|
| Cdkn2a | 2.636 | 0.000 | 0.002 |
| Prkdc | 0.744 | 0.001 | 0.012 |
| Cdkn1a | 1.649 | 0.000 | 0.000 |
| Abl1 | 0.668 | 0.003 | 0.024 |
| Top2a | 1.194 | 0.000 | 0.006 |
| Ccnb2 | 1.210 | 0.000 | 0.003 |
| Plk1 | 1.328 | 0.000 | 0.006 |
| Hipk2 | 2.086 | 0.000 | 0.000 |
| Brca1 | 1.813 | 0.000 | 0.006 |
| Prkcz | 0.719 | 0.000 | 0.000 |
| Ccnb1 | 0.752 | 0.009 | 0.048 |
| Ep300 | 0.982 | 0.002 | 0.020 |
| Atm | 0.601 | 0.001 | 0.008 |

**Role of BRCA1 in DNA Damage Response**

| Gene name | Log2 fold change | P value | Ajusted P value |
|---|---|---|---|
| Fancm | 1.016 | 0.002 | 0.021 |
| Ifng | -1.738 | 0.001 | 0.009 |
| Arid1a | 1.282 | 0.001 | 0.009 |
| Rbbp8 | 1.318 | 0.001 | 0.012 |
| Plk1 | 1.328 | 0.000 | 0.006 |
| Rfc1 | 1.541 | 0.001 | 0.009 |
| Rad50 | 1.435 | 0.003 | 0.024 |
| Smarca4 | 1.508 | 0.000 | 0.005 |
| Smarca2 | 1.018 | 0.005 | 0.034 |
| Cdkn1a | 1.649 | 0.000 | 0.000 |
| E2f7 | 2.030 | 0.003 | 0.022 |
| Smarcc2 | 0.826 | 0.005 | 0.032 |
| Smarcc1 | 1.091 | 0.001 | 0.012 |
| Stat1 | -0.638 | 0.004 | 0.027 |
| Brca1 | 1.813 | 0.000 | 0.006 |
| E2f8 | 1.603 | 0.009 | 0.048 |
| Atm | 0.601 | 0.001 | 0.008 |

**Cyclins and Cell Cycle Regulation**

| Gene name | Log2 fold change | P value | Ajusted P value |
|---|---|---|---|
| Cdkn2a | 2.636 | 0.000 | 0.002 |
| Hdac4 | 1.059 | 0.007 | 0.040 |
| Ppp2r5b | 0.753 | 0.004 | 0.027 |
| Abl1 | 0.668 | 0.003 | 0.024 |
| Ccnb2 | 1.210 | 0.000 | 0.003 |
| Ccnd1 | 2.948 | 0.008 | 0.045 |
| Ccnb1 | 0.752 | 0.009 | 0.048 |
| Hdac5 | 0.966 | 0.000 | 0.004 |
| Ccne1 | 0.873 | 0.000 | 0.006 |
| Cdkn1a | 1.649 | 0.000 | 0.000 |
| Ppm1l | -1.292 | 0.002 | 0.016 |
| E2f7 | 2.030 | 0.003 | 0.022 |
| Tgfb3 | 2.321 | 0.001 | 0.014 |
| Cdkn1b | -0.760 | 0.000 | 0.004 |
| E2f8 | 1.603 | 0.009 | 0.048 |
| Atm | 0.601 | 0.001 | 0.008 |

**ATM Signaling**

| Gene name | Log2 fold change | P value | Ajusted P value |
|---|---|---|---|
| Smc3 | 0.854 | 0.004 | 0.027 |
| Crebbp | 1.381 | 0.000 | 0.007 |
| Ppp2r5b | 0.753 | 0.004 | 0.027 |
| Abl1 | 0.668 | 0.003 | 0.024 |
| Rbbp8 | 1.318 | 0.001 | 0.012 |
| Ccnb2 | 1.210 | 0.000 | 0.003 |
| Rad50 | 1.435 | 0.003 | 0.024 |
| Ccnb1 | 0.752 | 0.009 | 0.048 |
| Smc1a | 1.189 | 0.001 | 0.012 |
| Ep300 | 0.982 | 0.002 | 0.020 |
| Herc2 | 0.915 | 0.000 | 0.006 |
| Smc2 | 1.098 | 0.000 | 0.007 |
| Trrap | 1.523 | 0.000 | 0.000 |
| Cdkn1a | 1.649 | 0.000 | 0.000 |
| Ppm1l | -1.292 | 0.002 | 0.016 |
| Trp53bp1 | 0.990 | 0.000 | 0.004 |
| Brca1 | 1.813 | 0.000 | 0.006 |
| Atm | 0.601 | 0.001 | 0.008 |

#### Category 2: genes implicated in immune cell function

**Fcγ Receptor-mediated Phagocytosis in Macrophages and Monocytes**

| Gene name | Log2 fold change | P value | Ajusted P value |
|---|---|---|---|
| Src | -0.948 | 0.000 | 0.001 |
| Gab2 | 3.494 | 0.000 | 0.004 |
| Ptk2b | 0.797 | 0.004 | 0.029 |
| Arpc1b | 0.674 | 0.000 | 0.000 |
| Crk | -0.647 | 0.000 | 0.000 |
| Tln1 | 1.632 | 0.000 | 0.003 |
| Nck1 | -0.776 | 0.000 | 0.005 |
| Fyb | 0.790 | 0.007 | 0.043 |
| Prkcz | 0.719 | 0.000 | 0.000 |
| Pld1 | 3.933 | 0.000 | 0.000 |

| | | | |
|---|---|---|---|
| Inpp5d | 0.616 | 0.001 | 0.011 |
| Pik3r3 | 1.379 | 0.000 | 0.005 |
| Pld4 | 3.314 | 0.002 | 0.020 |
| Myo5a | 1.128 | 0.007 | 0.043 |
| Arf6 | -0.591 | 0.000 | 0.000 |
| Yes1 | 2.992 | 0.000 | 0.002 |
| Gpld1 | -3.404 | 0.000 | 0.000 |
| Lyn | 2.117 | 0.001 | 0.009 |
| Pik3r2 | 0.931 | 0.001 | 0.009 |
| Prkcb | 1.448 | 0.000 | 0.001 |

**Antigen Presentation Pathway**

| Gene name | Log2 fold change | P value | Ajusted P value |
|---|---|---|---|
| H2-M3 | -0.655 | 0.001 | 0.008 |
| Psmb9' | -0.602 | 0.007 | 0.041 |
| Ifng' | -1.738 | 0.001 | 0.009 |
| H2-DMa | -0.993 | 0.001 | 0.011 |
| Gm8909 | -0.704 | 0.002 | 0.017 |
| H2-DMb1 | 1.646 | 0.000 | 0.006 |
| H2-Aa | 2.905 | 0.000 | 0.000 |
| Psmb8' | -0.604 | 0.003 | 0.023 |
| H2-Ab1 | 2.787 | 0.000 | 0.000 |
| Cd74' | 1.091 | 0.000 | 0.000 |
| Tap1' | -0.586 | 0.000 | 0.007 |
| H2-Eb1 | 2.703 | 0.000 | 0.000 |

**Th1 and Th2 Activation Pathway**

| Gene name | Log2 fold change | P value | Ajusted P value |
|---|---|---|---|
| Cd247 | -0.719 | 0.001 | 0.011 |
| Tnfsf4 | 1.914 | 0.000 | 0.001 |
| Chd4 | 1.342 | 0.000 | 0.004 |
| Gm8909 | -0.704 | 0.002 | 0.017 |
| Il1rl1 | -4.012 | 0.000 | 0.000 |
| Tgfbr3 | -1.247 | 0.000 | 0.000 |
| Havcr2 | 2.075 | 0.000 | 0.001 |
| H2-Aa | 2.905 | 0.000 | 0.000 |
| Cxcr3 | -0.921 | 0.001 | 0.014 |
| H2-Ab1 | 2.787 | 0.000 | 0.000 |
| Il18r1 | -1.979 | 0.000 | 0.000 |
| H2-DMa | -0.993 | 0.001 | 0.011 |
| H2-DMb1 | 1.646 | 0.000 | 0.006 |
| Pik3r2 | 0.931 | 0.001 | 0.009 |
| Stat1 | -0.638 | 0.004 | 0.027 |
| Atm | 0.601 | 0.001 | 0.008 |
| Klrc1 | 5.924 | 0.000 | 0.003 |
| Runx3 | 1.119 | 0.000 | 0.001 |
| Ifng | -1.738 | 0.001 | 0.009 |
| Tnfsf11 | -0.710 | 0.001 | 0.012 |
| Il12rb2 | 2.991 | 0.000 | 0.003 |
| Irf1 | -0.619 | 0.000 | 0.003 |
| Pik3r3 | 1.379 | 0.000 | 0.005 |
| Il17rb | -1.914 | 0.000 | 0.005 |
| S1pr1 | -0.890 | 0.003 | 0.024 |
| Il10rb | -0.616 | 0.000 | 0.004 |

| | | | |
|---|---|---|---|
| Nfatc2 | 0.653 | 0.001 | 0.013 |
| Jag1 | 3.256 | 0.000 | 0.000 |
| H2-Eb1 | 2.703 | 0.000 | 0.000 |

**Th1 Pathway**

| Gene name | Log2 fold change | P value | Ajusted P value |
|---|---|---|---|
| Cd247 | -0.719 | 0.001 | 0.011 |
| Runx3 | 1.119 | 0.000 | 0.001 |
| Ifng | -1.738 | 0.001 | 0.009 |
| Tnfsf11 | -0.710 | 0.001 | 0.012 |
| Gm8909 | -0.704 | 0.002 | 0.017 |
| Havcr2 | 2.075 | 0.000 | 0.001 |
| H2-Aa | 2.905 | 0.000 | 0.000 |
| Cxcr3 | -0.921 | 0.001 | 0.014 |
| Il12rb2 | 2.991 | 0.000 | 0.003 |
| H2-Ab1 | 2.787 | 0.000 | 0.000 |
| Irf1 | -0.619 | 0.000 | 0.003 |
| Il18r1 | -1.979 | 0.000 | 0.000 |
| Pik3r3 | 1.379 | 0.000 | 0.005 |
| H2-DMa | -0.993 | 0.001 | 0.011 |
| H2-DMb1 | 1.646 | 0.000 | 0.006 |
| Il10rb | -0.616 | 0.000 | 0.004 |
| Nfatc2 | 0.653 | 0.001 | 0.013 |
| Pik3r2 | 0.931 | 0.001 | 0.009 |
| Stat1 | -0.638 | 0.004 | 0.027 |
| H2-Eb1 | 2.703 | 0.000 | 0.000 |
| Atm | 0.601 | 0.001 | 0.008 |
| Klrc1 | 5.924 | 0.000 | 0.003 |

**Altered T Cell and B Cell Signaling in Rheumatoid Arthritis**

| Gene name | Log2 fold change | P value | Ajusted P value |
|---|---|---|---|
| Ifng | -1.738 | 0.001 | 0.009 |
| Map3k14 | 0.760 | 0.004 | 0.028 |
| Tnfsf11 | -0.710 | 0.001 | 0.012 |
| Traf3 | 0.732 | 0.006 | 0.038 |
| Spp1 | 3.639 | 0.002 | 0.021 |
| Slamf1 | -0.973 | 0.000 | 0.000 |
| Cd79b | -2.076 | 0.000 | 0.000 |
| Gm8909 | -0.704 | 0.002 | 0.017 |
| H2-Aa | 2.905 | 0.000 | 0.000 |
| H2-Ab1 | 2.787 | 0.000 | 0.000 |
| Fas | -0.715 | 0.001 | 0.009 |
| Tlr4 | 1.990 | 0.000 | 0.000 |
| H2-DMa | -0.993 | 0.001 | 0.011 |
| H2-DMb1 | 1.646 | 0.000 | 0.006 |
| Tlr1 | -0.869 | 0.000 | 0.006 |
| H2-Eb1 | 2.703 | 0.000 | 0.000 |
| Tnfsf13b | -2.039 | 0.000 | 0.002 |

**Calcium-induced T Lymphocyte Apoptosis**

| Gene name | Log2 fold change | P value | Ajusted P value |
|---|---|---|---|
| Cd247 | -0.719 | 0.001 | 0.011 |
| Itpr2 | 1.391 | 0.001 | 0.012 |

| | | | |
|---|---|---|---|
| Gm8909 | -0.704 | 0.002 | 0.017 |
| H2-Aa | 2.905 | 0.000 | 0.000 |
| H2-Ab1 | 2.787 | 0.000 | 0.000 |
| Itpr1 | 1.556 | 0.002 | 0.017 |
| Prkcz | 0.719 | 0.000 | 0.000 |
| Ep300 | 0.982 | 0.002 | 0.020 |
| Cabin1 | 0.853 | 0.002 | 0.021 |
| H2-DMa | -0.993 | 0.001 | 0.011 |
| H2-DMb1 | 1.646 | 0.000 | 0.006 |
| Nfatc2 | 0.653 | 0.001 | 0.013 |
| H2-Eb1 | 2.703 | 0.000 | 0.000 |
| Prkcb | 1.448 | 0.000 | 0.001 |

**Type I Diabetes Mellitus Signaling**

| Gene name | Log2 fold change | P value | Ajusted P value |
|---|---|---|---|
| Cd247 | -0.719 | 0.001 | 0.011 |
| Ifng | -1.738 | 0.001 | 0.009 |
| Map3k14 | 0.760 | 0.004 | 0.028 |
| Ica1 | 1.346 | 0.000 | 0.005 |
| Tnfrsf1a | -0.802 | 0.005 | 0.033 |
| Gm8909 | -0.704 | 0.002 | 0.017 |
| Pias1 | -0.590 | 0.001 | 0.012 |
| H2-Aa | 2.905 | 0.000 | 0.000 |
| H2-Ab1 | 2.787 | 0.000 | 0.000 |
| Il1r1 | 4.906 | 0.001 | 0.008 |
| Fas | -0.715 | 0.001 | 0.009 |
| Irf1 | -0.619 | 0.000 | 0.003 |
| H2-M3 | -0.655 | 0.001 | 0.008 |
| H2-DMa | -0.993 | 0.001 | 0.011 |
| H2-DMb1 | 1.646 | 0.000 | 0.006 |
| Socs2 | -1.309 | 0.000 | 0.007 |
| Stat1 | -0.638 | 0.004 | 0.027 |
| Cpe | -1.385 | 0.000 | 0.003 |
| H2-Eb1 | 2.703 | 0.000 | 0.000 |

#### Category 3: genes related to molecular mechanism of Cancer

**Molecular Mechanisms of Cancer**

| Gene name | Log2 fold change | P value | Ajusted P value |
|---|---|---|---|
| Cdkn2a | 2.636 | 0.000 | 0.002 |
| Gab2 | 3.494 | 0.000 | 0.004 |
| Rapgef1 | 0.923 | 0.000 | 0.007 |
| Naip1 | 0.921 | 0.001 | 0.014 |
| Abl1 | 0.668 | 0.003 | 0.024 |
| Ctnna1 | 0.779 | 0.004 | 0.029 |
| Crk | -0.647 | 0.000 | 0.000 |
| Ccnd1 | 2.948 | 0.008 | 0.045 |
| Rhoh | -0.863 | 0.000 | 0.001 |
| Fas | -0.715 | 0.001 | 0.009 |
| Prkcz | 0.719 | 0.000 | 0.000 |
| Ep300 | 0.982 | 0.002 | 0.020 |
| Myc | 0.881 | 0.000 | 0.000 |
| Zbtb17 | 0.834 | 0.001 | 0.014 |
| Camk2a | -2.046 | 0.000 | 0.001 |

| | | | |
|---|---|---|---|
| Bbc3 | 0.820 | 0.000 | 0.001 |
| Gna15 | -0.792 | 0.000 | 0.001 |
| Rhob | -0.755 | 0.001 | 0.012 |
| Bmpr1a | 1.605 | 0.000 | 0.000 |
| Smo | 2.209 | 0.000 | 0.000 |
| Pik3r2 | 0.931 | 0.001 | 0.009 |
| Hipk2 | 2.086 | 0.000 | 0.000 |
| Brca1 | 1.813 | 0.000 | 0.006 |
| Tab1 | 0.718 | 0.000 | 0.006 |
| E2f8 | 1.603 | 0.009 | 0.048 |
| Atm | 0.601 | 0.001 | 0.008 |
| Itgb1 | -0.837 | 0.001 | 0.008 |
| Src | -0.948 | 0.000 | 0.001 |
| Prkdc | 0.744 | 0.001 | 0.012 |
| Cdk13 | 0.864 | 0.005 | 0.035 |
| Crebbp | 1.381 | 0.000 | 0.007 |
| Prkar2a | 0.667 | 0.000 | 0.002 |
| Arhgef17 | 4.389 | 0.002 | 0.019 |
| Apc | 1.107 | 0.009 | 0.048 |
| Pik3r3 | 1.379 | 0.000 | 0.005 |
| Rasgrf2 | -1.372 | 0.000 | 0.003 |
| Plcb4 | 2.580 | 0.000 | 0.000 |
| Ccne1 | 0.873 | 0.000 | 0.006 |
| Cdh1 | 2.180 | 0.000 | 0.004 |
| E2f7 | 2.030 | 0.003 | 0.022 |
| Cdk11b | 0.954 | 0.001 | 0.010 |
| Cdkn1a | 1.649 | 0.000 | 0.000 |
| Tgfb3 | 2.321 | 0.001 | 0.014 |
| Rbpj | -0.616 | 0.000 | 0.001 |
| Cdkn1b | -0.760 | 0.000 | 0.004 |
| Bcl2l11 | 0.618 | 0.007 | 0.040 |
| Lrp1 | 5.203 | 0.000 | 0.001 |
| Prkcb | 1.448 | 0.000 | 0.001 |

#### Supplementary Table 4. Pearson Correlations between replicates for mLN RNA-seq

| | Treg WT1 | Treg WT2 | Treg WT3 |
|---|---|---|---|
| Treg WT1 | 1 | | |
| Treg WT2 | 0.942 | 1 | |
| Treg WT3 | 0.942 | 0.943 | 1 |

| | CD4 WT1 | CD4 WT2 | CD4 WT3 |
|---|---|---|---|
| CD4 WT1 | 1 | | |
| CD4 WT2 | 0.943 | 1 | |
| CD4 WT3 | 0.942 | 0.943 | 1 |

| | Treg DKO1 | Treg DKO2 | Treg DKO3 |
|---|---|---|---|
| Treg DKO1 | 1 | | |
| Treg DKO2 | 0.918 | 1 | |
| Treg DKO3 | 0.922 | 0.917 | 1 |

| | CD4 DKO1 | CD4 DKO2 | CD4 DKO3 |
|---|---|---|---|
| CD4 DKO1 | 1 | | |
| CD4 DKO2 | 0.913 | 1 | |
| CD4 DKO3 | 0.926 | 0.918 | 1 |

### Supplementary Table 5. Differentially expressed genes invovled in different pathways (mLNs)

#### Category 1: genes invovled in DNA damage, DNA repair and Cell cycle

##### Cell Cycle Control of Chromosomal Replication

| Gene name | Log2 fold change | P value | Ajusted P value |
|---|---|---|---|
| Mcm5 | 1.010 | 0.000 | 0.000 |
| Lig1 | 1.019 | 0.000 | 0.000 |
| Mcm6 | 0.986 | 0.000 | 0.000 |
| Cdc45 | 1.298 | 0.000 | 0.000 |
| Mcm8 | 1.075 | 0.000 | 0.000 |
| Pola1 | 0.796 | 0.000 | 0.000 |
| Cdc6 | 1.294 | 0.000 | 0.000 |
| Pole | 1.284 | 0.000 | 0.000 |
| Cdk6 | 1.056 | 0.000 | 0.002 |
| Dna2 | 1.453 | 0.000 | 0.000 |
| Cdk14 | 5.438 | 0.000 | 0.003 |
| Cdk1 | 1.497 | 0.000 | 0.000 |
| Mcm3 | 0.900 | 0.000 | 0.000 |
| Prim1 | 0.941 | 0.000 | 0.000 |
| Mcm2 | 0.658 | 0.000 | 0.000 |
| Top2a | 1.108 | 0.000 | 0.000 |
| Chek2 | 1.060 | 0.000 | 0.000 |
| Dbf4 | 0.799 | 0.000 | 0.000 |
| Mcm7 | 0.781 | 0.000 | 0.000 |
| Orc1 | 2.068 | 0.000 | 0.000 |

##### Cell Cycle: G2/M DNA Damage Checkpoint Regulation

| Gene name | Log2 fold change | P value | Ajusted P value |
|---|---|---|---|
| Cdkn2a | 3.986 | 0.000 | 0.003 |
| Cdc25c | 1.632 | 0.000 | 0.000 |
| Cks2 | 0.841 | 0.002 | 0.010 |
| Wee1 | 1.345 | 0.001 | 0.005 |
| Ccnb2 | 1.965 | 0.000 | 0.000 |
| Plk1 | 1.706 | 0.000 | 0.000 |
| Aurka | 1.339 | 0.000 | 0.000 |
| Cdk1 | 1.497 | 0.000 | 0.000 |
| Chek1 | 1.368 | 0.000 | 0.000 |
| Ccnb1 | 1.587 | 0.000 | 0.000 |
| Cdc25b | 1.480 | 0.001 | 0.008 |
| Cdkn1a | 1.297 | 0.000 | 0.001 |
| Top2a | 1.108 | 0.000 | 0.000 |
| Cks1b | 1.340 | 0.000 | 0.000 |
| Bora | 0.917 | 0.000 | 0.001 |
| Hipk2 | 1.016 | 0.000 | 0.000 |
| Brca1 | 1.359 | 0.000 | 0.000 |
| Chek2 | 1.060 | 0.000 | 0.000 |

##### Cyclins and Cell Cycle Regulation

| Gene name | Log2 fold change | P value | Ajusted P value |
|---|---|---|---|
| Cdkn2a | 3.986 | 0.000 | 0.003 |
| Raf1 | 0.666 | 0.000 | 0.000 |
| Ccne2 | 1.630 | 0.000 | 0.000 |

| | | | |
|---|---|---|---|
| Hdac4 | 0.726 | 0.000 | 0.000 |
| Tfdp1 | 0.655 | 0.000 | 0.000 |
| Wee1 | 1.345 | 0.001 | 0.005 |
| Cdk6 | 1.056 | 0.000 | 0.002 |
| Cdkn2c | 1.036 | 0.004 | 0.016 |
| Ccnb2 | 1.965 | 0.000 | 0.000 |
| Cdk1 | 1.497 | 0.000 | 0.000 |
| Ccnb1 | 1.587 | 0.000 | 0.000 |
| Hdac5 | 0.651 | 0.000 | 0.000 |
| Ccna2 | 1.675 | 0.000 | 0.000 |
| Ccne1 | 1.382 | 0.000 | 0.000 |
| Ppm1j | -1.198 | 0.008 | 0.031 |
| Cdkn1a | 1.297 | 0.000 | 0.001 |
| Ppm1l | -2.043 | 0.000 | 0.000 |
| E2f1 | 1.415 | 0.000 | 0.000 |
| E2f7 | 2.234 | 0.000 | 0.000 |
| Tgfb3 | 2.307 | 0.000 | 0.000 |
| E2f8 | 1.788 | 0.000 | 0.000 |

**Role of CHK Proteins in Cell Cycle Checkpoint Control**

| Gene name | Log2 fold change | P value | Ajusted P value |
|---|---|---|---|
| Cdc25c | 1.632 | 0.000 | 0.000 |
| Plk1 | 1.706 | 0.000 | 0.000 |
| Rfc1 | 0.680 | 0.000 | 0.000 |
| Rfc5 | 0.928 | 0.000 | 0.000 |
| Cdk1 | 1.497 | 0.000 | 0.000 |
| Chek1 | 1.368 | 0.000 | 0.000 |
| Rfc4 | 0.621 | 0.001 | 0.008 |
| Ppm1j | -1.198 | 0.008 | 0.031 |
| Cdkn1a | 1.297 | 0.000 | 0.001 |
| Ppm1l | -2.043 | 0.000 | 0.000 |
| E2f1 | 1.415 | 0.000 | 0.000 |
| E2f7 | 2.234 | 0.000 | 0.000 |
| Clspn | 1.526 | 0.000 | 0.000 |
| Brca1 | 1.359 | 0.000 | 0.000 |
| Check2 | 1.060 | 0.000 | 0.000 |
| E2f8 | 1.788 | 0.000 | 0.000 |
| Rfc3 | 0.670 | 0.000 | 0.000 |

**Role of BRCA1 in DNA Damage Response**

| Gene name | Log2 fold change | P value | Ajusted P value |
|---|---|---|---|
| Bard1 | 0.685 | 0.002 | 0.012 |
| Rbbp8 | 1.216 | 0.000 | 0.000 |
| Plk1 | 1.706 | 0.000 | 0.000 |
| Rfc1 | 0.680 | 0.000 | 0.000 |
| Rfc5 | 0.928 | 0.000 | 0.000 |
| Check1 | 1.368 | 0.000 | 0.000 |
| Rad51 | 1.300 | 0.000 | 0.000 |
| Rfc4 | 0.621 | 0.001 | 0.008 |
| Fancd2 | 0.750 | 0.000 | 0.002 |
| Cdkn1a | 1.297 | 0.000 | 0.001 |
| E2f1 | 1.415 | 0.000 | 0.000 |
| E2f7 | 2.234 | 0.000 | 0.000 |
| Brca2 | 0.731 | 0.000 | 0.002 |
| Brca1 | 1.359 | 0.000 | 0.000 |
| Brip1 | 1.293 | 0.000 | 0.000 |

| | | | |
|---|---|---|---|
| Stat1 | -0.777 | 0.000 | 0.001 |
| Fanca | 1.677 | 0.000 | 0.000 |
| Chek2 | 1.060 | 0.000 | 0.000 |
| E2f8 | 1.788 | 0.000 | 0.000 |
| Rfc3 | 0.670 | 0.000 | 0.000 |

#### Category 2: genes implicated in immune cell function

**Th1 and Th2 Activation Pathway**

| Gene name | Log2 fold change | P value | Ajusted P value |
|---|---|---|---|
| Cd247 | -0.846 | 0.000 | 0.000 |
| Ccr3 | 1.961 | 0.006 | 0.023 |
| Socs1 | -0.884 | 0.000 | 0.002 |
| Socs3 | -0.907 | 0.003 | 0.013 |
| Tnfsf4 | 2.973 | 0.000 | 0.001 |
| Tgfbr1 | -1.146 | 0.000 | 0.000 |
| Il12rb1 | -0.596 | 0.002 | 0.010 |
| H2-K1 | -0.707 | 0.000 | 0.000 |
| Klrd1 | 2.120 | 0.003 | 0.015 |
| Il1rl1 | -3.421 | 0.000 | 0.000 |
| Tgfbr3 | -1.266 | 0.010 | 0.036 |
| Pik3r1 | -0.753 | 0.000 | 0.000 |
| Pik3r5 | -0.761 | 0.002 | 0.009 |
| H2-Aa | 4.242 | 0.000 | 0.000 |
| Psen2 | -0.633 | 0.002 | 0.009 |
| H2-Ab1 | 5.073 | 0.000 | 0.000 |
| Il18r1 | -1.527 | 0.000 | 0.000 |
| Tgfbr2 | -0.769 | 0.004 | 0.017 |
| Notch2 | -0.809 | 0.000 | 0.000 |
| H2-DMA | -1.462 | 0.000 | 0.000 |
| H2-DMb1 | 1.410 | 0.002 | 0.010 |
| Cd274 | -0.723 | 0.000 | 0.000 |
| Stat1 | -0.777 | 0.000 | 0.001 |
| Klrc1 | 5.209 | 0.000 | 0.003 |
| Ccr1 | 3.059 | 0.000 | 0.000 |
| Runx3 | 0.745 | 0.000 | 0.000 |
| Il4ra | -0.999 | 0.000 | 0.003 |
| Dll1 | 3.783 | 0.010 | 0.039 |
| Cxcr4 | 0.959 | 0.000 | 0.000 |
| Il6ra | -0.890 | 0.000 | 0.001 |
| Acvr1 | 1.629 | 0.001 | 0.005 |
| Ifngr2 | -1.201 | 0.000 | 0.000 |
| Il12rb2 | 3.924 | 0.000 | 0.000 |
| Acvr1b | -0.784 | 0.001 | 0.003 |
| Itgb2 | -0.654 | 0.000 | 0.000 |
| Il17rb | -3.979 | 0.000 | 0.000 |
| Lta | -1.452 | 0.000 | 0.000 |
| Icos | 1.209 | 0.008 | 0.031 |
| Gfi1 | -0.637 | 0.002 | 0.010 |
| S1pr1 | -0.746 | 0.001 | 0.004 |
| Il2ra | -1.112 | 0.000 | 0.000 |
| Jag1 | 2.834 | 0.000 | 0.000 |
| Acvr2a | -1.503 | 0.000 | 0.000 |
| H2-Eb1 | 3.662 | 0.000 | 0.000 |

**T Helper Cell Differentiation**

| Gene name | Log2 fold change | P value | Ajusted P value |
|---|---|---|---|
| Tgfbr1 | -1.146 | 0.000 | 0.000 |
| Il12rb1 | -0.596 | 0.002 | 0.010 |
| H2-K1 | -0.707 | 0.000 | 0.000 |
| H2-Aa | 4.242 | 0.000 | 0.000 |
| H2-Ab1 | 5.073 | 0.000 | 0.000 |
| Bcl6 | 1.839 | 0.059 | 0.146 |
| Il18r1 | -1.527 | 0.000 | 0.000 |
| Tgfbr2 | -0.769 | 0.004 | 0.017 |
| H2-DMA | -1.462 | 0.000 | 0.000 |
| H2-DMb1 | 1.410 | 0.002 | 0.010 |
| Stat1 | -0.777 | 0.000 | 0.001 |
| Il17f | 4.602 | 0.002 | 0.012 |
| Il21 | 2.682 | 0.000 | 0.001 |
| Il4ra | -0.999 | 0.000 | 0.003 |
| Tnfrsf1a | -0.702 | 0.003 | 0.015 |
| Il6ra | -0.890 | 0.000 | 0.001 |
| Ifngr2 | -1.201 | 0.000 | 0.000 |
| Il12rb2 | 3.924 | 0.000 | 0.000 |
| Il17a | 4.094 | 0.005 | 0.021 |
| Il23r | 2.693 | 0.000 | 0.000 |
| Icos | 1.209 | 0.008 | 0.031 |
| Il2ra | -1.112 | 0.000 | 0.000 |
| Cxcr5 | 0.845 | 0.008 | 0.030 |
| Tnf | -0.823 | 0.002 | 0.009 |
| H2-Eb1 | 3.662 | 0.000 | 0.000 |

**Th2 Pathway**

| Gene name | Log2 fold change | P value | Ajusted P value |
|---|---|---|---|
| Cd247 | -0.846 | 0.000 | 0.000 |
| Ccr3 | 1.961 | 0.006 | 0.023 |
| Socs3 | -0.907 | 0.003 | 0.013 |
| Tnfsf4 | 2.973 | 0.000 | 0.001 |
| Tgfbr1 | -1.146 | 0.000 | 0.000 |
| Il12rb1 | -0.596 | 0.002 | 0.010 |
| H2-K1 | -0.707 | 0.000 | 0.000 |
| Il1rl1 | -3.421 | 0.000 | 0.000 |
| Pik3r1 | -0.753 | 0.000 | 0.000 |
| Tgfbr3 | -1.266 | 0.010 | 0.036 |
| Pik3r5 | -0.761 | 0.002 | 0.009 |
| H2-Aa | 4.242 | 0.000 | 0.000 |
| Psen2 | -0.633 | 0.002 | 0.009 |
| H2-Ab1 | 5.073 | 0.000 | 0.000 |
| Tgfbr2 | -0.769 | 0.004 | 0.017 |
| Notch2 | -0.809 | 0.000 | 0.000 |
| H2-DMA | -1.462 | 0.000 | 0.000 |
| H2-DMb1 | 1.410 | 0.002 | 0.010 |
| Ccr1 | 3.059 | 0.000 | 0.000 |
| Runx3 | 0.745 | 0.000 | 0.000 |
| Il4ra | -0.999 | 0.000 | 0.003 |
| Dll1 | 3.783 | 0.010 | 0.039 |
| Cxcr4 | 0.959 | 0.000 | 0.000 |
| Acvr1 | 1.629 | 0.001 | 0.005 |
| Il12rb2 | 3.924 | 0.000 | 0.000 |
| Acvr1b | -0.784 | 0.001 | 0.003 |
| Itgb2 | -0.654 | 0.000 | 0.000 |

| | | | |
|---|---|---|---|
| Il17rb | -3.979 | 0.000 | 0.000 |
| Icos | 1.209 | 0.008 | 0.031 |
| Gfi1 | -0.637 | 0.002 | 0.010 |
| S1pr1 | -0.746 | 0.001 | 0.004 |
| Il2ra | -1.112 | 0.000 | 0.000 |
| Jag1 | 2.834 | 0.000 | 0.000 |
| Acvr2a | -1.503 | 0.000 | 0.000 |
| H2-Eb1 | 3.662 | 0.000 | 0.000 |

**Aryl Hydrocarbon Receptor Signaling**

| Gene name | Log2 fold change | P value | Ajusted P value |
|---|---|---|---|
| Cdkn2a | 3.986 | 0.000 | 0.003 |
| Gstm5 | 5.233 | 0.000 | 0.000 |
| Pola1 | 0.796 | 0.000 | 0.000 |
| Rarg | -0.671 | 0.003 | 0.013 |
| Chek1 | 1.368 | 0.000 | 0.000 |
| Tgm2 | 1.456 | 0.001 | 0.006 |
| Ccna2 | 1.675 | 0.000 | 0.000 |
| Gstt2/Gstt2b | 1.491 | 0.000 | 0.000 |
| Aldh1l1 | -1.346 | 0.004 | 0.017 |
| Rara | -1.034 | 0.004 | 0.016 |
| Gstm4 | 0.698 | 0.009 | 0.035 |
| Chek2 | 1.060 | 0.000 | 0.000 |
| Aldh6a1 | -0.701 | 0.001 | 0.006 |
| Ahr | -1.330 | 0.000 | 0.000 |
| Aldh7a1 | 0.908 | 0.001 | 0.007 |
| Src | -2.318 | 0.000 | 0.000 |
| Ccne2 | 1.630 | 0.000 | 0.000 |
| Tfdp1 | 0.655 | 0.000 | 0.000 |
| Gstm1 | 2.589 | 0.000 | 0.000 |
| Cdk6 | 1.056 | 0.000 | 0.002 |
| Bax | 0.613 | 0.000 | 0.000 |
| Ccne1 | 1.382 | 0.000 | 0.000 |
| Mgst2 | -1.281 | 0.000 | 0.000 |
| Cdkn1a | 1.297 | 0.000 | 0.001 |
| E2f1 | 1.415 | 0.000 | 0.000 |
| Tgfb3 | 2.307 | 0.000 | 0.000 |
| Dhfr | 1.111 | 0.000 | 0.000 |
| Rxra | -0.732 | 0.010 | 0.037 |
| Tnf | -0.823 | 0.002 | 0.009 |
| Mcm7 | 0.781 | 0.000 | 0.000 |
| Hspb1 | 3.305 | 0.000 | 0.002 |

**Th1 Pathway**

| Gene name | Log2 fold change | P value | Ajusted P value |
|---|---|---|---|
| Cd247 | -0.846 | 0.000 | 0.000 |
| Socs1 | -0.884 | 0.000 | 0.002 |
| Socs3 | -0.907 | 0.003 | 0.013 |
| Il12rb1 | -0.596 | 0.002 | 0.010 |
| Klrd1 | 2.120 | 0.003 | 0.015 |
| H2-K1 | -0.707 | 0.000 | 0.000 |
| Pik3r1 | -0.753 | 0.000 | 0.000 |
| Pik3r5 | -0.761 | 0.002 | 0.009 |
| H2-Aa | 4.242 | 0.000 | 0.000 |
| Psen2 | -0.633 | 0.002 | 0.009 |
| H2-Ab1 | 5.073 | 0.000 | 0.000 |

| | | | |
|---|---|---|---|
| Il18r1 | -1.527 | 0.000 | 0.000 |
| Notch2 | -0.809 | 0.000 | 0.000 |
| H2-DMA | -1.462 | 0.000 | 0.000 |
| H2-DMb1 | 1.410 | 0.002 | 0.010 |
| Cd274 | -0.723 | 0.000 | 0.000 |
| Stat1 | -0.777 | 0.000 | 0.001 |
| Klrc1 | 5.209 | 0.000 | 0.003 |
| Runx3 | 0.745 | 0.000 | 0.000 |
| Dll1 | 3.783 | 0.010 | 0.039 |
| Il6ra | -0.890 | 0.000 | 0.001 |
| Ifngr2 | -1.201 | 0.000 | 0.000 |
| Il12rb2 | 3.924 | 0.000 | 0.000 |
| Itgb2 | -0.654 | 0.000 | 0.000 |
| Lta | -1.452 | 0.000 | 0.000 |
| Icos | 1.209 | 0.008 | 0.031 |
| H2-Eb1 | 3.662 | 0.000 | 0.000 |

#### Category 3: genes related to molecular mechanism of Cancer

**Molecular Mechanisms of Cancer**

| Gene name | Log2 fold change | P value | Ajusted P value |
|---|---|---|---|
| Cdkn2a | 3.986 | 0.000 | 0.003 |
| Gab2 | 3.045 | 0.000 | 0.000 |
| Raf1 | 0.666 | 0.000 | 0.000 |
| Naip1 | 0.905 | 0.004 | 0.017 |
| Tgfbr1 | -1.146 | 0.000 | 0.000 |
| Pik3r1 | -0.753 | 0.000 | 0.000 |
| Hras | 0.600 | 0.000 | 0.000 |
| Cdkn2c | 1.036 | 0.004 | 0.016 |
| Tgfbr2 | -0.769 | 0.004 | 0.017 |
| Rasgrf1 | 6.656 | 0.000 | 0.000 |
| Itga3 | 1.212 | 0.004 | 0.019 |
| Camk2a | -2.428 | 0.000 | 0.000 |
| Bbc3 | 0.613 | 0.014 | 0.048 |
| Rhob | -1.268 | 0.004 | 0.016 |
| Hipk2 | 1.016 | 0.000 | 0.000 |
| Brca1 | 1.359 | 0.000 | 0.000 |
| Ccne2 | 1.630 | 0.000 | 0.000 |
| Tfdp1 | 0.655 | 0.000 | 0.000 |
| Casp3 | 1.422 | 0.000 | 0.000 |
| Cdk6 | 1.056 | 0.000 | 0.002 |
| Cdk14 | 5.438 | 0.000 | 0.003 |
| Arhgef17 | 2.516 | 0.007 | 0.027 |
| Aurka | 1.339 | 0.000 | 0.000 |
| Rasgrf2 | -2.019 | 0.000 | 0.000 |
| Rhoq | -0.709 | 0.010 | 0.036 |
| Rnd3 | 5.180 | 0.002 | 0.009 |
| E2f1 | 1.415 | 0.000 | 0.000 |
| Tgfb3 | 2.307 | 0.000 | 0.000 |
| Lef1 | -0.871 | 0.004 | 0.016 |
| Map2k3 | 0.645 | 0.000 | 0.000 |
| Gnal | 1.847 | 0.007 | 0.029 |
| Pik3r5 | -0.761 | 0.002 | 0.009 |
| Psen2 | -0.633 | 0.002 | 0.009 |
| Hif1a | 1.286 | 0.000 | 0.000 |
| Gna14 | 2.724 | 0.000 | 0.000 |

| | | | |
|---|---|---|---|
| Rhoh | -0.864 | 0.000 | 0.000 |
| Chek1 | 1.368 | 0.000 | 0.000 |
| Bcl2 | -0.930 | 0.000 | 0.000 |
| Cdc25b | 1.480 | 0.001 | 0.008 |
| Syngap1 | 3.545 | 0.000 | 0.002 |
| Fancd2 | 0.750 | 0.000 | 0.002 |
| Smo | 1.836 | 0.002 | 0.009 |
| Rhou | 6.900 | 0.000 | 0.000 |
| Prkce | -0.645 | 0.001 | 0.008 |
| Casp8 | -0.600 | 0.000 | 0.000 |
| Chek2 | 1.060 | 0.000 | 0.000 |
| E2f8 | 1.788 | 0.000 | 0.000 |
| Bmp1 | 3.610 | 0.000 | 0.000 |
| Naip2 | 0.642 | 0.001 | 0.004 |
| Src | -2.318 | 0.000 | 0.000 |
| Pak4 | 0.771 | 0.002 | 0.010 |
| Cdc25c | 1.632 | 0.000 | 0.000 |
| Itga2 | 1.460 | 0.003 | 0.014 |
| Prkar2a | 1.098 | 0.000 | 0.000 |
| Gnai1 | 4.581 | 0.003 | 0.015 |
| Smad7 | -1.115 | 0.002 | 0.011 |
| Bax | 0.613 | 0.000 | 0.000 |
| Cdk1 | 1.497 | 0.000 | 0.000 |
| Plcb4 | 1.262 | 0.007 | 0.029 |
| Rhov | 3.372 | 0.001 | 0.005 |
| Ccne1 | 1.382 | 0.000 | 0.000 |
| Fzd4 | 7.381 | 0.000 | 0.000 |
| Foxo1 | -0.632 | 0.014 | 0.049 |
| Rasgrp1 | -0.910 | 0.000 | 0.001 |
| E2f7 | 2.234 | 0.000 | 0.000 |
| Cdkn1a | 1.297 | 0.000 | 0.001 |
| Bmp7 | -0.935 | 0.001 | 0.005 |
| Lrp1 | 3.829 | 0.000 | 0.000 |

**Pancreatic Adenocarcinoma Signaling**

| Gene name | Log2 fold change | P value | Ajusted P value |
|---|---|---|---|
| Cdkn2a | 3.986 | 0.000 | 0.003 |
| Raf1 | 0.666 | 0.000 | 0.000 |
| Tgfbr1 | -1.146 | 0.000 | 0.000 |
| Pik3r1 | -0.753 | 0.000 | 0.000 |
| Pik3r5 | -0.761 | 0.002 | 0.009 |
| Bcl2 | -0.930 | 0.000 | 0.000 |
| Rad51 | 1.300 | 0.000 | 0.000 |
| Tgfbr2 | -0.769 | 0.004 | 0.017 |
| Hmox1 | 1.122 | 0.003 | 0.013 |
| Gpld1 | -4.036 | 0.000 | 0.000 |
| Erbb2 | 6.036 | 0.000 | 0.001 |
| Stat1 | -0.777 | 0.000 | 0.001 |
| E2f8 | 1.788 | 0.000 | 0.000 |
| Egfr | 1.918 | 0.011 | 0.040 |
| Pld3 | 0.628 | 0.000 | 0.001 |
| Tfdp1 | 0.655 | 0.000 | 0.000 |
| Vegfc | 3.422 | 0.000 | 0.000 |
| Hbegf | -1.938 | 0.001 | 0.004 |
| Birc5 | 1.534 | 0.000 | 0.000 |
| Pld1 | 2.584 | 0.000 | 0.000 |
| Pld4 | 3.338 | 0.009 | 0.033 |

| | | | |
|---|---|---|---|
| Ccne1 | 1.382 | 0.000 | 0.000 |
| Cdkn1a | 1.297 | 0.000 | 0.001 |
| E2f1 | 1.415 | 0.000 | 0.000 |
| E2f7 | 2.234 | 0.000 | 0.000 |
| Tgfb3 | 2.307 | 0.000 | 0.000 |
| Brca2 | 0.731 | 0.000 | 0.002 |

**Hepatic Fibrosis/Hepatic Stellate Cell Activation**

| Gene name | Log2 fold change | P value | Ajusted P value |
|---|---|---|---|
| Igfbp4 | -1.858 | 0.001 | 0.007 |
| Fn1 | 4.053 | 0.000 | 0.000 |
| Tgfbr1 | -1.146 | 0.000 | 0.000 |
| Il1rl1 | -3.421 | 0.000 | 0.000 |
| Fgf2 | 3.595 | 0.000 | 0.000 |
| Mmp13 | 5.252 | 0.001 | 0.007 |
| Myl6b | 2.954 | 0.000 | 0.000 |
| Col15a1 | -1.244 | 0.000 | 0.001 |
| Bcl2 | -0.930 | 0.000 | 0.000 |
| Tgfbr2 | -0.769 | 0.004 | 0.017 |
| Il1r2 | 2.275 | 0.001 | 0.006 |
| Col1a2 | -3.621 | 0.001 | 0.004 |
| Timp1 | 4.484 | 0.012 | 0.042 |
| Serpine1 | 3.436 | 0.001 | 0.005 |
| Col18a1 | 4.162 | 0.000 | 0.001 |
| Klf12 | 3.458 | 0.000 | 0.000 |
| Stat1 | -0.777 | 0.000 | 0.001 |
| Pdgfrb | 5.579 | 0.000 | 0.000 |
| Egfr | 1.918 | 0.011 | 0.040 |
| Il4ra | -0.999 | 0.000 | 0.003 |
| Col5a2 | 3.762 | 0.006 | 0.026 |
| Tnfrsf1a | -0.702 | 0.003 | 0.015 |
| Il6ra | -0.890 | 0.000 | 0.001 |
| Smad7 | -1.115 | 0.002 | 0.011 |
| Ifngr2 | -1.201 | 0.000 | 0.000 |
| Vegfc | 3.422 | 0.000 | 0.000 |
| Bax | 0.613 | 0.000 | 0.000 |
| Il1r1 | 3.713 | 0.000 | 0.000 |
| Tlr4 | 2.647 | 0.000 | 0.000 |
| Col5a3 | 7.893 | 0.000 | 0.000 |
| Csf1 | 2.308 | 0.002 | 0.009 |
| Col23a1 | -1.362 | 0.008 | 0.032 |
| Tgfb3 | 2.307 | 0.000 | 0.000 |
| Tnf | -0.823 | 0.002 | 0.009 |
| Ccr7 | -0.932 | 0.000 | 0.002 |

#### Supplementary Table 6. CDR3 sequences for Treg cells and CD4$^{+}$ T cells from periphery

##### Treg WT1

| Sample Name | CDR3 Sequence | clone Count | best V Hit | best D Hit | best J Hit | Frequency |
|---|---|---|---|---|---|---|
| Treg WT1 | TGTGCCAGCAGTTTGCGGGGGGATGCAGAAACGCTGTATTTT | 4 | TRBV14*00 | TRBD2*00 | TRBJ2-3*00 | 1.10% |
| Treg WT1 | TGCACCTGCAGTGCAGACACAGAAGTCTTCTTT | 2 | TRBV1*00 | | TRBJ1-1*00 | 0.55% |
| Treg WT1 | TGTAGTTCTAGAGAGGGGGCCAACGAAAGATTATTTTTC | 2 | TRBV30*00 | TRBD1*00 | TRBJ1-4*00 | 0.55% |
| Treg WT1 | TGTGCAAGCAGCTTAGGGGGCAACCAGGCTCCGCTTTTT | 2 | TRBV16*00 | | TRBJ1-5*00 | 0.55% |
| Treg WT1 | TGTGCCAGCAGACAGGGGAGAGGTGCAGAAACGCTGTATTTT | 2 | TRBV15*00 | TRBD1*00 | TRBJ2-3*00 | 0.55% |
| Treg WT1 | TGTGCCAGCAGCCAGGGTTCTGGAAATACGCTCTATTTT | 2 | TRBV3*00 | TRBD1*00 | TRBJ1-3*00 | 0.55% |
| Treg WT1 | TGTGCCAGCAGTACGGTTAACCAAGACACCCAGTACTTT | 2 | TRBV26*00 | | TRBJ2-5*00 | 0.55% |
| Treg WT1 | TGTGCCAGCAGTATTCGTGAAACGCTGTATTTT | 2 | TRBV19*00 | | TRBJ2-3*00 | 0.55% |
| Treg WT1 | TGTGCCAGCAGTCAGGATGCAAACTCCGACTACACCTTC | 2 | TRBV14*00 | | TRBJ1-2*00 | 0.55% |
| Treg WT1 | TGTGCCAGCAGTTTTGGGGGGTCTTATGAACAGTACTTC | 2 | TRBV15*00 | TRBD2*00 | TRBJ2-7*00 | 0.55% |
| Treg WT1 | TGTGCCAGCGGTGATAGGAACTATGCTGAGCAGTTCTTC | 2 | TRBV13-2*00 | | TRBJ2-1*00 | 0.55% |
| Treg WT1 | TGTGGTGCCAACAGGGGAAACACAGAAGTCTTCTTT | 2 | TRBV20*00 | TRBD1*00 | TRBJ1-1*00 | 0.55% |
| Treg WT1 | TGTGGTGCTAGGGATCAGGCAAACTCCGACTACACCTTC | 2 | TRBV20*00 | | TRBJ1-2*00 | 0.55% |
| Treg WT1 | TGCACCTGCAGTAAACAGGGGTATGAACAGTACTTC | 1 | TRBV1*00 | TRBD1*00 | TRBJ2-7*00 | 0.28% |
| Treg WT1 | TGCACCTGCAGTAAACAGGGGTATGAACAGTACTTT | 1 | TRBV1*00 | TRBD1*00 | TRBJ2-7*00 | 0.28% |
| Treg WT1 | TGCACCTGCAGTACAGGGGGCCAAGACACCCAGTACTTT | 1 | TRBV1*00 | | TRBJ2-5*00 | 0.28% |
| Treg WT1 | TGCACCTGCAGTCCCAGACTGGGAAGCTCCTATGAACAGTACTTC | 1 | TRBV1*00 | TRBD2*00 | TRBJ2-7*00 | 0.28% |
| Treg WT1 | TGCACCTGCAGTCGGGGGTATGAACAGTACTTC | 1 | TRBV1*00 | TRBD1*00 | TRBJ2-7*00 | 0.28% |
| Treg WT1 | TGCACCTGCAGTGACAGGGGGAACCAAGACACCCAGTACTTT | 1 | TRBV1*00 | TRBD1*00 | TRBJ2-5*00 | 0.28% |
| Treg WT1 | TGCACCTGCAGTGACAGGGGGCGGGAAAGATTATTTTTC | 1 | TRBV1*00 | TRBD1*00 | TRBJ1-4*00 | 0.28% |
| Treg WT1 | TGCACCTGCAGTGAGACGGGTTCTGAGCAGTTCTTC | 1 | TRBV1*00 | | TRBJ2-1*00 | 0.28% |
| Treg WT1 | TGCACCTGCAGTGCAGACAGGGAAGTCTTCTTT | 1 | TRBV1*00 | | TRBJ1-1*00 | 0.28% |
| Treg WT1 | TGCACCTGCAGTGCAGACATCCAGGCTCCGCTTTTT | 1 | TRBV1*00 | | TRBJ1-5*00 | 0.28% |
| Treg WT1 | TGCACCTGCAGTGCAGACGGTAACCAGGCTCCGCTTTTT | 1 | TRBV1*00 | | TRBJ1-5*00 | 0.28% |
| Treg WT1 | TGCACCTGCAGTGCAGAGGGAGACACCCAGTACTTT | 1 | TRBV1*00 | | TRBJ2-5*00 | 0.28% |
| Treg WT1 | TGCACCTGCAGTGCAGATCAACAGACCGCCTTGTACTTT | 1 | TRBV1*00 | | TRBJ2-4*00 | 0.28% |
| Treg WT1 | TGCACCTGCAGTGCAGGAGACAGGGGTGACACCCAGTACTTT | 1 | TRBV1*00 | TRBD1*00 | TRBJ2-5*00 | 0.28% |
| Treg WT1 | TGCACCTGCAGTGCAGGGACAGGGGGCAGCGAAAGATTATTTTTC | 1 | TRBV1*00 | TRBD1*00 | TRBJ1-4*00 | 0.28% |
| Treg WT1 | TGCACCTGCAGTGCAGTCGGGACAGGGGTTGCTGAGCAGTTCTTC | 1 | TRBV1*00 | TRBD1*00 | TRBJ2-1*00 | 0.28% |
| Treg WT1 | TGCACCTGCAGTGCAGTGGGACAACAAGACACCCAGTACTTT | 1 | TRBV1*00 | TRBD1*00 | TRBJ2-5*00 | 0.28% |
| Treg WT1 | TGCACCTGCAGTGCGCCCCTTTATGCTGAGCAGTTCTTC | 1 | TRBV1*00 | TRBD1*00 | TRBJ2-1*00 | 0.28% |
| Treg WT1 | TGCACCTGCAGTGCGGACAGGGAAGTCTTCTTT | 1 | TRBV1*00 | TRBD1*00 | TRBJ1-1*00 | 0.28% |
| Treg WT1 | TGCACCTGCAGTGCGGGGAACCAAGACACCCAGTACTTT | 1 | TRBV1*00 | | TRBJ2-5*00 | 0.28% |
| Treg WT1 | TGCACCTGCAGTGGACGGGACAATGAGCAGTTCTTC | 1 | TRBV1*00 | TRBD1*00 | TRBJ2-1*00 | 0.28% |
| Treg WT1 | TGCACCTGCAGTGGGACTTCTTCTGGAAATACGCTCTATTTT | 1 | TRBV1*00 | TRBD2*00 | TRBJ1-3*00 | 0.28% |
| Treg WT1 | TGCACCTGCAGTGGGGGACAGCAGGCTCCGCTTTTT | 1 | TRBV1*00 | TRBD1*00 | TRBJ1-5*00 | 0.28% |
| Treg WT1 | TGCTCCAGCAGTCAATCGACTGGGGGAGACACCCAGTACTTT | 1 | TRBV23*00 | TRBD2*00 | TRBJ2-5*00 | 0.28% |
| Treg WT1 | TGCTCCAGCAGTCAATCTGGCAGCTATGAACAGTACTTC | 1 | TRBV23*00 | | TRBJ2-7*00 | 0.28% |
| Treg WT1 | TGTAGTTCTAGACGACAGAATGCAAACTCCGACTACACCTTC | 1 | TRBV30*00 | TRBD1*00 | TRBJ1-2*00 | 0.28% |
| Treg WT1 | TGTGCAAGCAGCCCTACAGCTAGTCAAAACACCTTGTACTTT | 1 | TRBV16*00 | | TRBJ2-4*00 | 0.28% |
| Treg WT1 | TGTGCAAGCAGCCTGAGGGGGGAGGACACCCAGTACTTT | 1 | TRBV16*00 | TRBD1*00 | TRBJ2-5*00 | 0.28% |
| Treg WT1 | TGTGCAAGCAGCTATGAACAGTACTTC | 1 | TRBV16*00 | | TRBJ2-7*00 | 0.28% |
| Treg WT1 | TGTGCAAGCAGCTCCTACAGGGGCCAAAACACCTTGTACTTT | 1 | TRBV16*00 | TRBD1*00 | TRBJ2-4*00 | 0.28% |
| Treg WT1 | TGTGCAAGCAGCTTAAACAAAAACTCCGACTACACCTTC | 1 | TRBV16*00 | | TRBJ1-2*00 | 0.28% |
| Treg WT1 | TGTGCAAGCAGCTTAAAGGGAGACACCCAGTACTTT | 1 | TRBV16*00 | | TRBJ2-5*00 | 0.28% |
| Treg WT1 | TGTGCAAGCAGCTTAAGAGGGGGCACCTCCGACTACACCTTC | 1 | TRBV16*00 | TRBD1*00 | TRBJ1-2*00 | 0.28% |
| Treg WT1 | TGTGCAAGCAGCTTACAAAACACAGAAGTCTTCTTT | 1 | TRBV16*00 | | TRBJ1-1*00 | 0.28% |
| Treg WT1 | TGTGCAAGCAGCTTACAGGGGGCAGGCAATGAACAGTACTTC | 1 | TRBV16*00 | TRBD1*00 | TRBJ2-7*00 | 0.28% |
| Treg WT1 | TGTGCAAGCAGCTTAGAGGCAGGGAACACCTTGTACTTT | 1 | TRBV16*00 | TRBD1*00 | TRBJ2-4*00 | 0.28% |
| Treg WT1 | TGTGCAAGCAGCTTAGAGGGACAGGGGGACACCCAGTACTTT | 1 | TRBV16*00 | TRBD1*00 | TRBJ2-5*00 | 0.28% |
| Treg WT1 | TGTGCAAGCAGCTTAGAGGGGGGCGGAGACACCCAGTACTTT | 1 | TRBV16*00 | TRBD2*00 | TRBJ2-5*00 | 0.28% |
| Treg WT1 | TGTGCAAGCAGCTTAGATGGGCTGGGGTATGAACAGTACTTC | 1 | TRBV16*00 | TRBD2*00 | TRBJ2-7*00 | 0.28% |
| Treg WT1 | TGTGCAAGCAGCTTAGCTGGGGACTATGCTGAGCAGTTCTTC | 1 | TRBV16*00 | TRBD2*00 | TRBJ2-1*00 | 0.28% |
| Treg WT1 | TGTGCAAGCAGCTTAGGACAACACACAGAAGTCTTCTTT | 1 | TRBV16*00 | | TRBJ1-1*00 | 0.28% |
| Treg WT1 | TGTGCAAGCAGCTTTTGGGGGGGCGATGAACAGTACTTC | 1 | TRBV16*00 | TRBD2*00 | TRBJ2-7*00 | 0.28% |
| Treg WT1 | TGTGCCAAGAATCAAGACACCCAGTACTTT | 1 | TRBV16*00 | | TRBJ2-5*00 | 0.28% |
| Treg WT1 | TGTGCCAGCAAGGGACAGCAAGACACCCAGTACTTT | 1 | TRBV13-3*00 | TRBD1*00 | TRBJ2-5*00 | 0.28% |
| Treg WT1 | TGTGCCAGCACCACTGGGGACTATGCTGAGCAGTTCTTC | 1 | TRBV13-1*00 | TRBD2*00 | TRBJ2-1*00 | 0.28% |
| Treg WT1 | TGTGCCAGCACCCGGGCCGGGACTAGGGAACAGTACTTC | 1 | TRBV15*00 | TRBD2*00 | TRBJ2-7*00 | 0.28% |
| Treg WT1 | TGTGCCAGCACCTCAGGGGATTCCGACTACACCTTC | 1 | TRBV15*00 | TRBD1*00 | TRBJ1-2*00 | 0.28% |
| Treg WT1 | TGTGCCAGCACGGACTGGGGGTCTAGTGCAGAAACGCTGTATTTT | 1 | TRBV26*00 | TRBD2*00 | TRBJ2-3*00 | 0.28% |
| Treg WT1 | TGTGCCAGCAGAACAAACTCCGACTACACCTTC | 1 | TRBV14*00 | | TRBJ1-2*00 | 0.28% |
| Treg WT1 | TGTGCCAGCAGACAGGATTCTGGAAATACGCTCTATTTT | 1 | TRBV13-1*00 | | TRBJ1-3*00 | 0.28% |
| Treg WT1 | TGTGCCAGCAGACTGGGGGGGCGAGACACCCAGTACTTT | 1 | TRBV14*00 | TRBD2*00 | TRBJ2-5*00 | 0.28% |
| Treg WT1 | TGTGCCAGCAGATACGAATATGCTGAGCAGTTCTTC | 1 | TRBV15*00 | | TRBJ2-1*00 | 0.28% |
| Treg WT1 | TGTGCCAGCAGATTTGGTGCAGAAACGCTGTATTTT | 1 | TRBV3*00 | | TRBJ2-3*00 | 0.28% |
| Treg WT1 | TGTGCCAGCAGCAAGCGGGTAAACTCCGACTACACCTTC | 1 | TRBV3*00 | | TRBJ1-2*00 | 0.28% |
| Treg WT1 | TGTGCCAGCAGCACCGGGACAGGGGGACAGGCTCCGCTTTTT | 1 | TRBV4*00 | TRBD1*00 | TRBJ1-5*00 | 0.28% |
| Treg WT1 | TGTGCCAGCAGCACCTGGGGGGGAGAAACGCTGTATTTT | 1 | TRBV4*00 | TRBD2*00 | TRBJ2-3*00 | 0.28% |
| Treg WT1 | TGTGCCAGCAGCATCGACAGGAACTATGCTGAGCAGTTCTTC | 1 | TRBV2*00 | TRBD1*00 | TRBJ2-1*00 | 0.28% |

| | | | | | | |
|---|---|---|---|---|---|---|
| Treg WT1 | TGTGCCAGCAGCCAACCGGGACAAAGCTCCTATGAACAGTACTTC | 1 | TRBV5*00 | TRBD1*00 | TRBJ2-7*00 | 0.28% |
| Treg WT1 | TGTGCCAGCAGCCAAGAAGGAGACACCCAGTACTTT | 1 | TRBV5*00 | | TRBJ2-5*00 | 0.28% |
| Treg WT1 | TGTGCCAGCAGCCAAGAAGGGTATAACTATGCTGAGCAGTTCTTC | 1 | TRBV5*00 | | TRBJ2-1*00 | 0.28% |
| Treg WT1 | TGTGCCAGCAGCCAAGACATCTATGCTGAGCAGTTCTTC | 1 | TRBV5*00 | | TRBJ2-1*00 | 0.28% |
| Treg WT1 | TGTGCCAGCAGCCAAGACGGCTGGGGGGCAGACACCCAGTACTTT | 1 | TRBV5*00 | TRBD2*00 | TRBJ2-5*00 | 0.28% |
| Treg WT1 | TGTGCCAGCAGCCAAGAGACAATACAGTACTTC | 1 | TRBV5*00 | | TRBJ2-7*00 | 0.28% |
| Treg WT1 | TGTGCCAGCAGCCAAGAGGGGGGTAATTCTGGAAATACGCTCTATTTT | 1 | TRBV2*00 | TRBD2*00 | TRBJ1-3*00 | 0.28% |
| Treg WT1 | TGTGCCAGCAGCCAAGGCTCAAACACAGAAGTCTTCTTT | 1 | TRBV2*00 | | TRBJ1-1*00 | 0.28% |
| Treg WT1 | TGTGCCAGCAGCCAAGGGGACAGTGCAGAAACGCTGTATTTT | 1 | TRBV2*00 | TRBD1*00 | TRBJ2-3*00 | 0.28% |
| Treg WT1 | TGTGCCAGCAGCCAAGGGGGCCCAGAAGTCTTCTTT | 1 | TRBV2*00 | TRBD1*00 | TRBJ1-1*00 | 0.28% |
| Treg WT1 | TGTGCCAGCAGCCAAGTAACTGGGGGGGGTGCTGAGCAGTTCTTC | 1 | TRBV5*00 | TRBD2*00 | TRBJ2-1*00 | 0.28% |
| Treg WT1 | TGTGCCAGCAGCCAAGTCCCATCTAACTATGCTGAGCAGTTCTTC | 1 | TRBV2*00 | | TRBJ2-1*00 | 0.28% |
| Treg WT1 | TGTGCCAGCAGCCAATGGGGGGATAGTGCAGAAACGCTGTATTTT | 1 | TRBV5*00 | TRBD2*00 | TRBJ2-3*00 | 0.28% |
| Treg WT1 | TGTGCCAGCAGCCACAGGGGGGGGCACAGAAGTCTTCTTT | 1 | TRBV2*00 | TRBD1*00 | TRBJ1-1*00 | 0.28% |
| Treg WT1 | TGTGCCAGCAGCCACTCCTATGAACAGTACTTC | 1 | TRBV15*00 | | TRBJ2-7*00 | 0.28% |
| Treg WT1 | TGTGCCAGCAGCCAGACCGGGGCTGAGCAGTTCTTC | 1 | TRBV5*00 | TRBD1*00 | TRBJ2-1*00 | 0.28% |
| Treg WT1 | TGTGCCAGCAGCCAGACTGGAGCAGAAACGCTGTATTTT | 1 | TRBV5*00 | TRBD2*00 | TRBJ2-3*00 | 0.28% |
| Treg WT1 | TGTGCCAGCAGCCAGACTGGGGGGAAAGACACCCAGTACTTT | 1 | TRBV2*00 | TRBD2*00 | TRBJ2-5*00 | 0.28% |
| Treg WT1 | TGTGCCAGCAGCCAGGGGGGCGCCACAGAAGTCTTCTTT | 1 | TRBV5*00 | TRBD2*00 | TRBJ1-1*00 | 0.28% |
| Treg WT1 | TGTGCCAGCAGCCAGTACAGAAACACAGAAGTCTTCTTT | 1 | TRBV5*00 | | TRBJ1-1*00 | 0.28% |
| Treg WT1 | TGTGCCAGCAGCCCCACGGGGGGCTATGCTGAGCAGTTCTTC | 1 | TRBV5*00 | TRBD2*00 | TRBJ2-1*00 | 0.28% |
| Treg WT1 | TGTGCCAGCAGCCCCCACCTGGGGGCTAGTGCAGAAACGCTGTATTTT | 1 | TRBV15*00 | TRBD2*00 | TRBJ2-3*00 | 0.28% |
| Treg WT1 | TGTGCCAGCAGCCCCCAGGGGTATGCTGAGCAGTTCTTC | 1 | TRBV2*00 | TRBD1*00 | TRBJ2-1*00 | 0.28% |
| Treg WT1 | TGTGCCAGCAGCCCCCGGGACATCCAAGACACCCAGTACTTT | 1 | TRBV5*00 | TRBD1*00 | TRBJ2-5*00 | 0.28% |
| Treg WT1 | TGTGCCAGCAGCCCCCTCTCCTATGAACAGTACTTC | 1 | TRBV4*00 | TRBD1*00 | TRBJ2-7*00 | 0.28% |
| Treg WT1 | TGTGCCAGCAGCCCCTCTGGGATATATGAACAGTACTTC | 1 | TRBV2*00 | TRBD2*00 | TRBJ2-7*00 | 0.28% |
| Treg WT1 | TGTGCCAGCAGCCCGGGACAAATCTATGAACAGTACTTC | 1 | TRBV5*00 | TRBD1*00 | TRBJ2-7*00 | 0.28% |
| Treg WT1 | TGTGCCAGCAGCCCGGGACAGGGGACCTCCGACTACACCCTC | 1 | TRBV13-3*00 | TRBD1*00 | TRBJ1-2*00 | 0.28% |
| Treg WT1 | TGTGCCAGCAGCCCTAGTGCAGAAACGCTGTATTTT | 1 | TRBV13-3*00 | | TRBJ2-3*00 | 0.28% |
| Treg WT1 | TGTGCCAGCAGCCCTCCGGGGGGGATAAACACCGGGCAGCTCTACTTT | 1 | TRBV5*00 | TRBD2*00 | TRBJ2-2*00 | 0.28% |
| Treg WT1 | TGTGCCAGCAGCCGACATGCAAACTCCGACTACACCTTC | 1 | TRBV3*00 | | TRBJ1-2*00 | 0.28% |
| Treg WT1 | TGTGCCAGCAGCCGACTGGGGATTGAACAGTACTTC | 1 | TRBV15*00 | TRBD2*00 | TRBJ2-7*00 | 0.28% |
| Treg WT1 | TGTGCCAGCAGCCGACTGGGGGGGCGCGACCAAGACACCCAGTACTTT | 1 | TRBV4*00 | TRBD2*00 | TRBJ2-5*00 | 0.28% |
| Treg WT1 | TGTGCCAGCAGCCGTCAAAACACCTTGTACTTT | 1 | TRBV3*00 | | TRBJ2-4*00 | 0.28% |
| Treg WT1 | TGTGCCAGCAGCCTAGTGCAGAAACGCTGTATTTT | 1 | TRBV26*00 | | TRBJ2-3*00 | 0.28% |
| Treg WT1 | TGTGCCAGCAGCCTCAAGAGTCAAAACACCTTGTACTTT | 1 | TRBV2*00 | | TRBJ2-4*00 | 0.28% |
| Treg WT1 | TGTGCCAGCAGCCTCACAGGAGCTCCTATGAACAGTACTTC | 1 | TRBV19*00 | | TRBJ2-7*00 | 0.28% |
| Treg WT1 | TGTGCCAGCAGCCTCGGGACTGGGGTTGCAGAAACGCTGTATTTT | 1 | TRBV5*00 | TRBD2*00 | TRBJ2-3*00 | 0.28% |
| Treg WT1 | TGTGCCAGCAGCGGACAAACAGAAGTCTTCTTT | 1 | TRBV15*00 | TRBD1*00 | TRBJ1-1*00 | 0.28% |
| Treg WT1 | TGTGCCAGCAGCGTCACAGCAAACACCGGGCAGCTCTACTTT | 1 | TRBV2*00 | | TRBJ2-2*00 | 0.28% |
| Treg WT1 | TGTGCCAGCAGCTAAGAGGGGGGCTCCGCTTTTT | 1 | TRBV4*00 | | TRBJ1-5*00 | 0.28% |
| Treg WT1 | TGTGCCAGCAGCTATGGGGGGCTAGGAAATACGCTCTATTTT | 1 | TRBV4*00 | TRBD2*00 | TRBJ1-3*00 | 0.28% |
| Treg WT1 | TGTGCCAGCAGCTCAGAGGCTGCTGAGCAGTTCTTC | 1 | TRBV3*00 | | TRBJ2-1*00 | 0.28% |
| Treg WT1 | TGTGCCAGCAGCTCCAGGGGGAACTCCGACTACACCTTC | 1 | TRBV25*00 | TRBD1*00 | TRBJ1-2*00 | 0.28% |
| Treg WT1 | TGTGCCAGCAGCTCCCGACTGGGGGAAACACCTTGTACTTT | 1 | TRBV25*00 | TRBD2*00 | TRBJ2-4*00 | 0.28% |
| Treg WT1 | TGTGCCAGCAGCTCGACAGGGGGGTATGCTGAGCAGTTCTTC | 1 | TRBV4*00 | TRBD1*00 | TRBJ2-1*00 | 0.28% |
| Treg WT1 | TGTGCCAGCAGCTGGACAACTAGTGCAGAAACGCTGTATTTT | 1 | TRBV3*00 | TRBD1*00 | TRBJ2-3*00 | 0.28% |
| Treg WT1 | TGTGCCAGCAGCTGGACAGGGGGCACCGGGCAGCTCTACTTT | 1 | TRBV25*00 | TRBD1*00 | TRBJ2-2*00 | 0.28% |
| Treg WT1 | TGTGCCAGCAGCTGGACTGGGGGGGACTATGCTGAGCAGTTCTTC | 1 | TRBV3*00 | TRBD2*00 | TRBJ2-1*00 | 0.28% |
| Treg WT1 | TGTGCCAGCAGCTTAAACGAAAGATTATTTTTC | 1 | TRBV3*00 | | TRBJ1-4*00 | 0.28% |
| Treg WT1 | TGTGCCAGCAGCTTAAGGACAGGGAGGGGGTGCAGAAACGCTGTATTTT | 1 | TRBV3*00 | TRBD1*00 | TRBJ2-3*00 | 0.28% |
| Treg WT1 | TGTGCCAGCAGCTTAAGGGGACAGAACTCCGACTACACCTTC | 1 | TRBV3*00 | TRBD1*00 | TRBJ1-2*00 | 0.28% |
| Treg WT1 | TGTGCCAGCAGCTTAAGGGGGGGCTCCGACTACACCTTC | 1 | TRBV3*00 | TRBD2*00 | TRBJ1-2*00 | 0.28% |
| Treg WT1 | TGTGCCAGCAGCTTACAGGGGGCGCACACAGAAGTCTTCTTT | 1 | TRBV3*00 | TRBD1*00 | TRBJ1-1*00 | 0.28% |
| Treg WT1 | TGTGCCAGCAGCTTACGGGGGGGGAACACCTTGTACTTT | 1 | TRBV3*00 | TRBD2*00 | TRBJ2-4*00 | 0.28% |
| Treg WT1 | TGTGCCAGCAGCTTACTGGTTAACCAAGACACCCAGTACTTT | 1 | TRBV3*00 | | TRBJ2-5*00 | 0.28% |
| Treg WT1 | TGTGCCAGCAGCTTAGAGGACACCGGGCAGCTCTACTTT | 1 | TRBV3*00 | | TRBJ2-2*00 | 0.28% |
| Treg WT1 | TGTGCCAGCAGCTTAGAGGGACAGGGGAGAGTCTTCTTT | 1 | TRBV3*00 | TRBD1*00 | TRBJ1-1*00 | 0.28% |
| Treg WT1 | TGTGCCAGCAGCTTAGAGGGAGCAAACTCCGACTACACCTTC | 1 | TRBV3*00 | | TRBJ1-2*00 | 0.28% |
| Treg WT1 | TGTGCCAGCAGCTTAGATGGCAGCAACACCGGGCAGCTCTACTTT | 1 | TRBV3*00 | | TRBJ2-2*00 | 0.28% |
| Treg WT1 | TGTGCCAGCAGCTTAGCAGGCAGTAACTCCGACTACACCTTC | 1 | TRBV3*00 | | TRBJ1-2*00 | 0.28% |
| Treg WT1 | TGTGCCAGCAGCTTAGCCAGGGGGGAACAGTACTTC | 1 | TRBV3*00 | TRBD1*00 | TRBJ2-7*00 | 0.28% |
| Treg WT1 | TGTGCCAGCAGCTTAGCCGGGGGCGGCACAGAAGTCTTCTTT | 1 | TRBV3*00 | TRBD1*00 | TRBJ1-1*00 | 0.28% |
| Treg WT1 | TGTGCCAGCAGCTTAGCGAACTCCGACTACACCTTC | 1 | TRBV3*00 | | TRBJ1-2*00 | 0.28% |
| Treg WT1 | TGTGCCAGCAGCTTAGCGCCCGGGGACTATGCTGAGCAGTTCTTC | 1 | TRBV3*00 | TRBD1*00 | TRBJ2-1*00 | 0.28% |
| Treg WT1 | TGTGCCAGCAGCTTAGGACAGGGGTCAGAAGTCTTCTTT | 1 | TRBV3*00 | TRBD1*00 | TRBJ1-1*00 | 0.28% |
| Treg WT1 | TGTGCCAGCAGCTTAGGCGGGTATGAACAGTACTTC | 1 | TRBV3*00 | | TRBJ2-7*00 | 0.28% |
| Treg WT1 | TGTGCCAGCAGCTTAGGGCACAACGAAAGATTATTTTTC | 1 | TRBV3*00 | | TRBJ1-4*00 | 0.28% |
| Treg WT1 | TGTGCCAGCAGCTTAGGGGGGCCCAACGAAAGATTATTTTTC | 1 | TRBV3*00 | TRBD1*00 | TRBJ1-4*00 | 0.28% |
| Treg WT1 | TGTGCCAGCAGCTTAGTTAACTCCGACTACACCTTC | 1 | TRBV3*00 | | TRBJ1-2*00 | 0.28% |
| Treg WT1 | TGTGCCAGCAGCTTAGTTGGGGTCTGGGGTGAGCAGTTCTTC | 1 | TRBV3*00 | TRBD2*00 | TRBJ2-1*00 | 0.28% |
| Treg WT1 | TGTGCCAGCAGCTTATACGGGGGCGCGGACACCCAGTACTTT | 1 | TRBV3*00 | TRBD1*00 | TRBJ2-5*00 | 0.28% |
| Treg WT1 | TGTGCCAGCAGCTTATTACAGGAAGTCTTCTTT | 1 | TRBV3*00 | | TRBJ1-1*00 | 0.28% |
| Treg WT1 | TGTGCCAGCAGCTTCAGGGTTGCTGAGCAGTTCTTC | 1 | TRBV3*00 | TRBD1*00 | TRBJ2-1*00 | 0.28% |
| Treg WT1 | TGTGCCAGCAGCTTCCTCGTCAACGAAAGATTATTTTTC | 1 | TRBV3*00 | | TRBJ1-4*00 | 0.28% |

| | | | | | | |
|---|---|---|---|---|---|---|
| Treg WT1 | TGTGCCAGCAGCTTCTGGGGGGACCAAGACACCCAGTACTTT | 1 | TRBV3*00 | TRBD2*00 | TRBJ2-5*00 | 0.28% |
| Treg WT1 | TGTGCCAGCAGCTTGACAGGGGACGAAAGATTATTTTTC | 1 | TRBV4*00 | TRBD1*00 | TRBJ1-4*00 | 0.28% |
| Treg WT1 | TGTGCCAGCAGCTTGACAGGGGGCTCCTATGAACAGTACTTC | 1 | TRBV3*00 | TRBD1*00 | TRBJ2-7*00 | 0.28% |
| Treg WT1 | TGTGCCAGCAGCTTGAGAGGATATGAACAGTACTTC | 1 | TRBV3*00 | | TRBJ2-7*00 | 0.28% |
| Treg WT1 | TGTGCCAGCAGCTTGATGGGGGCAGTCTTCTTT | 1 | TRBV3*00 | TRBD1*00 | TRBJ1-1*00 | 0.28% |
| Treg WT1 | TGTGCCAGCAGCTTGGACAGGGACTCCGACTACACCTTC | 1 | TRBV3*00 | TRBD1*00 | TRBJ1-2*00 | 0.28% |
| Treg WT1 | TGTGCCAGCAGCTTGGACCTGGGCTCCTATGAACAGTACTTC | 1 | TRBV4*00 | | TRBJ2-7*00 | 0.28% |
| Treg WT1 | TGTGCCAGCAGCTTGGGGGGCACAGAAGTCTTCTTT | 1 | TRBV3*00 | TRBD2*00 | TRBJ1-1*00 | 0.28% |
| Treg WT1 | TGTGCCAGCAGCTTTCAAAACACCTTGTACTTT | 1 | TRBV3*00 | | TRBJ2-4*00 | 0.28% |
| Treg WT1 | TGTGCCAGCAGGGACATAAACACAGAAGTCTTCTTT | 1 | TRBV13-3*00 | TRBD1*00 | TRBJ1-1*00 | 0.28% |
| Treg WT1 | TGTGCCAGCAGGGATTGGAGTCAAAACACCTTGTACTTT | 1 | TRBV13-3*00 | | TRBJ2-4*00 | 0.28% |
| Treg WT1 | TGTGCCAGCAGGGGGACAGGGAGCTCCGACTACACCTTC | 1 | TRBV15*00 | TRBD1*00 | TRBJ1-2*00 | 0.28% |
| Treg WT1 | TGTGCCAGCAGTAACCGGGACAGGGGCGCTGAGCAGTTCTTC | 1 | TRBV22*00 | TRBD1*00 | TRBJ2-1*00 | 0.28% |
| Treg WT1 | TGTGCCAGCAGTAAGGAAACGCTGTATTTT | 1 | TRBV19*00 | | TRBJ2-3*00 | 0.28% |
| Treg WT1 | TGTGCCAGCAGTAGATGGGGGGGGGATGCTGAGCAGTTCTTC | 1 | TRBV19*00 | TRBD2*00 | TRBJ2-1*00 | 0.28% |
| Treg WT1 | TGTGCCAGCAGTAGGGGGGGCTCCGACTACACCTTC | 1 | TRBV13-3*00 | TRBD2*00 | TRBJ1-2*00 | 0.28% |
| Treg WT1 | TGTGCCAGCAGTATAACGGCATGGGGCAACTCCGACTACACCTTC | 1 | TRBV19*00 | TRBD1*00 | TRBJ1-2*00 | 0.28% |
| Treg WT1 | TGTGCCAGCAGTATAAGGGGGGCTAGTGCAGAAACGCTGTATTTT | 1 | TRBV19*00 | TRBD1*00 | TRBJ2-3*00 | 0.28% |
| Treg WT1 | TGTGCCAGCAGTATAGAGGGGGCGGACACCCAGTACTTT | 1 | TRBV19*00 | TRBD1*00 | TRBJ2-5*00 | 0.28% |
| Treg WT1 | TGTGCCAGCAGTATCCCGACAGGGGGGGGCACAGAAGTCTTCTTT | 1 | TRBV19*00 | TRBD1*00 | TRBJ1-1*00 | 0.28% |
| Treg WT1 | TGTGCCAGCAGTATGGGAAACAACCAGGCTCCGCTTTTT | 1 | TRBV19*00 | | TRBJ1-5*00 | 0.28% |
| Treg WT1 | TGTGCCAGCAGTATTACGACAGACTATGCTGAGCAGTTCTTC | 1 | TRBV19*00 | TRBD1*00 | TRBJ2-1*00 | 0.28% |
| Treg WT1 | TGTGCCAGCAGTCCACTGGGGGGTGAACAGTACTTC | 1 | TRBV24*00 | TRBD2*00 | TRBJ2-7*00 | 0.28% |
| Treg WT1 | TGTGCCAGCAGTCCAGACAAGGACTCCGACTACACCTTC | 1 | TRBV14*00 | | TRBJ1-2*00 | 0.28% |
| Treg WT1 | TGTGCCAGCAGTCCAGGGGACTCTAGTGCAGAAACGCTGTATTTT | 1 | TRBV19*00 | TRBD1*00 | TRBJ2-3*00 | 0.28% |
| Treg WT1 | TGTGCCAGCAGTCTACACACAGAAGTCTTCTTT | 1 | TRBV26*00 | | TRBJ1-1*00 | 0.28% |
| Treg WT1 | TGTGCCAGCAGTCTACTGGGCCAAGACACCCAGTACTTT | 1 | TRBV13-1*00 | TRBD2*00 | TRBJ2-5*00 | 0.28% |
| Treg WT1 | TGTGCCAGCAGTCTCCGGGACACACAGTACTTC | 1 | TRBV26*00 | TRBD1*00 | TRBJ2-7*00 | 0.28% |
| Treg WT1 | TGTGCCAGCAGTCTCGACCCCCGCTCCTATGAACAGTACTTC | 1 | TRBV26*00 | TRBD1*00 | TRBJ2-7*00 | 0.28% |
| Treg WT1 | TGTGCCAGCAGTCTGAACAGGCAAAACACCTTC | 1 | TRBV26*00 | TRBD1*00 | TRBJ1-2*00 | 0.28% |
| Treg WT1 | TGTGCCAGCAGTCTGAGGAACTCCGACTACACCTTC | 1 | TRBV13-1*00 | | TRBJ1-2*00 | 0.28% |
| Treg WT1 | TGTGCCAGCAGTCTGGACAGATATGAACAGTACTTC | 1 | TRBV26*00 | TRBD1*00 | TRBJ2-7*00 | 0.28% |
| Treg WT1 | TGTGCCAGCAGTCTGTATGAACAGTACTTC | 1 | TRBV24*00 | | TRBJ2-7*00 | 0.28% |
| Treg WT1 | TGTGCCAGCAGTCTGTCCTATGAACAGTACTTC | 1 | TRBV26*00 | | TRBJ2-7*00 | 0.28% |
| Treg WT1 | TGTGCCAGCAGTCTGTCGGCCAGTCGGAACACCCAGTACTTT | 1 | TRBV26*00 | TRBD2*00 | TRBJ2-5*00 | 0.28% |
| Treg WT1 | TGTGCCAGCAGTCTGTCGGGAAACCAAGACACCCAGTACTTT | 1 | TRBV26*00 | | TRBJ2-5*00 | 0.28% |
| Treg WT1 | TGTGCCAGCAGTCTGTGGAGGGAGGAAGTCTTCTTT | 1 | TRBV26*00 | | TRBJ1-1*00 | 0.28% |
| Treg WT1 | TGTGCCAGCAGTCTGTTCAGGGGGAACTCCGACTACACCTTC | 1 | TRBV24*00 | TRBD1*00 | TRBJ1-2*00 | 0.28% |
| Treg WT1 | TGTGCCAGCAGTCTTCAGGGGGCAAACTCCGACTACACCTTC | 1 | TRBV24*00 | TRBD1*00 | TRBJ1-2*00 | 0.28% |
| Treg WT1 | TGTGCCAGCAGTGAAACTGGGGGGTGGTATGCTGAGCAGTTCTTC | 1 | TRBV13-1*00 | TRBD2*00 | TRBJ2-1*00 | 0.28% |
| Treg WT1 | TGTGCCAGCAGTGAAGCCCCGACAGACACCCAGTACTTT | 1 | TRBV13-3*00 | TRBD1*00 | TRBJ2-5*00 | 0.28% |
| Treg WT1 | TGTGCCAGCAGTGAATGGACTGGGGGGTTATGAACAGTACTTC | 1 | TRBV13-1*00 | TRBD2*00 | TRBJ2-7*00 | 0.28% |
| Treg WT1 | TGTGCCAGCAGTGACAATCAAAACACCTTGTACTTT | 1 | TRBV13-1*00 | | TRBJ2-4*00 | 0.28% |
| Treg WT1 | TGTGCCAGCAGTGACAATTATGCTGAGCAGTTCTTC | 1 | TRBV13-3*00 | | TRBJ2-1*00 | 0.28% |
| Treg WT1 | TGTGCCAGCAGTGAGAGACTGGTTAACCAAGACACCCAGTACTTT | 1 | TRBV13-1*00 | | TRBJ2-5*00 | 0.28% |
| Treg WT1 | TGTGCCAGCAGTGAGCTGACAAACTCCGACTACACCTTC | 1 | TRBV13-1*00 | | TRBJ1-2*00 | 0.28% |
| Treg WT1 | TGTGCCAGCAGTGAGGACAGGGGGCGGGACACCCAGTACTTT | 1 | TRBV13-1*00 | TRBD1*00 | TRBJ2-5*00 | 0.28% |
| Treg WT1 | TGTGCCAGCAGTGAGGGACAATTTAACCAAGACACCCAGTACTTT | 1 | TRBV13-3*00 | TRBD1*00 | TRBJ2-5*00 | 0.28% |
| Treg WT1 | TGTGCCAGCAGTGAGGGGGGGACCGACTACACCTTC | 1 | TRBV13-1*00 | TRBD2*00 | TRBJ1-2*00 | 0.28% |
| Treg WT1 | TGTGCCAGCAGTGATAACAATTATGAACAGTACTTC | 1 | TRBV13-3*00 | | TRBJ2-7*00 | 0.28% |
| Treg WT1 | TGTGCCAGCAGTGATAACTGGGGAACAGAAACGCTGTATTTT | 1 | TRBV13-1*00 | TRBD2*00 | TRBJ2-3*00 | 0.28% |
| Treg WT1 | TGTGCCAGCAGTGATACTGGGGGGGAAACGCTGTATTTT | 1 | TRBV13-1*00 | TRBD2*00 | TRBJ2-3*00 | 0.28% |
| Treg WT1 | TGTGCCAGCAGTGATAGGACAGGGTACACAGAAGTCTTCTTT | 1 | TRBV13-1*00 | TRBD1*00 | TRBJ1-1*00 | 0.28% |
| Treg WT1 | TGTGCCAGCAGTGATAGGGGTACTAGTGCAGAAACGCTGTATTTT | 1 | TRBV13-3*00 | TRBD1*00 | TRBJ2-3*00 | 0.28% |
| Treg WT1 | TGTGCCAGCAGTGATCAGGGTTCTGGAAATACGCTCTATTTT | 1 | TRBV13-1*00 | TRBD1*00 | TRBJ1-3*00 | 0.28% |
| Treg WT1 | TGTGCCAGCAGTGATCCGGGGGGGCGTGCAGAAACGCTGTATTTT | 1 | TRBV13-3*00 | TRBD2*00 | TRBJ2-3*00 | 0.28% |
| Treg WT1 | TGTGCCAGCAGTGATGACAACTCCTATGAACAGTACTTC | 1 | TRBV13-3*00 | | TRBJ2-7*00 | 0.28% |
| Treg WT1 | TGTGCCAGCAGTGATGACAGATACACAGAAGTCTTCTTT | 1 | TRBV13-1*00 | | TRBJ1-1*00 | 0.28% |
| Treg WT1 | TGTGCCAGCAGTGATGACAGTCAGGCTCCGCTTTTT | 1 | TRBV13-1*00 | | TRBJ1-5*00 | 0.28% |
| Treg WT1 | TGTGCCAGCAGTGATGAGAGGGTAAACACCTTGTACTTT | 1 | TRBV13-1*00 | | TRBJ2-4*00 | 0.28% |
| Treg WT1 | TGTGCCAGCAGTGATGCAGGACAGGGCACAGAAGTCTTCTTT | 1 | TRBV13-1*00 | TRBD1*00 | TRBJ1-1*00 | 0.28% |
| Treg WT1 | TGTGCCAGCAGTGATGCATTTAGTCAAAACACCTTGTACTTT | 1 | TRBV13-1*00 | | TRBJ2-4*00 | 0.28% |
| Treg WT1 | TGTGCCAGCAGTGATGCTGGGGAAAACACCTTGTACTTT | 1 | TRBV13-1*00 | TRBD2*00 | TRBJ2-4*00 | 0.28% |
| Treg WT1 | TGTGCCAGCAGTGATGCTGGGGAAGTCTTCTTT | 1 | TRBV13-1*00 | | TRBJ1-1*00 | 0.28% |
| Treg WT1 | TGTGCCAGCAGTGATGGAGGAACAGTACTTC | 1 | TRBV13-3*00 | | TRBJ2-7*00 | 0.28% |
| Treg WT1 | TGTGCCAGCAGTGATGGGACAAACACCGGGCAGCTCTACTTT | 1 | TRBV13-1*00 | | TRBJ2-2*00 | 0.28% |
| Treg WT1 | TGTGCCAGCAGTGATGGAGGAACTCCGACTACACCTTC | 1 | TRBV13-3*00 | | TRBJ1-2*00 | 0.28% |
| Treg WT1 | TGTGCCAGCAGTGATGGGGGGGACACCCAGTACTTT | 1 | TRBV13-3*00 | TRBD1*00 | TRBJ2-5*00 | 0.28% |
| Treg WT1 | TGTGCCAGCAGTGATGTACAGTATGCTGAGCAGTTCTTC | 1 | TRBV13-1*00 | | TRBJ2-1*00 | 0.28% |
| Treg WT1 | TGTGCCAGCAGTGATTACAACTATGCTGAGCAGTTCTTC | 1 | TRBV13-1*00 | | TRBJ2-1*00 | 0.28% |
| Treg WT1 | TGTGCCAGCAGTGCACAGGGCGGTAATTCGCCCCTCTACTTT | 1 | TRBV13-1*00 | TRBD1*00 | TRBJ1-6*00 | 0.28% |
| Treg WT1 | TGTGCCAGCAGTGCCGATGTCCAGTATAACTATGCTGAGCAGTTCTTC | 1 | TRBV13-1*00 | TRBD1*00 | TRBJ2-1*00 | 0.28% |
| Treg WT1 | TGTGCCAGCAGTGGAATTTATGAACAGTACTTC | 1 | TRBV13-1*00 | | TRBJ2-7*00 | 0.28% |
| Treg WT1 | TGTGCCAGCAGTGGACAAAGCTCCTATGAACAGTACTTC | 1 | TRBV13-1*00 | | TRBJ2-7*00 | 0.28% |
| Treg WT1 | TGTGCCAGCAGTGGGACAGACTATGCTGAGCAGTTCTTC | 1 | TRBV13-3*00 | TRBD1*00 | TRBJ2-1*00 | 0.28% |

| | | | | | | |
|---|---|---|---|---|---|---|
| Treg WT1 | TGTGCCAGCAGTGGGGGACAGGGGAGTGCAGAAACGCTGTATTTT | 1 | TRBV13-1*00 | TRBD1*00 | TRBJ2-3*00 | 0.28% |
| Treg WT1 | TGTGCCAGCAGTGTCACAGGGGAAGAACAGTACTTC | 1 | TRBV13-3*00 | TRBD1*00 | TRBJ2-7*00 | 0.28% |
| Treg WT1 | TGTGCCAGCAGTTCCGGGACAACAGAAACGCTGTATTTT | 1 | TRBV15*00 | TRBD1*00 | TRBJ2-3*00 | 0.28% |
| Treg WT1 | TGTGCCAGCAGTTCCGGTTATAACAACCAGGCTCCGCTTTTT | 1 | TRBV15*00 | | TRBJ1-5*00 | 0.28% |
| Treg WT1 | TGTGCCAGCAGTTGGACAGGGGACACCGGGCAGCTCTACTTT | 1 | TRBV13-3*00 | TRBD1*00 | TRBJ2-2*00 | 0.28% |
| Treg WT1 | TGTGCCAGCAGTTTAAGAGGATTTGAACAGTACTTC | 1 | TRBV15*00 | | TRBJ2-7*00 | 0.28% |
| Treg WT1 | TGTGCCAGCAGTTTAAGAGGGGCAGACACCCAGTACTTT | 1 | TRBV15*00 | TRBD1*00 | TRBJ2-5*00 | 0.28% |
| Treg WT1 | TGTGCCAGCAGTTTAGACCAAGACACCCAGTACTTT | 1 | TRBV15*00 | | TRBJ2-5*00 | 0.28% |
| Treg WT1 | TGTGCCAGCAGTTTAGACCGCGACACCCAGTACTTT | 1 | TRBV15*00 | | TRBJ2-5*00 | 0.28% |
| Treg WT1 | TGTGCCAGCAGTTTAGCGGAAGAGCAGTTCTTC | 1 | TRBV15*00 | | TRBJ2-1*00 | 0.28% |
| Treg WT1 | TGTGCCAGCAGTTTAGGCAGACCCTATGAACAGTACTTC | 1 | TRBV15*00 | | TRBJ2-7*00 | 0.28% |
| Treg WT1 | TGTGCCAGCAGTTTAGGGGAGAGGGAACAGTACTTC | 1 | TRBV15*00 | | TRBJ2-7*00 | 0.28% |
| Treg WT1 | TGTGCCAGCAGTTTTGGGGGGGCGGACACCTTGTACTTT | 1 | TRBV15*00 | TRBD2*00 | TRBJ2-4*00 | 0.28% |
| Treg WT1 | TGTGCCAGCGCAGAAAACTCCGACTACACCTTC | 1 | TRBV26*00 | | TRBJ1-2*00 | 0.28% |
| Treg WT1 | TGTGCCAGCGCCCGGGACATTGCAAACTCCGACTACACCTTC | 1 | TRBV13-2*00 | TRBD1*00 | TRBJ1-2*00 | 0.28% |
| Treg WT1 | TGTGCCAGCGCCGGGACAGGGGGCGCTGAGCAGTTCTTC | 1 | TRBV13-2*00 | TRBD1*00 | TRBJ2-1*00 | 0.28% |
| Treg WT1 | TGTGCCAGCGGAAGAGGTCAAAACACCTTGTACTTT | 1 | TRBV13-2*00 | | TRBJ2-4*00 | 0.28% |
| Treg WT1 | TGTGCCAGCGGAGGGTCCCTCAACGAAAGATTATTTTTC | 1 | TRBV13-2*00 | TRBD1*00 | TRBJ1-4*00 | 0.28% |
| Treg WT1 | TGTGCCAGCGGCAGGGACACCCAGTACTTT | 1 | TRBV13-2*00 | | TRBJ2-5*00 | 0.28% |
| Treg WT1 | TGTGCCAGCGGCGAACTCCGACTACACCTTC | 1 | TRBV13-2*00 | | TRBJ1-2*00 | 0.28% |
| Treg WT1 | TGTGCCAGCGGGACAGGGAACTATGCTGAGCAGTTCTTC | 1 | TRBV13-2*00 | TRBD1*00 | TRBJ2-1*00 | 0.28% |
| Treg WT1 | TGTGCCAGCGGGGAACTGGGGGGGTATGAACAGTACTTC | 1 | TRBV13-2*00 | TRBD2*00 | TRBJ2-7*00 | 0.28% |
| Treg WT1 | TGTGCCAGCGGGGGAGGGGGATTTGCTGAGCAGTTCTTC | 1 | TRBV13-2*00 | TRBD1*00 | TRBJ2-1*00 | 0.28% |
| Treg WT1 | TGTGCCAGCGGTATGGGACAAAACTCCGACTACACCTTC | 1 | TRBV13-2*00 | TRBD1*00 | TRBJ1-2*00 | 0.28% |
| Treg WT1 | TGTGCCAGCGGTCAGAGGGGGGCAAACTCCGACTACACCTTC | 1 | TRBV13-2*00 | TRBD1*00 | TRBJ1-2*00 | 0.28% |
| Treg WT1 | TGTGCCAGCGGTCTGGGGGGGAGTGCAGAAACGCTGTATTTT | 1 | TRBV13-2*00 | TRBD2*00 | TRBJ2-3*00 | 0.28% |
| Treg WT1 | TGTGCCAGCGGTGACATCAACCAGGCTCCGCTTTTT | 1 | TRBV13-2*00 | | TRBJ1-5*00 | 0.28% |
| Treg WT1 | TGTGCCAGCGGTGACTGGGGGGAAGACACCCAGTACTTT | 1 | TRBV13-2*00 | TRBD2*00 | TRBJ2-5*00 | 0.28% |
| Treg WT1 | TGTGCCAGCGGTGACTGGGGGGGCCTAAACACCTTGTACTTT | 1 | TRBV13-2*00 | TRBD2*00 | TRBJ2-4*00 | 0.28% |
| Treg WT1 | TGTGCCAGCGGTGAGAGGGGGGGCAAACTCCGACTACACCTTC | 1 | TRBV13-2*00 | TRBD1*00 | TRBJ1-2*00 | 0.28% |
| Treg WT1 | TGTGCCAGCGGTGATACGCTCTATTTT | 1 | TRBV13-2*00 | | TRBJ1-3*00 | 0.28% |
| Treg WT1 | TGTGCCAGCGGTGATAGGAACTATGCTAAGCAGTTCTTC | 1 | TRBV13-2*00 | | TRBJ2-1*00 | 0.28% |
| Treg WT1 | TGTGCCAGCGGTGATAGGAGTCAAAACACCTTGTACTTT | 1 | TRBV13-2*00 | | TRBJ2-4*00 | 0.28% |
| Treg WT1 | TGTGCCAGCGGTGATAGGTGGGCTCCGCTTTTT | 1 | TRBV13-2*00 | | TRBJ1-5*00 | 0.28% |
| Treg WT1 | TGTGCCAGCGGTGATCCAGCAAACTCCGACTACACCTTC | 1 | TRBV13-2*00 | | TRBJ1-2*00 | 0.28% |
| Treg WT1 | TGTGCCAGCGGTGATGAGCGGGGGGCGCAAGTCTTCTTT | 1 | TRBV13-2*00 | TRBD2*00 | TRBJ1-1*00 | 0.28% |
| Treg WT1 | TGTGCCAGCGGTGATGAGGCTGAGCAGTTCTTC | 1 | TRBV13-2*00 | | TRBJ2-1*00 | 0.28% |
| Treg WT1 | TGTGCCAGCGGTGATGAGGGACTGGTCTATGCTGAGCAGTTCTTC | 1 | TRBV13-2*00 | TRBD2*00 | TRBJ2-1*00 | 0.28% |
| Treg WT1 | TGTGCCAGCGGTGATGCAAATTCTGGAAATACGCTCTATTTT | 1 | TRBV13-2*00 | | TRBJ1-3*00 | 0.28% |
| Treg WT1 | TGTGCCAGCGGTGATGCAAGGGAGGCCGACTACACCTTC | 1 | TRBV13-2*00 | | TRBJ1-2*00 | 0.28% |
| Treg WT1 | TGTGCCAGCGGTGATGCAGGGGTCTCCGACTACACCTTC | 1 | TRBV13-2*00 | | TRBJ1-2*00 | 0.28% |
| Treg WT1 | TGTGCCAGCGGTGATGCCCTGGGAGGCACCTTGTACTTT | 1 | TRBV13-2*00 | TRBD2*00 | TRBJ2-4*00 | 0.28% |
| Treg WT1 | TGTGCCAGCGGTGATGCCGGGGGAACGCTGTATTTT | 1 | TRBV13-2*00 | TRBD1*00 | TRBJ2-3*00 | 0.28% |
| Treg WT1 | TGTGCCAGCGGTGATGCGGGCTCCGACTACACCTTC | 1 | TRBV13-2*00 | | TRBJ1-2*00 | 0.28% |
| Treg WT1 | TGTGCCAGCGGTGATGCTAACCAAGACACCCAGTACTTT | 1 | TRBV13-2*00 | | TRBJ2-5*00 | 0.28% |
| Treg WT1 | TGTGCCAGCGGTGATGCTGGGGGTTATGAACAGTACTTC | 1 | TRBV13-2*00 | TRBD2*00 | TRBJ2-7*00 | 0.28% |
| Treg WT1 | TGTGCCAGCGGTGATGGGGGGGCAAACACCGGGCAGCTCTACTTT | 1 | TRBV13-2*00 | TRBD1*00 | TRBJ2-2*00 | 0.28% |
| Treg WT1 | TGTGCCAGCGGTGATTCATATAATTCGCCCCTCTACTTT | 1 | TRBV13-2*00 | | TRBJ1-6*00 | 0.28% |
| Treg WT1 | TGTGCCAGCGGTGGGACAAATAACTATGCTGAGCAGTTCTTC | 1 | TRBV13-2*00 | TRBD1*00 | TRBJ2-1*00 | 0.28% |
| Treg WT1 | TGTGCCAGCGGTGGGGGACACCATAACTATGCTGAGCAGTTCTTC | 1 | TRBV13-2*00 | TRBD1*00 | TRBJ2-1*00 | 0.28% |
| Treg WT1 | TGTGCCAGCGGTGGTGACAGGGGGAACTCCGACTACACCTTC | 1 | TRBV13-2*00 | TRBD1*00 | TRBJ1-2*00 | 0.28% |
| Treg WT1 | TGTGCCAGCGGTGTTCTGGGGGGTAACTATGCTGAGCAGTTCTTC | 1 | TRBV13-2*00 | TRBD2*00 | TRBJ2-1*00 | 0.28% |
| Treg WT1 | TGTGCCAGCGTGGGGGGCTATGAACAGTACTTC | 1 | TRBV13-1*00 | TRBD2*00 | TRBJ2-7*00 | 0.28% |
| Treg WT1 | TGTGCCAGCTCCCTCAGGGGAAGGCACGAAAGATTATTTTTC | 1 | TRBV12-1*00 | TRBD1*00 | TRBJ1-4*00 | 0.28% |
| Treg WT1 | TGTGCCAGCTCCGGGACAGCAAACTCCGACTACACCTTC | 1 | TRBV18*00 | TRBD1*00 | TRBJ1-2*00 | 0.28% |
| Treg WT1 | TGTGCCAGCTCCTGGCCTGGGGGCTATGAACAGTACTTC | 1 | TRBV12-1*00 | TRBD2*00 | TRBJ2-7*00 | 0.28% |
| Treg WT1 | TGTGCCAGCTCGCGACTGGGGAGTCAAAACACCTTGTACTTT | 1 | TRBV12-1*00 | TRBD2*00 | TRBJ2-4*00 | 0.28% |
| Treg WT1 | TGTGCCAGCTCGGGGGCCTATGCTGAGCAGTTCTTC | 1 | TRBV12-1*00 | TRBD1*00 | TRBJ2-1*00 | 0.28% |
| Treg WT1 | TGTGCCAGCTCTCACCAGGGGAGGTACACCTTGTACTTT | 1 | TRBV12-1*00 | TRBD1*00 | TRBJ2-4*00 | 0.28% |
| Treg WT1 | TGTGCCAGCTCTCCCAATTCTGGAAATACGCTCTATTTT | 1 | TRBV12-2*00 | | TRBJ1-3*00 | 0.28% |
| Treg WT1 | TGTGCCAGCTCTCCCGGGGACTGGGGGGGCAGTGCAGAAACGCTGTATTTT | 1 | TRBV12-1*00 | TRBD2*00 | TRBJ2-3*00 | 0.28% |
| Treg WT1 | TGTGCCAGCTCTCGCTCTAGTGCAGAAACGCTGTATTTT | 1 | TRBV12-1*00 | | TRBJ2-3*00 | 0.28% |
| Treg WT1 | TGTGCCAGCTCTCTCAGGGACACCCAGTACTTT | 1 | TRBV12-1*00 | | TRBJ2-5*00 | 0.28% |
| Treg WT1 | TGTGCCAGCTCTCTCAGGGTATCAGAAGTCTTCTTT | 1 | TRBV12-1*00 | | TRBJ1-1*00 | 0.28% |
| Treg WT1 | TGTGCCAGCTCTCTCATGGGGGATGAACAGTACTTC | 1 | TRBV12-1*00 | TRBD2*00 | TRBJ2-7*00 | 0.28% |
| Treg WT1 | TGTGCCAGCTCTCTCCAGGGCCAAAACACCCAGTACTTT | 1 | TRBV12-1*00 | TRBD1*00 | TRBJ2-5*00 | 0.28% |
| Treg WT1 | TGTGCCAGCTCTCTCCGGGGGGTTAGTGCAGAAACGCTGTATTTT | 1 | TRBV12-1*00 | TRBD2*00 | TRBJ2-3*00 | 0.28% |
| Treg WT1 | TGTGCCAGCTCTCTCGAATACTCCGACTACACCTTC | 1 | TRBV12-1*00 | | TRBJ1-2*00 | 0.28% |
| Treg WT1 | TGTGCCAGCTCTCTCGACAGAAAAAAACACCTTGTACTTT | 1 | TRBV12-1*00 | | TRBJ2-4*00 | 0.28% |
| Treg WT1 | TGTGCCAGCTCTCTCGACTGGGGGGGCGGGGAACAGTACTTC | 1 | TRBV12-1*00 | TRBD2*00 | TRBJ2-7*00 | 0.28% |
| Treg WT1 | TGTGCCAGCTCTCTCGACTGGGGGGTTGAACAGTACTTC | 1 | TRBV12-1*00 | TRBD2*00 | TRBJ2-7*00 | 0.28% |
| Treg WT1 | TGTGCCAGCTCTCTCGAGTCATACTCCGACTACACCTTC | 1 | TRBV12-1*00 | | TRBJ1-2*00 | 0.28% |
| Treg WT1 | TGTGCCAGCTCTCTCGGCTGGGGGTATGAACAGTACTTC | 1 | TRBV12-1*00 | TRBD2*00 | TRBJ2-7*00 | 0.28% |
| Treg WT1 | TGTGCCAGCTCTCTCGGGACACCCTCCGACTACACCTTC | 1 | TRBV12-1*00 | TRBD1*00 | TRBJ1-2*00 | 0.28% |
| Treg WT1 | TGTGCCAGCTCTCTCGGGACTGGGGACTATGAACAGTACTTC | 1 | TRBV12-1*00 | TRBD2*00 | TRBJ2-7*00 | 0.28% |

| | | | | | | |
|---|---|---|---|---|---|---|
| Treg WT1 | TGTGCCAGCTCTCTCGGGGGGAACAACGAAAGATTATTTTTC | 1 | TRBV12-1*00 | TRBD1*00 | TRBJ1-4*00 | 0.28% |
| Treg WT1 | TGTGCCAGCTCTTACAGGGGAGACACCTTGTACTTT | 1 | TRBV12-1*00 | TRBD1*00 | TRBJ2-4*00 | 0.28% |
| Treg WT1 | TGTGCCAGCTGTGATTTTGAACAGTACTTC | 1 | TRBV13-2*00 | | TRBJ2-7*00 | 0.28% |
| Treg WT1 | TGTGCCATCAGAAACTCCGACTACACCTTC | 1 | TRBV4*00 | | TRBJ1-2*00 | 0.28% |
| Treg WT1 | TGTGCCTGGAATCTGGGCCAAGACACCCAGTACTTT | 1 | TRBV31*00 | TRBD2*00 | TRBJ2-5*00 | 0.28% |
| Treg WT1 | TGTGCCTGGACAGAAGTCTTCTTT | 1 | TRBV31*00 | | TRBJ1-1*00 | 0.28% |
| Treg WT1 | TGTGCCTGGACTGGGGGGCCCTATGAACAGTACTTC | 1 | TRBV31*00 | TRBD2*00 | TRBJ2-7*00 | 0.28% |
| Treg WT1 | TGTGCCTGGAGCTGGGGGGGTGCAGAAACGCTGTATTTT | 1 | TRBV31*00 | TRBD2*00 | TRBJ2-3*00 | 0.28% |
| Treg WT1 | TGTGCCTGGAGTCAGGGGGCGCGAGCTGAGCAGTTCTTC | 1 | TRBV31*00 | TRBD1*00 | TRBJ2-1*00 | 0.28% |
| Treg WT1 | TGTGCCTGGAGTCTAGACAGAACTCCGACTACACCTTC | 1 | TRBV31*00 | TRBD1*00 | TRBJ1-2*00 | 0.28% |
| Treg WT1 | TGTGCCTGGAGTCTAGGGGCAAACACAGAAGTCTTCTTT | 1 | TRBV31*00 | | TRBJ1-1*00 | 0.28% |
| Treg WT1 | TGTGCCTGGAGTCTAGGGGTTGAACAGTACTTC | 1 | TRBV31*00 | TRBD1*00 | TRBJ2-7*00 | 0.28% |
| Treg WT1 | TGTGCCTGGAGTCTGGGGGGGCAAGACACCCAGTACTTT | 1 | TRBV31*00 | TRBD2*00 | TRBJ2-5*00 | 0.28% |
| Treg WT1 | TGTGCCTGGAGTTTGGGGTTTAGTGCAGAAACGCTGTATTTT | 1 | TRBV31*00 | TRBD2*00 | TRBJ2-3*00 | 0.28% |
| Treg WT1 | TGTGCCTGGGGGGGACTGGGGGTCTATGCTGAGCAGTTCTTC | 1 | TRBV31*00 | TRBD2*00 | TRBJ2-1*00 | 0.28% |
| Treg WT1 | TGTGCTAGCACTAGGACAAACTATGCTGAGCAGTTCTTC | 1 | TRBV21*00 | TRBD1*00 | TRBJ2-1*00 | 0.28% |
| Treg WT1 | TGTGCTAGCAGAAGACGTGAAGTCTTCTTT | 1 | TRBV17*00 | | TRBJ1-1*00 | 0.28% |
| Treg WT1 | TGTGCTAGCAGCCATAAACAATCCTATAATTCGCCCCTCTACTTT | 1 | TRBV29*00 | | TRBJ1-6*00 | 0.28% |
| Treg WT1 | TGTGCTAGCAGCTACAGGGGAGAAAGATTATTTTTC | 1 | TRBV21*00 | TRBD1*00 | TRBJ1-4*00 | 0.28% |
| Treg WT1 | TGTGCTAGCAGTACCAGAAAAGTCTTCTTT | 1 | TRBV17*00 | | TRBJ1-1*00 | 0.28% |
| Treg WT1 | TGTGCTAGCAGTAGAAACTATGAACAGTACTTC | 1 | TRBV17*00 | | TRBJ2-7*00 | 0.28% |
| Treg WT1 | TGTGCTAGCAGTAGAAGACAGGGGCTCAACGAAAGATTATTTTTC | 1 | TRBV17*00 | TRBD1*00 | TRBJ1-4*00 | 0.28% |
| Treg WT1 | TGTGCTAGCAGTAGAGACTGGGGCCAAGACACCCAGTACTTT | 1 | TRBV17*00 | TRBD2*00 | TRBJ2-5*00 | 0.28% |
| Treg WT1 | TGTGCTAGCAGTAGAGAGGGAAACTCCGACTACACCTTC | 1 | TRBV17*00 | | TRBJ1-2*00 | 0.28% |
| Treg WT1 | TGTGCTAGCAGTAGAGATACCGGGCAGCTCTACTTT | 1 | TRBV17*00 | | TRBJ2-2*00 | 0.28% |
| Treg WT1 | TGTGCTAGCAGTAGAGGACTGGGGGGATTTGCTGAGCAGTTCTTC | 1 | TRBV17*00 | TRBD2*00 | TRBJ2-1*00 | 0.28% |
| Treg WT1 | TGTGCTAGCAGTAGAGGGTATGAACAGTACTTC | 1 | TRBV17*00 | | TRBJ2-7*00 | 0.28% |
| Treg WT1 | TGTGCTAGCAGTAGGATTGAACAGTACTTC | 1 | TRBV17*00 | | TRBJ2-7*00 | 0.28% |
| Treg WT1 | TGTGCTAGCAGTAGTTGGGGGGGCTACCAAGACACCCAGTACTTT | 1 | TRBV17*00 | TRBD2*00 | TRBJ2-5*00 | 0.28% |
| Treg WT1 | TGTGCTAGCAGTCAATCGAGGCAGGGGAGTGCAGAAACGCTGTATTTT | 1 | TRBV21*00 | TRBD1*00 | TRBJ2-3*00 | 0.28% |
| Treg WT1 | TGTGCTAGCAGTTGGGACAGCTCCTATGAACAGTACTTC | 1 | TRBV29*00 | TRBD1*00 | TRBJ2-7*00 | 0.28% |
| Treg WT1 | TGTGCTAGCAGTTTAGAGGGGTCCCCTTCTGGAAATACGCTCTATTTT | 1 | TRBV29*00 | TRBD1*00 | TRBJ1-3*00 | 0.28% |
| Treg WT1 | TGTGCTAGCAGTTTGCGGGGGAGCTCCTATGAACAGTACTTC | 1 | TRBV29*00 | | TRBJ2-7*00 | 0.28% |
| Treg WT1 | TGTGGTGCCATCGGGAGTAACCAAGACACCCAGTACTTT | 1 | TRBV20*00 | TRBD1*00 | TRBJ2-5*00 | 0.28% |
| Treg WT1 | TGTGGTGCCCACAGGGGTGGTGCAGAAACGCTGTATTTT | 1 | TRBV20*00 | TRBD1*00 | TRBJ2-3*00 | 0.28% |
| Treg WT1 | TGTGGTGCCCCTCCCTATGAACAGTACTTC | 1 | TRBV20*00 | | TRBJ2-7*00 | 0.28% |
| Treg WT1 | TGTGGTGCCCGGGACATCCAGGCTCCGCTTTTT | 1 | TRBV20*00 | TRBD1*00 | TRBJ1-5*00 | 0.28% |
| Treg WT1 | TGTGGTGCTAAGGGGAGCGAAAGATTATTTTTC | 1 | TRBV20*00 | TRBD1*00 | TRBJ1-4*00 | 0.28% |
| Treg WT1 | TGTGGTGCTAGGAACGGCGACACCTTGTACTTT | 1 | TRBV20*00 | | TRBJ2-4*00 | 0.28% |
| Treg WT1 | TGTGGTGCTAGGGAAACAACAAACTCCGACTACACCTTC | 1 | TRBV20*00 | | TRBJ1-2*00 | 0.28% |
| Treg WT1 | TGTGGTGCTAGGGACACATATAATTCGCCCCTCTACTTT | 1 | TRBV20*00 | | TRBJ1-6*00 | 0.28% |
| Treg WT1 | TGTGGTGCTAGGTCAGGGGAGGACACCCAGTACTTT | 1 | TRBV20*00 | TRBD1*00 | TRBJ2-5*00 | 0.28% |
| Treg WT1 | TGTGGTGCTAGGTGGGGGGGCGCTGAGCAGTTCTTC | 1 | TRBV20*00 | TRBD2*00 | TRBJ2-1*00 | 0.28% |
| Treg WT1 | TGTGGTGCTAGTTATAACCAAGACACCCAGTACTTT | 1 | TRBV20*00 | | TRBJ2-5*00 | 0.28% |
| Treg WT1 | TGTGGTGCTCCCAGTGCAGAAACGCTGTATTTT | 1 | TRBV20*00 | | TRBJ2-3*00 | 0.28% |
| Treg WT1 | TGTGGTGCTCGACAGGGAAACACAGAAGTCTTCTTT | 1 | TRBV20*00 | TRBD1*00 | TRBJ1-1*00 | 0.28% |
| Treg WT1 | TGTGGTGCTGGGGACTGGGGGGAAAACACCTTGTACTTT | 1 | TRBV20*00 | TRBD2*00 | TRBJ2-4*00 | 0.28% |
| Treg WT1 | TGTGTCTGGAGTCCAGCTAGTGCAGAAACGCTGTATTTT | 1 | TRBV31*00 | | TRBJ2-3*00 | 0.28% |
| Treg WT1 | TGTGCCAGCAGTATAGACAAAAACACCTTGTACTTT | 1 | TRBV19*00 | | TRBJ2-4*00 | 0.28% |
| Treg WT1 | TGTGCTAGCAGTTTATCGTCGCCAAACACAGAAGTCTTCTTT | 1 | TRBV29*00 | | TRBJ1-1*00 | 0.28% |
| Treg WT1 | TGTGCCAGCAGTCTCACAAACTCCGACTACACCTTC | 1 | TRBV26*00 | | TRBJ1-2*00 | 0.28% |
| Treg WT1 | TGTGGTGCTAGGGACAGGTATGCTGAGCAGTTCTTC | 1 | TRBV20*00 | | TRBJ2-1*00 | 0.28% |
| Treg WT1 | TGTGCCAGCAGTCGACAGAACTCCGACTACACCTTC | 1 | TRBV19*00 | TRBD1*00 | TRBJ1-2*00 | 0.28% |

#### Treg WT2

| Sample Name | CDR3 Sequence | clone Count | best V Hit | best D Hit | best J Hit | Frequency |
|---|---|---|---|---|---|---|
| Treg WT2 | TGTGCCAGCAGTATGGGGGCAAACTCCGACTACACCTTC | 4 | TRBV19*00 | | TRBJ1-2*00 | 0.95% |
| Treg WT2 | TGCACCTGCAGTGGGGACAGGGCTGAGCAGTTCTTC | 3 | TRBV1*00 | TRBD1*00 | TRBJ2-1*00 | 0.71% |
| Treg WT2 | TGTGCCAGCAGGGACAACACAGAAGTCTTCTTT | 3 | TRBV13-1*00 | | TRBJ1-1*00 | 0.71% |
| Treg WT2 | TGTGCCAGCAGTGGAGGGGGCGACACCCAGTACTTT | 3 | TRBV13-3*00 | TRBD1*00 | TRBJ2-5*00 | 0.71% |
| Treg WT2 | TGTGCCAGCAGTTTAGGACACTATGAACAGTACTTC | 3 | TRBV15*00 | | TRBJ2-7*00 | 0.71% |
| Treg WT2 | TGTGCCAGCGGAACTGGTAACTATGCTGAGCAGTTCTTC | 3 | TRBV13-2*00 | TRBD2*00 | TRBJ2-1*00 | 0.71% |
| Treg WT2 | TGTGCCAGCGGTGGGGACAGGGGTAACCAAGACACCCAGTACTTT | 3 | TRBV13-2*00 | TRBD1*00 | TRBJ2-5*00 | 0.71% |
| Treg WT2 | TGCACCTGCAGCCCCGGGACTGGCTATGCTGAGCAGTTCTTC | 2 | TRBV1*00 | TRBD2*00 | TRBJ2-1*00 | 0.47% |
| Treg WT2 | TGCACCTGCAGTGCTCGACAGGGGGACAACGAAAGATTATTTTTC | 2 | TRBV1*00 | TRBD1*00 | TRBJ1-4*00 | 0.47% |
| Treg WT2 | TGCACCTGCAGTGGACAGTATGCTGAGCAGTTCTTC | 2 | TRBV1*00 | TRBD1*00 | TRBJ2-1*00 | 0.47% |
| Treg WT2 | TGTGCCAGCACTGACAGGGATGCTGAGCAGTTCTTC | 2 | TRBV13-1*00 | TRBD1*00 | TRBJ2-1*00 | 0.47% |
| Treg WT2 | TGTGCCAGCAGCCCACAGAGGGGGGGTGCAGAAACGCTGTATTTT | 2 | TRBV19*00 | TRBD1*00 | TRBJ2-3*00 | 0.47% |
| Treg WT2 | TGTGCCAGCAGCCCCAGACTGGAAAACACCTTGTACTTT | 2 | TRBV3*00 | TRBD2*00 | TRBJ2-4*00 | 0.47% |
| Treg WT2 | TGTGCCAGCAGCCCGGGCTATGCTGAGCAGTTCTTC | 2 | TRBV4*00 | TRBD1*00 | TRBJ2-1*00 | 0.47% |
| Treg WT2 | TGTGCCAGCAGCTCCAGGGTGGAAGTCTTCTTT | 2 | TRBV13-3*00 | TRBD1*00 | TRBJ1-1*00 | 0.47% |
| Treg WT2 | TGTGCCAGCAGGGACAACTCCTATGAACAGTACTTC | 2 | TRBV15*00 | TRBD1*00 | TRBJ2-7*00 | 0.47% |
| Treg WT2 | TGTGCCAGCAGTCACAGCCAAGACACCCAGTACTTT | 2 | TRBV26*00 | | TRBJ2-5*00 | 0.47% |
| Treg WT2 | TGTGCCAGCAGTCTGTCTACAGGCACAGAAGTCTTCTTT | 2 | TRBV26*00 | | TRBJ1-1*00 | 0.47% |
| Treg WT2 | TGTGCCAGCAGTGCGCGTTACTTC | 2 | TRBV13-1*00 | | TRBJ2-7*00 | 0.47% |

| | | | | | | |
|---|---|---|---|---|---|---|
| Treg WT2 | TGTGCCAGCAGTGGCCGGGACAAATATGAGCAGTTCTTC | 2 | TRBV13-1*00 | TRBD1*00 | TRBJ2-1*00 | 0.47% |
| Treg WT2 | TGTGCCAGCGGTGATGCAGGGGGCGCCTATGAACAGTACTTC | 2 | TRBV13-2*00 | TRBD1*00 | TRBJ2-7*00 | 0.47% |
| Treg WT2 | TGTGCCAGCTCTCTCACAATTAACCAAGACACCCAGGACTTT | 2 | TRBV12-2*00 | | TRBJ2-5*00 | 0.47% |
| Treg WT2 | TGTGCCAGCTCTCTCGTTAACACCTTGTACTTT | 2 | TRBV12-1*00 | | TRBJ2-4*00 | 0.47% |
| Treg WT2 | TGTGCCTGGAGTCCCTTCTTGGGGAACCAAGACACCCAGTACTTT | 2 | TRBV31*00 | | TRBJ2-5*00 | 0.47% |
| Treg WT2 | TGTGCCTGGAGTCGGGCTGGGGACAGCTCCTATGAACAGTACTTC | 2 | TRBV31*00 | TRBD2*00 | TRBJ2-7*00 | 0.47% |
| Treg WT2 | TGTGCCTGGAGTCTGGGGCAGGTATGCTGAGCAGTTCTTC | 2 | TRBV31*00 | TRBD1*00 | TRBJ2-1*00 | 0.47% |
| Treg WT2 | TGTGCTAGCAGTAGACTGGGGGGGCGGGCTGAGCAGTTCTTC | 2 | TRBV17*00 | TRBD2*00 | TRBJ2-1*00 | 0.47% |
| Treg WT2 | TGTGGTGCTAGCAGGGGGAGCTCCTATGAACAGTACTTC | 2 | TRBV20*00 | TRBD1*00 | TRBJ2-7*00 | 0.47% |
| Treg WT2 | TGTGCCAGCAGCCGGGACATCTCCTATGAACAGTACTTC | 2 | TRBV19*00 | TRBD1*00 | TRBJ2-7*00 | 0.47% |
| Treg WT2 | TATGCCAGCAGCCAAGAAGGACTTTCCTATGAACAGTACTTC | 1 | TRBV5*00 | TRBD2*00 | TRBJ2-7*00 | 0.24% |
| Treg WT2 | TGCACCTGCAGGATGGGGATTGAACAGTACTTC | 1 | TRBV1*00 | TRBD2*00 | TRBJ2-7*00 | 0.24% |
| Treg WT2 | TGCACCTGCAGTCCCTTCTATGAACAGTACTTC | 1 | TRBV1*00 | | TRBJ2-7*00 | 0.24% |
| Treg WT2 | TGCACCTGCAGTCTCTCGAACTGGGGGAAAGACACCCAGTACTTT | 1 | TRBV1*00 | TRBD2*00 | TRBJ2-5*00 | 0.24% |
| Treg WT2 | TGCACCTGCAGTGCAAACACCGCAAACTCCGACTACACCTTC | 1 | TRBV1*00 | | TRBJ1-2*00 | 0.24% |
| Treg WT2 | TGCACCTGCAGTGCAACAAACACCGGGCAGCTCTACTTT | 1 | TRBV1*00 | | TRBJ2-2*00 | 0.24% |
| Treg WT2 | TGCACCTGCAGTGCACGGAGGGACTCCTATGAACAGTACTTC | 1 | TRBV1*00 | | TRBJ2-7*00 | 0.24% |
| Treg WT2 | TGCACCTGCAGTGCAGACACAAACACAGAAGTCTTCTTT | 1 | TRBV1*00 | | TRBJ1-1*00 | 0.24% |
| Treg WT2 | TGCACCTGCAGTGCAGACACCTATGAACAGTACTTC | 1 | TRBV1*00 | | TRBJ2-7*00 | 0.24% |
| Treg WT2 | TGCACCTGCAGTGCAGATATGGGTGGTGCAGAAACGCTGTATTTT | 1 | TRBV1*00 | | TRBJ2-3*00 | 0.24% |
| Treg WT2 | TGCACCTGCAGTGCAGATCAGGGAAACTCCGACTACACCTTC | 1 | TRBV1*00 | | TRBJ1-2*00 | 0.24% |
| Treg WT2 | TGCACCTGCAGTGCAGGCAACGAAAGATTATTTTTC | 1 | TRBV1*00 | | TRBJ1-4*00 | 0.24% |
| Treg WT2 | TGCACCTGCAGTGCAGGGACTGATGCTGAGCAGTTCTTC | 1 | TRBV1*00 | TRBD2*00 | TRBJ2-1*00 | 0.24% |
| Treg WT2 | TGCACCTGCAGTGCAGTAGGGATCTCCGACTACACCTTC | 1 | TRBV1*00 | | TRBJ1-2*00 | 0.24% |
| Treg WT2 | TGCACCTGCAGTGCAGTGGGGGGCTTTGCAAACTCCGACTACACCTTC | 1 | TRBV1*00 | TRBD2*00 | TRBJ1-2*00 | 0.24% |
| Treg WT2 | TGCACCTGCAGTGCCACTTATGAACAGTACTTC | 1 | TRBV1*00 | | TRBJ2-7*00 | 0.24% |
| Treg WT2 | TGCACCTGCAGTGCCCAACAAGACACCCAGTACTTT | 1 | TRBV1*00 | | TRBJ2-5*00 | 0.24% |
| Treg WT2 | TGCACCTGCAGTGCGCCACCCAAGACACCCAGTACTTT | 1 | TRBV1*00 | | TRBJ2-5*00 | 0.24% |
| Treg WT2 | TGCACCTGCAGTGCTCAGATTAACCAAGACACCCAGTACTTT | 1 | TRBV1*00 | | TRBJ2-5*00 | 0.24% |
| Treg WT2 | TGCTCCAGCAGTCAATCCAGGGACTATGCTGAGCAGTTCTTC | 1 | TRBV23*00 | TRBD1*00 | TRBJ2-1*00 | 0.24% |
| Treg WT2 | TGCTCCAGCAGTCAATCGATAGGGGCAAACTCCGACTACACCTTC | 1 | TRBV23*00 | | TRBJ1-2*00 | 0.24% |
| Treg WT2 | TGCTCCAGCAGTCGCAGGGGAAACACCTTGTACTTT | 1 | TRBV23*00 | TRBD1*00 | TRBJ2-4*00 | 0.24% |
| Treg WT2 | TGTACAAGCAGCTTAGCTGGGTATGAACAGTACTTC | 1 | TRBV16*00 | TRBD2*00 | TRBJ2-7*00 | 0.24% |
| Treg WT2 | TGTAGTTCTAGAGAGAGGGGCGAAAGATTATTTTTC | 1 | TRBV30*00 | TRBD1*00 | TRBJ1-4*00 | 0.24% |
| Treg WT2 | TGTAGTTCTAGTAGGGGGGCCATCTTT | 1 | TRBV30*00 | TRBD2*00 | TRBJ1-1*00 | 0.24% |
| Treg WT2 | TGTGCAAGCAGCCACAGTGCAGAAACGCTGTATTTT | 1 | TRBV16*00 | | TRBJ2-3*00 | 0.24% |
| Treg WT2 | TGTGCAAGCAGCCCAACAGGGGGCACAGAAGTCTTCTTT | 1 | TRBV16*00 | TRBD1*00 | TRBJ1-1*00 | 0.24% |
| Treg WT2 | TGTGCAAGCAGCCCCGACAGGGGGCGGCCCAACGAAAGATTATTTTTC | 1 | TRBV16*00 | TRBD1*00 | TRBJ1-4*00 | 0.24% |
| Treg WT2 | TGTGCAAGCAGCCGGACAGAAACGCTGTATTTT | 1 | TRBV16*00 | | TRBJ2-3*00 | 0.24% |
| Treg WT2 | TGTGCAAGCAGCGCGAGCTCCTATGAACAGTACTTC | 1 | TRBV16*00 | | TRBJ2-7*00 | 0.24% |
| Treg WT2 | TGTGCAAGCAGCGGGACTGGGGGCACCCAGTACTTT | 1 | TRBV16*00 | TRBD2*00 | TRBJ2-5*00 | 0.24% |
| Treg WT2 | TGTGCAAGCAGCTCAACCGACAACGAAAGATTATTTTTC | 1 | TRBV16*00 | | TRBJ1-4*00 | 0.24% |
| Treg WT2 | TGTGCAAGCAGCTCCAGGGGTGACACAGAAGTCTTCTTT | 1 | TRBV16*00 | TRBD1*00 | TRBJ1-1*00 | 0.24% |
| Treg WT2 | TGTGCAAGCAGCTCCCGTAATACGCTCTATTTT | 1 | TRBV16*00 | | TRBJ1-3*00 | 0.24% |
| Treg WT2 | TGTGCAAGCAGCTCCCTGGGGGGGAAAAACACCTTGTACTTT | 1 | TRBV16*00 | TRBD2*00 | TRBJ2-4*00 | 0.24% |
| Treg WT2 | TGTGCAAGCAGCTCGGGACTGGGGGGGCGTGCAGAAACGCTGTATTTT | 1 | TRBV16*00 | TRBD2*00 | TRBJ2-3*00 | 0.24% |
| Treg WT2 | TGTGCAAGCAGCTGGGGACAGGGGGTTTCTGGAAATACGCTCTATTTT | 1 | TRBV16*00 | TRBD1*00 | TRBJ1-3*00 | 0.24% |
| Treg WT2 | TGTGCAAGCAGCTGGGGATCAAACACCGGGCAGCTCTACTTT | 1 | TRBV16*00 | | TRBJ2-2*00 | 0.24% |
| Treg WT2 | TGTGCAAGCAGCTTAGAAGGCCAACAGTACTTC | 1 | TRBV16*00 | | TRBJ2-7*00 | 0.24% |
| Treg WT2 | TGTGCAAGCAGCTTAGACACAGAAGTCTTCTTT | 1 | TRBV16*00 | | TRBJ1-1*00 | 0.24% |
| Treg WT2 | TGTGCAAGCAGCTTAGACAGGGGCCAAGACACCCAGTACTTT | 1 | TRBV16*00 | TRBD1*00 | TRBJ2-5*00 | 0.24% |
| Treg WT2 | TGTGCAAGCAGCTTAGATAACTATGAACAGTACTTC | 1 | TRBV16*00 | | TRBJ2-7*00 | 0.24% |
| Treg WT2 | TGTGCAAGCAGCTTAGATGGGCGGCAAGACACCCAGTACTTT | 1 | TRBV16*00 | TRBD1*00 | TRBJ2-5*00 | 0.24% |
| Treg WT2 | TGTGCAAGCAGCTTAGCAGGGGGCGCTGAGCAGTTCTTC | 1 | TRBV16*00 | TRBD1*00 | TRBJ2-1*00 | 0.24% |
| Treg WT2 | TGTGCAAGCAGCTTAGGGACGGCAAACTCCGACTACACCTTC | 1 | TRBV16*00 | | TRBJ1-2*00 | 0.24% |
| Treg WT2 | TGTGCAAGCAGCTTAGGGCAGGGGGCCGGGCAGCTCTACTTT | 1 | TRBV16*00 | TRBD1*00 | TRBJ2-2*00 | 0.24% |
| Treg WT2 | TGTGCAAGCAGCTTAGGGGGCGCCTCCGACTACACCTTC | 1 | TRBV16*00 | TRBD1*00 | TRBJ1-2*00 | 0.24% |
| Treg WT2 | TGTGCAAGCAGCTTAGGGGGCGCTGAGCAGTTCTTC | 1 | TRBV16*00 | TRBD1*00 | TRBJ2-1*00 | 0.24% |
| Treg WT2 | TGTGCAAGCAGCTTAGGGGGGACAGAAGTCTTCTTT | 1 | TRBV16*00 | TRBD1*00 | TRBJ1-1*00 | 0.24% |
| Treg WT2 | TGTGCAAGCAGCTTAGGGGTTAACCAAGACACCCAGTACTTT | 1 | TRBV16*00 | | TRBJ2-5*00 | 0.24% |
| Treg WT2 | TGTGCAAGCAGCTTAGGTAACCAAGACACCCAGTACTTT | 1 | TRBV16*00 | | TRBJ2-5*00 | 0.24% |
| Treg WT2 | TGTGCAAGCAGCTTAGTCCCAGAAGTCTTCTTT | 1 | TRBV16*00 | | TRBJ1-1*00 | 0.24% |
| Treg WT2 | TGTGCAAGCAGCTTAGTGGGAATACAGTACTTC | 1 | TRBV16*00 | | TRBJ2-7*00 | 0.24% |
| Treg WT2 | TGTGCAAGCAGCTTAGTGGGGGGGTCCTATGAACAGTACTTC | 1 | TRBV16*00 | TRBD2*00 | TRBJ2-7*00 | 0.24% |
| Treg WT2 | TGTGCAAGCAGCTTAGTTGGGTATGAACAGTACTTC | 1 | TRBV16*00 | | TRBJ2-7*00 | 0.24% |
| Treg WT2 | TGTGCAAGCAGCTTCACCAGGGGGGGGAGTCTTCTTT | 1 | TRBV16*00 | TRBD1*00 | TRBJ1-1*00 | 0.24% |
| Treg WT2 | TGTGCAAGCAGCTTCATTCAAAACACCTTGTACTTT | 1 | TRBV16*00 | | TRBJ2-4*00 | 0.24% |
| Treg WT2 | TGTGCAAGCAGCTTGACAGGGGGCGAACAGTACTTC | 1 | TRBV16*00 | TRBD1*00 | TRBJ2-7*00 | 0.24% |
| Treg WT2 | TGTGCAAGCAGCTTGCAGGGTGAACAGTACTTC | 1 | TRBV16*00 | TRBD1*00 | TRBJ2-7*00 | 0.24% |
| Treg WT2 | TGTGCAAGCAGCTTGGGAGGGGGCACCCAGTACTTT | 1 | TRBV16*00 | TRBD1*00 | TRBJ2-5*00 | 0.24% |
| Treg WT2 | TGTGCAAGCAGGGGGGCGCGGAGAAGTCTTCTTT | 1 | TRBV16*00 | TRBD1*00 | TRBJ1-1*00 | 0.24% |
| Treg WT2 | TGTGCCACCAGGGGGGACTATGCTGAGCAGTTCTTC | 1 | TRBV13-1*00 | TRBD1*00 | TRBJ2-1*00 | 0.24% |
| Treg WT2 | TGTGCCAGCAACGGACAGGGAAGGGGTCAAAACACCTTGTACTTT | 1 | TRBV14*00 | TRBD1*00 | TRBJ2-4*00 | 0.24% |
| Treg WT2 | TGTGCCAGCAGACAGGGTGCAAACACAGAAGTCTTCTTT | 1 | TRBV13-3*00 | TRBD1*00 | TRBJ1-1*00 | 0.24% |
| Treg WT2 | TGTGCCAGCAGACCCCGGGACAATCAAAACACCTTGTACTTT | 1 | TRBV26*00 | TRBD1*00 | TRBJ2-4*00 | 0.24% |

| | | | | | | |
|---|---|---|---|---|---|---|
| Treg WT2 | TGTGCCAGCAGACTGGGGAACTATGCTGAGCAGTTCTTC | 1 | TRBV14*00 | TRBD2*00 | TRBJ2-1*00 | 0.24% |
| Treg WT2 | TGTGCCAGCAGAGAGGATGACACCCAGTACTTT | 1 | TRBV15*00 | | TRBJ2-5*00 | 0.24% |
| Treg WT2 | TGTGCCAGCAGATCAGGGGGGAGGGACACCCAGTACTTT | 1 | TRBV15*00 | TRBD1*00 | TRBJ2-5*00 | 0.24% |
| Treg WT2 | TGTGCCAGCAGCAAGACTGGGGGAAGACACCCAGTACTTT | 1 | TRBV3*00 | TRBD2*00 | TRBJ2-5*00 | 0.24% |
| Treg WT2 | TGTGCCAGCAGCCAAACAGCAAACTCCGACTACACCTTC | 1 | TRBV5*00 | | TRBJ1-2*00 | 0.24% |
| Treg WT2 | TGTGCCAGCAGCCAAACGGGGGCCCAAGACACCCAGTACTTT | 1 | TRBV5*00 | TRBD1*00 | TRBJ2-5*00 | 0.24% |
| Treg WT2 | TGTGCCAGCAGCCAAACTGGGCCCTATGAACAGTACTTC | 1 | TRBV2*00 | TRBD2*00 | TRBJ2-7*00 | 0.24% |
| Treg WT2 | TGTGCCAGCAGCCAACAGGGGAACACAGAAGTCTTCTTT | 1 | TRBV2*00 | TRBD1*00 | TRBJ1-1*00 | 0.24% |
| Treg WT2 | TGTGCCAGCAGCCAACCTCGGGACAACTATGCTGAGCAGTTCTTC | 1 | TRBV5*00 | TRBD1*00 | TRBJ2-1*00 | 0.24% |
| Treg WT2 | TGTGCCAGCAGCCAAGAAACTGGCTATGCTGAGCAGTTCTTC | 1 | TRBV5*00 | TRBD2*00 | TRBJ2-1*00 | 0.24% |
| Treg WT2 | TGTGCCAGCAGCCAAGACAATAGTCAAAACACCTTGTACTTT | 1 | TRBV5*00 | | TRBJ2-4*00 | 0.24% |
| Treg WT2 | TGTGCCAGCAGCCAAGACCCCAACTCCTATGAACAGTACTTC | 1 | TRBV2*00 | TRBD2*00 | TRBJ2-7*00 | 0.24% |
| Treg WT2 | TGTGCCAGCAGCCAAGACCCCAGGGGCTCCGCTTTTT | 1 | TRBV2*00 | TRBD2*00 | TRBJ1-5*00 | 0.24% |
| Treg WT2 | TGTGCCAGCAGCCAAGAGACACCAAACTCCGACTACACCTTC | 1 | TRBV2*00 | | TRBJ1-2*00 | 0.24% |
| Treg WT2 | TGTGCCAGCAGCCAAGAGCAGTCCTCTTATGCTGAGCAGTTCTTC | 1 | TRBV2*00 | TRBD2*00 | TRBJ2-1*00 | 0.24% |
| Treg WT2 | TGTGCCAGCAGCCAAGAGCATGAACAGTACTTC | 1 | TRBV2*00 | | TRBJ2-7*00 | 0.24% |
| Treg WT2 | TGTGCCAGCAGCCAAGAGGCCAGCTATGCTGAGCAGTTCTTC | 1 | TRBV2*00 | | TRBJ2-1*00 | 0.24% |
| Treg WT2 | TGTGCCAGCAGCCAAGAGGGACTGGCCTATGAACAGTACTTC | 1 | TRBV2*00 | TRBD2*00 | TRBJ2-7*00 | 0.24% |
| Treg WT2 | TGTGCCAGCAGCCAAGAGGGGGGAACAGAAGTCTTCTTT | 1 | TRBV5*00 | TRBD2*00 | TRBJ1-1*00 | 0.24% |
| Treg WT2 | TGTGCCAGCAGCCAAGATCCTTATAACCAAGACACCCAGTACTTT | 1 | TRBV5*00 | | TRBJ2-5*00 | 0.24% |
| Treg WT2 | TGTGCCAGCAGCCAAGATGGAGGGGACTCCGACTACACCTTC | 1 | TRBV2*00 | TRBD1*00 | TRBJ1-2*00 | 0.24% |
| Treg WT2 | TGTGCCAGCAGCCAAGATTGGACAAAAAACACCTTGTACTTT | 1 | TRBV2*00 | TRBD1*00 | TRBJ2-4*00 | 0.24% |
| Treg WT2 | TGTGCCAGCAGCCAAGCCCTGGGCTATGAACAGTACTTC | 1 | TRBV2*00 | TRBD1*00 | TRBJ2-7*00 | 0.24% |
| Treg WT2 | TGTGCCAGCAGCCAAGCTGGGGGCTCAGAAACGCTGTATTTT | 1 | TRBV5*00 | TRBD2*00 | TRBJ2-3*00 | 0.24% |
| Treg WT2 | TGTGCCAGCAGCCAAGGGGACAGGGTCAGTGCAGAAACGCTGTATTTT | 1 | TRBV5*00 | TRBD1*00 | TRBJ2-3*00 | 0.24% |
| Treg WT2 | TGTGCCAGCAGCCAATGGGACAGCCAAGACACCCAGTACTTT | 1 | TRBV5*00 | TRBD1*00 | TRBJ2-5*00 | 0.24% |
| Treg WT2 | TGTGCCAGCAGCCAATGGGATGCTGAGCAGTTCTTC | 1 | TRBV2*00 | | TRBJ2-1*00 | 0.24% |
| Treg WT2 | TGTGCCAGCAGCCACTGGGGGGGCTATGAACAGTACTTC | 1 | TRBV4*00 | TRBD2*00 | TRBJ2-7*00 | 0.24% |
| Treg WT2 | TGTGCCAGCAGCCAGAGACTGGGGGGAAACACCGGGCAGCTCTACTTT | 1 | TRBV5*00 | TRBD2*00 | TRBJ2-2*00 | 0.24% |
| Treg WT2 | TGTGCCAGCAGCCAGGACATTGCAAACTCCGACTACACCTTC | 1 | TRBV2*00 | TRBD1*00 | TRBJ1-2*00 | 0.24% |
| Treg WT2 | TGTGCCAGCAGCCATGGGGACCAAGACACCCAGTACTTT | 1 | TRBV18*00 | TRBD2*00 | TRBJ2-5*00 | 0.24% |
| Treg WT2 | TGTGCCAGCAGCCCAGGGGACTGGGGGAGCTCCTATGAACAGCACTTC | 1 | TRBV5*00 | TRBD2*00 | TRBJ2-7*00 | 0.24% |
| Treg WT2 | TGTGCCAGCAGCCCATATGAACAGTACTTC | 1 | TRBV4*00 | | TRBJ2-7*00 | 0.24% |
| Treg WT2 | TGTGCCAGCAGCCCCAGGGGTGCAGAAACGCTGTATTTT | 1 | TRBV5*00 | TRBD1*00 | TRBJ2-3*00 | 0.24% |
| Treg WT2 | TGTGCCAGCAGCCCCGACAGAGGGTCGCCCCTCTACTTT | 1 | TRBV5*00 | TRBD1*00 | TRBJ1-6*00 | 0.24% |
| Treg WT2 | TGTGCCAGCAGCCCCGGGACAGATAATTCGCCCCTCTACTTT | 1 | TRBV5*00 | TRBD1*00 | TRBJ1-6*00 | 0.24% |
| Treg WT2 | TGTGCCAGCAGCCCCGGGGGAACGCTGTATTTT | 1 | TRBV3*00 | TRBD1*00 | TRBJ2-3*00 | 0.24% |
| Treg WT2 | TGTGCCAGCAGCCCCTGGGACACCTATGCTGAGCAGTTCTTC | 1 | TRBV2*00 | TRBD1*00 | TRBJ2-1*00 | 0.24% |
| Treg WT2 | TGTGCCAGCAGCCCGACAGATTATGAACAGTACTTC | 1 | TRBV2*00 | TRBD1*00 | TRBJ2-7*00 | 0.24% |
| Treg WT2 | TGTGCCAGCAGCCCGACAGGTGGCGAACAGTACTTC | 1 | TRBV5*00 | TRBD1*00 | TRBJ2-7*00 | 0.24% |
| Treg WT2 | TGTGCCAGCAGCCCGGGACAATACTATGCTGAGCAGTTCTTC | 1 | TRBV4*00 | TRBD1*00 | TRBJ2-1*00 | 0.24% |
| Treg WT2 | TGTGCCAGCAGCCCGGGACAGGGCACAGAAGTCTTCTTT | 1 | TRBV4*00 | TRBD1*00 | TRBJ1-1*00 | 0.24% |
| Treg WT2 | TGTGCCAGCAGCCCGGGACAGGGGGCTGGAAATACGCTCTATTTT | 1 | TRBV2*00 | TRBD1*00 | TRBJ1-3*00 | 0.24% |
| Treg WT2 | TGTGCCAGCAGCCCGGGACTTTTAAACACCTTGTACTTT | 1 | TRBV2*00 | TRBD2*00 | TRBJ2-4*00 | 0.24% |
| Treg WT2 | TGTGCCAGCAGCCCTTGGACAGCTAGTCAAAACACCTTGTACTTT | 1 | TRBV5*00 | TRBD1*00 | TRBJ2-4*00 | 0.24% |
| Treg WT2 | TGTGCCAGCAGCCGACAGCAAAACACCTTGTACTTT | 1 | TRBV4*00 | TRBD1*00 | TRBJ2-4*00 | 0.24% |
| Treg WT2 | TGTGCCAGCAGCCGACTGGGGGGAGGTGCAGAAACGCTGTATTTT | 1 | TRBV3*00 | TRBD2*00 | TRBJ2-3*00 | 0.24% |
| Treg WT2 | TGTGCCAGCAGCCGCTGGGGGGGTCAAGACACCCAGTACTTT | 1 | TRBV4*00 | TRBD2*00 | TRBJ2-5*00 | 0.24% |
| Treg WT2 | TGTGCCAGCAGCCGGGACATGTATGCTGAGCAGTTCTTC | 1 | TRBV2*00 | TRBD1*00 | TRBJ2-1*00 | 0.24% |
| Treg WT2 | TGTGCCAGCAGCCGGGACCGAGAAGTCTTCTTT | 1 | TRBV3*00 | TRBD1*00 | TRBJ1-1*00 | 0.24% |
| Treg WT2 | TGTGCCAGCAGCCGGGACTGGGATGCTGAGCAGTTCTTC | 1 | TRBV14*00 | TRBD2*00 | TRBJ2-1*00 | 0.24% |
| Treg WT2 | TGTGCCAGCAGCCTTCGACAGAATTCGCCCCTCTACTTT | 1 | TRBV5*00 | TRBD1*00 | TRBJ1-6*00 | 0.24% |
| Treg WT2 | TGTGCCAGCAGCGGGACAGGGAACGAAAGATTATTTTTC | 1 | TRBV15*00 | TRBD1*00 | TRBJ1-4*00 | 0.24% |
| Treg WT2 | TGTGCCAGCAGCGTACAGGGGCAAGACACCCAGTACTTT | 1 | TRBV2*00 | TRBD1*00 | TRBJ2-5*00 | 0.24% |
| Treg WT2 | TGTGCCAGCAGCGTGGGGGGGCAAGACACCCAGTACTTT | 1 | TRBV4*00 | TRBD2*00 | TRBJ2-5*00 | 0.24% |
| Treg WT2 | TGTGCCAGCAGCTACTGGGGCGAAACGCTGTATTTT | 1 | TRBV3*00 | TRBD2*00 | TRBJ2-3*00 | 0.24% |
| Treg WT2 | TGTGCCAGCAGCTATCCTGGGGCCTCCTATGAACAGTACTTC | 1 | TRBV4*00 | TRBD2*00 | TRBJ2-7*00 | 0.24% |
| Treg WT2 | TGTGCCAGCAGCTCACAGGGGACTAGTGCAGAAACGCTGTATTTT | 1 | TRBV5*00 | TRBD1*00 | TRBJ2-3*00 | 0.24% |
| Treg WT2 | TGTGCCAGCAGCTCCACAGGGGAGGGAGTCTTCTTT | 1 | TRBV25*00 | TRBD1*00 | TRBJ1-1*00 | 0.24% |
| Treg WT2 | TGTGCCAGCAGCTCCAGGGGGGGGGCTGAGCAGTTCTTC | 1 | TRBV3*00 | TRBD1*00 | TRBJ2-1*00 | 0.24% |
| Treg WT2 | TGTGCCAGCAGCTCTAATCTGGGGAGCTCCTATGAACAGTACTTC | 1 | TRBV3*00 | | TRBJ2-7*00 | 0.24% |
| Treg WT2 | TGTGCCAGCAGCTCTACTGGGAGCCAAGACACCCAGTACTTT | 1 | TRBV4*00 | TRBD2*00 | TRBJ2-5*00 | 0.24% |
| Treg WT2 | TGTGCCAGCAGCTGGACTGGGGGCTATGCTGAGCAGTTCTTC | 1 | TRBV4*00 | TRBD2*00 | TRBJ2-1*00 | 0.24% |
| Treg WT2 | TGTGCCAGCAGCTGGGGACTGGGGGAAGACACCCAGTACTTT | 1 | TRBV4*00 | TRBD2*00 | TRBJ2-5*00 | 0.24% |
| Treg WT2 | TGTGCCAGCAGCTGGGTTTCTGGAAATACGCTCTATTTT | 1 | TRBV2*00 | | TRBJ1-3*00 | 0.24% |
| Treg WT2 | TGTGCCAGCAGCTTAAAAGGGAACTCCGACTACACCTTC | 1 | TRBV3*00 | | TRBJ1-2*00 | 0.24% |
| Treg WT2 | TGTGCCAGCAGCTTAAAGGGGAACTCCGACTACACCTTC | 1 | TRBV3*00 | TRBD1*00 | TRBJ1-2*00 | 0.24% |
| Treg WT2 | TGTGCCAGCAGCTTACAGGGCACCGGGCAGCTCTACTTT | 1 | TRBV3*00 | TRBD1*00 | TRBJ2-2*00 | 0.24% |
| Treg WT2 | TGTGCCAGCAGCTTACGCACAGGGGGTAACTCCGACTACACCTTC | 1 | TRBV3*00 | TRBD1*00 | TRBJ1-2*00 | 0.24% |
| Treg WT2 | TGTGCCAGCAGCTTACTGGGGGGGCAAAACACCTTGTACTTT | 1 | TRBV3*00 | TRBD2*00 | TRBJ2-4*00 | 0.24% |
| Treg WT2 | TGTGCCAGCAGCTTAGAAGACACCCAGTACTTT | 1 | TRBV3*00 | | TRBJ2-5*00 | 0.24% |
| Treg WT2 | TGTGCCAGCAGCTTAGAAGAGGGTTATGAACAGTACTTC | 1 | TRBV3*00 | | TRBJ2-7*00 | 0.24% |
| Treg WT2 | TGTGCCAGCAGCTTAGAAGGACAGGGGGCCTCCGACTACACCTTC | 1 | TRBV3*00 | TRBD1*00 | TRBJ1-2*00 | 0.24% |
| Treg WT2 | TGTGCCAGCAGCTTAGACAGCTATGAACAGTACTTC | 1 | TRBV3*00 | | TRBJ2-7*00 | 0.24% |
| Treg WT2 | TGTGCCAGCAGCTTAGACAGGGGCTATGAACAGTACTTC | 1 | TRBV3*00 | TRBD1*00 | TRBJ2-7*00 | 0.24% |

| | | | | | | |
|---|---|---|---|---|---|---|
| Treg WT2 | TGTGCCAGCAGCTTAGACCCAAACTCCGACTACACCTTC | 1 | TRBV3*00 | | TRBJ1-2*00 | 0.24% |
| Treg WT2 | TGTGCCAGCAGCTTAGAGCTGGGGGGGCTTGCTGAGCAGTTCTTC | 1 | TRBV25*00 | TRBD2*00 | TRBJ2-1*00 | 0.24% |
| Treg WT2 | TGTGCCAGCAGCTTAGCCAGGGTGAACTCCGACTACACCTTC | 1 | TRBV3*00 | TRBD1*00 | TRBJ1-2*00 | 0.24% |
| Treg WT2 | TGTGCCAGCAGCTTAGGAGTGGGCCGTCAAAACACCTTGTACTTT | 1 | TRBV3*00 | | TRBJ2-4*00 | 0.24% |
| Treg WT2 | TGTGCCAGCAGCTTAGGCCAGGCTCCGCTTTTT | 1 | TRBV3*00 | | TRBJ1-5*00 | 0.24% |
| Treg WT2 | TGTGCCAGCAGCTTAGGGGACAGAACAGAAGTCTTCTTT | 1 | TRBV3*00 | TRBD1*00 | TRBJ1-1*00 | 0.24% |
| Treg WT2 | TGTGCCAGCAGCTTAGGGGCGAACTCCGACTACACCTTC | 1 | TRBV3*00 | TRBD1*00 | TRBJ1-2*00 | 0.24% |
| Treg WT2 | TGTGCCAGCAGCTTAGGGGGAAACACCGGGCAGCTCTACTTC | 1 | TRBV3*00 | | TRBJ2-2*00 | 0.24% |
| Treg WT2 | TGTGCCAGCAGCTTAGGGGGACAGTCCTATGAACAGTACTTC | 1 | TRBV3*00 | TRBD1*00 | TRBJ2-7*00 | 0.24% |
| Treg WT2 | TGTGCCAGCAGCTTAGTGGCAGGGGGCTCCGACTACACCTTC | 1 | TRBV3*00 | TRBD1*00 | TRBJ1-2*00 | 0.24% |
| Treg WT2 | TGTGCCAGCAGCTTATCCAATAACCAAGACACCCAGTACTTT | 1 | TRBV3*00 | | TRBJ2-5*00 | 0.24% |
| Treg WT2 | TGTGCCAGCAGCTTCGACGGTGCTGAGCAGTTCTTC | 1 | TRBV3*00 | | TRBJ2-1*00 | 0.24% |
| Treg WT2 | TGTGCCAGCAGCTTCGGGACAGATAATTCGCCCCTCTACTTT | 1 | TRBV4*00 | TRBD1*00 | TRBJ1-6*00 | 0.24% |
| Treg WT2 | TGTGCCAGCAGCTTCGGGGGAGCAGAAACGCTGTATTTT | 1 | TRBV3*00 | TRBD1*00 | TRBJ2-3*00 | 0.24% |
| Treg WT2 | TGTGCCAGCAGCTTTAATTCTGGAAATACGCTCTATTTT | 1 | TRBV3*00 | | TRBJ1-3*00 | 0.24% |
| Treg WT2 | TGTGCCAGCAGCTTTCTGTATGCTGAGCAGTTCTTC | 1 | TRBV4*00 | | TRBJ2-1*00 | 0.24% |
| Treg WT2 | TGTGCCAGCAGGAAACAGGGTTCCGACTACACCTTC | 1 | TRBV22*00 | TRBD1*00 | TRBJ1-2*00 | 0.24% |
| Treg WT2 | TGTGCCAGCAGGCAGGGGCTTAACTATGCTGAGCAGTTCTTC | 1 | TRBV13-1*00 | TRBD1*00 | TRBJ2-1*00 | 0.24% |
| Treg WT2 | TGTGCCAGCAGGGAGGGGGCTTATAACTATGCTGAGCAGTTCTTC | 1 | TRBV13-1*00 | TRBD1*00 | TRBJ2-1*00 | 0.24% |
| Treg WT2 | TGTGCCAGCAGGGCGGGTGAACAGTACTTC | 1 | TRBV19*00 | | TRBJ2-7*00 | 0.24% |
| Treg WT2 | TGTGCCAGCAGGGGACAGGGAAGGCAAAACACCTTGTACTTT | 1 | TRBV15*00 | TRBD1*00 | TRBJ2-4*00 | 0.24% |
| Treg WT2 | TGTGCCAGCAGGGGACAGGTCTATGCTGAGCAGTTCTTC | 1 | TRBV3*00 | TRBD1*00 | TRBJ2-1*00 | 0.24% |
| Treg WT2 | TGTGCCAGCAGGGGGACAGTACAGTACTTC | 1 | TRBV3*00 | TRBD1*00 | TRBJ2-7*00 | 0.24% |
| Treg WT2 | TGTGCCAGCAGTAAAGAAGGGGGTGAAGTCTTCTTT | 1 | TRBV19*00 | TRBD1*00 | TRBJ1-1*00 | 0.24% |
| Treg WT2 | TGTGCCAGCAGTAGACTGGGGAGTGCAGAAACGCTGTATTTT | 1 | TRBV13-3*00 | TRBD2*00 | TRBJ2-3*00 | 0.24% |
| Treg WT2 | TGTGCCAGCAGTATAACTGGGGGGATCTATGCTGAGCAGTTCTTC | 1 | TRBV19*00 | TRBD2*00 | TRBJ2-1*00 | 0.24% |
| Treg WT2 | TGTGCCAGCAGTATAAGGCGGCAGGAACAGTACTTC | 1 | TRBV19*00 | | TRBJ2-7*00 | 0.24% |
| Treg WT2 | TGTGCCAGCAGTATACAAGTCAGTAACTATGCTGAGCAGTTCTTC | 1 | TRBV19*00 | | TRBJ2-1*00 | 0.24% |
| Treg WT2 | TGTGCCAGCAGTATACCAGGGGACGAAAGATTATTTTTC | 1 | TRBV19*00 | TRBD1*00 | TRBJ1-4*00 | 0.24% |
| Treg WT2 | TGTGCCAGCAGTATAGATTGGGACTCCGACTACACCTTC | 1 | TRBV19*00 | | TRBJ1-2*00 | 0.24% |
| Treg WT2 | TGTGCCAGCAGTATAGCGAGTGCAGAAACGCTGTATTTT | 1 | TRBV19*00 | | TRBJ2-3*00 | 0.24% |
| Treg WT2 | TGTGCCAGCAGTATCTCAGGGGGGGACCAAGACACCCAGTACTTT | 1 | TRBV19*00 | TRBD1*00 | TRBJ2-5*00 | 0.24% |
| Treg WT2 | TGTGCCAGCAGTATGAGGGCAAACTCCGACTACACCTTC | 1 | TRBV19*00 | | TRBJ1-2*00 | 0.24% |
| Treg WT2 | TGTGCCAGCAGTATGCTGGGGGGGCAAGACACCCAGTACTTT | 1 | TRBV19*00 | TRBD2*00 | TRBJ2-5*00 | 0.24% |
| Treg WT2 | TGTGCCAGCAGTATGGGGGGCTATGCTGAGCAGTTCTTC | 1 | TRBV19*00 | TRBD2*00 | TRBJ2-1*00 | 0.24% |
| Treg WT2 | TGTGCCAGCAGTCAACAGGGAGCAGAAGTCTTCTTT | 1 | TRBV26*00 | TRBD1*00 | TRBJ1-1*00 | 0.24% |
| Treg WT2 | TGTGCCAGCAGTCAATCAGGGGACTCCGACTACACCTTC | 1 | TRBV26*00 | TRBD1*00 | TRBJ1-2*00 | 0.24% |
| Treg WT2 | TGTGCCAGCAGTCCAGAGGCTGAGCAGTTCTTC | 1 | TRBV26*00 | | TRBJ2-1*00 | 0.24% |
| Treg WT2 | TGTGCCAGCAGTCCCCCGGCAGGGGGCTATGCTGAGCAGTTCTTC | 1 | TRBV26*00 | TRBD1*00 | TRBJ2-1*00 | 0.24% |
| Treg WT2 | TGTGCCAGCAGTCCCGCAGAAACGCTGTATTTT | 1 | TRBV26*00 | | TRBJ2-3*00 | 0.24% |
| Treg WT2 | TGTGCCAGCAGTCCCGGACTGGGTCAAAACACCTTGTACTTT | 1 | TRBV26*00 | TRBD2*00 | TRBJ2-4*00 | 0.24% |
| Treg WT2 | TGTGCCAGCAGTCCCGGGACAGAAGACACCCAGTACTTT | 1 | TRBV14*00 | TRBD1*00 | TRBJ2-5*00 | 0.24% |
| Treg WT2 | TGTGCCAGCAGTCCGACCGGGACCTATGCTGAGCAGTTCTTC | 1 | TRBV15*00 | TRBD1*00 | TRBJ2-1*00 | 0.24% |
| Treg WT2 | TGTGCCAGCAGTCCGGGGAACTCCGACTACACCTTC | 1 | TRBV15*00 | TRBD1*00 | TRBJ1-2*00 | 0.24% |
| Treg WT2 | TGTGCCAGCAGTCGACAGGGGAGCTCCTATGAACAGTACTTC | 1 | TRBV15*00 | TRBD1*00 | TRBJ2-7*00 | 0.24% |
| Treg WT2 | TGTGCCAGCAGTCGACATTCTGGAAATACGCTCTATTTT | 1 | TRBV26*00 | | TRBJ1-3*00 | 0.24% |
| Treg WT2 | TGTGCCAGCAGTCTAACTAGTCAAAACACCTTGTACTTT | 1 | TRBV24*00 | | TRBJ2-4*00 | 0.24% |
| Treg WT2 | TGTGCCAGCAGTCTCACAATTTCTGGAAATACGCTCTATTTT | 1 | TRBV26*00 | | TRBJ1-3*00 | 0.24% |
| Treg WT2 | TGTGCCAGCAGTCTCACAGGGGGTTCCGACTACACCTTC | 1 | TRBV26*00 | TRBD1*00 | TRBJ1-2*00 | 0.24% |
| Treg WT2 | TGTGCCAGCAGTCTCACGATCGGTGCAGAAACGCTGTATTTT | 1 | TRBV26*00 | | TRBJ2-3*00 | 0.24% |
| Treg WT2 | TGTGCCAGCAGTCTCAGGGGGGAAACGCTGTATTTT | 1 | TRBV19*00 | TRBD1*00 | TRBJ2-3*00 | 0.24% |
| Treg WT2 | TGTGCCAGCAGTCTCCAGGAAGGCTCCGACTACACCTTC | 1 | TRBV26*00 | | TRBJ1-2*00 | 0.24% |
| Treg WT2 | TGTGCCAGCAGTCTCCTGGATGAACAGTACTTC | 1 | TRBV26*00 | | TRBJ2-7*00 | 0.24% |
| Treg WT2 | TGTGCCAGCAGTCTCGACAGGGGGCGCGACTCCGACTACACCTTC | 1 | TRBV24*00 | TRBD1*00 | TRBJ1-2*00 | 0.24% |
| Treg WT2 | TGTGCCAGCAGTCTGGAGGGTTACACAGAAGTCTTCTTT | 1 | TRBV26*00 | | TRBJ1-1*00 | 0.24% |
| Treg WT2 | TGTGCCAGCAGTCTGGGACAGGGTGAACAGTACTTC | 1 | TRBV26*00 | TRBD1*00 | TRBJ2-7*00 | 0.24% |
| Treg WT2 | TGTGCCAGCAGTCTGGGCAACCAAGACACCCAGTACTTT | 1 | TRBV26*00 | | TRBJ2-5*00 | 0.24% |
| Treg WT2 | TGTGCCAGCAGTCTGTCAGGGAGCTCCTATGAACAGTACTTC | 1 | TRBV26*00 | | TRBJ2-7*00 | 0.24% |
| Treg WT2 | TGTGCCAGCAGTCTGTCGCGACAGGGGGCCGTCACCCAGTACTTT | 1 | TRBV26*00 | TRBD1*00 | TRBJ2-5*00 | 0.24% |
| Treg WT2 | TGTGCCAGCAGTCTGTCTGGGGGGGCGCGGGACACCCAGTACTTT | 1 | TRBV26*00 | TRBD2*00 | TRBJ2-5*00 | 0.24% |
| Treg WT2 | TGTGCCAGCAGTGAACTGGGGGAGGACACCTTGTACTTT | 1 | TRBV13-1*00 | TRBD2*00 | TRBJ2-4*00 | 0.24% |
| Treg WT2 | TGTGCCAGCAGTGAAGACGAACAGTACTTC | 1 | TRBV13-1*00 | | TRBJ2-7*00 | 0.24% |
| Treg WT2 | TGTGCCAGCAGTGAAGGGACAGGCTCCTATGAACAGTACTTC | 1 | TRBV13-3*00 | TRBD1*00 | TRBJ2-7*00 | 0.24% |
| Treg WT2 | TGTGCCAGCAGTGACAGGATTAATTCGCCCCTCTACTTT | 1 | TRBV13-3*00 | | TRBJ1-6*00 | 0.24% |
| Treg WT2 | TGTGCCAGCAGTGACATTGCAAACTCCGACTACACCTTC | 1 | TRBV13-3*00 | | TRBJ1-2*00 | 0.24% |
| Treg WT2 | TGTGCCAGCAGTGACCGGAGGAACTCCGACTACACCTTC | 1 | TRBV13-1*00 | | TRBJ1-2*00 | 0.24% |
| Treg WT2 | TGTGCCAGCAGTGAGGGAAACACAGAAGTCTTCTTT | 1 | TRBV13-1*00 | | TRBJ1-1*00 | 0.24% |
| Treg WT2 | TGTGCCAGCAGTGAGTTGGGGGGCCCCTCCTATGAACAGTACTTC | 1 | TRBV13-1*00 | TRBD2*00 | TRBJ2-7*00 | 0.24% |
| Treg WT2 | TGTGCCAGCAGTGATAGGGGAGGGGACACCCAGTACTTT | 1 | TRBV13-3*00 | TRBD1*00 | TRBJ2-5*00 | 0.24% |
| Treg WT2 | TGTGCCAGCAGTGATCAGGGGGGGAACACCTTGTACTTT | 1 | TRBV13-1*00 | TRBD1*00 | TRBJ2-4*00 | 0.24% |
| Treg WT2 | TGTGCCAGCAGTGATCGGACAGGGGACTATGCTGAGCAGTTCTTC | 1 | TRBV13-1*00 | TRBD1*00 | TRBJ2-1*00 | 0.24% |
| Treg WT2 | TGTGCCAGCAGTGATGACGGCAGGAGTGAACAGTACTTC | 1 | TRBV13-1*00 | | TRBJ2-7*00 | 0.24% |
| Treg WT2 | TGTGCCAGCAGTGATGAGGACTGGGGGTATGCTGAGCAGTTCTTC | 1 | TRBV13-1*00 | TRBD2*00 | TRBJ2-1*00 | 0.24% |
| Treg WT2 | TGTGCCAGCAGTGATGCAAATACAAACTCCGACTACACCTTC | 1 | TRBV13-1*00 | | TRBJ1-2*00 | 0.24% |
| Treg WT2 | TGTGCCAGCAGTGATGCAATGGGTAACTATGCTGAGCAGTTCTTC | 1 | TRBV13-1*00 | | TRBJ2-1*00 | 0.24% |

| | | | | | | |
|---|---|---|---|---|---|---|
| Treg WT2 | TGTGCCAGCAGTGATGCATCCTACACAGAAGTCTTCTTT | 1 | TRBV13-3*00 | | TRBJ1-1*00 | 0.24% |
| Treg WT2 | TGTGCCAGCAGTGATGCGGGGGACACCTTGTACTTT | 1 | TRBV13-1*00 | TRBD1*00 | TRBJ2-4*00 | 0.24% |
| Treg WT2 | TGTGCCAGCAGTGATGCTGGCACAAACACCGGGCAGCTCTACTTT | 1 | TRBV13-1*00 | | TRBJ2-2*00 | 0.24% |
| Treg WT2 | TGTGCCAGCAGTGATGCTGGGGGGGTCTATGAACAGTACTTC | 1 | TRBV13-1*00 | TRBD2*00 | TRBJ2-7*00 | 0.24% |
| Treg WT2 | TGTGCCAGCAGTGATGGCAGTCAAAACACCTTGTACTTT | 1 | TRBV13-1*00 | | TRBJ2-4*00 | 0.24% |
| Treg WT2 | TGTGCCAGCAGTGATTCCGGGGTGAAAGACACCCAGTACTTC | 1 | TRBV13-3*00 | TRBD1*00 | TRBJ2-5*00 | 0.24% |
| Treg WT2 | TGTGCCAGCAGTGATTCGGACTGGGACACCCAGTACTTT | 1 | TRBV13-3*00 | TRBD2*00 | TRBJ2-5*00 | 0.24% |
| Treg WT2 | TGTGCCAGCAGTGCCCTCACCGGGCAGCTCTACTTT | 1 | TRBV13-3*00 | | TRBJ2-2*00 | 0.24% |
| Treg WT2 | TGTGCCAGCAGTGTCGACAGGATTAACACGCAGTACTTT | 1 | TRBV13-1*00 | TRBD1*00 | TRBJ2-5*00 | 0.24% |
| Treg WT2 | TGTGCCAGCAGTGTGCTTGCTGAGCAGTTCTTC | 1 | TRBV13-3*00 | | TRBJ2-1*00 | 0.24% |
| Treg WT2 | TGTGCCAGCAGTGTGGGGTCTAGTGCAGAAACGCTGTATTTT | 1 | TRBV13-1*00 | TRBD2*00 | TRBJ2-3*00 | 0.24% |
| Treg WT2 | TGTGCCAGCAGTTACAGTAACTCCGACTACACCTTC | 1 | TRBV13-1*00 | | TRBJ1-2*00 | 0.24% |
| Treg WT2 | TGTGCCAGCAGTTCCGGGGGGTTCGGCACCGGGCAGCTCTACTTT | 1 | TRBV15*00 | TRBD2*00 | TRBJ2-2*00 | 0.24% |
| Treg WT2 | TGTGCCAGCAGTTCTGGGGTCTATGAACAGTACTTC | 1 | TRBV15*00 | TRBD2*00 | TRBJ2-7*00 | 0.24% |
| Treg WT2 | TGTGCCAGCAGTTGGGACAGCTCCTATGAACAGTACTTC | 1 | TRBV13-1*00 | TRBD1*00 | TRBJ2-7*00 | 0.24% |
| Treg WT2 | TGTGCCAGCAGTTGGGACTGGGAAGACACCCAGTACTTT | 1 | TRBV26*00 | TRBD2*00 | TRBJ2-5*00 | 0.24% |
| Treg WT2 | TGTGCCAGCAGTTGGGGGGGTGCTGAGCAGTTCTTC | 1 | TRBV15*00 | TRBD2*00 | TRBJ2-1*00 | 0.24% |
| Treg WT2 | TGTGCCAGCAGTTGTCCCGACAGGGATTGCTGAGCAGTTCTTC | 1 | TRBV22*00 | TRBD1*00 | TRBJ2-1*00 | 0.24% |
| Treg WT2 | TGTGCCAGCAGTTTACAGGGCGGGGCTGAGCAGTTCTTC | 1 | TRBV15*00 | TRBD1*00 | TRBJ2-1*00 | 0.24% |
| Treg WT2 | TGTGCCAGCAGTTTAGGGGCAGAAGTCTTCTTT | 1 | TRBV15*00 | | TRBJ1-1*00 | 0.24% |
| Treg WT2 | TGTGCCAGCAGTTTCAACACAGAAGTCTTCTTT | 1 | TRBV15*00 | | TRBJ1-1*00 | 0.24% |
| Treg WT2 | TGTGCCAGCAGTTTCGGGCCGATTTCCAACGAAAGATTATTTTTC | 1 | TRBV14*00 | | TRBJ1-4*00 | 0.24% |
| Treg WT2 | TGTGCCAGCAGTTTCTGGGGGAACTATGCTGAGCAGTTCTTC | 1 | TRBV14*00 | TRBD1*00 | TRBJ2-1*00 | 0.24% |
| Treg WT2 | TGTGCCAGCAGTTTCTTTGGATATGAACAGTACTTC | 1 | TRBV14*00 | | TRBJ2-7*00 | 0.24% |
| Treg WT2 | TGTGCCAGCAGTTTGAAGAGTGCAGAAACGCTGTATTTT | 1 | TRBV14*00 | | TRBJ2-3*00 | 0.24% |
| Treg WT2 | TGTGCCAGCAGTTTGAGACTGTACACCGGGCAGCTCTACTTT | 1 | TRBV15*00 | TRBD2*00 | TRBJ2-2*00 | 0.24% |
| Treg WT2 | TGTGCCAGCAGTTTGGAGGGCTCCGACTACACCTTC | 1 | TRBV14*00 | | TRBJ1-2*00 | 0.24% |
| Treg WT2 | TGTGCCAGCAGTTTGGGGAGCGGGCAGGCTCCGCTTTTT | 1 | TRBV15*00 | | TRBJ1-5*00 | 0.24% |
| Treg WT2 | TGTGCCAGCAGTTTTCAGAAAAACACCTTGTACTTT | 1 | TRBV19*00 | | TRBJ2-4*00 | 0.24% |
| Treg WT2 | TGTGCCAGCAGTTTTGGTGACACAGAAGTCTTCTTT | 1 | TRBV15*00 | | TRBJ1-1*00 | 0.24% |
| Treg WT2 | TGTGCCAGCCGGACAGTTAACCAAGACACCCAGTACTTT | 1 | TRBV4*00 | TRBD1*00 | TRBJ2-5*00 | 0.24% |
| Treg WT2 | TGTGCCAGCGGACAATCAGACACCCAGTACTTT | 1 | TRBV12-1*00 | TRBD1*00 | TRBJ2-5*00 | 0.24% |
| Treg WT2 | TGTGCCAGCGGCAATTCCTATAATTCGCCCCTCTACTTT | 1 | TRBV13-3*00 | | TRBJ1-6*00 | 0.24% |
| Treg WT2 | TGTGCCAGCGGCCACTCCTATGAACAGTACTTC | 1 | TRBV13-2*00 | | TRBJ2-7*00 | 0.24% |
| Treg WT2 | TGTGCCAGCGGGGGACAGGCAAACACAGAAGTCTTCTTT | 1 | TRBV13-2*00 | TRBD1*00 | TRBJ1-1*00 | 0.24% |
| Treg WT2 | TGTGCCAGCGGTCAGGGGACAGAAGTCTTCTTT | 1 | TRBV13-2*00 | TRBD1*00 | TRBJ1-1*00 | 0.24% |
| Treg WT2 | TGTGCCAGCGGTCGGACTGGGGGGGCGGGGGAACAGTACTTC | 1 | TRBV13-2*00 | TRBD2*00 | TRBJ2-7*00 | 0.24% |
| Treg WT2 | TGTGCCAGCGGTCGGGACACAGAAGTCTTCTTT | 1 | TRBV13-2*00 | | TRBJ1-1*00 | 0.24% |
| Treg WT2 | TGTGCCAGCGGTGACACAAAGAATTCGCCCCTCTACTTT | 1 | TRBV13-2*00 | | TRBJ1-6*00 | 0.24% |
| Treg WT2 | TGTGCCAGCGGTGACATCTATGAACAGTACTTC | 1 | TRBV13-2*00 | | TRBJ2-7*00 | 0.24% |
| Treg WT2 | TGTGCCAGCGGTGAGACTGGGGTTGAGGCTGAGCAGTTCTTC | 1 | TRBV13-2*00 | TRBD2*00 | TRBJ2-1*00 | 0.24% |
| Treg WT2 | TGTGCCAGCGGTGATATAGGCCAAGACACCCAGTACTTT | 1 | TRBV13-2*00 | | TRBJ2-5*00 | 0.24% |
| Treg WT2 | TGTGCCAGCGGTGATCCAGGGACCTATAATTCGCCCCTCTACTTT | 1 | TRBV13-2*00 | TRBD1*00 | TRBJ1-6*00 | 0.24% |
| Treg WT2 | TGTGCCAGCGGTGATCGGCTGCGAGAACAGTACTTC | 1 | TRBV13-2*00 | | TRBJ2-7*00 | 0.24% |
| Treg WT2 | TGTGCCAGCGGTGATGAGAACACAGAAGTCTTCTTT | 1 | TRBV13-2*00 | | TRBJ1-1*00 | 0.24% |
| Treg WT2 | TGTGCCAGCGGTGATGCAAGGGTCTATAATTCGCCCCTCTACTTT | 1 | TRBV13-2*00 | | TRBJ1-6*00 | 0.24% |
| Treg WT2 | TGTGCCAGCGGTGATGCAGGTGCTGAGCAGTTCTTC | 1 | TRBV13-2*00 | | TRBJ2-1*00 | 0.24% |
| Treg WT2 | TGTGCCAGCGGTGATGCCGGGACAGGGAACACCCAGTACTTT | 1 | TRBV13-2*00 | TRBD1*00 | TRBJ2-5*00 | 0.24% |
| Treg WT2 | TGTGCCAGCGGTGATGCGGGGAGTTCCTATAATTCGCCCCTCTACTTT | 1 | TRBV13-2*00 | | TRBJ1-6*00 | 0.24% |
| Treg WT2 | TGTGCCAGCGGTGATGCGGGGGAACAGTACTTC | 1 | TRBV13-2*00 | | TRBJ2-7*00 | 0.24% |
| Treg WT2 | TGTGCCAGCGGTGATGCTCGCACCGGGCAGCTCTACTTT | 1 | TRBV13-2*00 | | TRBJ2-2*00 | 0.24% |
| Treg WT2 | TGTGCCAGCGGTGATGTAGAAATCTATGAACAGTACTTC | 1 | TRBV13-2*00 | | TRBJ2-7*00 | 0.24% |
| Treg WT2 | TGTGCCAGCGGTGATGTCCTGGGGAGCTATGAACAGTACTTC | 1 | TRBV13-2*00 | TRBD2*00 | TRBJ2-7*00 | 0.24% |
| Treg WT2 | TGTGCCAGCGGTGCTATGGGGGCTCGAGACACCCAGTACTTT | 1 | TRBV13-2*00 | TRBD1*00 | TRBJ2-5*00 | 0.24% |
| Treg WT2 | TGTGCCAGCGGTGTTGACAGGGCCACCCAGTACTTT | 1 | TRBV13-2*00 | TRBD1*00 | TRBJ2-5*00 | 0.24% |
| Treg WT2 | TGTGCCAGCGGTTCAGGGGGCGCAGGCCAAGACACCCAGTACTTT | 1 | TRBV13-2*00 | TRBD1*00 | TRBJ2-5*00 | 0.24% |
| Treg WT2 | TGTGCCAGCGGTTTGGGGGGGGTCCAAGACACCCAGTACTTT | 1 | TRBV13-2*00 | TRBD2*00 | TRBJ2-5*00 | 0.24% |
| Treg WT2 | TGTGCCAGCTCCCCGGGACAGGGGAGCGAAAGATTATTTTTC | 1 | TRBV12-1*00 | TRBD1*00 | TRBJ1-4*00 | 0.24% |
| Treg WT2 | TGTGCCAGCTCCCTATATGAACAGTACTTC | 1 | TRBV12-2*00 | | TRBJ2-7*00 | 0.24% |
| Treg WT2 | TGTGCCAGCTCCCTGGGGGTCTCCTATGAACAGTACTTC | 1 | TRBV12-1*00 | TRBD2*00 | TRBJ2-7*00 | 0.24% |
| Treg WT2 | TGTGCCAGCTCGCAGGGGGCACAGAAGTCTTCTTT | 1 | TRBV12-1*00 | TRBD1*00 | TRBJ1-1*00 | 0.24% |
| Treg WT2 | TGTGCCAGCTCGCGACTGATTGAACAGTACTTC | 1 | TRBV12-1*00 | TRBD2*00 | TRBJ2-7*00 | 0.24% |
| Treg WT2 | TGTGCCAGCTCTACAGGGGCAAACTCCGACTACACCTTC | 1 | TRBV12-2*00 | TRBD1*00 | TRBJ1-2*00 | 0.24% |
| Treg WT2 | TGTGCCAGCTCTCCCAACTGGGCTAGTGCAGAAACGCTGTATTTT | 1 | TRBV12-1*00 | TRBD2*00 | TRBJ2-3*00 | 0.24% |
| Treg WT2 | TGTGCCAGCTCTCCCGGGACGGACTATGCTGAGCAGTTCTTC | 1 | TRBV12-1*00 | TRBD1*00 | TRBJ2-1*00 | 0.24% |
| Treg WT2 | TGTGCCAGCTCTCGGGACAGGGAGGACTCCGACTACACCTTC | 1 | TRBV18*00 | TRBD1*00 | TRBJ1-2*00 | 0.24% |
| Treg WT2 | TGTGCCAGCTCTCGTGGGGTCGGGCAGCTCTACTTT | 1 | TRBV12-1*00 | TRBD2*00 | TRBJ2-2*00 | 0.24% |
| Treg WT2 | TGTGCCAGCTCTCTCCAACTGGGGGGGGGGGCTGAGCAGTTCTTC | 1 | TRBV12-1*00 | TRBD2*00 | TRBJ2-1*00 | 0.24% |
| Treg WT2 | TGTGCCAGCTCTCTCCCCAGCTCCTATGAACAGTACTTC | 1 | TRBV12-1*00 | | TRBJ2-7*00 | 0.24% |
| Treg WT2 | TGTGCCAGCTCTCTCCGCAGGGGAAACAACCAGGCTCCGCTTTTT | 1 | TRBV12-1*00 | TRBD1*00 | TRBJ1-5*00 | 0.24% |
| Treg WT2 | TGTGCCAGCTCTCTCCGGGACAATAGTGCAGAAACGCTGTATTTT | 1 | TRBV12-2*00 | TRBD1*00 | TRBJ2-3*00 | 0.24% |
| Treg WT2 | TGTGCCAGCTCTCTCCGGGGAGAAAACACCTTGTACTTT | 1 | TRBV12-1*00 | | TRBJ2-4*00 | 0.24% |
| Treg WT2 | TGTGCCAGCTCTCTCGAACCTCATTCTGGAAATACGCTCTATTTT | 1 | TRBV12-2*00 | | TRBJ1-3*00 | 0.24% |
| Treg WT2 | TGTGCCAGCTCTCTCGACAGGCAGCAGTTCTTC | 1 | TRBV12-1*00 | | TRBJ2-1*00 | 0.24% |
| Treg WT2 | TGTGCCAGCTCTCTCGACAGGGACTCCGACTACACCTTC | 1 | TRBV12-2*00 | TRBD1*00 | TRBJ1-2*00 | 0.24% |

| | | | | | | |
|---|---|---|---|---|---|---|
| Treg WT2 | TGTGCCAGCTCTCTCGACATAAACACAGAAGTCTTCTTT | 1 | TRBV12-1*00 | | TRBJ1-1*00 | 0.24% |
| Treg WT2 | TGTGCCAGCTCTCTCGAGGGGGGCTCCTATGAACAGTACTTC | 1 | TRBV12-1*00 | | TRBJ2-7*00 | 0.24% |
| Treg WT2 | TGTGCCAGCTCTCTCGCCAGCTCCTATGAACAGTACTTC | 1 | TRBV12-1*00 | | TRBJ2-7*00 | 0.24% |
| Treg WT2 | TGTGCCAGCTCTCTCGCGGGGTCAAACACAGAAGTCTTCTTT | 1 | TRBV12-1*00 | | TRBJ1-1*00 | 0.24% |
| Treg WT2 | TGTGCCAGCTCTCTCGGACTGGGGGGGCTGGAACAGTACTTC | 1 | TRBV12-1*00 | TRBD2*00 | TRBJ2-7*00 | 0.24% |
| Treg WT2 | TGTGCCAGCTCTCTGCTGGGCCTCTATGCTGAGCAGTTCTTC | 1 | TRBV12-1*00 | TRBD2*00 | TRBJ2-1*00 | 0.24% |
| Treg WT2 | TGTGCCAGCTCTGCCACGACAGTTTTCGAAACGCTGTATTTT | 1 | TRBV12-1*00 | TRBD1*00 | TRBJ2-3*00 | 0.24% |
| Treg WT2 | TGTGCCTGGACAACGAACTATGCTGAGCAGTTCTTC | 1 | TRBV31*00 | | TRBJ2-1*00 | 0.24% |
| Treg WT2 | TGTGCCTGGACCGACAGGGACACCCAGTACTTT | 1 | TRBV31*00 | TRBD1*00 | TRBJ2-5*00 | 0.24% |
| Treg WT2 | TGTGCCTGGAGAGCCACCGGGCAGCTCTACTTT | 1 | TRBV31*00 | | TRBJ2-2*00 | 0.24% |
| Treg WT2 | TGTGCCTGGAGATCAGGGGCGGGCACCCAGTACTTT | 1 | TRBV31*00 | TRBD1*00 | TRBJ2-5*00 | 0.24% |
| Treg WT2 | TGTGCCTGGAGATCGGACAGAGCTCCTATGAACAGTACTTC | 1 | TRBV31*00 | TRBD1*00 | TRBJ2-7*00 | 0.24% |
| Treg WT2 | TGTGCCTGGAGCCTCCCAAACTCCGACTACACCTTC | 1 | TRBV31*00 | | TRBJ1-2*00 | 0.24% |
| Treg WT2 | TGTGCCTGGAGTAGGTGGGGAAACTATGCTGAGCAGTTCTTC | 1 | TRBV31*00 | TRBD2*00 | TRBJ2-1*00 | 0.24% |
| Treg WT2 | TGTGCCTGGAGTCCCAGAAACTATGCTGAGCAGTTCTTC | 1 | TRBV31*00 | | TRBJ2-1*00 | 0.24% |
| Treg WT2 | TGTGCCTGGAGTCCCGGGAGAGAGCAGTTCTTC | 1 | TRBV31*00 | TRBD1*00 | TRBJ2-1*00 | 0.24% |
| Treg WT2 | TGTGCCTGGAGTCCCTGACAGGGATCAACTCCGACTACACCTTC | 1 | TRBV31*00 | TRBD1*00 | TRBJ1-2*00 | 0.24% |
| Treg WT2 | TGTGCCTGGAGTCCTCCGCGACAGTCCTATAATTCGCCCCTCTACTTT | 1 | TRBV31*00 | TRBD1*00 | TRBJ1-6*00 | 0.24% |
| Treg WT2 | TGTGCCTGGAGTCGAGTTAACTATGCTGAGCAGTTCTTC | 1 | TRBV31*00 | | TRBJ2-1*00 | 0.24% |
| Treg WT2 | TGTGCCTGGAGTCGGGACAGCTATGCTGAGCAGTTCTTC | 1 | TRBV31*00 | TRBD1*00 | TRBJ2-1*00 | 0.24% |
| Treg WT2 | TGTGCCTGGAGTCTAACAGCCAACGAAAGATTATTTTTC | 1 | TRBV31*00 | | TRBJ1-4*00 | 0.24% |
| Treg WT2 | TGTGCCTGGAGTCTAGGACAGGGGAGGATCAACCAGGCTCCGCTTTTT | 1 | TRBV31*00 | TRBD1*00 | TRBJ1-5*00 | 0.24% |
| Treg WT2 | TGTGCCTGGAGTCTAGGGACAAACACAGAAGTCTTCTTT | 1 | TRBV31*00 | | TRBJ1-1*00 | 0.24% |
| Treg WT2 | TGTGCCTGGAGTCTCTGGGGGGGCCGTGAACAGTACTTC | 1 | TRBV31*00 | TRBD2*00 | TRBJ2-7*00 | 0.24% |
| Treg WT2 | TGTGCCTGGAGTCTGGGGGGGAGTGAACAGTACTTC | 1 | TRBV31*00 | TRBD2*00 | TRBJ2-7*00 | 0.24% |
| Treg WT2 | TGTGCCTGGAGTCTTAAGAGAACAGTACTTC | 1 | TRBV31*00 | | TRBJ2-7*00 | 0.24% |
| Treg WT2 | TGTGCCTGGAGTCTTCCAGGGGGGTATGCTGAGCAGTTCTTC | 1 | TRBV31*00 | TRBD1*00 | TRBJ2-1*00 | 0.24% |
| Treg WT2 | TGTGCCTGGAGTTCAGGGGGCGCCGACACCCAGTACTTT | 1 | TRBV31*00 | TRBD1*00 | TRBJ2-5*00 | 0.24% |
| Treg WT2 | TGTGCCTGGAGTTCTGGGGGGAGTCAAAACACCTTGTACTTT | 1 | TRBV31*00 | TRBD2*00 | TRBJ2-4*00 | 0.24% |
| Treg WT2 | TGTGCCTTGGGCAAACAAGACACCCAGTACTTT | 1 | TRBV13-1*00 | | TRBJ2-5*00 | 0.24% |
| Treg WT2 | TGTGCTAGCAATAGATTTGCAAACACAGAAGTCTTCTTT | 1 | TRBV21*00 | | TRBJ1-1*00 | 0.24% |
| Treg WT2 | TGTGCTAGCAGCAGACTGGGGAACTATGCTGAGCAGTTCTTC | 1 | TRBV17*00 | TRBD2*00 | TRBJ2-1*00 | 0.24% |
| Treg WT2 | TGTGCTAGCAGCCCAGGGAACACCGGGCAGCTCTACTTT | 1 | TRBV29*00 | TRBD1*00 | TRBJ2-2*00 | 0.24% |
| Treg WT2 | TGTGCTAGCAGTAAACATAACAACCAGGCTCCGCTTTTT | 1 | TRBV17*00 | | TRBJ1-5*00 | 0.24% |
| Treg WT2 | TGTGCTAGCAGTACAGGGGATGAACAGTACTTC | 1 | TRBV17*00 | TRBD1*00 | TRBJ2-7*00 | 0.24% |
| Treg WT2 | TGTGCTAGCAGTAGACTGGGAGGGCGGGCTGAGCAGTTCTTC | 1 | TRBV17*00 | TRBD2*00 | TRBJ2-1*00 | 0.24% |
| Treg WT2 | TGTGCTAGCAGTAGAGACTGGCAAGACACCCAGTACTTT | 1 | TRBV17*00 | | TRBJ2-5*00 | 0.24% |
| Treg WT2 | TGTGCTAGCAGTAGAGATGGGAATTCTGGAAATACGCTCTATTTT | 1 | TRBV17*00 | | TRBJ1-3*00 | 0.24% |
| Treg WT2 | TGTGCTAGCAGTAGTGCAGGGGGAGCTGAGCAGTTCTTC | 1 | TRBV17*00 | TRBD1*00 | TRBJ2-1*00 | 0.24% |
| Treg WT2 | TGTGCTAGCAGTCGACTGGACCAAGACACCCAGTACTTT | 1 | TRBV17*00 | TRBD2*00 | TRBJ2-5*00 | 0.24% |
| Treg WT2 | TGTGCTAGCAGTGATGGAAATTCTGGAAATACGCTCTATTTT | 1 | TRBV13-1*00 | | TRBJ1-3*00 | 0.24% |
| Treg WT2 | TGTGCTAGCAGTGGGACAGGGAAATATGCTGAGCAGTTCTTC | 1 | TRBV17*00 | TRBD1*00 | TRBJ2-1*00 | 0.24% |
| Treg WT2 | TGTGCTAGCAGTTCGAACAGGGTTAACCAAGACACCCAGTACTTT | 1 | TRBV29*00 | | TRBJ2-5*00 | 0.24% |
| Treg WT2 | TGTGCTAGCAGTTTACCGCAGGCAAACACCGGGCAGCTCTACTTT | 1 | TRBV29*00 | | TRBJ2-2*00 | 0.24% |
| Treg WT2 | TGTGCTAGCAGTTTAGGGGGGGGCGCTGAGCAGTTCTTC | 1 | TRBV29*00 | TRBD2*00 | TRBJ2-1*00 | 0.24% |
| Treg WT2 | TGTGCTAGCAGTTTCTGGGAAAGCTCCTATGAACAGTACTTC | 1 | TRBV29*00 | TRBD2*00 | TRBJ2-7*00 | 0.24% |
| Treg WT2 | TGTGGTGCTAGCAGGGGCAACCAGGCTCCGCTTTTT | 1 | TRBV20*00 | TRBD1*00 | TRBJ1-5*00 | 0.24% |
| Treg WT2 | TGTGGTGCTAGGACACAAGACACCCAGTACTTT | 1 | TRBV20*00 | | TRBJ2-5*00 | 0.24% |
| Treg WT2 | TGTGGTGCTAGGCGGTGGGACACCCAGTACTTT | 1 | TRBV20*00 | | TRBJ2-5*00 | 0.24% |
| Treg WT2 | TGTGGTGCTAGGGAACCCACAGGGAGAGTCTTCTTT | 1 | TRBV20*00 | TRBD1*00 | TRBJ1-1*00 | 0.24% |
| Treg WT2 | TGTGGTGCTAGGGAAGGGACAGGGGGTGAACAGTACTTC | 1 | TRBV20*00 | TRBD1*00 | TRBJ2-7*00 | 0.24% |
| Treg WT2 | TGTGGTGCTAGGGACACCGGGCAGCTCTACTTT | 1 | TRBV20*00 | | TRBJ2-2*00 | 0.24% |
| Treg WT2 | TGTGGTGCTAGGGAGACATCTAACCAAGACACCCAGTACTTT | 1 | TRBV20*00 | | TRBJ2-5*00 | 0.24% |
| Treg WT2 | TGTGGTGCTAGGGATCAGGGAGACACCCAGTACTTT | 1 | TRBV20*00 | | TRBJ2-5*00 | 0.24% |
| Treg WT2 | TGTGGTGCTAGGGGACAGGGGGCAGAAACGCTGTATTTT | 1 | TRBV20*00 | TRBD1*00 | TRBJ2-3*00 | 0.24% |
| Treg WT2 | TGTGGTGCTAGGGGACGAAACTATGCTGAGCAGTTCTTC | 1 | TRBV20*00 | | TRBJ2-1*00 | 0.24% |
| Treg WT2 | TGTGGTGCTAGGGGCCCAAACTCCGACTACACCTTC | 1 | TRBV20*00 | | TRBJ1-2*00 | 0.24% |
| Treg WT2 | TGTGGTGCTAGGGTGGGCCAAAACACCTTGTACTTT | 1 | TRBV20*00 | | TRBJ2-4*00 | 0.24% |
| Treg WT2 | TGTGGTGCTCGCTCCTGGGGGGGTGGTGAACAGTACTTC | 1 | TRBV20*00 | TRBD2*00 | TRBJ2-7*00 | 0.24% |
| Treg WT2 | TGTGCCAGCGGTGACAGTTATGAACAGTACTTC | 1 | TRBV13-2*00 | | TRBJ2-7*00 | 0.24% |
| Treg WT2 | TGTGCCAGCAGTCTGTCAACGTATGAACAGTACTTC | 1 | TRBV26*00 | | TRBJ2-7*00 | 0.24% |
| Treg WT2 | TGTGCTAGCAGTAGGGACAACAACGAAAGATTATTTTTC | 1 | TRBV17*00 | TRBD1*00 | TRBJ1-4*00 | 0.24% |
| Treg WT2 | TGTGCCAGCGGTGACAGGTATGAACAGTACTTC | 1 | TRBV13-2*00 | | TRBJ2-7*00 | 0.24% |
| Treg WT2 | TGTGCCAGCAGTAACCAAGACACCCAGTACTTT | 1 | TRBV15*00 | | TRBJ2-5*00 | 0.24% |
| Treg WT2 | TGTGCCAGCAGTCTGTCAGGGTATGAACAGTACTTC | 1 | TRBV26*00 | | TRBJ2-7*00 | 0.24% |
| Treg WT2 | TGTGCCAGCAGCTTAGGGACAGAAGTCTTCTTT | 1 | TRBV3*00 | | TRBJ1-1*00 | 0.24% |

#### Treg DKO

| Sample Name | CDR3 Sequence | clone Count | best V Hit | best D Hit | best J Hit | Frequency |
|---|---|---|---|---|---|---|
| Treg DKO | TGTGTCAGCAGTTTTGGGGTCTATGCTGAGCAGTTCTTC | 51 | TRBV14*00 | TRBD2*00 | TRBJ2-1*00 | 12.11% |
| Treg DKO | TGTGCCAGCAGCCAGGACTGGCAAGACACCCAGTACTTT | 19 | TRBV2*00 | TRBD2*00 | TRBJ2-5*00 | 4.51% |
| Treg DKO | TGTGCTAGCAGTTTCCAGGGTGCAGAAACGCTGTATTTT | 11 | TRBV29*00 | | TRBJ2-3*00 | 2.61% |
| Treg DKO | TGTGCCAGCAGTTTTGGGGTCTATGCTGAGCAGTTCTTC | 9 | TRBV14*00 | TRBD2*00 | TRBJ2-1*00 | 2.14% |
| Treg DKO | TGTGCCAGCAGTGATGCGGGCACGTATAACAACCAGGCTCCGCTTTTT | 6 | TRBV13-3*00 | | TRBJ1-5*00 | 1.43% |
| Treg DKO | TGTGCCAGCGGTGATCAGGGAAATACGCTCTATTTT | 4 | TRBV13-2*00 | | TRBJ1-3*00 | 0.95% |

| | | | | | | |
|---|---|---|---|---|---|---|
| Treg DKO | TGTGGTGCTAGGCCTGGGGGGTATGAACAGTACTTC | 4 | TRBV20*00 | TRBD2*00 | TRBJ2-7*00 | 0.95% |
| Treg DKO | TGCACCTGCAGTGCAGGGGCCAACCAGGCTCCGCTTTTT | 3 | TRBV1*00 | | TRBJ1-5*00 | 0.71% |
| Treg DKO | TGCTCCAGCAGTCAATCTCCAGGGAACTATGAACAGTACTTC | 3 | TRBV23*00 | TRBD1*00 | TRBJ2-7*00 | 0.71% |
| Treg DKO | TGTGCCTGGAGTCTAGTTGGGGGGGCGTCTGCTGAGCAGTTCTTC | 3 | TRBV31*00 | TRBD2*00 | TRBJ2-1*00 | 0.71% |
| Treg DKO | TGTGGTGCTAGGCTGACTGGACATAACTATGCTGAGCAGTTCTTC | 3 | TRBV20*00 | TRBD2*00 | TRBJ2-1*00 | 0.71% |
| Treg DKO | TGCACCTGCAGTGGGACTGGGGGTTATGAACAGTACTTC | 2 | TRBV1*00 | TRBD2*00 | TRBJ2-7*00 | 0.48% |
| Treg DKO | TGTGCAAGCAGCCGGGGACAGGACTCCTATGAACAGTACTTC | 2 | TRBV16*00 | TRBD1*00 | TRBJ2-7*00 | 0.48% |
| Treg DKO | TGTGCCAGCAGCCAAGACAGTTTTGCTGAGCAGTTCTTC | 2 | TRBV5*00 | | TRBJ2-1*00 | 0.48% |
| Treg DKO | TGTGCCAGCAGCCAAGTGGGGCGGCAAGACACCCAGTACTTT | 2 | TRBV5*00 | TRBD1*00 | TRBJ2-5*00 | 0.48% |
| Treg DKO | TGTGCCAGCAGCCAGGACTGGCAAGACACCCAGTACTTC | 2 | TRBV2*00 | TRBD2*00 | TRBJ2-5*00 | 0.48% |
| Treg DKO | TGTGCCAGCAGCCCAGGGCCCTCTAGTGCAGAAACGCTGTATTTT | 2 | TRBV2*00 | TRBD1*00 | TRBJ2-3*00 | 0.48% |
| Treg DKO | TGTGCCAGCAGCCCCTCGGGAAACACCTTGTACTTT | 2 | TRBV14*00 | TRBD1*00 | TRBJ2-4*00 | 0.48% |
| Treg DKO | TGTGCCAGCAGCTGGACAGTTTCCTATGAACAGTACTTC | 2 | TRBV25*00 | TRBD1*00 | TRBJ2-7*00 | 0.48% |
| Treg DKO | TGTGCCAGCAGCTTAGATACAAACTCCGACTACACCTTC | 2 | TRBV3*00 | | TRBJ1-2*00 | 0.48% |
| Treg DKO | TGTGCCAGCAGCTTGGGGGGGCGTGCAGAAACGCTGTATTTT | 2 | TRBV3*00 | TRBD2*00 | TRBJ2-3*00 | 0.48% |
| Treg DKO | TGTGCCAGCAGGGACATTCAAAACACCTTGTACTTT | 2 | TRBV15*00 | TRBD1*00 | TRBJ2-4*00 | 0.48% |
| Treg DKO | TGTGCCAGCAGGGGACAGAGCTCCTATGAACAGTACTTC | 2 | TRBV3*00 | TRBD1*00 | TRBJ2-7*00 | 0.48% |
| Treg DKO | TGTGCCAGCAGTCGGACAGGGGGGGACTATGCTGAGCAGTTCTTC | 2 | TRBV13-3*00 | TRBD1*00 | TRBJ2-1*00 | 0.48% |
| Treg DKO | TGTGCCAGCAGTGACAGGGCTCAAGACACCCAGTACTTT | 2 | TRBV13-3*00 | TRBD1*00 | TRBJ2-5*00 | 0.48% |
| Treg DKO | TGTGCCAGCAGTGAGCAATACTATGCTGAGCAGTTCTTC | 2 | TRBV13-1*00 | | TRBJ2-1*00 | 0.48% |
| Treg DKO | TGTGCCAGCAGTGAGGACACCAATAATTCGCCCCTCTACTTT | 2 | TRBV13-1*00 | TRBD1*00 | TRBJ1-6*00 | 0.48% |
| Treg DKO | TGTGCCAGCAGTGATCCCGACAGTTCTGGAAATACGCTCTATTTT | 2 | TRBV13-1*00 | TRBD1*00 | TRBJ1-3*00 | 0.48% |
| Treg DKO | TGTGCCAGCAGTGATGATAACTCCGACTACACCTTC | 2 | TRBV13-1*00 | | TRBJ1-2*00 | 0.48% |
| Treg DKO | TGTGCCAGCAGTGATGCGGGGTTCGAAAGATTATTTTTC | 2 | TRBV13-1*00 | | TRBJ1-4*00 | 0.48% |
| Treg DKO | TGTGCCAGCAGTGATTGGGGACATCGCTCCTATGAACAGTACTTC | 2 | TRBV13-1*00 | TRBD1*00 | TRBJ2-7*00 | 0.48% |
| Treg DKO | TGTGCCAGCAGTGCAGCAAACTCCGACTACACCTTC | 2 | TRBV13-1*00 | | TRBJ1-2*00 | 0.48% |
| Treg DKO | TGTGCCAGCAGTTTAAGGGCAGCAGAAACGCTGTATTTT | 2 | TRBV15*00 | | TRBJ2-3*00 | 0.48% |
| Treg DKO | TGTGCCAGCGGTGACAGCTATAATTCGCCCCTCTACTTT | 2 | TRBV13-2*00 | | TRBJ1-6*00 | 0.48% |
| Treg DKO | TGTGCCTGGAGTCGTGGGGGTAGTGCAGAAACGCTGTATTTT | 2 | TRBV31*00 | TRBD2*00 | TRBJ2-3*00 | 0.48% |
| Treg DKO | TGTGCCAGCTCTCTCGACGGTGGGGGGAAAGACACCCAGTACTTT | 2 | TRBV12-1*00 | TRBD2*00 | TRBJ2-5*00 | 0.48% |
| Treg DKO | TGTGCTAGCAGTCCCGCCCGTTCTGGAAATACGCTCTATTTT | 2 | TRBV29*00 | TRBD1*00 | TRBJ1-3*00 | 0.48% |
| Treg DKO | TGCACCTACAGTGCCCGGGACAGTGCAGAAACGCTGTATTTT | 1 | TRBV1*00 | TRBD1*00 | TRBJ2-3*00 | 0.24% |
| Treg DKO | TGCACCTGCAGCCCAGACTGGGGGGGCTATGAACAGTACTTC | 1 | TRBV1*00 | TRBD2*00 | TRBJ2-7*00 | 0.24% |
| Treg DKO | TGCACCTGCAGTCATCCCTCCTATGAACAGTACTTC | 1 | TRBV1*00 | | TRBJ2-7*00 | 0.24% |
| Treg DKO | TGCACCTGCAGTGCAGAAAGTCAAAACACCTTGTACTTT | 1 | TRBV1*00 | | TRBJ2-4*00 | 0.24% |
| Treg DKO | TGCACCTGCAGTGCAGAAGCTGGGGGTTATGAACAGTACTTC | 1 | TRBV1*00 | TRBD2*00 | TRBJ2-7*00 | 0.24% |
| Treg DKO | TGCACCTGCAGTGCAGGCCGGGACACCCAGTACTTT | 1 | TRBV1*00 | | TRBJ2-5*00 | 0.24% |
| Treg DKO | TGCACCTGCAGTGCCCGGACAGGGGGCTCCGACTACACCTTC | 1 | TRBV1*00 | TRBD1*00 | TRBJ1-2*00 | 0.24% |
| Treg DKO | TGCACCTGCAGTGCGCCGGGACAGAACTATGCTGAGCAGTTCTTC | 1 | TRBV1*00 | TRBD1*00 | TRBJ2-1*00 | 0.24% |
| Treg DKO | TGCACCTGCAGTGCGGGGGCCAACCAGGCTCCGCTTTTT | 1 | TRBV1*00 | TRBD1*00 | TRBJ1-5*00 | 0.24% |
| Treg DKO | TGCACCTGCAGTGTGCAGCCAAACTCCGACTACACCTTC | 1 | TRBV1*00 | | TRBJ1-2*00 | 0.24% |
| Treg DKO | TGCACCTGCAGTTCGTATGAACAGTACTTC | 1 | TRBV1*00 | | TRBJ2-7*00 | 0.24% |
| Treg DKO | TGCTCCAGCAGTCAATCAGGCAACGAAAGATTATTTTTC | 1 | TRBV23*00 | | TRBJ1-4*00 | 0.24% |
| Treg DKO | TGCTCCAGCAGTCAATCCTGGGGGGAATATGAACAGTACTTC | 1 | TRBV23*00 | TRBD2*00 | TRBJ2-7*00 | 0.24% |
| Treg DKO | TGCTCCAGCAGTCAATCTCCAGGGAACTATGAACAGTATTTC | 1 | TRBV23*00 | TRBD1*00 | TRBJ2-7*00 | 0.24% |
| Treg DKO | TGTAGTTCTAGAAAGCAGGGGGCCACAGAAGTCTTCTTT | 1 | TRBV30*00 | TRBD1*00 | TRBJ1-1*00 | 0.24% |
| Treg DKO | TGTAGTTCTAGAGATCGCAGATATGAACAGTACTTC | 1 | TRBV30*00 | | TRBJ2-7*00 | 0.24% |
| Treg DKO | TGTAGTTCTGTGGGACAGGGAACGCTGTATTTT | 1 | TRBV30*00 | TRBD1*00 | TRBJ2-3*00 | 0.24% |
| Treg DKO | TGTGCAAGCAGCACAGGGGGAGACAACGAAAGATTATTTTTC | 1 | TRBV16*00 | TRBD1*00 | TRBJ1-4*00 | 0.24% |
| Treg DKO | TGTGCAAGCAGCGAGGGTTACACAGAAGTCTTCTTT | 1 | TRBV16*00 | | TRBJ1-1*00 | 0.24% |
| Treg DKO | TGTGCAAGCAGCTCCGGACGAGAAGTCTTCTTT | 1 | TRBV16*00 | | TRBJ1-1*00 | 0.24% |
| Treg DKO | TGTGCAAGCAGCTGGGGACAGTCTGGAAATACGCTCTATTTT | 1 | TRBV16*00 | TRBD1*00 | TRBJ1-3*00 | 0.24% |
| Treg DKO | TGTGCAAGCAGCTTAAGTGACACAGAAGTCTTCTTT | 1 | TRBV16*00 | | TRBJ1-1*00 | 0.24% |
| Treg DKO | TGTGCAAGCAGCTTACGCAGGTCTAGCTCCTATGAACAGTACTTC | 1 | TRBV16*00 | | TRBJ2-7*00 | 0.24% |
| Treg DKO | TGTGCAAGCAGCTTAGAACAAAACACAGAAGTCTTCTTT | 1 | TRBV16*00 | | TRBJ1-1*00 | 0.24% |
| Treg DKO | TGTGCAAGCAGCTTAGAAGGGGACACCCAGTACTTT | 1 | TRBV16*00 | | TRBJ2-5*00 | 0.24% |
| Treg DKO | TGTGCAAGCAGCTTAGAAGGGGATACGCTCTATTTT | 1 | TRBV16*00 | TRBD1*00 | TRBJ1-3*00 | 0.24% |
| Treg DKO | TGTGCAAGCAGCTTAGACTATGAACAGTACTTC | 1 | TRBV16*00 | | TRBJ2-7*00 | 0.24% |
| Treg DKO | TGTGCAAGCAGCTTAGAGGGAAACACAGAAGTCTTCTTT | 1 | TRBV16*00 | | TRBJ1-1*00 | 0.24% |
| Treg DKO | TGTGCAAGCAGCTTAGAGTGGGGTGGAGACACCCAGTACTTT | 1 | TRBV16*00 | TRBD2*00 | TRBJ2-5*00 | 0.24% |
| Treg DKO | TGTGCAAGCAGCTTAGCAAACACCGGGCAGCTCTACTTT | 1 | TRBV16*00 | | TRBJ2-2*00 | 0.24% |
| Treg DKO | TGTGCAAGCAGCTTAGGACATCGAAACACAGAAGTCTTCTTT | 1 | TRBV16*00 | | TRBJ1-1*00 | 0.24% |
| Treg DKO | TGTGCAAGCAGCTTAGGGGACAGAGACACCCAGTACTTT | 1 | TRBV16*00 | TRBD1*00 | TRBJ2-5*00 | 0.24% |
| Treg DKO | TGTGCAAGCAGCTTAGTTGGGGGGGGATATGAACAGTACTTC | 1 | TRBV16*00 | TRBD2*00 | TRBJ2-7*00 | 0.24% |
| Treg DKO | TGTGCAAGCAGCTTGCAGAACACAGAAGTCTTCTTT | 1 | TRBV16*00 | | TRBJ1-1*00 | 0.24% |
| Treg DKO | TGTGCCAGCAACACTGGGGGTAGTGCAGAAACGCTGTATTTT | 1 | TRBV13-3*00 | TRBD2*00 | TRBJ2-3*00 | 0.24% |
| Treg DKO | TGTGCCAGCAACTTCAGGGATGAACAGTACTTC | 1 | TRBV4*00 | TRBD1*00 | TRBJ2-7*00 | 0.24% |
| Treg DKO | TGTGCCAGCACCAGAGACACCGGGCAGCTCTACTTT | 1 | TRBV15*00 | | TRBJ2-2*00 | 0.24% |
| Treg DKO | TGTGCCAGCACCCCGGGAAATACGCTCTATTTT | 1 | TRBV15*00 | | TRBJ1-3*00 | 0.24% |
| Treg DKO | TGTGCCAGCAGAAGACAGTATAACAACCAGGCTCCGCTTTTT | 1 | TRBV15*00 | TRBD1*00 | TRBJ1-5*00 | 0.24% |
| Treg DKO | TGTGCCAGCAGACTGGGGAGAGGCCAAGACACCCAGTACTTT | 1 | TRBV13-3*00 | TRBD2*00 | TRBJ2-5*00 | 0.24% |
| Treg DKO | TGTGCCAGCAGCAAGAGCTCCTATGAACAGTACTTC | 1 | TRBV13-1*00 | | TRBJ2-7*00 | 0.24% |
| Treg DKO | TGTGCCAGCAGCACCTCAGGGGGCGCTGAGCAGTTCTTC | 1 | TRBV4*00 | TRBD1*00 | TRBJ2-1*00 | 0.24% |
| Treg DKO | TGTGCCAGCAGCATAACAGGGAATGAACAGTACTTC | 1 | TRBV13-3*00 | TRBD1*00 | TRBJ2-7*00 | 0.24% |
| Treg DKO | TGTGCCAGCAGCCAAAAGACAGTGGGTAACTTC | 1 | TRBV2*00 | TRBD1*00 | TRBJ2-7*00 | 0.24% |

| | | | | | | |
|---|---|---|---|---|---|---|
| Treg DKO | TGTGCCAGCAGCCAACAGGGAAGGGGTGCAGAAACGCTGTATTTT | 1 | TRBV5*00 | TRBD1*00 | TRBJ2-3*00 | 0.24% |
| Treg DKO | TGTGCCAGCAGCCAACAGGGGTATGAACAGTACTTC | 1 | TRBV2*00 | TRBD1*00 | TRBJ2-7*00 | 0.24% |
| Treg DKO | TGTGCCAGCAGCCAACAGGTTTCCTATAATTCGCCCCTCTACTTT | 1 | TRBV5*00 | | TRBJ1-6*00 | 0.24% |
| Treg DKO | TGTGCCAGCAGCCAACGGGGGGACGAAAGATTATTTTTC | 1 | TRBV5*00 | TRBD2*00 | TRBJ1-4*00 | 0.24% |
| Treg DKO | TGTGCCAGCAGCCAAGAAGACACAGAAGTCTTCTTT | 1 | TRBV5*00 | | TRBJ1-1*00 | 0.24% |
| Treg DKO | TGTGCCAGCAGCCAAGAAGACTGGGGGGTCAACTATGCTGAGCAGTTCTTC | 1 | TRBV2*00 | TRBD2*00 | TRBJ2-1*00 | 0.24% |
| Treg DKO | TGTGCCAGCAGCCAAGACAATTATGCTGAGCAGTTCTTC | 1 | TRBV5*00 | | TRBJ2-1*00 | 0.24% |
| Treg DKO | TGTGCCAGCAGCCAAGACACTGTCTATGAACAGTACTTC | 1 | TRBV2*00 | | TRBJ2-7*00 | 0.24% |
| Treg DKO | TGTGCCAGCAGCCAAGACAGCAACACCGGGCAGCTCTACTTT | 1 | TRBV2*00 | | TRBJ2-2*00 | 0.24% |
| Treg DKO | TGTGCCAGCAGCCAAGACAGGGGTGGAGACACCCAATACTTT | 1 | TRBV5*00 | TRBD1*00 | TRBJ2-5*00 | 0.24% |
| Treg DKO | TGTGCCAGCAGCCAAGACCCAAACACCGGGCAGCTCTACTTT | 1 | TRBV5*00 | | TRBJ2-2*00 | 0.24% |
| Treg DKO | TGTGCCAGCAGCCAAGACGACTGGGGGGGCGCCGCGCTGTATTTT | 1 | TRBV5*00 | TRBD2*00 | TRBJ2-3*00 | 0.24% |
| Treg DKO | TGTGCCAGCAGCCAAGACTGGGGGAGTGCTGAGCAGTTCTTC | 1 | TRBV2*00 | TRBD2*00 | TRBJ2-1*00 | 0.24% |
| Treg DKO | TGTGCCAGCAGCCAAGACTGGGGTAACCAAGACACCCAGTACTTT | 1 | TRBV5*00 | TRBD2*00 | TRBJ2-5*00 | 0.24% |
| Treg DKO | TGTGCCAGCAGCCAAGACTGGTCTAGTCAAAACACCTTGTACTTT | 1 | TRBV5*00 | | TRBJ2-4*00 | 0.24% |
| Treg DKO | TGTGCCAGCAGCCAAGACTTTTACACCTTGTACTTT | 1 | TRBV5*00 | | TRBJ2-4*00 | 0.24% |
| Treg DKO | TGTGCCAGCAGCCAAGAGGACTATGAACAGTACTTC | 1 | TRBV5*00 | | TRBJ2-7*00 | 0.24% |
| Treg DKO | TGTGCCAGCAGCCAAGAGGCAGGAAACTATGCTGAGCAGTTCTTC | 1 | TRBV5*00 | | TRBJ2-1*00 | 0.24% |
| Treg DKO | TGTGCCAGCAGCCAAGAGGGAGGGGAACAGTACTTC | 1 | TRBV5*00 | | TRBJ2-7*00 | 0.24% |
| Treg DKO | TGTGCCAGCAGCCAAGTGGGGGAAACGCTGTATTTT | 1 | TRBV5*00 | TRBD2*00 | TRBJ2-3*00 | 0.24% |
| Treg DKO | TGTGCCAGCAGCCACACGACAGGGGATGAACAGTACTTC | 1 | TRBV5*00 | TRBD1*00 | TRBJ2-7*00 | 0.24% |
| Treg DKO | TGTGCCAGCAGCCACGACTGGGGGAATGAACAGTACTTC | 1 | TRBV5*00 | TRBD2*00 | TRBJ2-7*00 | 0.24% |
| Treg DKO | TGTGCCAGCAGCCAGACTGGGGGGGGCTATGAACAGTACTTC | 1 | TRBV4*00 | TRBD2*00 | TRBJ2-7*00 | 0.24% |
| Treg DKO | TGTGCCAGCAGCCAGACTGGGGGTGCAAACACCGGGCAGCTCTACTTT | 1 | TRBV5*00 | TRBD2*00 | TRBJ2-2*00 | 0.24% |
| Treg DKO | TGTGCCAGCAGCCAGGAAAAAGACACCCAGTACTTT | 1 | TRBV2*00 | | TRBJ2-5*00 | 0.24% |
| Treg DKO | TGTGCCAGCAGCCAGGGAGACACCCAGTACTTT | 1 | TRBV5*00 | | TRBJ2-5*00 | 0.24% |
| Treg DKO | TGTGCCAGCAGCCCAGGACAGTATGAACAGTACTTC | 1 | TRBV2*00 | TRBD1*00 | TRBJ2-7*00 | 0.24% |
| Treg DKO | TGTGCCAGCAGCCCAGGGGATAACCAAGACACCCAGTACTTT | 1 | TRBV2*00 | TRBD1*00 | TRBJ2-5*00 | 0.24% |
| Treg DKO | TGTGCCAGCAGCCCATACTTC | 1 | TRBV5*00 | | TRBJ2-7*00 | 0.24% |
| Treg DKO | TGTGCCAGCAGCCCCACCGGGACAGAAGTCTTCTTT | 1 | TRBV5*00 | TRBD1*00 | TRBJ1-1*00 | 0.24% |
| Treg DKO | TGTGCCAGCAGCCCCGCCAGCTTTGCAAACACAGAAGTCTTCTTT | 1 | TRBV3*00 | | TRBJ1-1*00 | 0.24% |
| Treg DKO | TGTGCCAGCAGCCCGACTGGGGGGAGTGAACAGTACTTC | 1 | TRBV2*00 | TRBD2*00 | TRBJ2-7*00 | 0.24% |
| Treg DKO | TGTGCCAGCAGCCCGAGACATCAAGACACCCAGTACTTT | 1 | TRBV5*00 | | TRBJ2-5*00 | 0.24% |
| Treg DKO | TGTGCCAGCAGCCCTACTGGGAACCAAGACACCCAGTACTTT | 1 | TRBV2*00 | TRBD2*00 | TRBJ2-5*00 | 0.24% |
| Treg DKO | TGTGCCAGCAGCCCTCGGGACAGCCCTAGTGCAGAAACGCTGTATTTT | 1 | TRBV4*00 | TRBD1*00 | TRBJ2-3*00 | 0.24% |
| Treg DKO | TGTGCCAGCAGCCGACAAGACCAGGCTCCGCTTTTT | 1 | TRBV22*00 | | TRBJ1-5*00 | 0.24% |
| Treg DKO | TGTGCCAGCAGCCGACAGCGGAGCGAAAGATTATTTTTC | 1 | TRBV5*00 | TRBD1*00 | TRBJ1-4*00 | 0.24% |
| Treg DKO | TGTGCCAGCAGCCGACTGGGGGGGGGTGAACAGTACTTC | 1 | TRBV3*00 | TRBD2*00 | TRBJ2-7*00 | 0.24% |
| Treg DKO | TGTGCCAGCAGCCGACTGGGGGTAACTATGCTGAGCAGTTCTTC | 1 | TRBV3*00 | TRBD2*00 | TRBJ2-1*00 | 0.24% |
| Treg DKO | TGTGCCAGCAGCCGGGACACCCAAGACACCCAGTACTTT | 1 | TRBV15*00 | TRBD1*00 | TRBJ2-5*00 | 0.24% |
| Treg DKO | TGTGCCAGCAGCCTCAAGGGATATGAACAGTACTTC | 1 | TRBV2*00 | | TRBJ2-7*00 | 0.24% |
| Treg DKO | TGTGCCAGCAGCCTCCGACAGTATAATTCGCCCCTCTACTTT | 1 | TRBV2*00 | TRBD1*00 | TRBJ1-6*00 | 0.24% |
| Treg DKO | TGTGCCAGCAGCCTTAAAGACACCCAGTACTTT | 1 | TRBV19*00 | | TRBJ2-5*00 | 0.24% |
| Treg DKO | TGTGCCAGCAGCGACCAAGACACCCAGTACTTT | 1 | TRBV3*00 | | TRBJ2-5*00 | 0.24% |
| Treg DKO | TGTGCCAGCAGCTAGACAGTATGCTGAGCAGTTCTTC | 1 | TRBV4*00 | TRBD1*00 | TRBJ2-1*00 | 0.24% |
| Treg DKO | TGTGCCAGCAGCTAT | 1 | TRBV4*00 | | TRBJ2-4*00 | 0.24% |
| Treg DKO | TGTGCCAGCAGCTCACAAGCAAACTCCGACTACACCTTC | 1 | TRBV3*00 | | TRBJ1-2*00 | 0.24% |
| Treg DKO | TGTGCCAGCAGCTCCAGGGGACCAAACACCGGGCAGCTCTACTTT | 1 | TRBV3*00 | TRBD1*00 | TRBJ2-2*00 | 0.24% |
| Treg DKO | TGTGCCAGCAGCTCCAGTAACTCCGACTACACCTTC | 1 | TRBV25*00 | | TRBJ1-2*00 | 0.24% |
| Treg DKO | TGTGCCAGCAGCTCCCGGGACAGCTCCTATGAACAGTACTTC | 1 | TRBV5*00 | TRBD1*00 | TRBJ2-7*00 | 0.24% |
| Treg DKO | TGTGCCAGCAGCTCCTCCGACACCCAGTACTTT | 1 | TRBV13-1*00 | | TRBJ2-5*00 | 0.24% |
| Treg DKO | TGTGCCAGCAGCTCCTGGTCGCAAAACACCTTGTACTTT | 1 | TRBV25*00 | | TRBJ2-4*00 | 0.24% |
| Treg DKO | TGTGCCAGCAGCTGGACAGGGTCCTATGAACAGTACTTC | 1 | TRBV25*00 | TRBD1*00 | TRBJ2-7*00 | 0.24% |
| Treg DKO | TGTGCCAGCAGCTGGGGGAGGGAACAGTACTTC | 1 | TRBV3*00 | TRBD1*00 | TRBJ2-7*00 | 0.24% |
| Treg DKO | TGTGCCAGCAGCTGGGGGTCCGTGCTGAGCAGTTCTTC | 1 | TRBV3*00 | TRBD1*00 | TRBJ2-1*00 | 0.24% |
| Treg DKO | TGTGCCAGCAGCTTAACAAACCAGGCTCCGCTTTTT | 1 | TRBV3*00 | | TRBJ1-5*00 | 0.24% |
| Treg DKO | TGTGCCAGCAGCTTACAGAACTATGAACAGTACTTC | 1 | TRBV3*00 | | TRBJ2-7*00 | 0.24% |
| Treg DKO | TGTGCCAGCAGCTTACAGGGGGACACCCAGTACTTT | 1 | TRBV3*00 | TRBD1*00 | TRBJ2-5*00 | 0.24% |
| Treg DKO | TGTGCCAGCAGCTTACAGGGGGCAAACTATGCTGAGCAGTTCTTC | 1 | TRBV3*00 | TRBD1*00 | TRBJ2-1*00 | 0.24% |
| Treg DKO | TGTGCCAGCAGCTTACGTAGTGCAGAAACGCTGTATTTT | 1 | TRBV3*00 | | TRBJ2-3*00 | 0.24% |
| Treg DKO | TGTGCCAGCAGCTTACTGGGGGGGCATGCTGAGCAGTTCTTC | 1 | TRBV3*00 | TRBD2*00 | TRBJ2-1*00 | 0.24% |
| Treg DKO | TGTGCCAGCAGCTTAGAACAGAATACGCTCTATTTT | 1 | TRBV3*00 | | TRBJ1-3*00 | 0.24% |
| Treg DKO | TGTGCCAGCAGCTTAGAAGACCAAGACACCCAGTACTTT | 1 | TRBV3*00 | | TRBJ2-5*00 | 0.24% |
| Treg DKO | TGTGCCAGCAGCTTAGACAGTCAAAACACCTTGTACTTT | 1 | TRBV3*00 | | TRBJ2-4*00 | 0.24% |
| Treg DKO | TGTGCCAGCAGCTTAGAGGGACAGAACTCCGACTACACCTTC | 1 | TRBV3*00 | TRBD1*00 | TRBJ1-2*00 | 0.24% |
| Treg DKO | TGTGCCAGCAGCTTAGATCAGAACACAGAAGTCTTCTTT | 1 | TRBV3*00 | | TRBJ1-1*00 | 0.24% |
| Treg DKO | TGTGCCAGCAGCTTAGGAGGGGGCGAACGGTACTTC | 1 | TRBV3*00 | TRBD1*00 | TRBJ2-7*00 | 0.24% |
| Treg DKO | TGTGCCAGCAGCTTAGGCGGAAACACCTTGTACTTT | 1 | TRBV3*00 | | TRBJ2-4*00 | 0.24% |
| Treg DKO | TGTGCCAGCAGCTTAGGGAGAAGGTATGCTGAGCAGTTCTTC | 1 | TRBV3*00 | | TRBJ2-1*00 | 0.24% |
| Treg DKO | TGTGCCAGCAGCTTAGTCAGCCCCTATGAACAGTACTTC | 1 | TRBV3*00 | | TRBJ2-7*00 | 0.24% |
| Treg DKO | TGTGCCAGCAGCTTAGTCGCAGGAGCAAACTCCGACTACACCTTC | 1 | TRBV3*00 | | TRBJ1-2*00 | 0.24% |
| Treg DKO | TGTGCCAGCAGCTTAGTGACAGGGGGCAACTCCGACTACACCTCC | 1 | TRBV3*00 | TRBD1*00 | TRBJ1-2*00 | 0.24% |
| Treg DKO | TGTGCCAGCAGCTTAGTGGGGCACCAAGACACCCAGTACTTT | 1 | TRBV3*00 | TRBD1*00 | TRBJ2-5*00 | 0.24% |
| Treg DKO | TGTGCCAGCAGCTTATTACACACAGAAGTCTTCTTT | 1 | TRBV3*00 | | TRBJ1-1*00 | 0.24% |
| Treg DKO | TGTGCCAGCAGCTTCAGGGGTCAAAACACCTTGTACTTT | 1 | TRBV3*00 | TRBD1*00 | TRBJ2-4*00 | 0.24% |

| | | | | | | |
|---|---|---|---|---|---|---|
| Treg DKO | TGTGCCAGCAGCTTCGCACAGACAGAAGTCTTCTTT | 1 | TRBV3*00 | | TRBJ1-1*00 | 0.24% |
| Treg DKO | TGTGCCAGCAGCTTCGTGGGCTCCGACTACACCTTC | 1 | TRBV3*00 | | TRBJ1-2*00 | 0.24% |
| Treg DKO | TGTGCCAGCAGGACCCAGGCAAACTCCGACTACACCTTC | 1 | TRBV13-3*00 | TRBD2*00 | TRBJ1-2*00 | 0.24% |
| Treg DKO | TGTGCCAGCAGGCAGGGCAACGAAAGATTATTTTTC | 1 | TRBV14*00 | TRBD1*00 | TRBJ1-4*00 | 0.24% |
| Treg DKO | TGTGCCAGCAGGCGACAGTATGAACAGTACTTC | 1 | TRBV19*00 | TRBD1*00 | TRBJ2-7*00 | 0.24% |
| Treg DKO | TGTGCCAGCAGGGAAGGCGGGAACCAAGACACCCAGTACTTT | 1 | TRBV3*00 | | TRBJ2-5*00 | 0.24% |
| Treg DKO | TGTGCCAGCAGGGACAACCAAGACACCCAGTACTTT | 1 | TRBV13-3*00 | | TRBJ2-5*00 | 0.24% |
| Treg DKO | TGTGCCAGCAGGGGTGACACAGAAGTCTTCTTT | 1 | TRBV13-1*00 | | TRBJ1-1*00 | 0.24% |
| Treg DKO | TGTGCCAGCAGTACAACAGGGTATGAACAGTACTTC | 1 | TRBV13-1*00 | TRBD1*00 | TRBJ2-7*00 | 0.24% |
| Treg DKO | TGTGCCAGCAGTATAGACAATAGTCAAAACACCTTGTACTTT | 1 | TRBV19*00 | | TRBJ2-4*00 | 0.24% |
| Treg DKO | TGTGCCAGCAGTATAGACAGTGCAGAAACGCTGTATTTT | 1 | TRBV19*00 | | TRBJ2-3*00 | 0.24% |
| Treg DKO | TGTGCCAGCAGTATAGGACAGAACACAGAAGTCTTCTTT | 1 | TRBV19*00 | TRBD1*00 | TRBJ1-1*00 | 0.24% |
| Treg DKO | TGTGCCAGCAGTATAGTTGGGGGGGCGTGAACAGTACTTC | 1 | TRBV19*00 | TRBD2*00 | TRBJ2-7*00 | 0.24% |
| Treg DKO | TGTGCCAGCAGTATCCAGGGAAACTCCGACTACACCTTC | 1 | TRBV19*00 | TRBD1*00 | TRBJ1-2*00 | 0.24% |
| Treg DKO | GTGCCAGCAGTATCCCGGGACTGGGGGGCCCTAGTGCAGAAACGCTGTATTT | 1 | TRBV19*00 | TRBD2*00 | TRBJ2-3*00 | 0.24% |
| Treg DKO | TGTGCCAGCAGTATCCTGACTGGGGCGAGCTCCTATGAACAGTACTTC | 1 | TRBV19*00 | TRBD2*00 | TRBJ2-7*00 | 0.24% |
| Treg DKO | TGTGCCAGCAGTCCAGAACAAGACACCCAGTACTTT | 1 | TRBV24*00 | | TRBJ2-5*00 | 0.24% |
| Treg DKO | TGTGCCAGCAGTCCTCAGGAAAACACCTTGTACTTT | 1 | TRBV13-3*00 | | TRBJ2-4*00 | 0.24% |
| Treg DKO | TGTGCCAGCAGTCCTGGGACGGAGGACACCCAGTACTTT | 1 | TRBV15*00 | TRBD1*00 | TRBJ2-5*00 | 0.24% |
| Treg DKO | TGTGCCAGCAGTCGAAACACAGAAGTCTTCTTT | 1 | TRBV19*00 | | TRBJ1-1*00 | 0.24% |
| Treg DKO | TGTGCCAGCAGTCGACACAACGAAAGATTATTTTTC | 1 | TRBV15*00 | | TRBJ1-4*00 | 0.24% |
| Treg DKO | TGTGCCAGCAGTCTAAGGAGTGTAGAAACGCTGTATTTT | 1 | TRBV26*00 | | TRBJ2-3*00 | 0.24% |
| Treg DKO | TGTGCCAGCAGTCTCTGGGGGCGCTATGCTGAGCAGTTCTTC | 1 | TRBV13-1*00 | TRBD1*00 | TRBJ2-1*00 | 0.24% |
| Treg DKO | TGTGCCAGCAGTCTGAGGCAAAACACCGGGCAGCTCTACTTT | 1 | TRBV26*00 | | TRBJ2-2*00 | 0.24% |
| Treg DKO | TGTGCCAGCAGTCTGATTGGTAACCAAGACACCCAGTACTTT | 1 | TRBV26*00 | | TRBJ2-5*00 | 0.24% |
| Treg DKO | TGTGCCAGCAGTCTGGGACAGGGGGCTGGAAATACGCTCTATTTT | 1 | TRBV26*00 | TRBD1*00 | TRBJ1-3*00 | 0.24% |
| Treg DKO | TGTGCCAGCAGTCTGGGGCTCTATGCTGAGCAGTTCTTC | 1 | TRBV13-1*00 | TRBD2*00 | TRBJ2-1*00 | 0.24% |
| Treg DKO | TGTGCCAGCAGTCTGTCGACAGCCAACGAAAGATTATTTTTC | 1 | TRBV26*00 | | TRBJ1-4*00 | 0.24% |
| Treg DKO | TGTGCCAGCAGTCTGTCTTCTCAAAACACCTTGTACTTT | 1 | TRBV26*00 | | TRBJ2-4*00 | 0.24% |
| Treg DKO | TGTGCCAGCAGTCTTCCGGGACAAAACACAGAAGTCTTCTTT | 1 | TRBV26*00 | TRBD1*00 | TRBJ1-1*00 | 0.24% |
| Treg DKO | TGTGCCAGCAGTGAAGCCGGGACGGGGGCCGGGCAGCTCTACTTT | 1 | TRBV13-3*00 | TRBD1*00 | TRBJ2-2*00 | 0.24% |
| Treg DKO | TGTGCCAGCAGTGAAGGGGGGGGAAGTCTTCTTT | 1 | TRBV13-1*00 | TRBD1*00 | TRBJ1-1*00 | 0.24% |
| Treg DKO | TGTGCCAGCAGTGAAGGTTCTGGAAATACGCTCTATTTT | 1 | TRBV13-3*00 | | TRBJ1-3*00 | 0.24% |
| Treg DKO | TGTGCCAGCAGTGACAAAACAAACACAGAAGTCTTCTTT | 1 | TRBV13-1*00 | | TRBJ1-1*00 | 0.24% |
| Treg DKO | TGTGCCAGCAGTGACATTAACCAAGACACCCAGTACTTT | 1 | TRBV13-3*00 | | TRBJ2-5*00 | 0.24% |
| Treg DKO | TGTGCCAGCAGTGACCAAGACACCCAGTACTTT | 1 | TRBV13-1*00 | | TRBJ2-5*00 | 0.24% |
| Treg DKO | TGTGCCAGCAGTGACGGGGGACAAAACACCTTGTACTTT | 1 | TRBV13-3*00 | TRBD1*00 | TRBJ2-4*00 | 0.24% |
| Treg DKO | TGTGCCAGCAGTGACTCCTATGCTGAGCAGTTCTTC | 1 | TRBV13-3*00 | | TRBJ2-1*00 | 0.24% |
| Treg DKO | TGTGCCAGCAGTGACTGGGGGGGCGATGCTAAGCAGTTCTTC | 1 | TRBV13-3*00 | TRBD2*00 | TRBJ2-1*00 | 0.24% |
| Treg DKO | TGTGCCAGCAGTGAGCGGGACTGGGGGTATGCTGAGCAGTTCTTC | 1 | TRBV13-3*00 | TRBD2*00 | TRBJ2-1*00 | 0.24% |
| Treg DKO | TGTGCCAGCAGTGAGGACACCAATAATTCACCCCTCTACTTT | 1 | TRBV13-1*00 | TRBD1*00 | TRBJ1-6*00 | 0.24% |
| Treg DKO | TGTGCCAGCAGTGAGGGACTGGGGGGGCAAGACACCCAGTACTTT | 1 | TRBV13-1*00 | TRBD2*00 | TRBJ2-5*00 | 0.24% |
| Treg DKO | TGTGCCAGCAGTGATCAGGGAGGCACAGAAGTCTTCTTT | 1 | TRBV13-1*00 | TRBD1*00 | TRBJ1-1*00 | 0.24% |
| Treg DKO | TGTGCCAGCAGTGATCGGGGGGGCGCCTATGAACAGTACTTC | 1 | TRBV13-3*00 | TRBD2*00 | TRBJ2-7*00 | 0.24% |
| Treg DKO | TGTGCCAGCAGTGATGACAACTATGAACAGTACTTC | 1 | TRBV13-1*00 | | TRBJ2-7*00 | 0.24% |
| Treg DKO | TGTGCCAGCAGTGATGAGGGCTTTAACCAAGACACCCAGTACTTT | 1 | TRBV13-1*00 | | TRBJ2-5*00 | 0.24% |
| Treg DKO | TGTGCCAGCAGTGATGGCAGAAACTCCGACTACACCTTC | 1 | TRBV13-1*00 | | TRBJ1-2*00 | 0.24% |
| Treg DKO | TGTGCCAGCAGTGATGGCCTGGGGGGGCCAGAACAGTACTTC | 1 | TRBV13-1*00 | TRBD2*00 | TRBJ2-7*00 | 0.24% |
| Treg DKO | TGTGCCAGCAGTGATGGGGGTAGTGCAGAAACGCTGTATTTT | 1 | TRBV13-3*00 | | TRBJ2-3*00 | 0.24% |
| Treg DKO | TGTGCCAGCAGTGATTCTGGGGGTCCTCCTGAGCAGTTCTTC | 1 | TRBV13-3*00 | TRBD2*00 | TRBJ2-1*00 | 0.24% |
| Treg DKO | TGTGCCAGCAGTGATTTGGGGGGCGCTGAACAGTACTTC | 1 | TRBV13-3*00 | TRBD2*00 | TRBJ2-7*00 | 0.24% |
| Treg DKO | TGTGCCAGCAGTGCCACAGGGGACTATGCTGAGCAGTTCTTC | 1 | TRBV13-3*00 | TRBD1*00 | TRBJ2-1*00 | 0.24% |
| Treg DKO | TGTGCCAGCAGTGGGACTGGGGGTAGTGCAGAAACGCTGTATTTT | 1 | TRBV13-3*00 | TRBD2*00 | TRBJ2-3*00 | 0.24% |
| Treg DKO | TGTGCCAGCAGTGGGGACAGATATAATTCGCCCCTCTACTTT | 1 | TRBV13-1*00 | TRBD1*00 | TRBJ1-6*00 | 0.24% |
| Treg DKO | TGTGCCAGCAGTTACAATGGAAATACGCTCTATTTT | 1 | TRBV15*00 | | TRBJ1-3*00 | 0.24% |
| Treg DKO | TGTGCCAGCAGTTATAATTCGCCCCTCTACTTT | 1 | TRBV14*00 | | TRBJ1-6*00 | 0.24% |
| Treg DKO | TGTGCCAGCAGTTCAGGCCTTTCCAACGAAAGATTATTTTTC | 1 | TRBV15*00 | | TRBJ1-4*00 | 0.24% |
| Treg DKO | TGTGCCAGCAGTTGGGACCTCACAGAAGTCTTCTTT | 1 | TRBV22*00 | | TRBJ1-1*00 | 0.24% |
| Treg DKO | TGTGCCAGCAGTTGGGACTTTAACTATGCTGAGCAGTTCTTC | 1 | TRBV13-3*00 | TRBD2*00 | TRBJ2-1*00 | 0.24% |
| Treg DKO | TGTGCCAGCAGTTTAAACATTGCAAACTCCGACTACACCTTC | 1 | TRBV15*00 | | TRBJ1-2*00 | 0.24% |
| Treg DKO | TGTGCCAGCAGTTTACGCAATTATGAACAGTACTTC | 1 | TRBV15*00 | | TRBJ2-7*00 | 0.24% |
| Treg DKO | TGTGCCAGCAGTTTAGGGGCCTATGAACAGTACTTC | 1 | TRBV15*00 | | TRBJ2-7*00 | 0.24% |
| Treg DKO | TGTGCCAGCAGTTTAGGGGTGGGTGCAGAAACGCTGTATTTT | 1 | TRBV15*00 | | TRBJ2-3*00 | 0.24% |
| Treg DKO | TGTGCCAGCAGTTTCAGGGACCAAGACACCCAGTACTTT | 1 | TRBV14*00 | | TRBJ2-5*00 | 0.24% |
| Treg DKO | TGTGCCAGCAGTTTCTGGGAGCGCTATGAACAGTACTTC | 1 | TRBV15*00 | TRBD2*00 | TRBJ2-7*00 | 0.24% |
| Treg DKO | TGTGCCAGCAGTTTGGGATTTAACTATGCTGAGCAGTTCTTC | 1 | TRBV15*00 | | TRBJ2-1*00 | 0.24% |
| Treg DKO | TGTGCCAGCAGTTTTGCCAGGGAGGCTGAGCAGTTCTTC | 1 | TRBV15*00 | TRBD1*00 | TRBJ2-1*00 | 0.24% |
| Treg DKO | TGTGCCAGCGGCCCCAGGGGAGAAGTCTTCTTT | 1 | TRBV13-2*00 | TRBD1*00 | TRBJ1-1*00 | 0.24% |
| Treg DKO | TGTGCCAGCGGGGGGACTGGGGGGGCGCCTGCTGAGCAGTTCTTC | 1 | TRBV13-2*00 | TRBD2*00 | TRBJ2-1*00 | 0.24% |
| Treg DKO | TGTGCCAGCGGTCCAGGGGGGGGCCACCAAGACACCCAGTACTTT | 1 | TRBV13-2*00 | TRBD2*00 | TRBJ2-5*00 | 0.24% |
| Treg DKO | TGTGCCAGCGGTCTGGGGGAAGACACCCAGTACTTT | 1 | TRBV13-2*00 | TRBD2*00 | TRBJ2-5*00 | 0.24% |
| Treg DKO | TGTGCCAGCGGTGACCCAGACATCTATGAACAGTACTTC | 1 | TRBV13-2*00 | TRBD2*00 | TRBJ2-7*00 | 0.24% |
| Treg DKO | TGTGCCAGCGGTGACGGGGACTATGCTGAGCAGTTCTTC | 1 | TRBV13-2*00 | | TRBJ2-1*00 | 0.24% |
| Treg DKO | TGTGCCAGCGGTGACTACAGCAACGAAAGATTATTTTTC | 1 | TRBV13-2*00 | | TRBJ1-4*00 | 0.24% |
| Treg DKO | TGTGCCAGCGGTGAGGACTGGGTCTACTTC | 1 | TRBV13-2*00 | TRBD2*00 | TRBJ2-7*00 | 0.24% |

| | | | | | | |
|---|---|---|---|---|---|---|
| Treg DKO | TGTGCCAGCGGTGAGGGGACTGGGGGTGCAGAAACGCTGTATTTT | 1 | TRBV13-2*00 | TRBD2*00 | TRBJ2-3*00 | 0.24% |
| Treg DKO | TGTGCCAGCGGTGAGGGGGGTCAAAACACCTTGTACTTT | 1 | TRBV13-2*00 | TRBD1*00 | TRBJ2-4*00 | 0.24% |
| Treg DKO | TGTGCCAGCGGTGATACGGACAGGGGGACCGACTACACCTTC | 1 | TRBV13-2*00 | TRBD1*00 | TRBJ1-2*00 | 0.24% |
| Treg DKO | TGTGCCAGCGGTGATAGTCCGGCAAACACAGAAGTCTTCTTT | 1 | TRBV13-2*00 | TRBD2*00 | TRBJ1-1*00 | 0.24% |
| Treg DKO | TGTGCCAGCGGTGATCCGGAAGTCTTCTTT | 1 | TRBV13-2*00 | | TRBJ1-1*00 | 0.24% |
| Treg DKO | TGTGCCAGCGGTGATGCGGGGGCGCAGGCTCCGCTTTTT | 1 | TRBV13-2*00 | TRBD1*00 | TRBJ1-5*00 | 0.24% |
| Treg DKO | TGTGCCAGCGGTGATGCGGGGGTAAAAACCTTGTACTTT | 1 | TRBV13-2*00 | TRBD1*00 | TRBJ2-4*00 | 0.24% |
| Treg DKO | TGTGCCAGCGGTGATGCTGGGGGGGAAGACACCCAGTACTTT | 1 | TRBV13-2*00 | TRBD2*00 | TRBJ2-5*00 | 0.24% |
| Treg DKO | TGTGCCAGCGGTGATGCTGGTTATGAACAGTACTTC | 1 | TRBV13-2*00 | | TRBJ2-7*00 | 0.24% |
| Treg DKO | TGTGCCAGCGGTGATGCTTGGTATGAACAGTACTTC | 1 | TRBV13-2*00 | | TRBJ2-7*00 | 0.24% |
| Treg DKO | TGTGCCAGCGGTGATGGGGGGCGTAGGGACACCCAGTACTTT | 1 | TRBV13-2*00 | TRBD1*00 | TRBJ2-5*00 | 0.24% |
| Treg DKO | TGTGCCAGCGGTGCAGAAACGCTGTATTTT | 1 | TRBV13-2*00 | | TRBJ2-3*00 | 0.24% |
| Treg DKO | TGTGCCAGCGGTTCACAAAACTCCGACTACACCTTC | 1 | TRBV13-2*00 | | TRBJ1-2*00 | 0.24% |
| Treg DKO | TGTGCCAGCTCCCCGACAGGGAACGAAAGATTATTTTTC | 1 | TRBV18*00 | TRBD1*00 | TRBJ1-4*00 | 0.24% |
| Treg DKO | TGTGCCAGCTCCCCTTTCCCTAGTCAAAACACCTTGTACTTT | 1 | TRBV12-2*00 | | TRBJ2-4*00 | 0.24% |
| Treg DKO | TGTGCCAGCTCCGGGACAAAAAACACCTTGTACTTT | 1 | TRBV12-1*00 | TRBD1*00 | TRBJ2-4*00 | 0.24% |
| Treg DKO | TGTGCCAGCTCTCCCGACAGGGGTAATTCGCCCCTCTACTTT | 1 | TRBV12-1*00 | TRBD1*00 | TRBJ1-6*00 | 0.24% |
| Treg DKO | TGTGCCAGCTCTCCCGGGACAGGGAATTCTGGAAATACGCTCTATTTT | 1 | TRBV12-1*00 | TRBD1*00 | TRBJ1-3*00 | 0.24% |
| Treg DKO | TGTGCCAGCTCTCCGACTCCCTCCTATGAACAGTACTTC | 1 | TRBV12-1*00 | | TRBJ2-7*00 | 0.24% |
| Treg DKO | TGTGCCAGCTCTCCGGGTAACTATGCTGAGCAGTTCTTC | 1 | TRBV12-1*00 | | TRBJ2-1*00 | 0.24% |
| Treg DKO | TGTGCCAGCTCTCTCAGGGGGAGGTCCGACTACACCTTC | 1 | TRBV12-1*00 | TRBD1*00 | TRBJ1-2*00 | 0.24% |
| Treg DKO | TGTGCCAGCTCTCTCGACTGGGGCGTTTATGAACAGTACTTC | 1 | TRBV12-1*00 | TRBD1*00 | TRBJ2-7*00 | 0.24% |
| Treg DKO | TGTGCCAGCTCTCTCGACTGGGGGGGCTCCTATGAACAGTACTTC | 1 | TRBV12-1*00 | TRBD2*00 | TRBJ2-7*00 | 0.24% |
| Treg DKO | TGTGCCAGCTCTCTCGAGAACTCCGACTACACCTTC | 1 | TRBV12-1*00 | | TRBJ1-2*00 | 0.24% |
| Treg DKO | TGTGCCAGCTCTCTCGATAACAACCAGGCTCCGCTTTTT | 1 | TRBV12-1*00 | | TRBJ1-5*00 | 0.24% |
| Treg DKO | TGTGCCAGCTCTCTCGCAGTCCGTGACACCCAGTACTTT | 1 | TRBV12-1*00 | TRBD2*00 | TRBJ2-5*00 | 0.24% |
| Treg DKO | TGTGCCAGCTCTCTCGGGGGACCCAACGAAAGATTATTTTTC | 1 | TRBV12-1*00 | TRBD1*00 | TRBJ1-4*00 | 0.24% |
| Treg DKO | TGTGCCAGCTCTCTCGTTTATGAACAGTACTTC | 1 | TRBV12-1*00 | | TRBJ2-7*00 | 0.24% |
| Treg DKO | TGTGCCAGCTCTCTGCAAGAAAACACCTTGTACTTT | 1 | TRBV12-1*00 | | TRBJ2-4*00 | 0.24% |
| Treg DKO | TGTGCCAGCTCTCTGGGGGAGGACACCTTGTACTTT | 1 | TRBV12-1*00 | TRBD1*00 | TRBJ2-4*00 | 0.24% |
| Treg DKO | TGTGCCAGCTCTGGGGGGCGGGGTGCAGAAACGCTGTATTTT | 1 | TRBV12-1*00 | TRBD2*00 | TRBJ2-3*00 | 0.24% |
| Treg DKO | TGTGCCTGGAGACAGGGACCTTCCTATAATTCGCCCCTCTACTTT | 1 | TRBV31*00 | TRBD1*00 | TRBJ1-6*00 | 0.24% |
| Treg DKO | TGTGCCTGGAGGCACATTACTAGTGCAGAAACGCTGTATTTT | 1 | TRBV31*00 | | TRBJ2-3*00 | 0.24% |
| Treg DKO | TGTGCCTGGAGTCCACAAGCAAACTCCGACTACACCTTC | 1 | TRBV31*00 | | TRBJ1-2*00 | 0.24% |
| Treg DKO | TGTGCCTGGAGTCCCGGACAGTATGCTGAGCAGTTCTTC | 1 | TRBV31*00 | TRBD1*00 | TRBJ2-1*00 | 0.24% |
| Treg DKO | TGTGCCTGGAGTCGACTGGGGGTAACTATGCTGAGCAGTTCTTC | 1 | TRBV31*00 | TRBD1*00 | TRBJ2-1*00 | 0.24% |
| Treg DKO | TGTGCCTGGAGTCGAGGGGCCCTCTATAATTCGCCCCTCTACTTT | 1 | TRBV31*00 | TRBD1*00 | TRBJ1-6*00 | 0.24% |
| Treg DKO | TGTGCCTGGAGTCGGGGGATCCACAGAAGTCTTCTTT | 1 | TRBV31*00 | | TRBJ1-1*00 | 0.24% |
| Treg DKO | TGTGCCTGGAGTCTAGGTGGGGGCGAACAGTACTTC | 1 | TRBV31*00 | TRBD1*00 | TRBJ2-7*00 | 0.24% |
| Treg DKO | TGTGCCTGGAGTCTGATGGGGGGTGGCACCGGGCAGCTCTACTTT | 1 | TRBV31*00 | TRBD2*00 | TRBJ2-2*00 | 0.24% |
| Treg DKO | TGTGCCTGGAGTCTGGGGACAACAAACTCCGACTACACCTTC | 1 | TRBV31*00 | TRBD1*00 | TRBJ1-2*00 | 0.24% |
| Treg DKO | TGTGCCTGGAGTCTGGGGGGAGACACCTTGTACTTT | 1 | TRBV31*00 | TRBD2*00 | TRBJ2-4*00 | 0.24% |
| Treg DKO | TGTGCCTGGGGGGACTGGGAAGACACCCAGTACTTT | 1 | TRBV31*00 | TRBD2*00 | TRBJ2-5*00 | 0.24% |
| Treg DKO | TGTGCTAGCAGACAGGGGGGTTCTGGAAATACGCTCTATTTT | 1 | TRBV29*00 | TRBD1*00 | TRBJ1-3*00 | 0.24% |
| Treg DKO | TGTGCTAGCAGTATGGCAAACACAGAAGTCTTCTTT | 1 | TRBV17*00 | | TRBJ1-1*00 | 0.24% |
| Treg DKO | TGTGCTAGCAGTCCCTCGGGGGTGGCTCCGCTTTTT | 1 | TRBV21*00 | TRBD1*00 | TRBJ1-5*00 | 0.24% |
| Treg DKO | TGTGCTAGCAGTCGACAAGCAAACACAGAAGTCTTCTTT | 1 | TRBV21*00 | | TRBJ1-1*00 | 0.24% |
| Treg DKO | TGTGCTAGCAGTCGACTGGGGGCCACCTTGTACTTT | 1 | TRBV17*00 | TRBD2*00 | TRBJ2-4*00 | 0.24% |
| Treg DKO | TGTGCTAGCAGTTTATACTCCCTCCACACCGGGCAGCTCTACTTT | 1 | TRBV29*00 | | TRBJ2-2*00 | 0.24% |
| Treg DKO | TGTGCTAGCAGTTTATCCCCTGGGCAGCTCTACTTT | 1 | TRBV29*00 | | TRBJ2-2*00 | 0.24% |
| Treg DKO | TGTGCTAGCAGTTTCCAGGGTGCAAACGCTGTATTTT | 1 | TRBV29*00 | TRBD1*00 | TRBJ2-3*00 | 0.24% |
| Treg DKO | TGTGGTGCTAATACAGGGAGCTCTAACTATGCTGAGCAGTTCTTC | 1 | TRBV20*00 | TRBD1*00 | TRBJ2-1*00 | 0.24% |
| Treg DKO | TGTGGTGCTAGAACAAGCTATGAACAGTACTTC | 1 | TRBV20*00 | | TRBJ2-7*00 | 0.24% |
| Treg DKO | TGTGGTGCTAGGGACTGGGCCTATGCTGAGCAGTTCTTC | 1 | TRBV20*00 | TRBD2*00 | TRBJ2-1*00 | 0.24% |
| Treg DKO | TGTGGTGCTAGTTCGACAGGGAACTCCGACTACACCTTC | 1 | TRBV20*00 | TRBD1*00 | TRBJ1-2*00 | 0.24% |
| Treg DKO | TGTGCCAGCAGTGATAACACAGTCTCCTATGAACAGTACTTC | 1 | TRBV13-1*00 | | TRBJ2-7*00 | 0.24% |
| Treg DKO | TGCACCTGCAGTGCAGGGGGGGGCTCCTATGAACAGTACTTC | 1 | TRBV1*00 | TRBD2*00 | TRBJ2-7*00 | 0.24% |
| Treg DKO | TGTGGTGCTAGGGACAGGTATGCTGAGCAGTTCTTC | 1 | TRBV20*00 | | TRBJ2-1*00 | 0.24% |
| Treg DKO | TGTGCCAGCAGTCTGTCAGGGTATGAACAGTACTTC | 1 | TRBV26*00 | | TRBJ2-7*00 | 0.24% |

#### Treg DKO*

| Sample Name | CDR3 Sequence | clone Count | best V Hit | best D Hit | best J Hit | Frequency |
|---|---|---|---|---|---|---|
| Treg DKO* | TGCTCCAGCGGGACACAAGACACCCAGTACTTT | 32 | TRBV23*00 | TRBD1*00 | TRBJ2-5*00 | 9.38% |
| Treg DKO* | TGTGCCAGCAGTGATGGTAATAACAACCAGGCTCCGCTTTTT | 29 | TRBV13-3*00 | | TRBJ1-5*00 | 8.50% |
| Treg DKO* | TGTGCCAGCGGTCCATCCTATGAACAGTACTTC | 28 | TRBV13-2*00 | | TRBJ2-7*00 | 8.21% |
| Treg DKO* | TGTGCCAGCTCTCTCGACAGTTCCTATAATTCGCCCCTCTACTTT | 14 | TRBV12-1*00 | | TRBJ1-6*00 | 4.11% |
| Treg DKO* | TGTGCCAGCAGCCAAGATAACCAGGGGGCGCTTGAACAGTACTTC | 11 | TRBV5*00 | TRBD1*00 | TRBJ2-7*00 | 3.23% |
| Treg DKO* | TGTGCCAGCTCTTACCGGGACACTGGAAATACGCTCTATTTT | 7 | TRBV12-2*00 | TRBD1*00 | TRBJ1-3*00 | 2.05% |
| Treg DKO* | TGTGCAAGCAACTTAGATTGGGCCAACGAAAGATTATTTTTC | 6 | TRBV16*00 | | TRBJ1-4*00 | 1.76% |
| Treg DKO* | TGTGCCAGCGGTCGGGACACCTATTCTGGAAATACGCTCTATTTT | 6 | TRBV13-2*00 | TRBD1*00 | TRBJ1-3*00 | 1.76% |
| Treg DKO* | TGCACCTGCAGTGCAGAGGGACTGGGGGGGCGCGAACAGTACTTC | 5 | TRBV1*00 | TRBD2*00 | TRBJ2-7*00 | 1.47% |
| Treg DKO* | TGTGCCAGCAGCTTAGGGGGGGGGCAAGACACCCAGTACTTT | 5 | TRBV3*00 | TRBD2*00 | TRBJ2-5*00 | 1.47% |
| Treg DKO* | TGCACCTGCAGTGCAGATCCTGGGTTTAACCAAGACACCCAGTACTTT | 4 | TRBV1*00 | TRBD2*00 | TRBJ2-5*00 | 1.17% |
| Treg DKO* | TGTGCCAGCAGTGGAGGGAAGAACACCTTGTACTTT | 4 | TRBV22*00 | | TRBJ2-4*00 | 1.17% |
| Treg DKO* | TGTGCCTGGAGTCTGAGGAACCAAGACACCCAGTACTTT | 4 | TRBV31*00 | | TRBJ2-5*00 | 1.17% |

| | | | | | | |
|---|---|---|---|---|---|---|
| Treg DKO* | TGTGCCAGCAGTCTGAATATAGGCTATGCTGAGCAGTTCTTC | 3 | TRBV26*00 | | TRBJ2-1*00 | 0.88% |
| Treg DKO* | TGCACCTGCAGTGTCGACAGGGCACAAGACACCCAGTACTTT | 2 | TRBV1*00 | TRBD1*00 | TRBJ2-5*00 | 0.59% |
| Treg DKO* | TGTGCAAGCAGCTTAGACAACGAAAGATTATTTTTC | 2 | TRBV16*00 | | TRBJ1-4*00 | 0.59% |
| Treg DKO* | TGTGCCAGCAGAGGGAATTCTGGAAATACGCTCTATTTT | 2 | TRBV19*00 | | TRBJ1-3*00 | 0.59% |
| Treg DKO* | TGTGCCAGCAGCCAAACAGGGCACTCCGACTACACCTTC | 2 | TRBV2*00 | TRBD1*00 | TRBJ1-2*00 | 0.59% |
| Treg DKO* | TGTGCCAGCAGCCAAGGACTCAGTCAAAACACCTTGTACTTT | 2 | TRBV5*00 | | TRBJ2-4*00 | 0.59% |
| Treg DKO* | TGTGCCAGCAGCCAGAGGGTGGGTGCAGAAACGCTGTATTTT | 2 | TRBV3*00 | | TRBJ2-3*00 | 0.59% |
| Treg DKO* | TGTGCCAGCAGCCCCCGACATCAAGACACCCAGTACTTT | 2 | TRBV5*00 | | TRBJ2-5*00 | 0.59% |
| Treg DKO* | TGTGCCAGCAGCTATTTGGGGTGGAACACCTTGTACTTT | 2 | TRBV4*00 | TRBD2*00 | TRBJ2-4*00 | 0.59% |
| Treg DKO* | TGTGCCAGCAGCTCGGGACAAAATTCGCCCCTCTACTTT | 2 | TRBV4*00 | TRBD1*00 | TRBJ1-6*00 | 0.59% |
| Treg DKO* | TGTGCCAGCAGCTTAGAAGACCCCGGGCAGCTCTACTTT | 2 | TRBV3*00 | | TRBJ2-2*00 | 0.59% |
| Treg DKO* | TGTGCCAGCAGTATAGACTGGCGAGAACAGTACTTC | 2 | TRBV19*00 | TRBD2*00 | TRBJ2-7*00 | 0.59% |
| Treg DKO* | TGTGCCAGCAGTCTGTCGACATTTAACCAAGACACCCAGTACTTT | 2 | TRBV26*00 | | TRBJ2-5*00 | 0.59% |
| Treg DKO* | TGTGCCAGCAGTGGACAAACTAGTGCAGAAACGCTGTATTTT | 2 | TRBV19*00 | TRBD1*00 | TRBJ2-3*00 | 0.59% |
| Treg DKO* | TGTGCCAGCAGTTTAGCAGCTGCTGAGCAGTTCTTC | 2 | TRBV15*00 | | TRBJ2-1*00 | 0.59% |
| Treg DKO* | TGTGCCAGCTCTCTCGACAACTATGCTGAGCAGTTCTTC | 2 | TRBV12-1*00 | | TRBJ2-1*00 | 0.59% |
| Treg DKO* | TGTGCCAGCTCTCTCGCAGAAACGCTGTATTTT | 2 | TRBV12-2*00 | | TRBJ2-3*00 | 0.59% |
| Treg DKO* | TGTGGTGCTAGGGAAACAGCAAACTCCGACTACACCTTC | 2 | TRBV20*00 | | TRBJ1-2*00 | 0.59% |
| Treg DKO* | TGTGCCAGCAGTCTAGAAAACACCGGGCAGCTCTACTTT | 2 | TRBV15*00 | | TRBJ2-2*00 | 0.59% |
| Treg DKO* | TGCACCTGCAGTCCCCAGGCAAACTCCGACTACACCTTC | 1 | TRBV1*00 | TRBD2*00 | TRBJ1-2*00 | 0.29% |
| Treg DKO* | TGCACCTGCAGTGCAGACTCTGCGCAGTTCTTC | 1 | TRBV1*00 | | TRBJ2-1*00 | 0.29% |
| Treg DKO* | TGCACCTGCAGTGCAGATCCTGGGTTTAACCAAGACACCCAGTACTTC | 1 | TRBV1*00 | TRBD2*00 | TRBJ2-5*00 | 0.29% |
| Treg DKO* | TGCACCTGCAGTGCAGATTCGCTCCGAGCTCCTATGAACAGTACTTC | 1 | TRBV1*00 | | TRBJ2-7*00 | 0.29% |
| Treg DKO* | TGCACCTGCAGTGCAGGGGATAATTCGCCCCTCTACTTT | 1 | TRBV1*00 | | TRBJ1-6*00 | 0.29% |
| Treg DKO* | TGCACCTGCAGTGCTGGGACAGGGAACGAAAGATTATTTTTC | 1 | TRBV1*00 | TRBD1*00 | TRBJ1-4*00 | 0.29% |
| Treg DKO* | TGCTCCAGCAGTCAATCGGGGGGGGCAGACACCCAGTACTTT | 1 | TRBV23*00 | TRBD2*00 | TRBJ2-5*00 | 0.29% |
| Treg DKO* | TGTGCAAGCAGCACTGGACATGCAAACACCGGGCAGCTCTACTTT | 1 | TRBV16*00 | TRBD1*00 | TRBJ2-2*00 | 0.29% |
| Treg DKO* | TGTGCAAGCAGCCCCGGGACAGGGGACACCTTGTACTTT | 1 | TRBV16*00 | TRBD1*00 | TRBJ2-4*00 | 0.29% |
| Treg DKO* | TGTGCAAGCAGCTACTGGGGGACTCCGACTACACCTTC | 1 | TRBV16*00 | TRBD2*00 | TRBJ1-2*00 | 0.29% |
| Treg DKO* | TGTGCAAGCAGCTCCCAGGGGGACTCCGACTACACCTTC | 1 | TRBV16*00 | TRBD1*00 | TRBJ1-2*00 | 0.29% |
| Treg DKO* | TGTGCAAGCAGCTCCGACAGGCAGGCTCCGCTTTTT | 1 | TRBV16*00 | TRBD1*00 | TRBJ1-5*00 | 0.29% |
| Treg DKO* | TGTGCAAGCAGCTCCTTTAACCAAGACACCCAGTACTTT | 1 | TRBV16*00 | | TRBJ2-5*00 | 0.29% |
| Treg DKO* | TGTGCAAGCAGCTTACAGGGCGAAAGATTATTTTTC | 1 | TRBV16*00 | TRBD1*00 | TRBJ1-4*00 | 0.29% |
| Treg DKO* | TGTGCAAGCAGCTTAGATCCCTATAACTATGCTGAGCAGTTCTTC | 1 | TRBV16*00 | | TRBJ2-1*00 | 0.29% |
| Treg DKO* | TGTGCAAGCAGCTTAGGACAGGGGAACTATGCTGAGCAGTTCTTC | 1 | TRBV16*00 | TRBD1*00 | TRBJ2-1*00 | 0.29% |
| Treg DKO* | TGTGCAAGCAGCTTAGTCACAAATACGCTCTATTTT | 1 | TRBV16*00 | | TRBJ1-3*00 | 0.29% |
| Treg DKO* | TGTGCAAGCAGCTTGGGAGGGCTCCAAGACACCCAGTACTTT | 1 | TRBV16*00 | | TRBJ2-5*00 | 0.29% |
| Treg DKO* | TGTGCAAGCAGCTTGGGGGGCACAAACACAGAAGTCTTCTTT | 1 | TRBV16*00 | TRBD2*00 | TRBJ1-1*00 | 0.29% |
| Treg DKO* | TGTGCCAGCACCCCGGGGGGAGGTGCAGAAACGCTGTATTTT | 1 | TRBV15*00 | TRBD1*00 | TRBJ2-3*00 | 0.29% |
| Treg DKO* | TGTGCCAGCACCCTGGGGGTCTATGCTGAGCAGTTCTTC | 1 | TRBV13-3*00 | TRBD2*00 | TRBJ2-1*00 | 0.29% |
| Treg DKO* | TGTGCCAGCACTCCTTCCTATAATTCGCCCCTCTACTTT | 1 | TRBV13-3*00 | | TRBJ1-6*00 | 0.29% |
| Treg DKO* | TGTGCCAGCAGAACGACAGGGGTGGCTGAGCAGTTCTTC | 1 | TRBV19*00 | TRBD1*00 | TRBJ2-1*00 | 0.29% |
| Treg DKO* | TGTGCCAGCAGCAACAGGGGCGAAACGCTGTATTTT | 1 | TRBV19*00 | TRBD1*00 | TRBJ2-3*00 | 0.29% |
| Treg DKO* | TGTGCCAGCAGCAAGGGGGGGGACACCCAGTACTTT | 1 | TRBV3*00 | TRBD2*00 | TRBJ2-5*00 | 0.29% |
| Treg DKO* | TGTGCCAGCAGCACCCAGTACTTT | 1 | TRBV19*00 | | TRBJ2-5*00 | 0.29% |
| Treg DKO* | TGTGCCAGCAGCCAAGAACAGAACTCCGACTACACCTTC | 1 | TRBV5*00 | | TRBJ1-2*00 | 0.29% |
| Treg DKO* | TGTGCCAGCAGCCAAGAACGGGGTTCCTATGAACAGTACTTC | 1 | TRBV2*00 | | TRBJ2-7*00 | 0.29% |
| Treg DKO* | TGTGCCAGCAGCCAAGAGGGCAATCAAAACACCTTGTACTTT | 1 | TRBV2*00 | | TRBJ2-4*00 | 0.29% |
| Treg DKO* | TGTGCCAGCAGCCAAGAGGGGGAGGGAGACACCCAGTACTTT | 1 | TRBV5*00 | TRBD1*00 | TRBJ2-5*00 | 0.29% |
| Treg DKO* | TGTGCCAGCAGCCAAGAGGGTAGTGCAGAAACGCTGTATTTT | 1 | TRBV5*00 | | TRBJ2-3*00 | 0.29% |
| Treg DKO* | TGTGCCAGCAGCCAAGATCTTCCAGACACCCAGTACTTT | 1 | TRBV2*00 | | TRBJ2-5*00 | 0.29% |
| Treg DKO* | TGTGCCAGCAGCCAAGCGGGACAGAAAAACACCTTGTACTTT | 1 | TRBV2*00 | TRBD1*00 | TRBJ2-4*00 | 0.29% |
| Treg DKO* | TGTGCCAGCAGCCAAGCTGGGACAACAGAAGTCTTCTTT | 1 | TRBV2*00 | TRBD1*00 | TRBJ1-1*00 | 0.29% |
| Treg DKO* | TGTGCCAGCAGCCAAGGGGACAACAACGAAAGATTATTTTTC | 1 | TRBV2*00 | TRBD1*00 | TRBJ1-4*00 | 0.29% |
| Treg DKO* | TGTGCCAGCAGCCAAGGGGACAACTATGCTGAGCAGTTCTTC | 1 | TRBV2*00 | TRBD1*00 | TRBJ2-1*00 | 0.29% |
| Treg DKO* | TGTGCCAGCAGCCAAGTCTCCTATGAACAGTACTTC | 1 | TRBV5*00 | | TRBJ2-7*00 | 0.29% |
| Treg DKO* | TGTGCCAGCAGCCAGACAGGGGTTGCTGAGCAGTTCTTC | 1 | TRBV5*00 | TRBD1*00 | TRBJ2-1*00 | 0.29% |
| Treg DKO* | TGTGCCAGCAGCCCAGGGGCAAACACAGAAGTCTTCTTT | 1 | TRBV4*00 | TRBD1*00 | TRBJ1-1*00 | 0.29% |
| Treg DKO* | TGTGCCAGCAGCCCCCGACTGGTTTATGAACAGTACTTC | 1 | TRBV13-1*00 | TRBD2*00 | TRBJ2-7*00 | 0.29% |
| Treg DKO* | TGTGCCAGCAGCCGCCGGGACAATAACCAGGCTCCGCTTTTT | 1 | TRBV2*00 | TRBD1*00 | TRBJ1-5*00 | 0.29% |
| Treg DKO* | TGTGCCAGCAGCCGGACAGCAAACTCCGACTACACCTTC | 1 | TRBV3*00 | TRBD1*00 | TRBJ1-2*00 | 0.29% |
| Treg DKO* | TGTGCCAGCAGCCTACAATTAACCAAGACACCCAGTACTTT | 1 | TRBV24*00 | | TRBJ2-5*00 | 0.29% |
| Treg DKO* | TGTGCCAGCAGCCTCCGGGACACAGAAGTCTTCTTT | 1 | TRBV2*00 | TRBD1*00 | TRBJ1-1*00 | 0.29% |
| Treg DKO* | TGTGCCAGCAGCGACAGGGGGCCGCTGTATTTT | 1 | TRBV4*00 | TRBD1*00 | TRBJ2-3*00 | 0.29% |
| Treg DKO* | TGTGCCAGCAGCTATGGGGGGGACTATGAACAGTACTTC | 1 | TRBV4*00 | TRBD2*00 | TRBJ2-7*00 | 0.29% |
| Treg DKO* | TGTGCCAGCAGCTCAGGGGTTGCTGAGCAGTTCTTC | 1 | TRBV25*00 | TRBD1*00 | TRBJ2-1*00 | 0.29% |
| Treg DKO* | TGTGCCAGCAGCTCGAAATATGAACAGTACTTC | 1 | TRBV3*00 | | TRBJ2-7*00 | 0.29% |
| Treg DKO* | TGTGCCAGCAGCTCGACACTTGCAAACACCGGGCAGCTCTACTTT | 1 | TRBV25*00 | | TRBJ2-2*00 | 0.29% |
| Treg DKO* | TGTGCCAGCAGCTCGGACTCCTACACCGGGCAGCTCTACTTT | 1 | TRBV25*00 | TRBD2*00 | TRBJ2-2*00 | 0.29% |
| Treg DKO* | TGTGCCAGCAGCTCGGGACAAGCAAACACAGAAGTCTTCTTT | 1 | TRBV25*00 | TRBD1*00 | TRBJ1-1*00 | 0.29% |
| Treg DKO* | TGTGCCAGCAGCTCGGGACAGGGCCCCGGGCAGCTCTACTTT | 1 | TRBV25*00 | TRBD1*00 | TRBJ2-2*00 | 0.29% |
| Treg DKO* | TGTGCCAGCAGCTTAACTGGACAATATGAACAGTACTTC | 1 | TRBV3*00 | TRBD1*00 | TRBJ2-7*00 | 0.29% |
| Treg DKO* | TGTGCCAGCAGCTTAAGCAGAAAAGAAGTCTTCTTT | 1 | TRBV25*00 | | TRBJ1-1*00 | 0.29% |
| Treg DKO* | TGTGCCAGCAGCTTAGAAGTCTTCTTT | 1 | TRBV3*00 | | TRBJ1-1*00 | 0.29% |
| Treg DKO* | TGTGCCAGCAGCTTAGAGGGTTCCGGGCAGCTCTACTTT | 1 | TRBV3*00 | | TRBJ2-2*00 | 0.29% |

| | | | | | | |
|---|---|---|---|---|---|---|
| Treg DKO* | TGTGCCAGCAGCTTAGCCCACACAGAAGTCTTCTTT | 1 | TRBV3*00 | | TRBJ1-1*00 | 0.29% |
| Treg DKO* | TGTGCCAGCAGCTTAGCCGGGACAATAAACACCTTGTACTTT | 1 | TRBV3*00 | TRBD1*00 | TRBJ2-4*00 | 0.29% |
| Treg DKO* | TGTGCCAGCAGCTTAGGGGCAAACACAGAAGTCTTCTTT | 1 | TRBV3*00 | | TRBJ1-1*00 | 0.29% |
| Treg DKO* | TGTGCCAGCAGCTTCACAGGGGGGATGGACACCCAGTACTTT | 1 | TRBV3*00 | TRBD1*00 | TRBJ2-5*00 | 0.29% |
| Treg DKO* | TGTGCCAGCAGCTTCGGGGAGGGCACAGAAGTCTTCTTT | 1 | TRBV3*00 | | TRBJ1-1*00 | 0.29% |
| Treg DKO* | TGTGCCAGCAGCTTGGGGACAGGGGGTTATGAACAGTACTTC | 1 | TRBV3*00 | TRBD1*00 | TRBJ2-7*00 | 0.29% |
| Treg DKO* | TGTGCCAGCAGCTTTCTGGACCAAGACACCCAGTACTTT | 1 | TRBV3*00 | | TRBJ2-5*00 | 0.29% |
| Treg DKO* | TGTGCCAGCAGGATAGTTAACCAAGACACCCAGTACTTT | 1 | TRBV19*00 | | TRBJ2-5*00 | 0.29% |
| Treg DKO* | TGTGCCAGCAGGGACATAAACTATGCTGAGCAGTTCTTC | 1 | TRBV13-3*00 | TRBD1*00 | TRBJ2-1*00 | 0.29% |
| Treg DKO* | TGTGCCAGCAGTAAAGAAAATTCGCCCCTCTACTTT | 1 | TRBV19*00 | | TRBJ1-6*00 | 0.29% |
| Treg DKO* | TGTGCCAGCAGTAAGGGGACAGACACAGAAGTCTTCTTT | 1 | TRBV19*00 | TRBD1*00 | TRBJ1-1*00 | 0.29% |
| Treg DKO* | TGTGCCAGCAGTATACTGGGGGGGGACACCCAGTACTTT | 1 | TRBV4*00 | TRBD2*00 | TRBJ2-5*00 | 0.29% |
| Treg DKO* | TGTGCCAGCAGTATATTCGGTAACTATGAACAGTACTTC | 1 | TRBV19*00 | | TRBJ2-7*00 | 0.29% |
| Treg DKO* | TGTGCCAGCAGTATGAGCACAGAAGTCTTCTTT | 1 | TRBV19*00 | | TRBJ1-1*00 | 0.29% |
| Treg DKO* | TGTGCCAGCAGTATTCCAAGAAACACCGGGCAGCTCTACTTT | 1 | TRBV19*00 | | TRBJ2-2*00 | 0.29% |
| Treg DKO* | TGTGCCAGCAGTCCCAATGCAGAAACGCTGTATTTT | 1 | TRBV15*00 | | TRBJ2-3*00 | 0.29% |
| Treg DKO* | TGTGCCAGCAGTCCGGACAGGGATGCTGAGCAGTTCTTC | 1 | TRBV26*00 | TRBD1*00 | TRBJ2-1*00 | 0.29% |
| Treg DKO* | TGTGCCAGCAGTCCGGACAGTGCAGAAACGCTGTATTTT | 1 | TRBV26*00 | | TRBJ2-3*00 | 0.29% |
| Treg DKO* | TGTGCCAGCAGTCCTCAGGGAGAAAGATTATTTTTC | 1 | TRBV15*00 | TRBD1*00 | TRBJ1-4*00 | 0.29% |
| Treg DKO* | TGTGCCAGCAGTCGACAGGGAAATACGCTCTATTTT | 1 | TRBV15*00 | TRBD1*00 | TRBJ1-3*00 | 0.29% |
| Treg DKO* | TGTGCCAGCAGTCGGGACAGCTATGCTGAGCAGTTCTTC | 1 | TRBV26*00 | TRBD1*00 | TRBJ2-1*00 | 0.29% |
| Treg DKO* | TGTGCCAGCAGTCTGCAGGGGGCGCTTGCTGAGCAGTTCTTC | 1 | TRBV26*00 | TRBD1*00 | TRBJ2-1*00 | 0.29% |
| Treg DKO* | TGTGCCAGCAGTCTGCCAGAAGTCTTCTTT | 1 | TRBV26*00 | | TRBJ1-1*00 | 0.29% |
| Treg DKO* | TGTGCCAGCAGTCTGTTCCATTCTGGAAATACGCTCTATTTT | 1 | TRBV26*00 | | TRBJ1-3*00 | 0.29% |
| Treg DKO* | TGTGCCAGCAGTGACGGCTATGAACAGTACTTC | 1 | TRBV13-3*00 | | TRBJ2-7*00 | 0.29% |
| Treg DKO* | TGTGCCAGCAGTGAGGGGAACTCCGACTACACCTTC | 1 | TRBV13-3*00 | | TRBJ1-2*00 | 0.29% |
| Treg DKO* | TGTGCCAGCAGTGATCCCGGGACAAGTCAAAACACCTTGTACTTT | 1 | TRBV13-1*00 | TRBD1*00 | TRBJ2-4*00 | 0.29% |
| Treg DKO* | TGTGCCAGCAGTGATGACAGGAATTCTGGAAATACGCTCTATTTT | 1 | TRBV13-3*00 | | TRBJ1-3*00 | 0.29% |
| Treg DKO* | TGTGCCAGCAGTGATGCACAGTCCAATCAAGTCTTCTTT | 1 | TRBV13-1*00 | TRBD2*00 | TRBJ1-1*00 | 0.29% |
| Treg DKO* | TGTGCCAGCAGTGATGCAGATACGCTCTATTTT | 1 | TRBV13-1*00 | | TRBJ1-3*00 | 0.29% |
| Treg DKO* | TGTGCCAGCAGTGATGCGGGACAACAAGACACCCAGTACTTT | 1 | TRBV13-3*00 | TRBD1*00 | TRBJ2-5*00 | 0.29% |
| Treg DKO* | TGTGCCAGCAGTGATTACAACCCAAACACAGAAGTCTTCTTT | 1 | TRBV13-1*00 | | TRBJ1-1*00 | 0.29% |
| Treg DKO* | TGTGCCAGCAGTGATTGGGGGGGCGCTGACACCCAGTACTTT | 1 | TRBV13-3*00 | TRBD2*00 | TRBJ2-5*00 | 0.29% |
| Treg DKO* | TGTGCCAGCAGTGCCGACAGGGGTGAACAGTACTTC | 1 | TRBV13-1*00 | TRBD1*00 | TRBJ2-7*00 | 0.29% |
| Treg DKO* | TGTGCCAGCAGTGGACAGAAATATAATTCGCCCCTCTACTTT | 1 | TRBV13-3*00 | TRBD1*00 | TRBJ1-6*00 | 0.29% |
| Treg DKO* | TGTGCCAGCAGTTCAGGGAACTCCGACTACACCTTC | 1 | TRBV15*00 | TRBD1*00 | TRBJ1-2*00 | 0.29% |
| Treg DKO* | TGTGCCAGCAGTTGGACAGGCGGGGACACCCAGTACTTT | 1 | TRBV22*00 | TRBD1*00 | TRBJ2-5*00 | 0.29% |
| Treg DKO* | TGTGCCAGCAGTTTAAGCCGGACAAACACAGAAGTCTTCTTT | 1 | TRBV15*00 | | TRBJ1-1*00 | 0.29% |
| Treg DKO* | TGTGCCAGCAGTTTAAGGGACACCGGGCAGCTCTACTTT | 1 | TRBV15*00 | | TRBJ2-2*00 | 0.29% |
| Treg DKO* | TGTGCCAGCAGTTTAGAAGGGGGAGGCTATGCTGAGCAGTTCTTC | 1 | TRBV15*00 | TRBD1*00 | TRBJ2-1*00 | 0.29% |
| Treg DKO* | TGTGCCAGCAGTTTAGGGGCAAGCTCCTATGAACAGTACTTC | 1 | TRBV15*00 | | TRBJ2-7*00 | 0.29% |
| Treg DKO* | TGTGCCAGCAGTTTCGGGGGGGCGGCCTATGAACAGTACTTC | 1 | TRBV14*00 | TRBD2*00 | TRBJ2-7*00 | 0.29% |
| Treg DKO* | TGTGCCAGCAGTTTGTGGGATAACCAAGACACCCAGTACTTT | 1 | TRBV15*00 | | TRBJ2-5*00 | 0.29% |
| Treg DKO* | TGTGCCAGCAGTTTTACAGAGTATGCTGAGCAGTTCTTC | 1 | TRBV15*00 | | TRBJ2-1*00 | 0.29% |
| Treg DKO* | TGTGCCAGCGGAACTGGGGGGGCGCGAGACACCCAGTACTTT | 1 | TRBV13-2*00 | TRBD2*00 | TRBJ2-5*00 | 0.29% |
| Treg DKO* | TGTGCCAGCGGCGAGGCCTTGTACTTT | 1 | TRBV13-2*00 | | TRBJ2-4*00 | 0.29% |
| Treg DKO* | TGTGCCAGCGGGCGGGGGGGCGGCTCCGACTACACCTTC | 1 | TRBV13-2*00 | TRBD2*00 | TRBJ1-2*00 | 0.29% |
| Treg DKO* | TGTGCCAGCGGGGGACAGACCCAGTACTTT | 1 | TRBV13-2*00 | TRBD1*00 | TRBJ2-5*00 | 0.29% |
| Treg DKO* | TGTGCCAGCGGTCGGGACACCTATTCTGGAAATACGCTCCATTTT | 1 | TRBV13-2*00 | TRBD1*00 | TRBJ1-3*00 | 0.29% |
| Treg DKO* | TGTGCCAGCGGTGAACCGGGACTGGGGAGGAGCTCCTATGAACAGTACTTC | 1 | TRBV13-2*00 | TRBD2*00 | TRBJ2-7*00 | 0.29% |
| Treg DKO* | TGTGCCAGCGGTGATCGCTGGGGGAACCAAGACACCCAGTACTTT | 1 | TRBV13-2*00 | TRBD2*00 | TRBJ2-5*00 | 0.29% |
| Treg DKO* | TGTGCCAGCGGTGATGCCTATAATTCGCCCCTCTACTTT | 1 | TRBV13-2*00 | | TRBJ1-6*00 | 0.29% |
| Treg DKO* | TGTGCCAGCGGTGATGCTGCGGGGGAACAGTACTTC | 1 | TRBV13-2*00 | | TRBJ2-7*00 | 0.29% |
| Treg DKO* | TGTGCCAGCGGTGATGGGGGGCGCGCAGAAACGCTGTATTTT | 1 | TRBV13-2*00 | TRBD1*00 | TRBJ2-3*00 | 0.29% |
| Treg DKO* | TGTGCCAGCGGTGATTTTAGCTCCTATGAACAGTACTTC | 1 | TRBV13-2*00 | | TRBJ2-7*00 | 0.29% |
| Treg DKO* | TGTGCCAGCGGTGCCCCGACAGGGGCAAACTCCGACTACACCTTC | 1 | TRBV13-2*00 | TRBD1*00 | TRBJ1-2*00 | 0.29% |
| Treg DKO* | TGTGCCAGCGGTGCGGACGTTAACCAAGACACCCAGTACTTT | 1 | TRBV13-2*00 | | TRBJ2-5*00 | 0.29% |
| Treg DKO* | TGTGCCAGCGGTGGGACAACCAACGAAAGATTATTTTTC | 1 | TRBV13-2*00 | TRBD1*00 | TRBJ1-4*00 | 0.29% |
| Treg DKO* | TGTGCCAGCGGTGGGGACAGCTATGCTGAGCAGTTCTTC | 1 | TRBV13-2*00 | TRBD1*00 | TRBJ2-1*00 | 0.29% |
| Treg DKO* | TGTGCCAGCGGTTCATCCTATGAACAGTACTTC | 1 | TRBV13-2*00 | | TRBJ2-7*00 | 0.29% |
| Treg DKO* | TGTGCCAGCTCCCCTGGAAATACGCTCTATTTT | 1 | TRBV12-1*00 | | TRBJ1-3*00 | 0.29% |
| Treg DKO* | TGTGCCAGCTCCCGCGTGGGGAGGACGCTGTATTTT | 1 | TRBV12-1*00 | TRBD2*00 | TRBJ2-3*00 | 0.29% |
| Treg DKO* | TGTGCCAGCTCCCGGACAGTTTCTGGAAATACGCTCTATTTT | 1 | TRBV12-1*00 | TRBD1*00 | TRBJ1-3*00 | 0.29% |
| Treg DKO* | TGTGCCAGCTCCCGTTATGCTGAGCAGTTCTTC | 1 | TRBV12-2*00 | | TRBJ2-1*00 | 0.29% |
| Treg DKO* | TGTGCCAGCTCCTCGGGACAATATAACTATGCTGAGCAGTTCTTC | 1 | TRBV12-1*00 | TRBD1*00 | TRBJ2-1*00 | 0.29% |
| Treg DKO* | TGTGCCAGCTCTCCGCAAAACCAGGCTCCGCTTTTT | 1 | TRBV12-1*00 | | TRBJ1-5*00 | 0.29% |
| Treg DKO* | TGTGCCAGCTCTCTCCGGACTGGGGGGAGAGCTGAGCAGTTCTTC | 1 | TRBV12-1*00 | TRBD2*00 | TRBJ2-1*00 | 0.29% |
| Treg DKO* | TGTGCCAGCTCTCTCGACATTAACCAAGACACCCAGTACTTT | 1 | TRBV12-1*00 | | TRBJ2-5*00 | 0.29% |
| Treg DKO* | TGTGCCAGCTCTCTCGACTGGGGGGGATATGAACAGTACTTC | 1 | TRBV12-1*00 | TRBD2*00 | TRBJ2-7*00 | 0.29% |
| Treg DKO* | TGTGCCAGCTCTCTGGGGACAGGGGCTCCGACTACACCTTC | 1 | TRBV12-1*00 | TRBD1*00 | TRBJ1-2*00 | 0.29% |
| Treg DKO* | TGTGCCATGACTGGGGGGAGTGCAGAAACGCTGTATTTT | 1 | TRBV13-1*00 | TRBD2*00 | TRBJ2-3*00 | 0.29% |
| Treg DKO* | TGTGCCTGGAGTAGGCAGGGGGCCAGTGCAGAAACGCTGTATTTT | 1 | TRBV31*00 | TRBD1*00 | TRBJ2-3*00 | 0.29% |
| Treg DKO* | TGTGCCTGGAGTCTCCAGGGGGCAAACTCCGACTACACCTTC | 1 | TRBV31*00 | TRBD1*00 | TRBJ1-2*00 | 0.29% |
| Treg DKO* | TGTGCCTGGAGTCTGAACACAGAAGTCTTCTTT | 1 | TRBV31*00 | | TRBJ1-1*00 | 0.29% |
| Treg DKO* | TGTGCCTGGAGTTCTGACAACTCCGACTACACCTTC | 1 | TRBV31*00 | | TRBJ1-2*00 | 0.29% |

| | | | | | | |
|---|---|---|---|---|---|---|
| Treg DKO* | TGTGCTAGCAGTTTAAGGGCAGGGACCACCCAGTACTTT | 1 | TRBV29*00 | TRBD1*00 | TRBJ2-5*00 | 0.29% |
| Treg DKO* | TGTGCTAGCAGTTTATCGGGGGCTCCGCTTTTT | 1 | TRBV29*00 | | TRBJ1-5*00 | 0.29% |
| Treg DKO* | TGTGCTAGCAGTTTATTCAGGGGGCACACCGGGCAGCTCTACTTT | 1 | TRBV29*00 | TRBD1*00 | TRBJ2-2*00 | 0.29% |
| Treg DKO* | TGTGCTAGCAGTTTCGCGAAAGACACCCAGTACTTT | 1 | TRBV29*00 | | TRBJ2-5*00 | 0.29% |
| Treg DKO* | TGTGCTAGCTCTCTCGACAACTATGCTGAGCAGTTCTTC | 1 | TRBV12-1*00 | | TRBJ2-1*00 | 0.29% |
| Treg DKO* | TGTGGTGCTAGGACCTATGCTGAGCAGTTCTTC | 1 | TRBV20*00 | | TRBJ2-1*00 | 0.29% |
| Treg DKO* | TGTGGTGCTAGGGAACAAAATTCCTATGAACAGTACTTC | 1 | TRBV20*00 | | TRBJ2-7*00 | 0.29% |
| Treg DKO* | TGTGGTGCTAGGGAAGGGGCCAACACAGAAGTCTTCTTT | 1 | TRBV20*00 | TRBD1*00 | TRBJ1-1*00 | 0.29% |
| Treg DKO* | TGTGGTGCTAGGGACAAAAACTCCGACTACACCTTC | 1 | TRBV20*00 | | TRBJ1-2*00 | 0.29% |
| Treg DKO* | TGTGGTGCTAGGGACAATTCTGGAAATACGCTCTATTTT | 1 | TRBV20*00 | | TRBJ1-3*00 | 0.29% |
| Treg DKO* | TGTGGTGCTAGGGATCAGGGGACATCAAACTCCGACTACACCTTC | 1 | TRBV20*00 | TRBD1*00 | TRBJ1-2*00 | 0.29% |
| Treg DKO* | TGTGGTGCTAGGGGAGGGGCTGGATCCGACTACACCTTC | 1 | TRBV20*00 | TRBD1*00 | TRBJ1-2*00 | 0.29% |
| Treg DKO* | TGTGGTGCTGGGACAGGAAACACCGGGCAGCTCTACTTT | 1 | TRBV20*00 | TRBD1*00 | TRBJ2-2*00 | 0.29% |
| Treg DKO* | TGTGGTGCTGGGTCCGGGACAGACCAAGACACCCAGTACTTT | 1 | TRBV20*00 | TRBD1*00 | TRBJ2-5*00 | 0.29% |
| Treg DKO* | TGTGGTGCTTATTCATATAACCAAGACACCCAGTACTTT | 1 | TRBV20*00 | | TRBJ2-5*00 | 0.29% |
| Treg DKO* | TGTGCCAGCGGTGATGGGGACACCCAGTACTTT | 1 | TRBV13-2*00 | | TRBJ2-5*00 | 0.29% |

#### CD4+YFP- WT1

| Sample Name | CDR3 Sequence | clone Count | best V Hit | best D Hit | best J Hit | Frequency |
|---|---|---|---|---|---|---|
| CD4+YFP- WT1 | TGTGCCAGCAGTCTGTCTTCATATAACTATGCTGAGCAGTTCTTC | 5 | TRBV26*00 | | TRBJ2-1*00 | 1.43% |
| CD4+YFP- WT1 | TGTGCCAGCAGCCAAGACAATTCTGGAAATACGCTCTATTTT | 4 | TRBV2*00 | | TRBJ1-3*00 | 1.14% |
| CD4+YFP- WT1 | TGTGCCAGCAGCCAAGAGATTAACCAAGACACCCAGTACTTT | 3 | TRBV5*00 | | TRBJ2-5*00 | 0.86% |
| CD4+YFP- WT1 | TGTGCCAGCGGTGATGGAAGTGCAGAAACGCTGTATTTT | 3 | TRBV13-2*00 | | TRBJ2-3*00 | 0.86% |
| CD4+YFP- WT1 | TGTGCTAGCAGTCAGGGACAATATGAACAGTACTTC | 3 | TRBV21*00 | TRBD1*00 | TRBJ2-7*00 | 0.86% |
| CD4+YFP- WT1 | TGCACCTGCAGTGCACAGGGCCCTAATGCTGAGCAGTTCTTC | 2 | TRBV1*00 | TRBD1*00 | TRBJ2-1*00 | 0.57% |
| CD4+YFP- WT1 | TGCACCTGCAGTGCAGAGGGCACCGGGCAGCTCTACTTT | 2 | TRBV1*00 | | TRBJ2-2*00 | 0.57% |
| CD4+YFP- WT1 | TGTGCCAGCAGCCAACAGGAAAACTCCGACTACACCTTC | 2 | TRBV2*00 | | TRBJ1-2*00 | 0.57% |
| CD4+YFP- WT1 | TGTGCCAGCAGCCCCGACGGGAACACCGGGCAGCTCTACTTT | 2 | TRBV4*00 | | TRBJ2-2*00 | 0.57% |
| CD4+YFP- WT1 | TGTGCCAGCAGCCCGGCTGGAAATACGCTCTATTTT | 2 | TRBV13-1*00 | TRBD1*00 | TRBJ1-3*00 | 0.57% |
| CD4+YFP- WT1 | TGTGCCAGCAGCCCTGGACAGGGCAGACAAAACACCTTGTACTTT | 2 | TRBV2*00 | TRBD1*00 | TRBJ2-4*00 | 0.57% |
| CD4+YFP- WT1 | TGTGCCAGCAGCTAAGAACGGGGGGGCTTGAACAGTACTTC | 2 | TRBV4*00 | TRBD2*00 | TRBJ2-7*00 | 0.57% |
| CD4+YFP- WT1 | TGTGCCAGCAGCTATGCAAACTCCGACTACACCTTC | 2 | TRBV4*00 | | TRBJ1-2*00 | 0.57% |
| CD4+YFP- WT1 | TGTGCCAGCAGTATAGCAGGAACTAGTGCAGAAACGCTGTATTTT | 2 | TRBV19*00 | | TRBJ2-3*00 | 0.57% |
| CD4+YFP- WT1 | TGTGCCAGCAGTCTGCTGGGGGGGCAAGACACCCAGTACTTT | 2 | TRBV26*00 | TRBD2*00 | TRBJ2-5*00 | 0.57% |
| CD4+YFP- WT1 | TGTGCCAGCAGTCTGTACAGTAAACACCGGGCAGCTCTACTTT | 2 | TRBV24*00 | | TRBJ2-2*00 | 0.57% |
| CD4+YFP- WT1 | TGTGCCAGCAGTGATAAGGGGGGGAATTCTGGAAATACGCTCTATTTT | 2 | TRBV13-1*00 | TRBD1*00 | TRBJ1-3*00 | 0.57% |
| CD4+YFP- WT1 | TGTGCCAGCAGTTTTCTGGGGGGGAACTATGCTGAGCAGTTCTTC | 2 | TRBV15*00 | TRBD2*00 | TRBJ2-1*00 | 0.57% |
| CD4+YFP- WT1 | TGTGCCAGCGGTGATGGTAGTGCAGAAACGCTGTATTTT | 2 | TRBV13-2*00 | | TRBJ2-3*00 | 0.57% |
| CD4+YFP- WT1 | TGTGCCAGCGGTGCCGGGGACTGGAGGAGCTCCTATGAACAGTACTTC | 2 | TRBV13-2*00 | TRBD2*00 | TRBJ2-7*00 | 0.57% |
| CD4+YFP- WT1 | TGTGCCAGCTCTCTGGACAACACCGGGCAGCTCTACTTT | 2 | TRBV12-1*00 | | TRBJ2-2*00 | 0.57% |
| CD4+YFP- WT1 | TGTGCTAGCAGTAGAGGGGGCTGGGAACAGTACTTC | 2 | TRBV17*00 | TRBD1*00 | TRBJ2-7*00 | 0.57% |
| CD4+YFP- WT1 | TGTGGTGCTAGGGATCAGGCAAACTCCGACTACACCTTC | 2 | TRBV20*00 | | TRBJ1-2*00 | 0.57% |
| CD4+YFP- WT1 | TGTGCCAGCAGCCAAGACAGCTATGAACAGTACTTC | 2 | TRBV2*00 | | TRBJ2-7*00 | 0.57% |
| CD4+YFP- WT1 | TGCACCTGCACCAGCCAAGACACCCAGTACTTT | 1 | TRBV1*00 | | TRBJ2-5*00 | 0.29% |
| CD4+YFP- WT1 | TGCACCTGCAGTGACAGGGGGCGCCAAGACACCCAGTACTTT | 1 | TRBV1*00 | TRBD1*00 | TRBJ2-5*00 | 0.29% |
| CD4+YFP- WT1 | TGCACCTGCAGTGACAGGGTTTCCAACGAAAGATTATTTTTC | 1 | TRBV1*00 | TRBD1*00 | TRBJ1-4*00 | 0.29% |
| CD4+YFP- WT1 | TGCACCTGCAGTGCAGAACTTTTTAACCAAGACACCCAGTACTTT | 1 | TRBV1*00 | | TRBJ2-5*00 | 0.29% |
| CD4+YFP- WT1 | TGCACCTGCAGTGCAGAAGACACCCAGTACTTT | 1 | TRBV1*00 | | TRBJ2-5*00 | 0.29% |
| CD4+YFP- WT1 | TGCACCTGCAGTGCAGAGACAGATTATGAACAGTACTTC | 1 | TRBV1*00 | TRBD1*00 | TRBJ2-7*00 | 0.29% |
| CD4+YFP- WT1 | TGCACCTGCAGTGCAGAGACTGGGGGCCCTGAGCAGTTCTTC | 1 | TRBV1*00 | TRBD2*00 | TRBJ2-1*00 | 0.29% |
| CD4+YFP- WT1 | TGCACCTGCAGTGCAGAGGGACTGGGGGAAACGCTGTATTTT | 1 | TRBV1*00 | TRBD2*00 | TRBJ2-3*00 | 0.29% |
| CD4+YFP- WT1 | TGCACCTGCAGTGCAGATAGAAATTCTGGAAATACGCTCTATTTT | 1 | TRBV1*00 | | TRBJ1-3*00 | 0.29% |
| CD4+YFP- WT1 | TGCACCTGCAGTGCAGATAGGCTGGGGGGTAGTGCAGAAACGCTGTATTTT | 1 | TRBV1*00 | TRBD2*00 | TRBJ2-3*00 | 0.29% |
| CD4+YFP- WT1 | TGCACCTGCAGTGCAGATAGGGGCATCCAGGCTCCGCTTTTT | 1 | TRBV1*00 | TRBD1*00 | TRBJ1-5*00 | 0.29% |
| CD4+YFP- WT1 | TGCACCTGCAGTGCAGATATGGGCCAGGAGTACTTT | 1 | TRBV1*00 | | TRBJ2-5*00 | 0.29% |
| CD4+YFP- WT1 | TGCACCTGCAGTGCAGATCACAGATATGAACAGTACTTC | 1 | TRBV1*00 | | TRBJ2-7*00 | 0.29% |
| CD4+YFP- WT1 | TGCACCTGCAGTGCAGATCAGGACACCTTGTACTTT | 1 | TRBV1*00 | | TRBJ2-4*00 | 0.29% |
| CD4+YFP- WT1 | TGCACCTGCAGTGCAGATCAGGGGGCAGAAGTCTTCTTT | 1 | TRBV1*00 | TRBD1*00 | TRBJ1-1*00 | 0.29% |
| CD4+YFP- WT1 | TGCACCTGCAGTGCAGATGGGGGGGGTGAACAGTACTTC | 1 | TRBV1*00 | TRBD2*00 | TRBJ2-7*00 | 0.29% |
| CD4+YFP- WT1 | TGCACCTGCAGTGCAGCCCATGAACAGTACTTC | 1 | TRBV1*00 | | TRBJ2-7*00 | 0.29% |
| CD4+YFP- WT1 | TGCACCTGCAGTGCAGGGAACTATGCTGAGCAGTTCTTC | 1 | TRBV1*00 | | TRBJ2-1*00 | 0.29% |
| CD4+YFP- WT1 | TGCACCTGCAGTGCAGGGGGGAACCAGGCTCCGCTTTTT | 1 | TRBV1*00 | TRBD1*00 | TRBJ1-5*00 | 0.29% |
| CD4+YFP- WT1 | TGCACCTGCAGTGCCCCCCCTGCCACCGGGCAGCTCTACTTT | 1 | TRBV1*00 | TRBD1*00 | TRBJ2-2*00 | 0.29% |
| CD4+YFP- WT1 | TGCACCTGCAGTGCCCGGACAGGGGCAAACTCCGACTACACCTTC | 1 | TRBV1*00 | TRBD1*00 | TRBJ1-2*00 | 0.29% |
| CD4+YFP- WT1 | TGCACCTGCAGTGCCTGGGGAAACTCCGACTACACCTTC | 1 | TRBV1*00 | TRBD2*00 | TRBJ1-2*00 | 0.29% |
| CD4+YFP- WT1 | TGCACCTGCAGTGCGCCCGGTTATGAACAGTACTTC | 1 | TRBV1*00 | TRBD1*00 | TRBJ2-7*00 | 0.29% |
| CD4+YFP- WT1 | TGCACCTGCAGTGCGGACAGGGGGAGTCAAAACACCTTGTACTTT | 1 | TRBV1*00 | TRBD1*00 | TRBJ2-4*00 | 0.29% |
| CD4+YFP- WT1 | TGCACCTGCAGTGCTTCGGGGAACCAGGCTCCGCTTTTT | 1 | TRBV1*00 | | TRBJ1-5*00 | 0.29% |
| CD4+YFP- WT1 | TGCACCTGCAGTGGGACACCTAAG | 1 | TRBV1*00 | TRBD1*00 | TRBJ2-4*00 | 0.29% |
| CD4+YFP- WT1 | TGCACCTGCAGTGGGACAGCAAACACAGAAGTCTTCTTT | 1 | TRBV1*00 | TRBD1*00 | TRBJ1-1*00 | 0.29% |
| CD4+YFP- WT1 | TGCACCTGCAGTGGGGACAGACAGGCTCCGCTTTTT | 1 | TRBV1*00 | TRBD1*00 | TRBJ1-5*00 | 0.29% |
| CD4+YFP- WT1 | TGCACCTGCAGTGGGTTCGGGACATCAGAAGTCTTCTTT | 1 | TRBV1*00 | TRBD1*00 | TRBJ1-1*00 | 0.29% |
| CD4+YFP- WT1 | TGCTCCAGCAGTGGGACAATCCTATGAACAGTACTTC | 1 | TRBV23*00 | TRBD1*00 | TRBJ2-7*00 | 0.29% |
| CD4+YFP- WT1 | TGTAGTTCTAGTGGGCACTATGAACAGTACTTC | 1 | TRBV30*00 | | TRBJ2-7*00 | 0.29% |
| CD4+YFP- WT1 | TGTGCAAGCAGCCCGACAGGTGCAGAAACGCTGTATTTT | 1 | TRBV16*00 | TRBD1*00 | TRBJ2-3*00 | 0.29% |

| | | | | | | |
|---|---|---|---|---|---|---|
| CD4+YFP- WT1 | TGTGCAAGCAGCCCTCAGGGGCGAGACACCCAGTACTTT | 1 | TRBV16*00 | TRBD1*00 | TRBJ2-5*00 | 0.29% |
| CD4+YFP- WT1 | TGTGCAAGCAGCCTAGAGGGTGAACAGTACTTC | 1 | TRBV16*00 | | TRBJ2-7*00 | 0.29% |
| CD4+YFP- WT1 | TGTGCAAGCAGCGTCGACAGGGGTGAACAGTACTTC | 1 | TRBV16*00 | TRBD1*00 | TRBJ2-7*00 | 0.29% |
| CD4+YFP- WT1 | TGTGCAAGCAGCTCAGGGGTTGCTGAGCAGTTCTTC | 1 | TRBV16*00 | TRBD1*00 | TRBJ2-1*00 | 0.29% |
| CD4+YFP- WT1 | TGTGCAAGCAGCTTACTGGGGAGCTCCTATGAACAGTACTTC | 1 | TRBV16*00 | | TRBJ2-7*00 | 0.29% |
| CD4+YFP- WT1 | TGTGCAAGCAGCTTAGAACCAAACTCCGACTACACCTTC | 1 | TRBV16*00 | | TRBJ1-2*00 | 0.29% |
| CD4+YFP- WT1 | TGTGCAAGCAGCTTAGACAGGGAAACGCTGTATTTT | 1 | TRBV16*00 | | TRBJ2-3*00 | 0.29% |
| CD4+YFP- WT1 | TGTGCAAGCAGCTTAGGGGGCCAAGACACCCAGTACTTT | 1 | TRBV16*00 | | TRBJ2-5*00 | 0.29% |
| CD4+YFP- WT1 | TGTGCAAGCAGCTTAGTAGACACCCAGTACTTT | 1 | TRBV16*00 | | TRBJ2-5*00 | 0.29% |
| CD4+YFP- WT1 | TGTGCAAGCAGCTTATGGGGAGCTCCTATGAACAGTACTTC | 1 | TRBV16*00 | | TRBJ2-7*00 | 0.29% |
| CD4+YFP- WT1 | TGTGCAAGCAGCTTCCGACAGGGGCAAGACACCTAGTACTTT | 1 | TRBV16*00 | TRBD1*00 | TRBJ2-5*00 | 0.29% |
| CD4+YFP- WT1 | TGTGCAAGCAGCTTCGCACTGGGGGACCAAGACACCCAGTACTTT | 1 | TRBV16*00 | TRBD2*00 | TRBJ2-5*00 | 0.29% |
| CD4+YFP- WT1 | TGTGCAAGCAGGCAGGGGGCAGAAACGCTGTATTTT | 1 | TRBV16*00 | TRBD1*00 | TRBJ2-3*00 | 0.29% |
| CD4+YFP- WT1 | TGTGCCACCGGGACAACAAACTCCGACTACACCTTC | 1 | TRBV13-2*00 | | TRBJ1-2*00 | 0.29% |
| CD4+YFP- WT1 | TGTGCCAGCAAAAGGGGGACCAACGAAAGATTATTTTTC | 1 | TRBV13-3*00 | TRBD1*00 | TRBJ1-4*00 | 0.29% |
| CD4+YFP- WT1 | TGTGCCAGCAACCCCAGCTCCTATGAACAGTACTTC | 1 | TRBV4*00 | | TRBJ2-7*00 | 0.29% |
| CD4+YFP- WT1 | TGTGCCAGCAACCCGACTGGGGAGAGGGCCTTGTACTTT | 1 | TRBV13-3*00 | TRBD2*00 | TRBJ2-4*00 | 0.29% |
| CD4+YFP- WT1 | TGTGCCAGCAATGACAGGGGGAACGAAAGATTATTTTTC | 1 | TRBV14*00 | TRBD1*00 | TRBJ1-4*00 | 0.29% |
| CD4+YFP- WT1 | TGTGCCAGCACCCAGAACACAGAAGTCTTCTTT | 1 | TRBV13-3*00 | TRBD2*00 | TRBJ1-1*00 | 0.29% |
| CD4+YFP- WT1 | TGTGCCAGCACCCGGGACAAAAACACCTTGTACTTT | 1 | TRBV19*00 | TRBD1*00 | TRBJ2-4*00 | 0.29% |
| CD4+YFP- WT1 | TGTGCCAGCACCTTAAACACCGGGCAGCTCTACTTT | 1 | TRBV3*00 | | TRBJ2-2*00 | 0.29% |
| CD4+YFP- WT1 | TGTGCCAGCAGAAGGACTGGGAACTATGCTGAGCAGTTCTTC | 1 | TRBV13-1*00 | TRBD2*00 | TRBJ2-1*00 | 0.29% |
| CD4+YFP- WT1 | TGTGCCAGCAGCAACAGGGAGGAAGTCTTCTTT | 1 | TRBV13-1*00 | TRBD1*00 | TRBJ1-1*00 | 0.29% |
| CD4+YFP- WT1 | TGTGCCAGCAGCCAACCGAATTCGCCCCTCTACTTT | 1 | TRBV19*00 | | TRBJ1-6*00 | 0.29% |
| CD4+YFP- WT1 | TGTGCCAGCAGCCAAGAACTTCTCTATGAACAGTACTTC | 1 | TRBV5*00 | | TRBJ2-7*00 | 0.29% |
| CD4+YFP- WT1 | TGTGCCAGCAGCCAAGAAGGGCTCCAAGACACCCAGTACTTT | 1 | TRBV5*00 | | TRBJ2-5*00 | 0.29% |
| CD4+YFP- WT1 | TGTGCCAGCAGCCAAGACAGTAACTATGCTGAGCAGTTCTTC | 1 | TRBV5*00 | | TRBJ2-1*00 | 0.29% |
| CD4+YFP- WT1 | TGTGCCAGCAGCCAAGACCCTGGGGGCAACACCGGGCAGCTCTACTTT | 1 | TRBV5*00 | TRBD2*00 | TRBJ2-2*00 | 0.29% |
| CD4+YFP- WT1 | TGTGCCAGCAGCCAAGACGACAACTATGAACAGTACTTC | 1 | TRBV5*00 | | TRBJ2-7*00 | 0.29% |
| CD4+YFP- WT1 | TGTGCCAGCAGCCAAGACGGAGCAAACACCGGGCAGCTCTACTTT | 1 | TRBV5*00 | | TRBJ2-2*00 | 0.29% |
| CD4+YFP- WT1 | TGTGCCAGCAGCCAAGACTGGGGGGACTATGCTGAGCAGTTCTTC | 1 | TRBV5*00 | TRBD2*00 | TRBJ2-1*00 | 0.29% |
| CD4+YFP- WT1 | TGTGCCAGCAGCCAAGAGAATTGGGGGAATGAACAGTACTTC | 1 | TRBV5*00 | TRBD2*00 | TRBJ2-7*00 | 0.29% |
| CD4+YFP- WT1 | TGTGCCAGCAGCCAAGAGACAGGGGACACCGGGCAGCTCTACTTT | 1 | TRBV5*00 | TRBD1*00 | TRBJ2-2*00 | 0.29% |
| CD4+YFP- WT1 | TGTGCCAGCAGCCAAGATCGACAGGGTAACTATGCTGAGCAGTTCTTC | 1 | TRBV5*00 | TRBD1*00 | TRBJ2-1*00 | 0.29% |
| CD4+YFP- WT1 | TGTGCCAGCAGCCAAGATGGCAGGGGGACTAGTCAAAACACCTTGTACTTT | 1 | TRBV5*00 | TRBD1*00 | TRBJ2-4*00 | 0.29% |
| CD4+YFP- WT1 | TGTGCCAGCAGCCAAGATGGGGAAGACACCCAGTACTTT | 1 | TRBV5*00 | | TRBJ2-5*00 | 0.29% |
| CD4+YFP- WT1 | TGTGCCAGCAGCCAAGATTCAGGAGACGAAAGATTATTTTTC | 1 | TRBV5*00 | | TRBJ1-4*00 | 0.29% |
| CD4+YFP- WT1 | TGTGCCAGCAGCCAAGGGTGGGGGGATGCTGAGCAGTTCTTC | 1 | TRBV5*00 | TRBD2*00 | TRBJ2-1*00 | 0.29% |
| CD4+YFP- WT1 | TGTGCCAGCAGCCAATATGAACAGTACTTC | 1 | TRBV4*00 | | TRBJ2-7*00 | 0.29% |
| CD4+YFP- WT1 | TGTGCCAGCAGCCACCAAGACACCCAGTACTTT | 1 | TRBV5*00 | | TRBJ2-5*00 | 0.29% |
| CD4+YFP- WT1 | TGTGCCAGCAGCCACCCCGGGACAGGGAGTGACACCCAGTACTTT | 1 | TRBV2*00 | TRBD1*00 | TRBJ2-5*00 | 0.29% |
| CD4+YFP- WT1 | TGTGCCAGCAGCCACCGGGGAAACCAAGACACCCAGTACTTT | 1 | TRBV2*00 | TRBD1*00 | TRBJ2-5*00 | 0.29% |
| CD4+YFP- WT1 | TGTGCCAGCAGCCAGACGGGCACAGAAGTCTTCTTT | 1 | TRBV2*00 | | TRBJ1-1*00 | 0.29% |
| CD4+YFP- WT1 | TGTGCCAGCAGCCAGAGGACAGGACCCACAGAAGTCTTCTTT | 1 | TRBV5*00 | TRBD1*00 | TRBJ1-1*00 | 0.29% |
| CD4+YFP- WT1 | TGTGCCAGCAGCCAGAGGGGTAGTCAAAACACCTTGTACTTT | 1 | TRBV4*00 | TRBD1*00 | TRBJ2-4*00 | 0.29% |
| CD4+YFP- WT1 | TGTGCCAGCAGCCAGGACAATACCTATGAACAGTACTTC | 1 | TRBV5*00 | TRBD1*00 | TRBJ2-7*00 | 0.29% |
| CD4+YFP- WT1 | TGTGCCAGCAGCCCAAGACAATTCTATGAACAGTACTTC | 1 | TRBV2*00 | | TRBJ2-7*00 | 0.29% |
| CD4+YFP- WT1 | TGTGCCAGCAGCCCCCCTGGGGGCTATGCTGAGCAGTTCTTC | 1 | TRBV2*00 | TRBD2*00 | TRBJ2-1*00 | 0.29% |
| CD4+YFP- WT1 | TGTGCCAGCAGCCCCGGGACAGGGATGGGTCAAAACACCTTGTACTTT | 1 | TRBV2*00 | TRBD1*00 | TRBJ2-4*00 | 0.29% |
| CD4+YFP- WT1 | TGTGCCAGCAGCCCGCCCGGGACTGGGGGGTGGAACAGTACTTC | 1 | TRBV5*00 | TRBD2*00 | TRBJ2-7*00 | 0.29% |
| CD4+YFP- WT1 | TGTGCCAGCAGCCCTCGGACTGGGGGGTATGAACAGTACTTC | 1 | TRBV2*00 | TRBD2*00 | TRBJ2-7*00 | 0.29% |
| CD4+YFP- WT1 | TGTGCCAGCAGCCGAGACACAGAAGTCTTCTTT | 1 | TRBV2*00 | | TRBJ1-1*00 | 0.29% |
| CD4+YFP- WT1 | TGTGCCAGCAGCCGCGGGACGGCAAACTCCGACTACACCTTC | 1 | TRBV2*00 | TRBD1*00 | TRBJ1-2*00 | 0.29% |
| CD4+YFP- WT1 | TGTGCCAGCAGCCGGACTGGGGGGGGTATGAACAGTACTTC | 1 | TRBV2*00 | TRBD2*00 | TRBJ2-7*00 | 0.29% |
| CD4+YFP- WT1 | TGTGCCAGCAGCCGGGACAGGGGAGACGAAAGATTATTTTTC | 1 | TRBV4*00 | TRBD1*00 | TRBJ1-4*00 | 0.29% |
| CD4+YFP- WT1 | TGTGCCAGCAGCCGGGGGGGGGCCCAAGACACCCAGTACTTT | 1 | TRBV2*00 | TRBD2*00 | TRBJ2-5*00 | 0.29% |
| CD4+YFP- WT1 | TGTGCCAGCAGCCTCTGACAGTTGCAAACTCCGACTACACCTTC | 1 | TRBV3*00 | TRBD1*00 | TRBJ1-2*00 | 0.29% |
| CD4+YFP- WT1 | TGTGCCAGCAGCCTTAACCAAGACACCCAGTACTTT | 1 | TRBV4*00 | | TRBJ2-5*00 | 0.29% |
| CD4+YFP- WT1 | TGTGCCAGCAGCGCGGACAGTTCTGGAAATACGCTCTATTTT | 1 | TRBV4*00 | TRBD1*00 | TRBJ1-3*00 | 0.29% |
| CD4+YFP- WT1 | TGTGCCAGCAGCGGAAGCTCCTATGAACAGTACTTC | 1 | TRBV15*00 | | TRBJ2-7*00 | 0.29% |
| CD4+YFP- WT1 | TGTGCCAGCAGCGGGACAGGGGTGAACACAGAAGTCTTCTTT | 1 | TRBV15*00 | TRBD1*00 | TRBJ1-1*00 | 0.29% |
| CD4+YFP- WT1 | TGTGCCAGCAGCTACAATCCAAACTCCGACTACACCTTC | 1 | TRBV4*00 | | TRBJ1-2*00 | 0.29% |
| CD4+YFP- WT1 | TGTGCCAGCAGCTACAGGGCCAACCAGGCTCCGCTTTTT | 1 | TRBV4*00 | TRBD1*00 | TRBJ1-5*00 | 0.29% |
| CD4+YFP- WT1 | TGTGCCAGCAGCTATCACAGAAACACCTTGTACTTT | 1 | TRBV4*00 | | TRBJ2-4*00 | 0.29% |
| CD4+YFP- WT1 | TGTGCCAGCAGCTATGCAAACTCCGACTACACCGTC | 1 | TRBV4*00 | | TRBJ1-2*00 | 0.29% |
| CD4+YFP- WT1 | TGTGCCAGCAGCTATGGGGACACAGAAGTCTTCTTT | 1 | TRBV4*00 | TRBD2*00 | TRBJ1-1*00 | 0.29% |
| CD4+YFP- WT1 | TGTGCCAGCAGCTATGGGGGAGCTCCTATGAACAGTACTTC | 1 | TRBV4*00 | TRBD2*00 | TRBJ2-7*00 | 0.29% |
| CD4+YFP- WT1 | TGTGCCAGCAGCTATGGGGGTGGCCAAGACACCCAGTACTTT | 1 | TRBV4*00 | TRBD2*00 | TRBJ2-5*00 | 0.29% |
| CD4+YFP- WT1 | TGTGCCAGCAGCTCCGACAAGTATGAACAGTACTTC | 1 | TRBV3*00 | | TRBJ2-7*00 | 0.29% |
| CD4+YFP- WT1 | TGTGCCAGCAGCTCCGGACTGGGACAAGACACCCAGTACTTT | 1 | TRBV3*00 | TRBD2*00 | TRBJ2-5*00 | 0.29% |
| CD4+YFP- WT1 | TGTGCCAGCAGCTCCGGGACACCAAACTCCGACTACACCTTC | 1 | TRBV25*00 | TRBD1*00 | TRBJ1-2*00 | 0.29% |
| CD4+YFP- WT1 | TGTGCCAGCAGCTCCTGGGGGGAAACGCTGTATTTT | 1 | TRBV4*00 | TRBD2*00 | TRBJ2-3*00 | 0.29% |
| CD4+YFP- WT1 | TGTGCCAGCAGCTCGGGACAGCATGTTGAGCAGTTCTTC | 1 | TRBV3*00 | TRBD1*00 | TRBJ2-1*00 | 0.29% |
| CD4+YFP- WT1 | TGTGCCAGCAGCTCGGGACTGGGGGGGCGAGAACAGTACTTC | 1 | TRBV4*00 | TRBD2*00 | TRBJ2-7*00 | 0.29% |
| CD4+YFP- WT1 | TGTGCCAGCAGCTCTACAAATAACCAAGACACCCAGTACTTT | 1 | TRBV25*00 | | TRBJ2-5*00 | 0.29% |

| | | | | | | |
|---|---|---|---|---|---|---|
| CD4+YFP- WT1 | TGTGCCAGCAGCTCTGTCAACCAGGCTCCGCTTTTT | 1 | TRBV4*00 | | TRBJ1-5*00 | 0.29% |
| CD4+YFP- WT1 | TGTGCCAGCAGCTGGCAGGGGGGTCAAAACACCTTGTACTTT | 1 | TRBV4*00 | TRBD1*00 | TRBJ2-4*00 | 0.29% |
| CD4+YFP- WT1 | TGTGCCAGCAGCTGGGACGACCAGGCTCCGCTTTTT | 1 | TRBV4*00 | TRBD1*00 | TRBJ1-5*00 | 0.29% |
| CD4+YFP- WT1 | TGTGCCAGCAGCTTAAGAGGGTCTAGTGCAGAAACGCTGTATTTT | 1 | TRBV3*00 | | TRBJ2-3*00 | 0.29% |
| CD4+YFP- WT1 | TGTGCCAGCAGCTTACAGGGCTTTTCCAACGAAAGATTATTTTTC | 1 | TRBV3*00 | TRBD1*00 | TRBJ1-4*00 | 0.29% |
| CD4+YFP- WT1 | TGTGCCAGCAGCTTACCCCCCTTGTACTTT | 1 | TRBV3*00 | | TRBJ2-4*00 | 0.29% |
| CD4+YFP- WT1 | TGTGCCAGCAGCTTAGAACAGGAAGTCTTCTTT | 1 | TRBV3*00 | | TRBJ1-1*00 | 0.29% |
| CD4+YFP- WT1 | TGTGCCAGCAGCTTAGACGGCTATGAACAGTACTTC | 1 | TRBV3*00 | | TRBJ2-7*00 | 0.29% |
| CD4+YFP- WT1 | TGTGCCAGCAGCTTAGACGGGGGTTCTGGAAATACGCTCTATTTT | 1 | TRBV3*00 | TRBD1*00 | TRBJ1-3*00 | 0.29% |
| CD4+YFP- WT1 | TGTGCCAGCAGCTTAGAGGGGGGGGGAGACACCCAGTACTTT | 1 | TRBV3*00 | TRBD2*00 | TRBJ2-5*00 | 0.29% |
| CD4+YFP- WT1 | TGTGCCAGCAGCTTAGCGGCAGGGGATTCTGGAAATACGCTCTATTTT | 1 | TRBV3*00 | TRBD1*00 | TRBJ1-3*00 | 0.29% |
| CD4+YFP- WT1 | TGTGCCAGCAGCTTAGGGAAGAGGGCAAACACAGAAGTCTTCTTT | 1 | TRBV3*00 | | TRBJ1-1*00 | 0.29% |
| CD4+YFP- WT1 | TGTGCCAGCAGCTTAGGTCAAAACACCTTGTACTTT | 1 | TRBV3*00 | | TRBJ2-4*00 | 0.29% |
| CD4+YFP- WT1 | TGTGCCAGCAGCTTAGTAGGGAAC | 1 | TRBV3*00 | | TRBJ2-7*00 | 0.29% |
| CD4+YFP- WT1 | TGTGCCAGCAGCTTATCTGGGGCAAACACCGGGCAGCTCTACTTT | 1 | TRBV3*00 | TRBD2*00 | TRBJ2-2*00 | 0.29% |
| CD4+YFP- WT1 | TGTGCCAGCAGCTTCAGGGGAGGAGACACCCAGTACTTT | 1 | TRBV4*00 | TRBD1*00 | TRBJ2-5*00 | 0.29% |
| CD4+YFP- WT1 | TGTGCCAGCAGCTTCCAAAACCAGGCTCCGCTTTTT | 1 | TRBV4*00 | | TRBJ1-5*00 | 0.29% |
| CD4+YFP- WT1 | TGTGCCAGCAGCTTCCGGGACTGGGGGGGCCTCCTATGAACAGTACTTC | 1 | TRBV4*00 | TRBD2*00 | TRBJ2-7*00 | 0.29% |
| CD4+YFP- WT1 | TGTGCCAGCAGCTTCCGGGGAAACTCCGACTACACCTTC | 1 | TRBV3*00 | TRBD1*00 | TRBJ1-2*00 | 0.29% |
| CD4+YFP- WT1 | TGTGCCAGCAGCTTCGGGACTTCTAGTCAAAACACCTTGTACTTT | 1 | TRBV4*00 | TRBD2*00 | TRBJ2-4*00 | 0.29% |
| CD4+YFP- WT1 | TGTGCCAGCAGCTTTCAGACAGAAGTCTTCTTT | 1 | TRBV3*00 | | TRBJ1-1*00 | 0.29% |
| CD4+YFP- WT1 | TGTGCCAGCAGGAGACAGGGGAACTCCGACTACACCTTC | 1 | TRBV13-1*00 | TRBD1*00 | TRBJ1-2*00 | 0.29% |
| CD4+YFP- WT1 | TGTGCCAGCAGGAGTTATGCTGAGCAGTTCTTC | 1 | TRBV3*00 | | TRBJ2-1*00 | 0.29% |
| CD4+YFP- WT1 | TGTGCCAGCAGGCGGGGACAGGGGGCGCTTGCTGAGCAGTTCTTC | 1 | TRBV4*00 | TRBD1*00 | TRBJ2-1*00 | 0.29% |
| CD4+YFP- WT1 | TGTGCCAGCAGTAAGAACTCCGACTACACCTTC | 1 | TRBV26*00 | | TRBJ1-2*00 | 0.29% |
| CD4+YFP- WT1 | TGTGCCAGCAGTAAGGGGGGGCTGGAACAGTACTTC | 1 | TRBV13-1*00 | TRBD2*00 | TRBJ2-7*00 | 0.29% |
| CD4+YFP- WT1 | TGTGCCAGCAGTAGGACAGGGGGCTATGCTGAGCAGTTCTTC | 1 | TRBV15*00 | TRBD1*00 | TRBJ2-1*00 | 0.29% |
| CD4+YFP- WT1 | TGTGCCAGCAGTAGGGACATAAACTCCGACTACACCTTC | 1 | TRBV19*00 | TRBD1*00 | TRBJ1-2*00 | 0.29% |
| CD4+YFP- WT1 | TGTGCCAGCAGTATAAGGGCACTTTCCAACGAAAGATTATTTTTC | 1 | TRBV19*00 | | TRBJ1-4*00 | 0.29% |
| CD4+YFP- WT1 | TGTGCCAGCAGTATAAGGGGAGGGAGCGAAAGATTATTTTTC | 1 | TRBV19*00 | TRBD1*00 | TRBJ1-4*00 | 0.29% |
| CD4+YFP- WT1 | TGTGCCAGCAGTATAATTCAAAACACCTTGTACTTT | 1 | TRBV19*00 | | TRBJ2-4*00 | 0.29% |
| CD4+YFP- WT1 | TGTGCCAGCAGTATAGACAGGGGAGACACCCAGTACTTT | 1 | TRBV19*00 | TRBD1*00 | TRBJ2-5*00 | 0.29% |
| CD4+YFP- WT1 | TGTGCCAGCAGTATAGACCTCTATGCTGAGCAGTTCTTC | 1 | TRBV19*00 | | TRBJ2-1*00 | 0.29% |
| CD4+YFP- WT1 | TGTGCCAGCAGTATAGGACATTCAAACACCGGGCAGCTCTACTTT | 1 | TRBV19*00 | | TRBJ2-2*00 | 0.29% |
| CD4+YFP- WT1 | TGTGCCAGCAGTATCCAAAACACCTTGTACTTT | 1 | TRBV19*00 | | TRBJ2-4*00 | 0.29% |
| CD4+YFP- WT1 | TGTGCCAGCAGTATGGGGAATTCTGGAAATACGCTCTATTTT | 1 | TRBV19*00 | | TRBJ1-3*00 | 0.29% |
| CD4+YFP- WT1 | TGTGCCAGCAGTATGGGGACAGTTGCTGAGCAGTTCTTC | 1 | TRBV19*00 | TRBD1*00 | TRBJ2-1*00 | 0.29% |
| CD4+YFP- WT1 | TGTGCCAGCAGTATTGGCTGGGGGGTTGAACAGTACTTC | 1 | TRBV19*00 | TRBD2*00 | TRBJ2-7*00 | 0.29% |
| CD4+YFP- WT1 | TGTGCCAGCAGTATTGGGAACTCCGACTACACCTTC | 1 | TRBV19*00 | | TRBJ1-2*00 | 0.29% |
| CD4+YFP- WT1 | TGTGCCAGCAGTATTTCCCATTCTGGAAATACGCTCTATTTT | 1 | TRBV19*00 | | TRBJ1-3*00 | 0.29% |
| CD4+YFP- WT1 | TGTGCCAGCAGTCAAATATATGAACAGTACTTC | 1 | TRBV26*00 | | TRBJ2-7*00 | 0.29% |
| CD4+YFP- WT1 | TGTGCCAGCAGTCAGGGGGACACCCAGTACTTT | 1 | TRBV21*00 | | TRBJ2-5*00 | 0.29% |
| CD4+YFP- WT1 | TGTGCCAGCAGTCAGGGGTTCTTCTTT | 1 | TRBV4*00 | TRBD1*00 | TRBJ1-1*00 | 0.29% |
| CD4+YFP- WT1 | TGTGCCAGCAGTCCCGACAATTATGCTGAGCAGTTCTTC | 1 | TRBV19*00 | | TRBJ2-1*00 | 0.29% |
| CD4+YFP- WT1 | TGTGCCAGCAGTCCCGGGACAGTCCGATTTTTC | 1 | TRBV13-1*00 | TRBD1*00 | TRBJ1-4*00 | 0.29% |
| CD4+YFP- WT1 | TGTGCCAGCAGTCCGACTGGGGGGGATGCTGAGCAGTTCTTC | 1 | TRBV15*00 | TRBD2*00 | TRBJ2-1*00 | 0.29% |
| CD4+YFP- WT1 | TGTGCCAGCAGTCCTAACACAGAAGTCTTCTTT | 1 | TRBV13-3*00 | | TRBJ1-1*00 | 0.29% |
| CD4+YFP- WT1 | TGTGCCAGCAGTCCTACTGGGGGGGAAAACACCTTGTACTTT | 1 | TRBV13-3*00 | TRBD2*00 | TRBJ2-4*00 | 0.29% |
| CD4+YFP- WT1 | TGTGCCAGCAGTCGACAATATGAACAGTACTTC | 1 | TRBV19*00 | | TRBJ2-7*00 | 0.29% |
| CD4+YFP- WT1 | TGTGCCAGCAGTCGACTGGGGGTCCAAGACACCCAGTACTTT | 1 | TRBV19*00 | TRBD2*00 | TRBJ2-5*00 | 0.29% |
| CD4+YFP- WT1 | TGTGCCAGCAGTCGAGGGGCTGAGCAGTTCTTC | 1 | TRBV26*00 | | TRBJ2-1*00 | 0.29% |
| CD4+YFP- WT1 | TGTGCCAGCAGTCGCCGGGACAGGGGTCGGGCTGAGCAGTTCTTC | 1 | TRBV26*00 | TRBD1*00 | TRBJ2-1*00 | 0.29% |
| CD4+YFP- WT1 | TGTGCCAGCAGTCGCGTTGCTGAGCAGTTCTTC | 1 | TRBV26*00 | | TRBJ2-1*00 | 0.29% |
| CD4+YFP- WT1 | TGTGCCAGCAGTCGGGAAATCTATGCTGAGCAGTTCTTC | 1 | TRBV24*00 | | TRBJ2-1*00 | 0.29% |
| CD4+YFP- WT1 | TGTGCCAGCAGTCTCAGGGGCGGCGAAAGATTATTTTTC | 1 | TRBV24*00 | TRBD1*00 | TRBJ1-4*00 | 0.29% |
| CD4+YFP- WT1 | TGTGCCAGCAGTCTCGGTAACTATGCTGAGCAGTTCTTC | 1 | TRBV26*00 | | TRBJ2-1*00 | 0.29% |
| CD4+YFP- WT1 | TGTGCCAGCAGTCTGACAGGGGGCTTTGGAAATACGCTCTATTTT | 1 | TRBV24*00 | TRBD1*00 | TRBJ1-3*00 | 0.29% |
| CD4+YFP- WT1 | TGTGCCAGCAGTCTGAGAAACTCCGACTACACCTTC | 1 | TRBV26*00 | | TRBJ1-2*00 | 0.29% |
| CD4+YFP- WT1 | TGTGCCAGCAGTCTGCAGGCAAACACAGAAGTCTTCTTT | 1 | TRBV24*00 | | TRBJ1-1*00 | 0.29% |
| CD4+YFP- WT1 | TGTGCCAGCAGTCTGGCAGGCTCCTATGAACAGTACTTC | 1 | TRBV26*00 | | TRBJ2-7*00 | 0.29% |
| CD4+YFP- WT1 | TGTGCCAGCAGTCTGGGACAGACAAACACCGGGCAGCTCTACTTT | 1 | TRBV26*00 | | TRBJ2-2*00 | 0.29% |
| CD4+YFP- WT1 | TGTGCCAGCAGTCTGTCCAGATATGAACAGTACTTC | 1 | TRBV26*00 | | TRBJ2-7*00 | 0.29% |
| CD4+YFP- WT1 | TGTGCCAGCAGTCTGTTGGCAGGGGAAGTCTTCTTT | 1 | TRBV26*00 | TRBD1*00 | TRBJ1-1*00 | 0.29% |
| CD4+YFP- WT1 | TGTGCCAGCAGTCTTAACCAAGACACCCAGTACTTT | 1 | TRBV26*00 | | TRBJ2-5*00 | 0.29% |
| CD4+YFP- WT1 | TGTGCCAGCAGTCTTGGAGGGAACCAAGACACCCAGTACTTT | 1 | TRBV26*00 | | TRBJ2-5*00 | 0.29% |
| CD4+YFP- WT1 | TGTGCCAGCAGTGAAACAGCAAACACAGAAGTCTTCTTT | 1 | TRBV13-1*00 | | TRBJ1-1*00 | 0.29% |
| CD4+YFP- WT1 | TGTGCCAGCAGTGAAAGACAGTATGAACAGTACTTC | 1 | TRBV13-3*00 | TRBD1*00 | TRBJ2-7*00 | 0.29% |
| CD4+YFP- WT1 | TGTGCCAGCAGTGACAAGAACTCCGACTACACCTTC | 1 | TRBV13-3*00 | | TRBJ1-2*00 | 0.29% |
| CD4+YFP- WT1 | TGTGCCAGCAGTGACAGGGGGCAAGACACCCAGTACTTT | 1 | TRBV13-3*00 | TRBD1*00 | TRBJ2-5*00 | 0.29% |
| CD4+YFP- WT1 | TGTGCCAGCAGTGACAGGGGGTCAAACTCCGACTACACCTTC | 1 | TRBV13-3*00 | TRBD1*00 | TRBJ1-2*00 | 0.29% |
| CD4+YFP- WT1 | TGTGCCAGCAGTGAGACACTTAACCAAGACACCCAGTACTTT | 1 | TRBV13-1*00 | | TRBJ2-5*00 | 0.29% |
| CD4+YFP- WT1 | TGTGCCAGCAGTGAGTGGGGTAGTCAAAACACCTTGTACTTT | 1 | TRBV13-1*00 | TRBD2*00 | TRBJ2-4*00 | 0.29% |
| CD4+YFP- WT1 | TGTGCCAGCAGTGATAACAAGTATGAACAGTACTTC | 1 | TRBV13-1*00 | | TRBJ2-7*00 | 0.29% |
| CD4+YFP- WT1 | TGTGCCAGCAGTGATCGGACTTCTGGAAATACGCTCTATTTT | 1 | TRBV13-3*00 | | TRBJ1-3*00 | 0.29% |
| CD4+YFP- WT1 | TGTGCCAGCAGTGATGACAGGGGATCCGACTACACCTTC | 1 | TRBV13-1*00 | TRBD1*00 | TRBJ1-2*00 | 0.29% |

| | | | | | | |
|---|---|---|---|---|---|---|
| CD4+YFP- WT1 | TGTGCCAGCAGTGATGAGAATTCTGGAAATACGCTCTATTTT | 1 | TRBV13-3*00 | | TRBJ1-3*00 | 0.29% |
| CD4+YFP- WT1 | TGTGCCAGCAGTGATGAGGGGGGGGGAGGAAGACACCCAGTACTTT | 1 | TRBV13-1*00 | TRBD2*00 | TRBJ2-5*00 | 0.29% |
| CD4+YFP- WT1 | TGTGCCAGCAGTGATGCAAAAGAAGTCTTCTTT | 1 | TRBV13-1*00 | | TRBJ1-1*00 | 0.29% |
| CD4+YFP- WT1 | TGTGCCAGCAGTGATGCACAGAATAACAACCAGGCTCCGCTTTTT | 1 | TRBV13-3*00 | | TRBJ1-5*00 | 0.29% |
| CD4+YFP- WT1 | TGTGCCAGCAGTGATGCGGGGACTGCCTATGAACAGTACTTC | 1 | TRBV13-1*00 | TRBD2*00 | TRBJ2-7*00 | 0.29% |
| CD4+YFP- WT1 | TGTGCCAGCAGTGATGCTGGGGGGGACACCCAGTACTTT | 1 | TRBV13-1*00 | TRBD2*00 | TRBJ2-5*00 | 0.29% |
| CD4+YFP- WT1 | TGTGCCAGCAGTGATGGCACCGGGCAGCTCTACTTT | 1 | TRBV13-1*00 | | TRBJ2-2*00 | 0.29% |
| CD4+YFP- WT1 | TGTGCCAGCAGTGATGGCACCTTGTACTTT | 1 | TRBV13-1*00 | | TRBJ2-4*00 | 0.29% |
| CD4+YFP- WT1 | TGTGCCAGCAGTGATGGCTCCACTTTTGCTGAGCAGTTCTTC | 1 | TRBV13-1*00 | | TRBJ2-1*00 | 0.29% |
| CD4+YFP- WT1 | TGTGCCAGCAGTGATGGGGGGAACTCCGACTACACCTTC | 1 | TRBV13-1*00 | TRBD1*00 | TRBJ1-2*00 | 0.29% |
| CD4+YFP- WT1 | TGTGCCAGCAGTGATGTCGCTGAGCAGTTCTTC | 1 | TRBV13-3*00 | | TRBJ2-1*00 | 0.29% |
| CD4+YFP- WT1 | TGTGCCAGCAGTGATTTGGATAGTGCAGAAACGCTGTATTTT | 1 | TRBV13-3*00 | | TRBJ2-3*00 | 0.29% |
| CD4+YFP- WT1 | TGTGCCAGCAGTGATTTGGCCACAGAAGTCTTCTTT | 1 | TRBV13-1*00 | | TRBJ1-1*00 | 0.29% |
| CD4+YFP- WT1 | TGTGCCAGCAGTGCAGGGGGCGAAAGATTATTTTTC | 1 | TRBV13-3*00 | TRBD1*00 | TRBJ1-4*00 | 0.29% |
| CD4+YFP- WT1 | TGTGCCAGCAGTGGACAGGGCCCCATGTCCTATGAACAGTACTTC | 1 | TRBV13-1*00 | TRBD1*00 | TRBJ2-7*00 | 0.29% |
| CD4+YFP- WT1 | TGTGCCAGCAGTGGAGAGGGCCCCATGTCCTATGAACAGTACTTC | 1 | TRBV13-1*00 | TRBD1*00 | TRBJ2-7*00 | 0.29% |
| CD4+YFP- WT1 | TGTGCCAGCAGTGGCAGGGATGCTGAGCAGTTCTTC | 1 | TRBV13-1*00 | TRBD1*00 | TRBJ2-1*00 | 0.29% |
| CD4+YFP- WT1 | TGTGCCAGCAGTGGGACTGGGGGCGGTGCAGAAACGCTGTATTTT | 1 | TRBV14*00 | TRBD2*00 | TRBJ2-3*00 | 0.29% |
| CD4+YFP- WT1 | TGTGCCAGCAGTGGGACTGGGGGTAGTCAAAACACCTTGTACTTT | 1 | TRBV13-3*00 | TRBD2*00 | TRBJ2-4*00 | 0.29% |
| CD4+YFP- WT1 | TGTGCCAGCAGTGGGGACAAACAGGCTCCGCTTTTT | 1 | TRBV13-3*00 | | TRBJ1-5*00 | 0.29% |
| CD4+YFP- WT1 | TGTGCCAGCAGTGGGGAGCAAGACACCCAGTACTTT | 1 | TRBV13-3*00 | | TRBJ2-5*00 | 0.29% |
| CD4+YFP- WT1 | TGTGCCAGCAGTGGGGGAAACTATGCTGAGCAGTTCTTC | 1 | TRBV13-3*00 | | TRBJ2-1*00 | 0.29% |
| CD4+YFP- WT1 | TGTGCCAGCAGTGTGGGGGATAACTATGCTGAGCAGTTCTTC | 1 | TRBV13-3*00 | TRBD2*00 | TRBJ2-1*00 | 0.29% |
| CD4+YFP- WT1 | TGTGCCAGCAGTTACAGGGCGAGAGCTGAGCAGTTCTTC | 1 | TRBV19*00 | TRBD1*00 | TRBJ2-1*00 | 0.29% |
| CD4+YFP- WT1 | TGTGCCAGCAGTTCACAGGGGGCCGGAGAACAGTACTTC | 1 | TRBV15*00 | TRBD1*00 | TRBJ2-7*00 | 0.29% |
| CD4+YFP- WT1 | TGTGCCAGCAGTTTAGAGACTGGTGCAGAAACGCTGTATTTT | 1 | TRBV15*00 | | TRBJ2-3*00 | 0.29% |
| CD4+YFP- WT1 | TGTGCCAGCAGTTTAGCAGGAGCAAACACCGGGCAGCTCTACTTT | 1 | TRBV15*00 | | TRBJ2-2*00 | 0.29% |
| CD4+YFP- WT1 | TGTGCCAGCAGTTTAGGGGGACAGGAAAACACCTTGTACTTT | 1 | TRBV15*00 | TRBD1*00 | TRBJ2-4*00 | 0.29% |
| CD4+YFP- WT1 | TGTGCCAGCAGTTTCGGACAGTATGAACAGTACTTC | 1 | TRBV14*00 | TRBD1*00 | TRBJ2-7*00 | 0.29% |
| CD4+YFP- WT1 | TGTGCCAGCAGTTTCGGGGCCAACCAGGCTCCGCTTTTT | 1 | TRBV15*00 | TRBD1*00 | TRBJ1-5*00 | 0.29% |
| CD4+YFP- WT1 | TGTGCCAGCAGTTTGAGGGGAGATGAACAGTACTTC | 1 | TRBV15*00 | TRBD1*00 | TRBJ2-7*00 | 0.29% |
| CD4+YFP- WT1 | TGTGCCAGCAGTTTTCCAGGGGGCTATGCTGAGCAGTTCTTC | 1 | TRBV14*00 | TRBD1*00 | TRBJ2-1*00 | 0.29% |
| CD4+YFP- WT1 | TGTGCCAGCAGTTTTTCGACCTCAGAAGTCTTCTTT | 1 | TRBV14*00 | | TRBJ1-1*00 | 0.29% |
| CD4+YFP- WT1 | TGTGCCAGCAGTTTTTCTGACAACCAGGCTCCGCTTTTT | 1 | TRBV15*00 | | TRBJ1-5*00 | 0.29% |
| CD4+YFP- WT1 | TGTGCCAGCGCCCCGGGGCAGGCGGAAGTCTTCTTT | 1 | TRBV13-2*00 | TRBD1*00 | TRBJ1-1*00 | 0.29% |
| CD4+YFP- WT1 | TGTGCCAGCGGACCAGGGAATAGTGCAGAAACGCTGTATTTT | 1 | TRBV13-2*00 | TRBD1*00 | TRBJ2-3*00 | 0.29% |
| CD4+YFP- WT1 | TGTGCCAGCGGGACTGGAGCTGAGCAGTTCTTC | 1 | TRBV13-2*00 | TRBD2*00 | TRBJ2-1*00 | 0.29% |
| CD4+YFP- WT1 | TGTGCCAGCGGGGGACTGGGGGGTGACACCCAGTACTTT | 1 | TRBV13-2*00 | TRBD2*00 | TRBJ2-5*00 | 0.29% |
| CD4+YFP- WT1 | TGTGCCAGCGGGGGGGACTGGGGTTATAACTATGCTGAGCAGTTCTTC | 1 | TRBV13-2*00 | TRBD2*00 | TRBJ2-1*00 | 0.29% |
| CD4+YFP- WT1 | TGTGCCAGCGGTGAAAACACCGGGCAGCTCTACTTT | 1 | TRBV13-2*00 | | TRBJ2-2*00 | 0.29% |
| CD4+YFP- WT1 | TGTGCCAGCGGTGACTGGGGGGGGTCCTATGAACAGTACTTC | 1 | TRBV13-2*00 | TRBD2*00 | TRBJ2-7*00 | 0.29% |
| CD4+YFP- WT1 | TGTGCCAGCGGTGAGGCGGGCCAAGACACCCAGTACTTT | 1 | TRBV13-2*00 | | TRBJ2-5*00 | 0.29% |
| CD4+YFP- WT1 | TGTGCCAGCGGTGATAGACTGGGGGGGCGCGCTGAGCAGTTCTTC | 1 | TRBV13-2*00 | TRBD2*00 | TRBJ2-1*00 | 0.29% |
| CD4+YFP- WT1 | TGTGCCAGCGGTGATCCAATATATGAACAGTACTTC | 1 | TRBV13-2*00 | | TRBJ2-7*00 | 0.29% |
| CD4+YFP- WT1 | TGTGCCAGCGGTGATCCGGACAGGCTCCAAGACACCCAGTACTTT | 1 | TRBV13-2*00 | TRBD1*00 | TRBJ2-5*00 | 0.29% |
| CD4+YFP- WT1 | TGTGCCAGCGGTGATCGGGGGGCGCACCATGAACAGTACTTC | 1 | TRBV13-2*00 | TRBD2*00 | TRBJ2-7*00 | 0.29% |
| CD4+YFP- WT1 | TGTGCCAGCGGTGATCTGGGGGCAGAAACGCTGTATTTT | 1 | TRBV13-2*00 | TRBD2*00 | TRBJ2-3*00 | 0.29% |
| CD4+YFP- WT1 | TGTGCCAGCGGTGATGACGGGGGATCCTATGAACAGTACTTC | 1 | TRBV13-2*00 | TRBD1*00 | TRBJ2-7*00 | 0.29% |
| CD4+YFP- WT1 | TGTGCCAGCGGTGATGCAATCGGGGACCAAGACACCCAGTACTTT | 1 | TRBV13-2*00 | | TRBJ2-5*00 | 0.29% |
| CD4+YFP- WT1 | TGTGCCAGCGGTGATGCAGACTCCGACTACACCTTC | 1 | TRBV13-2*00 | | TRBJ1-2*00 | 0.29% |
| CD4+YFP- WT1 | TGTGCCAGCGGTGATGCGGGCAGTAGTGCAGAAACGCTGTATTTT | 1 | TRBV13-2*00 | | TRBJ2-3*00 | 0.29% |
| CD4+YFP- WT1 | TGTGCCAGCGGTGATGCTGGGGGGAGCTCCTATGAACAGTACTTC | 1 | TRBV13-2*00 | TRBD2*00 | TRBJ2-7*00 | 0.29% |
| CD4+YFP- WT1 | TGTGCCAGCGGTGATGGCCAAGACACCCAGTACTTT | 1 | TRBV13-2*00 | | TRBJ2-5*00 | 0.29% |
| CD4+YFP- WT1 | TGTGCCAGCGGTGATGGGAATTCTGGAAATACGCTCTATTTT | 1 | TRBV13-2*00 | | TRBJ1-3*00 | 0.29% |
| CD4+YFP- WT1 | TGTGCCAGCGGTGATGGGACAAAAGACACCCAGTACTTT | 1 | TRBV13-2*00 | TRBD1*00 | TRBJ2-5*00 | 0.29% |
| CD4+YFP- WT1 | TGTGCCAGCGGTGATGGGACAGGGCAAGACACCCAGTACTTT | 1 | TRBV13-2*00 | TRBD1*00 | TRBJ2-5*00 | 0.29% |
| CD4+YFP- WT1 | TGTGCCAGCGGTGCATTTAACCAAGACACCCAGTACTTT | 1 | TRBV13-2*00 | | TRBJ2-5*00 | 0.29% |
| CD4+YFP- WT1 | TGTGCCAGCGGTGGACTGGGGGGATACTATGCTGAGCAGTTCTTC | 1 | TRBV13-2*00 | TRBD2*00 | TRBJ2-1*00 | 0.29% |
| CD4+YFP- WT1 | TGTGCCAGCGGTGGAGAAGTCTTCTTT | 1 | TRBV13-2*00 | | TRBJ1-1*00 | 0.29% |
| CD4+YFP- WT1 | TGTGCCAGCGGTGGGGACAGGAGAAACACCTTGTACTTT | 1 | TRBV13-2*00 | TRBD1*00 | TRBJ2-4*00 | 0.29% |
| CD4+YFP- WT1 | TGTGCCAGCTCGCGGGACAGGGAGGGCTCCGACTACACCTTC | 1 | TRBV12-1*00 | TRBD1*00 | TRBJ1-2*00 | 0.29% |
| CD4+YFP- WT1 | TGTGCCAGCTCTAAAACTAGTGCAGAAACGCTGTATTTT | 1 | TRBV19*00 | | TRBJ2-3*00 | 0.29% |
| CD4+YFP- WT1 | TGTGCCAGCTCTCCCGGGGACAGTGCAGAAACGCTGTATTTT | 1 | TRBV12-1*00 | TRBD1*00 | TRBJ2-3*00 | 0.29% |
| CD4+YFP- WT1 | TGTGCCAGCTCTCTAACTGGGGGGGTCTATGCTGAGCAGTTCTTC | 1 | TRBV12-1*00 | TRBD2*00 | TRBJ2-1*00 | 0.29% |
| CD4+YFP- WT1 | TGTGCCAGCTCTCTCAGGGGAGAAGAAACGCTGTATTTT | 1 | TRBV12-1*00 | TRBD1*00 | TRBJ2-3*00 | 0.29% |
| CD4+YFP- WT1 | TGTGCCAGCTCTCTCCAGATTAACCAAGACACCCAGTACTTT | 1 | TRBV12-1*00 | | TRBJ2-5*00 | 0.29% |
| CD4+YFP- WT1 | TGTGCCAGCTCTCTCCAGGGGGCAAACTCCGACTACACCTTC | 1 | TRBV12-1*00 | TRBD1*00 | TRBJ1-2*00 | 0.29% |
| CD4+YFP- WT1 | TGTGCCAGCTCTCTCGCTGGGGCCCACTATGTTGAGCAGTTCTTC | 1 | TRBV12-1*00 | TRBD2*00 | TRBJ2-1*00 | 0.29% |
| CD4+YFP- WT1 | TGTGCCTGGACAGGGGAGGTGGACACCCAGTACTTT | 1 | TRBV31*00 | TRBD1*00 | TRBJ2-5*00 | 0.29% |
| CD4+YFP- WT1 | TGTGCCTGGACGGAGGGGGGCGAACAGTACTTC | 1 | TRBV31*00 | TRBD2*00 | TRBJ2-7*00 | 0.29% |
| CD4+YFP- WT1 | TGTGCCTGGAGCCCAGGGACTGGGGGTGCTGAGCAGTTCTTC | 1 | TRBV31*00 | TRBD2*00 | TRBJ2-1*00 | 0.29% |
| CD4+YFP- WT1 | TGTGCCTGGAGGACAGGGGTTAGTCCAGAAACGCTGTATTTT | 1 | TRBV31*00 | TRBD1*00 | TRBJ2-3*00 | 0.29% |
| CD4+YFP- WT1 | TGTGCCTGGAGTACAGAGGACACCCAGTACTTT | 1 | TRBV31*00 | | TRBJ2-5*00 | 0.29% |
| CD4+YFP- WT1 | TGTGCCTGGAGTAGGGCAACTAGTGCAGAAACGCTGTATTTT | 1 | TRBV31*00 | | TRBJ2-3*00 | 0.29% |
| CD4+YFP- WT1 | TGTGCCTGGAGTCCCGACAGGGAACTCCGACTACACCTTC | 1 | TRBV31*00 | TRBD1*00 | TRBJ1-2*00 | 0.29% |

| | | | | | | |
|---|---|---|---|---|---|---|
| CD4+YFP- WT1 | TGTGCCTGGAGTCCCGGGACTGGGGGCTATGAACAGTACTTC | 1 | TRBV31*00 | TRBD2*00 | TRBJ2-7*00 | 0.29% |
| CD4+YFP- WT1 | TGTGCCTGGAGTCGACTGGGGGGGCGAGACACCCAGTACTTT | 1 | TRBV31*00 | TRBD2*00 | TRBJ2-5*00 | 0.29% |
| CD4+YFP- WT1 | TGTGCCTGGAGTCTAGAGGACAGAAACTCCGACTACACCTTC | 1 | TRBV31*00 | TRBD1*00 | TRBJ1-2*00 | 0.29% |
| CD4+YFP- WT1 | TGTGCCTGGAGTCTAGGGAATAACTATGCTGAGCAGTTCTTC | 1 | TRBV31*00 | | TRBJ2-1*00 | 0.29% |
| CD4+YFP- WT1 | TGTGCCTGGAGTCTAGGTGAACAGTACTTC | 1 | TRBV31*00 | | TRBJ2-7*00 | 0.29% |
| CD4+YFP- WT1 | TGTGCCTGGAGTCTCCGCCCATCTAGTGCAGAAACGCTGTATTTT | 1 | TRBV31*00 | TRBD1*00 | TRBJ2-3*00 | 0.29% |
| CD4+YFP- WT1 | TGTGCCTGGAGTCTCCGGGGGGCGGATGCTGAGCAGTTCTTC | 1 | TRBV31*00 | TRBD2*00 | TRBJ2-1*00 | 0.29% |
| CD4+YFP- WT1 | TGTGCCTGGAGTCTGGGGGGAGACACCCAGTACTTT | 1 | TRBV31*00 | TRBD2*00 | TRBJ2-5*00 | 0.29% |
| CD4+YFP- WT1 | TGTGCCTGGAGTCTTAAGGGGGCCTCCGACTACACCTTC | 1 | TRBV31*00 | TRBD1*00 | TRBJ1-2*00 | 0.29% |
| CD4+YFP- WT1 | TGTGCCTGGGGACTGGGTAGTGCAGAAACGCTGTATTTT | 1 | TRBV31*00 | TRBD2*00 | TRBJ2-3*00 | 0.29% |
| CD4+YFP- WT1 | TGTGCCTGGGGAGTAAACACCGGGCAGCTCTACTTT | 1 | TRBV31*00 | | TRBJ2-2*00 | 0.29% |
| CD4+YFP- WT1 | TGTGCTAGCAGTAGAGAAAACCAGGCTCCGCTTTTT | 1 | TRBV17*00 | | TRBJ1-5*00 | 0.29% |
| CD4+YFP- WT1 | TGTGCTAGCAGTAGAGATCCGAGTAGTGCAGAAACGCTGTATTTT | 1 | TRBV17*00 | | TRBJ2-3*00 | 0.29% |
| CD4+YFP- WT1 | TGTGCTAGCAGTAGAGGGACTGAACAGTACTTC | 1 | TRBV17*00 | | TRBJ2-7*00 | 0.29% |
| CD4+YFP- WT1 | TGTGCTAGCAGTCTCCGGGACAGGGGCTCCGACTACACCTTC | 1 | TRBV21*00 | TRBD1*00 | TRBJ1-2*00 | 0.29% |
| CD4+YFP- WT1 | TGTGCTAGCAGTGACACTTCTAGTCAAAACACCTTGTACTTT | 1 | TRBV17*00 | | TRBJ2-4*00 | 0.29% |
| CD4+YFP- WT1 | TGTGCTAGCAGTTTAAACAGTGAACAGTACTTC | 1 | TRBV29*00 | | TRBJ2-7*00 | 0.29% |
| CD4+YFP- WT1 | TGTGCTAGCAGTTTAGAGGTGAACACCGGGCAGCTCTACTTT | 1 | TRBV29*00 | | TRBJ2-2*00 | 0.29% |
| CD4+YFP- WT1 | TGTGCTAGCAGTTTAGGGCAGTCCCCTTCTGGAAATACGCTCTATTTT | 1 | TRBV29*00 | TRBD2*00 | TRBJ1-3*00 | 0.29% |
| CD4+YFP- WT1 | TGTGCTAGCAGTTTAGGGGACTATGAACAGTACTTC | 1 | TRBV29*00 | | TRBJ2-7*00 | 0.29% |
| CD4+YFP- WT1 | TGTGGTGCTAAACAGGGAGATGACACCCAGTACTTT | 1 | TRBV20*00 | TRBD1*00 | TRBJ2-5*00 | 0.29% |
| CD4+YFP- WT1 | TGTGGTGCTACCACAAACTCCGACTACACCTTC | 1 | TRBV20*00 | | TRBJ1-2*00 | 0.29% |
| CD4+YFP- WT1 | TGTGGTGCTAGCCGGGACATGAACTATGCTGAGCAGTTCTTC | 1 | TRBV20*00 | TRBD1*00 | TRBJ2-1*00 | 0.29% |
| CD4+YFP- WT1 | TGTGGTGCTAGGGAAGGGAGGGGAAACTCCGACTACACCTTC | 1 | TRBV20*00 | TRBD1*00 | TRBJ1-2*00 | 0.29% |
| CD4+YFP- WT1 | TGTGGTGCTAGGGAGAGGAGAGTCTTCTTT | 1 | TRBV20*00 | | TRBJ1-1*00 | 0.29% |
| CD4+YFP- WT1 | TGTGGTGCTAGGGAGGGCAATGCAGAAACGCTGTATTTT | 1 | TRBV20*00 | | TRBJ2-3*00 | 0.29% |
| CD4+YFP- WT1 | TGTGGTGCTAGGGCGGCAGGGGAGGACACCCAGTACTTT | 1 | TRBV20*00 | TRBD1*00 | TRBJ2-5*00 | 0.29% |
| CD4+YFP- WT1 | TGTGGTGCTAGGTGGGGAGACACCCAGTACTTT | 1 | TRBV20*00 | TRBD2*00 | TRBJ2-5*00 | 0.29% |
| CD4+YFP- WT1 | TGTGGTGCTCGGGACATTAATTCGCCCCTCTACTTT | 1 | TRBV20*00 | | TRBJ1-6*00 | 0.29% |
| CD4+YFP- WT1 | TGTGGTGCTGGAAGACTGGGGGGGCGAGCTGAGCAGTTCTTC | 1 | TRBV20*00 | TRBD2*00 | TRBJ2-1*00 | 0.29% |
| CD4+YFP- WT1 | TGTGGTGCTTCCCGGGACACTAACTATGCTGAGCAGTTCTTC | 1 | TRBV20*00 | TRBD1*00 | TRBJ2-1*00 | 0.29% |
| CD4+YFP- WT1 | TGTGTAAGCAGCTTAGAGGGGGGGGACACAGAAGTCTTCTTT | 1 | TRBV16*00 | TRBD2*00 | TRBJ1-1*00 | 0.29% |
| CD4+YFP- WT1 | TGTGCCAGCAGTTTGCGGGGGGATGCAGAAACGCTGTATTTT | 1 | TRBV14*00 | TRBD2*00 | TRBJ2-3*00 | 0.29% |
| CD4+YFP- WT1 | TGTGCCAGCAGCCAAGACTGGGGGGGCTATGCTGAGCAGTTCTTC | 1 | TRBV5*00 | TRBD2*00 | TRBJ2-1*00 | 0.29% |
| CD4+YFP- WT1 | TGTGCCAGCGGTGATCTGGGGGGGCAAGACACCCAGTACTTT | 1 | TRBV13-2*00 | TRBD2*00 | TRBJ2-5*00 | 0.29% |
| CD4+YFP- WT1 | TGTGCCAGCAGTATAGACAAAAACACCTTGTACTTT | 1 | TRBV19*00 | | TRBJ2-4*00 | 0.29% |
| CD4+YFP- WT1 | TGTGCTAGCAGTTTATCGTCGCCAAACACAGAAGTCTTCTTT | 1 | TRBV29*00 | | TRBJ1-1*00 | 0.29% |

#### CD4+YFP- WT2

| Sample Name | CDR3 Sequence | clone Count | best V Hit | best D Hit | best J Hit | Frequency |
|---|---|---|---|---|---|---|
| CD4+YFP- WT2 | TGTGCCAGCAGCCGCGGGACAGAAGTCTTCTTT | 3 | TRBV14*00 | | TRBJ1-1*00 | 0.90% |
| CD4+YFP- WT2 | TGTGCCAGCAGTGATCTGGGGGAAAACACCTTGTACTTT | 3 | TRBV13-3*00 | TRBD2*00 | TRBJ2-4*00 | 0.90% |
| CD4+YFP- WT2 | TGCTCCAGCAGTCAAGACAGGAATTCGCCCCTCTACTTT | 2 | TRBV23*00 | TRBD1*00 | TRBJ1-6*00 | 0.60% |
| CD4+YFP- WT2 | TGTGCAAGCAGCCTGGACTGGGGGGGCGGTGAACAGTACTTC | 2 | TRBV16*00 | TRBD2*00 | TRBJ2-7*00 | 0.60% |
| CD4+YFP- WT2 | TGTGCAAGCAGCTTAGATGGGGGGTATGAACAGTACTTC | 2 | TRBV16*00 | TRBD2*00 | TRBJ2-7*00 | 0.60% |
| CD4+YFP- WT2 | TGTGCCAGCAGCCCATCAGGGGTGGCAGAAGTCTTCTTT | 2 | TRBV5*00 | TRBD1*00 | TRBJ1-1*00 | 0.60% |
| CD4+YFP- WT2 | TGTGCCAGCAGCTACAGGAACCAAGACACCCAGTACTTT | 2 | TRBV4*00 | | TRBJ2-5*00 | 0.60% |
| CD4+YFP- WT2 | TGTGCCAGCAGTGAAGGGCTGGGGGGACTCTATGCTGAGCAGTTCTTC | 2 | TRBV13-1*00 | TRBD2*00 | TRBJ2-1*00 | 0.60% |
| CD4+YFP- WT2 | TGTGCCAGCAGTGATGACGGGGCTAGTGCAGAAACGCTGTATTTT | 2 | TRBV13-1*00 | | TRBJ2-3*00 | 0.60% |
| CD4+YFP- WT2 | TGTGCCAGCAGTGATGAGGACACCCAGTACTTT | 2 | TRBV13-1*00 | | TRBJ2-5*00 | 0.60% |
| CD4+YFP- WT2 | TGTGCCAGCAGTGATGGACAGAACCAGGCTCCGCTTTTT | 2 | TRBV13-1*00 | TRBD1*00 | TRBJ1-5*00 | 0.60% |
| CD4+YFP- WT2 | TGTGCCAGCAGTGCAGAAAACTCCGACTACACCTTC | 2 | TRBV22*00 | | TRBJ1-2*00 | 0.60% |
| CD4+YFP- WT2 | TGTGCCAGCAGTTGGGACAGGGGAGACACCCAGTACTTT | 2 | TRBV22*00 | TRBD1*00 | TRBJ2-5*00 | 0.60% |
| CD4+YFP- WT2 | TGTGCCAGCAGTTTACAGGGAGAGTCCTATGAACAGTACTTC | 2 | TRBV15*00 | TRBD1*00 | TRBJ2-7*00 | 0.60% |
| CD4+YFP- WT2 | TGTGCCAGCGGTGAAGGGAACCAAGACACCCAGTACTTT | 2 | TRBV13-2*00 | | TRBJ2-5*00 | 0.60% |
| CD4+YFP- WT2 | TGTGCCTGGAGTCCGGACTGGGGGGAATATGAACAGTACTTC | 2 | TRBV31*00 | TRBD2*00 | TRBJ2-7*00 | 0.60% |
| CD4+YFP- WT2 | TGTGCCTGGAGTCTAACAAACTCCGACTACACCTTC | 2 | TRBV31*00 | | TRBJ1-2*00 | 0.60% |
| CD4+YFP- WT2 | TGTGGTGCTAGGGGGCCGGGACAGGGGTGACACCCAGTACTTT | 2 | TRBV20*00 | TRBD1*00 | TRBJ2-5*00 | 0.60% |
| CD4+YFP- WT2 | TGCACCTGCAGGACAGGTACTAGTGCAGAAACGCTGTATTTT | 1 | TRBV1*00 | TRBD1*00 | TRBJ2-3*00 | 0.30% |
| CD4+YFP- WT2 | TGCACCTGCAGTCCAGGGGGCTATGAACAGTACTTC | 1 | TRBV1*00 | | TRBJ2-7*00 | 0.30% |
| CD4+YFP- WT2 | TGCACCTGCAGTCCCCGGGACAGGGATGAACAGTACTTC | 1 | TRBV1*00 | TRBD1*00 | TRBJ2-7*00 | 0.30% |
| CD4+YFP- WT2 | TGCACCTGCAGTGAGACAGACTATGCTGAGCAGTTCTTC | 1 | TRBV1*00 | TRBD1*00 | TRBJ2-1*00 | 0.30% |
| CD4+YFP- WT2 | TGCACCTGCAGTGAGTACAGTAACTATGCTGAGCAGTTCTTC | 1 | TRBV1*00 | | TRBJ2-1*00 | 0.30% |
| CD4+YFP- WT2 | TGCACCTGCAGTGCAAACTCCTATGAACAGTACTTC | 1 | TRBV1*00 | | TRBJ2-7*00 | 0.30% |
| CD4+YFP- WT2 | TGCACCTGCAGTGCACAGGGCTCCGACTACACCTTC | 1 | TRBV1*00 | TRBD1*00 | TRBJ1-2*00 | 0.30% |
| CD4+YFP- WT2 | TGCACCTGCAGTGCAGACAGATTCTATGCTGAGCAGTTCTTC | 1 | TRBV1*00 | | TRBJ2-1*00 | 0.30% |
| CD4+YFP- WT2 | TGCACCTGCAGTGCAGAGGCTGGTCAAAACACCTTGTACTTT | 1 | TRBV1*00 | | TRBJ2-4*00 | 0.30% |
| CD4+YFP- WT2 | TGCACCTGCAGTGCAGATCGGACTGGCGTTAACCAAGACACCCAGTACTTT | 1 | TRBV1*00 | TRBD2*00 | TRBJ2-5*00 | 0.30% |
| CD4+YFP- WT2 | TGCACCTGCAGTGCAGATGAACAGTACTTC | 1 | TRBV1*00 | | TRBJ2-7*00 | 0.30% |
| CD4+YFP- WT2 | TGCACCTGCAGTGCAGATGCTTTTAACCAAGACACCCAGTACTTT | 1 | TRBV1*00 | | TRBJ2-5*00 | 0.30% |
| CD4+YFP- WT2 | TGCACCTGCAGTGCAGATGGAGACACCCAGTACTTT | 1 | TRBV1*00 | | TRBJ2-5*00 | 0.30% |
| CD4+YFP- WT2 | TGCACCTGCAGTGCAGATGGTTATGAACAGTACTTC | 1 | TRBV1*00 | | TRBJ2-7*00 | 0.30% |
| CD4+YFP- WT2 | TGCACCTGCAGTGCAGATTCAGAAACGCTGTATTTT | 1 | TRBV1*00 | | TRBJ2-3*00 | 0.30% |
| CD4+YFP- WT2 | TGCACCTGCAGTGCAGCGGGGGGCCAAGACACCCAGTACTTT | 1 | TRBV1*00 | TRBD2*00 | TRBJ2-5*00 | 0.30% |
| CD4+YFP- WT2 | TGCACCTGCAGTGCAGCTGGGTACTCCGACTACACCTTC | 1 | TRBV1*00 | TRBD2*00 | TRBJ1-2*00 | 0.30% |

| | | | | | | |
|---|---|---|---|---|---|---|
| CD4+YFP- WT2 | TGCACCTGCAGTGCAGGAACTGGGGGGAGGGCTGAGCAGTTCTTC | 1 | TRBV1*00 | TRBD2*00 | TRBJ2-1*00 | 0.30% |
| CD4+YFP- WT2 | TGCACCTGCAGTGCAGGGGGGACCGGGCAGCTCTACTTT | 1 | TRBV1*00 | TRBD1*00 | TRBJ2-2*00 | 0.30% |
| CD4+YFP- WT2 | TGCACCTGCAGTGCAGTAGGGGATAGTGCAGAAACGCTGTATTTT | 1 | TRBV1*00 | TRBD1*00 | TRBJ2-3*00 | 0.30% |
| CD4+YFP- WT2 | TGCACCTGCAGTGCCAGACACAATTCTGGAAATACGCTCTATTTT | 1 | TRBV1*00 | | TRBJ1-3*00 | 0.30% |
| CD4+YFP- WT2 | TGCACCTGCAGTGCCCACAGTCAAAACACCTTGTACTTT | 1 | TRBV1*00 | | TRBJ2-4*00 | 0.30% |
| CD4+YFP- WT2 | TGCACCTGCAGTGCGCTGGGGGACCAAGACACCCAGTACTTT | 1 | TRBV1*00 | TRBD2*00 | TRBJ2-5*00 | 0.30% |
| CD4+YFP- WT2 | TGCACCTGCAGTGCTTCCGGAAACTCCGACTACACCTTC | 1 | TRBV1*00 | | TRBJ1-2*00 | 0.30% |
| CD4+YFP- WT2 | TGCACCTGCAGTGGGACAACTAGTGCAGAAACGCTGTATTTT | 1 | TRBV1*00 | TRBD1*00 | TRBJ2-3*00 | 0.30% |
| CD4+YFP- WT2 | TGCACCTGCAGTGGGCGGGACAAAGACACCCAGTACTTT | 1 | TRBV1*00 | TRBD1*00 | TRBJ2-5*00 | 0.30% |
| CD4+YFP- WT2 | TGCTCCAGCAGTTGGGACACTGCAAACTCCGACTACACCTTC | 1 | TRBV23*00 | TRBD1*00 | TRBJ1-2*00 | 0.30% |
| CD4+YFP- WT2 | TGTAGTTCTAACGGGGATTCTGGAAATACGCTCTATTTT | 1 | TRBV30*00 | | TRBJ1-3*00 | 0.30% |
| CD4+YFP- WT2 | TGTAGTTCTAGAGGGACAGCAGAAACGCTGTATTTT | 1 | TRBV30*00 | TRBD1*00 | TRBJ2-3*00 | 0.30% |
| CD4+YFP- WT2 | TGTAGTTCTAGTCGGCCTTATGAACAGTACTTC | 1 | TRBV30*00 | | TRBJ2-7*00 | 0.30% |
| CD4+YFP- WT2 | TGTGCAAGCAGCCCCGAAGGGGACGAAAGATTATTTTTC | 1 | TRBV16*00 | TRBD1*00 | TRBJ1-4*00 | 0.30% |
| CD4+YFP- WT2 | TGTGCAAGCAGCCCCGACTGGGGGGTTGCTGAGCAGTTCTTC | 1 | TRBV16*00 | TRBD2*00 | TRBJ2-1*00 | 0.30% |
| CD4+YFP- WT2 | TGTGCAAGCAGCCCCGGACAGGCAAACACAGAAGTCTTCTTT | 1 | TRBV16*00 | TRBD1*00 | TRBJ1-1*00 | 0.30% |
| CD4+YFP- WT2 | TGTGCAAGCAGCCGGGACTGGGGGCAAGACACCCAGTACTTT | 1 | TRBV16*00 | TRBD2*00 | TRBJ2-5*00 | 0.30% |
| CD4+YFP- WT2 | TGTGCAAGCAGCGGGACAGGAAAAGACACCCAGTACTTT | 1 | TRBV16*00 | TRBD1*00 | TRBJ2-5*00 | 0.30% |
| CD4+YFP- WT2 | TGTGCAAGCAGCGGGGGGGCGCAGAACACCTTGTACTTT | 1 | TRBV16*00 | TRBD2*00 | TRBJ2-4*00 | 0.30% |
| CD4+YFP- WT2 | TGTGCAAGCAGCTCACAGGGGTATGAACAGTACTTC | 1 | TRBV16*00 | TRBD1*00 | TRBJ2-7*00 | 0.30% |
| CD4+YFP- WT2 | TGTGCAAGCAGCTCAGAAGACACCCAGTACTTT | 1 | TRBV16*00 | | TRBJ2-5*00 | 0.30% |
| CD4+YFP- WT2 | TGTGCAAGCAGCTCCGACTGGGGGAGAGCTGAGCAGTTCTTC | 1 | TRBV16*00 | TRBD2*00 | TRBJ2-1*00 | 0.30% |
| CD4+YFP- WT2 | TGTGCAAGCAGCTCCGGGACAGGGGGCGTCGAAACGCTGTATTTT | 1 | TRBV16*00 | TRBD1*00 | TRBJ2-3*00 | 0.30% |
| CD4+YFP- WT2 | TGTGCAAGCAGCTCGCCCGGGACACCTTATGAACAGTACTTC | 1 | TRBV16*00 | TRBD1*00 | TRBJ2-7*00 | 0.30% |
| CD4+YFP- WT2 | TGTGCAAGCAGCTCGGACGGAAACCAAGACACCCAGTACTTT | 1 | TRBV16*00 | | TRBJ2-5*00 | 0.30% |
| CD4+YFP- WT2 | TGTGCAAGCAGCTTACTGGGGGGTGAACAGTACTTC | 1 | TRBV16*00 | TRBD2*00 | TRBJ2-7*00 | 0.30% |
| CD4+YFP- WT2 | TGTGCAAGCAGCTTAGAAGGGGATGAACAGTACTTC | 1 | TRBV16*00 | TRBD1*00 | TRBJ2-7*00 | 0.30% |
| CD4+YFP- WT2 | TGTGCAAGCAGCTTAGATACGGCCACCTTGTACTTT | 1 | TRBV16*00 | | TRBJ2-4*00 | 0.30% |
| CD4+YFP- WT2 | TGTGCAAGCAGCTTAGATAGGGACAGGGGAGACACCCAGTACTTT | 1 | TRBV16*00 | TRBD1*00 | TRBJ2-5*00 | 0.30% |
| CD4+YFP- WT2 | TGTGCAAGCAGCTTAGATCGACAAGACACCCAGTACTTT | 1 | TRBV16*00 | | TRBJ2-5*00 | 0.30% |
| CD4+YFP- WT2 | TGTGCAAGCAGCTTAGATGAGGGTGAACAGTACTTC | 1 | TRBV16*00 | | TRBJ2-7*00 | 0.30% |
| CD4+YFP- WT2 | TGTGCAAGCAGCTTAGATGGGTACTCCGACTACACCTTC | 1 | TRBV16*00 | | TRBJ1-2*00 | 0.30% |
| CD4+YFP- WT2 | TGTGCAAGCAGCTTAGCCGGGAACACAGAAGTCTTCTTT | 1 | TRBV16*00 | TRBD1*00 | TRBJ1-1*00 | 0.30% |
| CD4+YFP- WT2 | TGTGCAAGCAGCTTAGCGGGGAACTATGCTGAGCAGTTCTTC | 1 | TRBV16*00 | | TRBJ2-1*00 | 0.30% |
| CD4+YFP- WT2 | TGTGCAAGCAGCTTAGCTGGATATGAACAGTACTTC | 1 | TRBV16*00 | | TRBJ2-7*00 | 0.30% |
| CD4+YFP- WT2 | TGTGCAAGCAGCTTAGGGACAGCCGCAAACTCCGACTACACCTTC | 1 | TRBV16*00 | TRBD1*00 | TRBJ1-2*00 | 0.30% |
| CD4+YFP- WT2 | TGTGCAAGCAGCTTAGGGGGGAGCTCCTATGAACAGTACTTC | 1 | TRBV16*00 | | TRBJ2-7*00 | 0.30% |
| CD4+YFP- WT2 | TGTGCAAGCAGCTTAGTCGGGAACACAGAAGTCTTCTTT | 1 | TRBV16*00 | | TRBJ1-1*00 | 0.30% |
| CD4+YFP- WT2 | TGTGCAAGCAGCTTAGTGGCAGAAACGCTGTATTTT | 1 | TRBV16*00 | | TRBJ2-3*00 | 0.30% |
| CD4+YFP- WT2 | TGTGCAAGCAGCTTAGTTGGGTACTCCTATGAACAGTACTTC | 1 | TRBV16*00 | | TRBJ2-7*00 | 0.30% |
| CD4+YFP- WT2 | TGTGCAAGCAGTGCAGAAACGCTGTATTTT | 1 | TRBV16*00 | | TRBJ2-3*00 | 0.30% |
| CD4+YFP- WT2 | TGTGCCAGCAAACGGACAGATTCTGGAAATACGCTCTATTTT | 1 | TRBV26*00 | TRBD1*00 | TRBJ1-3*00 | 0.30% |
| CD4+YFP- WT2 | TGTGCCAGCAAGGACAATTCTGGAAATACGCTCTATTTT | 1 | TRBV13-1*00 | | TRBJ1-3*00 | 0.30% |
| CD4+YFP- WT2 | TGTGCCAGCACCCCCGGGGGAAACTCCGACTACACCTTC | 1 | TRBV19*00 | TRBD1*00 | TRBJ1-2*00 | 0.30% |
| CD4+YFP- WT2 | TGTGCCAGCACCCTGGACGGTAACTATGCTGAGCAGTTCTTC | 1 | TRBV15*00 | TRBD1*00 | TRBJ2-1*00 | 0.30% |
| CD4+YFP- WT2 | TGTGCCAGCAGAACCGGGACAGACGCAGAAGTCTTCTTT | 1 | TRBV15*00 | TRBD1*00 | TRBJ1-1*00 | 0.30% |
| CD4+YFP- WT2 | TGTGCCAGCAGCAGGGGGCTGGCCAACGAAAGATTATTTTTC | 1 | TRBV4*00 | TRBD1*00 | TRBJ1-4*00 | 0.30% |
| CD4+YFP- WT2 | TGTGCCAGCAGCCAAAAAACCGGGACACATAGTGCAGAAACGCTGTATTTT | 1 | TRBV5*00 | TRBD1*00 | TRBJ2-3*00 | 0.30% |
| CD4+YFP- WT2 | TGTGCCAGCAGCCAACTGGGGGGGCGGGAACAGTACTTC | 1 | TRBV2*00 | TRBD2*00 | TRBJ2-7*00 | 0.30% |
| CD4+YFP- WT2 | TGTGCCAGCAGCCAAGAAACAGACAACCAGGCTCCGCTTTTT | 1 | TRBV5*00 | | TRBJ1-5*00 | 0.30% |
| CD4+YFP- WT2 | TGTGCCAGCAGCCAAGACAGCTCCTATGAACAGTACTTC | 1 | TRBV5*00 | | TRBJ2-7*00 | 0.30% |
| CD4+YFP- WT2 | TGTGCCAGCAGCCAAGACGACTGGGGGAACTATGCTGAGCAGTTCTTC | 1 | TRBV2*00 | TRBD2*00 | TRBJ2-1*00 | 0.30% |
| CD4+YFP- WT2 | TGTGCCAGCAGCCAAGAGGGGACAGAAGTCTTCTTT | 1 | TRBV5*00 | | TRBJ1-1*00 | 0.30% |
| CD4+YFP- WT2 | TGTGCCAGCAGCCAAGAGGGGGGGCGAGAAGACACCCAGTACTTA | 1 | TRBV5*00 | TRBD2*00 | TRBJ2-5*00 | 0.30% |
| CD4+YFP- WT2 | TGTGCCAGCAGCCAAGATAACACCAATAACTATGCTGAGCAGTTCTTC | 1 | TRBV5*00 | | TRBJ2-1*00 | 0.30% |
| CD4+YFP- WT2 | TGTGCCAGCAGCCAAGATAACTCCGACTACACCTTC | 1 | TRBV2*00 | | TRBJ1-2*00 | 0.30% |
| CD4+YFP- WT2 | TGTGCCAGCAGCCAAGATCCCGGAAATACGCTCTATTTT | 1 | TRBV5*00 | | TRBJ1-3*00 | 0.30% |
| CD4+YFP- WT2 | TGTGCCAGCAGCCAAGATTTACAGCAGGCTCCGCTTTTT | 1 | TRBV5*00 | | TRBJ1-5*00 | 0.30% |
| CD4+YFP- WT2 | TGTGCCAGCAGCCAAGCAGGGGTTGAACAGTACTTC | 1 | TRBV5*00 | TRBD1*00 | TRBJ2-7*00 | 0.30% |
| CD4+YFP- WT2 | TGTGCCAGCAGCCAAGGCGGACACTCCAACGAAAGATTATTTTTC | 1 | TRBV5*00 | TRBD1*00 | TRBJ1-4*00 | 0.30% |
| CD4+YFP- WT2 | TGTGCCAGCAGCCAAGGGGGGGAGCTCCTATGAACAGTACTTC | 1 | TRBV2*00 | | TRBJ2-7*00 | 0.30% |
| CD4+YFP- WT2 | TGTGCCAGCAGCCAAGGGGGGGGGAGGAGACACCCAGTACTTT | 1 | TRBV2*00 | TRBD2*00 | TRBJ2-5*00 | 0.30% |
| CD4+YFP- WT2 | TGTGCCAGCAGCCAAGTGGGGGACAAAAATGCTGAGCAGTTCTTC | 1 | TRBV2*00 | TRBD1*00 | TRBJ2-1*00 | 0.30% |
| CD4+YFP- WT2 | TGTGCCAGCAGCCACAGGGGAGACACCCAGTACTTT | 1 | TRBV4*00 | TRBD1*00 | TRBJ2-5*00 | 0.30% |
| CD4+YFP- WT2 | TGTGCCAGCAGCCACAGGGGTGGCACCCAGTACTTT | 1 | TRBV4*00 | TRBD1*00 | TRBJ2-5*00 | 0.30% |
| CD4+YFP- WT2 | TGTGCCAGCAGCCAGGGACGGACCTATGAACAGTACTTC | 1 | TRBV5*00 | TRBD1*00 | TRBJ2-7*00 | 0.30% |
| CD4+YFP- WT2 | TGTGCCAGCAGCCCCGCTTGGGGGGGTGCAGAAACGCTGTATTTT | 1 | TRBV26*00 | TRBD2*00 | TRBJ2-3*00 | 0.30% |
| CD4+YFP- WT2 | TGTGCCAGCAGCCCGCAGAATTCTGGAAATACGCTCTATTTT | 1 | TRBV3*00 | | TRBJ1-3*00 | 0.30% |
| CD4+YFP- WT2 | TGTGCCAGCAGCCCGGGGGCGGGCACCGGGCAGCTCTACTTT | 1 | TRBV2*00 | TRBD1*00 | TRBJ2-2*00 | 0.30% |
| CD4+YFP- WT2 | TGTGCCAGCAGCCCTGACAATAACTCCGACTACACCTTC | 1 | TRBV2*00 | | TRBJ1-2*00 | 0.30% |
| CD4+YFP- WT2 | TGTGCCAGCAGCCCTTACTGGGGGGAAAACACCTTGTACTTT | 1 | TRBV4*00 | TRBD2*00 | TRBJ2-4*00 | 0.30% |
| CD4+YFP- WT2 | TGTGCCAGCAGCCGACAGAGTGCAGAAACGCTGTATTTT | 1 | TRBV3*00 | TRBD1*00 | TRBJ2-3*00 | 0.30% |
| CD4+YFP- WT2 | TGTGCCAGCAGCCGACGGGCAAACACAGAAGTCTTCTTT | 1 | TRBV2*00 | | TRBJ1-1*00 | 0.30% |
| CD4+YFP- WT2 | TGTGCCAGCAGCCGGGACAGAAACACCTTGTACTTT | 1 | TRBV4*00 | TRBD1*00 | TRBJ2-4*00 | 0.30% |
| CD4+YFP- WT2 | TGTGCCAGCAGCCGGGACAGGGACACAGAAGTCTTCTTT | 1 | TRBV2*00 | TRBD1*00 | TRBJ1-1*00 | 0.30% |

| | | | | | | |
|---|---|---|---|---|---|---|
| CD4+YFP- WT2 | TGTGCCAGCAGCCGGGACTGGGGGGAAGATGCTGAGCAGTTCTTC | 1 | TRBV4*00 | TRBD2*00 | TRBJ2-1*00 | 0.30% |
| CD4+YFP- WT2 | TGTGCCAGCAGCCGGGGACTGGGGGGAAGTGCAGAAACGCTGTATTTT | 1 | TRBV5*00 | TRBD2*00 | TRBJ2-3*00 | 0.30% |
| CD4+YFP- WT2 | TGTGCCAGCAGCCTACAGGGGAATACGCTCTATTTT | 1 | TRBV2*00 | TRBD1*00 | TRBJ1-3*00 | 0.30% |
| CD4+YFP- WT2 | TGTGCCAGCAGCCTCGACAGAAACTCCGACTACACCTTC | 1 | TRBV25*00 | TRBD1*00 | TRBJ1-2*00 | 0.30% |
| CD4+YFP- WT2 | TGTGCCAGCAGCCTCGGGATTAGTGCAGAAACGCTGTATTTT | 1 | TRBV4*00 | TRBD1*00 | TRBJ2-3*00 | 0.30% |
| CD4+YFP- WT2 | TGTGCCAGCAGCCTCGGGGGCGGAGCTGAGCAGTTCTTC | 1 | TRBV14*00 | TRBD1*00 | TRBJ2-1*00 | 0.30% |
| CD4+YFP- WT2 | TGTGCCAGCAGCCTCTCCGGGACAACAAACACCGGGCAGCTCTACTTT | 1 | TRBV5*00 | TRBD1*00 | TRBJ2-2*00 | 0.30% |
| CD4+YFP- WT2 | TGTGCCAGCAGCCTGACAACTAGTGCAGAAACGCTGTATTTT | 1 | TRBV13-1*00 | | TRBJ2-3*00 | 0.30% |
| CD4+YFP- WT2 | TGTGCCAGCAGCCTGACTGGGGGGGGTGAACAGTACTTC | 1 | TRBV2*00 | TRBD2*00 | TRBJ2-7*00 | 0.30% |
| CD4+YFP- WT2 | TGTGCCAGCAGCGAGACAGGGCAGGAACAGTACTTC | 1 | TRBV5*00 | TRBD1*00 | TRBJ2-7*00 | 0.30% |
| CD4+YFP- WT2 | TGTGCCAGCAGCGGGACTGGGGTTGCTGAGCAGTTCTTC | 1 | TRBV26*00 | TRBD2*00 | TRBJ2-1*00 | 0.30% |
| CD4+YFP- WT2 | TGTGCCAGCAGCGTCAGGGGAAGCCAGGCTCCGCTTTTT | 1 | TRBV4*00 | TRBD1*00 | TRBJ1-5*00 | 0.30% |
| CD4+YFP- WT2 | TGTGCCAGCAGCTACGCAGGAGACACCCAGTACTTT | 1 | TRBV4*00 | | TRBJ2-5*00 | 0.30% |
| CD4+YFP- WT2 | TGTGCCAGCAGCTCTGGGGGGTCCCAAGACACCCAGTACTTT | 1 | TRBV3*00 | TRBD2*00 | TRBJ2-5*00 | 0.30% |
| CD4+YFP- WT2 | TGTGCCAGCAGCTGGACTGGGTCAAACACCGGGCAGCTCTACTTT | 1 | TRBV4*00 | TRBD2*00 | TRBJ2-2*00 | 0.30% |
| CD4+YFP- WT2 | TGTGCCAGCAGCTTAAATTCTCTATAATTCGCCCCTCTACTTT | 1 | TRBV4*00 | | TRBJ1-6*00 | 0.30% |
| CD4+YFP- WT2 | TGTGCCAGCAGCTTAACCCCGGGACAGGATTCTGGAAATACGCTCTATTTT | 1 | TRBV3*00 | TRBD1*00 | TRBJ1-3*00 | 0.30% |
| CD4+YFP- WT2 | TGTGCCAGCAGCTTAAGCTGGGCTAGTGCAGAAACGCTGTATTTT | 1 | TRBV3*00 | TRBD2*00 | TRBJ2-3*00 | 0.30% |
| CD4+YFP- WT2 | TGTGCCAGCAGCTTAAGGGGTTCTGGAAATACGCTCTATTTT | 1 | TRBV3*00 | TRBD1*00 | TRBJ1-3*00 | 0.30% |
| CD4+YFP- WT2 | TGTGCCAGCAGCTTACAAAACCAAGACACCCAGTACTTT | 1 | TRBV3*00 | | TRBJ2-5*00 | 0.30% |
| CD4+YFP- WT2 | TGTGCCAGCAGCTTACAGAATTCTGGAAATACGCTCTATTTT | 1 | TRBV4*00 | | TRBJ1-3*00 | 0.30% |
| CD4+YFP- WT2 | TGTGCCAGCAGCTTACAGGGTGACGAACAGTACTTC | 1 | TRBV3*00 | TRBD1*00 | TRBJ2-7*00 | 0.30% |
| CD4+YFP- WT2 | TGTGCCAGCAGCTTAGAAAACAACCAGGCTCCGCTTTTT | 1 | TRBV3*00 | | TRBJ1-5*00 | 0.30% |
| CD4+YFP- WT2 | TGTGCCAGCAGCTTAGAACAGGCTAGTGCAGAAACGCTGTATTTT | 1 | TRBV3*00 | TRBD1*00 | TRBJ2-3*00 | 0.30% |
| CD4+YFP- WT2 | TGTGCCAGCAGCTTAGAATAGTCAAAACACCTTGTACTTT | 1 | TRBV3*00 | | TRBJ2-4*00 | 0.30% |
| CD4+YFP- WT2 | TGTGCCAGCAGCTTAGAGACAGGGGATGCTGAGCAGTTCTTC | 1 | TRBV3*00 | TRBD1*00 | TRBJ2-1*00 | 0.30% |
| CD4+YFP- WT2 | TGTGCCAGCAGCTTAGAGCAGAAAGACACCCAGTACTTT | 1 | TRBV25*00 | | TRBJ2-5*00 | 0.30% |
| CD4+YFP- WT2 | TGTGCCAGCAGCTTAGAGGGACAGGGAAGCACCGGGCAGCTCTACTTT | 1 | TRBV3*00 | TRBD1*00 | TRBJ2-2*00 | 0.30% |
| CD4+YFP- WT2 | TGTGCCAGCAGCTTAGATAACTCCGACTACACCTTC | 1 | TRBV3*00 | | TRBJ1-2*00 | 0.30% |
| CD4+YFP- WT2 | TGTGCCAGCAGCTTAGCCAGTTCAAACTCCGACTACACCTTC | 1 | TRBV3*00 | | TRBJ1-2*00 | 0.30% |
| CD4+YFP- WT2 | TGTGCCAGCAGCTTAGGACTGGGGGTTAACTATGCTGAGCAGTTCTTC | 1 | TRBV3*00 | TRBD2*00 | TRBJ2-1*00 | 0.30% |
| CD4+YFP- WT2 | TGTGCCAGCAGCTTAGGGACTGGGGGGGGCGGACACCCAGTACTTT | 1 | TRBV3*00 | TRBD2*00 | TRBJ2-5*00 | 0.30% |
| CD4+YFP- WT2 | TGTGCCAGCAGCTTAGTCAGTGCAGAAACGCTGTATTTT | 1 | TRBV3*00 | | TRBJ2-3*00 | 0.30% |
| CD4+YFP- WT2 | TGTGCCAGCAGCTTATCACCTAGTGCAGAAACGCTGTATTTT | 1 | TRBV3*00 | | TRBJ2-3*00 | 0.30% |
| CD4+YFP- WT2 | TGTGCCAGCAGCTTCAGGGGCTCCGACTACACCTTC | 1 | TRBV4*00 | TRBD1*00 | TRBJ1-2*00 | 0.30% |
| CD4+YFP- WT2 | TGTGCCAGCAGCTTGAGGGGCCTTTCTGGAAATACGCTCTATTTT | 1 | TRBV3*00 | TRBD1*00 | TRBJ1-3*00 | 0.30% |
| CD4+YFP- WT2 | TGTGCCAGCAGCTTTGGGGGGGGCGACAGAAACGCTGTATTTT | 1 | TRBV3*00 | TRBD2*00 | TRBJ2-3*00 | 0.30% |
| CD4+YFP- WT2 | TGTGCCAGCAGCTTTTCCGGGGCTGAGCAGTTCTTC | 1 | TRBV3*00 | TRBD1*00 | TRBJ2-1*00 | 0.30% |
| CD4+YFP- WT2 | TGTGCCAGCAGGACCGGGACAGGGGACGAAAGATTATTTTTC | 1 | TRBV15*00 | TRBD1*00 | TRBJ1-4*00 | 0.30% |
| CD4+YFP- WT2 | TGTGCCAGCAGGCAGGGAGACACCGGGCAGCTCTACTTT | 1 | TRBV15*00 | TRBD1*00 | TRBJ2-2*00 | 0.30% |
| CD4+YFP- WT2 | TGTGCCAGCAGGCCGGGACAGGGGGCTGAGCAGTTCTTC | 1 | TRBV19*00 | TRBD1*00 | TRBJ2-1*00 | 0.30% |
| CD4+YFP- WT2 | TGTGCCAGCAGGCGAGACACCCAGTACTTT | 1 | TRBV2*00 | | TRBJ2-5*00 | 0.30% |
| CD4+YFP- WT2 | TGTGCCAGCAGGGATAACCAAGACACCCAGTACTTT | 1 | TRBV26*00 | | TRBJ2-5*00 | 0.30% |
| CD4+YFP- WT2 | TGTGCCAGCAGGGGAGACTCCGACTACACCTTC | 1 | TRBV26*00 | | TRBJ1-2*00 | 0.30% |
| CD4+YFP- WT2 | TGTGCCAGCAGTAAGGACAGGAGGGCTGAGCAGTTCTTC | 1 | TRBV19*00 | TRBD1*00 | TRBJ2-1*00 | 0.30% |
| CD4+YFP- WT2 | TGTGCCAGCAGTAGACGGGCTGAGCAGTTCTTC | 1 | TRBV19*00 | | TRBJ2-1*00 | 0.30% |
| CD4+YFP- WT2 | TGTGCCAGCAGTATAGATGGAGCTGAGCAGTTCTTC | 1 | TRBV19*00 | | TRBJ2-1*00 | 0.30% |
| CD4+YFP- WT2 | TGTGCCAGCAGTATAGCTGCAGGGAATTCTGGAAATACGCTCTATTTT | 1 | TRBV19*00 | | TRBJ1-3*00 | 0.30% |
| CD4+YFP- WT2 | TGTGCCAGCAGTATAGGGGGGGAAGACAACCAGTACTTT | 1 | TRBV19*00 | TRBD2*00 | TRBJ2-5*00 | 0.30% |
| CD4+YFP- WT2 | TGTGCCAGCAGTATCCAGGACCAAGACACCCAGTACTTT | 1 | TRBV19*00 | | TRBJ2-5*00 | 0.30% |
| CD4+YFP- WT2 | TGTGCCAGCAGTATCGAGGACAGGGAAGTCTTCTTT | 1 | TRBV19*00 | TRBD1*00 | TRBJ1-1*00 | 0.30% |
| CD4+YFP- WT2 | TGTGCCAGCAGTATGAGGAATTCGCCCCTCTACTTT | 1 | TRBV19*00 | | TRBJ1-6*00 | 0.30% |
| CD4+YFP- WT2 | TGTGCCAGCAGTATGCGGACAGGACAGTACTTC | 1 | TRBV19*00 | TRBD1*00 | TRBJ2-7*00 | 0.30% |
| CD4+YFP- WT2 | TGTGCCAGCAGTATGGACTGGGGGGCGGAGACACCCAGTACTTT | 1 | TRBV19*00 | TRBD2*00 | TRBJ2-5*00 | 0.30% |
| CD4+YFP- WT2 | TGTGCCAGCAGTCACAGGGGAGACACCCAGTACTTT | 1 | TRBV14*00 | TRBD1*00 | TRBJ2-5*00 | 0.30% |
| CD4+YFP- WT2 | TGTGCCAGCAGTCAGGACAGGATTAGTGCAGAAACGCTGTATTTT | 1 | TRBV14*00 | TRBD1*00 | TRBJ2-3*00 | 0.30% |
| CD4+YFP- WT2 | TGTGCCAGCAGTCCAGGGAGCACCGGGCAGCTCTACTTT | 1 | TRBV26*00 | TRBD1*00 | TRBJ2-2*00 | 0.30% |
| CD4+YFP- WT2 | TGTGCCAGCAGTCCCGCCGGGGAAAACACCTTGTACTTT | 1 | TRBV13-1*00 | TRBD1*00 | TRBJ2-4*00 | 0.30% |
| CD4+YFP- WT2 | TGTGCCAGCAGTCCCTGGGGGGGCGTGGGTCAAAACACCTTGTACTTT | 1 | TRBV19*00 | TRBD2*00 | TRBJ2-4*00 | 0.30% |
| CD4+YFP- WT2 | TGTGCCAGCAGTCCTCAGGGTGACACCTTGTACTTT | 1 | TRBV15*00 | TRBD1*00 | TRBJ2-4*00 | 0.30% |
| CD4+YFP- WT2 | TGTGCCAGCAGTCGATGGGGGACTGATAACTATGCTGAGCAGTTCTTC | 1 | TRBV13-1*00 | TRBD2*00 | TRBJ2-1*00 | 0.30% |
| CD4+YFP- WT2 | TGTGCCAGCAGTCGGACAGGGCCAGACACCCAGTACTTT | 1 | TRBV13-3*00 | TRBD1*00 | TRBJ2-5*00 | 0.30% |
| CD4+YFP- WT2 | TGTGCCAGCAGTCGGACATATAACAACCAGGCTCCGCTTTTT | 1 | TRBV26*00 | TRBD1*00 | TRBJ1-5*00 | 0.30% |
| CD4+YFP- WT2 | TGTGCCAGCAGTCGGGACAATGCAAACTCCGACTACACCTTC | 1 | TRBV15*00 | TRBD1*00 | TRBJ1-2*00 | 0.30% |
| CD4+YFP- WT2 | TGTGCCAGCAGTCGGGACGGTGCAAACTCCGACTACACCTTC | 1 | TRBV13-1*00 | TRBD1*00 | TRBJ1-2*00 | 0.30% |
| CD4+YFP- WT2 | TGTGCCAGCAGTCTCAGGGGACAAGACACCCAGTACTTT | 1 | TRBV24*00 | TRBD1*00 | TRBJ2-5*00 | 0.30% |
| CD4+YFP- WT2 | TGTGCCAGCAGTCTGAACACCTATGAACAGTACTTC | 1 | TRBV26*00 | | TRBJ2-7*00 | 0.30% |
| CD4+YFP- WT2 | TGTGCCAGCAGTCTGACAGCAAACACCGGGCAGCTCTACTTT | 1 | TRBV24*00 | | TRBJ2-2*00 | 0.30% |
| CD4+YFP- WT2 | TGTGCCAGCAGTCTGACGGTCTATGCTGAGCAGTTCTTC | 1 | TRBV26*00 | | TRBJ2-1*00 | 0.30% |
| CD4+YFP- WT2 | TGTGCCAGCAGTCTGCGGGGGGGCGAAAGATTATTTTTC | 1 | TRBV24*00 | TRBD2*00 | TRBJ1-4*00 | 0.30% |
| CD4+YFP- WT2 | TGTGCCAGCAGTCTGGACACAGAAGTCTTCTTT | 1 | TRBV26*00 | | TRBJ1-1*00 | 0.30% |
| CD4+YFP- WT2 | TGTGCCAGCAGTGAAGTCAGGGGTGGCACAGAAGTCTTCTTT | 1 | TRBV13-1*00 | TRBD1*00 | TRBJ1-1*00 | 0.30% |
| CD4+YFP- WT2 | TGTGCCAGCAGTGACACAATTTCCAACGAAAGATTATTTTTC | 1 | TRBV13-3*00 | | TRBJ1-4*00 | 0.30% |
| CD4+YFP- WT2 | TGTGCCAGCAGTGACAGTGCAGAAACGCTGTATTTT | 1 | TRBV13-3*00 | | TRBJ2-3*00 | 0.30% |
| CD4+YFP- WT2 | TGTGCCAGCAGTGACCACACAAACTCCGACTACACCTTC | 1 | TRBV13-1*00 | | TRBJ1-2*00 | 0.30% |

| | | | | | | |
|---|---|---|---|---|---|---|
| CD4+YFP- WT2 | TGTGCCAGCAGTGACGAGACAAACACAGAAGTCTTCTTT | 1 | TRBV13-1*00 | | TRBJ1-1*00 | 0.30% |
| CD4+YFP- WT2 | TGTGCCAGCAGTGAGCGGGATGTCACCCAGTACTTT | 1 | TRBV13-3*00 | TRBD1*00 | TRBJ2-5*00 | 0.30% |
| CD4+YFP- WT2 | TGTGCCAGCAGTGAGGGCAGGGCAAACTCCGACTACACCTTC | 1 | TRBV13-1*00 | | TRBJ1-2*00 | 0.30% |
| CD4+YFP- WT2 | TGTGCCAGCAGTGAGGGGACAGGGGGCTCCTATGAACAGTACTTC | 1 | TRBV13-1*00 | TRBD1*00 | TRBJ2-7*00 | 0.30% |
| CD4+YFP- WT2 | TGTGCCAGCAGTGAGTTGGGACAGGGGGCCGAAAGATTATTTTTC | 1 | TRBV13-1*00 | TRBD1*00 | TRBJ1-4*00 | 0.30% |
| CD4+YFP- WT2 | TGTGCCAGCAGTGATACAGGGTATTCTGGAAATACGCTCTATTTT | 1 | TRBV13-1*00 | TRBD1*00 | TRBJ1-3*00 | 0.30% |
| CD4+YFP- WT2 | TGTGCCAGCAGTGATACATGGGGGTATGCTGAGCAGTTCTTC | 1 | TRBV13-1*00 | TRBD1*00 | TRBJ2-1*00 | 0.30% |
| CD4+YFP- WT2 | TGTGCCAGCAGTGATATGGATGGTAGTGCAGAAACGCTGTATTTT | 1 | TRBV13-3*00 | | TRBJ2-3*00 | 0.30% |
| CD4+YFP- WT2 | TGTGCCAGCAGTGATCGGGACAGGGTAGACACCCAGTACTTT | 1 | TRBV13-3*00 | TRBD1*00 | TRBJ2-5*00 | 0.30% |
| CD4+YFP- WT2 | TGTGCCAGCAGTGATCGGGGGTACTCCGACTACACCTTC | 1 | TRBV13-1*00 | TRBD1*00 | TRBJ1-2*00 | 0.30% |
| CD4+YFP- WT2 | TGTGCCAGCAGTGATGCAAGGACAGGGGAGGCTGAGCAGTTCTTC | 1 | TRBV13-1*00 | TRBD1*00 | TRBJ2-1*00 | 0.30% |
| CD4+YFP- WT2 | TGTGCCAGCAGTGATGCAGGCCTCTCCTATGAACAGTACTTC | 1 | TRBV13-1*00 | | TRBJ2-7*00 | 0.30% |
| CD4+YFP- WT2 | TGTGCCAGCAGTGATGGCGTCAACTATGCTGAGCAGTTCTTC | 1 | TRBV13-1*00 | | TRBJ2-1*00 | 0.30% |
| CD4+YFP- WT2 | TGTGCCAGCAGTGATGGGACAGGGATTAACCAAGACACCCAGTACTTT | 1 | TRBV13-3*00 | TRBD1*00 | TRBJ2-5*00 | 0.30% |
| CD4+YFP- WT2 | TGTGCCAGCAGTGATGGGACAGGGGGCCAAGACACCCAGTACTTT | 1 | TRBV13-1*00 | TRBD1*00 | TRBJ2-5*00 | 0.30% |
| CD4+YFP- WT2 | TGTGCCAGCAGTGATGGGACTGCTGACACCCAGTACTTT | 1 | TRBV13-3*00 | TRBD2*00 | TRBJ2-5*00 | 0.30% |
| CD4+YFP- WT2 | TGTGCCAGCAGTGATGGGGGGCGCGGGTATGCTGAGCAGTTCTTC | 1 | TRBV13-1*00 | TRBD1*00 | TRBJ2-1*00 | 0.30% |
| CD4+YFP- WT2 | TGTGCCAGCAGTGATGGGGGGGGTTATGAACAGTACTTC | 1 | TRBV13-1*00 | TRBD2*00 | TRBJ2-7*00 | 0.30% |
| CD4+YFP- WT2 | TGTGCCAGCAGTGATGGTTCTGGAAATACGCTCTATTTT | 1 | TRBV13-1*00 | | TRBJ1-3*00 | 0.30% |
| CD4+YFP- WT2 | TGTGCCAGCAGTGATGTGAACCAAGACACCCAGTACTTT | 1 | TRBV13-1*00 | | TRBJ2-5*00 | 0.30% |
| CD4+YFP- WT2 | TGTGCCAGCAGTGATTCAAGTCAAAACACCTTGTACTTT | 1 | TRBV13-1*00 | | TRBJ2-4*00 | 0.30% |
| CD4+YFP- WT2 | TGTGCCAGCAGTGATTCCCAAGACACCCAGTACTTT | 1 | TRBV13-1*00 | | TRBJ2-5*00 | 0.30% |
| CD4+YFP- WT2 | TGTGCCAGCAGTGCAGGAAACTCCGACTACACCTTC | 1 | TRBV13-3*00 | | TRBJ1-2*00 | 0.30% |
| CD4+YFP- WT2 | TGTGCCAGCAGTGCCTCCAGGGGGCGCTATGAACAGTACTTC | 1 | TRBV13-1*00 | TRBD1*00 | TRBJ2-7*00 | 0.30% |
| CD4+YFP- WT2 | TGTGCCAGCAGTGGGACAGGGGATTCGCCCCTCTACTTT | 1 | TRBV13-3*00 | TRBD1*00 | TRBJ1-6*00 | 0.30% |
| CD4+YFP- WT2 | TGTGCCAGCAGTGGGACAGGGGGCATCTCCGACTACACCTTC | 1 | TRBV13-1*00 | TRBD1*00 | TRBJ1-2*00 | 0.30% |
| CD4+YFP- WT2 | TGTGCCAGCAGTGGGGACAACCAGGCTCCGCTTTTT | 1 | TRBV13-1*00 | | TRBJ1-5*00 | 0.30% |
| CD4+YFP- WT2 | TGTGCCAGCAGTGGGGACTGGAATGCTGAGCAGTTCTTC | 1 | TRBV13-1*00 | TRBD2*00 | TRBJ2-1*00 | 0.30% |
| CD4+YFP- WT2 | TGTGCCAGCAGTGGGGGAGGCCAAGACACCCAGTACTTT | 1 | TRBV13-1*00 | | TRBJ2-5*00 | 0.30% |
| CD4+YFP- WT2 | TGTGCCAGCAGTTCTGGGGAAGACACCCAGTACTTT | 1 | TRBV15*00 | TRBD2*00 | TRBJ2-5*00 | 0.30% |
| CD4+YFP- WT2 | TGTGCCAGCAGTTCTGGGGGGCCTAGTGCAGAAACGCTGTATTTT | 1 | TRBV14*00 | TRBD2*00 | TRBJ2-3*00 | 0.30% |
| CD4+YFP- WT2 | TGTGCCAGCAGTTTACAGAACGAAAGATTATTTTTC | 1 | TRBV19*00 | | TRBJ1-4*00 | 0.30% |
| CD4+YFP- WT2 | TGTGCCAGCAGTTTAGCGGGGGGGAGAGAACAGTACTTC | 1 | TRBV15*00 | TRBD2*00 | TRBJ2-7*00 | 0.30% |
| CD4+YFP- WT2 | TGTGCCAGCAGTTTAGGCAGTTCTGGAAATACGCTCTATTTT | 1 | TRBV14*00 | | TRBJ1-3*00 | 0.30% |
| CD4+YFP- WT2 | TGTGCCAGCAGTTTAGGGTATGAACAGTACTTC | 1 | TRBV15*00 | | TRBJ2-7*00 | 0.30% |
| CD4+YFP- WT2 | TGTGCCAGCAGTTTATACAATTCTGGAAATACGCTCTATTTT | 1 | TRBV15*00 | | TRBJ1-3*00 | 0.30% |
| CD4+YFP- WT2 | TGTGCCAGCAGTTTATGGAACACAGAAGTCTTCTTT | 1 | TRBV15*00 | | TRBJ1-1*00 | 0.30% |
| CD4+YFP- WT2 | TGTGCCAGCAGTTTCAGTTATGCTGAGCAGTTCTTC | 1 | TRBV13-3*00 | | TRBJ2-1*00 | 0.30% |
| CD4+YFP- WT2 | TGTGCCAGCAGTTTCATGGGGAACACAGAAGTCTTCTTT | 1 | TRBV14*00 | TRBD2*00 | TRBJ1-1*00 | 0.30% |
| CD4+YFP- WT2 | TGTGCCAGCAGTTTCGACTGGGTTAACTATGCTGAGCAGTTCTTC | 1 | TRBV15*00 | TRBD2*00 | TRBJ2-1*00 | 0.30% |
| CD4+YFP- WT2 | TGTGCCAGCAGTTTCGAGGGCGAACAGTACTTC | 1 | TRBV15*00 | | TRBJ2-7*00 | 0.30% |
| CD4+YFP- WT2 | TGTGCCAGCAGTTTGGGGGGGGCCGCCGGGCAGCTCTACTTT | 1 | TRBV15*00 | TRBD2*00 | TRBJ2-2*00 | 0.30% |
| CD4+YFP- WT2 | TGTGCCAGCAGTTTTGGCCTCTCCTATGAACAGTACTTC | 1 | TRBV15*00 | | TRBJ2-7*00 | 0.30% |
| CD4+YFP- WT2 | TGTGCCAGCCGGGACAGGGGTGCAGAAACGCTGTATTTT | 1 | TRBV12-1*00 | TRBD1*00 | TRBJ2-3*00 | 0.30% |
| CD4+YFP- WT2 | TGTGCCAGCGCCCTGGGGGGGCGGGCTGAGCAGTTCTTC | 1 | TRBV12-1*00 | TRBD2*00 | TRBJ2-1*00 | 0.30% |
| CD4+YFP- WT2 | TGTGCCAGCGGAAATTCTGGAAATACGCTCTATTTT | 1 | TRBV13-2*00 | | TRBJ1-3*00 | 0.30% |
| CD4+YFP- WT2 | TGTGCCAGCGGACAGCAAGACACCCAGTACTTT | 1 | TRBV13-2*00 | | TRBJ2-5*00 | 0.30% |
| CD4+YFP- WT2 | TGTGCCAGCGGGGACAGGGGGCGGGGTGCTGAGCAGTTCTTC | 1 | TRBV13-2*00 | TRBD1*00 | TRBJ2-1*00 | 0.30% |
| CD4+YFP- WT2 | TGTGCCAGCGGTCTGGGGAGCTCCTATGAACAGTACTTC | 1 | TRBV13-2*00 | | TRBJ2-7*00 | 0.30% |
| CD4+YFP- WT2 | TGTGCCAGCGGTGAAACACTGGGGGTTGAACAGTACTTC | 1 | TRBV13-2*00 | TRBD2*00 | TRBJ2-7*00 | 0.30% |
| CD4+YFP- WT2 | TGTGCCAGCGGTGAATGGGACAGGGAGGAGTCCTATGAACAGTACTTC | 1 | TRBV13-2*00 | TRBD1*00 | TRBJ2-7*00 | 0.30% |
| CD4+YFP- WT2 | TGTGCCAGCGGTGACAATTCTGGAAATACGCTCTATTTT | 1 | TRBV13-2*00 | | TRBJ1-3*00 | 0.30% |
| CD4+YFP- WT2 | TGTGCCAGCGGTGACAGGGGCTCCTATGAACAGTACTTC | 1 | TRBV13-2*00 | | TRBJ2-7*00 | 0.30% |
| CD4+YFP- WT2 | TGTGCCAGCGGTGACCTCAGGGTTTCCAACGAAAGATTATTTTTC | 1 | TRBV13-2*00 | TRBD1*00 | TRBJ1-4*00 | 0.30% |
| CD4+YFP- WT2 | TGTGCCAGCGGTGAGCACAGGGGGCGCTATGAACAGTACTTC | 1 | TRBV13-2*00 | TRBD1*00 | TRBJ2-7*00 | 0.30% |
| CD4+YFP- WT2 | TGTGCCAGCGGTGAGCCACAGGGGTATGCTGAGCAGTTCTTC | 1 | TRBV13-2*00 | TRBD1*00 | TRBJ2-1*00 | 0.30% |
| CD4+YFP- WT2 | TGTGCCAGCGGTGAGCCCTCAGGGACGCCCCTCTACTTT | 1 | TRBV13-2*00 | TRBD1*00 | TRBJ1-6*00 | 0.30% |
| CD4+YFP- WT2 | TGTGCCAGCGGTGATGAGGGAGGGAACTCCGACTACACCTTC | 1 | TRBV13-2*00 | | TRBJ1-2*00 | 0.30% |
| CD4+YFP- WT2 | TGTGCCAGCGGTGATGATTCGCCCCTCTACTTT | 1 | TRBV13-2*00 | | TRBJ1-6*00 | 0.30% |
| CD4+YFP- WT2 | TGTGCCAGCGGTGATGCACAGGGGGCGGACACCCAGTACTTT | 1 | TRBV13-2*00 | TRBD1*00 | TRBJ2-5*00 | 0.30% |
| CD4+YFP- WT2 | TGTGCCAGCGGTGATGCAGGTGCAGAAACGCTGTATTTT | 1 | TRBV13-2*00 | | TRBJ2-3*00 | 0.30% |
| CD4+YFP- WT2 | TGTGCCAGCGGTGATGCCTGGGGGGCTGACACCCAGTACTTT | 1 | TRBV13-2*00 | TRBD2*00 | TRBJ2-5*00 | 0.30% |
| CD4+YFP- WT2 | TGTGCCAGCGGTGATGCTGGGAGCTCCTATGAACAGTACTTC | 1 | TRBV13-2*00 | | TRBJ2-7*00 | 0.30% |
| CD4+YFP- WT2 | TGTGCCAGCGGTGATGGGACATCAAACTCCGACTACACCTTC | 1 | TRBV13-2*00 | TRBD1*00 | TRBJ1-2*00 | 0.30% |
| CD4+YFP- WT2 | TGTGCCAGCGGTGATGGGGGGCGCTTTGAACAGTACTTC | 1 | TRBV13-2*00 | TRBD1*00 | TRBJ2-7*00 | 0.30% |
| CD4+YFP- WT2 | TGTGCCAGCGGTGATGGGGGGGGAGACACCCAGTACTTT | 1 | TRBV13-2*00 | TRBD2*00 | TRBJ2-5*00 | 0.30% |
| CD4+YFP- WT2 | TGTGCCAGCGGTGCCCAGGCCTCCGACTACACCTTC | 1 | TRBV13-2*00 | TRBD2*00 | TRBJ1-2*00 | 0.30% |
| CD4+YFP- WT2 | TGTGCCAGCGGTGCCGACAGGGCTAGTGCAGAAACGCTGTATTTT | 1 | TRBV13-2*00 | TRBD1*00 | TRBJ2-3*00 | 0.30% |
| CD4+YFP- WT2 | TGTGCCAGCGGTGCGGGGGGACAAAACACCTTGTACTTT | 1 | TRBV13-2*00 | TRBD2*00 | TRBJ2-4*00 | 0.30% |
| CD4+YFP- WT2 | TGTGCCAGCGGTGGGGGCGAGTACTTC | 1 | TRBV13-2*00 | TRBD1*00 | TRBJ2-7*00 | 0.30% |
| CD4+YFP- WT2 | TGTGCCAGCTCGCAGAACTCCGACTACACCTTC | 1 | TRBV12-1*00 | | TRBJ1-2*00 | 0.30% |
| CD4+YFP- WT2 | TGTGCCAGCTCTCGCGGGGGTTATGCTGAGCAGTTCTTC | 1 | TRBV12-1*00 | TRBD1*00 | TRBJ2-1*00 | 0.30% |
| CD4+YFP- WT2 | TGTGCCAGCTCTCTAGTAAACTCCGACTACACCTTC | 1 | TRBV12-1*00 | | TRBJ1-2*00 | 0.30% |
| CD4+YFP- WT2 | TGTGCCAGCTCTCTCGACGTCAGCTCCTATGAACAGTACTTC | 1 | TRBV12-1*00 | | TRBJ2-7*00 | 0.30% |
| CD4+YFP- WT2 | TGTGCCAGCTCTCTCGATGGGGACACCCAGTACTTT | 1 | TRBV12-2*00 | | TRBJ2-5*00 | 0.30% |

| | | | | | | |
|---|---|---|---|---|---|---|
| CD4+YFP- WT2 | TGTGCCAGCTCTCTCGCCTGGGGTAGTGCAGAAACGCTGTATTTT | 1 | TRBV12-1*00 | TRBD2*00 | TRBJ2-3*00 | 0.30% |
| CD4+YFP- WT2 | TGTGCCAGCTTCAAAAACACAGAAGTCTTCTTT | 1 | TRBV13-1*00 | | TRBJ1-1*00 | 0.30% |
| CD4+YFP- WT2 | TGTGCCAGGGACCTTCCTCCGCTGTATTTT | 1 | TRBV31*00 | TRBD1*00 | TRBJ2-3*00 | 0.30% |
| CD4+YFP- WT2 | TGTGCCCCAGGGGGGCCCGACTACACCTTC | 1 | TRBV13-3*00 | TRBD1*00 | TRBJ1-2*00 | 0.30% |
| CD4+YFP- WT2 | TGTGCCTCTAAAAGTCAAAACACCTTGTACTTT | 1 | TRBV31*00 | | TRBJ2-4*00 | 0.30% |
| CD4+YFP- WT2 | TGTGCCTGCCGGGACAGGGGAGACACCCAGTACTTT | 1 | TRBV31*00 | TRBD1*00 | TRBJ2-5*00 | 0.30% |
| CD4+YFP- WT2 | TGTGCCTGGAGCCCCGACACAGAAGTCTTCTTT | 1 | TRBV31*00 | | TRBJ1-1*00 | 0.30% |
| CD4+YFP- WT2 | TGTGCCTGGAGCCGGGACATCTCCTATGAACAGTACTTC | 1 | TRBV31*00 | TRBD1*00 | TRBJ2-7*00 | 0.30% |
| CD4+YFP- WT2 | TGTGCCTGGAGGACTGGGGGGGCTGAAACGCTGTATTTT | 1 | TRBV31*00 | TRBD2*00 | TRBJ2-3*00 | 0.30% |
| CD4+YFP- WT2 | TGTGCCTGGAGGGGACAGGGTAACACCGGGCAGCTCTACTTT | 1 | TRBV31*00 | TRBD1*00 | TRBJ2-2*00 | 0.30% |
| CD4+YFP- WT2 | TGTGCCTGGAGTCAGATCTATGCTGAGCAGTTCTTC | 1 | TRBV31*00 | | TRBJ2-1*00 | 0.30% |
| CD4+YFP- WT2 | TGTGCCTGGAGTCCAGGGGGCGACTCCGACTACACCTTC | 1 | TRBV31*00 | TRBD1*00 | TRBJ1-2*00 | 0.30% |
| CD4+YFP- WT2 | TGTGCCTGGAGTCGAGACAGGGCTGGAAATACGCTCTATTTT | 1 | TRBV31*00 | TRBD1*00 | TRBJ1-3*00 | 0.30% |
| CD4+YFP- WT2 | TGTGCCTGGAGTCTAAACTCCGACTACACCTTC | 1 | TRBV31*00 | | TRBJ1-2*00 | 0.30% |
| CD4+YFP- WT2 | TGTGCCTGGAGTCTAGAGGGGAGGAACTCCGACTACACCTTC | 1 | TRBV31*00 | TRBD1*00 | TRBJ1-2*00 | 0.30% |
| CD4+YFP- WT2 | TGTGCCTGGAGTCTAGGCAGCAACGAAAGATTATTTTTC | 1 | TRBV31*00 | | TRBJ1-4*00 | 0.30% |
| CD4+YFP- WT2 | TGTGCCTGGAGTCTAGTGGGTAATTCCTATAATTCGCCCCTCTACTTT | 1 | TRBV31*00 | | TRBJ1-6*00 | 0.30% |
| CD4+YFP- WT2 | TGTGCCTGGAGTGGCGGGAGACACCCAGTACTTT | 1 | TRBV31*00 | | TRBJ2-5*00 | 0.30% |
| CD4+YFP- WT2 | TGTGCCTGGGCAGGGTCCACAGAAGTCTTCTTT | 1 | TRBV31*00 | TRBD1*00 | TRBJ1-1*00 | 0.30% |
| CD4+YFP- WT2 | TGTGCCTGGGGACAGGGGAAAAACTCCGACTACACCTTC | 1 | TRBV31*00 | TRBD1*00 | TRBJ1-2*00 | 0.30% |
| CD4+YFP- WT2 | TGTGCTAGCACTGGTAGTCAAAACACCTTGTACTTT | 1 | TRBV21*00 | | TRBJ2-4*00 | 0.30% |
| CD4+YFP- WT2 | TGTGCTAGCAGCGGGACTGGGGACACCCAGTACTTT | 1 | TRBV17*00 | TRBD2*00 | TRBJ2-5*00 | 0.30% |
| CD4+YFP- WT2 | TGTGCTAGCAGCGGGGGGATTGCTGAGCAGTTCTTC | 1 | TRBV17*00 | TRBD2*00 | TRBJ2-1*00 | 0.30% |
| CD4+YFP- WT2 | TGTGCTAGCAGTAGAAGGACTGGGGGTTATGAACAGTACTTC | 1 | TRBV17*00 | TRBD2*00 | TRBJ2-7*00 | 0.30% |
| CD4+YFP- WT2 | TGTGCTAGCAGTAGAGGCAGGGACGGCTCCGACTACACCTTC | 1 | TRBV17*00 | TRBD1*00 | TRBJ1-2*00 | 0.30% |
| CD4+YFP- WT2 | TGTGCTAGCAGTAGTCTGGGGAACTATGCTGAGCAGTTCTCC | 1 | TRBV17*00 | TRBD2*00 | TRBJ2-1*00 | 0.30% |
| CD4+YFP- WT2 | TGTGCTAGCAGTCTTTATGCTGAGCAGTTCTTC | 1 | TRBV29*00 | | TRBJ2-1*00 | 0.30% |
| CD4+YFP- WT2 | TGTGCTAGCAGTTTCAACATCCAAGACACCCAGTACTTT | 1 | TRBV29*00 | | TRBJ2-5*00 | 0.30% |
| CD4+YFP- WT2 | TGTGCTAGCAGTTTCTGGGACACTAACTATGCTGAGCAGTTCTTC | 1 | TRBV29*00 | TRBD1*00 | TRBJ2-1*00 | 0.30% |
| CD4+YFP- WT2 | TGTGCTGGGAGTCTGGGGGTTTATGAACAGTACTTC | 1 | TRBV29*00 | TRBD2*00 | TRBJ2-7*00 | 0.30% |
| CD4+YFP- WT2 | TGTGGCAGCAGTTTAGACACAGAAGTCTTCTTT | 1 | TRBV15*00 | | TRBJ1-1*00 | 0.30% |
| CD4+YFP- WT2 | TGTGGTGCTAGGGAAGACACCCAGTACTTT | 1 | TRBV20*00 | | TRBJ2-5*00 | 0.30% |
| CD4+YFP- WT2 | TGTGGTGCTAGGGACAACTCCTATGAACAGTACTTC | 1 | TRBV20*00 | | TRBJ2-7*00 | 0.30% |
| CD4+YFP- WT2 | TGTGGTGCTAGGGACACTAACAACCAGGCTCCGCTTTTT | 1 | TRBV20*00 | | TRBJ1-5*00 | 0.30% |
| CD4+YFP- WT2 | TGTGGTGCTAGGGACAGTAGTGCAGAAACGCTGTATTTT | 1 | TRBV20*00 | | TRBJ2-3*00 | 0.30% |
| CD4+YFP- WT2 | TGTGGTGCTAGGGACATTAACCAAGACACCCAGTACTTT | 1 | TRBV20*00 | | TRBJ2-5*00 | 0.30% |
| CD4+YFP- WT2 | TGTGGTGCTAGGGACTGGGGGAACACCGGGCAGCTCTACTTT | 1 | TRBV20*00 | TRBD2*00 | TRBJ2-2*00 | 0.30% |
| CD4+YFP- WT2 | TGTGGTGCTAGGGATAGGACTGAGCAGTTCTTC | 1 | TRBV20*00 | | TRBJ2-1*00 | 0.30% |
| CD4+YFP- WT2 | TGTGGTGCTAGGGATATGGGCAGTCAAAACACCTTGTACTTT | 1 | TRBV20*00 | | TRBJ2-4*00 | 0.30% |
| CD4+YFP- WT2 | TGTGGTGCTAGGGGCAGTAGTCAAAACACCTTGTACTTT | 1 | TRBV20*00 | | TRBJ2-4*00 | 0.30% |
| CD4+YFP- WT2 | TGTGGTGCTAGGGGGACTGGGGCAGAAACGCTGTATTTT | 1 | TRBV20*00 | TRBD2*00 | TRBJ2-3*00 | 0.30% |
| CD4+YFP- WT2 | TGTGGTGCTAGGGGGGTTAACCAAGACACCCAGTACTTT | 1 | TRBV20*00 | | TRBJ2-5*00 | 0.30% |
| CD4+YFP- WT2 | TGTGGTGCTAGGGTTGGACTTAACTATGCTGAGCAGTTCTTC | 1 | TRBV20*00 | TRBD2*00 | TRBJ2-1*00 | 0.30% |
| CD4+YFP- WT2 | TGTGGTGCTATAGGACAGCAAGACACCCAGTACTTC | 1 | TRBV20*00 | TRBD1*00 | TRBJ2-5*00 | 0.30% |
| CD4+YFP- WT2 | TGTGGTGCTTTGACGGTAAACTCCGACTACACCTTC | 1 | TRBV20*00 | | TRBJ1-2*00 | 0.30% |
| CD4+YFP- WT2 | TGTGCCAGCAGCCAAGACAGCTATGAACAGTACTTC | 1 | TRBV5*00 | | TRBJ2-7*00 | 0.30% |
| CD4+YFP- WT2 | TGTGCAAGCAGCTTAGAATATGAACAGTACTTC | 1 | TRBV16*00 | | TRBJ2-7*00 | 0.30% |
| CD4+YFP- WT2 | TGTGCCAGCGGTGATTCCTATAATTCGCCCCTCTACTTT | 1 | TRBV13-2*00 | | TRBJ1-6*00 | 0.30% |
| CD4+YFP- WT2 | TGTGCCAGCAGTCTGGGAAATACGCTCTATTTT | 1 | TRBV26*00 | | TRBJ1-3*00 | 0.30% |
| CD4+YFP- WT2 | TGTGCCAGCAGTTTAGACACAGAAGTCTTCTTT | 1 | TRBV15*00 | | TRBJ1-1*00 | 0.30% |
| CD4+YFP- WT2 | TGTGCCAGCAGTCTGTCAACGTATGAACAGTACTTC | 1 | TRBV26*00 | | TRBJ2-7*00 | 0.30% |
| CD4+YFP- WT2 | TGTGCTAGCAGTAGGGACAACAACGAAAGATTATTTTTC | 1 | TRBV17*00 | TRBD1*00 | TRBJ1-4*00 | 0.30% |

#### CD4+YFP- DKO

| Sample Name | CDR3 Sequence | clone Count | best V Hit | best D Hit | best J Hit | Frequency |
|---|---|---|---|---|---|---|
| CD4+YFP- DKO | TGTGCCAGCAGCCAAGATAGAGACACCCAGTACTTT | 21 | TRBV2*00 | | TRBJ2-5*00 | 5.95% |
| CD4+YFP- DKO | TGTGCCAGCAGCCAAGATACGGGGGAAGACACCCAGTACTTT | 12 | TRBV5*00 | TRBD1*00 | TRBJ2-5*00 | 3.40% |
| CD4+YFP- DKO | TGTGCCAGCGGGGACTGGGGAAACACCGGGCAGCTCTACTTT | 5 | TRBV13-2*00 | TRBD2*00 | TRBJ2-2*00 | 1.42% |
| CD4+YFP- DKO | TGTGCAAGCAGCTTAGAGGACACCGGGCAGCTCTACTTT | 4 | TRBV16*00 | | TRBJ2-2*00 | 1.13% |
| CD4+YFP- DKO | TGCACCTGCAGTGCTCGGAACTCCGACTACACCTTC | 3 | TRBV1*00 | | TRBJ1-2*00 | 0.85% |
| CD4+YFP- DKO | TGTGCCAGCAGTCCGGGACTGGGGGGTCAAAACACCTTGTACTTT | 3 | TRBV26*00 | TRBD2*00 | TRBJ2-4*00 | 0.85% |
| CD4+YFP- DKO | TGTGCCAGCAGTGAAAGGGATGAAGTCTTCTTT | 3 | TRBV13-1*00 | | TRBJ1-1*00 | 0.85% |
| CD4+YFP- DKO | TGTGCCAGCAGTTCATCAAACACCGGGCAGCTCTACTTT | 3 | TRBV15*00 | | TRBJ2-2*00 | 0.85% |
| CD4+YFP- DKO | TGTGCCAGCAGTTCTCGACTGGACTATGAACAGTACTTC | 3 | TRBV22*00 | | TRBJ2-7*00 | 0.85% |
| CD4+YFP- DKO | TGCACCTGCAGTGCAAGTCAAAACACCTTGTACTTT | 2 | TRBV1*00 | | TRBJ2-4*00 | 0.57% |
| CD4+YFP- DKO | TGCACCTGCAGTGCAGATCCAGGGCAGCTCTACTTT | 2 | TRBV1*00 | | TRBJ2-2*00 | 0.57% |
| CD4+YFP- DKO | TGTGCCAGCAGCCAAGATAGTAGTCAAAACACCTTGTACTTT | 2 | TRBV2*00 | | TRBJ2-4*00 | 0.57% |
| CD4+YFP- DKO | TGTGCCAGCAGCTCGACTGGGGGGAACTATGCTGAGCAGTTCTTC | 2 | TRBV25*00 | TRBD2*00 | TRBJ2-1*00 | 0.57% |
| CD4+YFP- DKO | TGTGCCAGCAGCTGGGACACAAACCAAGACACCCAGTACTTT | 2 | TRBV3*00 | TRBD1*00 | TRBJ2-5*00 | 0.57% |
| CD4+YFP- DKO | TGTGCCAGCAGTATCTGGAGTGCAGAAACGCTGTATTTT | 2 | TRBV19*00 | | TRBJ2-3*00 | 0.57% |
| CD4+YFP- DKO | TGTGCCAGCAGTCTCCGGGACAGGAGCAACGAAAGATTATTTTTC | 2 | TRBV19*00 | TRBD1*00 | TRBJ1-4*00 | 0.57% |
| CD4+YFP- DKO | TGTGCCAGCAGTCTGACTGGGGGGAACACCGGGCAGCTCTACTTT | 2 | TRBV24*00 | TRBD2*00 | TRBJ2-2*00 | 0.57% |
| CD4+YFP- DKO | TGTGCCAGCAGTGATGACTGGGGGGCCTATGCTGAGCAGTTCTTC | 2 | TRBV13-1*00 | TRBD2*00 | TRBJ2-1*00 | 0.57% |
| CD4+YFP- DKO | TGTGCCAGCAGTGATGGGGGGGCGGACACCCAGTACTTT | 2 | TRBV13-1*00 | TRBD2*00 | TRBJ2-5*00 | 0.57% |
| CD4+YFP- DKO | TGTGCCAGCAGTGGACAGATTTCCAACGAAAGATTATTTTTC | 2 | TRBV13-1*00 | TRBD1*00 | TRBJ1-4*00 | 0.57% |

| | | | | | | |
|---|---|---|---|---|---|---|
| CD4+YFP- DKO | TGTGCCAGCAGTTTCGCCACAAACACCTTGTACTTT | 2 | TRBV14*00 | | TRBJ2-4*00 | 0.57% |
| CD4+YFP- DKO | TGTGCCAGCGGTGACCCCCGTGCAGAAACGCTGTATTTT | 2 | TRBV13-2*00 | TRBD1*00 | TRBJ2-3*00 | 0.57% |
| CD4+YFP- DKO | TGTGCCAGCTCTCTCGGGGAAGACACCCAGTACTTT | 2 | TRBV12-1*00 | | TRBJ2-5*00 | 0.57% |
| CD4+YFP- DKO | TGTGCCTGGAGTCTAGGACATTCTGGAAATACGCTCTATTTT | 2 | TRBV31*00 | | TRBJ1-3*00 | 0.57% |
| CD4+YFP- DKO | TGTGGTGCTAGGGACTGGGGACAAGACACCCAGTACTTT | 2 | TRBV20*00 | TRBD2*00 | TRBJ2-5*00 | 0.57% |
| CD4+YFP- DKO | TGCACCTGCAGTGCAAGGGGGATTAACCAAGACACCCAGTACTTT | 1 | TRBV1*00 | TRBD1*00 | TRBJ2-5*00 | 0.28% |
| CD4+YFP- DKO | TGCACCTGCAGTGCAGACAGGGTTGAACAGTACTTC | 1 | TRBV1*00 | TRBD1*00 | TRBJ2-7*00 | 0.28% |
| CD4+YFP- DKO | TGCACCTGCAGTGCAGACTGGGGGGGGAGCTCCTATGAACAGTACTTC | 1 | TRBV1*00 | TRBD2*00 | TRBJ2-7*00 | 0.28% |
| CD4+YFP- DKO | TGCACCTGCAGTGCAGATCTGGTCTCCTATGAACAGTACTTC | 1 | TRBV1*00 | | TRBJ2-7*00 | 0.28% |
| CD4+YFP- DKO | TGCACCTGCAGTGCAGATTTGAACAGTACTTC | 1 | TRBV1*00 | | TRBJ2-7*00 | 0.28% |
| CD4+YFP- DKO | TGCACCTGCAGTGCAGGACTGGGGGGGAGTGAACAGTACTTC | 1 | TRBV1*00 | TRBD2*00 | TRBJ2-7*00 | 0.28% |
| CD4+YFP- DKO | TGCACCTGCAGTGCAGGGATTGCAAACACAGAAGTCTTCTTT | 1 | TRBV1*00 | | TRBJ1-1*00 | 0.28% |
| CD4+YFP- DKO | TGCACCTGCAGTGCAGGGGGCGCGTCCTATGAACAGTACTTC | 1 | TRBV1*00 | TRBD1*00 | TRBJ2-7*00 | 0.28% |
| CD4+YFP- DKO | TGCACCTGCAGTGCAGTGGAGGGGGCCAAAGACACCCAGTACTTT | 1 | TRBV1*00 | TRBD1*00 | TRBJ2-5*00 | 0.28% |
| CD4+YFP- DKO | TGCACCTGCAGTGCAGTGGTGGGGGGCACCCAGTACTTT | 1 | TRBV1*00 | TRBD2*00 | TRBJ2-5*00 | 0.28% |
| CD4+YFP- DKO | TGCACCTGCAGTGCCACGGCAGAAGTCTTCTTT | 1 | TRBV1*00 | | TRBJ1-1*00 | 0.28% |
| CD4+YFP- DKO | TGCACCTGCAGTGCGAACAGGGGAAGAACCGGGCAGCTCTACTTT | 1 | TRBV1*00 | TRBD1*00 | TRBJ2-2*00 | 0.28% |
| CD4+YFP- DKO | TGCACCTGCAGTGCGGGGGACTATGCTGAGCAGTTCTTC | 1 | TRBV1*00 | TRBD1*00 | TRBJ2-1*00 | 0.28% |
| CD4+YFP- DKO | TGCACCTGCAGTGGGACGGATAACTATACTGAGCAGTTCTTC | 1 | TRBV1*00 | | TRBJ2-1*00 | 0.28% |
| CD4+YFP- DKO | TGCACCTGCAGTGGGGATTCCTATGAACAGTACTTC | 1 | TRBV1*00 | | TRBJ2-7*00 | 0.28% |
| CD4+YFP- DKO | TGCACCTGCAGTGGGGGGGAGGCTGAGCAGTTCTTC | 1 | TRBV1*00 | TRBD2*00 | TRBJ2-1*00 | 0.28% |
| CD4+YFP- DKO | TGCTCCAGCAGGGGGACAGCACATTACTTC | 1 | TRBV23*00 | TRBD1*00 | TRBJ2-7*00 | 0.28% |
| CD4+YFP- DKO | TGCTCCAGCAGTCAATCGGCAGGGGACCAAGACACCCAGTACTTT | 1 | TRBV23*00 | TRBD1*00 | TRBJ2-5*00 | 0.28% |
| CD4+YFP- DKO | TGTACCAGCGGTGGGACACCAAACTCCGACTACACCTTC | 1 | TRBV13-2*00 | TRBD1*00 | TRBJ1-2*00 | 0.28% |
| CD4+YFP- DKO | TGTGCAAGCAGACTGGGGGGATATGAACAGTACTTC | 1 | TRBV16*00 | TRBD2*00 | TRBJ2-7*00 | 0.28% |
| CD4+YFP- DKO | TGTGCAAGCAGCAGGACAGAAGAAGTCTTCTTT | 1 | TRBV16*00 | TRBD1*00 | TRBJ1-1*00 | 0.28% |
| CD4+YFP- DKO | TGTGCAAGCAGCTCAGACATCTATGCTGAGCAGTTCTTC | 1 | TRBV16*00 | | TRBJ2-1*00 | 0.28% |
| CD4+YFP- DKO | TGTGCAAGCAGCTCCCTGGGGGGTAACCAAGACACCCAGTACTTT | 1 | TRBV16*00 | TRBD2*00 | TRBJ2-5*00 | 0.28% |
| CD4+YFP- DKO | TGTGCAAGCAGCTCCCTTAACCAAGACACCCAGTACTTT | 1 | TRBV16*00 | | TRBJ2-5*00 | 0.28% |
| CD4+YFP- DKO | TGTGCAAGCAGCTGGGGGGGCGGTGCAGAAACGCTGTATTTT | 1 | TRBV16*00 | TRBD2*00 | TRBJ2-3*00 | 0.28% |
| CD4+YFP- DKO | TGTGCAAGCAGCTTAGAGAGGGGGCACACAGAAGTCTTCTTT | 1 | TRBV16*00 | TRBD1*00 | TRBJ1-1*00 | 0.28% |
| CD4+YFP- DKO | TGTGCAAGCAGCTTAGAGGGAAACTCCGACTACACCTTC | 1 | TRBV16*00 | | TRBJ1-2*00 | 0.28% |
| CD4+YFP- DKO | TGTGCAAGCAGCTTAGAGGGAGGCACCCAGTACTTT | 1 | TRBV16*00 | | TRBJ2-5*00 | 0.28% |
| CD4+YFP- DKO | TGTGCAAGCAGCTTAGAGGGATACACCTTGTACTTT | 1 | TRBV16*00 | | TRBJ2-4*00 | 0.28% |
| CD4+YFP- DKO | TGTGCAAGCAGCTTAGAGGGTCAAAACACCTTGTACTTT | 1 | TRBV16*00 | | TRBJ2-4*00 | 0.28% |
| CD4+YFP- DKO | TGTGCAAGCAGCTTAGATGAACATAACTCCGACTACACCTTC | 1 | TRBV16*00 | | TRBJ1-2*00 | 0.28% |
| CD4+YFP- DKO | TGTGCAAGCAGCTTAGGACAGGGAGAAGAACAGTACTTC | 1 | TRBV16*00 | TRBD1*00 | TRBJ2-7*00 | 0.28% |
| CD4+YFP- DKO | TGTGCAAGCAGCTTAGGAGGGGGGCAAGACACCCAGTACTTT | 1 | TRBV16*00 | TRBD1*00 | TRBJ2-5*00 | 0.28% |
| CD4+YFP- DKO | TGTGCAAGCAGCTTAGGGGCAAACACAGAAGTCTTCTTT | 1 | TRBV16*00 | | TRBJ1-1*00 | 0.28% |
| CD4+YFP- DKO | TGTGCAAGCAGCTTAGGGGGTAGTGCAGAAACGCTGTATTTT | 1 | TRBV16*00 | | TRBJ2-3*00 | 0.28% |
| CD4+YFP- DKO | TGTGCAAGCAGCTTAGTCGGTAACCAAGACACCCAGTACTTT | 1 | TRBV16*00 | | TRBJ2-5*00 | 0.28% |
| CD4+YFP- DKO | TGTGCAAGCAGCTTAGTTGGCAACCAGGCTCCGCTTTTT | 1 | TRBV16*00 | | TRBJ1-5*00 | 0.28% |
| CD4+YFP- DKO | TGTGCAAGCAGCTTCCAGGGGGAAAACACCTTGTACTTT | 1 | TRBV16*00 | TRBD1*00 | TRBJ2-4*00 | 0.28% |
| CD4+YFP- DKO | TGTGCAAGCAGCTTGGCAGCAAACACAGAAGTCTTCTTT | 1 | TRBV16*00 | | TRBJ1-1*00 | 0.28% |
| CD4+YFP- DKO | TGTGCAAGCAGCTTGGCCGGGAGAGACACCCAGTACTTT | 1 | TRBV16*00 | TRBD1*00 | TRBJ2-5*00 | 0.28% |
| CD4+YFP- DKO | TGTGCAAGCCGCAGCACATATAATTCGCCCCTCTACTTT | 1 | TRBV16*00 | | TRBJ1-6*00 | 0.28% |
| CD4+YFP- DKO | TGTGCCAGCAAAGACTGGGGAGTGCAGAAACGCTGTATTTT | 1 | TRBV13-1*00 | TRBD2*00 | TRBJ2-3*00 | 0.28% |
| CD4+YFP- DKO | TGTGCCAGCACCTTCCAAGACACCCAGTACTTT | 1 | TRBV15*00 | | TRBJ2-5*00 | 0.28% |
| CD4+YFP- DKO | TGTGCCAGCACTCAAGGGGGCTATGCTGAGCAGTTCTTC | 1 | TRBV13-1*00 | TRBD1*00 | TRBJ2-1*00 | 0.28% |
| CD4+YFP- DKO | TGTGCCAGCAGAAGAAGGCTCCGCTTTTT | 1 | TRBV15*00 | | TRBJ1-5*00 | 0.28% |
| CD4+YFP- DKO | TGTGCCAGCAGATCAAACACAGAAGTCTTCTTT | 1 | TRBV4*00 | | TRBJ1-1*00 | 0.28% |
| CD4+YFP- DKO | TGTGCCAGCAGCAGACTGGGGGGGGAGCTCCTATGAACAGTACTTC | 1 | TRBV3*00 | TRBD2*00 | TRBJ2-7*00 | 0.28% |
| CD4+YFP- DKO | TGTGCCAGCAGCCAAATCCAGGGTAACCAAGACACCCAGTACTTT | 1 | TRBV5*00 | TRBD1*00 | TRBJ2-5*00 | 0.28% |
| CD4+YFP- DKO | TGTGCCAGCAGCCAACAATCTAGTCAAAACACCTTGTACTTT | 1 | TRBV2*00 | | TRBJ2-4*00 | 0.28% |
| CD4+YFP- DKO | TGTGCCAGCAGCCAAGAAGTGAACACCCAGTACTTT | 1 | TRBV5*00 | | TRBJ2-5*00 | 0.28% |
| CD4+YFP- DKO | TGTGCCAGCAGCCAAGATAGGGGGAGAGACACCCAGTACTTT | 1 | TRBV5*00 | TRBD1*00 | TRBJ2-5*00 | 0.28% |
| CD4+YFP- DKO | TGTGCCAGCAGCCAAGATCGACAATATGAACAGTACTTC | 1 | TRBV5*00 | | TRBJ2-7*00 | 0.28% |
| CD4+YFP- DKO | TGTGCCAGCAGCCAAGATCGGAACACCGGGCAGCTCTACTTT | 1 | TRBV5*00 | | TRBJ2-2*00 | 0.28% |
| CD4+YFP- DKO | TGTGCCAGCAGCCAAGATGGGGACAACAAGACACCCAGTACTTT | 1 | TRBV2*00 | TRBD1*00 | TRBJ2-5*00 | 0.28% |
| CD4+YFP- DKO | TGTGCCAGCAGCCAAGCGTACAGTTTTAACCAAGACACCCAGTACTTT | 1 | TRBV2*00 | | TRBJ2-5*00 | 0.28% |
| CD4+YFP- DKO | TGTGCCAGCAGCCAAGGGAGGAATTCTGGAAATACGCTCTATTTT | 1 | TRBV2*00 | | TRBJ1-3*00 | 0.28% |
| CD4+YFP- DKO | TGTGCCAGCAGCCAAGGGGGACACTCTTCTGGAAATACGCTCTATTTT | 1 | TRBV2*00 | TRBD1*00 | TRBJ1-3*00 | 0.28% |
| CD4+YFP- DKO | TGTGCCAGCAGCCACAAACAATAGTCAAAACACCTTGTACTTT | 1 | TRBV2*00 | | TRBJ2-4*00 | 0.28% |
| CD4+YFP- DKO | TGTGCCAGCAGCCAGACATCTAGTGCAGAAACGCTGTATTTT | 1 | TRBV2*00 | | TRBJ2-3*00 | 0.28% |
| CD4+YFP- DKO | TGTGCCAGCAGCCAGCTAAACTCCGACTACACCTTC | 1 | TRBV2*00 | | TRBJ1-2*00 | 0.28% |
| CD4+YFP- DKO | TGTGCCAGCAGCCAGGAATATGAACAGTACTTC | 1 | TRBV2*00 | | TRBJ2-7*00 | 0.28% |
| CD4+YFP- DKO | TGTGCCAGCAGCCCACAGGGCTCTGGAAATACGCTCTATTTT | 1 | TRBV4*00 | TRBD1*00 | TRBJ1-3*00 | 0.28% |
| CD4+YFP- DKO | TGTGCCAGCAGCCCCGGGACAGGGACCAACGAAAGATTATTTTTC | 1 | TRBV4*00 | TRBD1*00 | TRBJ1-4*00 | 0.28% |
| CD4+YFP- DKO | TGTGCCAGCAGCCCCTCAGGGTTGTATGAACAGTACTTC | 1 | TRBV2*00 | TRBD1*00 | TRBJ2-7*00 | 0.28% |
| CD4+YFP- DKO | TGTGCCAGCAGCCCCTCCTCCTATGAACAGTACTTC | 1 | TRBV2*00 | | TRBJ2-7*00 | 0.28% |
| CD4+YFP- DKO | TGTGCCAGCAGCCCGGGACACCAAGACACCCAGTACTTT | 1 | TRBV3*00 | TRBD1*00 | TRBJ2-5*00 | 0.28% |
| CD4+YFP- DKO | TGTGCCAGCAGCCCTCACAGGTTCTAGTGCAGAAACGCTGCATTTT | 1 | TRBV3*00 | TRBD1*00 | TRBJ2-3*00 | 0.28% |
| CD4+YFP- DKO | TGTGCCAGCAGCCGGACTGGGGGGTTTGCTGAGCAGTTCTTC | 1 | TRBV4*00 | TRBD2*00 | TRBJ2-1*00 | 0.28% |
| CD4+YFP- DKO | TGTGCCAGCAGCCTAAATTCCTATAATTCGCCCCTCTACTTT | 1 | TRBV15*00 | | TRBJ1-6*00 | 0.28% |
| CD4+YFP- DKO | TGTGCCAGCAGCCTACAGGTCTCCTATGAACAGTACTTC | 1 | TRBV2*00 | TRBD1*00 | TRBJ2-7*00 | 0.28% |

| | | | | | | |
|---|---|---|---|---|---|---|
| CD4+YFP- DKO | TGTGCCAGCAGCCTCAGAGGTGGCCAAGACACCCAGTACTTT | 1 | TRBV2*00 | | TRBJ2-5*00 | 0.28% |
| CD4+YFP- DKO | TGTGCCAGCAGCCTGGGGGGGCAAGACACCCAGTACTTT | 1 | TRBV5*00 | TRBD2*00 | TRBJ2-5*00 | 0.28% |
| CD4+YFP- DKO | TGTGCCAGCAGCGGACAGGGAAACGAAAGATTATTTTTC | 1 | TRBV13-3*00 | TRBD1*00 | TRBJ1-4*00 | 0.28% |
| CD4+YFP- DKO | TGTGCCAGCAGCTAATGGGACGGAAGTGCAGAAACGCTGTATTTT | 1 | TRBV4*00 | TRBD1*00 | TRBJ2-3*00 | 0.28% |
| CD4+YFP- DKO | TGTGCCAGCAGCTACAGAATTCAAAACACCTTGTACTTT | 1 | TRBV4*00 | | TRBJ2-4*00 | 0.28% |
| CD4+YFP- DKO | TGTGCCAGCAGCTCCAGCTCCTATGAACAGTACTTC | 1 | TRBV3*00 | | TRBJ2-7*00 | 0.28% |
| CD4+YFP- DKO | TGTGCCAGCAGCTCGACTGGGGGAAAAACACCTTGTACTTT | 1 | TRBV3*00 | TRBD2*00 | TRBJ2-4*00 | 0.28% |
| CD4+YFP- DKO | TGTGCCAGCAGCTCGAGCTCCTATGAACAGTACTTC | 1 | TRBV25*00 | | TRBJ2-7*00 | 0.28% |
| CD4+YFP- DKO | TGTGCCAGCAGCTCGTCCAGGGGTAGTGCAGAAACGCTGTATTTT | 1 | TRBV3*00 | TRBD1*00 | TRBJ2-3*00 | 0.28% |
| CD4+YFP- DKO | TGTGCCAGCAGCTCTACTGGGTCTAGTGCAGAAACGCTGTATTTT | 1 | TRBV3*00 | TRBD2*00 | TRBJ2-3*00 | 0.28% |
| CD4+YFP- DKO | TGTGCCAGCAGCTGGGACAGTGCAGAAACGCTGTATTTT | 1 | TRBV3*00 | TRBD1*00 | TRBJ2-3*00 | 0.28% |
| CD4+YFP- DKO | TGTGCCAGCAGCTTAACTGGGGGGGGAGACACCCAGTACTTT | 1 | TRBV3*00 | TRBD2*00 | TRBJ2-5*00 | 0.28% |
| CD4+YFP- DKO | TGTGCCAGCAGCTTACGGGGGGACTATGCTGAGCAGTTCTTC | 1 | TRBV3*00 | TRBD2*00 | TRBJ2-1*00 | 0.28% |
| CD4+YFP- DKO | TGTGCCAGCAGCTTAGAAGGGGCAAACACCGGGCAGCTCTACTTT | 1 | TRBV3*00 | | TRBJ2-2*00 | 0.28% |
| CD4+YFP- DKO | TGTGCCAGCAGCTTAGACAGCTATGCTGAGCAGTTCTTC | 1 | TRBV3*00 | | TRBJ2-1*00 | 0.28% |
| CD4+YFP- DKO | TGTGCCAGCAGCTTAGAGACAGATTATGAACAGTACTTC | 1 | TRBV3*00 | TRBD1*00 | TRBJ2-7*00 | 0.28% |
| CD4+YFP- DKO | TGTGCCAGCAGCTTAGAGCAAAACTCCGACTACACCTTC | 1 | TRBV3*00 | | TRBJ1-2*00 | 0.28% |
| CD4+YFP- DKO | TGTGCCAGCAGCTTAGAGGGAGACGGAAATACGCTCTATTTT | 1 | TRBV3*00 | | TRBJ1-3*00 | 0.28% |
| CD4+YFP- DKO | TGTGCCAGCAGCTTAGCGCCTACTAGTCAAAACACCTTGTACTTT | 1 | TRBV3*00 | | TRBJ2-4*00 | 0.28% |
| CD4+YFP- DKO | TGTGCCAGCAGCTTATTGGACACCGGGCAGCTCTACTTT | 1 | TRBV3*00 | | TRBJ2-2*00 | 0.28% |
| CD4+YFP- DKO | TGTGCCAGCAGCTTCCTTAACTATGCTGAGCAGTTCTTC | 1 | TRBV3*00 | | TRBJ2-1*00 | 0.28% |
| CD4+YFP- DKO | TGTGCCAGCAGCTTTAACCAAGACACCCAGTACTTT | 1 | TRBV13-3*00 | | TRBJ2-5*00 | 0.28% |
| CD4+YFP- DKO | TGTGCCAGCAGCTTTCAGGGGGCGCAAGTCTTCTTT | 1 | TRBV4*00 | TRBD1*00 | TRBJ1-1*00 | 0.28% |
| CD4+YFP- DKO | TGTGCCAGCAGGACCCGGGGGCCCACCGGGCAGCTCTACTTT | 1 | TRBV13-3*00 | TRBD1*00 | TRBJ2-2*00 | 0.28% |
| CD4+YFP- DKO | TGTGCCAGCAGGCTAAGAACAGAAGTCTTCTTT | 1 | TRBV4*00 | | TRBJ1-1*00 | 0.28% |
| CD4+YFP- DKO | TGTGCCAGCAGGGGGGAAAGGGGTGCAGAAACGCTGTATTTT | 1 | TRBV13-3*00 | TRBD1*00 | TRBJ2-3*00 | 0.28% |
| CD4+YFP- DKO | TGTGCCAGCAGTAACTGGGGAACCTATGAACAGTACTTC | 1 | TRBV13-3*00 | TRBD2*00 | TRBJ2-7*00 | 0.28% |
| CD4+YFP- DKO | TGTGCCAGCAGTAAGACAATTTCCTATAATTCGCCCCTCTACTTT | 1 | TRBV19*00 | | TRBJ1-6*00 | 0.28% |
| CD4+YFP- DKO | TGTGCCAGCAGTAGGACAGACGGCACCGGGCAGCTCTACTTT | 1 | TRBV14*00 | TRBD1*00 | TRBJ2-2*00 | 0.28% |
| CD4+YFP- DKO | TGTGCCAGCAGTAGGACCCAAAACACCTTGTACTTT | 1 | TRBV14*00 | | TRBJ2-4*00 | 0.28% |
| CD4+YFP- DKO | TGTGCCAGCAGTAGGTACCAAGACACCCAGTACTTT | 1 | TRBV19*00 | | TRBJ2-5*00 | 0.28% |
| CD4+YFP- DKO | TGTGCCAGCAGTATAAACTGGGATGCTGAGCAGTTCTTC | 1 | TRBV19*00 | TRBD2*00 | TRBJ2-1*00 | 0.28% |
| CD4+YFP- DKO | TGTGCCAGCAGTATAATCTCTAGTGCAGAAACGCTGTATTTT | 1 | TRBV19*00 | | TRBJ2-3*00 | 0.28% |
| CD4+YFP- DKO | TGTGCCAGCAGTATAATGGGAGAGGAACAGTACTTC | 1 | TRBV19*00 | | TRBJ2-7*00 | 0.28% |
| CD4+YFP- DKO | TGTGCCAGCAGTATAGGGACTGGGGGTGCAGAAACGCTGTATTTT | 1 | TRBV19*00 | TRBD2*00 | TRBJ2-3*00 | 0.28% |
| CD4+YFP- DKO | TGTGCCAGCAGTATGAGGGCCGAAAGATTATTTTTC | 1 | TRBV19*00 | | TRBJ1-4*00 | 0.28% |
| CD4+YFP- DKO | TGTGCCAGCAGTATGTCCTGGGCAAACACCGGGCAGCTCTACTTT | 1 | TRBV19*00 | | TRBJ2-2*00 | 0.28% |
| CD4+YFP- DKO | TGTGCCAGCAGTCAAGACCAAGACACCCAGTACTTT | 1 | TRBV2*00 | | TRBJ2-5*00 | 0.28% |
| CD4+YFP- DKO | TGTGCCAGCAGTCCAACAGCAAACTCCGACTACACCTTC | 1 | TRBV13-1*00 | | TRBJ1-2*00 | 0.28% |
| CD4+YFP- DKO | TGTGCCAGCAGTCCCCTGGGGGGCGGGAACACCTTGTACTTT | 1 | TRBV26*00 | TRBD2*00 | TRBJ2-4*00 | 0.28% |
| CD4+YFP- DKO | TGTGCCAGCAGTCCCGCCTATGAACAGTACTTC | 1 | TRBV19*00 | | TRBJ2-7*00 | 0.28% |
| CD4+YFP- DKO | TGTGCCAGCAGTCGACAAAATTCTGGAAATACGCGCTATTTT | 1 | TRBV19*00 | | TRBJ1-3*00 | 0.28% |
| CD4+YFP- DKO | TGTGCCAGCAGTCGCCGGGACATCTATGCTGAGCAGTTCTTC | 1 | TRBV26*00 | TRBD1*00 | TRBJ2-1*00 | 0.28% |
| CD4+YFP- DKO | TGTGCCAGCAGTCGGACAGAAGACACCCAGTACTTT | 1 | TRBV24*00 | TRBD1*00 | TRBJ2-5*00 | 0.28% |
| CD4+YFP- DKO | TGTGCCAGCAGTCGGACAGGGGGCGCCGACACCCAGTACTTT | 1 | TRBV14*00 | TRBD1*00 | TRBJ2-5*00 | 0.28% |
| CD4+YFP- DKO | TGTGCCAGCAGTCGGGACAGGGGTGAAGACACCCAGTACTTT | 1 | TRBV13-3*00 | TRBD1*00 | TRBJ2-5*00 | 0.28% |
| CD4+YFP- DKO | TGTGCCAGCAGTCTCAGAATCAATTCTGGAAATACGCTCTATTTT | 1 | TRBV26*00 | | TRBJ1-3*00 | 0.28% |
| CD4+YFP- DKO | TGTGCCAGCAGTCTGACAGGGGGCTATGCTGAGCAGTTCTTC | 1 | TRBV26*00 | TRBD1*00 | TRBJ2-1*00 | 0.28% |
| CD4+YFP- DKO | TGTGCCAGCAGTCTGCCAGGGCCCAACGAAAGATTATTTTTC | 1 | TRBV26*00 | TRBD1*00 | TRBJ1-4*00 | 0.28% |
| CD4+YFP- DKO | TGTGCCAGCAGTCTGGACAGCTATGAACAGTACTTC | 1 | TRBV26*00 | TRBD1*00 | TRBJ2-7*00 | 0.28% |
| CD4+YFP- DKO | TGTGCCAGCAGTCTGGCAGGGGACTCCGACTACACCTTC | 1 | TRBV26*00 | TRBD1*00 | TRBJ1-2*00 | 0.28% |
| CD4+YFP- DKO | TGTGCCAGCAGTCTGGGAAACCAAGACACCCAGTACTTT | 1 | TRBV26*00 | | TRBJ2-5*00 | 0.28% |
| CD4+YFP- DKO | TGTGCCAGCAGTCTGGGGTATGCTGAGCAGTTCTTC | 1 | TRBV26*00 | | TRBJ2-1*00 | 0.28% |
| CD4+YFP- DKO | TGTGCCAGCAGTCTGTCGAGTCACACAGAAGTCTTCTTT | 1 | TRBV26*00 | | TRBJ1-1*00 | 0.28% |
| CD4+YFP- DKO | TGTGCCAGCAGTCTTTCAGGAACCTATGAACAGTACTTC | 1 | TRBV24*00 | | TRBJ2-7*00 | 0.28% |
| CD4+YFP- DKO | TGTGCCAGCAGTGAAAAACTGGGGGGAGAGGACACCCAGTACTTT | 1 | TRBV13-3*00 | TRBD2*00 | TRBJ2-5*00 | 0.28% |
| CD4+YFP- DKO | TGTGCCAGCAGTGAACTGGGTAGTCAAAACACCTTGTACTTT | 1 | TRBV14*00 | TRBD2*00 | TRBJ2-4*00 | 0.28% |
| CD4+YFP- DKO | TGTGCCAGCAGTGACAGGGGAGGATTGGAACAGTACTTC | 1 | TRBV13-1*00 | TRBD1*00 | TRBJ2-7*00 | 0.28% |
| CD4+YFP- DKO | TGTGCCAGCAGTGACCCCGACAGGGGTGAACAGTACTTC | 1 | TRBV13-1*00 | TRBD1*00 | TRBJ2-7*00 | 0.28% |
| CD4+YFP- DKO | TGTGCCAGCAGTGACGACAAAAGTCAAAACACCTTGTACTTT | 1 | TRBV13-1*00 | | TRBJ2-4*00 | 0.28% |
| CD4+YFP- DKO | TGTGCCAGCAGTGAGGACAGGGGTGAACAGTACTTC | 1 | TRBV13-1*00 | TRBD1*00 | TRBJ2-7*00 | 0.28% |
| CD4+YFP- DKO | TGTGCCAGCAGTGAGGGACATTATGAACAGTACTTC | 1 | TRBV13-3*00 | TRBD1*00 | TRBJ2-7*00 | 0.28% |
| CD4+YFP- DKO | TGTGCCAGCAGTGAGGGTGCAGAAACGCTGTATTTT | 1 | TRBV13-1*00 | | TRBJ2-3*00 | 0.28% |
| CD4+YFP- DKO | TGTGCCAGCAGTGATACCAGAAACTCCGACTACACCTTC | 1 | TRBV13-3*00 | | TRBJ1-2*00 | 0.28% |
| CD4+YFP- DKO | TGTGCCAGCAGTGATCCAGACACAGAAGTCTTCTTT | 1 | TRBV13-1*00 | | TRBJ1-1*00 | 0.28% |
| CD4+YFP- DKO | TGTGCCAGCAGTGATCCGGGGAGTCAAAACACCTTGTACTTT | 1 | TRBV13-1*00 | TRBD1*00 | TRBJ2-4*00 | 0.28% |
| CD4+YFP- DKO | TGTGCCAGCAGTGATGACCGAGACACCCAGTACTTT | 1 | TRBV13-3*00 | | TRBJ2-5*00 | 0.28% |
| CD4+YFP- DKO | TGTGCCAGCAGTGATGAGTATGCTGAGCAGTTCTTC | 1 | TRBV13-3*00 | | TRBJ2-1*00 | 0.28% |
| CD4+YFP- DKO | TGTGCCAGCAGTGATGCAAGGGTCTATGCTGAGCAGTTCTTC | 1 | TRBV13-1*00 | | TRBJ2-1*00 | 0.28% |
| CD4+YFP- DKO | TGTGCCAGCAGTGATGCAGGACAGTATGAACAGTACTTC | 1 | TRBV13-1*00 | TRBD1*00 | TRBJ2-7*00 | 0.28% |
| CD4+YFP- DKO | TGTGCCAGCAGTGATGCCCGGGACAACTATGCTGAGCAGTTCTTC | 1 | TRBV13-1*00 | TRBD1*00 | TRBJ2-1*00 | 0.28% |
| CD4+YFP- DKO | TGTGCCAGCAGTGATGCGGACTCCGACTACACCTTC | 1 | TRBV13-1*00 | | TRBJ1-2*00 | 0.28% |
| CD4+YFP- DKO | TGTGCCAGCAGTGATGCTGGGGGGGTCTATGCTGAGCAGTTCTTC | 1 | TRBV13-1*00 | TRBD2*00 | TRBJ2-1*00 | 0.28% |
| CD4+YFP- DKO | TGTGCCAGCAGTGATGGGGGGAGGGAGAACACCTTGTACTTT | 1 | TRBV13-1*00 | TRBD1*00 | TRBJ2-4*00 | 0.28% |
| CD4+YFP- DKO | TGTGCCAGCAGTGATGTACGCGGTCAAAACACCTTGTACTTT | 1 | TRBV13-1*00 | | TRBJ2-4*00 | 0.28% |

| | | | | | | |
|---|---|---|---|---|---|---|
| CD4+YFP- DKO | TGTGCCAGCAGTGATTTCAGTGAACAGTACTTC | 1 | TRBV13-1*00 | | TRBJ2-7*00 | 0.28% |
| CD4+YFP- DKO | TGTGCCAGCAGTGGACCCGCAGAAACGCTGTATTTT | 1 | TRBV15*00 | | TRBJ2-3*00 | 0.28% |
| CD4+YFP- DKO | TGTGCCAGCAGTGGACTGGGGGGTGCAGAAACGCTGTATTTT | 1 | TRBV13-1*00 | TRBD2*00 | TRBJ2-3*00 | 0.28% |
| CD4+YFP- DKO | TGTGCCAGCAGTGGCAACTCCGACTACACCTTC | 1 | TRBV13-1*00 | | TRBJ1-2*00 | 0.28% |
| CD4+YFP- DKO | TGTGCCAGCAGTGGGACTGGAGGAGACACCCAGTACTTT | 1 | TRBV13-3*00 | TRBD2*00 | TRBJ2-5*00 | 0.28% |
| CD4+YFP- DKO | TGTGCCAGCAGTGGGACTGGGTCAAACACCTTGTACTTT | 1 | TRBV13-1*00 | TRBD2*00 | TRBJ2-4*00 | 0.28% |
| CD4+YFP- DKO | TGTGCCAGCAGTGGGGGGAGTCAAAACACCTTGTACTTT | 1 | TRBV13-1*00 | TRBD1*00 | TRBJ2-4*00 | 0.28% |
| CD4+YFP- DKO | TGTGCCAGCAGTGGTGGAAATTCTGGAAATACGCTCTATTTT | 1 | TRBV13-3*00 | | TRBJ1-3*00 | 0.28% |
| CD4+YFP- DKO | TGTGCCAGCAGTGTCGAGGGGGGCCAAGACACCCAGTACTTT | 1 | TRBV13-1*00 | TRBD1*00 | TRBJ2-5*00 | 0.28% |
| CD4+YFP- DKO | TGTGCCAGCAGTGTGGGGGACCAAGACACCCAGTACTTT | 1 | TRBV13-1*00 | TRBD2*00 | TRBJ2-5*00 | 0.28% |
| CD4+YFP- DKO | TGTGCCAGCAGTTCAGCCAACTCCGACTACACCTTC | 1 | TRBV14*00 | | TRBJ1-2*00 | 0.28% |
| CD4+YFP- DKO | TGTGCCAGCAGTTTAAGCTATGAACAGTACTTC | 1 | TRBV15*00 | | TRBJ2-7*00 | 0.28% |
| CD4+YFP- DKO | TGTGCCAGCAGTTTACTGGGGGATATGAACAGTACTTC | 1 | TRBV15*00 | TRBD2*00 | TRBJ2-7*00 | 0.28% |
| CD4+YFP- DKO | TGTGCCAGCAGTTTAGAAGAAACGCTGTATTTT | 1 | TRBV15*00 | | TRBJ2-3*00 | 0.28% |
| CD4+YFP- DKO | TGTGCCAGCAGTTTAGACCTTAACCAAGACACCCAGTACTTT | 1 | TRBV15*00 | | TRBJ2-5*00 | 0.28% |
| CD4+YFP- DKO | TGTGCCAGCAGTTTAGGACAGGGCCTTTATGAACAGTACTTC | 1 | TRBV15*00 | TRBD1*00 | TRBJ2-7*00 | 0.28% |
| CD4+YFP- DKO | TGTGCCAGCAGTTTAGGCAATCAAGACACCCAGTACTTT | 1 | TRBV15*00 | | TRBJ2-5*00 | 0.28% |
| CD4+YFP- DKO | TGTGCCAGCAGTTTAGTCAGGGGGGAACAGTACTTC | 1 | TRBV15*00 | TRBD1*00 | TRBJ2-7*00 | 0.28% |
| CD4+YFP- DKO | TGTGCCAGCAGTTTATCAAATGCAAACACCGGGCAGCTCTACTTT | 1 | TRBV15*00 | | TRBJ2-2*00 | 0.28% |
| CD4+YFP- DKO | TGTGCCAGCAGTTTATCAGGGGAGGAACAGTACTTC | 1 | TRBV15*00 | TRBD1*00 | TRBJ2-7*00 | 0.28% |
| CD4+YFP- DKO | TGTGCCAGCAGTTTCCAATCCTATAATTCGCCCCTCTACTTT | 1 | TRBV14*00 | | TRBJ1-6*00 | 0.28% |
| CD4+YFP- DKO | TGTGCCAGCAGTTTCGCGGACCAAGACACCCAGTACTTT | 1 | TRBV15*00 | | TRBJ2-5*00 | 0.28% |
| CD4+YFP- DKO | TGTGCCAGCAGTTTCTCCGGGAGCTCCTATGAACAGTACTTC | 1 | TRBV14*00 | | TRBJ2-7*00 | 0.28% |
| CD4+YFP- DKO | TGTGCCAGCAGTTTGGACTGGGGGAGCTATGCTGAGCAGTTCTTC | 1 | TRBV15*00 | TRBD2*00 | TRBJ2-1*00 | 0.28% |
| CD4+YFP- DKO | TGTGCCAGCAGTTTGGCAAACACCGGGCAGCTCTACTTT | 1 | TRBV15*00 | | TRBJ2-2*00 | 0.28% |
| CD4+YFP- DKO | TGTGCCAGCAGTTTGTCAGGGGGCAACGAAAGATTATTTTTC | 1 | TRBV26*00 | TRBD1*00 | TRBJ1-4*00 | 0.28% |
| CD4+YFP- DKO | TGTGCCAGCAGTTTTTGGCTCGAAAGATTATTTTTC | 1 | TRBV14*00 | | TRBJ1-4*00 | 0.28% |
| CD4+YFP- DKO | TGTGCCAGCGAGGGTTTCTCCTATGAACAGTACTTC | 1 | TRBV19*00 | | TRBJ2-7*00 | 0.28% |
| CD4+YFP- DKO | TGTGCCAGCGCCAGAAACAACCAGGCTCCGCTTTTT | 1 | TRBV13-3*00 | | TRBJ1-5*00 | 0.28% |
| CD4+YFP- DKO | TGTGCCAGCGGATGGACAGGGAATTCTGGAAATACGCTCTATTTT | 1 | TRBV13-2*00 | TRBD1*00 | TRBJ1-3*00 | 0.28% |
| CD4+YFP- DKO | TGTGCCAGCGGCCTTCAGGGGGGAGAACAGTACTTC | 1 | TRBV13-2*00 | TRBD1*00 | TRBJ2-7*00 | 0.28% |
| CD4+YFP- DKO | TGTGCCAGCGGGACAGAAGTCTTCTTT | 1 | TRBV13-2*00 | | TRBJ1-1*00 | 0.28% |
| CD4+YFP- DKO | TGTGCCAGCGGGGACTGGGGAAACACCGGGCAGCTCTACATT | 1 | TRBV13-2*00 | TRBD2*00 | TRBJ2-2*00 | 0.28% |
| CD4+YFP- DKO | TGTGCCAGCGGGGACTGGGGAAACACCGGGCAGCTCTTCTTT | 1 | TRBV13-2*00 | TRBD2*00 | TRBJ2-2*00 | 0.28% |
| CD4+YFP- DKO | TGTGCCAGCGGGGGGACAGGAAGTGCAGAAACGCTGTATTTT | 1 | TRBV13-2*00 | TRBD1*00 | TRBJ2-3*00 | 0.28% |
| CD4+YFP- DKO | TGTGCCAGCGGTCACAGAAATTCTGGAAATACGCTCTATTTT | 1 | TRBV13-2*00 | | TRBJ1-3*00 | 0.28% |
| CD4+YFP- DKO | TGTGCCAGCGGTCTACAGACCTATGAACAGTACTTC | 1 | TRBV13-2*00 | | TRBJ2-7*00 | 0.28% |
| CD4+YFP- DKO | TGTGCCAGCGGTGAACAATTCTATGCTGAGCAGTTCTTC | 1 | TRBV13-2*00 | | TRBJ2-1*00 | 0.28% |
| CD4+YFP- DKO | TGTGCCAGCGGTGAACTGGGGAGTCAAAACACCTTGTACTTT | 1 | TRBV13-2*00 | TRBD2*00 | TRBJ2-4*00 | 0.28% |
| CD4+YFP- DKO | TGTGCCAGCGGTGACCCGGGGAGGGGTGCAGAAACGCTGTATTTT | 1 | TRBV13-2*00 | TRBD1*00 | TRBJ2-3*00 | 0.28% |
| CD4+YFP- DKO | TGTGCCAGCGGTGATAACAACCAGGCTCCGCTTTTT | 1 | TRBV13-2*00 | | TRBJ1-5*00 | 0.28% |
| CD4+YFP- DKO | TGTGCCAGCGGTGATACCAGTTCTGGAAATACGCTCTATTTT | 1 | TRBV13-2*00 | | TRBJ1-3*00 | 0.28% |
| CD4+YFP- DKO | TGTGCCAGCGGTGATAGGACATATAGTGCAGAAACGCTGTATTTT | 1 | TRBV13-2*00 | TRBD1*00 | TRBJ2-3*00 | 0.28% |
| CD4+YFP- DKO | TGTGCCAGCGGTGATCACAGTCAAGACACCCAGTACTTT | 1 | TRBV13-2*00 | | TRBJ2-5*00 | 0.28% |
| CD4+YFP- DKO | TGTGCCAGCGGTGATGACAGGAGGGAAGTCTTCTTT | 1 | TRBV13-2*00 | TRBD1*00 | TRBJ1-1*00 | 0.28% |
| CD4+YFP- DKO | TGTGCCAGCGGTGATGCAAACCAGGCTCCGCTTTTT | 1 | TRBV13-2*00 | | TRBJ1-5*00 | 0.28% |
| CD4+YFP- DKO | TGTGCCAGCGGTGATGCAAGGGGCACAGAAGTCTTCTTT | 1 | TRBV13-2*00 | TRBD1*00 | TRBJ1-1*00 | 0.28% |
| CD4+YFP- DKO | TGTGCCAGCGGTGATGCAGGGGGCGACTCCGACTACACCTTC | 1 | TRBV13-2*00 | TRBD1*00 | TRBJ1-2*00 | 0.28% |
| CD4+YFP- DKO | TGTGCCAGCGGTGATGCCGGGACTGGGGGAGACACCCAGTACTTT | 1 | TRBV13-2*00 | TRBD2*00 | TRBJ2-5*00 | 0.28% |
| CD4+YFP- DKO | TGTGCCAGCGGTGATGCGAGAGGCAACTCCGACTACACCTTC | 1 | TRBV13-2*00 | | TRBJ1-2*00 | 0.28% |
| CD4+YFP- DKO | TGTGCCAGCGGTGATGCGGGGGGCTCCAACGAAAGATTATTTTTC | 1 | TRBV13-2*00 | TRBD2*00 | TRBJ1-4*00 | 0.28% |
| CD4+YFP- DKO | TGTGCCAGCGGTGATGGACAGGGAAACACAGAAGTCTTCTTT | 1 | TRBV13-2*00 | TRBD1*00 | TRBJ1-1*00 | 0.28% |
| CD4+YFP- DKO | TGTGCCAGCGGTGATGGGACAGGGTATGCTGAGCAGTTCTTC | 1 | TRBV13-2*00 | TRBD1*00 | TRBJ2-1*00 | 0.28% |
| CD4+YFP- DKO | TGTGCCAGCGGTGATGGGGGGTCTAGTGCAGAAACGCTGTATTTT | 1 | TRBV13-2*00 | TRBD1*00 | TRBJ2-3*00 | 0.28% |
| CD4+YFP- DKO | TGTGCCAGCGGTGATTGGGGGGTGAGGGCTGAGCAGTTCTTC | 1 | TRBV13-2*00 | TRBD2*00 | TRBJ2-1*00 | 0.28% |
| CD4+YFP- DKO | TGTGCCAGCGGTGCCGGGACAGCAAACTCCGACTACACCTTC | 1 | TRBV13-2*00 | TRBD1*00 | TRBJ1-2*00 | 0.28% |
| CD4+YFP- DKO | TGTGCCAGCGGTGGACTGGGGGGTGAGCAGTTCTTC | 1 | TRBV13-2*00 | TRBD2*00 | TRBJ2-1*00 | 0.28% |
| CD4+YFP- DKO | TGTGCCAGCGGTGGAGGGACAGCAAACTCCGACTACACCTTC | 1 | TRBV13-2*00 | TRBD1*00 | TRBJ1-2*00 | 0.28% |
| CD4+YFP- DKO | TGTGCCAGCGGTGGGGGAGATAACTATGCTGAGCAGTTCTTC | 1 | TRBV13-2*00 | | TRBJ2-1*00 | 0.28% |
| CD4+YFP- DKO | TGTGCCAGCTCTCAACAAAACCAAGACACCCAGTACTTT | 1 | TRBV12-1*00 | | TRBJ2-5*00 | 0.28% |
| CD4+YFP- DKO | TGTGCCAGCTCTCCCCGGGACAGTTCCTATAATTCGCCCCTCTACTTT | 1 | TRBV12-2*00 | TRBD1*00 | TRBJ1-6*00 | 0.28% |
| CD4+YFP- DKO | TGTGCCAGCTCTCTCACAAACCAAGACACCCAGTACTTT | 1 | TRBV12-1*00 | | TRBJ2-5*00 | 0.28% |
| CD4+YFP- DKO | TGTGCCAGCTCTCTCCTCTATGAACAGTACTTC | 1 | TRBV12-1*00 | | TRBJ2-7*00 | 0.28% |
| CD4+YFP- DKO | TGTGCCAGCTCTCTCGAGGGGGCGCCGAGTGCAGAAACGCTGTATTTT | 1 | TRBV12-1*00 | TRBD1*00 | TRBJ2-3*00 | 0.28% |
| CD4+YFP- DKO | TGTGCCAGCTCTCTCGATAATTCGCCCCTCTACTTT | 1 | TRBV12-2*00 | | TRBJ1-6*00 | 0.28% |
| CD4+YFP- DKO | TGTGCCAGCTCTCTCGTTGAACAGTACTTC | 1 | TRBV12-1*00 | | TRBJ2-7*00 | 0.28% |
| CD4+YFP- DKO | TGTGCCTGGACAGTCCAAGACACCCAGTACTTT | 1 | TRBV31*00 | | TRBJ2-5*00 | 0.28% |
| CD4+YFP- DKO | TGTGCCTGGAGACCGGGACAGGGAAACTCCGACTACACCTTC | 1 | TRBV31*00 | TRBD1*00 | TRBJ1-2*00 | 0.28% |
| CD4+YFP- DKO | TGTGCCTGGAGCCCAGGGGGGGGCGACACAGAAGTCTTCTTT | 1 | TRBV31*00 | TRBD2*00 | TRBJ1-1*00 | 0.28% |
| CD4+YFP- DKO | TGTGCCTGGAGCCCCGGGACTGGGGTAGTGCAGAAACGCTGTATTTT | 1 | TRBV31*00 | TRBD2*00 | TRBJ2-3*00 | 0.28% |
| CD4+YFP- DKO | TGTGCCTGGAGCCCCTTGGGGGGGAGGCTTGAACAGTACTTC | 1 | TRBV31*00 | TRBD2*00 | TRBJ2-7*00 | 0.28% |
| CD4+YFP- DKO | TGTGCCTGGAGCCTGGACCATAACAACCAGGCTCCGCTTTTT | 1 | TRBV31*00 | | TRBJ1-5*00 | 0.28% |
| CD4+YFP- DKO | TGTGCCTGGAGTCAACAAAACACCTTGTACTTT | 1 | TRBV31*00 | | TRBJ2-4*00 | 0.28% |
| CD4+YFP- DKO | TGTGCCTGGAGTCAAGACACCCAGTACTTT | 1 | TRBV31*00 | | TRBJ2-5*00 | 0.28% |
| CD4+YFP- DKO | TGTGCCTGGAGTCAGCGACAGCCAAACTCCGACTACACCTTC | 1 | TRBV31*00 | TRBD1*00 | TRBJ1-2*00 | 0.28% |

| | | | | | | |
|---|---|---|---|---|---|---|
| CD4+YFP- DKO | TGTGCCTGGAGTCCGCAGGGGGGCTCCGACTACACCTTC | 1 | TRBV31*00 | TRBD1*00 | TRBJ1-2*00 | 0.28% |
| CD4+YFP- DKO | TGTGCCTGGAGTCGACTGGTAGTGCAGAAACGCTGTATTTT | 1 | TRBV31*00 | | TRBJ2-3*00 | 0.28% |
| CD4+YFP- DKO | TGTGCCTGGAGTCTAACGCAGTATGAACAGTACTTC | 1 | TRBV31*00 | | TRBJ2-7*00 | 0.28% |
| CD4+YFP- DKO | TGTGCCTGGAGTCTAAGGGGTAGTGCAGAAACGCTGTATTTT | 1 | TRBV31*00 | TRBD1*00 | TRBJ2-3*00 | 0.28% |
| CD4+YFP- DKO | TGTGCCTGGAGTCTACAGGGGGCGGACACAGAAGTCTTCTTT | 1 | TRBV31*00 | TRBD1*00 | TRBJ1-1*00 | 0.28% |
| CD4+YFP- DKO | TGTGCCTGGAGTCTAGAGGGGGGGGCTGAGCAGTTCTTC | 1 | TRBV31*00 | TRBD2*00 | TRBJ2-1*00 | 0.28% |
| CD4+YFP- DKO | TGTGCCTGGAGTCTAGCTGTTAACTATGCTGAGCAGTTCTTC | 1 | TRBV31*00 | | TRBJ2-1*00 | 0.28% |
| CD4+YFP- DKO | TGTGCCTGGAGTCTCCCCAATTCTGGAAATACGCTCTATTTT | 1 | TRBV31*00 | | TRBJ1-3*00 | 0.28% |
| CD4+YFP- DKO | TGTGCCTGGAGTCTGACTGGTAACTATGCTGAGCAGTTCTTC | 1 | TRBV31*00 | TRBD2*00 | TRBJ2-1*00 | 0.28% |
| CD4+YFP- DKO | TGTGCCTGGAGTCTGGGGGGCTCCTATGAACAGTACTTC | 1 | TRBV31*00 | TRBD2*00 | TRBJ2-7*00 | 0.28% |
| CD4+YFP- DKO | TGTGCCTGGGCCTCCGGGACAGCCTATGAACAGTACTTC | 1 | TRBV31*00 | TRBD1*00 | TRBJ2-7*00 | 0.28% |
| CD4+YFP- DKO | TGTGCTAGCAGTAGAGATGGGACTGGCAAAGACACCCAGTACTTT | 1 | TRBV17*00 | TRBD2*00 | TRBJ2-5*00 | 0.28% |
| CD4+YFP- DKO | TGTGCTAGCAGTAGAGTGGGACAGGGGTATGAACAGTACTTC | 1 | TRBV17*00 | TRBD1*00 | TRBJ2-7*00 | 0.28% |
| CD4+YFP- DKO | TGTGCTAGCAGTAGCTATTCTGGAAATACGCTCTATTTT | 1 | TRBV17*00 | | TRBJ1-3*00 | 0.28% |
| CD4+YFP- DKO | TGTGCTAGCAGTAGGGACAGGTGGAGTTCCGACTACACCTTC | 1 | TRBV17*00 | TRBD1*00 | TRBJ1-2*00 | 0.28% |
| CD4+YFP- DKO | TGTGCTAGCAGTAGGGGGGACACCTTGTACTTT | 1 | TRBV17*00 | TRBD1*00 | TRBJ2-4*00 | 0.28% |
| CD4+YFP- DKO | TGTGCTAGCAGTCTCTGGGGGGACTATGCTGAGCAGTTCTTC | 1 | TRBV17*00 | TRBD2*00 | TRBJ2-1*00 | 0.28% |
| CD4+YFP- DKO | TGTGCTAGCAGTTCTAGGGCTGAGCAGTTCTTC | 1 | TRBV29*00 | | TRBJ2-1*00 | 0.28% |
| CD4+YFP- DKO | TGTGCTAGCAGTTTAGGGGGGGGACGTGCAGAAACGCTGTATTTT | 1 | TRBV29*00 | TRBD2*00 | TRBJ2-3*00 | 0.28% |
| CD4+YFP- DKO | TGTGCTAGCAGTTTATCAGGGGGCCTTAACTATGCTGAGCAGTTCTTC | 1 | TRBV29*00 | TRBD1*00 | TRBJ2-1*00 | 0.28% |
| CD4+YFP- DKO | TGTGCTAGCAGTTTATTCCTATGAACAGTACTTC | 1 | TRBV29*00 | | TRBJ2-7*00 | 0.28% |
| CD4+YFP- DKO | TGTGCTAGCAGTTTCCGGGGCCCGTCCGACTACACCTTC | 1 | TRBV29*00 | TRBD1*00 | TRBJ1-2*00 | 0.28% |
| CD4+YFP- DKO | TGTGCTAGCAGTTTTAGTGCAGAAACGCTGTATTTT | 1 | TRBV29*00 | | TRBJ2-3*00 | 0.28% |
| CD4+YFP- DKO | TGTGCTAGCGGACAGAACGCAAACACCGGGCAGCTCTACTTT | 1 | TRBV29*00 | TRBD1*00 | TRBJ2-2*00 | 0.28% |
| CD4+YFP- DKO | TGTGGTGCTACGGGGGACTCCTATGAACAGTACTTC | 1 | TRBV20*00 | TRBD1*00 | TRBJ2-7*00 | 0.28% |
| CD4+YFP- DKO | TGTGGTGCTAGAAAGGGCTATGCTGAGCAGTTCTTC | 1 | TRBV20*00 | | TRBJ2-1*00 | 0.28% |
| CD4+YFP- DKO | TGTGGTGCTAGGACAACAAACTCCGACTACACCTTC | 1 | TRBV20*00 | | TRBJ1-2*00 | 0.28% |
| CD4+YFP- DKO | TGTGGTGCTAGGAGGGTTAGCTCCTATGAACAGTACTTC | 1 | TRBV20*00 | | TRBJ2-7*00 | 0.28% |
| CD4+YFP- DKO | TGTGGTGCTAGGCTGGGGGGTGGAGACACCCAGTACTTT | 1 | TRBV20*00 | TRBD2*00 | TRBJ2-5*00 | 0.28% |
| CD4+YFP- DKO | TGTGGTGCTAGGGACTGGGGAAACACCGGGCAGCTCTACTTT | 1 | TRBV20*00 | TRBD2*00 | TRBJ2-2*00 | 0.28% |
| CD4+YFP- DKO | TGTGGTGCTAGGGATAACCAAGACACCCAGTACTTT | 1 | TRBV20*00 | | TRBJ2-5*00 | 0.28% |
| CD4+YFP- DKO | TGTGGTGCTAGGGATCTGGGCGAAAGATTATTTTTC | 1 | TRBV20*00 | | TRBJ1-4*00 | 0.28% |
| CD4+YFP- DKO | TGTGGTGCTAGGGATCTTCTGGGGGCAGAAACGCTGTATTTT | 1 | TRBV20*00 | TRBD2*00 | TRBJ2-3*00 | 0.28% |
| CD4+YFP- DKO | TGTGGTGCTAGGGCCAGGGCCAACACAGAAGTCTTCTTT | 1 | TRBV20*00 | TRBD1*00 | TRBJ1-1*00 | 0.28% |
| CD4+YFP- DKO | TGTGGTGCTAGGGGGACAGTGGGCTCCGACTACACCTTC | 1 | TRBV20*00 | TRBD1*00 | TRBJ1-2*00 | 0.28% |
| CD4+YFP- DKO | TGTGGTGCTAGTCAGAATTCTGGAAATACGCTCTATTTT | 1 | TRBV20*00 | | TRBJ1-3*00 | 0.28% |
| CD4+YFP- DKO | TGTGCCAGCTCTCTCGACGGTGGGGGGAAAGACACCCAGTACTTT | 1 | TRBV12-1*00 | TRBD2*00 | TRBJ2-5*00 | 0.28% |
| CD4+YFP- DKO | TGTGCTAGCAGTCCCGCCCGTTCTGGAAATACGCTCTATTTT | 1 | TRBV29*00 | TRBD1*00 | TRBJ1-3*00 | 0.28% |
| CD4+YFP- DKO | TGTGCCAGCAGCCAAGACTGGGGGGGCTATGCTGAGCAGTTCTTC | 1 | TRBV2*00 | TRBD2*00 | TRBJ2-1*00 | 0.28% |
| CD4+YFP- DKO | TGTGCTAGCAGTAGTTACAACTATGCTGAGCAGTTCTTC | 1 | TRBV17*00 | | TRBJ2-1*00 | 0.28% |
| CD4+YFP- DKO | TGTGGTGCTAGGGACTGGGGCTCCTATGAACAGTACTTC | 1 | TRBV20*00 | TRBD2*00 | TRBJ2-7*00 | 0.28% |
| CD4+YFP- DKO | TGTGCCAGCAGTGATAACACAGTCTCCTATGAACAGTACTTC | 1 | TRBV13-3*00 | | TRBJ2-7*00 | 0.28% |
| CD4+YFP- DKO | TGTGCCAGCGGGACAACAAACTCCGACTACACCTTC | 1 | TRBV13-3*00 | TRBD1*00 | TRBJ1-2*00 | 0.28% |
| CD4+YFP- DKO | TGTGCCAGCGGTGACAGTTATGAACAGTACTTC | 1 | TRBV13-2*00 | | TRBJ2-7*00 | 0.28% |

**CD4+YFP- DKO***

| Sample Name | CDR3 Sequence | clone Count | best V Hit | best D Hit | best J Hit | Frequency |
|---|---|---|---|---|---|---|
| CD4+YFP- DKO* | TGTGCCAGCAGTCTAGAAAACACCGGGCAGCTCTACTTT | 629 | TRBV15*00 | | TRBJ2-2*00 | 83.53% |
| CD4+YFP- DKO* | TGTGCCAGCAGCTCTGACAATTATGAACAGTACTTC | 3 | TRBV25*00 | | TRBJ2-7*00 | 0.40% |
| CD4+YFP- DKO* | TGCACCTGCAGTGCAGACTGGAGCTCCTATGAACAGTACTTC | 2 | TRBV1*00 | | TRBJ2-7*00 | 0.27% |
| CD4+YFP- DKO* | TGTGCCAGCAGCCCCGGACAAATCCAAGACACCCAGTACTTT | 2 | TRBV4*00 | TRBD1*00 | TRBJ2-5*00 | 0.27% |
| CD4+YFP- DKO* | TGTGCCAGCAGTGATCAGGGGGGAGTCTTCTTT | 2 | TRBV13-3*00 | TRBD1*00 | TRBJ1-1*00 | 0.27% |
| CD4+YFP- DKO* | TGTGCCAGCAGTGATGTGGTTAACCAAGACACCCAGTACTTT | 2 | TRBV13-1*00 | | TRBJ2-5*00 | 0.27% |
| CD4+YFP- DKO* | TGTGCCAGCGGTGATCCAGACTGGGGGTCAACCCAGTACTTT | 2 | TRBV13-2*00 | TRBD2*00 | TRBJ2-5*00 | 0.27% |
| CD4+YFP- DKO* | TGTGCTAGCAGTTTAGGGGGACAGGGGAAAGACACCCAGTACTTT | 2 | TRBV29*00 | TRBD1*00 | TRBJ2-5*00 | 0.27% |
| CD4+YFP- DKO* | TGTGCCAGCAGCCTCCAAGACACCCAGTACTTT | 2 | TRBV3*00 | | TRBJ2-5*00 | 0.27% |
| CD4+YFP- DKO* | TGCACCTGCAGTGCAGACACCGGGCAGCTCTACTTT | 1 | TRBV1*00 | | TRBJ2-2*00 | 0.13% |
| CD4+YFP- DKO* | TGCACCTGCAGTGCAGAGAACTGGGGGGATGAACAGTACTTC | 1 | TRBV1*00 | TRBD2*00 | TRBJ2-7*00 | 0.13% |
| CD4+YFP- DKO* | TGCACCTGCAGTGCAGAGGGACTCCAAGACACCCAGTACTTT | 1 | TRBV1*00 | TRBD2*00 | TRBJ2-5*00 | 0.13% |
| CD4+YFP- DKO* | TGCACCTGCAGTGCAGGGGCTGCTGAGCAGTTCTTC | 1 | TRBV1*00 | | TRBJ2-1*00 | 0.13% |
| CD4+YFP- DKO* | TGCTCCAGCAGTCAATCGAACTATGCTGAGCAGTTCTTC | 1 | TRBV23*00 | | TRBJ2-1*00 | 0.13% |
| CD4+YFP- DKO* | TGTGCAAGCAGCATCTATGCTGAGCAGTTCTTC | 1 | TRBV16*00 | | TRBJ2-1*00 | 0.13% |
| CD4+YFP- DKO* | TGTGCAAGCAGCCCACAGGGGGCTGGGTCCGACTACACCTTC | 1 | TRBV16*00 | TRBD1*00 | TRBJ1-2*00 | 0.13% |
| CD4+YFP- DKO* | TGTGCAAGCAGCCCCCTTGCTGAGCAGTTCTTC | 1 | TRBV16*00 | TRBD1*00 | TRBJ2-1*00 | 0.13% |
| CD4+YFP- DKO* | TGTGCAAGCAGCCCGCCCCGGTGGTATGAACAGTACTTC | 1 | TRBV16*00 | TRBD1*00 | TRBJ2-7*00 | 0.13% |
| CD4+YFP- DKO* | TGTGCAAGCAGCCCGGACAGTGCAGAAACGCTGTATTTT | 1 | TRBV16*00 | | TRBJ2-3*00 | 0.13% |
| CD4+YFP- DKO* | TGTGCAAGCAGCTCCGACCAAGACACCCAGTACTTT | 1 | TRBV16*00 | | TRBJ2-5*00 | 0.13% |
| CD4+YFP- DKO* | TGTGCAAGCAGCTCGACGGTCACAGAAGTCTTCTTT | 1 | TRBV16*00 | | TRBJ1-1*00 | 0.13% |
| CD4+YFP- DKO* | TGTGCAAGCAGCTTAGAAGACACAGAAGTCTTCTTT | 1 | TRBV16*00 | | TRBJ1-1*00 | 0.13% |
| CD4+YFP- DKO* | TGTGCCAGAAGGGACATCCAAGACACCCAGTACTTT | 1 | TRBV13-3*00 | TRBD1*00 | TRBJ2-5*00 | 0.13% |
| CD4+YFP- DKO* | TGTGCCAGCAGAACCACAATTAACCAAGACACCCAGTACTTT | 1 | TRBV19*00 | | TRBJ2-5*00 | 0.13% |
| CD4+YFP- DKO* | TGTGCCAGCAGAGCGGGACTGGGGGCTAGTGCAGAAACGCTGTATTTT | 1 | TRBV15*00 | TRBD2*00 | TRBJ2-3*00 | 0.13% |
| CD4+YFP- DKO* | TGTGCCAGCAGCAGGGGTAGTCAAAACACCTTGTACTTT | 1 | TRBV15*00 | TRBD1*00 | TRBJ2-4*00 | 0.13% |
| CD4+YFP- DKO* | TGTGCCAGCAGCCAACCTGGGAGGGCTCCGCTTTTT | 1 | TRBV5*00 | TRBD2*00 | TRBJ1-5*00 | 0.13% |
| CD4+YFP- DKO* | TGTGCCAGCAGCCAAGAAGACAATAACTATGCTGAGCAGTTCTTC | 1 | TRBV5*00 | | TRBJ2-1*00 | 0.13% |

| | | | | | | |
|---|---|---|---|---|---|---|
| CD4+YFP- DKO* | TGTGCCAGCAGCCAAGACCTGGGGGGAATTTATGAACAGTACTTC | 1 | TRBV5*00 | TRBD2*00 | TRBJ2-7*00 | 0.13% |
| CD4+YFP- DKO* | TGTGCCAGCAGCCAAGAGCAGGGGAGGGGAGACACCCAGTACTTT | 1 | TRBV5*00 | TRBD1*00 | TRBJ2-5*00 | 0.13% |
| CD4+YFP- DKO* | TGTGCCAGCAGCCAAGAGGCTGGGGGGCGAGAACAGTACTTC | 1 | TRBV2*00 | TRBD2*00 | TRBJ2-7*00 | 0.13% |
| CD4+YFP- DKO* | TGTGCCAGCAGCCAAGAGGGGGGGGCGCACCGGGCAGCTCTACTTT | 1 | TRBV2*00 | TRBD2*00 | TRBJ2-2*00 | 0.13% |
| CD4+YFP- DKO* | TGTGCCAGCAGCCAAGGATCTGGGCGTGCAGAAACGCTGTATTTT | 1 | TRBV5*00 | | TRBJ2-3*00 | 0.13% |
| CD4+YFP- DKO* | TGTGCCAGCAGCCAAGGCTATGAACAGTACTTC | 1 | TRBV5*00 | | TRBJ2-7*00 | 0.13% |
| CD4+YFP- DKO* | TGTGCCAGCAGCCAAGTTGTCGTCGACACAGAAGTCTTCTTT | 1 | TRBV2*00 | | TRBJ1-1*00 | 0.13% |
| CD4+YFP- DKO* | TGTGCCAGCAGCCCAGGGGGGTATGAACAGTACTTC | 1 | TRBV14*00 | TRBD1*00 | TRBJ2-7*00 | 0.13% |
| CD4+YFP- DKO* | TGTGCCAGCAGCCCGACAGGGAGTTCTGGAAATACGCTCTATTTT | 1 | TRBV3*00 | TRBD1*00 | TRBJ1-3*00 | 0.13% |
| CD4+YFP- DKO* | TGTGCCAGCAGCCCGGGACAGTACTATGCTGAGCAGTTCTTC | 1 | TRBV15*00 | TRBD1*00 | TRBJ2-1*00 | 0.13% |
| CD4+YFP- DKO* | TGTGCCAGCAGCCCTGGGGTAGTGCAGAAACGCTGTATTTT | 1 | TRBV5*00 | TRBD2*00 | TRBJ2-3*00 | 0.13% |
| CD4+YFP- DKO* | TGTGCCAGCAGCCGGGACAGGGGGAGTGCAGAAACGCTGTATTTT | 1 | TRBV5*00 | TRBD1*00 | TRBJ2-3*00 | 0.13% |
| CD4+YFP- DKO* | TGTGCCAGCAGCCGTAACCAAGACACCCAGTACTTT | 1 | TRBV19*00 | | TRBJ2-5*00 | 0.13% |
| CD4+YFP- DKO* | TGTGCCAGCAGCCTACAGAACCAGGCTCCGCTTTTT | 1 | TRBV4*00 | | TRBJ1-5*00 | 0.13% |
| CD4+YFP- DKO* | TGTGCCAGCAGCTACAACAACGAAAGATTATTTTTC | 1 | TRBV4*00 | | TRBJ1-4*00 | 0.13% |
| CD4+YFP- DKO* | TGTGCCAGCAGCTACCAAGACACCCAGTACTTT | 1 | TRBV5*00 | | TRBJ2-5*00 | 0.13% |
| CD4+YFP- DKO* | TGTGCCAGCAGCTCCACAGGGGACACCCAGTACTTT | 1 | TRBV14*00 | TRBD1*00 | TRBJ2-5*00 | 0.13% |
| CD4+YFP- DKO* | TGTGCCAGCAGCTGGGGGGGCGCAGAAACGCTGTATTTT | 1 | TRBV4*00 | TRBD2*00 | TRBJ2-3*00 | 0.13% |
| CD4+YFP- DKO* | TGTGCCAGCAGGGACACCAACCAGGCTCCGCTTTTT | 1 | TRBV13-1*00 | TRBD1*00 | TRBJ1-5*00 | 0.13% |
| CD4+YFP- DKO* | TGTGCCAGCAGTATAACCGTTAACCAAGACACCCAGTACTTT | 1 | TRBV19*00 | | TRBJ2-5*00 | 0.13% |
| CD4+YFP- DKO* | TGTGCCAGCAGTATACCGGCAGAAGTCTTCTTT | 1 | TRBV19*00 | | TRBJ1-1*00 | 0.13% |
| CD4+YFP- DKO* | TGTGCCAGCAGTATAGAGGGGGGGCGAACAGTACTTC | 1 | TRBV19*00 | TRBD2*00 | TRBJ2-7*00 | 0.13% |
| CD4+YFP- DKO* | TGTGCCAGCAGTATAGATGGGGAGGGCACCCAGTACTTT | 1 | TRBV19*00 | TRBD2*00 | TRBJ2-5*00 | 0.13% |
| CD4+YFP- DKO* | TGTGCCAGCAGTATCCCACCATCAAACACCGGGCAGCTCTACTTT | 1 | TRBV19*00 | | TRBJ2-2*00 | 0.13% |
| CD4+YFP- DKO* | TGTGCCAGCAGTCCAGAAACACCGGGCAGCTCTACTTT | 1 | TRBV15*00 | | TRBJ2-2*00 | 0.13% |
| CD4+YFP- DKO* | TGTGCCAGCAGTCCCGGACACCTTATGAACAGTACTTC | 1 | TRBV22*00 | TRBD1*00 | TRBJ2-7*00 | 0.13% |
| CD4+YFP- DKO* | TGTGCCAGCAGTCCGACAACTAGTGCAGAAACGCTGTATTTT | 1 | TRBV26*00 | | TRBJ2-3*00 | 0.13% |
| CD4+YFP- DKO* | TGTGCCAGCAGTCTCTTGGGGGACTACACCTTC | 1 | TRBV24*00 | TRBD2*00 | TRBJ1-2*00 | 0.13% |
| CD4+YFP- DKO* | TGTGCCAGCAGTCTGACAGGGGATAATTCGCCCCTCTACTTT | 1 | TRBV26*00 | TRBD1*00 | TRBJ1-6*00 | 0.13% |
| CD4+YFP- DKO* | TGTGCCAGCAGTCTGGACTCCGACTACACCTTC | 1 | TRBV26*00 | | TRBJ1-2*00 | 0.13% |
| CD4+YFP- DKO* | TGTGCCAGCAGTCTGGGGGACAGAGAAGTCTTCTTT | 1 | TRBV26*00 | | TRBJ1-1*00 | 0.13% |
| CD4+YFP- DKO* | TGTGCCAGCAGTCTGTTGGGGGACACCGGGCAGCTCTACTTT | 1 | TRBV26*00 | TRBD2*00 | TRBJ2-2*00 | 0.13% |
| CD4+YFP- DKO* | TGTGCCAGCAGTGAACAGGGGGCCAACGAAAGATTATTTTTC | 1 | TRBV13-3*00 | TRBD1*00 | TRBJ1-4*00 | 0.13% |
| CD4+YFP- DKO* | TGTGCCAGCAGTGACACGACAGGGGGCAACTCCGACTACACCTTC | 1 | TRBV13-3*00 | TRBD1*00 | TRBJ1-2*00 | 0.13% |
| CD4+YFP- DKO* | TGTGCCAGCAGTGACGGGTACTCCGACTACACCTTC | 1 | TRBV13-1*00 | | TRBJ1-2*00 | 0.13% |
| CD4+YFP- DKO* | TGTGCCAGCAGTGAGATTGGGGACACCGGGCAGCTCTACTTT | 1 | TRBV13-3*00 | TRBD2*00 | TRBJ2-2*00 | 0.13% |
| CD4+YFP- DKO* | TGTGCCAGCAGTGAGGGGGGCTGGGCTGAGCAGTTCTTC | 1 | TRBV13-1*00 | TRBD2*00 | TRBJ2-1*00 | 0.13% |
| CD4+YFP- DKO* | TGTGCCAGCAGTGATAACTCCGACTACACCTTC | 1 | TRBV13-3*00 | | TRBJ1-2*00 | 0.13% |
| CD4+YFP- DKO* | TGTGCCAGCAGTGATAGGTATTCTGGAAATACGCTCTATTTT | 1 | TRBV13-1*00 | | TRBJ1-3*00 | 0.13% |
| CD4+YFP- DKO* | TGTGCCAGCAGTGATCCGGGGGCTGGCTCCGACTACACCTTC | 1 | TRBV13-1*00 | TRBD1*00 | TRBJ1-2*00 | 0.13% |
| CD4+YFP- DKO* | TGTGCCAGCAGTGATGCCAATTCTGGAAATACGCTCTATTTT | 1 | TRBV13-1*00 | | TRBJ1-3*00 | 0.13% |
| CD4+YFP- DKO* | TGTGCCAGCAGTGATGTGGACAGGGAGGGCACCCAGTACTTT | 1 | TRBV13-3*00 | TRBD1*00 | TRBJ2-5*00 | 0.13% |
| CD4+YFP- DKO* | TGTGCCAGCAGTGATTACGGAACTAACGAAAGATTATTTTTC | 1 | TRBV13-1*00 | | TRBJ1-4*00 | 0.13% |
| CD4+YFP- DKO* | TGTGCCAGCAGTGATTCCTGGCAGGGCACCGGGCAGCTCTACTTT | 1 | TRBV13-3*00 | TRBD1*00 | TRBJ2-2*00 | 0.13% |
| CD4+YFP- DKO* | TGTGCCAGCAGTGGAGACAGGAGCTCCTATGAACAGTACTTC | 1 | TRBV13-3*00 | | TRBJ2-7*00 | 0.13% |
| CD4+YFP- DKO* | TGTGCCAGCAGTGGAGGACAGAACACAGAAGTCTTCTTT | 1 | TRBV13-1*00 | TRBD1*00 | TRBJ1-1*00 | 0.13% |
| CD4+YFP- DKO* | TGTGCCAGCAGTGGCAACGAAAGATTATTTTTC | 1 | TRBV13-1*00 | | TRBJ1-4*00 | 0.13% |
| CD4+YFP- DKO* | TGTGCCAGCAGTTCAGCTAGTCAAAACACCTTGTACTTT | 1 | TRBV22*00 | | TRBJ2-4*00 | 0.13% |
| CD4+YFP- DKO* | TGTGCCAGCAGTTCGGACTGGGGGGATGCTGAGCAGTTCTTC | 1 | TRBV26*00 | TRBD2*00 | TRBJ2-1*00 | 0.13% |
| CD4+YFP- DKO* | TGTGCCAGCAGTTCTGGGGGGGCTGAAGACACCCAGTACTTT | 1 | TRBV15*00 | TRBD2*00 | TRBJ2-5*00 | 0.13% |
| CD4+YFP- DKO* | TGTGCCAGCAGTTGACTAGTGCAGAAACGCTGTATTTT | 1 | TRBV19*00 | | TRBJ2-3*00 | 0.13% |
| CD4+YFP- DKO* | TGTGCCAGCAGTTTACAGTACACCGGGCAGCTCTACTTT | 1 | TRBV14*00 | | TRBJ2-2*00 | 0.13% |
| CD4+YFP- DKO* | TGTGCCAGCAGTTTAGACTCCGACTACACCTTC | 1 | TRBV15*00 | | TRBJ1-2*00 | 0.13% |
| CD4+YFP- DKO* | TGTGCCAGCAGTTTCGGCAGGTATGAACAGTACTTC | 1 | TRBV14*00 | | TRBJ2-7*00 | 0.13% |
| CD4+YFP- DKO* | TGTGCCAGCAGTTTCTGGAGTCAAAACACCTTGTACTTT | 1 | TRBV14*00 | | TRBJ2-4*00 | 0.13% |
| CD4+YFP- DKO* | TGTGCCAGCAGTTTGGCCAGGGGTAACCAAGACACCCAGTACTTT | 1 | TRBV14*00 | TRBD1*00 | TRBJ2-5*00 | 0.13% |
| CD4+YFP- DKO* | TGTGCCAGCAGTTTGTATGCTGAGCAGTTCTTC | 1 | TRBV14*00 | | TRBJ2-1*00 | 0.13% |
| CD4+YFP- DKO* | TGTGCCAGCCGAGGGGGGATTTCCAACGAAAGATTATTTTTC | 1 | TRBV12-1*00 | TRBD1*00 | TRBJ1-4*00 | 0.13% |
| CD4+YFP- DKO* | TGTGCCAGCGGCACTTCCTATAATTCGCCCCTCTACTTT | 1 | TRBV4*00 | | TRBJ1-6*00 | 0.13% |
| CD4+YFP- DKO* | TGTGCCAGCGGGGACTGGGGGTCTAGTCAAAACACCTTGTACTTT | 1 | TRBV13-2*00 | TRBD2*00 | TRBJ2-4*00 | 0.13% |
| CD4+YFP- DKO* | TGTGCCAGCGGTCCAACAAACTCCGACTACACCTTC | 1 | TRBV13-2*00 | | TRBJ1-2*00 | 0.13% |
| CD4+YFP- DKO* | TGTGCCAGCGGTGATGCAGGGTATAACAACCAGGCTCCGCTTTTT | 1 | TRBV13-2*00 | | TRBJ1-5*00 | 0.13% |
| CD4+YFP- DKO* | TGTGCCAGCGGTGATGCCGGACAGAACACAGAAGTCTTCTTT | 1 | TRBV13-2*00 | TRBD1*00 | TRBJ1-1*00 | 0.13% |
| CD4+YFP- DKO* | TGTGCCAGCGGTGATGCCGGGGGAAGGCAAAACACCTTGTACTTT | 1 | TRBV13-2*00 | TRBD1*00 | TRBJ2-4*00 | 0.13% |
| CD4+YFP- DKO* | TGTGCCAGCGGTGATGCGAGCAGAAACTCCGACTACACCTTC | 1 | TRBV13-2*00 | | TRBJ1-2*00 | 0.13% |
| CD4+YFP- DKO* | TGTGCCAGCGGTGATGTGGGGGCTCCGCTTTTT | 1 | TRBV13-2*00 | | TRBJ1-5*00 | 0.13% |
| CD4+YFP- DKO* | TGTGCCAGCGGTGATTACAATAGTGCAGAAACGCTGTATTTT | 1 | TRBV13-2*00 | | TRBJ2-3*00 | 0.13% |
| CD4+YFP- DKO* | TGTGCCAGCGGTGATTGGGTTTACCAAGACACCCAGTACTTT | 1 | TRBV13-2*00 | | TRBJ2-5*00 | 0.13% |
| CD4+YFP- DKO* | TGTGCCAGCGGTGGGGGTAACCAAGACACCCAGTACTTT | 1 | TRBV13-2*00 | | TRBJ2-5*00 | 0.13% |
| CD4+YFP- DKO* | TGTGCCAGCGGTGTCAACACCCTTGAACAGTACTTC | 1 | TRBV13-2*00 | | TRBJ2-7*00 | 0.13% |
| CD4+YFP- DKO* | TGTGCCAGCTCTCTCGAGCCCAGGAGCTCCTATGAACAGTACTTC | 1 | TRBV12-1*00 | | TRBJ2-7*00 | 0.13% |
| CD4+YFP- DKO* | TGTGCCTGGACCCCCACCAATGCTGAGCAGTTCTTC | 1 | TRBV31*00 | TRBD2*00 | TRBJ2-1*00 | 0.13% |
| CD4+YFP- DKO* | TGTGCCTGGAGCCAGGACAGGGGGCCCGACTACACCTTC | 1 | TRBV31*00 | TRBD1*00 | TRBJ1-2*00 | 0.13% |
| CD4+YFP- DKO* | TGTGCCTGGAGTCTAAACAAAAACACAGAAGTCTTCTTT | 1 | TRBV31*00 | | TRBJ1-1*00 | 0.13% |
| CD4+YFP- DKO* | TGTGCCTGGAGTCTAGGCAGGGCCTATAATTCGCCCCTCTACTTT | 1 | TRBV31*00 | TRBD1*00 | TRBJ1-6*00 | 0.13% |

| | | | | | | |
|---|---|---|---|---|---|---|
| CD4+YFP- DKO* | TGTGCCTGGTCGGACAATTCTGGAAATACGCTCTATTTT | 1 | TRBV31*00 | | TRBJ1-3*00 | 0.13% |
| CD4+YFP- DKO* | TGTGCTAGCAGTCCCGACAGGTATGAACAGTACTTC | 1 | TRBV17*00 | TRBD1*00 | TRBJ2-7*00 | 0.13% |
| CD4+YFP- DKO* | TGTGCTAGCAGTTACGGGGACAGGCAAGACACCAAGTACTTT | 1 | TRBV29*00 | TRBD1*00 | TRBJ2-5*00 | 0.13% |
| CD4+YFP- DKO* | TGTGCTAGCAGTTTGGCCAGGGCAAACTCCGACTACACCTTC | 1 | TRBV29*00 | | TRBJ1-2*00 | 0.13% |
| CD4+YFP- DKO* | TGTGCTAGCAGTTTGGGGCCAGGGGGGAACTCCGACTACACCTTC | 1 | TRBV29*00 | TRBD1*00 | TRBJ1-2*00 | 0.13% |
| CD4+YFP- DKO* | TGTGCTAGCGCCTATGAACAGTACTTC | 1 | TRBV21*00 | | TRBJ2-7*00 | 0.13% |
| CD4+YFP- DKO* | TGTGGTGCTAGAACAACTTATAACAACCAGGCTCCGCTTTTT | 1 | TRBV20*00 | | TRBJ1-5*00 | 0.13% |
| CD4+YFP- DKO* | TGTGGTGCTAGCAGGGACTATGCTGAGCAGTTCTTC | 1 | TRBV20*00 | TRBD1*00 | TRBJ2-1*00 | 0.13% |
| CD4+YFP- DKO* | TGTGGTGCTAGGAAGTCTAGTGCAGAAACGCTGTATTTT | 1 | TRBV20*00 | | TRBJ2-3*00 | 0.13% |
| CD4+YFP- DKO* | TGTGGTGCTAGGACAGGGGGCTATGCTGAGCAGTTCTTC | 1 | TRBV20*00 | TRBD1*00 | TRBJ2-1*00 | 0.13% |
| CD4+YFP- DKO* | TGTGGTGCTAGGCCTAGTGCAGAAACGCTGTATTTT | 1 | TRBV20*00 | | TRBJ2-3*00 | 0.13% |
| CD4+YFP- DKO* | TGTGGTGCTAGGGAGGCCCGGACGGACACCCAGTACTTT | 1 | TRBV20*00 | TRBD1*00 | TRBJ2-5*00 | 0.13% |
| CD4+YFP- DKO* | TGTGGTGCTAGTGGGCGGTATGCTGAGCAGTTCTTC | 1 | TRBV20*00 | TRBD1*00 | TRBJ2-1*00 | 0.13% |

#### Supplementary Table 7. CDR3 sequences for Treg cells and $CD4^+$ T cells from mLNs

##### Treg WT1

| Sample name | CDR3 Sequence | clone Count | best V Hit | best D Hit | best J Hit | Frequency |
|---|---|---|---|---|---|---|
| Treg WT1 | TGTGCCAGCAGTAACGGGGGCCAATATGAACAGTACTTC | 3 | TRBV19*00 | TRBD1*00 | TRBJ2-7*00 | 1.18% |
| Treg WT1 | TGTGCTAGCAGTTTAGACTGGGGGTCCACCGGGCAGCTCTACTTT | 2 | TRBV29*00 | TRBD2*00 | TRBJ2-2*00 | 0.78% |
| Treg WT1 | TGTGCCAGCAGTATGCCGGGGGGAGGGAATGCTGAGCAGTTCTTC | 2 | TRBV19*00 | TRBD2*00 | TRBJ2-1*00 | 0.78% |
| Treg WT1 | TGTGCCAGCAGCTTGAGGACATCTAGTGCAGAAACGCTGTATTTT | 2 | TRBV3*00 | TRBD1*00 | TRBJ2-3*00 | 0.78% |
| Treg WT1 | TGTGCCAGCAGCCGGGGGCACCCTAGTGCAGAAACGCTGTATTTT | 2 | TRBV5*00 | TRBD1*00 | TRBJ2-3*00 | 0.78% |
| Treg WT1 | TGTGCCAGCCAGGGACTGGGGAGTGCAGAAACGCTGTATTTT | 2 | TRBV3*00 | TRBD2*00 | TRBJ2-3*00 | 0.78% |
| Treg WT1 | TGTGCCAGCAGTTTAGGGGGTTCTGGAAATACGCTCTATTTT | 2 | TRBV15*00 | | TRBJ1-3*00 | 0.78% |
| Treg WT1 | TGTGCCAGCAGTGATGGGGGTAACCAAGACACCCAGTACTTT | 2 | TRBV13-1*00 | | TRBJ2-5*00 | 0.78% |
| Treg WT1 | TGTGCCAGCAGCTTACTGGCGAGCTCCTATGAACAGTACTTC | 2 | TRBV3*00 | | TRBJ2-7*00 | 0.78% |
| Treg WT1 | TGTGCCAGCAGCCGCGCATCAAACACCGGGCAGCTCTACTTT | 2 | TRBV4*00 | | TRBJ2-2*00 | 0.78% |
| Treg WT1 | TGTGCCAGCAGCCGGGACAATAACTATGCTGAGCAGTTCTTC | 2 | TRBV19*00 | TRBD1*00 | TRBJ2-1*00 | 0.78% |
| Treg WT1 | TGTGCTAGCAGTTTATCGGTGAAGGACACCCAGTACTTT | 2 | TRBV29*00 | | TRBJ2-5*00 | 0.78% |
| Treg WT1 | TGTGCTAGCAGTTTAGGGGCTGGAAATACGCTCTATTTT | 2 | TRBV29*00 | | TRBJ1-3*00 | 0.78% |
| Treg WT1 | TGTGCCAGCTCTCAACAGGGAGGCGAAAGATTATTTTTC | 2 | TRBV12-1*00 | TRBD1*00 | TRBJ1-4*00 | 0.78% |
| Treg WT1 | TGTGCCAGCGGTGATTGGAACACCGGGCAGCTCTACTTT | 2 | TRBV13-2*00 | | TRBJ2-2*00 | 0.78% |
| Treg WT1 | TGTGCCAGCAGTCCCCGGAACAGCTATGAACAGTACTTC | 2 | TRBV26*00 | TRBD1*00 | TRBJ2-7*00 | 0.78% |
| Treg WT1 | TGTGCCAGCAGTATAGCAGGTGCAGAAACGCTGTATTTT | 2 | TRBV19*00 | | TRBJ2-3*00 | 0.78% |
| Treg WT1 | TGTGCCAGCAGCTTAACGGCAAACTCCGACTACACCTTC | 2 | TRBV3*00 | | TRBJ1-2*00 | 0.78% |
| Treg WT1 | TGTGCCAGCGCCGTTTAACTATGCTGAGCAGTTCTTC | 2 | TRBV19*00 | | TRBJ2-1*00 | 0.78% |
| Treg WT1 | TGTGCCAGCTCTCTCGATGCTGAGGAACAGTACTTC | 2 | TRBV12-2*00 | | TRBJ2-7*00 | 0.78% |
| Treg WT1 | TGTGCCAGCGGTGATCACAACTCCGACTACACCTTC | 2 | TRBV13-2*00 | | TRBJ1-2*00 | 0.78% |
| Treg WT1 | TGTGCCAGCAGTGATCACAGTTATGAACAGTACTTC | 2 | TRBV13-3*00 | | TRBJ2-7*00 | 0.78% |
| Treg WT1 | TGTGCCAGCAGCTTAGTAGGGGTTGAACAGTACTTC | 2 | TRBV3*00 | TRBD1*00 | TRBJ2-7*00 | 0.78% |
| Treg WT1 | TGTGCCAGCAGCCGGACAGGAAATACGCTCTATTTT | 2 | TRBV2*00 | TRBD1*00 | TRBJ1-3*00 | 0.78% |
| Treg WT1 | TGTGCCAGCAGCAGACAAACCGGGCAGCTCTACTTT | 2 | TRBV2*00 | | TRBJ2-2*00 | 0.78% |
| Treg WT1 | TGTGGTGCTAGCCGTAACCAAGACACCCAGTACTTT | 2 | TRBV20*00 | | TRBJ2-5*00 | 0.78% |
| Treg WT1 | TGTGCCAGCAGCCCACAGGAAAGATTATTTTTC | 2 | TRBV3*00 | | TRBJ1-4*00 | 0.78% |
| Treg WT1 | TGTGCCAGCAGTCGGGGACAGGGGAGGGGTGCAGAAACGCTGTATTTT | 1 | TRBV26*00 | TRBD1*00 | TRBJ2-3*00 | 0.39% |
| Treg WT1 | TGTGCAAGCAGCTTGGACTCTAATTCCTATAATTCGCCCCTCTACTTT | 1 | TRBV16*00 | | TRBJ1-6*00 | 0.39% |
| Treg WT1 | TGTGCCAGCTCTCTAGGGACAGCCGAGCTCCTATGAACAGTACTTC | 1 | TRBV12-2*00 | TRBD1*00 | TRBJ2-7*00 | 0.39% |
| Treg WT1 | TGTGCTAGCAGTAGACGGACAGGGAGCTCCTATGAACAGTACTTC | 1 | TRBV17*00 | TRBD1*00 | TRBJ2-7*00 | 0.39% |
| Treg WT1 | TGTGCTAGCAGTAGAACCCACAGATCCAACGAAAGATTATTTTTC | 1 | TRBV17*00 | | TRBJ1-4*00 | 0.39% |
| Treg WT1 | TGTGCCAGCTCTCTCGGTTGGGATTCTGGAAATACGCTCTATTTT | 1 | TRBV12-1*00 | | TRBJ1-3*00 | 0.39% |
| Treg WT1 | TGTGCCAGCCGCCCTCGGGACAGTTCTGGAAATACGCTCTATTTT | 1 | TRBV19*00 | TRBD1*00 | TRBJ1-3*00 | 0.39% |
| Treg WT1 | TGTGCCAGCGGTGGCGGACTGGGGCTGGGAGAAACGCTGTATTTT | 1 | TRBV13-2*00 | TRBD2*00 | TRBJ2-3*00 | 0.39% |
| Treg WT1 | TGTGCCAGCGGTGATGCACAGGGCAACTATGCTGAGCAGTTCTTC | 1 | TRBV13-2*00 | TRBD1*00 | TRBJ2-1*00 | 0.39% |
| Treg WT1 | TGTGCCAGCAGTTTCGTCACAGCTAGTGCAGAAACGCTGTATTTT | 1 | TRBV14*00 | | TRBJ2-3*00 | 0.39% |
| Treg WT1 | TGTGCCAGCAGTCTGAGTGGGGCTAGTGCAGAAACGCTGTATTTT | 1 | TRBV26*00 | TRBD2*00 | TRBJ2-3*00 | 0.39% |
| Treg WT1 | TGTGCCAGCAGTGATCTGGGGGCTAGTGCAGAAACGCTGTATTTT | 1 | TRBV13-3*00 | TRBD2*00 | TRBJ2-3*00 | 0.39% |
| Treg WT1 | TGTGCCAGCAGTGATCCTGGGGGGAACTATGCTGAGCAGTTCTTC | 1 | TRBV13-3*00 | TRBD2*00 | TRBJ2-1*00 | 0.39% |
| Treg WT1 | TGTGCCAGCAGTGACCGGGACAGGGGGCATGCTGAGCAGTTCTTC | 1 | TRBV13-3*00 | TRBD1*00 | TRBJ2-1*00 | 0.39% |
| Treg WT1 | TGTGCCAGCAGTATAGACAGATCAAACACCGGGCAGCTCTACTTT | 1 | TRBV19*00 | | TRBJ2-2*00 | 0.39% |
| Treg WT1 | TGTGCCAGCAGCTATCGGGGGTCTAGTGCAGAAACGCTGTATTTT | 1 | TRBV4*00 | TRBD1*00 | TRBJ2-3*00 | 0.39% |
| Treg WT1 | TGTGCCAGCAGCCTCGACAGGGCCACAAACTCCGACTACACCTTC | 1 | TRBV26*00 | TRBD1*00 | TRBJ1-2*00 | 0.39% |
| Treg WT1 | TGTGCCAGCAGCCAGGACTGGGGGGGAGCAGAAACGCTGTATTTT | 1 | TRBV5*00 | TRBD2*00 | TRBJ2-3*00 | 0.39% |
| Treg WT1 | TGTGCCAGCAGCCAAGATCCGGGGGGACAAGACACCCAGTACTTT | 1 | TRBV2*00 | TRBD2*00 | TRBJ2-5*00 | 0.39% |
| Treg WT1 | TGTGCCAGCAGCCAAGATCCGGGGAACCAAGACACCCAGTACTTT | 1 | TRBV5*00 | | TRBJ2-5*00 | 0.39% |
| Treg WT1 | TGTGCCAGCAGCCAAGAAGGGGCAAACACCGGGCAGCTCTACTTT | 1 | TRBV5*00 | | TRBJ2-2*00 | 0.39% |
| Treg WT1 | TGTGCCAGCAGCCAAGAAAGTAGTAACCAAGACACCCAGTACTTT | 1 | TRBV5*00 | | TRBJ2-5*00 | 0.39% |
| Treg WT1 | TGTGCCAGCAGGCCTCACTGGGGGAGCTCCTATGAACAGTACTTC | 1 | TRBV14*00 | TRBD2*00 | TRBJ2-7*00 | 0.39% |
| Treg WT1 | TGTGCAAGCAGCTTATTACAGGGGGCCAACGAAAGATTATTTTTC | 1 | TRBV16*00 | TRBD1*00 | TRBJ1-4*00 | 0.39% |
| Treg WT1 | TGTGCAAGCAGCTTAGACAGGGGGAGTGCAGAAACGCTGTATTTT | 1 | TRBV16*00 | TRBD1*00 | TRBJ2-3*00 | 0.39% |
| Treg WT1 | TGTGCAAGCAGCTTAACGGGGGGCCGTGCAGAAACGCTGTATTTT | 1 | TRBV16*00 | TRBD2*00 | TRBJ2-3*00 | 0.39% |
| Treg WT1 | TGCTCCAGCAGTCAACCGGGACTGGGGTGGGCTGAGCAGTTCTTC | 1 | TRBV23*00 | TRBD2*00 | TRBJ2-1*00 | 0.39% |
| Treg WT1 | TGCACCTGCAGTGCAGATTGGGGGGCTGGAAATACGCTCTATTTT | 1 | TRBV1*00 | TRBD2*00 | TRBJ1-3*00 | 0.39% |
| Treg WT1 | TGTGCCTGGAGTCTGGAGGGGGCCTCAAGACACCCAGTACTTT | 1 | TRBV31*00 | TRBD1*00 | TRBJ2-5*00 | 0.39% |
| Treg WT1 | TGTGCTAGCAGTCTCAGGGGGCGCAGAGACACCCAGTACTTT | 1 | TRBV21*00 | TRBD1*00 | TRBJ2-5*00 | 0.39% |
| Treg WT1 | TGTGCTAGCAGCCCGACTTTTAACCAAGACACCCAGTACTTT | 1 | TRBV17*00 | | TRBJ2-5*00 | 0.39% |
| Treg WT1 | TGTGCTAGCAAGAAGAATTCCTATAATTCGCCCCTCTACTTT | 1 | TRBV29*00 | | TRBJ1-6*00 | 0.39% |
| Treg WT1 | TGTGCCAGCGGGAGGGGGAATTCCAACGAAAGATTATTTTTC | 1 | TRBV13-2*00 | TRBD1*00 | TRBJ1-4*00 | 0.39% |
| Treg WT1 | TGTGCCAGCAGTTTTGGACAGAGCTCCTATGAACAGTACTTC | 1 | TRBV15*00 | TRBD1*00 | TRBJ2-7*00 | 0.39% |
| Treg WT1 | TGTGCCAGCAGTTTAGTGGGTTCTGGAAATACGCTCTATTTT | 1 | TRBV15*00 | | TRBJ1-3*00 | 0.39% |
| Treg WT1 | TGTGCCAGCAGTTCCGGGACAGGGGGCGAAAGATTATTTTTC | 1 | TRBV15*00 | TRBD1*00 | TRBJ1-4*00 | 0.39% |
| Treg WT1 | TGTGCCAGCAGTTGGGGACAGATCTATGCTGAGCAGTTCTTC | 1 | TRBV15*00 | TRBD1*00 | TRBJ2-1*00 | 0.39% |
| Treg WT1 | TGTGCCAGCAGTCTCGACAATTCTGGAAATACGCTCTATTTT | 1 | TRBV26*00 | | TRBJ1-3*00 | 0.39% |
| Treg WT1 | TGTGCCAGCAGTCTGGGACAGGGCAACGAAAGATTATTTTTC | 1 | TRBV26*00 | TRBD1*00 | TRBJ1-4*00 | 0.39% |
| Treg WT1 | TGTGCCAGCAGTCCCAGGGGTAACTATGCTGAGCAGTTCTTC | 1 | TRBV14*00 | TRBD1*00 | TRBJ2-1*00 | 0.39% |
| Treg WT1 | TGTGCCAGCAGTGTGGGGGAAGACCAAGACACCCAGTACTTT | 1 | TRBV13-3*00 | TRBD2*00 | TRBJ2-5*00 | 0.39% |
| Treg WT1 | TGTGCCAGCAGTGATTCGGGGGGTGCAAAAACGCTGTATTTT | 1 | TRBV13-1*00 | TRBD1*00 | TRBJ2-3*00 | 0.39% |
| Treg WT1 | TGTGCCAGCAGTGATCCAGGTTCTGGAAATACGCTCTATTTT | 1 | TRBV13-3*00 | | TRBJ1-3*00 | 0.39% |
| Treg WT1 | TGTGCCAGCAGTGATCAGGGAAGGTACTCCGACTACACCTTC | 1 | TRBV13-1*00 | TRBD1*00 | TRBJ1-2*00 | 0.39% |
| Treg WT1 | TGTGCCAGCAGTGATGCTGGGGGCTCCTATGAACAGTACTTC | 1 | TRBV13-1*00 | TRBD2*00 | TRBJ2-7*00 | 0.39% |
| Treg WT1 | TGTGCCAGCAGTGATGCGGCAGGGACCACAGAAGTCTTCTTT | 1 | TRBV13-1*00 | TRBD1*00 | TRBJ1-1*00 | 0.39% |
| Treg WT1 | TGTGCCAGCAGTGAATCAGGGACGAGGGACACCCAGTACTTT | 1 | TRBV13-1*00 | TRBD1*00 | TRBJ2-5*00 | 0.39% |
| Treg WT1 | TGTGCCAGCAGCTTCCGGGACAGGGGTGACACCCAGTACTTT | 1 | TRBV3*00 | TRBD1*00 | TRBJ2-5*00 | 0.39% |
| Treg WT1 | TGTGCCAGCAGCTTCGACAGTTCTGGAAATACGCTCTATTTT | 1 | TRBV3*00 | TRBD1*00 | TRBJ1-3*00 | 0.39% |
| Treg WT1 | TGTGCCAGCAGCTTATCCGGGGTCTATGCTGAGCAGTTCTTC | 1 | TRBV3*00 | TRBD1*00 | TRBJ2-1*00 | 0.39% |
| Treg WT1 | TGTGCCAGCAGCTTACGGGGGTTTCAAAACACCTTGTACTTT | 1 | TRBV3*00 | TRBD1*00 | TRBJ2-4*00 | 0.39% |
| Treg WT1 | TGTGCCAGCAGCTTAGGGACAGGGATCTCCGACTACACCTTC | 1 | TRBV3*00 | TRBD1*00 | TRBJ1-2*00 | 0.39% |
| Treg WT1 | TGTGCCAGCAGCTCAGACAGGGGCTATGCTGAGCAGTTCTTC | 1 | TRBV3*00 | TRBD1*00 | TRBJ2-1*00 | 0.39% |

| | | | | | | |
|---|---|---|---|---|---|---|
| Treg WT1 | TGTGCCAGCAGCCCCTCCGGGACAAACACAGAAGTCTTCTTT | 1 | TRBV5*00 | TRBD1*00 | TRBJ1-1*00 | 0.39% |
| Treg WT1 | TGTGCCAGCAGCCCGGGACAGTACCAAGACACCCAGTACTTT | 1 | TRBV5*00 | TRBD1*00 | TRBJ2-5*00 | 0.39% |
| Treg WT1 | TGTGCCAGCAGCCACTGGGAGAGTCAAAACACCTTGTACTTT | 1 | TRBV13-3*00 | TRBD2*00 | TRBJ2-4*00 | 0.39% |
| Treg WT1 | TGTGCCAGCAGCCAAGACGGACAAAACTCCGACTACACCTTC | 1 | TRBV2*00 | TRBD1*00 | TRBJ1-2*00 | 0.39% |
| Treg WT1 | TGTGCCAGCAGCCAAGAGGGGGGCTATGCTGAGCAGTTCTTC | 1 | TRBV2*00 | TRBD2*00 | TRBJ2-1*00 | 0.39% |
| Treg WT1 | TGTGCCAGCAGCCAAGAGACAGGGAACTCCGACTACACCTTC | 1 | TRBV2*00 | TRBD1*00 | TRBJ1-2*00 | 0.39% |
| Treg WT1 | TGTGCCAGCAGCCAAGAACAGGGAGGCTCCGACTACACCTTC | 1 | TRBV5*00 | TRBD1*00 | TRBJ1-2*00 | 0.39% |
| Treg WT1 | TGTGCCAGCAGCCAAGAACAGGGAGGAGCTGAGCAGTTCTTC | 1 | TRBV5*00 | TRBD1*00 | TRBJ2-1*00 | 0.39% |
| Treg WT1 | TGTGCCAGCAACCCGGGACTGGGGGTTGCTGAGCAGTTCTTC | 1 | TRBV15*00 | TRBD2*00 | TRBJ2-1*00 | 0.39% |
| Treg WT1 | TGTGCAAGCAGCTTAGCGGGACACTATGCTGAGCAGTTCTTC | 1 | TRBV16*00 | TRBD1*00 | TRBJ2-1*00 | 0.39% |
| Treg WT1 | TGTGCAAGCAGCTTAGATCCAAGTCAAAACACCTTGTACTTT | 1 | TRBV16*00 | | TRBJ2-4*00 | 0.39% |
| Treg WT1 | TGTGCAAGCAGCTTAGAGGCCCCCGCGGACACCCAGTACTTT | 1 | TRBV16*00 | TRBD1*00 | TRBJ2-5*00 | 0.39% |
| Treg WT1 | TGTGCAAGCAGCCCCGACACGAACACCGGGCAGCTCTACTTT | 1 | TRBV16*00 | | TRBJ2-2*00 | 0.39% |
| Treg WT1 | TGTGCAAGCAGCCGGACAACAAACACCGGGCAGCTCTACTTT | 1 | TRBV16*00 | TRBD1*00 | TRBJ2-2*00 | 0.39% |
| Treg WT1 | TGTGGTGCTAGGGATCTGGGGGGGCGCTCTGAGCAGTTCTTC | 1 | TRBV20*00 | TRBD2*00 | TRBJ2-1*00 | 0.39% |
| Treg WT1 | TGCTCCAGCAGTCAATCTGGGGGGTCGGACACCCAGTACTTT | 1 | TRBV23*00 | TRBD2*00 | TRBJ2-5*00 | 0.39% |
| Treg WT1 | TGCACCTGCAGTGCGCCAGGGAACTATGCTGAGCAGTTCTTC | 1 | TRBV1*00 | TRBD1*00 | TRBJ2-1*00 | 0.39% |
| Treg WT1 | TGCACCTGCAGTGCAGGGCGGAACTATGCTGAGCAGTTCTTC | 1 | TRBV1*00 | | TRBJ2-1*00 | 0.39% |
| Treg WT1 | TGCACCTGCAGTGCAGGACATTCTGGAAATACGCTCTATTTT | 1 | TRBV1*00 | | TRBJ1-3*00 | 0.39% |
| Treg WT1 | TGCACCTGCAGTGCAGATCCGGCAAACTCCGACTACACCTTC | 1 | TRBV1*00 | | TRBJ1-2*00 | 0.39% |
| Treg WT1 | TGCACCTGCAGTGCAGACAATTCTGGAAATACGCTCTATTTT | 1 | TRBV1*00 | | TRBJ1-3*00 | 0.39% |
| Treg WT1 | TGCACCTGCAGTAAACAGCATTCTGGAAATACGCTCTATTTT | 1 | TRBV1*00 | | TRBJ1-3*00 | 0.39% |
| Treg WT1 | TGTGCTAGCAGTTCAGTTAACCAAGACACCCAGTACTTT | 1 | TRBV29*00 | | TRBJ2-5*00 | 0.39% |
| Treg WT1 | TGTGCTAGCAGTCTCACAGGGAACTCCGACTACACCTTC | 1 | TRBV21*00 | TRBD1*00 | TRBJ1-2*00 | 0.39% |
| Treg WT1 | TGTGCCTGGAGTCTGGCAGCAAACTCCGACTACACCTTC | 1 | TRBV31*00 | | TRBJ1-2*00 | 0.39% |
| Treg WT1 | TGTGCCTGGAGTCTATTTGGGGGGGCTGAGCAGTTCTTC | 1 | TRBV31*00 | TRBD2*00 | TRBJ2-1*00 | 0.39% |
| Treg WT1 | TGTGCCTGGAGTCTACAGGGAAACTCCGACTACACCTTC | 1 | TRBV31*00 | TRBD1*00 | TRBJ1-2*00 | 0.39% |
| Treg WT1 | TGTGCCTGGAGTCACTGGAGCTCCTATGAACAGTACTTC | 1 | TRBV31*00 | | TRBJ2-7*00 | 0.39% |
| Treg WT1 | TGTGCCTGGAGTAGGGGGGCAAACTCCGACTACACCTTC | 1 | TRBV31*00 | TRBD1*00 | TRBJ1-2*00 | 0.39% |
| Treg WT1 | TGTGCCAGCTCTCTCCTGGGGGGCTATGAACAGTACTTC | 1 | TRBV12-2*00 | TRBD2*00 | TRBJ2-7*00 | 0.39% |
| Treg WT1 | TGTGCCAGCTCTCTCGCCCAGTATGCTGAGCAGTTCTTC | 1 | TRBV12-1*00 | TRBD2*00 | TRBJ2-1*00 | 0.39% |
| Treg WT1 | TGTGCCAGCTCTCTCGACGGGAACTCCGACTACACCTTC | 1 | TRBV12-1*00 | | TRBJ1-2*00 | 0.39% |
| Treg WT1 | TGTGCCAGCTCTCTCGAACGGGGTGAAACGCTGTATTTT | 1 | TRBV12-1*00 | | TRBJ2-3*00 | 0.39% |
| Treg WT1 | TGTGCCAGCGGTGATGACTTTCAAAACACCTTGTACTTT | 1 | TRBV13-2*00 | | TRBJ2-4*00 | 0.39% |
| Treg WT1 | TGTGCCAGCGGTGATAACTGGTCCTATGAACAGTACTTC | 1 | TRBV13-2*00 | TRBD2*00 | TRBJ2-7*00 | 0.39% |
| Treg WT1 | TGTGCCAGCGGTGACCACATCCAAGACACCCAGTACTTT | 1 | TRBV13-2*00 | | TRBJ2-5*00 | 0.39% |
| Treg WT1 | TGTGCCAGCGGTGAGGGGGGTGCAGAAACGCTGTATTTT | 1 | TRBV13-2*00 | TRBD1*00 | TRBJ2-3*00 | 0.39% |
| Treg WT1 | TGTGCCAGCAGTTTGGACTGGGGGTATGAACAGTACTTC | 1 | TRBV15*00 | TRBD2*00 | TRBJ2-7*00 | 0.39% |
| Treg WT1 | TGTGCCAGCAGTTTAGGGGGGAACGAAAGATTATTTTTC | 1 | TRBV15*00 | TRBD1*00 | TRBJ1-4*00 | 0.39% |
| Treg WT1 | TGTGCCAGCAGTTCGGATTCTGGAAATACGCTCTATTTT | 1 | TRBV15*00 | | TRBJ1-3*00 | 0.39% |
| Treg WT1 | TGTGCCAGCAGTCTCTACAGTCAAGACACCCAGTACTTT | 1 | TRBV24*00 | | TRBJ2-5*00 | 0.39% |
| Treg WT1 | TGTGCCAGCAGTCTGGGCAGCTCCTATGAACAGTACTTC | 1 | TRBV26*00 | | TRBJ2-7*00 | 0.39% |
| Treg WT1 | TGTGCCAGCAGTCTGAACAGGGGGCAGGAACAGTACTTC | 1 | TRBV26*00 | TRBD1*00 | TRBJ2-7*00 | 0.39% |
| Treg WT1 | TGTGCCAGCAGTCAAACAGGGAACGAAAGATTATTTTTC | 1 | TRBV24*00 | TRBD1*00 | TRBJ1-4*00 | 0.39% |
| Treg WT1 | TGTGCCAGCAGTGGGCACAGGGGGCGTGAACAGTACTTC | 1 | TRBV13-1*00 | TRBD1*00 | TRBJ2-7*00 | 0.39% |
| Treg WT1 | TGTGCCAGCAGTGATGGGGGGAACTCCGACTACACCTTC | 1 | TRBV13-1*00 | TRBD1*00 | TRBJ1-2*00 | 0.39% |
| Treg WT1 | TGTGCCAGCAGTGACTGGGGGGAAAACACCTTGTACTTT | 1 | TRBV13-1*00 | TRBD2*00 | TRBJ2-4*00 | 0.39% |
| Treg WT1 | TGTGCCAGCAGTGACAGGGACGACGAAAGATTATTTTTC | 1 | TRBV13-1*00 | TRBD1*00 | TRBJ1-4*00 | 0.39% |
| Treg WT1 | TGTGCCAGCAGTGAGGGGGGGGTTGCTGAGCAGTTCTTC | 1 | TRBV13-3*00 | TRBD2*00 | TRBJ2-1*00 | 0.39% |
| Treg WT1 | TGTGCCAGCAGCTTCCGGGACAGCCAGGCTCCGCTTTTT | 1 | TRBV3*00 | TRBD1*00 | TRBJ1-5*00 | 0.39% |
| Treg WT1 | TGTGCCAGCAGCTTGGCAGGGTTCGGGCAGCTCTACTTT | 1 | TRBV3*00 | TRBD1*00 | TRBJ2-2*00 | 0.39% |
| Treg WT1 | TGTGCCAGCAGCTTGGAGGGCGATGCTGAGCAGTTCTTC | 1 | TRBV3*00 | TRBD1*00 | TRBJ2-1*00 | 0.39% |
| Treg WT1 | TGTGCCAGCAGCTTAGACTTAAACACAGAAGTCTTCTTT | 1 | TRBV3*00 | | TRBJ1-1*00 | 0.39% |
| Treg WT1 | TGTGCCAGCAGCTTAGACAGGAACTCCGACTACACCTTC | 1 | TRBV3*00 | TRBD1*00 | TRBJ1-2*00 | 0.39% |
| Treg WT1 | TGTGCCAGCAGCTTAGAGGGACAACAGGCTCCGCTTTTT | 1 | TRBV3*00 | TRBD1*00 | TRBJ1-5*00 | 0.39% |
| Treg WT1 | TGTGCCAGCAGCTGGGGGGGGCCGGCTGAGCAGTTCTTC | 1 | TRBV4*00 | TRBD2*00 | TRBJ2-1*00 | 0.39% |
| Treg WT1 | TGTGCCAGCAGCTACAGAAACACCGGGCAGCTCTACTTT | 1 | TRBV3*00 | | TRBJ2-2*00 | 0.39% |
| Treg WT1 | TGTGCCAGCAGCCCCCCGGGTGGGGACACCCAGTACTTT | 1 | TRBV2*00 | TRBD1*00 | TRBJ2-5*00 | 0.39% |
| Treg WT1 | TGTGCCAGCAGCCCCCAGAGCTCCTATGAACAGTACTTC | 1 | TRBV5*00 | | TRBJ2-7*00 | 0.39% |
| Treg WT1 | TGTGCCAGCAGCCCGACACCAAACACAGAAGTCTTCTTT | 1 | TRBV26*00 | | TRBJ1-1*00 | 0.39% |
| Treg WT1 | TGTGCCAGCAGCCGGGACTGGGATGCTGAGCAGTTCTTC | 1 | TRBV2*00 | TRBD2*00 | TRBJ2-1*00 | 0.39% |
| Treg WT1 | TGTGCCAGCAGCCGGGACACAAACACAGAAGTCTTCTTT | 1 | TRBV2*00 | TRBD1*00 | TRBJ1-1*00 | 0.39% |
| Treg WT1 | TGTGCCAGCAGCCAGGGACAGCAAGACACCCAGTACTTT | 1 | TRBV5*00 | TRBD1*00 | TRBJ2-5*00 | 0.39% |
| Treg WT1 | TGTGCCAGCAGCCAAGACCAGGAAGGCACCCAGTACTTT | 1 | TRBV5*00 | | TRBJ2-5*00 | 0.39% |
| Treg WT1 | TGTGCCAGCAGCCAAGACAGGAACTCCGACTACACCTTC | 1 | TRBV5*00 | | TRBJ1-2*00 | 0.39% |
| Treg WT1 | TGTGCCAGCAGCCAAGAGAGAAACACAGAAGTCTTCTTT | 1 | TRBV5*00 | | TRBJ1-1*00 | 0.39% |
| Treg WT1 | TGTGCCAGCAGCGACAGTAACCAAGACACCCAGTACTTT | 1 | TRBV14*00 | TRBD1*00 | TRBJ2-5*00 | 0.39% |
| Treg WT1 | TGTGCCAGCAGGGACAATTCTGGAAATACGCTCTATTTT | 1 | TRBV13-3*00 | | TRBJ1-3*00 | 0.39% |
| Treg WT1 | TGTGCCAGCAAAAACCTTAACCAAGACACCCAGTACTTT | 1 | TRBV13-1*00 | | TRBJ2-5*00 | 0.39% |
| Treg WT1 | TGTGCAAGCAGCTTCCGGGCAAACTCCGACTACACCTTC | 1 | TRBV16*00 | | TRBJ1-2*00 | 0.39% |
| Treg WT1 | TGTGCAAGCAGCTTAGGGACTGGAAATACGCTCTATTTT | 1 | TRBV16*00 | | TRBJ1-3*00 | 0.39% |
| Treg WT1 | TGTGCAAGCAGCTTAGATACCAACTATGAACAGTACTTC | 1 | TRBV16*00 | | TRBJ2-7*00 | 0.39% |
| Treg WT1 | TGTGGTGCTAGCGGGACTGGGCAAGACACCCAGTACTTC | 1 | TRBV20*00 | TRBD2*00 | TRBJ2-5*00 | 0.39% |
| Treg WT1 | TGTGGTGCCGGGGGGGGTGGTGCAGAAACGCTGTATTTT | 1 | TRBV20*00 | TRBD2*00 | TRBJ2-3*00 | 0.39% |
| Treg WT1 | TGCACCTGTGGACAGTCAAACACCGGGCAGCTCTACTTT | 1 | TRBV1*00 | TRBD1*00 | TRBJ2-2*00 | 0.39% |
| Treg WT1 | TGCACCTGCAGTGCTGGGGGGCCCTATGAACAGTACTTC | 1 | TRBV1*00 | TRBD2*00 | TRBJ2-7*00 | 0.39% |
| Treg WT1 | TGCACCTGCAGTGATAAGGGGCAAGACACCCAGTACTTT | 1 | TRBV1*00 | TRBD1*00 | TRBJ2-5*00 | 0.39% |
| Treg WT1 | TGTGCCAGCAGTACTCAAACACCGGGCAGCTCTACTTT | 1 | TRBV19*00 | | TRBJ2-2*00 | 0.39% |
| Treg WT1 | TGTGCCAGCTCTCTCTTCGGGGGGGAACAGTACTTC | 1 | TRBV12-1*00 | TRBD2*00 | TRBJ2-7*00 | 0.39% |
| Treg WT1 | TGTGCCAGCTCTCTCGGGAACTCCGACTACACCTTC | 1 | TRBV12-1*00 | | TRBJ1-2*00 | 0.39% |
| Treg WT1 | TGTGCCAGCTCCCTGGGGTTCTATGAACAGTACTTC | 1 | TRBV12-1*00 | TRBD2*00 | TRBJ2-7*00 | 0.39% |
| Treg WT1 | TGTGCCAGCGGTGATGCTAACTCCGACTACACCTTC | 1 | TRBV13-2*00 | | TRBJ1-2*00 | 0.39% |
| Treg WT1 | TGTGCCAGCGGTGAGGGGGCGTATGAACAGTACTTC | 1 | TRBV13-2*00 | TRBD1*00 | TRBJ2-7*00 | 0.39% |
| Treg WT1 | TGTGCCAGCACCGGTTCTGGAAATACGCTCTATTTT | 1 | TRBV19*00 | | TRBJ1-3*00 | 0.39% |
| Treg WT1 | TGTGCCAGCAGTTTCTGGGGCACAGAAGTCTTCTTT | 1 | TRBV22*00 | TRBD2*00 | TRBJ1-1*00 | 0.39% |

| | | | | | | |
|---|---|---|---|---|---|---|
| Treg WT1 | TGTGCCAGCAGTTTGAGTCAAAACACCTTGTACTTT | 1 | TRBV14*00 | | TRBJ2-4*00 | 0.39% |
| Treg WT1 | TGTGCCAGCAGTCTCAACACCGGGCAGCTCTACTTT | 1 | TRBV26*00 | | TRBJ2-2*00 | 0.39% |
| Treg WT1 | TGTGCCAGCAGTCTGATAAACACAGAAGTCTTCTTT | 1 | TRBV26*00 | | TRBJ1-1*00 | 0.39% |
| Treg WT1 | TGTGCCAGCAGTCCCAGTCAAAACACCTTGTACTTT | 1 | TRBV15*00 | | TRBJ2-4*00 | 0.39% |
| Treg WT1 | TGTGCCAGCAGTCATTCTGGAAATACGCTCTATTTT | 1 | TRBV24*00 | | TRBJ1-3*00 | 0.39% |
| Treg WT1 | TGTGCCAGCAGTGGGACTGGGGGTTCCCAGTACTTT | 1 | TRBV13-1*00 | TRBD2*00 | TRBJ2-5*00 | 0.39% |
| Treg WT1 | TGTGCCAGCAGTGATGCAGGGGGAGAACAGTACTTC | 1 | TRBV13-1*00 | TRBD1*00 | TRBJ2-7*00 | 0.39% |
| Treg WT1 | TGTGCCAGCAGTGAGGGGGGGGATGAACAGTACTTC | 1 | TRBV13-1*00 | TRBD2*00 | TRBJ2-7*00 | 0.39% |
| Treg WT1 | TGTGCCAGCAGTGAAGGGGGGGAGTGAACAGTACTTC | 1 | TRBV13-3*00 | TRBD1*00 | TRBJ2-7*00 | 0.39% |
| Treg WT1 | TGTGCCAGCAGTATCAGCTGGGGTGAACAGTACTTC | 1 | TRBV19*00 | TRBD2*00 | TRBJ2-7*00 | 0.39% |
| Treg WT1 | TGTGCCAGCAGTATGGGAAACTCCGACTACACCTTC | 1 | TRBV19*00 | | TRBJ1-2*00 | 0.39% |
| Treg WT1 | TGTGCCAGCAGCTTTCAGGGGCAAGAAGTCTTCTTT | 1 | TRBV3*00 | TRBD1*00 | TRBJ1-1*00 | 0.39% |
| Treg WT1 | TGTGCCAGCAGCTTGACATACACAGAAGTCTTCTTT | 1 | TRBV25*00 | | TRBJ1-1*00 | 0.39% |
| Treg WT1 | TGTGCCAGCAGCTTGAGCTCCTATGAACAGTACTTC | 1 | TRBV3*00 | | TRBJ2-7*00 | 0.39% |
| Treg WT1 | TGTGCCAGCAGCTTAGCTGGGTATGAACAGTACTTC | 1 | TRBV3*00 | | TRBJ2-7*00 | 0.39% |
| Treg WT1 | TGTGCCAGCAGCTTAGCAGGATATGAACAGTACTTC | 1 | TRBV3*00 | | TRBJ2-7*00 | 0.39% |
| Treg WT1 | TGTGCCAGCAGCTTAGGGGATTCCGACTACACCTTC | 1 | TRBV3*00 | | TRBJ1-2*00 | 0.39% |
| Treg WT1 | TGTGCCAGCAGCTCCTGGGGGAACACCTTGTACTTT | 1 | TRBV3*00 | TRBD2*00 | TRBJ2-4*00 | 0.39% |
| Treg WT1 | TGTGCCAGCAGCTCCAGGTACTCCGACTACACCTTC | 1 | TRBV3*00 | | TRBJ1-2*00 | 0.39% |
| Treg WT1 | TGTGCCAGCAGCTCCAGAAACTCCGACTACACCTTC | 1 | TRBV3*00 | | TRBJ1-2*00 | 0.39% |
| Treg WT1 | TGTGCCAGCAGCTACAGCCCTGGGGCTCCGCTTTTT | 1 | TRBV4*00 | TRBD1*00 | TRBJ1-5*00 | 0.39% |
| Treg WT1 | TGTGCCAGCAGCCTTAGTGCAGAAACGCTGTATTTT | 1 | TRBV3*00 | | TRBJ2-3*00 | 0.39% |
| Treg WT1 | TGTGCCAGCAGCCCCGGGTATGCTGAGCAGTTCTTC | 1 | TRBV15*00 | TRBD1*00 | TRBJ2-1*00 | 0.39% |
| Treg WT1 | TGTGCCAGCAGCCCCGACAGGGTTGAACAGTACTTC | 1 | TRBV2*00 | TRBD1*00 | TRBJ2-7*00 | 0.39% |
| Treg WT1 | TGTGCCAGCAGCCCCAGGGACCAGGCTCCGCTTTTT | 1 | TRBV2*00 | TRBD1*00 | TRBJ1-5*00 | 0.39% |
| Treg WT1 | TGTGCCAGCAGCCGGGGTCAAAACACCTTGTACTTT | 1 | TRBV3*00 | | TRBJ2-4*00 | 0.39% |
| Treg WT1 | TGTGCCAGCAGCCGGGACAACGAAAGATTATTTTTC | 1 | TRBV2*00 | | TRBJ1-4*00 | 0.39% |
| Treg WT1 | TGTGCCAGCAGCCGACATCAAAACACCTTGTACTTT | 1 | TRBV2*00 | | TRBJ2-4*00 | 0.39% |
| Treg WT1 | TGTGCCAGCAGCCGACAGAACGAAAGATTATTTTTC | 1 | TRBV3*00 | TRBD1*00 | TRBJ1-4*00 | 0.39% |
| Treg WT1 | TGTGCCAGCAGCCAAGAGCAAAACACCTTGTACTTT | 1 | TRBV2*00 | | TRBJ2-4*00 | 0.39% |
| Treg WT1 | TGTGCCAGCAGCCAAGAAAACACAGAAGTCTTCTTT | 1 | TRBV5*00 | | TRBJ1-1*00 | 0.39% |
| Treg WT1 | TGTGCCAGCAGCGCCCGACAGGGGGACTACACCTTC | 1 | TRBV2*00 | TRBD1*00 | TRBJ1-2*00 | 0.39% |
| Treg WT1 | TGTGCCAGCAGGGAGACAGACTATGAACAGTACTTC | 1 | TRBV3*00 | TRBD1*00 | TRBJ2-7*00 | 0.39% |
| Treg WT1 | TGTGCCAGCAGACTGGGGGTAGGTGAACAGTACTTC | 1 | TRBV13-3*00 | TRBD2*00 | TRBJ2-7*00 | 0.39% |
| Treg WT1 | TGTGCAAGCAGCTTAGGGAACACAGAAGTCTTCTTT | 1 | TRBV16*00 | | TRBJ1-1*00 | 0.39% |
| Treg WT1 | TGTGCAAGCAGCTTAGATCAAGGGGGGCAGTACTTC | 1 | TRBV16*00 | TRBD1*00 | TRBJ2-7*00 | 0.39% |
| Treg WT1 | TGTGCAAGCAGCTTAGAGGACACAGAAGTCTTCTTT | 1 | TRBV16*00 | | TRBJ1-1*00 | 0.39% |
| Treg WT1 | TGTGGTGCTAGGACTGGGTATGCTGAGCAGTTCTTC | 1 | TRBV20*00 | TRBD2*00 | TRBJ2-1*00 | 0.39% |
| Treg WT1 | TGCTCCAGCAGTCAATCGGGGGACACCCAGTACTTT | 1 | TRBV23*00 | | TRBJ2-5*00 | 0.39% |
| Treg WT1 | TGCACCTGCAGTGCAGATCCGGACACCTTGTACTTT | 1 | TRBV1*00 | | TRBJ2-4*00 | 0.39% |
| Treg WT1 | TGTGCCAGCTCTCTGGAAAACACCTTGTACTTT | 1 | TRBV12-1*00 | | TRBJ2-4*00 | 0.39% |
| Treg WT1 | TGTGCCAGCAGTTTCCAAGCTGAGCAGTTCTTC | 1 | TRBV15*00 | | TRBJ2-1*00 | 0.39% |
| Treg WT1 | TGTGCCAGCAGTTTGGATGCTGAGCAGTTCTTC | 1 | TRBV26*00 | | TRBJ2-1*00 | 0.39% |
| Treg WT1 | TGTGCCAGCAGTTTAGCCGGAGAAGTCTTCTTT | 1 | TRBV15*00 | | TRBJ1-1*00 | 0.39% |
| Treg WT1 | TGTGCCAGCAGTCGCTCCTATGAACAGTACTTC | 1 | TRBV15*00 | | TRBJ2-7*00 | 0.39% |
| Treg WT1 | TGTGCCAGCAGTGCCTCCTATGAACAGTACTTC | 1 | TRBV15*00 | | TRBJ2-7*00 | 0.39% |
| Treg WT1 | TGTGCCAGCAGTGGGGACTCCGACTACACCTTC | 1 | TRBV13-1*00 | | TRBJ1-2*00 | 0.39% |
| Treg WT1 | TGTGCCAGCAGCTTTCAAAACACCTTGTACTTT | 1 | TRBV3*00 | | TRBJ2-4*00 | 0.39% |
| Treg WT1 | TGTGCCAGCAGCTTGCGGGAAACGCTGTATTTT | 1 | TRBV3*00 | | TRBJ2-3*00 | 0.39% |
| Treg WT1 | TGTGCCAGCAGCTTAAGGGACACCCAGTACTTT | 1 | TRBV3*00 | | TRBJ2-5*00 | 0.39% |
| Treg WT1 | TGTGCCAGCAGCTTAAAGTATGAACAGTACTTC | 1 | TRBV3*00 | | TRBJ2-7*00 | 0.39% |
| Treg WT1 | TGTGCCAGCAGCTGGGATGCTGAGCAGTTCTTC | 1 | TRBV3*00 | | TRBJ2-1*00 | 0.39% |
| Treg WT1 | TGTGCCAGCAGCCCGCAAGACACCCAGTACTTT | 1 | TRBV15*00 | | TRBJ2-5*00 | 0.39% |
| Treg WT1 | TGTGCCAGCAGCCAAGAAGAGGAACAGTACTTC | 1 | TRBV5*00 | | TRBJ2-7*00 | 0.39% |
| Treg WT1 | TGTGCCAGCAGCGTTGGAAACACCTTGTACTTT | 1 | TRBV15*00 | | TRBJ2-4*00 | 0.39% |
| Treg WT1 | TGTGCCAGCAGCACTGGGGAAACGCTGTATTTT | 1 | TRBV15*00 | TRBD2*00 | TRBJ2-3*00 | 0.39% |
| Treg WT1 | TGTGGTGCTCAAAACTATGCTGAGCAGTTCTTC | 1 | TRBV20*00 | | TRBJ2-1*00 | 0.39% |
| Treg WT1 | TGCACCTGCAGTGGCACCGGGCAGCTCTACTTT | 1 | TRBV1*00 | | TRBJ2-2*00 | 0.39% |
| Treg WT1 | TGTGCCTGGAGTCTAGGACACCTTGTACTTT | 1 | TRBV31*00 | | TRBJ2-4*00 | 0.39% |
| Treg WT1 | TGTGCTAGCAGTCAAGACACCCAGTACTTT | 1 | TRBV29*00 | | TRBJ2-5*00 | 0.39% |
| Treg WT1 | TGTGCCAGCAGCCGGACAGAAGTCTTCTTT | 1 | TRBV3*00 | | TRBJ1-1*00 | 0.39% |
| Treg WT1 | TGTGCCAGCAAAAACACAGAAGTCTTCTTT | 1 | TRBV19*00 | | TRBJ1-1*00 | 0.39% |

#### Treg WT2

| Sample name | CDR3 Sequence | clone Count | best V Hit | best D Hit | best J Hit | Frequency |
|---|---|---|---|---|---|---|
| Treg WT2 | TGTGCCAGCAGCCAAGGGACAAACACAGAAGTCTTCTTT | 3 | TRBV5*00 | | TRBJ1-1*00 | 1.20% |
| Treg WT2 | TGTGCCAGCAGGCAACAGAACACAGAAGTCTTCTTT | 3 | TRBV19*00 | | TRBJ1-1*00 | 1.20% |
| Treg WT2 | TGTGCCAGCAGCCAAGAACAGGGGAACACCGGGCAGCTCTACTTT | 2 | TRBV5*00 | TRBD1*00 | TRBJ2-2*00 | 0.80% |
| Treg WT2 | TGTGCCAGCAGTTCCTGGGGGCGGACCGGGCAGCTCTACTTT | 2 | TRBV2*00 | TRBD1*00 | TRBJ2-2*00 | 0.80% |
| Treg WT2 | TGTGCCAGCAGCTTAGTAGGGAACACCGGGCAGCTCTACTTT | 2 | TRBV3*00 | | TRBJ2-2*00 | 0.80% |
| Treg WT2 | TGCACCTGCAGTGAACAGCATTCTGGAAATACGCTCTATTTT | 2 | TRBV1*00 | | TRBJ1-3*00 | 0.80% |
| Treg WT2 | TGTGCCAGCTCTTTCGACATACAAAACACCTTGTACTTT | 2 | TRBV12-1*00 | | TRBJ2-4*00 | 0.80% |
| Treg WT2 | TGTGCCAGCAGTTGGGGGGACTATGCTGAGCAGTTCTTC | 2 | TRBV15*00 | TRBD2*00 | TRBJ2-1*00 | 0.80% |
| Treg WT2 | TGTGCCAGCAGTGGGGACAGGGGAGATGAACAGTACTTC | 2 | TRBV13-1*00 | TRBD1*00 | TRBJ2-7*00 | 0.80% |
| Treg WT2 | TGTGCCAGCAGCGGACAGAGTCAAAACACCTTGTACTTT | 2 | TRBV3*00 | TRBD1*00 | TRBJ2-4*00 | 0.80% |
| Treg WT2 | TGTGCCAGCTCTCTGGGGAACTCCGACTACACCTTC | 2 | TRBV12-1*00 | | TRBJ1-2*00 | 0.80% |
| Treg WT2 | TGTGCCAGCGGTGCCCTATATGCTGAGCAGTTCTTC | 2 | TRBV13-2*00 | | TRBJ2-1*00 | 0.80% |
| Treg WT2 | TGTGCCAGCAGTTTAGACAGTTATGAACAGTACTTC | 2 | TRBV15*00 | | TRBJ2-7*00 | 0.80% |
| Treg WT2 | TGTGCCAGCAGTCTGCCAGATGCTGAGCAGTTCTTC | 2 | TRBV26*00 | | TRBJ2-1*00 | 0.80% |
| Treg WT2 | TGTGCCAGCAGTACTAGTGCAGAAACGCTGTATTTT | 2 | TRBV26*00 | | TRBJ2-3*00 | 0.80% |
| Treg WT2 | TGTGCCAGCAGCCTCGACAAGACTGAGCAGTTCTTC | 2 | TRBV5*00 | | TRBJ2-1*00 | 0.80% |
| Treg WT2 | TGTGCCAGCAGCTCGGAGAATACGCTCTATTTT | 2 | TRBV2*00 | | TRBJ1-3*00 | 0.80% |

| | | | | | | |
|---|---|---|---|---|---|---|
| Treg WT2 | GTGCCAGCAGCCAAGATCACAGGGTCGGGAACAACCAGGCTCCGCTTTT | 1 | TRBV5*00 | TRBD1*00 | TRBJ1-5*00 | 0.40% |
| Treg WT2 | GTGCCAGCAGCCAAGAGGGACTGGGGACTAGTGCAGAAACGCTGCATTT | 1 | TRBV5*00 | TRBD2*00 | TRBJ2-3*00 | 0.40% |
| Treg WT2 | TGTGCCAGCGGTGATGCACCCGGGACAGCAAACTCCGACTACACCTTC | 1 | TRBV13-2*00 | TRBD1*00 | TRBJ1-2*00 | 0.40% |
| Treg WT2 | TGTGCCAGCAGCCGACAGGGTTCTTCCTATAATTCGCCCCTCTACTTT | 1 | TRBV5*00 | TRBD1*00 | TRBJ1-6*00 | 0.40% |
| Treg WT2 | TGTGCCAGCAGCATACAGGGGGGTAATTCTGGAAATACGCTCTATTTT | 1 | TRBV3*00 | TRBD1*00 | TRBJ1-3*00 | 0.40% |
| Treg WT2 | TGTGCTAGCAGTTTACAGGACTGGAACACCGGGCAGCTCTACTTT | 1 | TRBV29*00 | TRBD2*00 | TRBJ2-2*00 | 0.40% |
| Treg WT2 | TGTGCCTGGAGTCCTAGGGGAACTAGTGCAGAAACGCTGTATTTT | 1 | TRBV31*00 | TRBD1*00 | TRBJ2-3*00 | 0.40% |
| Treg WT2 | TGTGCCAGCTCTCTCTGGGGGAGTTCCACCGGGCAGCTCTACTTT | 1 | TRBV12-1*00 | TRBD2*00 | TRBJ2-2*00 | 0.40% |
| Treg WT2 | TGTGCCAGCTCTCTCGAACAGGGGAGTGCAGAAACGCTGTATTTT | 1 | TRBV12-1*00 | TRBD1*00 | TRBJ2-3*00 | 0.40% |
| Treg WT2 | TGTGCCAGCTCTCCTCCGGGACAGAGCTCCTATGAACAGTACTTC | 1 | TRBV12-1*00 | TRBD1*00 | TRBJ2-7*00 | 0.40% |
| Treg WT2 | TGTGCCAGCTCTCCATCAGGGGAGGCCTATGCTGAGCAGTTCTTC | 1 | TRBV12-1*00 | TRBD1*00 | TRBJ2-1*00 | 0.40% |
| Treg WT2 | TGTGCCAGCGGTGATGCTGGGGGGGCGCGGAACACCCAGTACTTT | 1 | TRBV13-2*00 | TRBD2*00 | TRBJ2-5*00 | 0.40% |
| Treg WT2 | TGTGCCAGCGGTGATGCTGGGAGTCACACCGGGCAGCTCTACTTT | 1 | TRBV13-2*00 | | TRBJ2-2*00 | 0.40% |
| Treg WT2 | TGTGCCAGCAGTTTCCAACAGGGGGCGGGGGCTGAGCAGTTCTTC | 1 | TRBV14*00 | TRBD1*00 | TRBJ2-1*00 | 0.40% |
| Treg WT2 | TGTGCCAGCAGTGCATGGGGGACTAGTGCAGAAACGCTGTATTTT | 1 | TRBV19*00 | TRBD2*00 | TRBJ2-3*00 | 0.40% |
| Treg WT2 | TGTGCCAGCAGTGATGCAAGGCAGGGAGGCTCCGACTACACCTTC | 1 | TRBV13-3*00 | TRBD1*00 | TRBJ1-2*00 | 0.40% |
| Treg WT2 | TGTGCCAGCAGTGATGACTGGGGTAGTCAAAACACCTTGTACTTT | 1 | TRBV13-1*00 | TRBD2*00 | TRBJ2-4*00 | 0.40% |
| Treg WT2 | TGTGCCAGCAGTATTCCTGGGGGTAACTATGCTGAGCAGTTCTTC | 1 | TRBV19*00 | TRBD2*00 | TRBJ2-1*00 | 0.40% |
| Treg WT2 | TGTGCCAGCAGTATAGTCCGGGACACAAACTCCGACTACACCTTC | 1 | TRBV19*00 | TRBD1*00 | TRBJ1-2*00 | 0.40% |
| Treg WT2 | TGTGCCAGCAGTATAGCGGGGACAGGGGTCCATGAACAGTACTTC | 1 | TRBV19*00 | TRBD1*00 | TRBJ2-7*00 | 0.40% |
| Treg WT2 | TGTGCCAGCAGCTTACGGACTGGGGGGGCGCGAGAACAGTACTTC | 1 | TRBV3*00 | TRBD2*00 | TRBJ2-7*00 | 0.40% |
| Treg WT2 | TGTGCCAGCAGCTTAGAGGGGACAGCAAACACAGAAGTCTTCTTT | 1 | TRBV3*00 | TRBD1*00 | TRBJ1-1*00 | 0.40% |
| Treg WT2 | TGTGCCAGCAGCTCTGACCGGGACTGGGGAGAAACGCTGTATTTT | 1 | TRBV25*00 | TRBD2*00 | TRBJ2-3*00 | 0.40% |
| Treg WT2 | TGTGCCAGCAGCCAGGGACTGTCTAGTCAAAACACCTTGTACTTT | 1 | TRBV3*00 | TRBD2*00 | TRBJ2-4*00 | 0.40% |
| Treg WT2 | TGTGCCAGCAGCCAAGGGACTGGAAACTATGCTGAGCAGTTCTTC | 1 | TRBV5*00 | TRBD2*00 | TRBJ2-1*00 | 0.40% |
| Treg WT2 | TGTGCAAGCAGCTTAGGGTGGGTCTCTGGAAATACGCTCTATTTT | 1 | TRBV16*00 | | TRBJ1-3*00 | 0.40% |
| Treg WT2 | TGTGGTGCTGGAGGGACAGGGGTTAGTGCAGAAACGCTGTATTTT | 1 | TRBV20*00 | TRBD1*00 | TRBJ2-3*00 | 0.40% |
| Treg WT2 | TGCACCTGCAGTGCAGATGACAGGAACAACCAGGCTCCGCTTTTT | 1 | TRBV1*00 | TRBD1*00 | TRBJ1-5*00 | 0.40% |
| Treg WT2 | TGCACCTGCAGTGCAGAGACACATTCTGGAAATACGCTCTATTTT | 1 | TRBV1*00 | | TRBJ1-3*00 | 0.40% |
| Treg WT2 | TGTGCTAGCAGTTTAGGACAGAGGGACTATGAACAGTACTTC | 1 | TRBV29*00 | TRBD1*00 | TRBJ2-7*00 | 0.40% |
| Treg WT2 | TGTGCCTGGAGTCTCGGACTGGGGGGGCATGAACAGTACTTC | 1 | TRBV31*00 | TRBD2*00 | TRBJ2-7*00 | 0.40% |
| Treg WT2 | TGTGCCTGGAGTCTAGCAGGGGAGAGAGACACCCAGTACTTT | 1 | TRBV31*00 | TRBD1*00 | TRBJ2-5*00 | 0.40% |
| Treg WT2 | TGTGCCTGGAGGCGGACTGGGGGGGCCTATGAACAGTACTTC | 1 | TRBV31*00 | TRBD2*00 | TRBJ2-7*00 | 0.40% |
| Treg WT2 | TGTGCCAGCTCTCTGTGGGGGGGTCAAAACACCTTGTACTTT | 1 | TRBV12-1*00 | TRBD2*00 | TRBJ2-4*00 | 0.40% |
| Treg WT2 | TGTGCCAGCTCCCCACAGGGTTCTGGAAATACGCTCTATTTT | 1 | TRBV12-1*00 | TRBD1*00 | TRBJ1-3*00 | 0.40% |
| Treg WT2 | TGTGCCAGCGCCTGGACAGATTCTGGAAATACGCTCTATTTT | 1 | TRBV13-1*00 | TRBD1*00 | TRBJ1-3*00 | 0.40% |
| Treg WT2 | TGTGCCAGCGGTGATGCTGGGAGTCAAAACACCTTGTACTTT | 1 | TRBV13-2*00 | | TRBJ2-4*00 | 0.40% |
| Treg WT2 | TGTGCCAGCGGTGATGCCTGGGGGGCGTATGAACAGTACTTC | 1 | TRBV13-2*00 | TRBD2*00 | TRBJ2-7*00 | 0.40% |
| Treg WT2 | TGTGCCAGCGGTGATGCGGGGGTCTCCTATGAACAGTACTTC | 1 | TRBV13-2*00 | TRBD1*00 | TRBJ2-7*00 | 0.40% |
| Treg WT2 | TGTGCCAGCACCCCGACTGGGGGGACCGGGCAGCTCTACTTT | 1 | TRBV13-3*00 | TRBD2*00 | TRBJ2-2*00 | 0.40% |
| Treg WT2 | TGTGCCAGCAGTTTCCCTCGACAGAATTCGCCCCTCTACTTT | 1 | TRBV19*00 | TRBD1*00 | TRBJ1-6*00 | 0.40% |
| Treg WT2 | TGTGCCAGCAGTTTGGCGGCGAGCTATGCTGAGCAGTTCTTC | 1 | TRBV15*00 | | TRBJ2-1*00 | 0.40% |
| Treg WT2 | TGTGCCAGCAGTTTGGACTGGGGGGCGTATGAACAGTACTTC | 1 | TRBV14*00 | TRBD2*00 | TRBJ2-7*00 | 0.40% |
| Treg WT2 | TGTGCCAGCAGTCTTGGGACATCAAACTCCGACTACACCTTC | 1 | TRBV15*00 | TRBD1*00 | TRBJ1-2*00 | 0.40% |
| Treg WT2 | TGTGCCAGCAGTCGACTGGGGGGGCGGGCTGAGCAGTTCTTC | 1 | TRBV14*00 | TRBD2*00 | TRBJ2-1*00 | 0.40% |
| Treg WT2 | TGTGCCAGCAGTGGACAGGGTTCTGGAAATACGCTCTATTTT | 1 | TRBV13-1*00 | TRBD1*00 | TRBJ1-3*00 | 0.40% |
| Treg WT2 | TGTGCCAGCAGTGATGCAGCGAGCTCCTATGAACAGCACTTC | 1 | TRBV13-3*00 | | TRBJ2-7*00 | 0.40% |
| Treg WT2 | TGTGCCAGCAGTGATGCAAGGGCAGGGTTTGAACAGTACTTC | 1 | TRBV13-1*00 | TRBD1*00 | TRBJ2-7*00 | 0.40% |
| Treg WT2 | TGTGCCAGCAGTGATAGGGGGAACTATGCTGAGCAGTTCTTC | 1 | TRBV13-1*00 | TRBD1*00 | TRBJ2-1*00 | 0.40% |
| Treg WT2 | TGTGCCAGCAGTATCGGGACTGGGGGGAACACCTTGTACTTT | 1 | TRBV15*00 | TRBD2*00 | TRBJ2-4*00 | 0.40% |
| Treg WT2 | TGTGCCAGCAGTATGCGGGGGAGCACCGGGCAGCTCTACTTT | 1 | TRBV19*00 | TRBD1*00 | TRBJ2-2*00 | 0.40% |
| Treg WT2 | TGTGCCAGCAGTATGACGGGACCGGGAGCTGAGCAGTTCTTC | 1 | TRBV19*00 | TRBD1*00 | TRBJ2-1*00 | 0.40% |
| Treg WT2 | TGTGCCAGCAGTATAGAACAGGGGAATTATGAACAGTACTTC | 1 | TRBV19*00 | TRBD1*00 | TRBJ2-7*00 | 0.40% |
| Treg WT2 | TGTGCCAGCAGTACGACAGATTCTGGAAATACGCTCTATTTT | 1 | TRBV13-3*00 | TRBD1*00 | TRBJ1-3*00 | 0.40% |
| Treg WT2 | TGTGCCAGCAGTAGGGGACAGGGGGGCTCCGACTACACCTTC | 1 | TRBV19*00 | TRBD1*00 | TRBJ1-2*00 | 0.40% |
| Treg WT2 | TGTGCCAGCAGCTTGGGACTGGGCCCCTATGAACAGTACTTC | 1 | TRBV3*00 | TRBD2*00 | TRBJ2-7*00 | 0.40% |
| Treg WT2 | TGTGCCAGCAGCTTACCTGGGAACACCGGGCAGCTCTACTTT | 1 | TRBV3*00 | TRBD2*00 | TRBJ2-2*00 | 0.40% |
| Treg WT2 | TGTGCCAGCAGCTTACGGTCTAGTCAAAACACCTTGTACTTT | 1 | TRBV3*00 | | TRBJ2-4*00 | 0.40% |
| Treg WT2 | TGTGCCAGCAGCTTACACAGGGGGCTCTCCGACTACACCTTC | 1 | TRBV3*00 | TRBD1*00 | TRBJ1-2*00 | 0.40% |
| Treg WT2 | TGTGCCAGCAGCTTAGGGCTGGGGGGGCGTGAACAGTACTTC | 1 | TRBV3*00 | TRBD2*00 | TRBJ2-7*00 | 0.40% |
| Treg WT2 | TGTGCCAGCAGCTTAGGGCAGGCAAACTCCGACTACACCTTC | 1 | TRBV3*00 | | TRBJ1-2*00 | 0.40% |
| Treg WT2 | TGTGCCAGCAGCTTAGAGGGAAGTGCAGAAACGCTGTATTTT | 1 | TRBV3*00 | | TRBJ2-3*00 | 0.40% |
| Treg WT2 | TGTGCCAGCAGCTCACAGGGGAACACCGGGCAGCTCTACTTT | 1 | TRBV3*00 | TRBD1*00 | TRBJ2-2*00 | 0.40% |
| Treg WT2 | TGTGCCAGCAGCTGGACTTTTAACCAAGACACCCAGTACTTT | 1 | TRBV4*00 | TRBD2*00 | TRBJ2-5*00 | 0.40% |
| Treg WT2 | TGTGCCAGCAGCCTACAGGGAGAGGCAGAAACGCTGTATTTT | 1 | TRBV5*00 | TRBD1*00 | TRBJ2-3*00 | 0.40% |
| Treg WT2 | TGTGCCAGCAGCCCCCGGGACAGGGGGCGGGAACAGTACTTC | 1 | TRBV5*00 | TRBD1*00 | TRBJ2-7*00 | 0.40% |
| Treg WT2 | TGTGCCAGCAGCCGGACAATTTCTGGAAATACGCTCTATTTT | 1 | TRBV3*00 | TRBD1*00 | TRBJ1-3*00 | 0.40% |
| Treg WT2 | TGTGCCAGCAGCCAAGGGGAAAACACCGGGCAGCTCTACTTT | 1 | TRBV2*00 | | TRBJ2-2*00 | 0.40% |
| Treg WT2 | TGTGCCAGCAGCCAAGGGATTAACCAAGACACCCAGTACTTT | 1 | TRBV5*00 | | TRBJ2-5*00 | 0.40% |
| Treg WT2 | TGTGCCAGCAGCCAAGATTACAACCAAGACACCCAGTACTTT | 1 | TRBV5*00 | | TRBJ2-5*00 | 0.40% |
| Treg WT2 | TGTGCCAGCAGCCAAGACGGGACAATCACAGAAGTCTTCTTT | 1 | TRBV2*00 | TRBD1*00 | TRBJ1-1*00 | 0.40% |
| Treg WT2 | TGTGCCAGCAGCCAAGACATAAACACCGGGCAGCTCTACTTT | 1 | TRBV2*00 | | TRBJ2-2*00 | 0.40% |
| Treg WT2 | TGTGCCAGCAGCGGCTGGGGGGGTGCAGAAACGCTGTATTTT | 1 | TRBV2*00 | TRBD2*00 | TRBJ2-3*00 | 0.40% |
| Treg WT2 | TGTGCAAGCAGCTTCGGGGGTAGTGCAGAAACGCTGTATTTT | 1 | TRBV16*00 | TRBD1*00 | TRBJ2-3*00 | 0.40% |

| | | | | | | |
|---|---|---|---|---|---|---|
| Treg WT2 | TGTGCAAGCAGCTTAGAGAGGGCCCCCTATGAACAGTACTTC | 1 | TRBV16*00 | | TRBJ2-7*00 | 0.40% |
| Treg WT2 | TGTGCAAGCAGCTCACAGGGAAACCAAGACACCCAGTACTTT | 1 | TRBV16*00 | TRBD1*00 | TRBJ2-5*00 | 0.40% |
| Treg WT2 | TGTGGTGCTAGGGATCAGTTTTCCAACGAAAGATTATTTTTC | 1 | TRBV20*00 | | TRBJ1-4*00 | 0.40% |
| Treg WT2 | TGCTCCAGCAGTCCCTCGGGGGGCACCGGGCAGCTCTACTTT | 1 | TRBV23*00 | TRBD2*00 | TRBJ2-2*00 | 0.40% |
| Treg WT2 | TGCACCTGCACCCCCGGGGCTAGTGCAGAAACGCTGTATTTT | 1 | TRBV1*00 | TRBD1*00 | TRBJ2-3*00 | 0.40% |
| Treg WT2 | TGCACCTGCAGTGCACAGATCCCTGGAAATACGCTCTATTTT | 1 | TRBV1*00 | | TRBJ1-3*00 | 0.40% |
| Treg WT2 | TGCACCTGCAGTGCAGGGTTTTCCAACGAAAGATTATTTTTC | 1 | TRBV1*00 | | TRBJ1-4*00 | 0.40% |
| Treg WT2 | TGCACCTGCAGTGCAGATCCGAGCAACGAAAGATTATTTTTC | 1 | TRBV1*00 | | TRBJ1-4*00 | 0.40% |
| Treg WT2 | TGCACCTGCAGTGGACCGGGACAGGGGGCTGAGCAGTTCTTC | 1 | TRBV1*00 | TRBD1*00 | TRBJ2-1*00 | 0.40% |
| Treg WT2 | TGTGCTAGCAGTATCCTTAGTGCAGAAACGCTGTATTTT | 1 | TRBV17*00 | | TRBJ2-3*00 | 0.40% |
| Treg WT2 | TGTGCTAGCAGTAGTTGGGGTCAAAACACCTTGTACTTT | 1 | TRBV17*00 | | TRBJ2-4*00 | 0.40% |
| Treg WT2 | TGTGCTAGCAGTAGACAGGCGGGGACAGAAGTCTTCTTT | 1 | TRBV17*00 | | TRBJ1-1*00 | 0.40% |
| Treg WT2 | TGTGCCTGGAGTTTCAGGGGAAGCGAAAGATTATTTTTC | 1 | TRBV31*00 | TRBD1*00 | TRBJ1-4*00 | 0.40% |
| Treg WT2 | TGTGCCTGGAGTCTGGGGGGGCTGGCTGAGCAGTTCTTC | 1 | TRBV31*00 | TRBD2*00 | TRBJ2-1*00 | 0.40% |
| Treg WT2 | TGTGCCTGGAGTCTAGCCAGAAACTCCGACTACACCTTC | 1 | TRBV31*00 | | TRBJ1-2*00 | 0.40% |
| Treg WT2 | TGTGCCTGGAGTCCTTGGGACTCCTATGAACAGTACTTC | 1 | TRBV31*00 | | TRBJ2-7*00 | 0.40% |
| Treg WT2 | TGTGCCTGGAGTCCGGACTGGGTAGCTGAGCAGTTCTTC | 1 | TRBV31*00 | TRBD2*00 | TRBJ2-1*00 | 0.40% |
| Treg WT2 | TGTGCCAGCTCTCTCCAGAACACCGGGCAGCTCTACTTT | 1 | TRBV12-1*00 | | TRBJ2-2*00 | 0.40% |
| Treg WT2 | TGTGCCAGCTCTCTCGAAAACTCCTATGAACAGTACTTC | 1 | TRBV12-1*00 | | TRBJ2-7*00 | 0.40% |
| Treg WT2 | TGTGCCAGCTCTCTACCATCTGGAAATACGCTCTATTTT | 1 | TRBV12-2*00 | | TRBJ1-3*00 | 0.40% |
| Treg WT2 | TGTGCCAGCTCTCGACTGGGTAGGGCTGAGCAGTTCTTC | 1 | TRBV12-1*00 | TRBD2*00 | TRBJ2-1*00 | 0.40% |
| Treg WT2 | TGTGCCAGCCACAGATTTTCTGGAAATACGCTCTATTTT | 1 | TRBV12-2*00 | | TRBJ1-3*00 | 0.40% |
| Treg WT2 | TGTGCCAGCGGTGTGGGACAATATGCTGAGCAGTTCTTC | 1 | TRBV13-2*00 | TRBD1*00 | TRBJ2-1*00 | 0.40% |
| Treg WT2 | TGTGCCAGCGGTGGGACAGGACATGCTGAGCAGTTCTTC | 1 | TRBV13-2*00 | TRBD1*00 | TRBJ2-1*00 | 0.40% |
| Treg WT2 | TGTGCCAGCGGTGAACGGGGTGCAGAAACGCTGTATTTT | 1 | TRBV13-2*00 | | TRBJ2-3*00 | 0.40% |
| Treg WT2 | TGTGCCAGCGGGACTGGGGTCTATGCTGAGCAGTTCTTC | 1 | TRBV13-2*00 | TRBD2*00 | TRBJ2-1*00 | 0.40% |
| Treg WT2 | TGTGCCAGCAGTTTTGCAAACACCGGGCAGCTCTACTTT | 1 | TRBV14*00 | | TRBJ2-2*00 | 0.40% |
| Treg WT2 | TGTGCCAGCAGTTTCGGGACAGTTTATGAACAGTACTTT | 1 | TRBV15*00 | TRBD1*00 | TRBJ2-7*00 | 0.40% |
| Treg WT2 | TGTGCCAGCAGTTTCAGGGAAGACTCCGACTACACCTTC | 1 | TRBV13-3*00 | TRBD1*00 | TRBJ1-2*00 | 0.40% |
| Treg WT2 | TGTGCCAGCAGTTCCATTAACAACCAGGCTCCGCTTTTT | 1 | TRBV14*00 | | TRBJ1-5*00 | 0.40% |
| Treg WT2 | TGTGCCAGCAGTCTGTCAGGGTTCTCCGACTACACCTTC | 1 | TRBV26*00 | | TRBJ1-2*00 | 0.40% |
| Treg WT2 | TGTGCCAGCAGTCTGGTCAACCAAGACACCCAGTACTTC | 1 | TRBV26*00 | | TRBJ2-5*00 | 0.40% |
| Treg WT2 | TGTGCCAGCAGTCTGATTAACCAAGACACCCAGTACTTT | 1 | TRBV24*00 | | TRBJ2-5*00 | 0.40% |
| Treg WT2 | TGTGCCAGCAGTCTACAGGGGAGGGACACCCAGTACTTT | 1 | TRBV24*00 | TRBD1*00 | TRBJ2-5*00 | 0.40% |
| Treg WT2 | TGTGCCAGCAGTCCCGTGGGTGCAGAAACGCTGTATTTT | 1 | TRBV15*00 | | TRBJ2-3*00 | 0.40% |
| Treg WT2 | TGTGCCAGCAGTCCCGGGGGCAACTCCGACTACACCTTC | 1 | TRBV26*00 | TRBD1*00 | TRBJ1-2*00 | 0.40% |
| Treg WT2 | TGTGCCAGCAGTCGGGACTGGGCCTATGAACAGTACTTC | 1 | TRBV19*00 | TRBD2*00 | TRBJ2-7*00 | 0.40% |
| Treg WT2 | TGTGCCAGCAGTGTCGACAGGGGCTATGAACAGTACTTC | 1 | TRBV22*00 | | TRBJ2-7*00 | 0.40% |
| Treg WT2 | TGTGCCAGCAGTGTGGGTAACCAAGACACCCAGTACTTT | 1 | TRBV13-3*00 | | TRBJ2-5*00 | 0.40% |
| Treg WT2 | TGTGCCAGCAGTGGTCGACAGGACTGGCCCCTCTACTTT | 1 | TRBV13-3*00 | TRBD1*00 | TRBJ1-6*00 | 0.40% |
| Treg WT2 | TGTGCCAGCAGTGGGGACAGTGCAGAAACGCTGTATTTT | 1 | TRBV13-1*00 | TRBD1*00 | TRBJ2-3*00 | 0.40% |
| Treg WT2 | TGTGCCAGCAGTGGACATAACTATGCTGAGCAGTTCTTC | 1 | TRBV26*00 | | TRBJ2-1*00 | 0.40% |
| Treg WT2 | TGTGCCAGCAGTGATGCAGGGACAGGGGAAGTCTTCTTT | 1 | TRBV13-1*00 | TRBD1*00 | TRBJ1-1*00 | 0.40% |
| Treg WT2 | TGTGCCAGCAGTGATGGTGGGGCGGACACCCAGTACTTT | 1 | TRBV13-1*00 | TRBD1*00 | TRBJ2-5*00 | 0.40% |
| Treg WT2 | TGTGCCAGCAGTGATGGGCCCACCGGGCAGCTCTACTTT | 1 | TRBV13-1*00 | | TRBJ2-2*00 | 0.40% |
| Treg WT2 | TGTGCCAGCAGTGATGGGCAGCAAGACACCCAGTACTTT | 1 | TRBV13-1*00 | | TRBJ2-5*00 | 0.40% |
| Treg WT2 | TGTGCCAGCAGTGACAGGGGAGGAAACACCTTGTACTTT | 1 | TRBV13-1*00 | TRBD1*00 | TRBJ2-4*00 | 0.40% |
| Treg WT2 | TGTGCCAGCAGTGACAACTATAATTCGCCCCTCTACTTT | 1 | TRBV13-3*00 | | TRBJ1-6*00 | 0.40% |
| Treg WT2 | TGTGCCAGCAGTGAAAGGGGGAACACAGAAGTCTTCTTT | 1 | TRBV13-1*00 | TRBD1*00 | TRBJ1-1*00 | 0.40% |
| Treg WT2 | TGTGCCAGCAGTAGGGACAAAAACTCCGACTACACCTTC | 1 | TRBV19*00 | TRBD1*00 | TRBJ1-2*00 | 0.40% |
| Treg WT2 | TGTGCCAGCAGCTTTGCTGGGACAGAAACGCTGTATTTT | 1 | TRBV3*00 | TRBD2*00 | TRBJ2-3*00 | 0.40% |
| Treg WT2 | TGTGCCAGCAGCTTTGACAGGGAAGACACCCAGTACTTT | 1 | TRBV3*00 | TRBD1*00 | TRBJ2-5*00 | 0.40% |
| Treg WT2 | TGTGCCAGCAGCTTTACTAGTCAAAACACCTTGTACTTT | 1 | TRBV3*00 | | TRBJ2-4*00 | 0.40% |
| Treg WT2 | TGTGCCAGCAGCTTCACAGGTCAAAACACCTTGTACTTT | 1 | TRBV3*00 | | TRBJ2-4*00 | 0.40% |
| Treg WT2 | TGTGCCAGCAGCTTAGTTACAAACTCCGACTACACCTTC | 1 | TRBV3*00 | | TRBJ1-2*00 | 0.40% |
| Treg WT2 | TGTGCCAGCAGCTTAGAGGGGAACGAAAGATTATTTTTC | 1 | TRBV3*00 | TRBD1*00 | TRBJ1-4*00 | 0.40% |
| Treg WT2 | TGTGCCAGCAGCTTAATTGGGGGAAACACCTTGTACTTT | 1 | TRBV3*00 | TRBD2*00 | TRBJ2-4*00 | 0.40% |
| Treg WT2 | TGTGCCAGCAGCTTAATCAGCCTTTATGAACAGTACTTC | 1 | TRBV3*00 | | TRBJ2-7*00 | 0.40% |
| Treg WT2 | TGTGCCAGCAGCTTAACTGCAAACTCCGACTACACCTTC | 1 | TRBV3*00 | | TRBJ1-2*00 | 0.40% |
| Treg WT2 | TGTGCCAGCAGCTTAAAGCAAAACACAGAAGTCTTCTTT | 1 | TRBV3*00 | | TRBJ1-1*00 | 0.40% |
| Treg WT2 | TGTGCCAGCAGCTGGACTGGTCAAAACACCTTGTACTTT | 1 | TRBV3*00 | | TRBJ2-4*00 | 0.40% |
| Treg WT2 | TGTGCCAGCAGCTACCTGGGTCAAGACACCCAGTACTTT | 1 | TRBV4*00 | TRBD2*00 | TRBJ2-5*00 | 0.40% |
| Treg WT2 | TGTGCCAGCAGCCTGACAGCAAACTCCGACTACACCTTC | 1 | TRBV2*00 | | TRBJ1-2*00 | 0.40% |
| Treg WT2 | TGTGCCAGCAGCCAAGAATCAAACACAGAAGTCTTCTTT | 1 | TRBV2*00 | | TRBJ1-1*00 | 0.40% |
| Treg WT2 | TGTGCCAGCAGCCAAAATAGTCAAAACACCTTGTACTTT | 1 | TRBV5*00 | | TRBJ2-4*00 | 0.40% |
| Treg WT2 | TGTGCCAGCAGCAGGACAGGGATTGCTGAGCAGTTCTTC | 1 | TRBV26*00 | TRBD1*00 | TRBJ2-1*00 | 0.40% |
| Treg WT2 | TGTGCCAGCAGCAGATCAGGGTACTCCGACTACACCTTC | 1 | TRBV3*00 | TRBD1*00 | TRBJ1-2*00 | 0.40% |
| Treg WT2 | TGTGCCAGCAGGCGGGCGGCAAACACAGAAGTCTTCTTT | 1 | TRBV3*00 | TRBD1*00 | TRBJ1-1*00 | 0.40% |
| Treg WT2 | TGTGCCAGCAGATCAACAGGGACAGACACCCAGTACTTT | 1 | TRBV19*00 | TRBD1*00 | TRBJ2-5*00 | 0.40% |
| Treg WT2 | TGTGCCAGCAGAAGACACGCAAACTCCGACTACACCTTC | 1 | TRBV15*00 | | TRBJ1-2*00 | 0.40% |
| Treg WT2 | TGTGCAAGCAGCTTAGATCGCTGGTATGAACAGTACTTC | 1 | TRBV16*00 | | TRBJ2-7*00 | 0.40% |
| Treg WT2 | TGTGCAAGCAGCTTAGACAGTCAAAACACCTTGTACTTT | 1 | TRBV16*00 | | TRBJ2-4*00 | 0.40% |
| Treg WT2 | TGTGCAAGCAGCTTAGACAACACCGGGCAGCTCTACTTT | 1 | TRBV16*00 | | TRBJ2-2*00 | 0.40% |
| Treg WT2 | TGTGGTGCTGCCCGGGACAGGGACACAGAAGTCTTCTTT | 1 | TRBV20*00 | TRBD1*00 | TRBJ1-1*00 | 0.40% |

| | | | | | | |
|---|---|---|---|---|---|---|
| Treg WT2 | TGTGGTGCTAGGGTCAGGGGGCGATCCGACTACACCTTC | 1 | TRBV20*00 | TRBD1*00 | TRBJ1-2*00 | 0.40% |
| Treg WT2 | TGTGGTGCTAGGGGCAGGGTGGACTCCGACTACACCTTC | 1 | TRBV20*00 | TRBD1*00 | TRBJ1-2*00 | 0.40% |
| Treg WT2 | TGTGGTGCTAGGGGACAAAACAACCAGGCTCCGCTTTTT | 1 | TRBV20*00 | | TRBJ1-5*00 | 0.40% |
| Treg WT2 | TGTGGTGCCCGGACAGAAAACACCGGGCAGCTCTACTTT | 1 | TRBV20*00 | TRBD1*00 | TRBJ2-2*00 | 0.40% |
| Treg WT2 | TGCTCCAGCAGTCAGCAGGGGGGAAACACCTTGTACTTT | 1 | TRBV23*00 | TRBD1*00 | TRBJ2-4*00 | 0.40% |
| Treg WT2 | TGCACCTGCGGCAGGGCTAGTGCAGAAACGCTGTATTTT | 1 | TRBV1*00 | TRBD1*00 | TRBJ2-3*00 | 0.40% |
| Treg WT2 | TGCACCTGCAGTGCTACTAGTCAAAACACCTTGTACTTT | 1 | TRBV1*00 | | TRBJ2-4*00 | 0.40% |
| Treg WT2 | TGCACCTGCAGTGCAGACTATAATTCGCCCCTCTACTTT | 1 | TRBV1*00 | | TRBJ1-6*00 | 0.40% |
| Treg WT2 | TGCACCTGCAGTGGGGACAGTGCAGAAACGCTGTATTTT | 1 | TRBV1*00 | TRBD1*00 | TRBJ2-3*00 | 0.40% |
| Treg WT2 | TGCACCTGCAGTGGACAGGGGGACTCCGACTACACCTTC | 1 | TRBV1*00 | TRBD1*00 | TRBJ1-2*00 | 0.40% |
| Treg WT2 | TGTGCTAGCAGTTTAATTGGGAACACCTTGTACTTT | 1 | TRBV29*00 | | TRBJ2-4*00 | 0.40% |
| Treg WT2 | TGTGCTAGCAGTCACAGGGGAAACACCTTGTACTTT | 1 | TRBV21*00 | TRBD1*00 | TRBJ2-4*00 | 0.40% |
| Treg WT2 | TGTGCTAGCAGCCTGAACTATGCTGAGCAGTTCTTC | 1 | TRBV17*00 | | TRBJ2-1*00 | 0.40% |
| Treg WT2 | TGTGCTAGCAGCCAGACAGGGGGAGAACAGTACTTC | 1 | TRBV17*00 | TRBD1*00 | TRBJ2-7*00 | 0.40% |
| Treg WT2 | TGTGCTAGCAGAAGGGACAGGATTGAACAGTACTTC | 1 | TRBV21*00 | TRBD1*00 | TRBJ2-7*00 | 0.40% |
| Treg WT2 | TGTGCCTGGGGACAGGGGAGAAATACGCTCTATTTT | 1 | TRBV31*00 | TRBD1*00 | TRBJ1-3*00 | 0.40% |
| Treg WT2 | TGTGCCTGGAGTCCCAACGAAGACACCCAGTACTTT | 1 | TRBV31*00 | | TRBJ2-5*00 | 0.40% |
| Treg WT2 | TGTGCCAGCTCTCTTGGCTCCTATGAACAGTACTTC | 1 | TRBV12-1*00 | | TRBJ2-7*00 | 0.40% |
| Treg WT2 | TGTGCCAGCTCTCTCGAGGGCTATGAACAGTACTTC | 1 | TRBV12-1*00 | | TRBJ2-7*00 | 0.40% |
| Treg WT2 | TGTGCCAGCTCTCTGCGGGGGACAGAAGTCTTCTTT | 1 | TRBV12-1*00 | TRBD1*00 | TRBJ1-1*00 | 0.40% |
| Treg WT2 | TGTGCCAGCTCTCGACAGCAAGACACCCAGTACTTT | 1 | TRBV12-1*00 | TRBD1*00 | TRBJ2-5*00 | 0.40% |
| Treg WT2 | TGTGCCAGCGGTGAGACATATGCTGAGCAGTTCTTC | 1 | TRBV13-2*00 | | TRBJ2-1*00 | 0.40% |
| Treg WT2 | TGTGCCAGCAGTTTTGAGACAGGGGCTCCGCTTTTT | 1 | TRBV15*00 | TRBD1*00 | TRBJ1-5*00 | 0.40% |
| Treg WT2 | TGTGCCAGCAGTGATGCAGGGGGAGAACAGTACTTC | 1 | TRBV13-1*00 | TRBD1*00 | TRBJ2-7*00 | 0.40% |
| Treg WT2 | TGTGCCAGCAGTATATGGGACACAGAAGTCTTCTTT | 1 | TRBV19*00 | | TRBJ1-1*00 | 0.40% |
| Treg WT2 | TGTGCCAGCAGTAGGAACACCGGGCAGCTCTACTTT | 1 | TRBV13-1*00 | | TRBJ2-2*00 | 0.40% |
| Treg WT2 | TGTGCCAGCAGCTTTAGCAAAAACACCTTGTACTTT | 1 | TRBV4*00 | | TRBJ2-4*00 | 0.40% |
| Treg WT2 | TGTGCCAGCAGCTTCGGGGCAGAAACGCTGTATTTT | 1 | TRBV3*00 | | TRBJ2-3*00 | 0.40% |
| Treg WT2 | TGTGCCAGCAGCTTGACTGGGGGTGCGCTGTATTTT | 1 | TRBV3*00 | TRBD2*00 | TRBJ2-3*00 | 0.40% |
| Treg WT2 | TGTGCCAGCAGCTTATACAGAAACACCTTGTACTTT | 1 | TRBV3*00 | | TRBJ2-4*00 | 0.40% |
| Treg WT2 | TGTGCCAGCAGCTTAGCCCAGATTGAACAGTACTTC | 1 | TRBV3*00 | | TRBJ2-7*00 | 0.40% |
| Treg WT2 | TGTGCCAGCAGCTTAGCCACCGGGCAGCTCTACTTT | 1 | TRBV3*00 | | TRBJ2-2*00 | 0.40% |
| Treg WT2 | TGTGCCAGCAGCTTAGGCAACGAAAGATTATTTTTC | 1 | TRBV3*00 | | TRBJ1-4*00 | 0.40% |
| Treg WT2 | TGTGCCAGCAGCTTAGGGGGGAACACCTTGTACTTT | 1 | TRBV3*00 | TRBD1*00 | TRBJ2-4*00 | 0.40% |
| Treg WT2 | TGTGCCAGCAGCTTAGAGACTTATGAACAGTACTTC | 1 | TRBV3*00 | | TRBJ2-7*00 | 0.40% |
| Treg WT2 | TGTGCCAGCAGCCTCGACAGTGCTGAGCAGTTCTCC | 1 | TRBV3*00 | TRBD1*00 | TRBJ2-1*00 | 0.40% |
| Treg WT2 | TGTGCCAGCAGCCCCCGGGATGCTGAGCAGTTCTTC | 1 | TRBV3*00 | TRBD1*00 | TRBJ2-1*00 | 0.40% |
| Treg WT2 | TGTGCCAGCAGCCGACAGGGCTATGAACAGTACTTC | 1 | TRBV3*00 | TRBD1*00 | TRBJ2-7*00 | 0.40% |
| Treg WT2 | TGTGCCAGCAGCCAACAGCAAGACACCCAGTACTTT | 1 | TRBV5*00 | | TRBJ2-5*00 | 0.40% |
| Treg WT2 | TGTGCCAGCAGCACCAGGGCATACACCTTGTACTTT | 1 | TRBV4*00 | TRBD1*00 | TRBJ2-4*00 | 0.40% |
| Treg WT2 | TGTGCCAGCAGGTCGGACTGGGGGACCCAGTACTTT | 1 | TRBV15*00 | TRBD2*00 | TRBJ2-5*00 | 0.40% |
| Treg WT2 | TGTGCCAGCAGGATGGACAGTGCTGAGCAGTTCTTC | 1 | TRBV13-3*00 | TRBD1*00 | TRBJ2-1*00 | 0.40% |
| Treg WT2 | TGTGCCAGCAGGACAGGGGGGATGGAAGTCTTCTTT | 1 | TRBV26*00 | TRBD1*00 | TRBJ1-1*00 | 0.40% |
| Treg WT2 | TGTGCAAGCAGCTTAGATAAAGATGAACAGTACTTC | 1 | TRBV16*00 | | TRBJ2-7*00 | 0.40% |
| Treg WT2 | TGTGGTGCTGGGAGACAAAATGCTGAGCAGTTCTTC | 1 | TRBV20*00 | | TRBJ2-1*00 | 0.40% |
| Treg WT2 | TGTGGTGCTAGGGATCGACAAGACACCCAGTACTTT | 1 | TRBV20*00 | | TRBJ2-5*00 | 0.40% |
| Treg WT2 | TGCACCTGCAGTGGGACAAACGAAAGATTATTTTTC | 1 | TRBV1*00 | TRBD1*00 | TRBJ1-4*00 | 0.40% |
| Treg WT2 | TGCACCTGCAGTGACCAGGGACAGGCTCCGCTTTTT | 1 | TRBV1*00 | TRBD1*00 | TRBJ1-5*00 | 0.40% |
| Treg WT2 | TGTGCCAGCGGTGATACCACAGAAGTCTTCTTT | 1 | TRBV13-2*00 | | TRBJ1-1*00 | 0.40% |
| Treg WT2 | TGTGCCAGCGGTGACAAGACAGAAGTCTTCTTT | 1 | TRBV13-2*00 | | TRBJ1-1*00 | 0.40% |
| Treg WT2 | TGTGCCAGCAGTTTTGACACAGAAGTCTTCTTT | 1 | TRBV15*00 | | TRBJ1-1*00 | 0.40% |
| Treg WT2 | TGTGCCAGCAGTCAGGGCACAGAAGTCTTCTTT | 1 | TRBV15*00 | TRBD1*00 | TRBJ1-1*00 | 0.40% |
| Treg WT2 | TGTGCCAGCAGTGATCGGGACACCCAGTACTTT | 1 | TRBV13-3*00 | | TRBJ2-5*00 | 0.40% |
| Treg WT2 | TGTGCCAGCAGTGATCAAGACACCCAGTACTTT | 1 | TRBV13-3*00 | | TRBJ2-5*00 | 0.40% |
| Treg WT2 | TGTGCCAGCAGCTTCAGGGCAGAAGTCTTCTTT | 1 | TRBV3*00 | TRBD1*00 | TRBJ1-1*00 | 0.40% |
| Treg WT2 | TGTGCCAGCAGCTTAGACACAGAAGTCTTCTTT | 1 | TRBV3*00 | | TRBJ1-1*00 | 0.40% |
| Treg WT2 | TGTGCCAGCAGCTCCTCCTATGAACAGTACTTC | 1 | TRBV3*00 | | TRBJ2-7*00 | 0.40% |
| Treg WT2 | TGTGCCAGCAGCCTGACATATGAACAGTACTTC | 1 | TRBV4*00 | | TRBJ2-7*00 | 0.40% |
| Treg WT2 | TGTGCCAGCAGCGACAACTCCGACTACACCTTC | 1 | TRBV3*00 | | TRBJ1-2*00 | 0.40% |
| Treg WT2 | TGTGCCAGCAGAAACTATGCTGAGCAGTTCTTC | 1 | TRBV15*00 | | TRBJ2-1*00 | 0.40% |
| Treg WT2 | TGTGCAAGCAGCTTGGAAGACACCCAGTACTTT | 1 | TRBV16*00 | | TRBJ2-5*00 | 0.40% |
| Treg WT2 | TGTGCCAGCGGTGGCTATGAACAGTACTTC | 1 | TRBV13-2*00 | | TRBJ2-7*00 | 0.40% |
| Treg WT2 | TGTGCCAGCAGGTAACCGGCTCCGCTTTTT | 1 | TRBV19*00 | | TRBJ1-5*00 | 0.40% |
| Treg WT2 | TGTGCCAGCAGCTGGGAAGTCTTCTTT | 1 | TRBV4*00 | | TRBJ1-1*00 | 0.40% |
| Treg WT2 | TGTGCCAGCCCTATGAACAGTACTTC | 1 | TRBV22*00 | | TRBJ2-7*00 | 0.40% |
| Treg WT2 | TGTGCCCCCGGCCGCTGTATTTT | 1 | TRBV4*00 | TRBD1*00 | TRBJ2-3*00 | 0.40% |

#### Treg WT3

| Sample name | CDR3 Sequence | clone Count | best V Hit | best D Hit | best J Hit | Frequency |
|---|---|---|---|---|---|---|
| Treg WT3 | TGTGCTAGCAGTAGCCCCCCCACCACAAGCTCCTATGAACAGTACTTC | 2 | TRBV17*00 | TRBD2*00 | TRBJ2-7*00 | 1.21% |
| Treg WT3 | TGTGCTAGCAGTTTGACAGGGCGGGGCACAGAAGTCTTCTTT | 2 | TRBV29*00 | TRBD1*00 | TRBJ1-1*00 | 1.21% |
| Treg WT3 | TGTGCTAGCAGTTTAAGGGGAGGACAAAACACCTTGTACTTT | 2 | TRBV29*00 | TRBD1*00 | TRBJ2-4*00 | 1.21% |
| Treg WT3 | TGTGCCAGCTCTTACAATTCCTATAATTCGCCCCTCTACTTT | 2 | TRBV12-2*00 | | TRBJ1-6*00 | 1.21% |
| Treg WT3 | TGTGCCAGCTCTCGGGACAGTAACTATGCTGAGCAGTTCTTC | 2 | TRBV12-1*00 | TRBD1*00 | TRBJ2-1*00 | 1.21% |
| Treg WT3 | TGTGCCAGCAGCCAAGGGACAGCAAACTCCGACTACACCTTC | 2 | TRBV2*00 | TRBD1*00 | TRBJ1-2*00 | 1.21% |

| | | | | | | |
|---|---|---|---|---|---|---|
| Treg WT3 | TGCACCTGCAGTGCACAAAATTCTGGAAATACGCTCTATTTT | 2 | TRBV1*00 | | TRBJ1-3*00 | 1.21% |
| Treg WT3 | TGTGCCAGCAGTATCAATTCTGGAAATACGCTCTATTTT | 2 | TRBV19*00 | | TRBJ1-3*00 | 1.21% |
| Treg WT3 | TGTGCCAGCAGCCAAGAGGGGAACTCCGACTACACCTTC | 2 | TRBV2*00 | | TRBJ1-2*00 | 1.21% |
| Treg WT3 | TGTGCCAGCAGCAGGGCTGCAAACTCCGACTACACCTTC | 2 | TRBV3*00 | | TRBJ1-2*00 | 1.21% |
| Treg WT3 | TGTGCCAGCTCCCCGACAAATTATGAACAGTACTTC | 2 | TRBV18*00 | | TRBJ2-7*00 | 1.21% |
| Treg WT3 | TGTGCCAGCAGCTTAGGTCAAAACACCTTGTACTTT | 2 | TRBV3*00 | | TRBJ2-4*00 | 1.21% |
| Treg WT3 | TGTGCCAGCAGCTACAGGAATTCGCCCCTCTACTTT | 2 | TRBV4*00 | | TRBJ1-6*00 | 1.21% |
| Treg WT3 | TGTGCCAGCAGCCCCGACACCTATGAACAGTACTTC | 2 | TRBV26*00 | | TRBJ2-7*00 | 1.21% |
| Treg WT3 | TGCACCTGCAGTGGGGCCGGGCAGCTCTACTTT | 2 | TRBV1*00 | | TRBJ2-2*00 | 1.21% |
| Treg WT3 | TGTGCCAGCAGCTTACGAGAACAGTACTTC | 2 | TRBV3*00 | | TRBJ2-7*00 | 1.21% |
| Treg WT3 | GTGCCTGGAGTCGAGGCTGGGGGGGCGTCAGTCAAAACACCTTGTACTT | 1 | TRBV31*00 | TRBD2*00 | TRBJ2-4*00 | 0.61% |
| Treg WT3 | TGTGCTAGCAGTTTACCGGGACAGAACAACCAGGCTCCGCTTTTT | 1 | TRBV29*00 | TRBD1*00 | TRBJ1-5*00 | 0.61% |
| Treg WT3 | TGTGCTAGCAGTAGAGCCGGGACAGCAAACTCCGACTACACCTTC | 1 | TRBV17*00 | TRBD1*00 | TRBJ1-2*00 | 0.61% |
| Treg WT3 | TGTGCCAGCGGTGATGTGGGGGGGGTCTATGCTGAGCAGTTCTTC | 1 | TRBV13-2*00 | TRBD2*00 | TRBJ2-1*00 | 0.61% |
| Treg WT3 | TGTGCCAGCGGCCTGACTGGGGGGAGCTCCTATGAACAGTACTTC | 1 | TRBV13-2*00 | TRBD2*00 | TRBJ2-7*00 | 0.61% |
| Treg WT3 | TGTGCCAGCAGTCCCGGACTGGGGGGTGCAGAAACGCTGTATTTT | 1 | TRBV19*00 | TRBD2*00 | TRBJ2-3*00 | 0.61% |
| Treg WT3 | TGTGCCAGCAGTCCCGACCCCTATTCTGGAAATACGCTCTATTTT | 1 | TRBV14*00 | TRBD1*00 | TRBJ1-3*00 | 0.61% |
| Treg WT3 | TGTGCCAGCAGTCCAGGGGGCCATTCTGGAAATACGCTCTATTTT | 1 | TRBV4*00 | TRBD1*00 | TRBJ1-3*00 | 0.61% |
| Treg WT3 | TGTGCCAGCAGTGATGCTGGGGGAAACTATGCTGAGCAGTTCTTC | 1 | TRBV13-1*00 | TRBD2*00 | TRBJ2-1*00 | 0.61% |
| Treg WT3 | TGTGCCAGCAGTATACTGGGGGGGCGCGCAGAAACGCTGTATTTT | 1 | TRBV19*00 | TRBD2*00 | TRBJ2-3*00 | 0.61% |
| Treg WT3 | TGTGCCAGCAGTATAGGGGCAGGGAGCTCCTATGAACAGTACTTC | 1 | TRBV19*00 | | TRBJ2-7*00 | 0.61% |
| Treg WT3 | TGTGCCAGCAGCTTGAGGGGGTCTAGTGCAGAAACGCTGTATTTT | 1 | TRBV3*00 | TRBD1*00 | TRBJ2-3*00 | 0.61% |
| Treg WT3 | TGTGCCAGCAGCTTAGTCGGGGGGGTAAACTCCGACTACACCTTC | 1 | TRBV3*00 | TRBD2*00 | TRBJ1-2*00 | 0.61% |
| Treg WT3 | TGTGCCAGCAGCCGCCGACAGGGAAGTGCAGAAACGCTGTATTTT | 1 | TRBV3*00 | TRBD1*00 | TRBJ2-3*00 | 0.61% |
| Treg WT3 | TGTGCCAGCAGCAGGACTGGGGGAAACACCGGGCAGCTCTACTTT | 1 | TRBV4*00 | TRBD2*00 | TRBJ2-2*00 | 0.61% |
| Treg WT3 | TGCACCTGCAGTGCCGGGGGCCCTAGTGCAGAAACGCTGTATTTT | 1 | TRBV1*00 | TRBD1*00 | TRBJ2-3*00 | 0.61% |
| Treg WT3 | TGCACCTGCAGTGCAGGAGACAGAAACTATGCTGAGCAGTTCTTC | 1 | TRBV1*00 | TRBD1*00 | TRBJ2-1*00 | 0.61% |
| Treg WT3 | TGCACCTGCAGTGCAGATGAGGGTAGTGCAGAAACGCTGTATTTT | 1 | TRBV1*00 | | TRBJ2-3*00 | 0.61% |
| Treg WT3 | TGTGCAAGCAGCTTCCGACAGGGGGAGGCGAAAGATTATTTTTC | 1 | TRBV16*00 | TRBD1*00 | TRBJ1-4*00 | 0.61% |
| Treg WT3 | TGTGCCAGCAGCCCGGGAGGGGAACTATGCTGAGCAGTTCTTC | 1 | TRBV4*00 | TRBD1*00 | TRBJ2-1*00 | 0.61% |
| Treg WT3 | TGTGCTAGCAGTTTCCGGGACAACCAAGACACCCAGTACTTC | 1 | TRBV29*00 | TRBD1*00 | TRBJ2-5*00 | 0.61% |
| Treg WT3 | TGTGCTAGCAGTTTACCGACAATCTATGCTGAGCAGTTCTTC | 1 | TRBV29*00 | | TRBJ2-1*00 | 0.61% |
| Treg WT3 | TGTGCTAGCAGCGCCGGGACAGCCTATGCTGAGCAGTTCTTC | 1 | TRBV21*00 | TRBD1*00 | TRBJ2-1*00 | 0.61% |
| Treg WT3 | TGTGCCTGGAGTCTATCATCTAGTGCAGAAACGCTGTATTTT | 1 | TRBV31*00 | | TRBJ2-3*00 | 0.61% |
| Treg WT3 | TGTGCCTGGAGTCCCGGGACACCAAACTCCGACTACACCTTC | 1 | TRBV31*00 | TRBD1*00 | TRBJ1-2*00 | 0.61% |
| Treg WT3 | TGTGCCAGCTCTCTCCCGGGGACAGGGGTGGCCTTGTACTTT | 1 | TRBV12-1*00 | TRBD1*00 | TRBJ2-4*00 | 0.61% |
| Treg WT3 | TGTGCCAGCTCTCTACAACTTCTCTATGCTGAGCAGTTCTTC | 1 | TRBV12-1*00 | | TRBJ2-1*00 | 0.61% |
| Treg WT3 | TGTGCCAGCGGTGATGCAGGGGCAAACTCCGACTACACCTTC | 1 | TRBV13-2*00 | | TRBJ1-2*00 | 0.61% |
| Treg WT3 | TGTGCCAGCGGTGAGACAACTAGTCAAAACACCTTGTACTTT | 1 | TRBV13-2*00 | | TRBJ2-4*00 | 0.61% |
| Treg WT3 | TGTGCCAGCATCGACAGTTATTCTGGAAATACGCTCTATTTT | 1 | TRBV19*00 | TRBD1*00 | TRBJ1-3*00 | 0.61% |
| Treg WT3 | TGTGCCAGCAGTTTACTGGGGGGTAGTGCTGAGCAGTTCTTC | 1 | TRBV15*00 | TRBD2*00 | TRBJ2-1*00 | 0.61% |
| Treg WT3 | TGTGCCAGCAGTCTGGGACAGTGGAACTCCGACTACACCTTC | 1 | TRBV26*00 | TRBD1*00 | TRBJ1-2*00 | 0.61% |
| Treg WT3 | TGTGCCAGCAGTCTGACGGGAAGAGGCTCCGACTACACCTTC | 1 | TRBV26*00 | TRBD1*00 | TRBJ1-2*00 | 0.61% |
| Treg WT3 | TGTGCCAGCAGTCACAGAAATTCTGGAAATACGCTCTATTTT | 1 | TRBV13-1*00 | | TRBJ1-3*00 | 0.61% |
| Treg WT3 | TGTGCCAGCAGTGATCCGGGAAGTGCAGAAACGCTGTATTTT | 1 | TRBV13-3*00 | TRBD1*00 | TRBJ2-3*00 | 0.61% |
| Treg WT3 | TGTGCCAGCAGTGAGGGACCTCTTGCAGAAACGCTGTATTTT | 1 | TRBV19*00 | TRBD1*00 | TRBJ2-3*00 | 0.61% |
| Treg WT3 | TGTGCCAGCAGTATTGGGAGAGACCAAGACACCCAGTACTTT | 1 | TRBV19*00 | | TRBJ2-5*00 | 0.61% |
| Treg WT3 | TGTGCCAGCAGCTTTGACTGGGGGGGCGCTGAACAGTACTTC | 1 | TRBV3*00 | TRBD2*00 | TRBJ2-7*00 | 0.61% |
| Treg WT3 | TGTGCCAGCAGCTTAGGACCCTCAAACTCCGACTACACCTTC | 1 | TRBV3*00 | | TRBJ1-2*00 | 0.61% |
| Treg WT3 | TGTGCCAGCAGCTTAGATGGGGGGGCGCGTGAACAGTACTTC | 1 | TRBV3*00 | TRBD2*00 | TRBJ2-7*00 | 0.61% |
| Treg WT3 | TGTGCCAGCAGCCCCGACAGTAACCAAGACACCCAGTACTTT | 1 | TRBV2*00 | TRBD1*00 | TRBJ2-5*00 | 0.61% |
| Treg WT3 | TGTGCCAGCAGCCGGACTGGGGGGCAAAACACCTTGTACTTT | 1 | TRBV2*00 | TRBD2*00 | TRBJ2-4*00 | 0.61% |
| Treg WT3 | TGTGCCAGCAGCCGAGGGCGGCGTCAAAACACCTTGTACTTT | 1 | TRBV3*00 | TRBD1*00 | TRBJ2-4*00 | 0.61% |
| Treg WT3 | TGTGCCAGCAGCCAGCAGGGGAACCAAGACACCCAGTACTTT | 1 | TRBV5*00 | TRBD1*00 | TRBJ2-5*00 | 0.61% |
| Treg WT3 | TGTGCCAGCAGCATCAGGGGCGGGCAAGACACCCAGTACTTT | 1 | TRBV3*00 | TRBD1*00 | TRBJ2-5*00 | 0.61% |
| Treg WT3 | TGTGCCAGCAGACTGGGACTTTTCTATGCTGAGCAGTTCTTC | 1 | TRBV13-3*00 | TRBD2*00 | TRBJ2-1*00 | 0.61% |
| Treg WT3 | TGTGCCAGCAACCAGGGGGCTAGTGCAGAAACGCTGTATTTT | 1 | TRBV13-3*00 | TRBD1*00 | TRBJ2-3*00 | 0.61% |
| Treg WT3 | TGTGCCAGCAAAAAGACAGGGGATGACTCCGACTACACCTTC | 1 | TRBV13-1*00 | TRBD1*00 | TRBJ1-2*00 | 0.61% |
| Treg WT3 | TGTGCAAGCAGTCCGGAGGGGGCAAACTCCGACTACACCTTC | 1 | TRBV16*00 | TRBD1*00 | TRBJ1-2*00 | 0.61% |
| Treg WT3 | TGTGCAAGCAGCTGGCAGGGAAGTGCAGAAACGCTGTATTTT | 1 | TRBV16*00 | TRBD1*00 | TRBJ2-3*00 | 0.61% |
| Treg WT3 | TGTGCAAGCAGCAGGACAATTAACCAAGACACCCAGCACTTT | 1 | TRBV16*00 | TRBD1*00 | TRBJ2-5*00 | 0.61% |
| Treg WT3 | TGCACCTGCAGTGCAGGGAATTCTGGAAATACGCTCTATTTT | 1 | TRBV1*00 | | TRBJ1-3*00 | 0.61% |
| Treg WT3 | TGTGCTAGCAGTTTGAGGGGAGGGTCCGACTACACCTTC | 1 | TRBV29*00 | TRBD1*00 | TRBJ1-2*00 | 0.61% |
| Treg WT3 | TGTGCTAGCAGTCCTGGGGGGATAGACACCCAGTACTTT | 1 | TRBV29*00 | TRBD2*00 | TRBJ2-5*00 | 0.61% |
| Treg WT3 | TGTGCTAGCAGGGACAGTAGTGCAGAAACGCTGTATTTT | 1 | TRBV21*00 | TRBD1*00 | TRBJ2-3*00 | 0.61% |
| Treg WT3 | TGTGCCAGCTCTCCTGCAAACACCGGGCAGCTCTACTTT | 1 | TRBV12-1*00 | | TRBJ2-2*00 | 0.61% |
| Treg WT3 | TGTGCCAGCTCTCCGCAGAACACCGGGCAGCTCTACTTT | 1 | TRBV12-1*00 | | TRBJ2-2*00 | 0.61% |
| Treg WT3 | TGTGCCAGCTCTCGACAAGCAAACTCCGACTACACCTTC | 1 | TRBV12-1*00 | | TRBJ1-2*00 | 0.61% |
| Treg WT3 | TGTGCCAGCGGTTGGGGGTACCAAGACACCCAGTACTTT | 1 | TRBV13-2*00 | TRBD2*00 | TRBJ2-5*00 | 0.61% |
| Treg WT3 | TGTGCCAGCGGTCGACAGGCAGGCTCCGACTACACCTTC | 1 | TRBV13-2*00 | TRBD1*00 | TRBJ1-2*00 | 0.61% |
| Treg WT3 | TGTGCCAGCGGTGCCCCTAGTGCAGAAACGCTGTATTTT | 1 | TRBV13-2*00 | | TRBJ2-3*00 | 0.61% |
| Treg WT3 | TGTGCCAGCGGGGGACAGACAAACTCCGACTACACCTTC | 1 | TRBV13-2*00 | TRBD1*00 | TRBJ1-2*00 | 0.61% |
| Treg WT3 | TGTGCCAGCAGTTTTCACAGGGAAAGTGAACAGTACTTC | 1 | TRBV15*00 | TRBD1*00 | TRBJ2-7*00 | 0.61% |
| Treg WT3 | TGTGCCAGCAGTTTTGGGAACAACCAGGCTCCGCTTTTT | 1 | TRBV15*00 | | TRBJ1-5*00 | 0.61% |
| Treg WT3 | TGTGCCAGCAGTTTGTTGGACCAAGACACCCAGTACTTT | 1 | TRBV14*00 | | TRBJ2-5*00 | 0.61% |
| Treg WT3 | TGTGCCAGCAGTTGGGGGGCTGCAGAAACGCTGTATTTT | 1 | TRBV15*00 | TRBD2*00 | TRBJ2-3*00 | 0.61% |
| Treg WT3 | TGTGCCAGCAGTCTCCCCTACCAAGACACCCAGTACTTT | 1 | TRBV24*00 | TRBD1*00 | TRBJ2-5*00 | 0.61% |
| Treg WT3 | TGTGCCAGCAGTCTGTCGAGGCCAAATACGCTCTATTTT | 1 | TRBV26*00 | | TRBJ1-3*00 | 0.61% |
| Treg WT3 | TGTGCCAGCAGTCTGAGGCAGAACTATGAACAGTACTTC | 1 | TRBV26*00 | | TRBJ2-7*00 | 0.61% |
| Treg WT3 | TGTGCCAGCAGTGGTCAAAACTATGCTGAGCAGTTCTTC | 1 | TRBV13-1*00 | | TRBJ2-1*00 | 0.61% |
| Treg WT3 | TGTGCCAGCAGTGGGACAGCAAACACAGAAGTCTTCTTT | 1 | TRBV13-1*00 | TRBD1*00 | TRBJ1-1*00 | 0.61% |
| Treg WT3 | TGTGCCAGCAGTGATCGCAGGGGGCGAGAACAGTACTTC | 1 | TRBV13-1*00 | TRBD1*00 | TRBJ2-7*00 | 0.61% |
| Treg WT3 | TGTGCCAGCAGTGATGAGGGGGAAGACACCCAGTACTTT | 1 | TRBV13-3*00 | TRBD1*00 | TRBJ2-5*00 | 0.61% |
| Treg WT3 | TGTGCCAGCAGTGAAATTTCTGGAAATACGCTCTATTTT | 1 | TRBV13-1*00 | | TRBJ1-3*00 | 0.61% |
| Treg WT3 | TGTGCCAGCAGTGAAAGTTTCGACCAGGCTCCGCTTTTT | 1 | TRBV13-1*00 | | TRBJ1-5*00 | 0.61% |

| | | | | | | |
|---|---|---|---|---|---|---|
| Treg WT3 | TGTGCCAGCAGTATGGGGATTGCAGAAACGCTGTATTTT | 1 | TRBV19*00 | | TRBJ2-3*00 | 0.61% |
| Treg WT3 | TGTGCCAGCAGCTTGGGGGGGCGAGCTGAGCAGTTCTTC | 1 | TRBV3*00 | TRBD2*00 | TRBJ2-1*00 | 0.61% |
| Treg WT3 | TGTGCCAGCAGCTTATTGGGGAACTCCGACTACACCTTC | 1 | TRBV3*00 | TRBD2*00 | TRBJ1-2*00 | 0.61% |
| Treg WT3 | TGTGCCAGCAGCTTATCTAGAAATTCGCCCCTCTACTTT | 1 | TRBV3*00 | | TRBJ1-6*00 | 0.61% |
| Treg WT3 | TGTGCCAGCAGCTGGGGGCGTCAAAACACCTTGTACTTT | 1 | TRBV15*00 | TRBD2*00 | TRBJ2-4*00 | 0.61% |
| Treg WT3 | TGTGCCAGCAGCCCTTTCCCCTCCTATGAACAGTACTTC | 1 | TRBV3*00 | | TRBJ2-7*00 | 0.61% |
| Treg WT3 | TGTGCCAGCAGCCCCTGGGGGGGCTATGAACAGTACTTC | 1 | TRBV2*00 | TRBD2*00 | TRBJ2-7*00 | 0.61% |
| Treg WT3 | TGTGCCAGCAGCCCCCTGGGGTCCTATGAACAGTACTTC | 1 | TRBV2*00 | TRBD2*00 | TRBJ2-7*00 | 0.61% |
| Treg WT3 | TGTGCCAGCAGCCAAGGGGGGTCTTATGAACAGTACTTC | 1 | TRBV5*00 | TRBD1*00 | TRBJ2-7*00 | 0.61% |
| Treg WT3 | TGTGCCAGCAGGCGACTGGGGTCCTATGAACAGTACTTC | 1 | TRBV3*00 | TRBD2*00 | TRBJ2-7*00 | 0.61% |
| Treg WT3 | TGTGCCAGCAGGAAATCTAGTGCAGAAACGCTGTATTTT | 1 | TRBV15*00 | | TRBJ2-3*00 | 0.61% |
| Treg WT3 | TGTGCCAGCAGAGGGACTGGGGGAACTGAGCAGTTCTTC | 1 | TRBV26*00 | TRBD2*00 | TRBJ2-1*00 | 0.61% |
| Treg WT3 | TGTGCAAGCAGCTTAGCAGGGGGCACAGAAGTCTTCTTT | 1 | TRBV16*00 | TRBD1*00 | TRBJ1-1*00 | 0.61% |
| Treg WT3 | TGTGGTGCTGGGGGGAATAACTATGCTGAGCAGTTCTTC | 1 | TRBV20*00 | | TRBJ2-1*00 | 0.61% |
| Treg WT3 | TGTGGTGCTAGTGACAGTAACTATGCTGAGCAGTTCTTC | 1 | TRBV20*00 | TRBD1*00 | TRBJ2-1*00 | 0.61% |
| Treg WT3 | TGTGGTGCTAGCGGGACAGTCTATGCTGAGCAGTTCTTC | 1 | TRBV20*00 | TRBD1*00 | TRBJ2-1*00 | 0.61% |
| Treg WT3 | TGTGGTGCTAGGGGGACAGGGGCAGAAACGCTGTATTTT | 1 | TRBV20*00 | TRBD1*00 | TRBJ2-3*00 | 0.61% |
| Treg WT3 | TGTGGTGCTAGGACAGGGGGCTATGCTGAGCAGTTCTTC | 1 | TRBV20*00 | TRBD1*00 | TRBJ2-1*00 | 0.61% |
| Treg WT3 | TGTGGTGCTAGAGGAACAGGAGGGTCCGACTACACCTTC | 1 | TRBV20*00 | TRBD1*00 | TRBJ1-2*00 | 0.61% |
| Treg WT3 | TGCACCTGCAGTGCGACAGGGATCACAGAAGTCTTCTTT | 1 | TRBV1*00 | TRBD1*00 | TRBJ1-1*00 | 0.61% |
| Treg WT3 | TGCACCTGCAGTGCAGATGAGGGGGACACCCAGTACTTT | 1 | TRBV1*00 | TRBD1*00 | TRBJ2-5*00 | 0.61% |
| Treg WT3 | TGTGCCAGCTCTCTTGACATCTCCGACTACACCTTC | 1 | TRBV12-2*00 | | TRBJ1-2*00 | 0.61% |
| Treg WT3 | TGTGCCAGCTCTCTCCACTATGCTGAGCAGTTCTTC | 1 | TRBV12-1*00 | | TRBJ2-1*00 | 0.61% |
| Treg WT3 | TGTGCCAGCTCTCTCGACAACTATGAACAGTACTTC | 1 | TRBV12-1*00 | | TRBJ2-7*00 | 0.61% |
| Treg WT3 | TGTGCCAGCTCTCAGAACCAAGACACCCAGTACTTT | 1 | TRBV12-2*00 | | TRBJ2-5*00 | 0.61% |
| Treg WT3 | TGTGCCAGCTCACCCCTGGGTGGGGAACAGTACTTC | 1 | TRBV12-1*00 | TRBD1*00 | TRBJ2-7*00 | 0.61% |
| Treg WT3 | TGTGCCAGCGGTGGGACTGGGGTCACCCAGTACTTT | 1 | TRBV13-2*00 | TRBD2*00 | TRBJ2-5*00 | 0.61% |
| Treg WT3 | TGTGCCAGCGGTGATCGCTGGGTTGAACAGTACTTC | 1 | TRBV13-2*00 | TRBD2*00 | TRBJ2-7*00 | 0.61% |
| Treg WT3 | TGTGCCAGCATGACAACAAACTCCGACTACACCTTC | 1 | TRBV2*00 | | TRBJ1-2*00 | 0.61% |
| Treg WT3 | TGTGCCAGCAGTTTCAACACCGGGCAGCTCTACTTT | 1 | TRBV15*00 | | TRBJ2-2*00 | 0.61% |
| Treg WT3 | TGTGCCAGCAGTTTAGGTGGAAATACGCTCTATTTT | 1 | TRBV15*00 | | TRBJ1-3*00 | 0.61% |
| Treg WT3 | TGTGCCAGCAGTGCACAGAGACAGGCTCCGCTTTTT | 1 | TRBV13-3*00 | | TRBJ1-5*00 | 0.61% |
| Treg WT3 | TGTGCCAGCAGTGGGACAAATTCGCCCCTCTACTTT | 1 | TRBV13-1*00 | TRBD1*00 | TRBJ1-6*00 | 0.61% |
| Treg WT3 | TGTGCCAGCAGTGATGCACGGGGGAAAGTCTTCTTT | 1 | TRBV13-1*00 | TRBD1*00 | TRBJ1-1*00 | 0.61% |
| Treg WT3 | TGTGCCAGCAGTGAGGGACACTCCGACTACACCTTC | 1 | TRBV13-3*00 | TRBD1*00 | TRBJ1-2*00 | 0.61% |
| Treg WT3 | TGTGCCAGCAGTGAGGGAGAGAACACCTTGTACTTT | 1 | TRBV13-1*00 | | TRBJ2-4*00 | 0.61% |
| Treg WT3 | TGTGCCAGCAGTACTAGTGCAGAAACGCTGTATTTT | 1 | TRBV19*00 | | TRBJ2-3*00 | 0.61% |
| Treg WT3 | TGTGCCAGCAGTAAGGACATCTATGAACAGTACTTC | 1 | TRBV19*00 | TRBD1*00 | TRBJ2-7*00 | 0.61% |
| Treg WT3 | TGTGCCAGCAGCTTATCTAACTATGAACAGTACTTC | 1 | TRBV3*00 | | TRBJ2-7*00 | 0.61% |
| Treg WT3 | TGTGCCAGCAGCTTACGGGGGTGGGAACAGTACTTC | 1 | TRBV3*00 | TRBD1*00 | TRBJ2-7*00 | 0.61% |
| Treg WT3 | TGTGCCAGCAGCTTAACTGGGAACACCTTGTACTTT | 1 | TRBV3*00 | TRBD2*00 | TRBJ2-4*00 | 0.61% |
| Treg WT3 | TGTGCCAGCAGCTTAAAGTCCTATGAACAGTACTTC | 1 | TRBV3*00 | | TRBJ2-7*00 | 0.61% |
| Treg WT3 | TGTGCCAGCAGCTACAGAAACGAAAGATTATTTTTC | 1 | TRBV4*00 | | TRBJ1-4*00 | 0.61% |
| Treg WT3 | TGTGCCAGCAGCCTCGGACAGTATGAACAGTACTTC | 1 | TRBV15*00 | TRBD1*00 | TRBJ2-7*00 | 0.61% |
| Treg WT3 | TGTGCCAGCAGCCGACAGGGTGCTGAGCAGTTCTTC | 1 | TRBV26*00 | TRBD1*00 | TRBJ2-1*00 | 0.61% |
| Treg WT3 | TGTGCCAGCAGCCAGGCAAACTCCGACTACACCTTC | 1 | TRBV2*00 | | TRBJ1-2*00 | 0.61% |
| Treg WT3 | TGTGCCAGCAGCCAAGTTTATGCTGAGCAGTTCTTC | 1 | TRBV5*00 | | TRBJ2-1*00 | 0.61% |
| Treg WT3 | TGTGCCAGCAGCAGACAACAAAACACCTTGTACTTT | 1 | TRBV3*00 | | TRBJ2-4*00 | 0.61% |
| Treg WT3 | TGTGCCAGCAACAAAAACACCGGGCAGCTCTACTTT | 1 | TRBV15*00 | | TRBJ2-2*00 | 0.61% |
| Treg WT3 | TGTGCAAGCAGCACTCAGGTCACAGAAGTCTTCTTT | 1 | TRBV16*00 | | TRBJ1-1*00 | 0.61% |
| Treg WT3 | TGTGGTGCTCGGGACATCAACGAAAGATTATTTTTC | 1 | TRBV20*00 | | TRBJ1-4*00 | 0.61% |
| Treg WT3 | TGTGGTGCTAGTCAAAACTATGCTGAGCAGTTCTTC | 1 | TRBV20*00 | | TRBJ2-1*00 | 0.61% |
| Treg WT3 | TGCACCTGCAGTGCAGACAACCAGGCTCCGCTTTTT | 1 | TRBV1*00 | | TRBJ1-5*00 | 0.61% |
| Treg WT3 | TGTGCCAGCTCTCTCGCTGGGGAACAGTACTTC | 1 | TRBV12-2*00 | TRBD2*00 | TRBJ2-7*00 | 0.61% |
| Treg WT3 | TGTGCCAGCGGTGATGAACAGGAACAGTACTTC | 1 | TRBV13-2*00 | | TRBJ2-7*00 | 0.61% |
| Treg WT3 | TGTGCCAGCAGCTTCTCCTATGAACAGTACTTC | 1 | TRBV3*00 | | TRBJ2-7*00 | 0.61% |
| Treg WT3 | TGTGCCAGCAGCTGGGGGGACACCCAGTACTTT | 1 | TRBV4*00 | TRBD1*00 | TRBJ2-5*00 | 0.61% |
| Treg WT3 | TGTGCCAGCAGCGGGACAGGGGCTTTC | 1 | TRBV26*00 | TRBD1*00 | TRBJ2-7*00 | 0.61% |

#### Treg DKO1

| Sample name | CDR3 Sequence | clone Count | best V Hit | best D Hit | best J Hit | Frequency |
|---|---|---|---|---|---|---|
| Treg DKO1 | TGTGCCAGCTCTCTCGGGGGGGGAATGAACACCCAGTACTTT | 69 | TRBV12-1*00 | TRBD2*00 | TRBJ2-5*00 | 25.37% |
| Treg DKO1 | TGTGCCAGCAGTATAGGGGTTGAGAACACAGAAGTCTTCTTT | 47 | TRBV19*00 | | TRBJ1-1*00 | 17.28% |
| Treg DKO1 | TGTGCCAGCAGTTTTAACAACCAGGCTCCGCTTTTT | 12 | TRBV15*00 | | TRBJ1-5*00 | 4.41% |
| Treg DKO1 | TGTGCCAGCTCTCTCGAGGGGGTAGCAAACACCGGGCAGCTCTACTTT | 7 | TRBV12-1*00 | | TRBJ2-2*00 | 2.57% |
| Treg DKO1 | TGTGCCAGCGGTGCAGGGGGAGATGAACAGTACTTC | 4 | TRBV13-2*00 | TRBD1*00 | TRBJ2-7*00 | 1.47% |
| Treg DKO1 | TGTGCCAGCGGTGATGCGGCATACAACCAGGCTCCGCTTTTT | 3 | TRBV13-2*00 | | TRBJ1-5*00 | 1.10% |
| Treg DKO1 | TGTGCCAGCAGCCAAGGGACAAACACAGAAGTCTTCTTT | 3 | TRBV5*00 | | TRBJ1-1*00 | 1.10% |
| Treg DKO1 | TGTGCAAGCAGCTTAAACAGAAACTCCGACTACACCTTC | 3 | TRBV16*00 | | TRBJ1-2*00 | 1.10% |
| Treg DKO1 | TGTGCCAGCAGCGCAAACTCCTATGAACAGTACTTC | 3 | TRBV15*00 | | TRBJ2-7*00 | 1.10% |
| Treg DKO1 | TGTGCCAGCAGCAGGGACAGGGACGTCTTCTTT | 3 | TRBV15*00 | TRBD1*00 | TRBJ1-1*00 | 1.10% |
| Treg DKO1 | TGTGCCAGCAGCCAACAGGGATAAATTCTGGAAATACGCTCTATTTT | 2 | TRBV2*00 | TRBD1*00 | TRBJ1-3*00 | 0.74% |
| Treg DKO1 | TGTGCCAGCTCTCTCGGGGCTGGGGGTGCAGAAACGCTGTATTTT | 2 | TRBV12-1*00 | TRBD2*00 | TRBJ2-3*00 | 0.74% |
| Treg DKO1 | TGTGCCAGCAGCCAACTGGGTCCTAGTGCAGAAACGCTGTATTTT | 2 | TRBV5*00 | TRBD2*00 | TRBJ2-3*00 | 0.74% |
| Treg DKO1 | TGTGGTGCTAGGGACTGGGGACCTAGTCAAAACACCTTGTACTTT | 2 | TRBV20*00 | TRBD2*00 | TRBJ2-4*00 | 0.74% |
| Treg DKO1 | TGCACCTGCAGTCCCGACTGGGGGAGTGCAGAAACGCTGTATTTT | 2 | TRBV1*00 | TRBD2*00 | TRBJ2-3*00 | 0.74% |
| Treg DKO1 | TGTGCCAGCTCTCTCGAGGGGGCTGGAAATACGCTCTATTTT | 2 | TRBV12-1*00 | | TRBJ1-3*00 | 0.74% |
| Treg DKO1 | TGTGCCAGCGGTGGGCAGAACTATGCTGAGCAGTTCTTC | 2 | TRBV13-2*00 | | TRBJ2-1*00 | 0.74% |
| Treg DKO1 | GTGCCAGCAGCCAAGCCGGATCCGGGGCCACAAACTCCGACTACACCTT | 1 | TRBV2*00 | TRBD1*00 | TRBJ1-2*00 | 0.37% |
| Treg DKO1 | TGCACCTGCAGTGCAGGCCCCCGCCAGAGCTCCTATGAACAGTACTTC | 1 | TRBV1*00 | TRBD1*00 | TRBJ2-7*00 | 0.37% |
| Treg DKO1 | TGTGCCTGGAGCCGGGACAGCAAAAAATCTCCTATGAACAGTACTTC | 1 | TRBV31*00 | TRBD1*00 | TRBJ2-7*00 | 0.37% |
| Treg DKO1 | TGTGCTAGCAGTAGAGGGACTGGGGGCCTTGCTGAGCAGTTCTTC | 1 | TRBV17*00 | TRBD2*00 | TRBJ2-1*00 | 0.37% |

| | | | | | | |
|---|---|---|---|---|---|---|
| Treg DKO1 | TGTGCCAGCTCCCGGGACTGGGTCCTCTATGCTGAGCAGTTCTTC | 1 | TRBV12-1*00 | TRBD2*00 | TRBJ2-1*00 | 0.37% |
| Treg DKO1 | TGTGCCAGCGGTGATGCGACTGGGGGTAAAGACACCCAGTACTTT | 1 | TRBV13-2*00 | TRBD2*00 | TRBJ2-5*00 | 0.37% |
| Treg DKO1 | TGTGCCAGCAGCCCCCGGACATTTAACCAAGACACCCAGTACTTT | 1 | TRBV2*00 | TRBD1*00 | TRBJ2-5*00 | 0.37% |
| Treg DKO1 | TGTGCCAGCAGCCACGACTGGGGGGTCCAAGACACCCAGTACTTT | 1 | TRBV5*00 | TRBD2*00 | TRBJ2-5*00 | 0.37% |
| Treg DKO1 | TGTGGTGCTAGGAAATGGGGGACCAACTATGCTGAGCAGTTCTTC | 1 | TRBV20*00 | TRBD2*00 | TRBJ2-1*00 | 0.37% |
| Treg DKO1 | TGTGCCTGGAGTCTAATGGTTAACCAAGACACCCAGTACTTT | 1 | TRBV31*00 | | TRBJ2-5*00 | 0.37% |
| Treg DKO1 | TGTGCCAGCTCTCTCTCCTGGGGGGCCGGGCAGCTCTACTTT | 1 | TRBV13-1*00 | TRBD2*00 | TRBJ2-2*00 | 0.37% |
| Treg DKO1 | TGTGCCAGCTCTCTCGCTGGGGGGGCCCATGAACAGTACTTC | 1 | TRBV12-1*00 | TRBD2*00 | TRBJ2-7*00 | 0.37% |
| Treg DKO1 | TGTGCCAGCTCTCCGGGACAGGCAAACTCCGACTACACCTTC | 1 | TRBV12-2*00 | TRBD1*00 | TRBJ1-2*00 | 0.37% |
| Treg DKO1 | TGTGCCAGCGGTCGGACAGCAAACACCGGGCAGCTCTACTTT | 1 | TRBV13-2*00 | TRBD1*00 | TRBJ2-2*00 | 0.37% |
| Treg DKO1 | TGTGCCAGCGGTGATGGCAGTAACCAAGACACCCAGTACTTT | 1 | TRBV13-2*00 | | TRBJ2-5*00 | 0.37% |
| Treg DKO1 | TGTGCCAGCGGGGACAGGGGTAGTGCAGAAACGCTGTATTTT | 1 | TRBV13-2*00 | TRBD1*00 | TRBJ2-3*00 | 0.37% |
| Treg DKO1 | TGTGCCAGCGACTGGGGGCTTAACCAAGACACCCAGTACTTT | 1 | TRBV15*00 | TRBD2*00 | TRBJ2-5*00 | 0.37% |
| Treg DKO1 | TGTGCCAGCACTCCACACAATTCTGGAAATACGCTCTATTTT | 1 | TRBV26*00 | | TRBJ1-3*00 | 0.37% |
| Treg DKO1 | TGTGCCAGCAGTTTCGGTAGGGGAGGCGACACCCAGTACTTT | 1 | TRBV14*00 | TRBD1*00 | TRBJ2-5*00 | 0.37% |
| Treg DKO1 | TGTGCCAGCAGTTTAGGACAAAACCAAGACACCCAGTACTTT | 1 | TRBV15*00 | | TRBJ2-5*00 | 0.37% |
| Treg DKO1 | TGTGCCAGCAGTTTAAACTGGGGGCAAGACACCCAGTACTTT | 1 | TRBV15*00 | TRBD2*00 | TRBJ2-5*00 | 0.37% |
| Treg DKO1 | TGTGCCAGCAGTCTGGGGGACAGAAACACAGAAGTCTTCTTT | 1 | TRBV26*00 | TRBD1*00 | TRBJ1-1*00 | 0.37% |
| Treg DKO1 | TGTGCCAGCAGTCTGGACAGTGCAAACTCCGACTACACCTTC | 1 | TRBV13-1*00 | TRBD1*00 | TRBJ1-2*00 | 0.37% |
| Treg DKO1 | TGTGCCAGCAGTCCCCAGGGAAGTCAAAACACCTTGTACTTT | 1 | TRBV26*00 | TRBD1*00 | TRBJ2-4*00 | 0.37% |
| Treg DKO1 | TGTGCCAGCAGTCCCAGGGTTTCTGGAAATACGCTCTATTTT | 1 | TRBV26*00 | TRBD1*00 | TRBJ1-3*00 | 0.37% |
| Treg DKO1 | TGTGCCAGCAGTGGACTGGGGGTCTATGCTGAGCAGTTCTTC | 1 | TRBV13-3*00 | TRBD2*00 | TRBJ2-1*00 | 0.37% |
| Treg DKO1 | TGTGCCAGCAGTGATTCCGACAACAACCAGGCTCCGCTTTTT | 1 | TRBV13-1*00 | | TRBJ1-5*00 | 0.37% |
| Treg DKO1 | TGTGCCAGCAGTGATGCGGGGTACAACCAGGCTCCGCTTTTT | 1 | TRBV13-1*00 | | TRBJ1-5*00 | 0.37% |
| Treg DKO1 | TGTGCCAGCAGTGATGGGGGTAACTATGCTGAGCAGTTCTTC | 1 | TRBV13-1*00 | | TRBJ2-1*00 | 0.37% |
| Treg DKO1 | TGTGCCAGCAGTGACTGGGGGAACTATGCTGAGCAGTTCTTC | 1 | TRBV13-1*00 | TRBD2*00 | TRBJ2-1*00 | 0.37% |
| Treg DKO1 | TGTGCCAGCAGCTTAGACTGGGGGGGTGCTGAGCAGTTCTTC | 1 | TRBV3*00 | TRBD2*00 | TRBJ2-1*00 | 0.37% |
| Treg DKO1 | TGTGCCAGCAGCCCCACAGGAGGCCAAGACACCCAGTACTTT | 1 | TRBV4*00 | TRBD1*00 | TRBJ2-5*00 | 0.37% |
| Treg DKO1 | TGTGCCAGCAGCGGGGACAGGGGAAACTCCGACTACACCTTC | 1 | TRBV13-3*00 | TRBD1*00 | TRBJ1-2*00 | 0.37% |
| Treg DKO1 | TGTGCCAGCAGCGGACAGGCAAACACCGGGCAGCTCTACTTT | 1 | TRBV15*00 | TRBD1*00 | TRBJ2-2*00 | 0.37% |
| Treg DKO1 | TGTGCAAGCAGCTTACAGGGGGCCAACGAAAGATTATTTTTC | 1 | TRBV16*00 | TRBD1*00 | TRBJ1-4*00 | 0.37% |
| Treg DKO1 | TGTGGTGCTAGGAGGAGCAGGGGTGCAGAAACGCTGTATTTT | 1 | TRBV20*00 | TRBD1*00 | TRBJ2-3*00 | 0.37% |
| Treg DKO1 | TGCACCTGCAGTGGGACAGGGGGCCACTCCGACTACACCTTC | 1 | TRBV1*00 | TRBD1*00 | TRBJ1-2*00 | 0.37% |
| Treg DKO1 | TGTGCCAGCTCTCTCGTCTGGGGGGGGGAACAGTACTTC | 1 | TRBV12-1*00 | TRBD2*00 | TRBJ2-7*00 | 0.37% |
| Treg DKO1 | TGTGCCAGCAGTTTCGGGACAGCAGAAACGCTGTATTTT | 1 | TRBV15*00 | TRBD1*00 | TRBJ2-3*00 | 0.37% |
| Treg DKO1 | TGTGCCAGCAGTTTCGACGCAAACACAGAAGTCTTCTTT | 1 | TRBV15*00 | | TRBJ1-1*00 | 0.37% |
| Treg DKO1 | TGTGCCAGCAGTCTGTCAGGACTCCCCGACTACACCTTC | 1 | TRBV26*00 | TRBD2*00 | TRBJ1-2*00 | 0.37% |
| Treg DKO1 | TGTGCCAGCAGTGGGCGGGGTGCAGAAACGCTGTATTTT | 1 | TRBV13-1*00 | | TRBJ2-3*00 | 0.37% |
| Treg DKO1 | TGTGCCAGCAGTGGACTGAACCAAGACACCCAGTACTTT | 1 | TRBV13-1*00 | TRBD2*00 | TRBJ2-5*00 | 0.37% |
| Treg DKO1 | TGTGCCAGCAGTGATATCGGGAGAGCTGAGCAGTTCTTC | 1 | TRBV13-1*00 | TRBD1*00 | TRBJ2-1*00 | 0.37% |
| Treg DKO1 | TGTGCCAGCAGTGACTGGGGGGGAGACACCCAGTACTTT | 1 | TRBV13-3*00 | TRBD2*00 | TRBJ2-5*00 | 0.37% |
| Treg DKO1 | TGTGCCAGCAGTGACGGGGCGGCAGAAACGCTGTATTTT | 1 | TRBV13-1*00 | TRBD1*00 | TRBJ2-3*00 | 0.37% |
| Treg DKO1 | TGTGCCAGCAGTGACAATAACCAAGACACCCAGTACTTT | 1 | TRBV15*00 | | TRBJ2-5*00 | 0.37% |
| Treg DKO1 | TGTGCCAGCAGTGAAGCGGACTCCTATGAACAGTACTTC | 1 | TRBV13-1*00 | | TRBJ2-7*00 | 0.37% |
| Treg DKO1 | TGTGCCAGCAGTATTCAGGGCCAAGACACCCAGTACTTT | 1 | TRBV19*00 | TRBD1*00 | TRBJ2-5*00 | 0.37% |
| Treg DKO1 | TGTGCCAGCAGTATGGGGAGCTCCTATGAACAGTACTTC | 1 | TRBV19*00 | | TRBJ2-7*00 | 0.37% |
| Treg DKO1 | TGTGCCAGCAGTATAACACCAAACTCCGACTACACCTTC | 1 | TRBV19*00 | | TRBJ1-2*00 | 0.37% |
| Treg DKO1 | TGTGCCAGCAGTAACAAAGCAAACTCCGACTACACCTTC | 1 | TRBV19*00 | | TRBJ1-2*00 | 0.37% |
| Treg DKO1 | TGTGCCAGCAGCTTAGCTGCAGAAAACACCTTGTACTTT | 1 | TRBV3*00 | | TRBJ2-4*00 | 0.37% |
| Treg DKO1 | TGTGCCAGCAGCTTAGGGGACTCCTATGAACAGTACTTC | 1 | TRBV3*00 | | TRBJ2-7*00 | 0.37% |
| Treg DKO1 | TGTGCCAGCAGCCCCAACGGGGGCACAGAAGTCTTCTTT | 1 | TRBV26*00 | TRBD1*00 | TRBJ1-1*00 | 0.37% |
| Treg DKO1 | TGTGCCAGCAGCCCGGGACACCAAGACACCCAGTACTTT | 1 | TRBV15*00 | TRBD1*00 | TRBJ2-5*00 | 0.37% |
| Treg DKO1 | TGTGCCAGCAGCCCGGACAGCAACCAGGCTCCGCTTTTT | 1 | TRBV5*00 | TRBD1*00 | TRBJ1-5*00 | 0.37% |
| Treg DKO1 | TGTGCCAGCAGCCGGGTTTCTGGAAATACGCTCTATTTT | 1 | TRBV15*00 | TRBD1*00 | TRBJ1-3*00 | 0.37% |
| Treg DKO1 | TGTGCCAGCAGCCAAGTGGGGGTCTATGAACAGTACTTC | 1 | TRBV2*00 | TRBD2*00 | TRBJ2-7*00 | 0.37% |
| Treg DKO1 | TGTGCCAGCAGCCAAGAAGGAAACACAGAAGTCTTCTTT | 1 | TRBV5*00 | | TRBJ1-1*00 | 0.37% |
| Treg DKO1 | TGTGCCAGCAGCAGACAGAACACCGGGCAGCTCTACTTT | 1 | TRBV19*00 | TRBD1*00 | TRBJ2-2*00 | 0.37% |
| Treg DKO1 | TGTGCCAGCAAGGAGGGGGCCTCCTATGAACAGTACTTC | 1 | TRBV13-1*00 | TRBD1*00 | TRBJ2-7*00 | 0.37% |
| Treg DKO1 | TGTGCAAGCAGCTTAGTAGGGGGGGGCTGAGCAGTTCTTC | 1 | TRBV16*00 | TRBD1*00 | TRBJ2-1*00 | 0.37% |
| Treg DKO1 | TGTGGTGCTAGGGACTGGGGTGCAGAAACGCTGTATTTT | 1 | TRBV20*00 | TRBD2*00 | TRBJ2-3*00 | 0.37% |
| Treg DKO1 | TGCACCTGCAGTGCCTTACAAAACTCCGACTACACCTTC | 1 | TRBV1*00 | | TRBJ1-2*00 | 0.37% |
| Treg DKO1 | TGCACCTGCAGTGCCCGGGACCCCCAACCCTTGTACTTT | 1 | TRBV1*00 | TRBD1*00 | TRBJ2-4*00 | 0.37% |
| Treg DKO1 | TGTGCCAGCAGCCCTGACAGGGGGACACCTTGTACTTT | 1 | TRBV4*00 | TRBD1*00 | TRBJ2-4*00 | 0.37% |
| Treg DKO1 | TGTGCCTGGAGTTATAACAACCAGGCTCCGCTTTTTG | 1 | TRBV31*00 | | TRBJ1-5*00 | 0.37% |
| Treg DKO1 | TGTGCCAGTGAAGGGACAGGAAACACCTTGTACTTT | 1 | TRBV14*00 | TRBD1*00 | TRBJ2-4*00 | 0.37% |
| Treg DKO1 | TGTGCCAGCTCACGGGACAGGAGGGAACAGTACTTC | 1 | TRBV12-1*00 | TRBD1*00 | TRBJ2-7*00 | 0.37% |
| Treg DKO1 | TGTGCCAGCGGTTCTCAGGGGAACACCCAGTACTTT | 1 | TRBV13-2*00 | TRBD1*00 | TRBJ2-5*00 | 0.37% |
| Treg DKO1 | TGTGCCAGCGGTGCCAGCCAAGACACCCAGTACTTC | 1 | TRBV13-2*00 | | TRBJ2-5*00 | 0.37% |
| Treg DKO1 | TGTGCCAGCAGTTTAGAACGAAACACCTTGTACTTT | 1 | TRBV15*00 | | TRBJ2-4*00 | 0.37% |
| Treg DKO1 | TGTGCCAGCAGTCCCCCGGAAATACGCTCTATTTT | 1 | TRBV13-3*00 | TRBD2*00 | TRBJ1-3*00 | 0.37% |
| Treg DKO1 | TGTGCCAGCAGTGCCCAAAATTCGCCCCTCTACTTT | 1 | TRBV13-1*00 | | TRBJ1-6*00 | 0.37% |
| Treg DKO1 | TGTGCCAGCAGTGATGCAAGGGAGGCTCCGCTTTTT | 1 | TRBV13-3*00 | | TRBJ1-5*00 | 0.37% |
| Treg DKO1 | TGTGCCAGCAGTATCGACGGAGACACCCAGTACTTT | 1 | TRBV19*00 | | TRBJ2-5*00 | 0.37% |
| Treg DKO1 | TGTGCCAGCAGCTTTGGGGAGGCCACCTTGTACTTT | 1 | TRBV3*00 | TRBD2*00 | TRBJ2-4*00 | 0.37% |

| | | | | | | |
|---|---|---|---|---|---|---|
| Treg DKO1 | TGTGCCAGCAGCTCGGACACCTATGAACAGTACTTC | 1 | TRBV3*00 | TRBD1*00 | TRBJ2-7*00 | 0.37% |
| Treg DKO1 | TGTGCCAGCAGCGGGGACTTAAACACCTTGTACTTT | 1 | TRBV2*00 | TRBD2*00 | TRBJ2-4*00 | 0.37% |
| Treg DKO1 | TGTGCCAGCAGGGGGGCAAACTCCGACTACACCTTC | 1 | TRBV13-1*00 | | TRBJ1-2*00 | 0.37% |
| Treg DKO1 | TGTGCCAGCAGGACTGGGGGAGACACCCAGTACTTT | 1 | TRBV14*00 | TRBD2*00 | TRBJ2-5*00 | 0.37% |
| Treg DKO1 | TGTGCCAGCAGGACAGCAAACTCCGACTACACCTTC | 1 | TRBV14*00 | | TRBJ1-2*00 | 0.37% |
| Treg DKO1 | TGTGCCAGCAGAGTGGGGTATGCTGAGCAGTTCTTC | 1 | TRBV19*00 | TRBD2*00 | TRBJ2-1*00 | 0.37% |
| Treg DKO1 | TGTGCCAGCAGAGGGACATCCTATGAACAGTACTTC | 1 | TRBV13-1*00 | TRBD1*00 | TRBJ2-7*00 | 0.37% |
| Treg DKO1 | TGTGCCAGCAGAGACAACAACGAAAGATTATTTTTC | 1 | TRBV15*00 | | TRBJ1-4*00 | 0.37% |
| Treg DKO1 | TGTGCAAGCAGCTTAGCAGTCTATGAACAGTACTTC | 1 | TRBV16*00 | | TRBJ2-7*00 | 0.37% |
| Treg DKO1 | TGTGCAAGCAGCCGGGATCAAAACACCTTGTACTTT | 1 | TRBV16*00 | TRBD1*00 | TRBJ2-4*00 | 0.37% |
| Treg DKO1 | TGTGGTGCTAGGAGAGGAAACTCCGACTACACCTTC | 1 | TRBV20*00 | | TRBJ1-2*00 | 0.37% |
| Treg DKO1 | TGCACCTGCAGTGCTAACACCGGGCAGCTCTACTTT | 1 | TRBV1*00 | | TRBJ2-2*00 | 0.37% |
| Treg DKO1 | TGTGCCAGCTCTCCCCCCGCTGAGCAGTTCTTC | 1 | TRBV12-1*00 | TRBD1*00 | TRBJ2-1*00 | 0.37% |
| Treg DKO1 | TGTGCCAGCTCCCGGAACCAGGCTCCGCTTTTT | 1 | TRBV12-2*00 | | TRBJ1-5*00 | 0.37% |
| Treg DKO1 | TGTGCCAGCGGTGACAGATATGAACAGTACTTC | 1 | TRBV13-2*00 | | TRBJ2-7*00 | 0.37% |
| Treg DKO1 | TGTGCCAGCAGTTCACAAGACACCCAGTACTTT | 1 | TRBV15*00 | | TRBJ2-5*00 | 0.37% |
| Treg DKO1 | TGTGCCAGCAGTTGGGGAACAGAAGTCTTCTTT | 1 | TRBV15*00 | | TRBJ1-1*00 | 0.37% |
| Treg DKO1 | TGTGCCAGCAGTCTGGAGAATACGCTCTATTTT | 1 | TRBV24*00 | | TRBJ1-3*00 | 0.37% |
| Treg DKO1 | TGTGCCAGCAGTACAAACTCCGACTACACCTTC | 1 | TRBV26*00 | | TRBJ1-2*00 | 0.37% |
| Treg DKO1 | TGTGCCAGCAGCTACAACGAAAGATTATTTTTC | 1 | TRBV3*00 | | TRBJ1-4*00 | 0.37% |
| Treg DKO1 | TGTGCCAGCAGCGCAGACACAGAAGTCTTCTTT | 1 | TRBV3*00 | | TRBJ1-1*00 | 0.37% |
| Treg DKO1 | TGTGCAACTGGGACAACAGAAACGCTGTATTTT | 1 | TRBV16*00 | TRBD1*00 | TRBJ2-3*00 | 0.37% |
| Treg DKO1 | TGTGCAAGCAGCTTCTCTTATGAACAGTACTTC | 1 | TRBV16*00 | | TRBJ2-7*00 | 0.37% |
| Treg DKO1 | TGTGCAAGCAGCTTAGGTGACACCCAGTACTTT | 1 | TRBV16*00 | | TRBJ2-5*00 | 0.37% |
| Treg DKO1 | TGTGCAAGCAGCCCTTATGCTGAGCAGTTCTTC | 1 | TRBV16*00 | | TRBJ2-1*00 | 0.37% |
| Treg DKO1 | TGTGCCAGCAGCCACAGTGAAGTCTTCTTT | 1 | TRBV4*00 | | TRBJ1-1*00 | 0.37% |

#### Treg DKO2

| Sample name | CDR3 Sequence | clone Count | best V Hit | best D Hit | best J Hit | Frequency |
|---|---|---|---|---|---|---|
| Treg DKO2 | TGTGCCAGCAGCCAAGATCTCAGAAACACCTTGTACTTT | 140 | TRBV5*00 | | TRBJ2-4*00 | 35.44% |
| Treg DKO2 | TGTGCCAGCGGTGATGTGGACTGGGGGAGTGCAGAAACGCTGTATTTT | 104 | TRBV13-2*00 | TRBD2*00 | TRBJ2-3*00 | 26.33% |
| Treg DKO2 | TGTGCCAGCGGTGATATGGACAGGGCAAACTCCGACTACACCTTC | 19 | TRBV13-2*00 | TRBD1*00 | TRBJ1-2*00 | 4.81% |
| Treg DKO2 | TGTGCCAGCAGCCCGACAACTGCAAACACAGAAGTCTTCTTT | 14 | TRBV5*00 | | TRBJ1-1*00 | 3.54% |
| Treg DKO2 | TGTAGTTCCCACAAAAACGAAAGATTATTTTTC | 10 | TRBV30*00 | | TRBJ1-4*00 | 2.53% |
| Treg DKO2 | TGTGCCAGCAGTTTCCCAAGCAGTTCTTC | 5 | TRBV15*00 | | TRBJ2-1*00 | 1.27% |
| Treg DKO2 | TGTGCAAGCAGCTTACCTAACCAAGACACCCAGTACTTT | 4 | TRBV16*00 | | TRBJ2-5*00 | 1.01% |
| Treg DKO2 | TGTGCCAGCGGTGAAGGGGACCGTAGTGCAGAAACGCTGTATTTT | 2 | TRBV13-2*00 | TRBD1*00 | TRBJ2-3*00 | 0.51% |
| Treg DKO2 | TGTGCCAGCGGTACCGACAGGGGTGCAGAAACGCTGTATTTT | 2 | TRBV13-2*00 | TRBD1*00 | TRBJ2-3*00 | 0.51% |
| Treg DKO2 | TGTGCCAGCAGTACGAGGGGAAACCAAGACACCCAGTACTTT | 2 | TRBV13-1*00 | TRBD1*00 | TRBJ2-5*00 | 0.51% |
| Treg DKO2 | TGTGCTAGCAGTACGGCAAACACCGGGCAGCTCTACTTT | 2 | TRBV17*00 | | TRBJ2-2*00 | 0.51% |
| Treg DKO2 | TGTGCCAGCTCTCTCGCATATGAACAGTACTTC | 2 | TRBV12-1*00 | | TRBJ2-7*00 | 0.51% |
| Treg DKO2 | GTGCCAGCAGCCAAGACCACTGGGGGGGTGGTCAAAACACCTTGTACTT | 1 | TRBV5*00 | TRBD2*00 | TRBJ2-4*00 | 0.25% |
| Treg DKO2 | TGTGCCAGCGGTGATGTAGACTGGGGGGGTGCAGAAACGCTGTATTTT | 1 | TRBV13-2*00 | TRBD2*00 | TRBJ2-3*00 | 0.25% |
| Treg DKO2 | TGTGCAAGCAGCTTTGGGCTGGGGGTTAACCAAGACACCCAGTACTTT | 1 | TRBV16*00 | TRBD2*00 | TRBJ2-5*00 | 0.25% |
| Treg DKO2 | TGTGCCAGCTCTCTTCGCCGGAGAAACACCGGGCAGCTCTACTTT | 1 | TRBV12-1*00 | | TRBJ2-2*00 | 0.25% |
| Treg DKO2 | TGTGCCAGCGGTGATGCTGGACAGGGTTATGCTGAGCAGTTCTTC | 1 | TRBV13-2*00 | TRBD1*00 | TRBJ2-1*00 | 0.25% |
| Treg DKO2 | TGTGCCAGCGGTGATGCCACAGGGGCCAACGAAAGATTATTTTTC | 1 | TRBV13-2*00 | TRBD1*00 | TRBJ1-4*00 | 0.25% |
| Treg DKO2 | TGTGCCAGCGGTGATATGGACAGGGCAAGCTCCGACTACACCTTC | 1 | TRBV13-2*00 | TRBD1*00 | TRBJ1-2*00 | 0.25% |
| Treg DKO2 | TGTGCCAGCGGTGATATGGACAGGGCAAACTCCGACTACACCGTC | 1 | TRBV13-2*00 | TRBD1*00 | TRBJ1-2*00 | 0.25% |
| Treg DKO2 | TGTGCCAGCAGTATAAGGCTGGGTAGTGCAGAAACGCTGTATTTT | 1 | TRBV19*00 | TRBD2*00 | TRBJ2-3*00 | 0.25% |
| Treg DKO2 | TGTGCCAGCAGCTGGGACTGGGGGAACCAAGACACCCAGTACTTT | 1 | TRBV25*00 | TRBD2*00 | TRBJ2-5*00 | 0.25% |
| Treg DKO2 | TGTGCCAGCAGCCCCTCCCCAACTAGTGCAGAAACGCTGTATTTT | 1 | TRBV2*00 | TRBD2*00 | TRBJ2-3*00 | 0.25% |
| Treg DKO2 | TGTGCCAGCAGCCCCGGAAGAATTTCCAACGAAAGATTATTTTTC | 1 | TRBV5*00 | | TRBJ1-4*00 | 0.25% |
| Treg DKO2 | TGTGCCAGCAGCCACCCGGGACAGGGGAGCGAAAGATTATTTTTC | 1 | TRBV2*00 | TRBD1*00 | TRBJ1-4*00 | 0.25% |
| Treg DKO2 | TGTGCCAGCAGCCAAGATGTGGACACCTATTCGCCCCTCTACTTT | 1 | TRBV5*00 | TRBD1*00 | TRBJ1-6*00 | 0.25% |
| Treg DKO2 | TGTGCAAGCAGCTTACGGACATCAAACACCGGGCAGCTCTACTTT | 1 | TRBV16*00 | TRBD1*00 | TRBJ2-2*00 | 0.25% |
| Treg DKO2 | TGTGCAAGCAGCCTTGGGGGGTCTAGTGCAGAAACGCTGTATTTT | 1 | TRBV16*00 | TRBD2*00 | TRBJ2-3*00 | 0.25% |
| Treg DKO2 | TGTGCTAGCAGTACGGGACAAGAAAACTCCGACTACACCTGC | 1 | TRBV21*00 | TRBD1*00 | TRBJ1-2*00 | 0.25% |
| Treg DKO2 | TGTGCCTGGAGTTCCCAGGGGGCGCGGGACACCCAGTACTTT | 1 | TRBV31*00 | TRBD1*00 | TRBJ2-5*00 | 0.25% |
| Treg DKO2 | TGTGCCTGGAGTCTCAGGACATCTGGAAATACGCTCTATTTT | 1 | TRBV31*00 | TRBD1*00 | TRBJ1-3*00 | 0.25% |
| Treg DKO2 | TGTGCCGTCCGACAGGGTTTAAACTATGCTGAGCAGTTCTTC | 1 | TRBV26*00 | TRBD1*00 | TRBJ2-1*00 | 0.25% |
| Treg DKO2 | TGTGCCAGCTCTCTCGGCAATTCTGGAAATACGCTCTATTTT | 1 | TRBV12-1*00 | | TRBJ1-3*00 | 0.25% |
| Treg DKO2 | TGTGCCAGCCAAAGGAAGGCTAGTGCAGAAACGCTGTATTTT | 1 | TRBV4*00 | | TRBJ2-3*00 | 0.25% |
| Treg DKO2 | TGTGCCAGCGGTGACGCAGGGAGCTCCTATGAACAGTACTTC | 1 | TRBV13-2*00 | | TRBJ2-7*00 | 0.25% |
| Treg DKO2 | TGTGCCAGCGGAAGACAGGGTGGTGCAGAAACGCTGTATTTT | 1 | TRBV13-2*00 | TRBD1*00 | TRBJ2-3*00 | 0.25% |
| Treg DKO2 | TGTGCCAGCAGTGATGGGACAGTCTATGCTGAGCAGTTCTTC | 1 | TRBV13-3*00 | TRBD1*00 | TRBJ2-1*00 | 0.25% |
| Treg DKO2 | TGTGCCAGCAGCTCGGGACAGATCTATGCTGAGCAGTTCTTC | 1 | TRBV25*00 | TRBD1*00 | TRBJ2-1*00 | 0.25% |
| Treg DKO2 | TGTGCCAGCAGCCGCAGGGGTGCAAACACAGAAGTCTTCTTT | 1 | TRBV15*00 | TRBD1*00 | TRBJ1-1*00 | 0.25% |
| Treg DKO2 | TGTGCCAGCAGCCAAGGGGGTTTTGCAGAAACGCTGTATTTT | 1 | TRBV2*00 | | TRBJ2-3*00 | 0.25% |
| Treg DKO2 | TGTGCCAGCAGCCAAGACTGGGGGGGCTATGAACAGTACTTC | 1 | TRBV2*00 | TRBD2*00 | TRBJ2-7*00 | 0.25% |
| Treg DKO2 | TGTGCCAGCAGCCAAGACAGGAACCAAGACACCCAGTACTTT | 1 | TRBV2*00 | | TRBJ2-5*00 | 0.25% |
| Treg DKO2 | TGTGCCAGCAGAACAGGGGAGAGAGCAGAAACGCTGTATTTT | 1 | TRBV13-3*00 | TRBD1*00 | TRBJ2-3*00 | 0.25% |
| Treg DKO2 | TGTGCAAGCAGCTTAGGCGGTAGTGCAGAAACGCTGTATTTT | 1 | TRBV16*00 | | TRBJ2-3*00 | 0.25% |
| Treg DKO2 | TGTGGTGCTAGGGTCGGGCGTAACTATGCTGAGCAGTTCTTC | 1 | TRBV20*00 | TRBD1*00 | TRBJ2-1*00 | 0.25% |

| | | | | | | |
|---|---|---|---|---|---|---|
| Treg DKO2 | TGCACCTGCAGTGCAGATAGGAACACCGGGCAGCTCTACTTT | 1 | TRBV1*00 | | TRBJ2-2*00 | 0.25% |
| Treg DKO2 | TGCACCTGCAGTGGGGGACAGGGGAACTCCGACTACACCTTC | 1 | TRBV1*00 | TRBD1*00 | TRBJ1-2*00 | 0.25% |
| Treg DKO2 | TGTGCTAGCAGTTTTGCAGGGGGTAACACCTTGTACTTT | 1 | TRBV29*00 | TRBD1*00 | TRBJ2-4*00 | 0.25% |
| Treg DKO2 | TGTGCCTGGAGTGACAATTCTGGAAATACGCTCTATTTT | 1 | TRBV31*00 | | TRBJ1-3*00 | 0.25% |
| Treg DKO2 | TGTGCCTGGAGCCCCGGGAGCTCCTATGAACAGTACTTC | 1 | TRBV31*00 | | TRBJ2-7*00 | 0.25% |
| Treg DKO2 | TGTGCCAGCTCTCTACAGGGAGGTTATGAACAGTACTTC | 1 | TRBV12-1*00 | TRBD1*00 | TRBJ2-7*00 | 0.25% |
| Treg DKO2 | TGTGCCAGCTCGGGGCTTAACCAAGACACCCAGTACTTT | 1 | TRBV12-1*00 | TRBD1*00 | TRBJ2-5*00 | 0.25% |
| Treg DKO2 | TGTGCCAGCAGTTTGAGGCTGAAAAACACCTTGTACTTT | 1 | TRBV15*00 | | TRBJ2-4*00 | 0.25% |
| Treg DKO2 | TGTGCCAGCAGTTGGGACAAAGGAAATACGCTCTATTTT | 1 | TRBV13-1*00 | TRBD1*00 | TRBJ1-3*00 | 0.25% |
| Treg DKO2 | TGTGCCAGCAGTCGACAGGGCTCCTATGAACAGTACTTC | 1 | TRBV15*00 | TRBD1*00 | TRBJ2-7*00 | 0.25% |
| Treg DKO2 | TGTGCCAGCAGTCACTGGGGTGCAGAAACGCTGTATTTT | 1 | TRBV14*00 | TRBD2*00 | TRBJ2-3*00 | 0.25% |
| Treg DKO2 | TGTGCCAGCAGTGGGGGTAACCAAGACACCCAGTACTTT | 1 | TRBV13-1*00 | | TRBJ2-5*00 | 0.25% |
| Treg DKO2 | TGTGCCAGCAGTGATGCGGGGAGGGACACCCAGTACTTT | 1 | TRBV13-1*00 | | TRBJ2-5*00 | 0.25% |
| Treg DKO2 | TGTGCCAGCAGTGACAGCAACAACCAGGCTCCGCTTTTT | 1 | TRBV13-1*00 | | TRBJ1-5*00 | 0.25% |
| Treg DKO2 | TGTGCCAGCAGTATACCAGACCAAGACACCCAGTACTTT | 1 | TRBV19*00 | | TRBJ2-5*00 | 0.25% |
| Treg DKO2 | TGTGCCAGCAGTACATCAAACACCGGGCAGCTCTACTTT | 1 | TRBV19*00 | | TRBJ2-2*00 | 0.25% |
| Treg DKO2 | TGTGCCAGCAGCCGGGGACAGCCCTATGAACAGTACTTC | 1 | TRBV15*00 | TRBD1*00 | TRBJ2-7*00 | 0.25% |
| Treg DKO2 | TGTGCCAGCAGCCACAGGGGTGACACAGAAGTCTTCTTT | 1 | TRBV26*00 | TRBD1*00 | TRBJ1-1*00 | 0.25% |
| Treg DKO2 | TGTGCCAGCAGCCAAGACACTGGAAATACGCTCTATTTT | 1 | TRBV2*00 | | TRBJ1-3*00 | 0.25% |
| Treg DKO2 | TGTGCCAGCAGCCAAGACACAAAAGACACCCAGTACTTT | 1 | TRBV5*00 | | TRBJ2-5*00 | 0.25% |
| Treg DKO2 | TGTGCAAGCAGCTTCTGGAGTCAAAACACCTTGTACTTT | 1 | TRBV16*00 | | TRBJ2-4*00 | 0.25% |
| Treg DKO2 | TGTGCAAGCAGCTCGACAGGGGGCTCCGACTACACCTTC | 1 | TRBV16*00 | TRBD1*00 | TRBJ1-2*00 | 0.25% |
| Treg DKO2 | TGTGCAAGCAGCCGACAGAACCAAGACACCCAGTACTTT | 1 | TRBV16*00 | TRBD1*00 | TRBJ2-5*00 | 0.25% |
| Treg DKO2 | TGTGGTGCTAGCCCGGGGGCGGGTGCTGAGCAGTTCTTC | 1 | TRBV20*00 | TRBD1*00 | TRBJ2-1*00 | 0.25% |
| Treg DKO2 | TGCACCTGCAGTGCAGGGGACAGGGTAGAACAGTACTTC | 1 | TRBV1*00 | TRBD1*00 | TRBJ2-7*00 | 0.25% |
| Treg DKO2 | TGCACCTGCAGTGCAGATGCAAACTCCGACTACACCTTC | 1 | TRBV1*00 | | TRBJ1-2*00 | 0.25% |
| Treg DKO2 | TGTGCCAGCAGTATACCGACAACTATGAACAGTACTTC | 1 | TRBV19*00 | | TRBJ2-7*00 | 0.25% |
| Treg DKO2 | TGTGCAAGCAGCTCCCGCAAACACAGAAGTCTTCTTT | 1 | TRBV16*00 | | TRBJ1-1*00 | 0.25% |
| Treg DKO2 | TGTGCTAGCAGTTTAACCCAGTATGAACAGTACTTC | 1 | TRBV29*00 | TRBD2*00 | TRBJ2-7*00 | 0.25% |
| Treg DKO2 | TGTGCCTGGAAGACTGGAGGAGCTGAGCAGTTCTTC | 1 | TRBV31*00 | TRBD2*00 | TRBJ2-1*00 | 0.25% |
| Treg DKO2 | TGTGCCAGCTCTCTGCAGGGCAGGGAACAGTACTTC | 1 | TRBV12-1*00 | TRBD1*00 | TRBJ2-7*00 | 0.25% |
| Treg DKO2 | TGTGCCAGCTCTCCCACCACCGGGCAGCTCTACTTT | 1 | TRBV12-1*00 | | TRBJ2-2*00 | 0.25% |
| Treg DKO2 | TGTGCCAGCTCGCGGACAAAAAACACCTTGTACTTT | 1 | TRBV12-1*00 | TRBD1*00 | TRBJ2-4*00 | 0.25% |
| Treg DKO2 | TGTGCCAGCACCCAGGGGACCGGGCAGCTCTACTTT | 1 | TRBV15*00 | TRBD1*00 | TRBJ2-2*00 | 0.25% |
| Treg DKO2 | TGTGCCAGCAGTTTCGCCAGGGCTGAGCAGTTCTTC | 1 | TRBV15*00 | | TRBJ2-1*00 | 0.25% |
| Treg DKO2 | TGTGCCAGCAGTTCAGGGAACTCCGACTACACCTTC | 1 | TRBV15*00 | TRBD1*00 | TRBJ1-2*00 | 0.25% |
| Treg DKO2 | TGTGCCAGCAGTTACAACAACCAGGCTCCGCTTTTT | 1 | TRBV15*00 | | TRBJ1-5*00 | 0.25% |
| Treg DKO2 | TGTGCCAGCAGTGCAAACACCGGGCAGCTCTACTTT | 1 | TRBV13-1*00 | | TRBJ2-2*00 | 0.25% |
| Treg DKO2 | TGTGCCAGCAGCCCCGCCTATGCTGAGCAGTTCTTC | 1 | TRBV2*00 | | TRBJ2-1*00 | 0.25% |
| Treg DKO2 | TGTGCCAGCAGCCAAGACAGGGACGAACAGTACTTC | 1 | TRBV5*00 | TRBD1*00 | TRBJ2-7*00 | 0.25% |
| Treg DKO2 | TGTGCCAGCAGCCAAAGGAGGGACACCCAGTACTTT | 1 | TRBV2*00 | | TRBJ2-5*00 | 0.25% |
| Treg DKO2 | TGTGCCAGCAGCACAGGAGGTGCTGAGCAGTTCTTC | 1 | TRBV13-3*00 | TRBD1*00 | TRBJ2-1*00 | 0.25% |
| Treg DKO2 | TGTGCCAGCAGAGGCTGGGGGTATGAACAGTACTTC | 1 | TRBV22*00 | TRBD2*00 | TRBJ2-7*00 | 0.25% |
| Treg DKO2 | TGTGCAAGCAGCTTAGAGGGGGAGGAACAGTACTTC | 1 | TRBV16*00 | TRBD1*00 | TRBJ2-7*00 | 0.25% |
| Treg DKO2 | TGTGCAAGCAGCCGACAGGGCGAAAGATTATTTTTC | 1 | TRBV16*00 | TRBD1*00 | TRBJ1-4*00 | 0.25% |
| Treg DKO2 | TGTGCCAGCAGTCCCCGGGCTGAGCAGTTCTTC | 1 | TRBV26*00 | TRBD1*00 | TRBJ2-1*00 | 0.25% |
| Treg DKO2 | TGTGCCAGCAGTCAGGGTGCTGAGCAGTTCTTC | 1 | TRBV15*00 | TRBD1*00 | TRBJ2-1*00 | 0.25% |
| Treg DKO2 | TGTGCCAGCAGTGTCTCCTATGAACAGTACTTC | 1 | TRBV26*00 | | TRBJ2-7*00 | 0.25% |
| Treg DKO2 | TGTGCCAGCAGTGCAAATGCTGAGCAGTTCTTC | 1 | TRBV13-1*00 | | TRBJ2-1*00 | 0.25% |
| Treg DKO2 | TGTGCCAGCAGTAACAGGGGAGAAGTCTTCTTT | 1 | TRBV15*00 | TRBD1*00 | TRBJ1-1*00 | 0.25% |
| Treg DKO2 | TGTGCCAGCAGCTTAGTCACAGAAGTCTTCTTT | 1 | TRBV3*00 | | TRBJ1-1*00 | 0.25% |
| Treg DKO2 | TGTGCAAGCAGCTTAGCGGTTGAACAGTACTTC | 1 | TRBV16*00 | | TRBJ2-7*00 | 0.25% |
| Treg DKO2 | TGTGGTGCTTCTAGTCAAAACACCTTGTACTTT | 1 | TRBV20*00 | | TRBJ2-4*00 | 0.25% |
| Treg DKO2 | TGTGCCAGCAGACACTCCGACTACACCTTC | 1 | TRBV13-3*00 | | TRBJ1-2*00 | 0.25% |
| Treg DKO2 | TGTGCCAGCAGAAACTCCGACTACACCTTC | 1 | TRBV19*00 | | TRBJ1-2*00 | 0.25% |
| Treg DKO2 | TGTGGTGCTAGGGATGAGACGCTGTATTTT | 1 | TRBV20*00 | | TRBJ2-3*00 | 0.25% |

#### Treg DKO3

| Sample name | CDR3 Sequence | clone Count | best V Hit | best D Hit | best J Hit | Frequency |
|---|---|---|---|---|---|---|
| Treg DKO3 | TGTGCCAGCAGCCAAAGACAGGGAGCCAACGAAAGATTATTTTTC | 31 | TRBV5*00 | TRBD1*00 | TRBJ1-4*00 | 17.82% |
| Treg DKO3 | TGTGGTGCTAGGGGTGCAAACACAGAAGTCTTCTTT | 9 | TRBV20*00 | | TRBJ1-1*00 | 5.17% |
| Treg DKO3 | TGTGCCAGCAGTCTCAGGGGTCAAAACACCTTGTACTTT | 8 | TRBV26*00 | TRBD1*00 | TRBJ2-4*00 | 4.60% |
| Treg DKO3 | TGTGCCAGCAGTGGAACAATTAACCAAGACACCCAGTACTTT | 5 | TRBV13-1*00 | | TRBJ2-5*00 | 2.87% |
| Treg DKO3 | TGTGCCAGCAGCTTAGCTGGAAATACGCTCTATTTT | 5 | TRBV3*00 | | TRBJ1-3*00 | 2.87% |
| Treg DKO3 | TGTGCCAGCAGCCAACTGGGAGCAAACACCGGGCAGCTCTACTTT | 4 | TRBV5*00 | TRBD2*00 | TRBJ2-2*00 | 2.30% |
| Treg DKO3 | TGTGCCAGCAGTGACAGGGTCGACTACACCTTC | 4 | TRBV15*00 | TRBD1*00 | TRBJ1-2*00 | 2.30% |
| Treg DKO3 | TGTGCCAGCAGCTCACAGCGGGGTGCAGAAACGCTGTATTTT | 3 | TRBV3*00 | | TRBJ2-3*00 | 1.72% |
| Treg DKO3 | TGTGCCAGCAGCCAAGGACTGGGTAGTGCAGAAACGCTGTATTTT | 2 | TRBV2*00 | TRBD2*00 | TRBJ2-3*00 | 1.15% |
| Treg DKO3 | TGTGCCAGCTCCTCCGGGACAAACACCGGGCAGCTCTACTTT | 2 | TRBV12-2*00 | TRBD1*00 | TRBJ2-2*00 | 1.15% |
| Treg DKO3 | TGTGCAAGCAGCCCCTACTATAGTGCAGAAACGCTGTATTTT | 2 | TRBV16*00 | | TRBJ2-3*00 | 1.15% |
| Treg DKO3 | TGCACCTGCAGTGTTCAGGGGGCGCAGGACACCCAGTACTTT | 2 | TRBV1*00 | TRBD1*00 | TRBJ2-5*00 | 1.15% |
| Treg DKO3 | TGTGCCAGCAGTTTCACTAACTATGCTGAGCAGTTCTTC | 2 | TRBV14*00 | | TRBJ2-1*00 | 1.15% |
| Treg DKO3 | TGTGCCAGCAGTTTGGGACAAAACTCCGACTACACCTTC | 2 | TRBV15*00 | TRBD1*00 | TRBJ1-2*00 | 1.15% |
| Treg DKO3 | TGTGCCTGGAGTAGAGGGGGCGGGGAACAGTACTTC | 2 | TRBV31*00 | TRBD1*00 | TRBJ2-7*00 | 1.15% |

| | | | | | | |
|---|---|---|---|---|---|---|
| Treg DKO3 | TGTGCCAGCAGGACAGTACCTTGGGAACAGTACTTC | 2 | TRBV14*00 | TRBD1*00 | TRBJ2-7*00 | 1.15% |
| Treg DKO3 | TGTGCCAGCGGCGACGAAAGATTATTTTTC | 2 | TRBV13-2*00 | | TRBJ1-4*00 | 1.15% |
| Treg DKO3 | TGTGCCAGCAGTGATGTCGGCAGGGGTAACTATGCTGAGCAGTTCTTC | 1 | TRBV13-3*00 | TRBD1*00 | TRBJ2-1*00 | 0.57% |
| Treg DKO3 | TGTGCTAGCAGTTTGACAGGTTCCTATAATTCGCCCCTCTACTTT | 1 | TRBV29*00 | TRBD1*00 | TRBJ1-6*00 | 0.57% |
| Treg DKO3 | TGTGCCAGCTCTCTTCAGGGGAATTCTGGAAATACGCTCTATTTT | 1 | TRBV12-2*00 | TRBD1*00 | TRBJ1-3*00 | 0.57% |
| Treg DKO3 | TGTGCCAGCGGTGAGGTTTTGGGGGGTGCAGAAACGCTGTATTTT | 1 | TRBV13-2*00 | TRBD2*00 | TRBJ2-3*00 | 0.57% |
| Treg DKO3 | TGTGCCAGCAGTCTGCCGGGAGGGAGTCAAAACACCTTGTACTTT | 1 | TRBV24*00 | TRBD1*00 | TRBJ2-4*00 | 0.57% |
| Treg DKO3 | TGTGCCAGCAGTGAAGGGACACTTTCTGGAAATACGCTCTATTTT | 1 | TRBV13-1*00 | TRBD1*00 | TRBJ1-3*00 | 0.57% |
| Treg DKO3 | TGTGCCAGCAGCTTGATTGGGGGGAACACCGGGCAGCTCTACTTT | 1 | TRBV3*00 | TRBD2*00 | TRBJ2-2*00 | 0.57% |
| Treg DKO3 | TGTGCCAGCAGCTCCCGACAGGGTGGGGACGAAAGATTATTTTTC | 1 | TRBV25*00 | TRBD1*00 | TRBJ1-4*00 | 0.57% |
| Treg DKO3 | TGTGCCAGCAGCTATGTGGGGTCTAGTGCAGAAACGCTGTATTTT | 1 | TRBV4*00 | TRBD2*00 | TRBJ2-3*00 | 0.57% |
| Treg DKO3 | TGTGCCAGCAGCCCATACTGGGGGGCGCCTATGAACAGTACTTC | 1 | TRBV5*00 | TRBD2*00 | TRBJ2-7*00 | 0.57% |
| Treg DKO3 | TGTGCCTGGAGTGGGACTGGGGGGTATGCTGAGCAGTTCTTC | 1 | TRBV31*00 | TRBD2*00 | TRBJ2-1*00 | 0.57% |
| Treg DKO3 | TGTGCCAGCTCTCCCCGGCATTCTGGAAATACGCTCTATTTT | 1 | TRBV12-2*00 | TRBD1*00 | TRBJ1-3*00 | 0.57% |
| Treg DKO3 | TGTGCCAGCTCTCCGCAGGGTTCTGGAAATACGCTCTATTTT | 1 | TRBV12-1*00 | TRBD1*00 | TRBJ1-3*00 | 0.57% |
| Treg DKO3 | TGTGCCAGCTCTCGGGACAGTGCAAACTCCGACTACACCTTC | 1 | TRBV12-1*00 | TRBD1*00 | TRBJ1-2*00 | 0.57% |
| Treg DKO3 | TGTGCCAGCTCCCCTCAAGCAAACACCGGGCAGCTCTACTTT | 1 | TRBV18*00 | | TRBJ2-2*00 | 0.57% |
| Treg DKO3 | TGTGCCAGCGGTGATCCGGGGGGGCGCGATGAACAGTACTTC | 1 | TRBV13-2*00 | TRBD2*00 | TRBJ2-7*00 | 0.57% |
| Treg DKO3 | TGTGCCAGCGGTGATGCGGGGGGGGTTTATGAACAGTACTTC | 1 | TRBV13-2*00 | TRBD2*00 | TRBJ2-7*00 | 0.57% |
| Treg DKO3 | TGTGCCAGCGGTGATGCACAGGACAACGAAAGATTATTTTTC | 1 | TRBV13-2*00 | | TRBJ1-4*00 | 0.57% |
| Treg DKO3 | TGTGCCAGCGGTGATGAACAGGGAATATGGGAACAGTACTTC | 1 | TRBV13-2*00 | TRBD1*00 | TRBJ2-7*00 | 0.57% |
| Treg DKO3 | TGTGCCAGCAGTTTAGGGACAGCAAACTCCGACTACACCTTC | 1 | TRBV15*00 | TRBD1*00 | TRBJ1-2*00 | 0.57% |
| Treg DKO3 | TGTGCCAGCAGTTCGACAGGGGGCAACTCCGACTACACCTTC | 1 | TRBV14*00 | TRBD1*00 | TRBJ1-2*00 | 0.57% |
| Treg DKO3 | TGTGCCAGCAGTCTTCTAATTAACCAAGACACCCAGTACTTT | 1 | TRBV26*00 | | TRBJ2-5*00 | 0.57% |
| Treg DKO3 | TGTGCCAGCAGTCTGCCGGGACAGGGTGCTGAGCAGTTCTTC | 1 | TRBV26*00 | TRBD1*00 | TRBJ2-1*00 | 0.57% |
| Treg DKO3 | TGTGCCAGCAGTGTGGGACGGTACTATGCTGAGCAGTTCTTC | 1 | TRBV13-1*00 | TRBD1*00 | TRBJ2-1*00 | 0.57% |
| Treg DKO3 | TGTGCCAGCAGTGATGCTAAAAACACCGGGCAGCTCTACTTT | 1 | TRBV13-1*00 | | TRBJ2-2*00 | 0.57% |
| Treg DKO3 | TGTGCCAGCAGCTTAGAGCGGGGCTCCTATGAACAGTACTTC | 1 | TRBV3*00 | | TRBJ2-7*00 | 0.57% |
| Treg DKO3 | TGTGCCAGCAGCTATGGGGACAGGGGCACAGAAGTCTTCTTT | 1 | TRBV3*00 | TRBD1*00 | TRBJ1-1*00 | 0.57% |
| Treg DKO3 | TGTGCCAGCAGCCCCCGGACCAGTGCAGAAACGCTGTATTTT | 1 | TRBV5*00 | TRBD1*00 | TRBJ2-3*00 | 0.57% |
| Treg DKO3 | TGTGCCAGCAGCCCGAGGGGGGTCTCCTATGAACAGTACTTC | 1 | TRBV2*00 | TRBD1*00 | TRBJ2-7*00 | 0.57% |
| Treg DKO3 | TGTGCCAGCAGCCAAACCACTTCTGGAAATACGCTCTATTTT | 1 | TRBV5*00 | | TRBJ1-3*00 | 0.57% |
| Treg DKO3 | TGTGCCAGCAGACCCGGGACAGGGAGGGAAACGCTGTATTTT | 1 | TRBV13-1*00 | TRBD1*00 | TRBJ2-3*00 | 0.57% |
| Treg DKO3 | TGTGCCAGCAGAGGACAAAATAACTATGCTGAGCAGTTCTTC | 1 | TRBV13-1*00 | TRBD1*00 | TRBJ2-1*00 | 0.57% |
| Treg DKO3 | TGTGCAAGCAGCTTAGGTCAGGGAGGCACAGAAGTCTTCTTT | 1 | TRBV16*00 | TRBD1*00 | TRBJ1-1*00 | 0.57% |
| Treg DKO3 | TGTGCAAGCAGCTTAGGGGGGGTCAACGAAAGATTATTTTTC | 1 | TRBV16*00 | TRBD2*00 | TRBJ1-4*00 | 0.57% |
| Treg DKO3 | TGTGGTGCTAGGGACAATTCCTATAATTCGCCCCTCTACTTT | 1 | TRBV20*00 | | TRBJ1-6*00 | 0.57% |
| Treg DKO3 | TGCACCTGCAGTGCAGGGGGCAGGAACGAAAGATTATTTTTC | 1 | TRBV1*00 | TRBD1*00 | TRBJ1-4*00 | 0.57% |
| Treg DKO3 | TGCACCTGCAGTGCAGGGACATCAAACTCCGACTACACCTTC | 1 | TRBV1*00 | TRBD1*00 | TRBJ1-2*00 | 0.57% |
| Treg DKO3 | TGTGCTAGCAGTAGAGCGAGCTCCTATGAACAGTACTTC | 1 | TRBV17*00 | | TRBJ2-7*00 | 0.57% |
| Treg DKO3 | TGTGCTAGCAGCTCTGGGGGGGAGGACACCCAGTACTTT | 1 | TRBV21*00 | TRBD2*00 | TRBJ2-5*00 | 0.57% |
| Treg DKO3 | TGTGCCAGCTCTCTCGACTACAACCAGGCTCCGCTTTTT | 1 | TRBV12-1*00 | | TRBJ1-5*00 | 0.57% |
| Treg DKO3 | TGTGCCAGCTCTCCTGGACAAGCAGAAACGCTGTATTTT | 1 | TRBV12-1*00 | TRBD1*00 | TRBJ2-3*00 | 0.57% |
| Treg DKO3 | TGTGCCAGCTCCCCCGACTGGGGCTATGAACAGTACTTC | 1 | TRBV12-1*00 | TRBD2*00 | TRBJ2-7*00 | 0.57% |
| Treg DKO3 | TGTGCCAGCCGACACCCTAGTGCAGAAACGCTGTATTTT | 1 | TRBV19*00 | | TRBJ2-3*00 | 0.57% |
| Treg DKO3 | TGTGCCAGCGGCCGGGACAATGCAGAAACGCTGTATTTT | 1 | TRBV13-2*00 | TRBD1*00 | TRBJ2-3*00 | 0.57% |
| Treg DKO3 | TGTGCCAGCGGAGGACAGCCAAACACAGAAGTCTTCTTT | 1 | TRBV13-2*00 | TRBD1*00 | TRBJ1-1*00 | 0.57% |
| Treg DKO3 | TGTGCCAGCAGTTTAGGAGGGGGCGCTGAGCAGTTCTTC | 1 | TRBV15*00 | TRBD1*00 | TRBJ2-1*00 | 0.57% |
| Treg DKO3 | TGTGCCAGCAGTTCCGGGACAGGGGTCACCTTGTACTTT | 1 | TRBV3*00 | TRBD1*00 | TRBJ2-4*00 | 0.57% |
| Treg DKO3 | TGTGCCAGCAGTCTCGGACGGAACACAGAAGTCTTCTTT | 1 | TRBV26*00 | | TRBJ1-1*00 | 0.57% |
| Treg DKO3 | TGTGCCAGCAGTCGGGACAGCTCCTATGAACAGTACTTC | 1 | TRBV15*00 | TRBD1*00 | TRBJ2-7*00 | 0.57% |
| Treg DKO3 | TGTGCCAGCAGTCAGGGCAACACCGGGCAGCTCTACTTT | 1 | TRBV26*00 | | TRBJ2-2*00 | 0.57% |
| Treg DKO3 | TGTGCCAGCAGTGGAACAGGGATGACAGAAGTCTTCTTT | 1 | TRBV13-3*00 | TRBD1*00 | TRBJ1-1*00 | 0.57% |
| Treg DKO3 | TGTGCCAGCAGTGATGCAGGCCAAAACACCTTGTACTTT | 1 | TRBV13-1*00 | | TRBJ2-4*00 | 0.57% |
| Treg DKO3 | TGTGCCAGCAGTGATGCAGGGTACTCCGACTACACCTTC | 1 | TRBV13-3*00 | | TRBJ1-2*00 | 0.57% |
| Treg DKO3 | TGTGCCAGCAGTGATGCAGAGGGGAACACCTTGTACTTT | 1 | TRBV13-1*00 | TRBD1*00 | TRBJ2-4*00 | 0.57% |
| Treg DKO3 | TGTGCCAGCAGCTCTCACAGGGGGCGGGAACAGTACTTC | 1 | TRBV25*00 | TRBD1*00 | TRBJ2-7*00 | 0.57% |
| Treg DKO3 | TGTGCCAGCAGCCCCTCTAGTCAAAACACCTTGTACTTT | 1 | TRBV2*00 | | TRBJ2-4*00 | 0.57% |
| Treg DKO3 | TGTGCCAGCAGCCAAGATCATGGCTCCGACTACACCTTC | 1 | TRBV2*00 | | TRBJ1-2*00 | 0.57% |
| Treg DKO3 | TGTGCCAGCAGACAATATTCTGGAAATACGCTCTATTTT | 1 | TRBV3*00 | | TRBJ1-3*00 | 0.57% |
| Treg DKO3 | TGTGCAAGCAGCTTACGACAAAACTCCGACTACACCTTC | 1 | TRBV16*00 | | TRBJ1-2*00 | 0.57% |
| Treg DKO3 | TGTGCAAGCAGCTGGACAGGGCAAGACACCCAGTACTTC | 1 | TRBV16*00 | TRBD1*00 | TRBJ2-5*00 | 0.57% |
| Treg DKO3 | TGTGCAAGCAGCCTTTCCTATAATTCGCCCCTCTACTTT | 1 | TRBV16*00 | | TRBJ1-6*00 | 0.57% |
| Treg DKO3 | TGCACCTGCAGTGCAGACACAGGGGGAGAAGTCTTCTTT | 1 | TRBV1*00 | TRBD1*00 | TRBJ1-1*00 | 0.57% |
| Treg DKO3 | TGTGCCAGCAGTTTATGGGAACTCCGACTACACCTTC | 1 | TRBV15*00 | | TRBJ1-2*00 | 0.57% |
| Treg DKO3 | TGTGCTAGCAGTTTCTCCTGGATTGAACAGTACTTC | 1 | TRBV29*00 | | TRBJ2-7*00 | 0.57% |
| Treg DKO3 | TGTGCTAGCAGTAGAGACAGTGCTGAGCAGTTCTTC | 1 | TRBV17*00 | | TRBJ2-1*00 | 0.57% |
| Treg DKO3 | TGTGCTAGCAGACTGGGGCAAAACACCTTGTACTTT | 1 | TRBV17*00 | TRBD2*00 | TRBJ2-4*00 | 0.57% |
| Treg DKO3 | TGTGCCTGGAGCCGGACAGGGGATGAACAGTACTTC | 1 | TRBV31*00 | TRBD1*00 | TRBJ2-7*00 | 0.57% |
| Treg DKO3 | TGTGCCAGTGTGGGAAACACCGGGCAGCTCTACTTT | 1 | TRBV12-2*00 | | TRBJ2-2*00 | 0.57% |
| Treg DKO3 | TGTGCCAGCTCTCTCGAACAGTATGAACAGTACTTC | 1 | TRBV12-1*00 | | TRBJ2-7*00 | 0.57% |
| Treg DKO3 | TGTGCCAGCGCACAGGTCTATGCTGAGCAGTTCTTC | 1 | TRBV13-2*00 | TRBD1*00 | TRBJ2-1*00 | 0.57% |
| Treg DKO3 | TGTGCCAGCGGGACTGGAGCAGAAACGCTGTATTTT | 1 | TRBV13-2*00 | TRBD2*00 | TRBJ2-3*00 | 0.57% |
| Treg DKO3 | TGTGCCAGCAGTTTCGGCGCCACAGAAGTCTTCTTT | 1 | TRBV15*00 | TRBD1*00 | TRBJ1-1*00 | 0.57% |

| | | | | | | |
|---|---|---|---|---|---|---|
| Treg DKO3 | TGTGCCAGCAGTGACAGAAACTCCGACTACACCTTC | 1 | TRBV13-1*00 | | TRBJ1-2*00 | 0.57% |
| Treg DKO3 | TGTGCCAGCAGCTTAGACACCGGGCAGCTCTACTTT | 1 | TRBV3*00 | | TRBJ2-2*00 | 0.57% |
| Treg DKO3 | TGTGCCAGCAGGACAGGGAACACAGAAGTCTTCTTT | 1 | TRBV15*00 | TRBD1*00 | TRBJ1-1*00 | 0.57% |
| Treg DKO3 | TGTGCAAGCAGCTTAGACAAATATGAACAGTACTTC | 1 | TRBV16*00 | | TRBJ2-7*00 | 0.57% |
| Treg DKO3 | TGTGGTGCTAGGGCTGGGACCCAGGCTCCGCTTTTT | 1 | TRBV20*00 | TRBD1*00 | TRBJ1-5*00 | 0.57% |
| Treg DKO3 | TGCACCTGCAGTGCACCCCCCTATGAACAGTACTTC | 1 | TRBV1*00 | | TRBJ2-7*00 | 0.57% |
| Treg DKO3 | TGTGCCAGCGGTGGCCTAGGGCAGCTCTACTTT | 1 | TRBV13-2*00 | | TRBJ2-2*00 | 0.57% |
| Treg DKO3 | TGTGCCAGCGGGAGCTCCTATGAACAGTACTTC | 1 | TRBV13-2*00 | | TRBJ2-7*00 | 0.57% |
| Treg DKO3 | TGTGCCAGCAGTGCAGCCTATGAACAGTACTTC | 1 | TRBV13-3*00 | | TRBJ2-7*00 | 0.57% |
| Treg DKO3 | TGTGCCAGCAGTATGGAAGACACCCAGTACTTT | 1 | TRBV19*00 | | TRBJ2-5*00 | 0.57% |
| Treg DKO3 | TGTGCCAGCAAGAACTATGCTGAGCAGTTCTTC | 1 | TRBV15*00 | | TRBJ2-1*00 | 0.57% |
| Treg DKO3 | TGTGCAAGCAGCTTGTTGGACACCCAGTACTTT | 1 | TRBV16*00 | | TRBJ2-5*00 | 0.57% |
| Treg DKO3 | TGTGGTGCTAGGGATCTCGAAAGATTATTTTTC | 1 | TRBV20*00 | | TRBJ1-4*00 | 0.57% |
| Treg DKO3 | TGTGCCAGCTCTCTAGGGGAACAGTACTTC | 1 | TRBV12-2*00 | | TRBJ2-7*00 | 0.57% |
| Treg DKO3 | TGTGCCAGCAGTCTCTATGAACAGTACTTC | 1 | TRBV26*00 | | TRBJ2-7*00 | 0.57% |

#### CD4+YFP- WT1

| Sample name | CDR3 Sequence | clone Count | best V Hit | best D Hit | best J Hit | Frequency |
|---|---|---|---|---|---|---|
| CD4+YFP- WT1 | TGTGCCAGCAGTGACCCCGGGACGGCCAACGAAAGATTATTTTTC | 2 | TRBV13-1*00 | TRBD1*00 | TRBJ1-4*00 | 0.96% |
| CD4+YFP- WT1 | TGTGCCAGCTCTCTCGGGAATTCTGGAAATACGCTCTATTTT | 2 | TRBV12-1*00 | | TRBJ1-3*00 | 0.96% |
| CD4+YFP- WT1 | TGTGCCAGCAGCCTCACAGGGTGGGGACACCCAGTACTTT | 2 | TRBV4*00 | TRBD1*00 | TRBJ2-5*00 | 0.96% |
| CD4+YFP- WT1 | TGTGCCAGCAGCCAAGGGGGTGGGGACACCCAGTACTTT | 2 | TRBV5*00 | | TRBJ2-5*00 | 0.96% |
| CD4+YFP- WT1 | TGTGCAAGCAGCTTACCTGGAGTGTATGAACAGTACTTC | 2 | TRBV16*00 | | TRBJ2-7*00 | 0.96% |
| CD4+YFP- WT1 | TGTGCCAGCAGTTACAACTCCTATGAACAGTACTTC | 2 | TRBV19*00 | | TRBJ2-7*00 | 0.96% |
| CD4+YFP- WT1 | TGTGCCAGCAGGGGGAGTCAAAACACCTTGTACTTT | 2 | TRBV26*00 | | TRBJ2-4*00 | 0.96% |
| CD4+YFP- WT1 | TGTGCCAGCGGTGACTATGAACAGTACTTC | 2 | TRBV13-2*00 | | TRBJ2-7*00 | 0.96% |
| CD4+YFP- WT1 | TGTGCCAGCAGCCAAGACAGAGGGGTCTTC | 2 | TRBV2*00 | TRBD1*00 | TRBJ2-7*00 | 0.96% |
| CD4+YFP- WT1 | TGTGCTAGCAGGACTGGGGGGGCGAGTTATAACTATGCTGAGCAGTTCTTC | 1 | TRBV17*00 | TRBD2*00 | TRBJ2-1*00 | 0.48% |
| CD4+YFP- WT1 | TGTGCCAGCAGTATAGCTCGCAGTAATTCTGGAAATACGCTCTATTTT | 1 | TRBV19*00 | | TRBJ1-3*00 | 0.48% |
| CD4+YFP- WT1 | TGTGCCAGCAGCTTATCTGGGGGGGCTAGTCAAAACACCTTGTACTTT | 1 | TRBV3*00 | TRBD2*00 | TRBJ2-4*00 | 0.48% |
| CD4+YFP- WT1 | TGTGCCAGCAGCCAAGGACTGGGGGTAACTATGCTGAGCAGTTCTTC | 1 | TRBV2*00 | TRBD2*00 | TRBJ2-1*00 | 0.48% |
| CD4+YFP- WT1 | TGTGCTAGCAGCCGACAGGGGCTTTCTGGAAATACGCTCTATTTT | 1 | TRBV4*00 | TRBD1*00 | TRBJ1-3*00 | 0.48% |
| CD4+YFP- WT1 | TGTGCCTGGAGTCTAAGCACATCTAGTGCAGAAACGCTGTATTTT | 1 | TRBV31*00 | | TRBJ2-3*00 | 0.48% |
| CD4+YFP- WT1 | TGTGCCAGCAGTTTAGAGGCTGGGGTCTATGCTGAGCAGTTCTTC | 1 | TRBV15*00 | TRBD2*00 | TRBJ2-1*00 | 0.48% |
| CD4+YFP- WT1 | TGTGCCAGCAGTCTGTTGGACAATAACTATGCTGAGCAGTTCTTC | 1 | TRBV24*00 | TRBD1*00 | TRBJ2-1*00 | 0.48% |
| CD4+YFP- WT1 | TGTGCCAGCAGTCCCCGAGGGACAACAAACTCCGACTACACCTTC | 1 | TRBV26*00 | TRBD1*00 | TRBJ1-2*00 | 0.48% |
| CD4+YFP- WT1 | TGTGCCAGCAGTCGACGGGCCTCTTCTGGAAATACGCTCTATTTT | 1 | TRBV26*00 | | TRBJ1-3*00 | 0.48% |
| CD4+YFP- WT1 | TGTGCCAGCAGTGGTGACAGTTTTAACCAAGACACCCAGTACTTT | 1 | TRBV13-1*00 | TRBD1*00 | TRBJ2-5*00 | 0.48% |
| CD4+YFP- WT1 | TGTGCCAGCAGTGGGGTCCCCAATTCTGGAAATACGCTCTATTTT | 1 | TRBV13-1*00 | TRBD1*00 | TRBJ1-3*00 | 0.48% |
| CD4+YFP- WT1 | TGTGCCAGCAGTGATGCACAGGGGCGCTCCTATGAACAGTACTTC | 1 | TRBV13-1*00 | TRBD1*00 | TRBJ2-7*00 | 0.48% |
| CD4+YFP- WT1 | TGTGCCAGCAGTATAGGCTGGGGGAGCTCCTATGAACAGTACTTC | 1 | TRBV19*00 | TRBD2*00 | TRBJ2-7*00 | 0.48% |
| CD4+YFP- WT1 | TGTGCCAGCAGCTTAGCCGGGAATAACTATGCTGAGCAGTTCTTC | 1 | TRBV3*00 | TRBD1*00 | TRBJ2-1*00 | 0.48% |
| CD4+YFP- WT1 | TGTGCCAGCAGCCCGACTGGGGGTAGTGCAGAAACGCTGTATTTT | 1 | TRBV3*00 | TRBD2*00 | TRBJ2-3*00 | 0.48% |
| CD4+YFP- WT1 | TGTGCCAGCAGCCAAGTCTGGGGGGACCAAGACACCCAGTACTTT | 1 | TRBV5*00 | TRBD2*00 | TRBJ2-5*00 | 0.48% |
| CD4+YFP- WT1 | TGTGCCAGCAGCCAAGGGGGGGGGCGCTTATGCTGAGCAGTTCTTC | 1 | TRBV2*00 | TRBD2*00 | TRBJ2-1*00 | 0.48% |
| CD4+YFP- WT1 | TGTGCCAGCAGCCAAGGGACATCTAACAACCAGGCTCCGCTTTTT | 1 | TRBV2*00 | TRBD1*00 | TRBJ1-5*00 | 0.48% |
| CD4+YFP- WT1 | TGTGCCAGCAGCCAAGATCTGGGCAGCTCCTATGAACAGTACTTC | 1 | TRBV2*00 | | TRBJ2-7*00 | 0.48% |
| CD4+YFP- WT1 | TGTGCCAGCAGCCAAACTGGGCATAACTATGCTGAGCAGTTCTTC | 1 | TRBV5*00 | TRBD2*00 | TRBJ2-1*00 | 0.48% |
| CD4+YFP- WT1 | TGTGCAAGCAGCTTAGCAGGGGGTAGTGCAGAAACGCTGTATTTT | 1 | TRBV16*00 | TRBD1*00 | TRBJ2-3*00 | 0.48% |
| CD4+YFP- WT1 | TGTGCAAGCAGCCCGACAGGGGCTAGTCAAAACACCTTGTACTTT | 1 | TRBV16*00 | TRBD1*00 | TRBJ2-4*00 | 0.48% |
| CD4+YFP- WT1 | TGCACCTGCAGTGCCCCCGGGGGCGGGTATGCTGAGCAGTTCTTC | 1 | TRBV1*00 | TRBD1*00 | TRBJ2-1*00 | 0.48% |
| CD4+YFP- WT1 | TGCACCTGCAGTGCGCAGGGAGATTCTGGAAATACGCTCTATTTT | 1 | TRBV1*00 | TRBD1*00 | TRBJ1-3*00 | 0.48% |
| CD4+YFP- WT1 | TGCACCTGCAGTGCGGGGACAGCAAACACCGGGCAGCTCTACTTT | 1 | TRBV1*00 | TRBD1*00 | TRBJ2-2*00 | 0.48% |
| CD4+YFP- WT1 | TGCACCTGCAGTGCAGACAGGGGGCCAAACTCCGACTACACCTTC | 1 | TRBV1*00 | TRBD1*00 | TRBJ1-2*00 | 0.48% |
| CD4+YFP- WT1 | TGTGCTAGCAGTTTATCGGGACAGCAAGACACCCAGTACTTT | 1 | TRBV29*00 | TRBD1*00 | TRBJ2-5*00 | 0.48% |
| CD4+YFP- WT1 | TGTGCTAGCAGTCTGGACAATTCTGGAAATACGCTCTATTTT | 1 | TRBV29*00 | | TRBJ1-3*00 | 0.48% |
| CD4+YFP- WT1 | TGTGCCTGGAGTCTAACAGGGGGCTATGCTGAGCAGTTCTTC | 1 | TRBV31*00 | TRBD1*00 | TRBJ2-1*00 | 0.48% |
| CD4+YFP- WT1 | TGTGCCAGCGGTGATGAGGGGGGCCGAGCTGAGCAGTTCTTC | 1 | TRBV13-2*00 | TRBD2*00 | TRBJ2-1*00 | 0.48% |
| CD4+YFP- WT1 | TGTGCCAGCGGGGGACAAAATTCTGGAAATACGCTCTATTTT | 1 | TRBV26*00 | TRBD1*00 | TRBJ1-3*00 | 0.48% |
| CD4+YFP- WT1 | TGTGCCAGCACCCGGGACAGGAACTATGCTGAGCAGTTCTTC | 1 | TRBV13-3*00 | TRBD1*00 | TRBJ2-1*00 | 0.48% |
| CD4+YFP- WT1 | TGTGCCAGCAGTTTCGGGGGTAACCAAGACACCCAGTACTTT | 1 | TRBV15*00 | TRBD1*00 | TRBJ2-5*00 | 0.48% |
| CD4+YFP- WT1 | TGTGCCAGCAGTTTCAGGGGAACCACCGGGCAGCTCTACTTT | 1 | TRBV14*00 | TRBD1*00 | TRBJ2-2*00 | 0.48% |
| CD4+YFP- WT1 | TGTGCCAGCAGTTTAGGGGACAGGGATTATGAACAGTACTTC | 1 | TRBV15*00 | TRBD1*00 | TRBJ2-7*00 | 0.48% |
| CD4+YFP- WT1 | TGTGCCAGCAGTTTAGGAGGGGGCCAAGACACCCAGTACTTT | 1 | TRBV15*00 | TRBD1*00 | TRBJ2-5*00 | 0.48% |
| CD4+YFP- WT1 | TGTGCCAGCAGTCGGCCGGGACAGGGGGGAGAACAGTACTTC | 1 | TRBV26*00 | TRBD1*00 | TRBJ2-7*00 | 0.48% |
| CD4+YFP- WT1 | TGTGCCAGCAGTCGGGGGATAGTGCAGAAACGCTGTATTTT | 1 | TRBV13-3*00 | TRBD2*00 | TRBJ2-3*00 | 0.48% |
| CD4+YFP- WT1 | TGTGCCAGCAGTGGGACCGCAAACACCGGGCAGCTCTACTTT | 1 | TRBV13-1*00 | | TRBJ2-2*00 | 0.48% |
| CD4+YFP- WT1 | TGTGCCAGCAGTGATCTGGGAAACACCGGGCAGCTCTACTTT | 1 | TRBV13-1*00 | TRBD2*00 | TRBJ2-2*00 | 0.48% |
| CD4+YFP- WT1 | TGTGCCAGCAGTGATGTGTATTCTGGAAATACGCTCTATTTT | 1 | TRBV13-1*00 | | TRBJ1-3*00 | 0.48% |
| CD4+YFP- WT1 | TGTGCCAGCAGTGATGCGGGACAGCCTTATGAACAGTACTTC | 1 | TRBV13-1*00 | TRBD1*00 | TRBJ2-7*00 | 0.48% |
| CD4+YFP- WT1 | TGTGCCAGCAGTGATGAATCTTCTGGAAATACGCTCTATTTT | 1 | TRBV13-3*00 | | TRBJ1-3*00 | 0.48% |
| CD4+YFP- WT1 | TGTGCCAGCAGTGACATAGCAAACACCGGGCAGCTCTACTTT | 1 | TRBV13-1*00 | | TRBJ2-2*00 | 0.48% |
| CD4+YFP- WT1 | TGTGCCAGCAGTAGACAGGGGGCAGAAGCTGAGCAGTTCTTC | 1 | TRBV19*00 | TRBD1*00 | TRBJ2-1*00 | 0.48% |
| CD4+YFP- WT1 | TGTGCCAGCAGCTTTCACTTTAACTATGCTGAGCAGTTCTTC | 1 | TRBV3*00 | | TRBJ2-1*00 | 0.48% |
| CD4+YFP- WT1 | TGTGCCAGCAGCTTCTGGGGGGGTCAAGACACCCAGTACTTT | 1 | TRBV3*00 | TRBD2*00 | TRBJ2-5*00 | 0.48% |
| CD4+YFP- WT1 | TGTGCCAGCAGCTTCACCATTAACCAAGACACCCAGTACTTT | 1 | TRBV4*00 | | TRBJ2-5*00 | 0.48% |
| CD4+YFP- WT1 | TGTGCCAGCAGCTTGGCCAATGCAAACTCCGACTACACCTTC | 1 | TRBV3*00 | | TRBJ1-2*00 | 0.48% |
| CD4+YFP- WT1 | TGTGCCAGCAGCTTAGTACAGGCAAACTCCGACTACACCTTC | 1 | TRBV3*00 | | TRBJ1-2*00 | 0.48% |
| CD4+YFP- WT1 | TGTGCCAGCAGCTTAGCTGGGGGGGTAGGTGAACAGTACTTC | 1 | TRBV3*00 | TRBD2*00 | TRBJ2-7*00 | 0.48% |
| CD4+YFP- WT1 | TGTGCCAGCAGCTTAACAGATAACAACCAGGCTCCGCTTTTT | 1 | TRBV3*00 | | TRBJ1-5*00 | 0.48% |
| CD4+YFP- WT1 | TGTGCCAGCAGCTATCGGGACTGGGGGGGCGCCTTGTACTTT | 1 | TRBV4*00 | TRBD2*00 | TRBJ2-4*00 | 0.48% |
| CD4+YFP- WT1 | TGTGCCAGCAGCTAACGGGGGCGAGGGGCTGAGCAGTTCTTC | 1 | TRBV4*00 | TRBD1*00 | TRBJ2-1*00 | 0.48% |
| CD4+YFP- WT1 | TGTGCCAGCAGCCCGACTGATAACTATGCTGAGCAGTTCTTC | 1 | TRBV2*00 | TRBD2*00 | TRBJ2-1*00 | 0.48% |

| | | | | | | |
|---|---|---|---|---|---|---|
| CD4+YFP- WT1 | TGTGCCAGCAGCCGCCTCGGGACGTCGGCTGAGCAGTTCTTC | 1 | TRBV5*00 | TRBD1*00 | TRBJ2-1*00 | 0.48% |
| CD4+YFP- WT1 | TGTGCCAGCAGCCGCCAGGGGGAAAGGGAAACGCTGTATTTT | 1 | TRBV2*00 | TRBD1*00 | TRBJ2-3*00 | 0.48% |
| CD4+YFP- WT1 | TGTGCCAGCAGCCAAGTGGGGAACACCGGGCAGCTCTACTTT | 1 | TRBV5*00 | TRBD2*00 | TRBJ2-2*00 | 0.48% |
| CD4+YFP- WT1 | TGTGCCAGCAGCCAAGCAGGGAACACCGGGCAGCTCTACTTT | 1 | TRBV2*00 | TRBD1*00 | TRBJ2-2*00 | 0.48% |
| CD4+YFP- WT1 | TGTGCCAGCAGCCAAGGGACACCAAACTCCGACTACACCTTC | 1 | TRBV2*00 | TRBD1*00 | TRBJ1-2*00 | 0.48% |
| CD4+YFP- WT1 | TGTGCCAGCAGCCAAGGGACAAGCAACTCCGACTACACCTTC | 1 | TRBV2*00 | TRBD1*00 | TRBJ1-2*00 | 0.48% |
| CD4+YFP- WT1 | TGTGCCAGCAGCCAAGACGGGGGCGCCTATGAACAGTACTTC | 1 | TRBV2*00 | TRBD1*00 | TRBJ2-7*00 | 0.48% |
| CD4+YFP- WT1 | TGTGCCAGCAGCCAAACTGGGGGGGGTTATGAACAGTACTTC | 1 | TRBV5*00 | TRBD2*00 | TRBJ2-7*00 | 0.48% |
| CD4+YFP- WT1 | TGTGCCAGCAGCAGGACAGCAAACACCGGGCAGCTCTCCTTT | 1 | TRBV3*00 | TRBD1*00 | TRBJ2-2*00 | 0.48% |
| CD4+YFP- WT1 | TGTGCAAGCAGCTTAGAGCTGGGGGGGGTTGAACAGTACTTC | 1 | TRBV16*00 | TRBD2*00 | TRBJ2-7*00 | 0.48% |
| CD4+YFP- WT1 | TGTGCAAGCAGCTTAGAGGGGTCAAACTCCGACTACACCTTC | 1 | TRBV16*00 | | TRBJ1-2*00 | 0.48% |
| CD4+YFP- WT1 | TGTGGTGCTAGGTCTGGGGGGGACTATGCTGAGCAGTTCTTC | 1 | TRBV20*00 | TRBD2*00 | TRBJ2-1*00 | 0.48% |
| CD4+YFP- WT1 | TGTGGTGCTAGGGATTGGTCTAGTGCAGAAACGCTGTATTTT | 1 | TRBV20*00 | | TRBJ2-3*00 | 0.48% |
| CD4+YFP- WT1 | TGCACCTGCAGTGCAGCCCCACGTGCAGAAACGCTGTATTTT | 1 | TRBV1*00 | TRBD2*00 | TRBJ2-3*00 | 0.48% |
| CD4+YFP- WT1 | TGCACCTGCAGTGACAGAAATTCTGGAAATACGCTCTATTTT | 1 | TRBV1*00 | | TRBJ1-3*00 | 0.48% |
| CD4+YFP- WT1 | TGCACCTGCAAGGGGACAGATAGTGCAGAAACGCTGTATTTT | 1 | TRBV1*00 | TRBD1*00 | TRBJ2-3*00 | 0.48% |
| CD4+YFP- WT1 | TGTGCCAGCAGTGGGGGGAAACACCGGGCAGCTCTACTTT | 1 | TRBV13-1*00 | TRBD1*00 | TRBJ2-2*00 | 0.48% |
| CD4+YFP- WT1 | TGTGCTAGCAGTAGGGACAGCTATGCTGAGCAGTTCTTC | 1 | TRBV17*00 | TRBD1*00 | TRBJ2-1*00 | 0.48% |
| CD4+YFP- WT1 | TGTGCCTGGAGTCTAGGGGGGGAAGACACCCAGTACTTT | 1 | TRBV31*00 | TRBD2*00 | TRBJ2-5*00 | 0.48% |
| CD4+YFP- WT1 | TGTGCCAGCCTCATGCCAAGCTCCTATGAACAGTACTTC | 1 | TRBV13-1*00 | | TRBJ2-7*00 | 0.48% |
| CD4+YFP- WT1 | TGTGCCAGCCGGGACAGGGTCTATGCTGAGCAGTTCTTC | 1 | TRBV14*00 | TRBD1*00 | TRBJ2-1*00 | 0.48% |
| CD4+YFP- WT1 | TGTGCCAGCGGTGATGAGGGGTTTGACACCCAGTACTTT | 1 | TRBV13-2*00 | TRBD1*00 | TRBJ2-5*00 | 0.48% |
| CD4+YFP- WT1 | TGTGCCAGCGGGGGTGCTAGTGCAGAAACGCTGTATTTT | 1 | TRBV13-1*00 | TRBD1*00 | TRBJ2-3*00 | 0.48% |
| CD4+YFP- WT1 | TGTGCCAGCGGGGACAATTCTGGAAATACGCTCCATTTT | 1 | TRBV13-2*00 | | TRBJ1-3*00 | 0.48% |
| CD4+YFP- WT1 | TGTGCCAGCAGTTTTCGGGGCGGAGACACCCAGTACTTT | 1 | TRBV15*00 | TRBD1*00 | TRBJ2-5*00 | 0.48% |
| CD4+YFP- WT1 | TGTGCCAGCAGTTTCTGGGGTCTTTATGAACAGTACTTC | 1 | TRBV15*00 | TRBD2*00 | TRBJ2-7*00 | 0.48% |
| CD4+YFP- WT1 | TGTGCCAGCAGTTACAATGCAAACTCCGACTACACCTTC | 1 | TRBV19*00 | | TRBJ1-2*00 | 0.48% |
| CD4+YFP- WT1 | TGTGCCAGCAGTCTTGGCAAAACCGGGCAGCTCTACTTT | 1 | TRBV26*00 | | TRBJ2-2*00 | 0.48% |
| CD4+YFP- WT1 | TGTGCCAGCAGTCTGTCCACAAACTCCGACTACACCTTC | 1 | TRBV26*00 | | TRBJ1-2*00 | 0.48% |
| CD4+YFP- WT1 | TGTGCCAGCAGTCGGGAAAACTATGCTGAGCAGTTCTTC | 1 | TRBV14*00 | TRBD1*00 | TRBJ2-1*00 | 0.48% |
| CD4+YFP- WT1 | TGTGCCAGCAGTGATTGGGACCAAGACACCCAGTACTTT | 1 | TRBV13-3*00 | | TRBJ2-5*00 | 0.48% |
| CD4+YFP- WT1 | TGTGCCAGCAGTGATTGGGACAGTTATGAACAGTACTTC | 1 | TRBV13-3*00 | TRBD1*00 | TRBJ2-7*00 | 0.48% |
| CD4+YFP- WT1 | TGTGCCAGCAGTGATCTGGGTCAAAACACCTTGTACTTT | 1 | TRBV13-1*00 | | TRBJ2-4*00 | 0.48% |
| CD4+YFP- WT1 | TGTGCCAGCAGTGATCGTGCAAACACAGAAGTCTTCTTT | 1 | TRBV13-1*00 | | TRBJ1-1*00 | 0.48% |
| CD4+YFP- WT1 | TGTGCCAGCAGTGATGCGAGGGACTCCGACTACACCTTC | 1 | TRBV13-3*00 | | TRBJ1-2*00 | 0.48% |
| CD4+YFP- WT1 | TGTGCCAGCAGTGATGCAGGGCCCGGGCAGCTCTACTTT | 1 | TRBV13-3*00 | | TRBJ2-2*00 | 0.48% |
| CD4+YFP- WT1 | TGTGCCAGCAGTGATAATTCTGGAAATACGCTCTATTTT | 1 | TRBV13-1*00 | | TRBJ1-3*00 | 0.48% |
| CD4+YFP- WT1 | TGTGCCAGCAGTGACAGGGGGCAAGACACCCAGTACTTT | 1 | TRBV13-1*00 | TRBD1*00 | TRBJ2-5*00 | 0.48% |
| CD4+YFP- WT1 | TGTGCCAGCAGTGACAGGGAAAACTCCGACTACACCTTC | 1 | TRBV13-3*00 | TRBD1*00 | TRBJ1-2*00 | 0.48% |
| CD4+YFP- WT1 | TGTGCCAGCAGTGAACTGGGGGGGCCTGAACAGTACTTC | 1 | TRBV13-1*00 | TRBD2*00 | TRBJ2-7*00 | 0.48% |
| CD4+YFP- WT1 | TGTGCCAGCAGTGAAGGGGGGCCCTATGAACAGTACTTC | 1 | TRBV13-3*00 | TRBD2*00 | TRBJ2-7*00 | 0.48% |
| CD4+YFP- WT1 | TGTGCCAGCAGTATGGAGAAAAACACAGAAGTCTTCTTT | 1 | TRBV19*00 | | TRBJ1-1*00 | 0.48% |
| CD4+YFP- WT1 | TGTGCCAGCAGCTTCGGAACAAACTCCGACTACACCTTC | 1 | TRBV3*00 | | TRBJ1-2*00 | 0.48% |
| CD4+YFP- WT1 | TGTGCCAGCAGCTTAGACTGGGGTGCTGAGCAGTTCTTC | 1 | TRBV3*00 | TRBD2*00 | TRBJ2-1*00 | 0.48% |
| CD4+YFP- WT1 | TGTGCCAGCAGCTTAGAGGGCGGCTCCGACTACACCTTC | 1 | TRBV3*00 | TRBD1*00 | TRBJ1-2*00 | 0.48% |
| CD4+YFP- WT1 | TGTGCCAGCAGCTGGGACTGGGTCTATGAACAGTACTTC | 1 | TRBV15*00 | TRBD2*00 | TRBJ2-7*00 | 0.48% |
| CD4+YFP- WT1 | TGTGCCAGCAGCTATCTGGGTTCTGCTGAGCAGTTCTTC | 1 | TRBV4*00 | TRBD2*00 | TRBJ2-1*00 | 0.48% |
| CD4+YFP- WT1 | TGTGCCAGCAGCCCCCGGACAAATGCTGAGCAGTTCTTC | 1 | TRBV5*00 | TRBD1*00 | TRBJ2-1*00 | 0.48% |
| CD4+YFP- WT1 | TGTGCCAGCAGCCCGACAGGGCCCGGGCAGCTCTACTTT | 1 | TRBV4*00 | TRBD1*00 | TRBJ2-2*00 | 0.48% |
| CD4+YFP- WT1 | TGTGCCAGCAGCCACTGGGGGAGAAACACCTTGTACTTT | 1 | TRBV4*00 | TRBD2*00 | TRBJ2-4*00 | 0.48% |
| CD4+YFP- WT1 | TGTGCCAGCAGCCACAGGGGGAACTCCGACTACACCTTC | 1 | TRBV2*00 | TRBD1*00 | TRBJ1-2*00 | 0.48% |
| CD4+YFP- WT1 | TGTGCCAGCAGCCAAGATGACTCCTATGAACAGTACTTC | 1 | TRBV5*00 | | TRBJ2-7*00 | 0.48% |
| CD4+YFP- WT1 | TGTGCCAGCAGCCAAGACGGACAGGGGGAAGTCTTCTTT | 1 | TRBV2*00 | TRBD1*00 | TRBJ1-1*00 | 0.48% |
| CD4+YFP- WT1 | TGTGCCAGCAGCCAAGACAGATCCGGGCAGCTCTACTTT | 1 | TRBV5*00 | | TRBJ2-2*00 | 0.48% |
| CD4+YFP- WT1 | TGTGCCAGCAGCCAAACAGGGGGCTCCGACTACACCTTC | 1 | TRBV2*00 | TRBD1*00 | TRBJ1-2*00 | 0.48% |
| CD4+YFP- WT1 | TGTGCCAGCAGCACGACACAGGGGTGGGAACAGTACTTC | 1 | TRBV2*00 | TRBD1*00 | TRBJ2-7*00 | 0.48% |
| CD4+YFP- WT1 | TGTGCCAGCAGCAGACAGGGGGCGCTTGAACAGTACTTC | 1 | TRBV4*00 | TRBD1*00 | TRBJ2-7*00 | 0.48% |
| CD4+YFP- WT1 | TGTGCCAGCAGGGACAATTCTGGAAATACGCTCTATTTT | 1 | TRBV13-3*00 | | TRBJ1-3*00 | 0.48% |
| CD4+YFP- WT1 | TGTGCCAGCAACCAGGGGGCGAACACAGAAGTCTTCTTT | 1 | TRBV13-1*00 | TRBD1*00 | TRBJ1-1*00 | 0.48% |
| CD4+YFP- WT1 | TGTGCAAGCAGCTTAGTAGAGAACCAGGCTCCGCTTTTT | 1 | TRBV16*00 | | TRBJ1-5*00 | 0.48% |
| CD4+YFP- WT1 | TGTGCAAGCAGCTTAGCGGGGTCCTATGAACAGTACTTC | 1 | TRBV16*00 | | TRBJ2-7*00 | 0.48% |
| CD4+YFP- WT1 | TGTGCAAGCAGCTTAGAGAGGGGCTCCGACTACACCTTC | 1 | TRBV16*00 | TRBD1*00 | TRBJ1-2*00 | 0.48% |
| CD4+YFP- WT1 | TGTGCAAGCAGCCGGGACAACAACGAAAGATTATTTTTC | 1 | TRBV16*00 | TRBD1*00 | TRBJ1-4*00 | 0.48% |
| CD4+YFP- WT1 | TGTGCAAGCAGCACAAATTCTGGAAATACGCTCTATTTT | 1 | TRBV16*00 | | TRBJ1-3*00 | 0.48% |
| CD4+YFP- WT1 | TGTGCAAGCAGACCAGGGGGCTATGCTGAGCAGTTCTTC | 1 | TRBV16*00 | TRBD1*00 | TRBJ2-1*00 | 0.48% |
| CD4+YFP- WT1 | TGTGGTGCTAGGTCGACAGTTCAAGACACCCAGTACTTT | 1 | TRBV20*00 | TRBD1*00 | TRBJ2-5*00 | 0.48% |
| CD4+YFP- WT1 | TGTGGTGCTAGGTCAGGCAACTATGCTGAGCAGTTCTTC | 1 | TRBV20*00 | | TRBJ2-1*00 | 0.48% |
| CD4+YFP- WT1 | TGTGGTGCTAGGGATTGGGGCGCAGAAACGCTGTATTTT | 1 | TRBV20*00 | TRBD1*00 | TRBJ2-3*00 | 0.48% |
| CD4+YFP- WT1 | TGTGGTGCTAGGACAGGGCGGAACACAGAAGTCTTCTTT | 1 | TRBV20*00 | TRBD1*00 | TRBJ1-1*00 | 0.48% |
| CD4+YFP- WT1 | TGCTCCAGCCCGGGACAAACAAACACAGAAGTCTTCTTT | 1 | TRBV23*00 | TRBD1*00 | TRBJ1-1*00 | 0.48% |
| CD4+YFP- WT1 | TGCACCTGCAGAGGGGACAGCTATGCTGAGCAGTTCTTC | 1 | TRBV1*00 | TRBD1*00 | TRBJ2-1*00 | 0.48% |
| CD4+YFP- WT1 | TGCACCTGCAGTGCATAACCAACCAGGCTCCGCTTTTT | 1 | TRBV1*00 | | TRBJ1-5*00 | 0.48% |
| CD4+YFP- WT1 | TGTGCCTGCACTGGGGAACTATGCTGAGCAGTTCTTC | 1 | TRBV31*00 | TRBD2*00 | TRBJ2-1*00 | 0.48% |
| CD4+YFP- WT1 | TGTGCCAGCAGCCAAGTCACAGCAGGCTCCGCTTTTT | 1 | TRBV2*00 | | TRBJ1-5*00 | 0.48% |
| CD4+YFP- WT1 | TGTGCTAGCAGTAGAGGGACCCTCGAACAGTACTTC | 1 | TRBV17*00 | | TRBJ2-7*00 | 0.48% |
| CD4+YFP- WT1 | TGTGCTAGCAGCCCACTCCCCGAAAGATTATTTTTC | 1 | TRBV21*00 | TRBD2*00 | TRBJ1-4*00 | 0.48% |
| CD4+YFP- WT1 | TGTGCCTGGACAGAACATAATTCGCCCCTCTACTTT | 1 | TRBV31*00 | | TRBJ1-6*00 | 0.48% |
| CD4+YFP- WT1 | TGTGCCTGGAGTCTTGGAAACTCCGACTACACCATC | 1 | TRBV31*00 | | TRBJ1-2*00 | 0.48% |
| CD4+YFP- WT1 | TGTGCCTGGAGTCCAGGGGACTCCGACTACACCTTC | 1 | TRBV31*00 | TRBD1*00 | TRBJ1-2*00 | 0.48% |
| CD4+YFP- WT1 | TGTGCCTGGAGGAATGCAAACACAGAAGTCTTCTTT | 1 | TRBV31*00 | | TRBJ1-1*00 | 0.48% |
| CD4+YFP- WT1 | TGTGCCAGCTCTCTCCAGGGTTATGAACAGTACTTC | 1 | TRBV12-2*00 | TRBD1*00 | TRBJ2-7*00 | 0.48% |
| CD4+YFP- WT1 | TGTGCCAGCTCTCTCGACAGATATGAACAGTACTTC | 1 | TRBV12-1*00 | | TRBJ2-7*00 | 0.48% |
| CD4+YFP- WT1 | TGTGCCAGCGGTGATGCTGACTCCGACTACACCTTC | 1 | TRBV13-2*00 | | TRBJ1-2*00 | 0.48% |
| CD4+YFP- WT1 | TGTGCCAGCGGGAACAGCAACCAGGCTCCGCTTTTT | 1 | TRBV13-2*00 | | TRBJ1-5*00 | 0.48% |
| CD4+YFP- WT1 | TGTGCCAGCGGAGGGTCTGGAAATACGCTCTATTTT | 1 | TRBV13-2*00 | | TRBJ1-3*00 | 0.48% |

| | | | | | | |
|---|---|---|---|---|---|---|
| CD4+YFP- WT1 | TGTGCCAGCAGTTACATGAACACAGAAGTCTTCTTT | 1 | TRBV26*00 | | TRBJ1-1*00 | 0.48% |
| CD4+YFP- WT1 | TGTGCCAGCAGTCTACAAAAAAACACCTTGTACTTT | 1 | TRBV26*00 | | TRBJ2-4*00 | 0.48% |
| CD4+YFP- WT1 | TGTGCCAGCAGTCCGCAATATGCTGAGCAGTTCTTC | 1 | TRBV26*00 | | TRBJ2-1*00 | 0.48% |
| CD4+YFP- WT1 | TGTGCCAGCAGTCGGGACTATGCTGAGCAGTTCTTC | 1 | TRBV14*00 | | TRBJ2-1*00 | 0.48% |
| CD4+YFP- WT1 | TGTGCCAGCAGTGATCCGAACTCCGACTACACCTTC | 1 | TRBV13-1*00 | | TRBJ1-2*00 | 0.48% |
| CD4+YFP- WT1 | TGTGCCAGCAGTGATGGCCTTTATGAACAGTACTTC | 1 | TRBV13-1*00 | | TRBJ2-7*00 | 0.48% |
| CD4+YFP- WT1 | TGTGCCAGCAGTGATGGCCGCAGGACCTTGTACTTT | 1 | TRBV13-1*00 | | TRBJ2-4*00 | 0.48% |
| CD4+YFP- WT1 | TGTGCCAGCAGTGATGAAGGGGCAGGAGTCTTCTTT | 1 | TRBV13-1*00 | TRBD1*00 | TRBJ1-1*00 | 0.48% |
| CD4+YFP- WT1 | TGTGCCAGCAGTGAAAGTGCAGAAACGCTGTATTTT | 1 | TRBV13-1*00 | | TRBJ2-3*00 | 0.48% |
| CD4+YFP- WT1 | TGTGCCAGCAGCTTAGGGCCCTATGAACAGTACTTC | 1 | TRBV3*00 | | TRBJ2-7*00 | 0.48% |
| CD4+YFP- WT1 | TGTGCCAGCAGCTCGGACAGGGTTGAACAGTACTTC | 1 | TRBV3*00 | TRBD1*00 | TRBJ2-7*00 | 0.48% |
| CD4+YFP- WT1 | TGTGCCAGCAGCTCGGACAGAAACACCTTGTACTTT | 1 | TRBV4*00 | TRBD1*00 | TRBJ2-4*00 | 0.48% |
| CD4+YFP- WT1 | TGTGCCAGCAGCTCGACAAAAAACACCTTGTACTTT | 1 | TRBV4*00 | | TRBJ2-4*00 | 0.48% |
| CD4+YFP- WT1 | TGTGCCAGCAGCCTTAGTCAAAACACCTTGTACTTT | 1 | TRBV3*00 | | TRBJ2-4*00 | 0.48% |
| CD4+YFP- WT1 | TGTGCCAGCAGCCCCCTAACCTATGAACAGTACTTC | 1 | TRBV4*00 | TRBD1*00 | TRBJ2-7*00 | 0.48% |
| CD4+YFP- WT1 | TGTGCCAGCAGCCCGAGCTCCTATGAACAGTACTTC | 1 | TRBV3*00 | | TRBJ2-7*00 | 0.48% |
| CD4+YFP- WT1 | TGTGCCAGCAGCCGGGACACCGGGCAGCTCTACTTT | 1 | TRBV4*00 | | TRBJ2-2*00 | 0.48% |
| CD4+YFP- WT1 | TGTGCCAGCAGCCAAGACTGGGACACCCAGTACTTT | 1 | TRBV2*00 | | TRBJ2-5*00 | 0.48% |
| CD4+YFP- WT1 | TGTGCCAGCAGGGACAGTGCTTATGAACAGTACTTC | 1 | TRBV13-3*00 | TRBD1*00 | TRBJ2-7*00 | 0.48% |
| CD4+YFP- WT1 | TGTGCCAGGGACAAAAACACCGGGCAGCTCTACTTT | 1 | TRBV13-1*00 | TRBD1*00 | TRBJ2-2*00 | 0.48% |
| CD4+YFP- WT1 | TGTGCAAGCAGCTTACACTCCTATGAACAGTACTTC | 1 | TRBV16*00 | | TRBJ2-7*00 | 0.48% |
| CD4+YFP- WT1 | TGTGCAAGCAGCTCCAGGGAGGACACCCAGTACTTT | 1 | TRBV16*00 | TRBD1*00 | TRBJ2-5*00 | 0.48% |
| CD4+YFP- WT1 | TGTGCAAGCAGCCCGGGCTATGCTGAGCAGTTCTTC | 1 | TRBV16*00 | TRBD1*00 | TRBJ2-1*00 | 0.48% |
| CD4+YFP- WT1 | TGTGCAAGCAGGCCTAGTGCAGAAACGCTGTATTTT | 1 | TRBV16*00 | | TRBJ2-3*00 | 0.48% |
| CD4+YFP- WT1 | TGTGGTGCTAGGAGTGGCCAAGACACCCAGTACTTT | 1 | TRBV20*00 | | TRBJ2-5*00 | 0.48% |
| CD4+YFP- WT1 | TGCTCCAGCAGGGGGGGCTCTGACACCCAGTACTTT | 1 | TRBV23*00 | TRBD2*00 | TRBJ2-5*00 | 0.48% |
| CD4+YFP- WT1 | TGCACCTGCAGTGCTTCCAACCAGGCTCCGCTTTTT | 1 | TRBV1*00 | | TRBJ1-5*00 | 0.48% |
| CD4+YFP- WT1 | TGCACCTGCAGTGCGGGGGGTTATGAACAGTACTTC | 1 | TRBV1*00 | TRBD2*00 | TRBJ2-7*00 | 0.48% |
| CD4+YFP- WT1 | TGCACCTGCAGTGCGATTGGTTATGAACAGTACTTC | 1 | TRBV1*00 | | TRBJ2-7*00 | 0.48% |
| CD4+YFP- WT1 | TGCACCTGCAGTGCAGATGGGGGGGGTCAGTACTTC | 1 | TRBV1*00 | TRBD2*00 | TRBJ2-7*00 | 0.48% |
| CD4+YFP- WT1 | TGCACCTGCAGTGCAGATAACTCCGACTACACCTTC | 1 | TRBV1*00 | | TRBJ1-2*00 | 0.48% |
| CD4+YFP- WT1 | TGTGCCTGGAGTCTAGGCGAAAGATTATTTTTC | 1 | TRBV31*00 | | TRBJ1-4*00 | 0.48% |
| CD4+YFP- WT1 | TGTGCCTGGAGGACTAACTCCGACTACACCTTC | 1 | TRBV31*00 | | TRBJ1-2*00 | 0.48% |
| CD4+YFP- WT1 | TGTGCCTGGAGAGGGGACTCCGACTACACCTTC | 1 | TRBV31*00 | TRBD1*00 | TRBJ1-2*00 | 0.48% |
| CD4+YFP- WT1 | TGTGCCAGCCGGACATCCTATGAACAGTACTTC | 1 | TRBV19*00 | TRBD1*00 | TRBJ2-7*00 | 0.48% |
| CD4+YFP- WT1 | TGTGCCAGCGGTGATGCTTCTCAACAGTACTTC | 1 | TRBV13-2*00 | | TRBJ2-7*00 | 0.48% |
| CD4+YFP- WT1 | TGTGCCAGCAGTTTAGGGGGGGAAGTCTTCTTT | 1 | TRBV15*00 | TRBD1*00 | TRBJ1-1*00 | 0.48% |
| CD4+YFP- WT1 | TGTGCCAGCAGTTTAGACGGGGAAGTCTTCTTT | 1 | TRBV15*00 | | TRBJ1-1*00 | 0.48% |
| CD4+YFP- WT1 | TGTGCCAGCAGTTCCCCTGGGCAGCTCTACTTT | 1 | TRBV15*00 | TRBD1*00 | TRBJ2-2*00 | 0.48% |
| CD4+YFP- WT1 | TGTGCCAGCAGTGATCGGGACACCCAGTACTTT | 1 | TRBV13-1*00 | | TRBJ2-5*00 | 0.48% |
| CD4+YFP- WT1 | TGTGCCAGCAGTGACAGGGGAGAAGTCTTCTTT | 1 | TRBV13-3*00 | TRBD1*00 | TRBJ1-1*00 | 0.48% |
| CD4+YFP- WT1 | TGTGCCAGCAGTATGAACGAAAGATTATTTTTC | 1 | TRBV19*00 | | TRBJ1-4*00 | 0.48% |
| CD4+YFP- WT1 | TGTGCCAGCAGTATAAGGGATGGGCAGTACTTC | 1 | TRBV19*00 | | TRBJ2-7*00 | 0.48% |
| CD4+YFP- WT1 | TGTGCCAGCAGCCAAGACGAAACGCTGTATTTT | 1 | TRBV2*00 | | TRBJ2-3*00 | 0.48% |
| CD4+YFP- WT1 | TGTGCAAGCAGCTTGGGGGTTGAACAGTACTTC | 1 | TRBV16*00 | TRBD1*00 | TRBJ2-7*00 | 0.48% |
| CD4+YFP- WT1 | TGCACCTGCAGTGCAGAGGCTGAGCAGTTCTTC | 1 | TRBV1*00 | | TRBJ2-1*00 | 0.48% |
| CD4+YFP- WT1 | TGTGCCAGCTCCCGGTGACACCCAGTACTTT | 1 | TRBV18*00 | | TRBJ2-5*00 | 0.48% |
| CD4+YFP- WT1 | TGTGCCAGCGGTGAAGACACCCAGTACTTT | 1 | TRBV13-2*00 | | TRBJ2-5*00 | 0.48% |
| CD4+YFP- WT1 | TGTGCCAGCAGCGCTTCTGAACAGTACTTC | 1 | TRBV4*00 | | TRBJ2-7*00 | 0.48% |
| CD4+YFP- WT1 | TGTGCCTGGAGTCTAGAAGTCTTCTTT | 1 | TRBV31*00 | | TRBJ1-1*00 | 0.48% |

#### CD4+YFP- WT2

| Sample name | CDR3 Sequence | clone Count | best V Hit | best D Hit | best J Hit | Frequency |
|---|---|---|---|---|---|---|
| CD4+YFP- WT2 | TGTGCTAGCAGTAGATTGGGGGGGCAAGACACCCAGTACTTT | 2 | TRBV17*00 | TRBD2*00 | TRBJ2-5*00 | 0.70% |
| CD4+YFP- WT2 | TGTGCTAGCAGTAGACTGGGGAACTATGCTGAGCAGTTCTTC | 2 | TRBV17*00 | TRBD2*00 | TRBJ2-1*00 | 0.70% |
| CD4+YFP- WT2 | TGTGCCTGGAGGGGGACAGGGGGAAACACAGAAGTCTTCTTT | 2 | TRBV31*00 | TRBD1*00 | TRBJ1-1*00 | 0.70% |
| CD4+YFP- WT2 | TGTGCCAGCAGTTTCTTGGGGATCACCGGGCAGCTCTACTTT | 2 | TRBV14*00 | TRBD2*00 | TRBJ2-2*00 | 0.70% |
| CD4+YFP- WT2 | TGTGCCAGCAGCCAACAAGATTCTGGAAATACGCTCTATTTT | 2 | TRBV2*00 | | TRBJ1-3*00 | 0.70% |
| CD4+YFP- WT2 | TGTGCCAGCAGCTTAGAACACAACTCCGACTACACCTTC | 2 | TRBV3*00 | | TRBJ1-2*00 | 0.70% |
| CD4+YFP- WT2 | TGTGCCAGCAGCCCACTGGACCAAGACACCCAGTACTTT | 2 | TRBV13-3*00 | TRBD2*00 | TRBJ2-5*00 | 0.70% |
| CD4+YFP- WT2 | TGTGCAAGCAGCTCGTATAACAACCAGGCTCCGCTTTTT | 2 | TRBV16*00 | | TRBJ1-5*00 | 0.70% |
| CD4+YFP- WT2 | TGCACCTGCAGTGCCGACAGGAACTCCGACTACACCTTC | 2 | TRBV1*00 | TRBD1*00 | TRBJ1-2*00 | 0.70% |
| CD4+YFP- WT2 | TGTGCCAGCAGTCCGGACCAAGACACCCAGTACTTT | 2 | TRBV4*00 | | TRBJ2-5*00 | 0.70% |
| CD4+YFP- WT2 | TGTGCCAGCAGTCATCCTAATTCGCCCCTCTACTTT | 2 | TRBV26*00 | | TRBJ1-6*00 | 0.70% |
| CD4+YFP- WT2 | TGTGCCAGCAGCGACAGCAACCAGGCTCCGCTTTTT | 2 | TRBV2*00 | TRBD1*00 | TRBJ1-5*00 | 0.70% |
| CD4+YFP- WT2 | TGTGGTGCTAGGGCGGGTGCAGAAACGCTGTATTTT | 2 | TRBV20*00 | | TRBJ2-3*00 | 0.70% |
| CD4+YFP- WT2 | TGTGCCAGCAGCTTAGACTCCGACTACACCTTC | 2 | TRBV3*00 | | TRBJ1-2*00 | 0.70% |
| CD4+YFP- WT2 | TGTGCCAGCAGCCGGACAACTGAGCAGTTCTTC | 2 | TRBV14*00 | TRBD1*00 | TRBJ2-1*00 | 0.70% |
| CD4+YFP- WT2 | TGTGGTGCTAGGGATGATGCTGAGCAGTTCTTC | 2 | TRBV20*00 | | TRBJ2-1*00 | 0.70% |
| CD4+YFP- WT2 | TGTGCCAGCGGTGAGGGGGACAGATCTAGTGCAGAAACGCTGTATTTT | 1 | TRBV13-2*00 | TRBD1*00 | TRBJ2-3*00 | 0.35% |
| CD4+YFP- WT2 | TGTGCCAGCGGAGTTCGGACTGGGGGGGCCTCCTATGAACAGTACTTC | 1 | TRBV13-2*00 | TRBD2*00 | TRBJ2-7*00 | 0.35% |
| CD4+YFP- WT2 | TGTGCCAGCAGTGATTGGGACAGCTCCTATAATTCGCCCCTCTACTTT | 1 | TRBV13-3*00 | TRBD1*00 | TRBJ1-6*00 | 0.35% |
| CD4+YFP- WT2 | TGTGCCAGCAGCTTAGTGGCAGGGAATTCTGGAAATACGCTCTATTTT | 1 | TRBV3*00 | | TRBJ1-3*00 | 0.35% |
| CD4+YFP- WT2 | TGTGCCAGCAGCCCCCGACTGGGGACTAGTGCAGAAACGCTGTATTTT | 1 | TRBV5*00 | TRBD2*00 | TRBJ2-3*00 | 0.35% |
| CD4+YFP- WT2 | TGTGCCAGCAGCCAAGATGCCTGGGGGAGCTCCTATGAACAGTACTTC | 1 | TRBV5*00 | TRBD2*00 | TRBJ2-7*00 | 0.35% |
| CD4+YFP- WT2 | TGTGCAAGCAGCCCGGGACAGGGTTATTCTGGAAATACGCTCTATTTT | 1 | TRBV16*00 | TRBD1*00 | TRBJ1-3*00 | 0.35% |
| CD4+YFP- WT2 | TGTGCCAGCTCTCTCGGGGGAAAATAACTATGCTGAGCAGTTCTTC | 1 | TRBV12-1*00 | | TRBJ2-1*00 | 0.35% |
| CD4+YFP- WT2 | TGTGCCTGGAGTCTAGAAGTTAATTCTGGAAATACGCTCTATTTT | 1 | TRBV31*00 | | TRBJ1-3*00 | 0.35% |
| CD4+YFP- WT2 | TGTGCCTGGAGCCGGGACATTTATAACTATGCTGAGCAGTTCTTC | 1 | TRBV31*00 | TRBD1*00 | TRBJ2-1*00 | 0.35% |
| CD4+YFP- WT2 | TGTGCCTGGAGCCAAGGGACACCTTCTGGAAATACGCTCTATTTT | 1 | TRBV31*00 | TRBD1*00 | TRBJ1-3*00 | 0.35% |

| | | | | | | |
|---|---|---|---|---|---|---|
| CD4+YFP- WT2 | TGTGCCAGCGGTGATGGGGCGGCAAACACCGGGCAGCTCTACTTT | 1 | TRBV13-2*00 | TRBD1*00 | TRBJ2-2*00 | 0.35% |
| CD4+YFP- WT2 | TGTGCCAGCGGTGATGAGGCCCTTGCAAACTCCGACTACACCTTC | 1 | TRBV13-2*00 | | TRBJ1-2*00 | 0.35% |
| CD4+YFP- WT2 | TGTGCCAGCAGTTACAGGGTCAGGAGCTCCTATGAACAGTACTTC | 1 | TRBV14*00 | TRBD1*00 | TRBJ2-7*00 | 0.35% |
| CD4+YFP- WT2 | TGTGCCAGCAGTGATGTGGGGGCTAGTGCAGAAACGCTGTATTTT | 1 | TRBV13-1*00 | TRBD2*00 | TRBJ2-3*00 | 0.35% |
| CD4+YFP- WT2 | TGTGCCAGCAGTGATGTACGGTTTAACCAAGACACCCAGTACTTT | 1 | TRBV13-3*00 | | TRBJ2-5*00 | 0.35% |
| CD4+YFP- WT2 | TGTGCCAGCAGTGATGCTCGGGGAGAGGGAGACACCCAGTACTTT | 1 | TRBV13-1*00 | TRBD1*00 | TRBJ2-5*00 | 0.35% |
| CD4+YFP- WT2 | TGTGCCAGCAGTGATGCCGAGGGGAACAACCAGGCTCCGCTTTTT | 1 | TRBV13-1*00 | TRBD1*00 | TRBJ1-5*00 | 0.35% |
| CD4+YFP- WT2 | TGTGCCAGCAGTGAAGGGACAGGGGGCATCACAGAAGTCTTCTTT | 1 | TRBV13-1*00 | TRBD1*00 | TRBJ1-1*00 | 0.35% |
| CD4+YFP- WT2 | TGTGCCAGCAGTATAAGGGGCAGGGGAGGGGCTGAGCAGTTCTTC | 1 | TRBV19*00 | TRBD1*00 | TRBJ2-1*00 | 0.35% |
| CD4+YFP- WT2 | TGTGCCAGCAGTATAAGGGACATTGCAAACTCCGACTACACCTTC | 1 | TRBV19*00 | TRBD1*00 | TRBJ1-2*00 | 0.35% |
| CD4+YFP- WT2 | TGTGCCAGCAGCTTAGGGAGGACAGCAAACACAGAAGTCTTCTTT | 1 | TRBV3*00 | TRBD1*00 | TRBJ1-1*00 | 0.35% |
| CD4+YFP- WT2 | TGTGCCAGCAGCTTAGATCAGTCTGGAAACTCCGACTACACCTTC | 1 | TRBV3*00 | | TRBJ1-2*00 | 0.35% |
| CD4+YFP- WT2 | TGTGCCAGCAGCTACAGACAGGGAAGTGCAGAAACGCTGTATTTT | 1 | TRBV4*00 | TRBD1*00 | TRBJ2-3*00 | 0.35% |
| CD4+YFP- WT2 | TGTGCCAGCAGCCTACAGGGGTTTTCTGGAAATACGCTCTATTTT | 1 | TRBV26*00 | TRBD1*00 | TRBJ1-3*00 | 0.35% |
| CD4+YFP- WT2 | TGTGCCAGCAGCCAACCGGGACAGGGGGCTGACACCCAGTACTTT | 1 | TRBV2*00 | TRBD1*00 | TRBJ2-5*00 | 0.35% |
| CD4+YFP- WT2 | TGTGCCAGCAGCCAAGTCCGGGGTACAAACTCCGACTACACCTTC | 1 | TRBV5*00 | TRBD1*00 | TRBJ1-2*00 | 0.35% |
| CD4+YFP- WT2 | TGTGCCAGCAGCGCGGGACAATATAACAACCAGGCTCCGCTTTTT | 1 | TRBV2*00 | TRBD1*00 | TRBJ1-5*00 | 0.35% |
| CD4+YFP- WT2 | TGTGCCAGCAGCAGCCGGGACAATAGTGCAGAAACGCTGTATTTT | 1 | TRBV13-3*00 | TRBD1*00 | TRBJ2-3*00 | 0.35% |
| CD4+YFP- WT2 | TGTGCAAGCAGCTTAGAGGGGGGGGCGTCCTATGAACAGTACTTC | 1 | TRBV16*00 | TRBD2*00 | TRBJ2-7*00 | 0.35% |
| CD4+YFP- WT2 | TGTGCAAGCAGCTTAGAGGACATCACCTATGCTGAGCAGTTCTTC | 1 | TRBV16*00 | TRBD1*00 | TRBJ2-1*00 | 0.35% |
| CD4+YFP- WT2 | TGTGCAAGCAGCTCGGGACAAACTAGTGCAGAAACGCTGTATTTT | 1 | TRBV16*00 | TRBD1*00 | TRBJ2-3*00 | 0.35% |
| CD4+YFP- WT2 | TGCACCTGCAGTGCAGGGGGGGCGTCTGGAAATACGCTCTATTTT | 1 | TRBV1*00 | TRBD2*00 | TRBJ1-3*00 | 0.35% |
| CD4+YFP- WT2 | TGCACCTGCAGTGCAGATTCGGGGGGGCAGGCTGAGCAGTTCTTC | 1 | TRBV1*00 | TRBD2*00 | TRBJ2-1*00 | 0.35% |
| CD4+YFP- WT2 | TGCACCTGCAGTGCAGAAGGACAGGGGAACACAGAAGTCTTCTTT | 1 | TRBV1*00 | TRBD1*00 | TRBJ1-1*00 | 0.35% |
| CD4+YFP- WT2 | TGTGCCAGCAGCTTATCGGGACCAACTATGCTGAGCAGTTCTTC | 1 | TRBV3*00 | TRBD1*00 | TRBJ2-1*00 | 0.35% |
| CD4+YFP- WT2 | TGTGCCTGGAGTCACTGGGGGGAACTATGCTGAGCAGTTCTTC | 1 | TRBV31*00 | TRBD2*00 | TRBJ2-1*00 | 0.35% |
| CD4+YFP- WT2 | TGTGCTAGCAGTTTATCGGGGGGCTCCTATGAACAGTACTTC | 1 | TRBV29*00 | | TRBJ2-7*00 | 0.35% |
| CD4+YFP- WT2 | TGTGCCTGGAGTCTGGAGGGCCCCAACGAAAGATTATTTTTC | 1 | TRBV31*00 | | TRBJ1-4*00 | 0.35% |
| CD4+YFP- WT2 | TGTGCCTGGAGTCTAGGGCAGGGCGCAGAAACGCTGTATTTT | 1 | TRBV31*00 | TRBD1*00 | TRBJ2-3*00 | 0.35% |
| CD4+YFP- WT2 | TGTGCCTGGAGTCTAGGGGGTAGTCAAAACACCTTGTACTTT | 1 | TRBV31*00 | TRBD1*00 | TRBJ2-4*00 | 0.35% |
| CD4+YFP- WT2 | TGTGCCTGGAGTCTAGAGGGGGCGAGAGACACCCAGTACTTT | 1 | TRBV31*00 | TRBD1*00 | TRBJ2-5*00 | 0.35% |
| CD4+YFP- WT2 | TGTGCCTGGAGATCAGGGAATTCTGGAAATACGCTCTATTTT | 1 | TRBV31*00 | | TRBJ1-3*00 | 0.35% |
| CD4+YFP- WT2 | TGTGCCAGCTCTCTTGGACTGGGGGGAGACACCCAGTACTTT | 1 | TRBV12-1*00 | TRBD2*00 | TRBJ2-5*00 | 0.35% |
| CD4+YFP- WT2 | TGTGCCAGCTCTCTCGAACATTCTGGAAATACGCTCTATTTT | 1 | TRBV12-2*00 | | TRBJ1-3*00 | 0.35% |
| CD4+YFP- WT2 | TGTGCCAGCTCTCCCACATCTAGTGCAGAAACGCTGTATTTT | 1 | TRBV12-1*00 | | TRBJ2-3*00 | 0.35% |
| CD4+YFP- WT2 | TGTGCCAGCTCTACCGACTGGAGCTCCTATGAACAGTACTTC | 1 | TRBV12-1*00 | | TRBJ2-7*00 | 0.35% |
| CD4+YFP- WT2 | TGTGCCAGCTCCGGGACTGGTAGTGCAGAAACGCTGTATTTT | 1 | TRBV12-1*00 | TRBD2*00 | TRBJ2-3*00 | 0.35% |
| CD4+YFP- WT2 | TGTGCCAGCCGCCGGGGAGCGAATCAAAACACCTTGTACTTT | 1 | TRBV19*00 | TRBD1*00 | TRBJ2-4*00 | 0.35% |
| CD4+YFP- WT2 | TGTGCCAGCGGTCCCGACAGGGGCGGTGCTGAGCAGTTCTTC | 1 | TRBV13-2*00 | TRBD1*00 | TRBJ2-1*00 | 0.35% |
| CD4+YFP- WT2 | TGTGCCAGCGGTGGGACAGATTCTGGAAATACGCTCTATTTT | 1 | TRBV13-2*00 | TRBD1*00 | TRBJ1-3*00 | 0.35% |
| CD4+YFP- WT2 | TGTGCCAGCGGTGATGCTCAGGGGAGGTATGAACAGTACTTC | 1 | TRBV13-2*00 | TRBD1*00 | TRBJ2-7*00 | 0.35% |
| CD4+YFP- WT2 | TGTGCCAGCGGTGATGGGGGGCTCCAAGACACCCAGTACTTT | 1 | TRBV13-2*00 | TRBD1*00 | TRBJ2-5*00 | 0.35% |
| CD4+YFP- WT2 | TGTGCCAGCGGTGATGGGACAAACACCGGGCAGCTCTACTTT | 1 | TRBV13-2*00 | | TRBJ2-2*00 | 0.35% |
| CD4+YFP- WT2 | TGTGCCAGCGGTGATGAAGGGGGGCAAGACACCCAGTACTTT | 1 | TRBV13-2*00 | TRBD1*00 | TRBJ2-5*00 | 0.35% |
| CD4+YFP- WT2 | TGTGCCAGCGGTGAAGGGACAAACACCGGGCAGCTCTACTTT | 1 | TRBV13-2*00 | | TRBJ2-2*00 | 0.35% |
| CD4+YFP- WT2 | TGTGCCAGCACCCGGACAGCTAGTGCAGAAACGCTGTATTTT | 1 | TRBV13-3*00 | TRBD1*00 | TRBJ2-3*00 | 0.35% |
| CD4+YFP- WT2 | TGTGCCAGCAGTTTGAGGGGAAACCAAGACACCCAGTACTTT | 1 | TRBV14*00 | TRBD1*00 | TRBJ2-5*00 | 0.35% |
| CD4+YFP- WT2 | TGTGCCAGCAGTCTTTCGTATAACTATGCTGAGCAGTTCTTC | 1 | TRBV26*00 | | TRBJ2-1*00 | 0.35% |
| CD4+YFP- WT2 | TGTGCCAGCAGTCTCGACTGGGGGGCATATGAACAGTACTTC | 1 | TRBV19*00 | TRBD2*00 | TRBJ2-7*00 | 0.35% |
| CD4+YFP- WT2 | TGTGCCAGCAGTCTGTCGAGGCAGACTTATGAACAGTACTTC | 1 | TRBV26*00 | | TRBJ2-7*00 | 0.35% |
| CD4+YFP- WT2 | TGTGCCAGCAGTCTGCAGGGGATTGGAAATACGCTCTATTTT | 1 | TRBV26*00 | TRBD1*00 | TRBJ1-3*00 | 0.35% |
| CD4+YFP- WT2 | TGTGCCAGCAGTCTACCAGGGGAGGGAAATACGCTCTATTTT | 1 | TRBV26*00 | TRBD1*00 | TRBJ1-3*00 | 0.35% |
| CD4+YFP- WT2 | TGTGCCAGCAGTCCTACAGGCCCCTATGCTGAGCAGTTCTTC | 1 | TRBV19*00 | TRBD1*00 | TRBJ2-1*00 | 0.35% |
| CD4+YFP- WT2 | TGTGCCAGCAGTCCGGGACAGAACTATGCTGAGCAGTTCTTC | 1 | TRBV14*00 | TRBD1*00 | TRBJ2-1*00 | 0.35% |
| CD4+YFP- WT2 | TGTGCCAGCAGTCGGACAGATTCTGGAAATACGCTCTATTTT | 1 | TRBV19*00 | TRBD1*00 | TRBJ1-3*00 | 0.35% |
| CD4+YFP- WT2 | TGTGCCAGCAGTCGACAATTTAACCAAGACACCCAGTACTTT | 1 | TRBV26*00 | | TRBJ2-5*00 | 0.35% |
| CD4+YFP- WT2 | TGTGCCAGCAGTGTCTGGGGGGGCTCCTATGAACAGTACTTC | 1 | TRBV13-3*00 | TRBD2*00 | TRBJ2-7*00 | 0.35% |
| CD4+YFP- WT2 | TGTGCCAGCAGTGGGGGACAGGGGTATGCTGAGCAGTTCTTC | 1 | TRBV13-3*00 | TRBD1*00 | TRBJ2-1*00 | 0.35% |
| CD4+YFP- WT2 | TGTGCCAGCAGTGATCGGGGGGCGGCAGAAACGCTGTATTTT | 1 | TRBV13-1*00 | TRBD2*00 | TRBJ2-3*00 | 0.35% |
| CD4+YFP- WT2 | TGTGCCAGCAGTGAGCAGGGGCGGGATGCTGAGCAGTTCTTC | 1 | TRBV13-3*00 | TRBD1*00 | TRBJ2-1*00 | 0.35% |
| CD4+YFP- WT2 | TGTGCCAGCAGTGAGGGGTTTAACCAAGACACCCAGTACTTT | 1 | TRBV13-1*00 | | TRBJ2-5*00 | 0.35% |
| CD4+YFP- WT2 | TGTGCCAGCAGTGAACGACTGGGGGGTGCTGAGCAGTTCTTC | 1 | TRBV13-1*00 | TRBD2*00 | TRBJ2-1*00 | 0.35% |
| CD4+YFP- WT2 | TGTGCCAGCAGTGAAGAAGGAAGGCAAAACACCTTGTACTTT | 1 | TRBV13-3*00 | | TRBJ2-4*00 | 0.35% |
| CD4+YFP- WT2 | TGTGCCAGCAGTATCGGGGGTTCTGGAAATACGCTCTATTTT | 1 | TRBV19*00 | TRBD1*00 | TRBJ1-3*00 | 0.35% |
| CD4+YFP- WT2 | TGTGCCAGCAGTATCATGGGGAGTGCAGAAACGCTGTATTTT | 1 | TRBV19*00 | TRBD2*00 | TRBJ2-3*00 | 0.35% |
| CD4+YFP- WT2 | TGTGCCAGCAGTATGAGGGGAAACACCGGGCAGCTCTACTTT | 1 | TRBV19*00 | TRBD1*00 | TRBJ2-2*00 | 0.35% |
| CD4+YFP- WT2 | TGTGCCAGCAGCTTGACAGATTCTGGAAATACGCTCTATTTT | 1 | TRBV3*00 | TRBD1*00 | TRBJ1-3*00 | 0.35% |
| CD4+YFP- WT2 | TGTGCCAGCAGCTTAGTCGGGAGTGCAGAAACGCTGTATTTT | 1 | TRBV3*00 | | TRBJ2-3*00 | 0.35% |
| CD4+YFP- WT2 | TGTGCCAGCAGCTCGACCGATGCAAACTCCGACTACACCTTC | 1 | TRBV4*00 | | TRBJ1-2*00 | 0.35% |
| CD4+YFP- WT2 | TGTGCCAGCAGCTACTCCGGGGAGGGAAACACCTTGTACTTT | 1 | TRBV4*00 | TRBD1*00 | TRBJ2-4*00 | 0.35% |
| CD4+YFP- WT2 | TGTGCCAGCAGCCTGGGGCCTAGTGCAGAAACGCTGTATTTT | 1 | TRBV3*00 | TRBD2*00 | TRBJ2-3*00 | 0.35% |
| CD4+YFP- WT2 | TGTGCCAGCAGCCCCGGACAGGGGGGCACAGAAGTCTTCTTT | 1 | TRBV15*00 | TRBD1*00 | TRBJ1-1*00 | 0.35% |
| CD4+YFP- WT2 | TGTGCCAGCAGCCCGGAGATTTCCAACGAAAGATTATTTTTC | 1 | TRBV5*00 | | TRBJ1-4*00 | 0.35% |
| CD4+YFP- WT2 | TGTGCCAGCAGCCCATATTCCTATAATTCGCCCCTCTACTTT | 1 | TRBV5*00 | | TRBJ1-6*00 | 0.35% |

| | | | | | | |
|---|---|---|---|---|---|---|
| CD4+YFP- WT2 | TGTGCCAGCAGCCGGGACAATAGTGCAGAAACGCTGTATTTT | 1 | TRBV13-3*00 | TRBD1*00 | TRBJ2-3*00 | 0.35% |
| CD4+YFP- WT2 | TGTGCCAGCAGCCAACCAGAAAACACCGGGCAGCTCTACTTT | 1 | TRBV2*00 | | TRBJ2-2*00 | 0.35% |
| CD4+YFP- WT2 | TGTGCCAGCAGCCAAGGGTTTAACTATGCTGAGCAGTTCTTC | 1 | TRBV5*00 | | TRBJ2-1*00 | 0.35% |
| CD4+YFP- WT2 | TGTGCCAGCAGCCAAGGGACACCAAACTCCGACTACACCTTC | 1 | TRBV2*00 | TRBD1*00 | TRBJ1-2*00 | 0.35% |
| CD4+YFP- WT2 | TGTGCCAGCAGCCAAGGGACAGACTATGCTGAGCAGTTCTTC | 1 | TRBV2*00 | TRBD1*00 | TRBJ2-1*00 | 0.35% |
| CD4+YFP- WT2 | TGTGCCAGCAGCCAAGGACATACCAACGAAAGATTATTTTTC | 1 | TRBV5*00 | | TRBJ1-4*00 | 0.35% |
| CD4+YFP- WT2 | TGTGCCAGCAGCCAAGATCCCGGGGGGGTTGAACAGTACTTC | 1 | TRBV2*00 | TRBD2*00 | TRBJ2-7*00 | 0.35% |
| CD4+YFP- WT2 | TGTGCCAGCAGCCAAGAAAATTCTGGAAATACGCTCTATTTT | 1 | TRBV2*00 | | TRBJ1-3*00 | 0.35% |
| CD4+YFP- WT2 | TGTGCCAGCAGCCAAAATTCCTATAATTCGCCCCTCTACTTT | 1 | TRBV5*00 | | TRBJ1-6*00 | 0.35% |
| CD4+YFP- WT2 | TGTGCCAGCAGCATCGACAATTCTGGAAATACGCTCTATTTT | 1 | TRBV3*00 | | TRBJ1-3*00 | 0.35% |
| CD4+YFP- WT2 | TGTGCAAGCAGCTTATCTGGAAACACCGGGCAGCTCTACTTT | 1 | TRBV16*00 | | TRBJ2-2*00 | 0.35% |
| CD4+YFP- WT2 | TGTGCAAGCAGCTTACAGGGGTCCTATGCTGAGCAGTTCTTC | 1 | TRBV16*00 | TRBD1*00 | TRBJ2-1*00 | 0.35% |
| CD4+YFP- WT2 | TGTGCAAGCAGCTTAGTCCAGGGAAGCACAGAAGTCTTCTTT | 1 | TRBV16*00 | TRBD1*00 | TRBJ1-1*00 | 0.35% |
| CD4+YFP- WT2 | TGTGCAAGCAGCTTAGCGGTTAGTGCAGAAACGCTGTATTTT | 1 | TRBV16*00 | | TRBJ2-3*00 | 0.35% |
| CD4+YFP- WT2 | TGTGCAAGCAGCTTAGGACAGGGGAACTCCGACTACACCTTC | 1 | TRBV16*00 | TRBD1*00 | TRBJ1-2*00 | 0.35% |
| CD4+YFP- WT2 | TGTGCAAGCAGCTTAGATACTGGGCAAGACACCCAGTACTTT | 1 | TRBV16*00 | TRBD2*00 | TRBJ2-5*00 | 0.35% |
| CD4+YFP- WT2 | TGTGCAAGCAGCCGACAGGGGACAAACTCCGACTACACCTTC | 1 | TRBV16*00 | TRBD1*00 | TRBJ1-2*00 | 0.35% |
| CD4+YFP- WT2 | TGTGGTGCTAGGGGGGGAACAACAAACTCCGACTACACCTTC | 1 | TRBV20*00 | | TRBJ1-2*00 | 0.35% |
| CD4+YFP- WT2 | TGTGGTGCTAGGACAGGGGGCTATAATTCGCCCCTCTACTTT | 1 | TRBV20*00 | TRBD1*00 | TRBJ1-6*00 | 0.35% |
| CD4+YFP- WT2 | TGCACCTGCAGTGCCCCGGGACTGGGTGCTGAGCAGTTCTTC | 1 | TRBV1*00 | TRBD2*00 | TRBJ2-1*00 | 0.35% |
| CD4+YFP- WT2 | TGCACCTGCAGTGCAGGGGTTAACCAAGACACCCAGTACTTT | 1 | TRBV1*00 | | TRBJ2-5*00 | 0.35% |
| CD4+YFP- WT2 | TGCACCTGCAGTGCAGGGGGAGGCTATGCTGAGCAGTTCTTC | 1 | TRBV1*00 | | TRBJ2-1*00 | 0.35% |
| CD4+YFP- WT2 | TGCACCTGCAGTGCAGATCGGGGGCGGGCTGAGCAGTTCTTC | 1 | TRBV1*00 | TRBD1*00 | TRBJ2-1*00 | 0.35% |
| CD4+YFP- WT2 | TGTGCTAGCAGTTTTTCAACAAACTCCGACTACACCTTC | 1 | TRBV29*00 | | TRBJ1-2*00 | 0.35% |
| CD4+YFP- WT2 | TGTGCTAGCAGCCGACAGGGGGCAGAAACGCTGTATTTT | 1 | TRBV3*00 | TRBD1*00 | TRBJ2-3*00 | 0.35% |
| CD4+YFP- WT2 | TGTGCTAGCAGCCACAGGGGAGGCTCCGACTACACCTTC | 1 | TRBV17*00 | TRBD1*00 | TRBJ1-2*00 | 0.35% |
| CD4+YFP- WT2 | TGTGCCTGGGGGGGGACAGGGGGATCCGACTACACCTTC | 1 | TRBV31*00 | TRBD1*00 | TRBJ1-2*00 | 0.35% |
| CD4+YFP- WT2 | TGTGCCTGGACTGGAAGGAGCTCCTATGAACAGTACTTC | 1 | TRBV31*00 | | TRBJ2-7*00 | 0.35% |
| CD4+YFP- WT2 | TGTGCCTGGAGTCTTATTTCCAACGAAAGATTATTTTTC | 1 | TRBV31*00 | | TRBJ1-4*00 | 0.35% |
| CD4+YFP- WT2 | TGTGCCTGGAGTCGGACCGGTGCAGAAACGCTGTATTTT | 1 | TRBV31*00 | | TRBJ2-3*00 | 0.35% |
| CD4+YFP- WT2 | TGTGCCAGCTCTCTCGATCCAAACTCCGACTACACCTTC | 1 | TRBV12-1*00 | | TRBJ1-2*00 | 0.35% |
| CD4+YFP- WT2 | TGTGCCAGCGGTGATGTGGGGGCCGGGCAGCTCTACTTT | 1 | TRBV13-2*00 | TRBD2*00 | TRBJ2-2*00 | 0.35% |
| CD4+YFP- WT2 | TGTGCCAGCGGTGATGCTGGGACAGAAACGCTGTATTTT | 1 | TRBV13-2*00 | | TRBJ2-3*00 | 0.35% |
| CD4+YFP- WT2 | TGTGCCAGCGGTGATGGGGGGGCAAACACCTTGTACTTT | 1 | TRBV13-2*00 | TRBD2*00 | TRBJ2-4*00 | 0.35% |
| CD4+YFP- WT2 | TGTGCCAGCGGTGAAGACTGGGGGGACACCCAGTACTTT | 1 | TRBV13-2*00 | TRBD2*00 | TRBJ2-5*00 | 0.35% |
| CD4+YFP- WT2 | TGTGCCAGCAGTTTTTACAACTATGCTGAGCAGTTCTTC | 1 | TRBV14*00 | | TRBJ2-1*00 | 0.35% |
| CD4+YFP- WT2 | TGTGCCAGCAGTTTAGCGACAACAGAAACGCTGTATTTT | 1 | TRBV15*00 | | TRBJ2-3*00 | 0.35% |
| CD4+YFP- WT2 | TGTGCCAGCAGTTTAGCAGGGGAAGACACCCAGTACTTT | 1 | TRBV14*00 | TRBD1*00 | TRBJ2-5*00 | 0.35% |
| CD4+YFP- WT2 | TGTGCCAGCAGTTTAGAGGGCAACCAGGCTCCGCTTTTT | 1 | TRBV15*00 | | TRBJ1-5*00 | 0.35% |
| CD4+YFP- WT2 | TGTGCCAGCAGTCTTCAGTACCAAGACACCCAGTACTTT | 1 | TRBV26*00 | | TRBJ2-5*00 | 0.35% |
| CD4+YFP- WT2 | TGTGCCAGCAGTCTCCCAGGGGATTCGCCCCTCTACTTT | 1 | TRBV26*00 | TRBD1*00 | TRBJ1-6*00 | 0.35% |
| CD4+YFP- WT2 | TGTGCCAGCAGTCTGGGTGGCCAAGACACCCAGTACTTT | 1 | TRBV24*00 | | TRBJ2-5*00 | 0.35% |
| CD4+YFP- WT2 | TGTGCCAGCAGTCTGGACAGACATGCTGAGCAGTTCTTC | 1 | TRBV26*00 | TRBD1*00 | TRBJ2-1*00 | 0.35% |
| CD4+YFP- WT2 | TGTGCCAGCAGTCCCAATTCTGGAAATACGCTCTATTTT | 1 | TRBV26*00 | | TRBJ1-3*00 | 0.35% |
| CD4+YFP- WT2 | TGTGCCAGCAGTCCGGGACAGGGGGCTGAGCAGTTCTTC | 1 | TRBV14*00 | TRBD1*00 | TRBJ2-1*00 | 0.35% |
| CD4+YFP- WT2 | TGTGCCAGCAGTCGGGGGTACTATGCTGAGCAGTTCTTC | 1 | TRBV13-3*00 | TRBD1*00 | TRBJ2-1*00 | 0.35% |
| CD4+YFP- WT2 | TGTGCCAGCAGTCGGACAGGGGGCCCCGACTACACCTTC | 1 | TRBV24*00 | TRBD1*00 | TRBJ1-2*00 | 0.35% |
| CD4+YFP- WT2 | TGTGCCAGCAGTGTCGCAGGGGAAAACACCTTGTACTTT | 1 | TRBV13-1*00 | TRBD1*00 | TRBJ2-4*00 | 0.35% |
| CD4+YFP- WT2 | TGTGCCAGCAGTGTCACAGATAACTCCGACTACACCTTC | 1 | TRBV13-1*00 | | TRBJ1-2*00 | 0.35% |
| CD4+YFP- WT2 | TGTGCCAGCAGTGTCAGGGTGTACTCCGACTACACCTTC | 1 | TRBV13-3*00 | TRBD1*00 | TRBJ1-2*00 | 0.35% |
| CD4+YFP- WT2 | TGTGCCAGCAGTGGGTTTAACCAAGACACCCAGTACTTT | 1 | TRBV5*00 | | TRBJ2-5*00 | 0.35% |
| CD4+YFP- WT2 | TGTGCCAGCAGTGGGGGGACAGGCTCCGACTACACCTTC | 1 | TRBV13-1*00 | TRBD1*00 | TRBJ1-2*00 | 0.35% |
| CD4+YFP- WT2 | TGTGCCAGCAGTGGGGAAAACTATGCTGAGCAGTTCTTC | 1 | TRBV13-1*00 | | TRBJ2-1*00 | 0.35% |
| CD4+YFP- WT2 | TGTGCCAGCAGTGGACAGAGTCAAAACACCTTGTACTTT | 1 | TRBV13-1*00 | TRBD1*00 | TRBJ2-4*00 | 0.35% |
| CD4+YFP- WT2 | TGTGCCAGCAGTGATCTCCGGAACCAGGCTCCGCTTTTT | 1 | TRBV13-1*00 | | TRBJ1-5*00 | 0.35% |
| CD4+YFP- WT2 | TGTGCCAGCAGTGATGGACAGGGTGACACCCAGTACTTT | 1 | TRBV13-1*00 | TRBD1*00 | TRBJ2-5*00 | 0.35% |
| CD4+YFP- WT2 | TGTGCCAGCAGTGATGATAACAACCAGGCTCCGCTTTTT | 1 | TRBV13-3*00 | | TRBJ1-5*00 | 0.35% |
| CD4+YFP- WT2 | TGTGCCAGCAGTGACAGGGGGGACTCCGACTACACCTTC | 1 | TRBV13-3*00 | TRBD1*00 | TRBJ1-2*00 | 0.35% |
| CD4+YFP- WT2 | TGTGCCAGCAGTGAACGGGGCGCGGACACCCAGTACTTT | 1 | TRBV13-3*00 | TRBD1*00 | TRBJ2-5*00 | 0.35% |
| CD4+YFP- WT2 | TGTGCCAGCAGTGAACAGGGGGGCGGCACCCAGTACTTT | 1 | TRBV13-1*00 | TRBD1*00 | TRBJ2-5*00 | 0.35% |
| CD4+YFP- WT2 | TGTGCCAGCAGTATAGAGGGCCGCACAGAAGTCTTCTTT | 1 | TRBV19*00 | | TRBJ1-1*00 | 0.35% |
| CD4+YFP- WT2 | TGTGCCAGCAGTAGGGACAGCACCGGGCAGCTCTACTTT | 1 | TRBV19*00 | TRBD1*00 | TRBJ2-2*00 | 0.35% |
| CD4+YFP- WT2 | TGTGCCAGCAGCTTTGGACAGAACTCCGACTACACCTTC | 1 | TRBV3*00 | TRBD1*00 | TRBJ1-2*00 | 0.35% |
| CD4+YFP- WT2 | TGTGCCAGCAGCTTTGAGAGCTCCTATGAACAGTACTTC | 1 | TRBV3*00 | | TRBJ2-7*00 | 0.35% |
| CD4+YFP- WT2 | TGTGCCAGCAGCTTTACTGGGAAAGACACCCAGTACTTT | 1 | TRBV3*00 | TRBD2*00 | TRBJ2-5*00 | 0.35% |
| CD4+YFP- WT2 | TGTGCCAGCAGCTTAGCAGGGAACGAAAGATTATTTTTC | 1 | TRBV3*00 | | TRBJ1-4*00 | 0.35% |
| CD4+YFP- WT2 | TGTGCCAGCAGCTTAGATAGGGATGCTGAGCAGTTCTTC | 1 | TRBV3*00 | | TRBJ2-1*00 | 0.35% |
| CD4+YFP- WT2 | TGTGCCAGCAGCTTAGAACATAACTCCGACTACACCTTC | 1 | TRBV3*00 | | TRBJ1-2*00 | 0.35% |
| CD4+YFP- WT2 | TGTGCCAGCAGCTTAAAGAGTCAAAACACCTTGTACTTT | 1 | TRBV3*00 | | TRBJ2-4*00 | 0.35% |
| CD4+YFP- WT2 | TGTGCCAGCAGCTCGACAACAAACTCCGACTACACCTTC | 1 | TRBV25*00 | | TRBJ1-2*00 | 0.35% |
| CD4+YFP- WT2 | TGTGCCAGCAGCTCACTGGGGGGAGACACCCAGTACTTT | 1 | TRBV4*00 | TRBD2*00 | TRBJ2-5*00 | 0.35% |
| CD4+YFP- WT2 | TGTGCCAGCAGCTGGGGGGTCTATGCTGAGCAGTTCTTC | 1 | TRBV25*00 | TRBD2*00 | TRBJ2-1*00 | 0.35% |
| CD4+YFP- WT2 | TGTGCCAGCAGCTATGGGAAAGGGAACACCTTGTACTTT | 1 | TRBV4*00 | TRBD1*00 | TRBJ2-4*00 | 0.35% |
| CD4+YFP- WT2 | TGTGCCAGCAGCTACCAGGATAATTCGCCCCTCTACTTT | 1 | TRBV4*00 | | TRBJ1-6*00 | 0.35% |

| | | | | | | |
|---|---|---|---|---|---|---|
| CD4+YFP- WT2 | TGTGCCAGCAGCCTCGGGACAGATGCTGAGCAGTTCTTC | 1 | TRBV15*00 | TRBD1*00 | TRBJ2-1*00 | 0.35% |
| CD4+YFP- WT2 | TGTGCCAGCAGCCTGGGGGGGCCCTATGAACAGTACTTC | 1 | TRBV15*00 | TRBD2*00 | TRBJ2-7*00 | 0.35% |
| CD4+YFP- WT2 | TGTGCCAGCAGCCCCCGGGCTGGAAATACGCTCTATTTT | 1 | TRBV2*00 | TRBD1*00 | TRBJ1-3*00 | 0.35% |
| CD4+YFP- WT2 | TGTGCCAGCAGCCGGGACAACAACGAAAGATTATTTTTC | 1 | TRBV4*00 | TRBD1*00 | TRBJ1-4*00 | 0.35% |
| CD4+YFP- WT2 | TGTGCCAGCAGCCGACAGGGGGAAGACACCCAGTACTTT | 1 | TRBV2*00 | TRBD1*00 | TRBJ2-5*00 | 0.35% |
| CD4+YFP- WT2 | TGTGCCAGCAGCCAAGATCAGGGGTATGAACAGTACTTC | 1 | TRBV2*00 | TRBD1*00 | TRBJ2-7*00 | 0.35% |
| CD4+YFP- WT2 | TGTGCCAGCAGCCAAGATACACAGACAGAAGTCTTCTTT | 1 | TRBV2*00 | | TRBJ1-1*00 | 0.35% |
| CD4+YFP- WT2 | TGTGCCAGCAGCGGGACAGCCAACGAAAGATTATTTTTC | 1 | TRBV13-1*00 | TRBD1*00 | TRBJ1-4*00 | 0.35% |
| CD4+YFP- WT2 | TGTGCCAGCAGCGGACTGGGGCAAAACACCTTGTACTTT | 1 | TRBV3*00 | TRBD2*00 | TRBJ2-4*00 | 0.35% |
| CD4+YFP- WT2 | TGTGCCAGCAGCAATGGGGACTCCTATGAACAGTACTTC | 1 | TRBV3*00 | TRBD2*00 | TRBJ2-7*00 | 0.35% |
| CD4+YFP- WT2 | TGTGCCAGCAGGGGACAGGGCTATGCTGAGCAGTTCTTC | 1 | TRBV13-1*00 | TRBD1*00 | TRBJ2-1*00 | 0.35% |
| CD4+YFP- WT2 | TGTGCCAGCAGGGACACCTATAATTCGCCCCTCTACTTT | 1 | TRBV19*00 | TRBD1*00 | TRBJ1-6*00 | 0.35% |
| CD4+YFP- WT2 | TGTGCCAGCAGGACAGGGAGCTCCTATGAACAGTACTTC | 1 | TRBV15*00 | TRBD1*00 | TRBJ2-7*00 | 0.35% |
| CD4+YFP- WT2 | TGTGCCAGCAGACGGGGAGGCCAAGACACCCAGTACTTT | 1 | TRBV19*00 | | TRBJ2-5*00 | 0.35% |
| CD4+YFP- WT2 | TGTGCCAGCAACCGACAGGTCTATGCTGAGCAGTTCTTC | 1 | TRBV13-3*00 | TRBD1*00 | TRBJ2-1*00 | 0.35% |
| CD4+YFP- WT2 | TGTGCAAGCAGCTTGACAGGGTACACAGAAGTCTTCTTT | 1 | TRBV16*00 | TRBD1*00 | TRBJ1-1*00 | 0.35% |
| CD4+YFP- WT2 | TGTGCAAGCAGCTTAGGGGGGGTTGACACCCAGTACTTT | 1 | TRBV16*00 | TRBD2*00 | TRBJ2-5*00 | 0.35% |
| CD4+YFP- WT2 | TGTGCAAGCAGCTTAACTAGTCAAAACACCTTGTACTTT | 1 | TRBV16*00 | | TRBJ2-4*00 | 0.35% |
| CD4+YFP- WT2 | TGTGCAAGCAGCTGGGGGGGGGCGCCTGAACAGTACTTC | 1 | TRBV16*00 | TRBD2*00 | TRBJ2-7*00 | 0.35% |
| CD4+YFP- WT2 | TGTGCAAGCAGCCCCTCAAACACCGGGCAGCTCTACTTT | 1 | TRBV16*00 | | TRBJ2-2*00 | 0.35% |
| CD4+YFP- WT2 | TGTGCAAGCAGCCGGGACAGAAACACAGAAGTCTTCTTT | 1 | TRBV16*00 | TRBD1*00 | TRBJ1-1*00 | 0.35% |
| CD4+YFP- WT2 | TGTGGTGCTAGTTGGACTAGTGCAGAAACGCTGTATTTT | 1 | TRBV20*00 | | TRBJ2-3*00 | 0.35% |
| CD4+YFP- WT2 | TGCACCTGCAGTGTCCGACAACAAGACACCCAGTACTTT | 1 | TRBV1*00 | | TRBJ2-5*00 | 0.35% |
| CD4+YFP- WT2 | TGCACCTGCAGTGCTCGACTGGGAGAAACGCTGTATTTT | 1 | TRBV1*00 | TRBD2*00 | TRBJ2-3*00 | 0.35% |
| CD4+YFP- WT2 | TGCACCTGCAGTGCTCAGGCAAACTCCGACTACACCTTC | 1 | TRBV1*00 | | TRBJ1-2*00 | 0.35% |
| CD4+YFP- WT2 | TGCACCTGCAGTGCCCTACAGAACTCCGACTACACCTTC | 1 | TRBV1*00 | | TRBJ1-2*00 | 0.35% |
| CD4+YFP- WT2 | TGCACCTGCAGTGCCAGGGCGGGGGCTGAGCAGTTCTTC | 1 | TRBV1*00 | TRBD1*00 | TRBJ2-1*00 | 0.35% |
| CD4+YFP- WT2 | TGCACCTGCAGTGCGTCAGGAAACTCCGACTACACCTTC | 1 | TRBV1*00 | | TRBJ1-2*00 | 0.35% |
| CD4+YFP- WT2 | TGCACCTGCAGTGCAGTGGGCAACCAGGCTCCGCTTTTT | 1 | TRBV1*00 | | TRBJ1-5*00 | 0.35% |
| CD4+YFP- WT2 | TGCACCTGCAGTGCAGGGGACGATGCTGAGCAGTTCTTC | 1 | TRBV1*00 | TRBD1*00 | TRBJ2-1*00 | 0.35% |
| CD4+YFP- WT2 | TGTGCTAGCAGTTTAGGACAATATGAACAGTACTTC | 1 | TRBV29*00 | TRBD1*00 | TRBJ2-7*00 | 0.35% |
| CD4+YFP- WT2 | TGTGCTAGCAGTACAGCAAACTCCGACTACACCTTC | 1 | TRBV21*00 | | TRBJ1-2*00 | 0.35% |
| CD4+YFP- WT2 | TGTGCTAGCAGTAGTGGACAGCAGGCTCCGCTTTTT | 1 | TRBV17*00 | TRBD1*00 | TRBJ1-5*00 | 0.35% |
| CD4+YFP- WT2 | TGTGCCTGGAGTCTTAGACAGTATGAACAGTACTTC | 1 | TRBV31*00 | TRBD1*00 | TRBJ2-7*00 | 0.35% |
| CD4+YFP- WT2 | TGTGCCTGGAGTCTAGGGTGGGAAAGATTATTTTTC | 1 | TRBV31*00 | | TRBJ1-4*00 | 0.35% |
| CD4+YFP- WT2 | TGTGCCTGGAGTCTAGAGGGGGATACGCTCTATTTT | 1 | TRBV31*00 | TRBD1*00 | TRBJ1-3*00 | 0.35% |
| CD4+YFP- WT2 | TGTGCCTGGAGCCATTCAAACTCCGACTACACCTTC | 1 | TRBV31*00 | | TRBJ1-2*00 | 0.35% |
| CD4+YFP- WT2 | TGTGCCACCGGGACTGGGTTTGAAACGCTGTATTTT | 1 | TRBV13-3*00 | TRBD2*00 | TRBJ2-3*00 | 0.35% |
| CD4+YFP- WT2 | TGTGCCACGTCGGCAGGGGGGCAGGAACAGTACTTC | 1 | TRBV12-2*00 | TRBD1*00 | TRBJ2-7*00 | 0.35% |
| CD4+YFP- WT2 | TGTGCCAGCTCTCTCGTTGACCAGGCTCCGCTTTTT | 1 | TRBV12-1*00 | | TRBJ1-5*00 | 0.35% |
| CD4+YFP- WT2 | TGTGCCAGCTCTCTCGACAATTATGAACAGTACTTC | 1 | TRBV12-1*00 | | TRBJ2-7*00 | 0.35% |
| CD4+YFP- WT2 | TGTGCCAGCTCTCTCGAGTACTCCGACTACACCTTC | 1 | TRBV12-1*00 | | TRBJ1-2*00 | 0.35% |
| CD4+YFP- WT2 | TGTGCCAGCTCTCGAGACAACTATGAACAGTACTTC | 1 | TRBV18*00 | | TRBJ2-7*00 | 0.35% |
| CD4+YFP- WT2 | TGTGCCAGCGGTGGCAGGGCCTCCGACTACACCTTC | 1 | TRBV13-2*00 | TRBD1*00 | TRBJ1-2*00 | 0.35% |
| CD4+YFP- WT2 | TGTGCCAGCGGTGATGAAAGGGACACCTTGTACTTT | 1 | TRBV13-2*00 | | TRBJ2-4*00 | 0.35% |
| CD4+YFP- WT2 | TGTGCCAGCAGTTTTGGGACCCAGGCTCCGCTTTTT | 1 | TRBV15*00 | TRBD1*00 | TRBJ1-5*00 | 0.35% |
| CD4+YFP- WT2 | TGTGCCAGCAGTTTTAACCAAGACACCCAGTACTTT | 1 | TRBV14*00 | | TRBJ2-5*00 | 0.35% |
| CD4+YFP- WT2 | TGTGCCAGCAGTCTCGGTCCCTATGAACAGTACTTC | 1 | TRBV19*00 | | TRBJ2-7*00 | 0.35% |
| CD4+YFP- WT2 | TGTGCCAGCAGTCTCGGCAGGGGCGAACAGTACTTC | 1 | TRBV26*00 | TRBD1*00 | TRBJ2-7*00 | 0.35% |
| CD4+YFP- WT2 | TGTGCCAGCAGTCTGGGGGGAGGGGAACAGTACTTC | 1 | TRBV26*00 | TRBD1*00 | TRBJ2-7*00 | 0.35% |
| CD4+YFP- WT2 | TGTGCCAGCAGTGATCCAACCTATGAACAGTACTTC | 1 | TRBV13-3*00 | | TRBJ2-7*00 | 0.35% |
| CD4+YFP- WT2 | TGTGCCAGCAGTGATAACAACCAGGCTCCGCTTTTT | 1 | TRBV13-1*00 | | TRBJ1-5*00 | 0.35% |
| CD4+YFP- WT2 | TGTGCCAGCAGTGACAACACCGGGCAGCTCTACTTT | 1 | TRBV13-3*00 | | TRBJ2-2*00 | 0.35% |
| CD4+YFP- WT2 | TGTGCCAGCAGTGAGAGGGGCTATGAACAGTACTTC | 1 | TRBV13-1*00 | TRBD1*00 | TRBJ2-7*00 | 0.35% |
| CD4+YFP- WT2 | TGTGCCAGCAGTGAAGCAGGGGAAAGATTATTTTTC | 1 | TRBV13-1*00 | | TRBJ1-4*00 | 0.35% |
| CD4+YFP- WT2 | TGTGCCAGCAGCTTACAACAAAACACCTTGTACTTT | 1 | TRBV3*00 | | TRBJ2-4*00 | 0.35% |
| CD4+YFP- WT2 | TGTGCCAGCAGCTTAGGGGGTGAAAGATTATTTTTC | 1 | TRBV3*00 | | TRBJ1-4*00 | 0.35% |
| CD4+YFP- WT2 | TGTGCCAGCAGCTTAGACAACACAGAAGTCTTCTTT | 1 | TRBV3*00 | | TRBJ1-1*00 | 0.35% |
| CD4+YFP- WT2 | TGTGCCAGCAGCTGGGACAGGAGCACCTTGTACTTT | 1 | TRBV4*00 | TRBD1*00 | TRBJ2-4*00 | 0.35% |
| CD4+YFP- WT2 | TGTGCCAGCAGCCCGGACATCTATGAACAGTACTTC | 1 | TRBV4*00 | TRBD1*00 | TRBJ2-7*00 | 0.35% |
| CD4+YFP- WT2 | TGTGCCAGCAGCCGAACAAACACAGAAGTCTTCTTT | 1 | TRBV4*00 | | TRBJ1-1*00 | 0.35% |
| CD4+YFP- WT2 | TGTGCCAGCAGCGGGACAGCAGAAACGCTGTATTTT | 1 | TRBV15*00 | TRBD1*00 | TRBJ2-3*00 | 0.35% |
| CD4+YFP- WT2 | TGTGCCAGCAGGACAGCAAACTCCGACTACACCTTC | 1 | TRBV13-1*00 | | TRBJ1-2*00 | 0.35% |
| CD4+YFP- WT2 | TGTGCCAGCAGGAGACAGCAAGACACCCAGTACTTT | 1 | TRBV19*00 | TRBD1*00 | TRBJ2-5*00 | 0.35% |
| CD4+YFP- WT2 | TGTGCCAGCAGAACGCAGGGGGCTGAGCAGTTCTTC | 1 | TRBV15*00 | TRBD1*00 | TRBJ2-1*00 | 0.35% |
| CD4+YFP- WT2 | TGTGCCAGGACACTTTCTGGAAATACGCTCTATTTT | 1 | TRBV13-3*00 | | TRBJ1-3*00 | 0.35% |
| CD4+YFP- WT2 | TGTGCAAGCAGCTTAGGCCAGTATGAACAGTACTTC | 1 | TRBV16*00 | | TRBJ2-7*00 | 0.35% |
| CD4+YFP- WT2 | TGTGCAAGCAGCTTAGACCAAGACACCCAGTACTTT | 1 | TRBV16*00 | | TRBJ2-5*00 | 0.35% |
| CD4+YFP- WT2 | TGTGCAAGCAGCTCCGGGACAGAAGAAGTCTTCTTT | 1 | TRBV16*00 | TRBD1*00 | TRBJ1-1*00 | 0.35% |
| CD4+YFP- WT2 | TGTGCAAGCAGCCCCCGCAGCACAGAAGTCTTCTTT | 1 | TRBV16*00 | | TRBJ1-1*00 | 0.35% |
| CD4+YFP- WT2 | TGTGGTGCTCAGGGGAACCAAGACACCCAGTACTTT | 1 | TRBV20*00 | TRBD1*00 | TRBJ2-5*00 | 0.35% |
| CD4+YFP- WT2 | TGTGGTGCTAGTAATTCTGGAAATACGCTCCATTTT | 1 | TRBV20*00 | | TRBJ1-3*00 | 0.35% |
| CD4+YFP- WT2 | TGTGGTGCTAGGACTAGTCAAAACACCTTGTACTTT | 1 | TRBV20*00 | | TRBJ2-4*00 | 0.35% |
| CD4+YFP- WT2 | TGCTCCAGCAGTCAATCGGGGGGCACCCAGTACTTT | 1 | TRBV23*00 | TRBD1*00 | TRBJ2-5*00 | 0.35% |

| | | | | | | |
|---|---|---|---|---|---|---|
| CD4+YFP- WT2 | TGCACCTGCAGTGCAGGTGGGTATGAACAGTACTTC | 1 | TRBV1*00 | | TRBJ2-7*00 | 0.35% |
| CD4+YFP- WT2 | TGCACCTGCAGTGCAGGAGAGGGAGAAGTCTTCTTT | 1 | TRBV1*00 | | TRBJ1-1*00 | 0.35% |
| CD4+YFP- WT2 | TGCACCTGCAGTGCAGATCTCTATGAACAGTACTTC | 1 | TRBV1*00 | | TRBJ2-7*00 | 0.35% |
| CD4+YFP- WT2 | TGCACCTGCAGTGCAGAAGGGTGGGAACAGTACTTC | 1 | TRBV1*00 | | TRBJ2-7*00 | 0.35% |
| CD4+YFP- WT2 | TGTGCCAGCAGTGAGAACAAGACACCCAGTACTTT | 1 | TRBV13-1*00 | | TRBJ2-5*00 | 0.35% |
| CD4+YFP- WT2 | TGTGCCTGGAGTAGGAAAGACACCCAGTACTTT | 1 | TRBV31*00 | | TRBJ2-5*00 | 0.35% |
| CD4+YFP- WT2 | TGTGCCAGCTCTTACTCCTATGAACAGTACTTC | 1 | TRBV12-1*00 | | TRBJ2-7*00 | 0.35% |
| CD4+YFP- WT2 | TGTGCCAGCAGTCTGGGAAACACCCAGTACTTT | 1 | TRBV24*00 | | TRBJ2-5*00 | 0.35% |
| CD4+YFP- WT2 | TGTGCCAGCAGTCCCGGACAGGGGCAGTACTTC | 1 | TRBV24*00 | TRBD1*00 | TRBJ2-7*00 | 0.35% |
| CD4+YFP- WT2 | TGTGCCAGCAGTACAGGGGCAGAAGTCTTCTTT | 1 | TRBV13-3*00 | TRBD1*00 | TRBJ1-1*00 | 0.35% |
| CD4+YFP- WT2 | TGTGCCAGCAGTAGACAGTATGAACAGTACTTC | 1 | TRBV15*00 | TRBD1*00 | TRBJ2-7*00 | 0.35% |
| CD4+YFP- WT2 | TGTGCCAGCAGCTTAGACACAGAAGTCTTCTTT | 1 | TRBV3*00 | | TRBJ1-1*00 | 0.35% |
| CD4+YFP- WT2 | TGTGCCAGCAGCTCCGGGGCTGAGCAGTTCTTC | 1 | TRBV3*00 | TRBD1*00 | TRBJ2-1*00 | 0.35% |
| CD4+YFP- WT2 | TGTGCCAGCAGGGAACAAACAGAAGTCTTCTTT | 1 | TRBV13-1*00 | | TRBJ1-1*00 | 0.35% |
| CD4+YFP- WT2 | TGTGCAAGCAGCCTGGGGGGTGAACAGTACTTC | 1 | TRBV16*00 | TRBD2*00 | TRBJ2-7*00 | 0.35% |
| CD4+YFP- WT2 | TGTGGTGCTAGGGGGGGGGAATGAACAGTACTTC | 1 | TRBV20*00 | TRBD1*00 | TRBJ2-7*00 | 0.35% |
| CD4+YFP- WT2 | TGTAGTTCTCGCAGTTCCTATGAACAGTACTTC | 1 | TRBV30*00 | | TRBJ2-7*00 | 0.35% |
| CD4+YFP- WT2 | TGTGCCAGCACTGGCTATGAACAGTACTTC | 1 | TRBV13-1*00 | | TRBJ2-7*00 | 0.35% |
| CD4+YFP- WT2 | TGTGCCAGCAGCCACCGCACCTTGTACTTT | 1 | TRBV19*00 | | TRBJ2-4*00 | 0.35% |
| CD4+YFP- WT2 | TGTGGTGCTAGGGGAGACACCCAGTACTTT | 1 | TRBV20*00 | | TRBJ2-5*00 | 0.35% |
| CD4+YFP- WT2 | TGCTCCAGCAGTCACCAAGAAGTCTTCTTT | 1 | TRBV23*00 | | TRBJ1-1*00 | 0.35% |
| CD4+YFP- WT2 | TGTGCCTTCCGGGACAAGAGCAGTTCTTC | 1 | TRBV4*00 | TRBD1*00 | TRBJ2-1*00 | 0.35% |

#### CD4+YFP- WT3

| Sample name | CDR3 Sequence | clone Count | best V Hit | best D Hit | best J Hit | Frequency |
|---|---|---|---|---|---|---|
| CD4+YFP- WT3 | TGTGCCAGCAGTATAGCCGGGTTTCCTATAATTCGCCCCTCTACTTT | 2 | TRBV19*00 | | TRBJ1-6*00 | 1.46% |
| CD4+YFP- WT3 | TGTGCCAGCAGCCGGGACAGAAACCAAGACACCCAGTACTTT | 2 | TRBV4*00 | TRBD1*00 | TRBJ2-5*00 | 1.46% |
| CD4+YFP- WT3 | TGCACCTGCAGTGCAGATCACAACACCGGGCAGCTCTACTTT | 2 | TRBV1*00 | | TRBJ2-2*00 | 1.46% |
| CD4+YFP- WT3 | TGCACCTGCAGTGGGACAGGTAGCTCCTATGAACAGTACTTC | 2 | TRBV1*00 | TRBD1*00 | TRBJ2-7*00 | 1.46% |
| CD4+YFP- WT3 | TGTGCTAGCAGTTTGACAGAATTCTATGAACAGTACTTC | 2 | TRBV29*00 | TRBD1*00 | TRBJ2-7*00 | 1.46% |
| CD4+YFP- WT3 | TGTGCCAGCAGCCAACTGGACACCGGGCAGCTCTACTTT | 2 | TRBV2*00 | | TRBJ2-2*00 | 1.46% |
| CD4+YFP- WT3 | TGTGCCAGCAGTGGAAACAAGACACCCAGTACTTT | 2 | TRBV19*00 | | TRBJ2-5*00 | 1.46% |
| CD4+YFP- WT3 | TGTGCCAGCAGAAACACCGGGCAGCTCTACTTT | 2 | TRBV13-1*00 | | TRBJ2-2*00 | 1.46% |
| CD4+YFP- WT3 | TGTGCAAGCAGCTTAAAAGACACCCAGTACTTT | 2 | TRBV16*00 | | TRBJ2-5*00 | 1.46% |
| CD4+YFP- WT3 | TGTGCAAGCAGCTTATCCTGGGGGGGTAGTAACCAAGACACCCAGTACTTT | 1 | TRBV16*00 | TRBD2*00 | TRBJ2-5*00 | 0.73% |
| CD4+YFP- WT3 | TGTGCTAGCAGTTTAAGCTGGGGGGCAAACCAAGACACCCAGTACTTT | 1 | TRBV29*00 | TRBD2*00 | TRBJ2-5*00 | 0.73% |
| CD4+YFP- WT3 | TGTGCCAGCGGTGATCGGGACAGGGGTTCTGGAAATACGCTCTATTTT | 1 | TRBV13-2*00 | TRBD1*00 | TRBJ1-3*00 | 0.73% |
| CD4+YFP- WT3 | TGTGCCAGCAGTCCACAGGACTGGGGAAGTGCAGAAACGCTGTATTTT | 1 | TRBV26*00 | TRBD2*00 | TRBJ2-3*00 | 0.73% |
| CD4+YFP- WT3 | TGTGCCAGCAGTGAAGGGACTGGGGGTAGTGCAGAAACGCTGTATTTT | 1 | TRBV13-3*00 | TRBD2*00 | TRBJ2-3*00 | 0.73% |
| CD4+YFP- WT3 | TGTGCCAGCGGGGACAGGGGTTCCTATAATTCGCCCCTCTACTTT | 1 | TRBV13-2*00 | TRBD1*00 | TRBJ1-6*00 | 0.73% |
| CD4+YFP- WT3 | TGTGCCAGCAGTTCGACAGGGTATAACTATGCTGAGCAGTTCTTC | 1 | TRBV15*00 | TRBD1*00 | TRBJ2-1*00 | 0.73% |
| CD4+YFP- WT3 | TGTGCCAGCAGTCTGTCGGGAGGGGGCCAAGACACCCAGTACTTT | 1 | TRBV26*00 | TRBD1*00 | TRBJ2-5*00 | 0.73% |
| CD4+YFP- WT3 | TGTGCCAGCAGTCCCTCCCGGGACAGTCAAAACACCTTGTACTTT | 1 | TRBV14*00 | TRBD1*00 | TRBJ2-4*00 | 0.73% |
| CD4+YFP- WT3 | TGTGCCAGCAGTCCCGGGACAGGGGCTGGAAATACGCTCTATTTT | 1 | TRBV14*00 | TRBD1*00 | TRBJ1-3*00 | 0.73% |
| CD4+YFP- WT3 | TGTGCCAGCAGTGATGAGACAGATTCTGGAAATACGCTCTATTTT | 1 | TRBV13-1*00 | TRBD1*00 | TRBJ1-3*00 | 0.73% |
| CD4+YFP- WT3 | TGTGCCAGCAGTATTCCCCCCAATAGTGCAGAAACGCTGTATTTT | 1 | TRBV19*00 | TRBD2*00 | TRBJ2-3*00 | 0.73% |
| CD4+YFP- WT3 | TGTGCCAGCAGTATAGGAGGGGGTAACTATGCTGAGCAGTTCTTC | 1 | TRBV19*00 | TRBD1*00 | TRBJ2-1*00 | 0.73% |
| CD4+YFP- WT3 | TGTGCCAGCAGCTTAGAGGGGCCCGGTGCAGAAACGCTGTATTTT | 1 | TRBV3*00 | TRBD1*00 | TRBJ2-3*00 | 0.73% |
| CD4+YFP- WT3 | TGTGCCAGCAGCCAAGCCGGGACAGCAAACTCCGACTACACCTTC | 1 | TRBV2*00 | TRBD1*00 | TRBJ1-2*00 | 0.73% |
| CD4+YFP- WT3 | TGTGCCAGCAGCCAAGAGGGGGGCTACACCGGGCAGCTCTACTTT | 1 | TRBV2*00 | TRBD2*00 | TRBJ2-2*00 | 0.73% |
| CD4+YFP- WT3 | TGTGCCAGCAGCCAAGAGAGCGTTAACCAAGACACCCAGTACTTT | 1 | TRBV5*00 | | TRBJ2-5*00 | 0.73% |
| CD4+YFP- WT3 | TGTGCAAGCAGCTTAGTTGGGGTCAGTGCAGAAACGCTGTATTTT | 1 | TRBV16*00 | TRBD2*00 | TRBJ2-3*00 | 0.73% |
| CD4+YFP- WT3 | TGTGCAAGCAGCTTAGTGGGGCAGAACTATGCTGAGCAGTTCTTC | 1 | TRBV16*00 | TRBD1*00 | TRBJ2-1*00 | 0.73% |
| CD4+YFP- WT3 | TGCACCTGCAGTGCCCCCGAAAATAACCAAGACACCCAGTACTTT | 1 | TRBV1*00 | | TRBJ2-5*00 | 0.73% |
| CD4+YFP- WT3 | TGTGCTAGCAGTCCCTACAATAACCAAGACACCCAGTACTTT | 1 | TRBV17*00 | | TRBJ2-5*00 | 0.73% |
| CD4+YFP- WT3 | TGTGCCAGCGGTGGGAGACAGGGGAAAGACACCCAGTACTTT | 1 | TRBV13-2*00 | TRBD1*00 | TRBJ2-5*00 | 0.73% |
| CD4+YFP- WT3 | TGTGCCAGCGGTGACCCGGGGGGCTCCTATGAACAGTACTTC | 1 | TRBV13-2*00 | TRBD1*00 | TRBJ2-7*00 | 0.73% |
| CD4+YFP- WT3 | TGTGCCAGCGGTGAGGGTACAAACACCGGGCAGCTCTACTTT | 1 | TRBV13-2*00 | | TRBJ2-2*00 | 0.73% |
| CD4+YFP- WT3 | TGTGCCAGCAGTTTGGAGGACAGGAAGGCTGAGCAGTTCTTC | 1 | TRBV14*00 | TRBD1*00 | TRBJ2-1*00 | 0.73% |
| CD4+YFP- WT3 | TGTGCCAGCAGTTCGACAGCAAACACCGGGCAGCTCTACTTT | 1 | TRBV14*00 | | TRBJ2-2*00 | 0.73% |
| CD4+YFP- WT3 | TGTGCCAGCAGTCCGGGCCCTTCAAACACAGAAGTCTTCTTT | 1 | TRBV14*00 | TRBD1*00 | TRBJ1-1*00 | 0.73% |
| CD4+YFP- WT3 | TGTGCCAGCAGTGCGGGTCATTCTGGAAATACGCTCTATTTT | 1 | TRBV13-1*00 | | TRBJ1-3*00 | 0.73% |
| CD4+YFP- WT3 | TGTGCCAGCAGTGGGACTGGGGGGAGGGACACCCAGTACTTT | 1 | TRBV13-3*00 | TRBD2*00 | TRBJ2-5*00 | 0.73% |
| CD4+YFP- WT3 | TGTGCCAGCAGTGGACTGGGGAGTGCAGAAACGCTGTATTTT | 1 | TRBV13-1*00 | TRBD2*00 | TRBJ2-3*00 | 0.73% |
| CD4+YFP- WT3 | TGTGCCAGCAGTGATACCGCAAACACCGGGCAGCTCTACTTT | 1 | TRBV13-1*00 | | TRBJ2-2*00 | 0.73% |
| CD4+YFP- WT3 | TGTGCCAGCAGTGAGCGACAGAACTATGCTGAGCAGTTCTTC | 1 | TRBV13-3*00 | TRBD1*00 | TRBJ2-1*00 | 0.73% |
| CD4+YFP- WT3 | TGTGCCAGCAGTGAGGGGGACAGGGGGGCTGAGCAGTTCTTC | 1 | TRBV13-3*00 | TRBD1*00 | TRBJ2-1*00 | 0.73% |
| CD4+YFP- WT3 | TGTGCCAGCAGTATGAGGGGGGGTGCAGAAACGCTGTATTTT | 1 | TRBV19*00 | TRBD1*00 | TRBJ2-3*00 | 0.73% |
| CD4+YFP- WT3 | TGTGCCAGCAGTATAGCGCGGGGGGATGCTGAGCAGTTCTTC | 1 | TRBV19*00 | TRBD2*00 | TRBJ2-1*00 | 0.73% |
| CD4+YFP- WT3 | TGTGCCAGCAGTAAGGACTGGGCTAATGCTGAGCAGTTCTTC | 1 | TRBV19*00 | TRBD2*00 | TRBJ2-1*00 | 0.73% |
| CD4+YFP- WT3 | TGTGCCAGCAGCTTAGGGACAGCCCGGGACACCCAGTACTTT | 1 | TRBV3*00 | TRBD1*00 | TRBJ2-5*00 | 0.73% |
| CD4+YFP- WT3 | TGTGCCAGCAGCTCGGACTGGGGGCAAGACACCCAGTACTTT | 1 | TRBV25*00 | TRBD2*00 | TRBJ2-5*00 | 0.73% |
| CD4+YFP- WT3 | TGTGCCAGCAGCTCGACAGGGCACTCCTATGAACAGTACTTC | 1 | TRBV25*00 | TRBD1*00 | TRBJ2-7*00 | 0.73% |
| CD4+YFP- WT3 | TGTGCCAGCAGCCCTCGGCCTGGGGAAAACACCTTGTACTTT | 1 | TRBV4*00 | TRBD2*00 | TRBJ2-4*00 | 0.73% |

| | | | | | | |
|---|---|---|---|---|---|---|
| CD4+YFP- WT3 | TGTGCCAGCAGCCCCTCAGGGAGGTCCTATGAACAGTACTTC | 1 | TRBV15*00 | TRBD1*00 | TRBJ2-7*00 | 0.73% |
| CD4+YFP- WT3 | TGTGCAAGCAGCTTACCCTCTTCTGGAAATACGCTCTATTTT | 1 | TRBV16*00 | | TRBJ1-3*00 | 0.73% |
| CD4+YFP- WT3 | TGTGCAAGCAGCTTAGGACTGGGGAACTATGAACAGTACTTC | 1 | TRBV16*00 | TRBD2*00 | TRBJ2-7*00 | 0.73% |
| CD4+YFP- WT3 | TGTGCAAGCAGCCCTAAAAATAACTATGCTGAGCAGTTCTTC | 1 | TRBV16*00 | | TRBJ2-1*00 | 0.73% |
| CD4+YFP- WT3 | TGCTCCAGCAGTCAAAACAGTAACTATGCTGAGCAGTTCTTC | 1 | TRBV23*00 | | TRBJ2-1*00 | 0.73% |
| CD4+YFP- WT3 | TGTGCCTGGAGTCCTCAAATGCAAACTCCGACTACACCTTC | 1 | TRBV31*00 | | TRBJ1-2*00 | 0.73% |
| CD4+YFP- WT3 | TGTGCTAGCAGTAGAGATGACAAATATGAACAGTACTTC | 1 | TRBV17*00 | | TRBJ2-7*00 | 0.73% |
| CD4+YFP- WT3 | TGTGCTAGCAGCAGACAGGCAAACACAGAAGTCTTCTTT | 1 | TRBV17*00 | | TRBJ1-1*00 | 0.73% |
| CD4+YFP- WT3 | TGTGCCTGGGCACAGGGGGCAAACTCCGACTACACCTTC | 1 | TRBV31*00 | TRBD1*00 | TRBJ1-2*00 | 0.73% |
| CD4+YFP- WT3 | TGTGCCTGGAGTCCACAGGGGAACTCCGACTACACCTTC | 1 | TRBV31*00 | TRBD1*00 | TRBJ1-2*00 | 0.73% |
| CD4+YFP- WT3 | TGTGCCAGCTCTCTCGATCAGAACTCCGACTACACCTTC | 1 | TRBV12-1*00 | | TRBJ1-2*00 | 0.73% |
| CD4+YFP- WT3 | TGTGCCAGCTCAGGGGGTAACCAAGACACCCAGTACTTT | 1 | TRBV14*00 | TRBD1*00 | TRBJ2-5*00 | 0.73% |
| CD4+YFP- WT3 | TGTGCCAGCGGTGATGCGTGGGAACAGGCTCCGCTTTTT | 1 | TRBV13-2*00 | | TRBJ1-5*00 | 0.73% |
| CD4+YFP- WT3 | TGTGCCAGCGGTGATGCAAAGGGGGACACCCAGTACTTT | 1 | TRBV13-2*00 | TRBD1*00 | TRBJ2-5*00 | 0.73% |
| CD4+YFP- WT3 | TGTGCCAGCGGTGAGACTGGGAAAGACACCCAGTACTTT | 1 | TRBV13-2*00 | TRBD2*00 | TRBJ2-5*00 | 0.73% |
| CD4+YFP- WT3 | TGTGCCAGCACTGGGGGGGTCCAAGACACCCAGTACTTT | 1 | TRBV5*00 | TRBD2*00 | TRBJ2-5*00 | 0.73% |
| CD4+YFP- WT3 | TGTGCCAGCAGTTACGACGGCAACCAGGCTCCGCTTTTT | 1 | TRBV15*00 | | TRBJ1-5*00 | 0.73% |
| CD4+YFP- WT3 | TGTGCCAGCAGTCTCAGGGGTCAAAACACCTTGTACTTT | 1 | TRBV26*00 | TRBD1*00 | TRBJ2-4*00 | 0.73% |
| CD4+YFP- WT3 | TGTGCCAGCAGTCTGTCCGGCACCTATGAACAGTACTTC | 1 | TRBV26*00 | | TRBJ2-7*00 | 0.73% |
| CD4+YFP- WT3 | TGTGCCAGCAGTCTGTCGCGCAGCTATGAACAGTACTTC | 1 | TRBV26*00 | | TRBJ2-7*00 | 0.73% |
| CD4+YFP- WT3 | TGTGCCAGCAGTCGACAGGGCTATGCTGAGCAGTTCTTC | 1 | TRBV13-1*00 | TRBD1*00 | TRBJ2-1*00 | 0.73% |
| CD4+YFP- WT3 | TGTGCCAGCAGTGGTCAAGCAAACTCCGACTACACCTTC | 1 | TRBV13-1*00 | | TRBJ1-2*00 | 0.73% |
| CD4+YFP- WT3 | TGTGCCAGCAGTATACGGGGTAGCACAGAAGTCTTCTTT | 1 | TRBV19*00 | | TRBJ1-1*00 | 0.73% |
| CD4+YFP- WT3 | TGTGCCAGCAGTAGGCCAGACAACCAGGCTCCGCTTTTT | 1 | TRBV15*00 | | TRBJ1-5*00 | 0.73% |
| CD4+YFP- WT3 | TGTGCCAGCAGTAAGGGGGTCTATGCTGAGCAGTTCTTC | 1 | TRBV19*00 | TRBD1*00 | TRBJ2-1*00 | 0.73% |
| CD4+YFP- WT3 | TGTGCCAGCAGCTTCAATTCTGGAAATACGCTCTATTTT | 1 | TRBV3*00 | | TRBJ1-3*00 | 0.73% |
| CD4+YFP- WT3 | TGTGCCAGCAGCTTGACTGGGGGTTATGAACAGTACTTC | 1 | TRBV3*00 | TRBD2*00 | TRBJ2-7*00 | 0.73% |
| CD4+YFP- WT3 | TGTGCCAGCAGCTACAGGGGGGTTGCTGAGCAGTTCTTC | 1 | TRBV4*00 | TRBD1*00 | TRBJ2-1*00 | 0.73% |
| CD4+YFP- WT3 | TGTGCCAGCAGCCGGGACAGAGGGGACACCCAGTACTTT | 1 | TRBV3*00 | TRBD1*00 | TRBJ2-5*00 | 0.73% |
| CD4+YFP- WT3 | TGTGCCAGCAGCCGACTGGAGCAAGACACCCAGTACTTT | 1 | TRBV26*00 | TRBD2*00 | TRBJ2-5*00 | 0.73% |
| CD4+YFP- WT3 | TGTGCCAGCAGCCAAGCCAACTATGCTGAGCAGTTCTTC | 1 | TRBV2*00 | | TRBJ2-1*00 | 0.73% |
| CD4+YFP- WT3 | TGTGCCAGCAGCCAAGATGGGGGGAATGAACAGTACTTC | 1 | TRBV5*00 | TRBD2*00 | TRBJ2-7*00 | 0.73% |
| CD4+YFP- WT3 | TGTGCCAGCAGCCAAGAAAGGAACTCCGACTACACCTTC | 1 | TRBV5*00 | | TRBJ1-2*00 | 0.73% |
| CD4+YFP- WT3 | TGTGCCAGCAGGGCCGGGACAAACTCCGACTACACCTTC | 1 | TRBV13-1*00 | TRBD1*00 | TRBJ1-2*00 | 0.73% |
| CD4+YFP- WT3 | TGTGCCAGCAGGGGAGCAAACACCGGGCAGCTCTACTTT | 1 | TRBV15*00 | | TRBJ2-2*00 | 0.73% |
| CD4+YFP- WT3 | TGTGCCAGCAGATCCTATTCTGGAAATACGCTCTATTTT | 1 | TRBV13-3*00 | | TRBJ1-3*00 | 0.73% |
| CD4+YFP- WT3 | TGTGCAAGCAGCTTGGAAAAAAACACAGAAGTCTTCTTT | 1 | TRBV16*00 | | TRBJ1-1*00 | 0.73% |
| CD4+YFP- WT3 | TGTGCAAGCAGCTTAGGGGGGGTCTATGAACAGTACTTC | 1 | TRBV16*00 | TRBD2*00 | TRBJ2-7*00 | 0.73% |
| CD4+YFP- WT3 | TGTGGTGCTAGTTGGGCTAGTGCAGAAACGCTGTATTTT | 1 | TRBV20*00 | | TRBJ2-3*00 | 0.73% |
| CD4+YFP- WT3 | TGTGGTGCTAGGGGACAGAACTATGCTGAGCAGTTCTTC | 1 | TRBV20*00 | TRBD1*00 | TRBJ2-1*00 | 0.73% |
| CD4+YFP- WT3 | TGCACCTGCAGTGGGGGTTCTGGAAATACGCTCTATTTT | 1 | TRBV1*00 | | TRBJ1-3*00 | 0.73% |
| CD4+YFP- WT3 | TGTGCTAGCAGTTTATCAGATTATGAACAGTACTTC | 1 | TRBV29*00 | | TRBJ2-7*00 | 0.73% |
| CD4+YFP- WT3 | TGTGCTAGCAGCAAGAGACAAAACACCTTGTACTTT | 1 | TRBV29*00 | | TRBJ2-4*00 | 0.73% |
| CD4+YFP- WT3 | TGTGCCTGGACAGTGTTCAACGAAAGATTATTTTTC | 1 | TRBV31*00 | | TRBJ1-4*00 | 0.73% |
| CD4+YFP- WT3 | TGTGCCTGGAGTCTGGGGCGAGACACCCAGTACTTT | 1 | TRBV31*00 | TRBD1*00 | TRBJ2-5*00 | 0.73% |
| CD4+YFP- WT3 | TGTGCCAGCGTGGGGGGGGGCTATGAACAGTACTTC | 1 | TRBV26*00 | TRBD2*00 | TRBJ2-7*00 | 0.73% |
| CD4+YFP- WT3 | TGTGCCAGCGGGACAGCAAACTCCGACTACACCTTC | 1 | TRBV13-2*00 | | TRBJ1-2*00 | 0.73% |
| CD4+YFP- WT3 | TGTGCCAGCAGTCTGGGGATCTATGAACAGTACTTC | 1 | TRBV26*00 | | TRBJ2-7*00 | 0.73% |
| CD4+YFP- WT3 | TGTGCCAGCAGTCGGGACACCGGGCAGCTCTACTTT | 1 | TRBV13-1*00 | | TRBJ2-2*00 | 0.73% |
| CD4+YFP- WT3 | TGTGCCAGCAGTGATACAAACTCCGACTACACCTTC | 1 | TRBV13-1*00 | | TRBJ1-2*00 | 0.73% |
| CD4+YFP- WT3 | TGTGCCAGCAGTGAAAGGGAAGACACCCAGTACTTT | 1 | TRBV13-1*00 | | TRBJ2-5*00 | 0.73% |
| CD4+YFP- WT3 | TGTGCCAGCAGTATTTCAGACCAGGCTCCGCTTTTT | 1 | TRBV19*00 | | TRBJ1-5*00 | 0.73% |
| CD4+YFP- WT3 | TGTGCCAGCAGCTTTAACCAAGACACCCAGTACTTT | 1 | TRBV4*00 | | TRBJ2-5*00 | 0.73% |
| CD4+YFP- WT3 | TGTGCCAGCAGCCCTCAGAACTATGAACAGTACTTC | 1 | TRBV2*00 | | TRBJ2-7*00 | 0.73% |
| CD4+YFP- WT3 | TGTGCCAGCAGCCCCGGGACAGCAGAAGTCTTCTTT | 1 | TRBV4*00 | TRBD1*00 | TRBJ1-1*00 | 0.73% |
| CD4+YFP- WT3 | TGTGCCAGCAGCAGACAGAACTCCGACTACACCTTC | 1 | TRBV26*00 | TRBD1*00 | TRBJ1-2*00 | 0.73% |
| CD4+YFP- WT3 | TGTGCCAGCAGGAATTCTGGAAATACGCTCTATTTT | 1 | TRBV13-1*00 | | TRBJ1-3*00 | 0.73% |
| CD4+YFP- WT3 | TGTGCAAGCAGCTTAGAGATAAACACCTTGTACTTT | 1 | TRBV16*00 | | TRBJ2-4*00 | 0.73% |
| CD4+YFP- WT3 | TGTGGTGCTAGGGCTCAGGGAAACACCTTGTACTTT | 1 | TRBV20*00 | TRBD1*00 | TRBJ2-4*00 | 0.73% |
| CD4+YFP- WT3 | TGCACCTGCAGTGCCCGTAACACAGAAGTCTTCTTT | 1 | TRBV1*00 | | TRBJ1-1*00 | 0.73% |
| CD4+YFP- WT3 | TGCACCTGCAGTGCAGATGGAAATACGCTCTATTTT | 1 | TRBV1*00 | | TRBJ1-3*00 | 0.73% |
| CD4+YFP- WT3 | TGCACCTGCAGTGCAGACCAAGACACCCAGTACTTT | 1 | TRBV1*00 | | TRBJ2-5*00 | 0.73% |
| CD4+YFP- WT3 | TGCACCTGCAGTACAGGGGACCAGGCTCCGCTTTTT | 1 | TRBV1*00 | | TRBJ1-5*00 | 0.73% |
| CD4+YFP- WT3 | TGTGCCTGGAGTCTCAAGGGAGAAGTCTTCTTT | 1 | TRBV31*00 | | TRBJ1-1*00 | 0.73% |
| CD4+YFP- WT3 | TGTGCCAGCTGGGGCAACCAGGCTCCGCTTTTT | 1 | TRBV4*00 | TRBD2*00 | TRBJ1-5*00 | 0.73% |
| CD4+YFP- WT3 | TGTGCCAGCGCTAGAAACTCCGACTACACCTTC | 1 | TRBV13-2*00 | | TRBJ1-2*00 | 0.73% |
| CD4+YFP- WT3 | TGTGCCAGCGGTGACCAAGACACCCAGTACTTT | 1 | TRBV13-2*00 | | TRBJ2-5*00 | 0.73% |
| CD4+YFP- WT3 | TGTGCCAGCAGTGATGCTGTCGGGGCCCCCTTC | 1 | TRBV13-1*00 | TRBD1*00 | TRBJ1-2*00 | 0.73% |
| CD4+YFP- WT3 | TGTGCCAGCAGCTGGGGGGACACCCAGTACTTT | 1 | TRBV4*00 | TRBD1*00 | TRBJ2-5*00 | 0.73% |
| CD4+YFP- WT3 | TGTGCAAGCAGCTTACATTATGAACAGTACTTC | 1 | TRBV16*00 | | TRBJ2-7*00 | 0.73% |
| CD4+YFP- WT3 | TGTGCAAGCAGCGGGCAGGGGGAAGTCTTCTTT | 1 | TRBV16*00 | TRBD1*00 | TRBJ1-1*00 | 0.73% |
| CD4+YFP- WT3 | TGTAGTTCTTTTAACCAAGACACCCAGTACTTT | 1 | TRBV30*00 | | TRBJ2-5*00 | 0.73% |
| CD4+YFP- WT3 | TGTAGTTCTAGGGACAGGGGGCAGCTCTACTTT | 1 | TRBV30*00 | TRBD1*00 | TRBJ2-2*00 | 0.73% |
| CD4+YFP- WT3 | TGTGCCTGGAGTCTAGGGACGCTGTATTTT | 1 | TRBV31*00 | | TRBJ2-3*00 | 0.73% |

| | | | | | | |
|---|---|---|---|---|---|---|
| CD4+YFP- WT3 | TGTGCCAGCGCCAACGAAAGATTATTTTTC | 1 | TRBV13-3*00 | | TRBJ1-4*00 | 0.73% |
| CD4+YFP- WT3 | TGTGCCAGCAGCCACACAGAAGTCTTCTTT | 1 | TRBV4*00 | | TRBJ1-1*00 | 0.73% |
| CD4+YFP- WT3 | TGTGCCAGCGGTGATGAACAGTACTTC | 1 | TRBV13-2*00 | | TRBJ2-7*00 | 0.73% |
| CD4+YFP- WT3 | TGTGCCAGCGGATATGAACAGTACTTC | 1 | TRBV13-3*00 | | TRBJ2-7*00 | 0.73% |
| CD4+YFP- WT3 | TGTGCCAGCAGTTTGTACTTC | 1 | TRBV13-3*00 | | TRBJ2-7*00 | 0.73% |

#### CD4+YFP- DKO1

| Sample name | CDR3 Sequence | clone Count | best V Hit | best D Hit | best J Hit | Frequency |
|---|---|---|---|---|---|---|
| CD4+YFP- DKO1 | TGTGCCAGCAGGGAGACTGGCTATGCTGAGCAGTTCTTC | 20 | TRBV13-3*00 | TRBD2*00 | TRBJ2-1*00 | 6.97% |
| CD4+YFP- DKO1 | TGTGCCAGCAGCCAGGGGGACACCGGGCAGCTCTACTTT | 7 | TRBV2*00 | TRBD1*00 | TRBJ2-2*00 | 2.44% |
| CD4+YFP- DKO1 | TGTGCTAGCAGTCGACAGGGAAACACCGGGCAGCTCTACTTT | 6 | TRBV21*00 | TRBD1*00 | TRBJ2-2*00 | 2.09% |
| CD4+YFP- DKO1 | TGTGCCAGCTCTCTCGGGGGGGGAATGAACACCCAGTACTTT | 5 | TRBV12-1*00 | TRBD2*00 | TRBJ2-5*00 | 1.74% |
| CD4+YFP- DKO1 | TGTGCCAGCAGCCCCTGGGGGGCCTTTAACCAAGACACCCAGTACTTT | 3 | TRBV2*00 | TRBD2*00 | TRBJ2-5*00 | 1.05% |
| CD4+YFP- DKO1 | TGTGGTGCTCAACAGGGGAACTATGCTGAGCAGTTCTTC | 3 | TRBV20*00 | TRBD1*00 | TRBJ2-1*00 | 1.05% |
| CD4+YFP- DKO1 | TGTGCCAGCGGTGATGCAAGGGGGAGTCAAAACACCTTGTACTTT | 2 | TRBV13-2*00 | TRBD1*00 | TRBJ2-4*00 | 0.70% |
| CD4+YFP- DKO1 | TGCACCTGCAGTGCAGACAGGGTCAACACCGGGCAGCTCTACTTT | 2 | TRBV1*00 | TRBD1*00 | TRBJ2-2*00 | 0.70% |
| CD4+YFP- DKO1 | TGTGCCTGGAGTCTCGGCATCTCCAACGAAAGATTATTTTTC | 2 | TRBV31*00 | | TRBJ1-4*00 | 0.70% |
| CD4+YFP- DKO1 | TGTGCCAGCGGTGATGCAGCCAACTATGCTGAGCAGTTCTTC | 2 | TRBV13-2*00 | | TRBJ2-1*00 | 0.70% |
| CD4+YFP- DKO1 | TGTGCCAGCAGCCAAGACTGGGGGGTCTATGAACAGTACTTC | 2 | TRBV5*00 | TRBD2*00 | TRBJ2-7*00 | 0.70% |
| CD4+YFP- DKO1 | TGTGCCAGCAGTGATGCACGGGAAGACACCCAGTACTTT | 2 | TRBV13-1*00 | | TRBJ2-5*00 | 0.70% |
| CD4+YFP- DKO1 | TGTGCAAGCAGCCGGGACTTCCAAGACACCCAGTACTTT | 2 | TRBV16*00 | TRBD2*00 | TRBJ2-5*00 | 0.70% |
| CD4+YFP- DKO1 | TGTGCAAGCAGACATAGTGCAGAAACGCTGTATTTT | 2 | TRBV16*00 | | TRBJ2-3*00 | 0.70% |
| CD4+YFP- DKO1 | TGCACCTGCAGTGCCAGGGTTGGGAACAGTACTTC | 2 | TRBV1*00 | TRBD1*00 | TRBJ2-7*00 | 0.70% |
| CD4+YFP- DKO1 | CACCTGCAGTGCAGGGGACTGGGGGCCATCTAGTGCAGAAACGCTGTATT | 1 | TRBV1*00 | TRBD2*00 | TRBJ2-3*00 | 0.35% |
| CD4+YFP- DKO1 | GTGCCAGCAGCCAAGGTGGGGGGGCGCGGAGTGCAGAAACGCTGTATTT | 1 | TRBV2*00 | TRBD2*00 | TRBJ2-3*00 | 0.35% |
| CD4+YFP- DKO1 | TGTGCCAGCAGCCAAGATGGGGGGGCAAACACCGGGCAGCTCTACTTT | 1 | TRBV2*00 | TRBD2*00 | TRBJ2-2*00 | 0.35% |
| CD4+YFP- DKO1 | TGTGCAAGCAGCTTAGAGGTGGGGATTAACCAAGACACCCAGTACTTT | 1 | TRBV16*00 | TRBD2*00 | TRBJ2-5*00 | 0.35% |
| CD4+YFP- DKO1 | TGTGCCAGCAGCCCCCGGGACAAATAACTATGCTGAGCAGTTCTTC | 1 | TRBV5*00 | TRBD1*00 | TRBJ2-1*00 | 0.35% |
| CD4+YFP- DKO1 | TGTGCCAGCTCTCCCAACAGCAGGGGTGCAGAAACGCTGTATTTT | 1 | TRBV12-1*00 | TRBD1*00 | TRBJ2-3*00 | 0.35% |
| CD4+YFP- DKO1 | TGTGCCAGCGGTGGTGGGGGGGCGAACTATGCTGAGCAGTTCTTC | 1 | TRBV13-2*00 | TRBD2*00 | TRBJ2-1*00 | 0.35% |
| CD4+YFP- DKO1 | TGTGCCAGCGGTGATGCAGGCACCCACAACCAGGCTCCGCTTTTT | 1 | TRBV13-2*00 | | TRBJ1-5*00 | 0.35% |
| CD4+YFP- DKO1 | TGTGCCAGCGGTGATGGGACTGGGGGGGTGGGAACGCTGTATTTT | 1 | TRBV13-2*00 | TRBD2*00 | TRBJ2-3*00 | 0.35% |
| CD4+YFP- DKO1 | TGTGCCAGCGGTGATAGGGCAACTAGTGCAGAAACGCTGTATTTT | 1 | TRBV13-2*00 | | TRBJ2-3*00 | 0.35% |
| CD4+YFP- DKO1 | TGTGCCAGCAGTTTCGGCAGGGGAAGTGCAGAAACGCTGTATTTT | 1 | TRBV14*00 | TRBD1*00 | TRBJ2-3*00 | 0.35% |
| CD4+YFP- DKO1 | TGTGCCAGCAGTCTGTCAGGACTGGGGGGTAACACCCAGTACTTC | 1 | TRBV26*00 | TRBD2*00 | TRBJ2-5*00 | 0.35% |
| CD4+YFP- DKO1 | TGTGCCAGCAGTCTGGGACAGTTTGCAAACTCCGACTACACCTTC | 1 | TRBV26*00 | | TRBJ1-2*00 | 0.35% |
| CD4+YFP- DKO1 | TGTGCCAGCAGTCTGACTGGGGGGGATTTTGCTGAGCAGTTCTTC | 1 | TRBV24*00 | TRBD2*00 | TRBJ2-1*00 | 0.35% |
| CD4+YFP- DKO1 | TGTGCCAGCAGTGATCGGACAGGGGGCGACGAAAGATTATTTTTC | 1 | TRBV13-1*00 | TRBD1*00 | TRBJ1-4*00 | 0.35% |
| CD4+YFP- DKO1 | TGTGCCAGCAGTGAGAGGGGGCCGAGTCAAAACACCTTGTACTTT | 1 | TRBV13-1*00 | TRBD1*00 | TRBJ2-4*00 | 0.35% |
| CD4+YFP- DKO1 | TGTGCCAGCAGTATCGCCTGGGGGGGTGCAGAAACGCTGTATTTT | 1 | TRBV19*00 | TRBD2*00 | TRBJ2-3*00 | 0.35% |
| CD4+YFP- DKO1 | TGTGCCAGCAGCTTAGCCAGGGGTAACAACCAGGCTCCGCTTTTT | 1 | TRBV3*00 | TRBD1*00 | TRBJ1-5*00 | 0.35% |
| CD4+YFP- DKO1 | TGTGCCAGCAGCTTAGACTGGGGTAGTGCAGAAACGCTGTATTTT | 1 | TRBV3*00 | TRBD2*00 | TRBJ2-3*00 | 0.35% |
| CD4+YFP- DKO1 | TGTGCCAGCAGCTACGGACTGGGGAGCTCCTATGAACAGTACTTC | 1 | TRBV4*00 | TRBD2*00 | TRBJ2-7*00 | 0.35% |
| CD4+YFP- DKO1 | TGTGCCAGCAGCCGGACAGGGGTTAGTGCAGAAACGCTGTATTTT | 1 | TRBV3*00 | TRBD1*00 | TRBJ2-3*00 | 0.35% |
| CD4+YFP- DKO1 | TGTGCCAGCAGCCAGGGGGGGGCTAGTGCAGAAACGCTGTATTTT | 1 | TRBV2*00 | TRBD2*00 | TRBJ2-3*00 | 0.35% |
| CD4+YFP- DKO1 | TGTGCCAGCAGCCAACTGGGGGGGTACCAAGACACCCAGTACTTT | 1 | TRBV2*00 | TRBD2*00 | TRBJ2-5*00 | 0.35% |
| CD4+YFP- DKO1 | TGTGCCAGCAGCCAAGCACAGGGGGCTGGAAATACGCTCTATTTT | 1 | TRBV2*00 | TRBD1*00 | TRBJ1-3*00 | 0.35% |
| CD4+YFP- DKO1 | TGTGCCAGCAGCCAAGGGACAAATAACTATGCTGAGCAGTTCTTC | 1 | TRBV5*00 | TRBD1*00 | TRBJ2-1*00 | 0.35% |
| CD4+YFP- DKO1 | TGTGCCAGCAGCAAGGACAGGGCGAACACCGGGCAGCTCTACTTT | 1 | TRBV3*00 | TRBD1*00 | TRBJ2-2*00 | 0.35% |
| CD4+YFP- DKO1 | TGTGCAAGCAGCTTAGGAGACAGGGGCTATGCTGAGCAGTTCTTC | 1 | TRBV16*00 | TRBD1*00 | TRBJ2-1*00 | 0.35% |
| CD4+YFP- DKO1 | TGTGCAAGCAGCTTAGATTACTGGGGCCAAGACACCCAGTACTTT | 1 | TRBV16*00 | TRBD2*00 | TRBJ2-5*00 | 0.35% |
| CD4+YFP- DKO1 | TGTGCAAGCAGCCCCCTCGGGGGGAACTATGCTGAGCAGTTCTTC | 1 | TRBV16*00 | TRBD2*00 | TRBJ2-1*00 | 0.35% |
| CD4+YFP- DKO1 | TGCACCTGCAGTGCGGGACTGGGGTATTATGCTGAGCAGTTCTTC | 1 | TRBV1*00 | TRBD2*00 | TRBJ2-1*00 | 0.35% |
| CD4+YFP- DKO1 | TGTGCCAGCAGCTAAGAGGCAATTCTGGAAATACGCTCTATTTT | 1 | TRBV4*00 | | TRBJ1-3*00 | 0.35% |
| CD4+YFP- DKO1 | TGTGCTAGCAGTTTGGCAGGGACCAACGAAAGATTATTTTTC | 1 | TRBV29*00 | TRBD1*00 | TRBJ1-4*00 | 0.35% |
| CD4+YFP- DKO1 | TGTGCTAGCAGTTTAGGAGCTTCTGGAAATACGCTCTATTTT | 1 | TRBV29*00 | | TRBJ1-3*00 | 0.35% |
| CD4+YFP- DKO1 | TGTGCTAGCAGTAGATTGGGGGTCCTTGCTGAGCAGTTCTTC | 1 | TRBV17*00 | TRBD2*00 | TRBJ2-1*00 | 0.35% |
| CD4+YFP- DKO1 | TGTGCTAGCAGTAGAGGTGGGGTCACCGGGCAGCTCTACTTT | 1 | TRBV17*00 | TRBD2*00 | TRBJ2-2*00 | 0.35% |
| CD4+YFP- DKO1 | TGTGCTAGCAGCCCGGGACAGGGGAACGAAAGATTATTTTTC | 1 | TRBV21*00 | TRBD1*00 | TRBJ1-4*00 | 0.35% |
| CD4+YFP- DKO1 | TGTGCTAGCAGATCCGGGACAACAAACTCCGACTACACCTTC | 1 | TRBV21*00 | TRBD1*00 | TRBJ1-2*00 | 0.35% |
| CD4+YFP- DKO1 | TGTGCCTGGAGTTCTGGGGTTAGTGCAGAAACGCTGTATTTT | 1 | TRBV31*00 | TRBD2*00 | TRBJ2-3*00 | 0.35% |
| CD4+YFP- DKO1 | TGTGCCTGGAGTCTGACAGTTTCTGGAAATACGCTCTATTTT | 1 | TRBV31*00 | TRBD1*00 | TRBJ1-3*00 | 0.35% |
| CD4+YFP- DKO1 | TGTGCCTGGAGTCGGGGGGGGACCAACTCCGACTACACCTTC | 1 | TRBV31*00 | TRBD2*00 | TRBJ1-2*00 | 0.35% |
| CD4+YFP- DKO1 | TGTGCCTGGAGTCAACAGGGGAACTATGCTGAGCAGTTCTTC | 1 | TRBV31*00 | TRBD1*00 | TRBJ2-1*00 | 0.35% |
| CD4+YFP- DKO1 | TGTGCCTGGAACCTGGGGGGCCTACTGGCTGAGCAGTTCTTC | 1 | TRBV31*00 | TRBD2*00 | TRBJ2-1*00 | 0.35% |
| CD4+YFP- DKO1 | TGTGCCAGCTCTCTCTGGGGGAGTGCAGAAACGCTGTATTTT | 1 | TRBV12-2*00 | TRBD2*00 | TRBJ2-3*00 | 0.35% |
| CD4+YFP- DKO1 | TGTGCCAGCTCTCTCGACTGGGCCACCGGGCAGCTCTACTTT | 1 | TRBV12-1*00 | TRBD2*00 | TRBJ2-2*00 | 0.35% |
| CD4+YFP- DKO1 | TGTGCCAGCTCTCCCACACCAAACACCGGGCAGCTCTACTTT | 1 | TRBV12-1*00 | | TRBJ2-2*00 | 0.35% |
| CD4+YFP- DKO1 | TGTGCCAGCTCCCCGACAGTTAACCAAGACACCCAGTACTTT | 1 | TRBV12-1*00 | | TRBJ2-5*00 | 0.35% |
| CD4+YFP- DKO1 | TGTGCCAGCGGTCCCCCCCTTAACCAAGACACCCAGTACTTT | 1 | TRBV13-2*00 | TRBD2*00 | TRBJ2-5*00 | 0.35% |
| CD4+YFP- DKO1 | TGTGCCAGCGGTGATCTGGGGGGGGCTTATGAACAGTACTTC | 1 | TRBV13-2*00 | TRBD2*00 | TRBJ2-7*00 | 0.35% |
| CD4+YFP- DKO1 | TGTGCCAGCGGTGATCCAGGGGGCAACGAAAGATTATTTTTC | 1 | TRBV13-2*00 | TRBD1*00 | TRBJ1-4*00 | 0.35% |
| CD4+YFP- DKO1 | TGTGCCAGCGGTGATGCTCCAAACACCGGGCAGCTCTACTTT | 1 | TRBV13-2*00 | | TRBJ2-2*00 | 0.35% |
| CD4+YFP- DKO1 | TGTGCCAGCGGTGATGCCGACGGTCAAGACACCCAGTACTTT | 1 | TRBV13-2*00 | | TRBJ2-5*00 | 0.35% |

| | | | | | | |
|---|---|---|---|---|---|---|
| CD4+YFP- DKO1 | TGTGCCAGCGGTGATGAGACTGGAGGGGACACCCAGTACTTT | 1 | TRBV13-2*00 | TRBD2*00 | TRBJ2-5*00 | 0.35% |
| CD4+YFP- DKO1 | TGTGCCAGCGGGGACAGGGGGCTTGGAAATACGCTCTATTTT | 1 | TRBV13-2*00 | TRBD1*00 | TRBJ1-3*00 | 0.35% |
| CD4+YFP- DKO1 | TGTGCCAGCAGTTTCAGGAGGGGGAACACAGAAGTCTTCTTT | 1 | TRBV14*00 | TRBD1*00 | TRBJ1-1*00 | 0.35% |
| CD4+YFP- DKO1 | TGTGCCAGCAGTTTGGGGAATTCTGGAAATACGCCCTATTTT | 1 | TRBV15*00 | | TRBJ1-3*00 | 0.35% |
| CD4+YFP- DKO1 | TGTGCCAGCAGTTTAGACAGTGACACCGGGCAGCTCTACTTT | 1 | TRBV15*00 | | TRBJ2-2*00 | 0.35% |
| CD4+YFP- DKO1 | TGTGCCAGCAGTTCGGGACCTTCTGGAAATACGCTCTATTTT | 1 | TRBV15*00 | TRBD1*00 | TRBJ1-3*00 | 0.35% |
| CD4+YFP- DKO1 | TGTGCCAGCAGTCTCTGGGTTAACCAAGACACCCAGTACTTT | 1 | TRBV14*00 | | TRBJ2-5*00 | 0.35% |
| CD4+YFP- DKO1 | TGTGCCAGCAGTCTGGACACCTACTCCTATGAACAGTACTTC | 1 | TRBV24*00 | | TRBJ2-7*00 | 0.35% |
| CD4+YFP- DKO1 | TGTGCCAGCAGTGATCGGCTGGGCTCCTATGAACAGTACTTC | 1 | TRBV13-3*00 | | TRBJ2-7*00 | 0.35% |
| CD4+YFP- DKO1 | TGTGCCAGCAGTGATCAGGGGGCCTATGCTGAGCAGTTCTTC | 1 | TRBV13-3*00 | TRBD1*00 | TRBJ2-1*00 | 0.35% |
| CD4+YFP- DKO1 | TGTGCCAGCAGTGATGCGGGGGGGCGCATTGAACAGTACTTC | 1 | TRBV13-1*00 | TRBD2*00 | TRBJ2-7*00 | 0.35% |
| CD4+YFP- DKO1 | TGTGCCAGCAGTGATAGGACTGGGGCGTATGAACAGTACTTC | 1 | TRBV13-1*00 | TRBD2*00 | TRBJ2-7*00 | 0.35% |
| CD4+YFP- DKO1 | TGTGCCAGCAGTGAAGGGGCAGGGGCCTATGAACAGTACTTC | 1 | TRBV13-1*00 | TRBD1*00 | TRBJ2-7*00 | 0.35% |
| CD4+YFP- DKO1 | TGTGCCAGCAGTGAAGGGGGGAGTGCAGAAACGCTGTATTTT | 1 | TRBV13-1*00 | TRBD1*00 | TRBJ2-3*00 | 0.35% |
| CD4+YFP- DKO1 | TGTGCCAGCAGTATATCAGGGACCGGCACAGAAGTCTTCTTT | 1 | TRBV19*00 | TRBD1*00 | TRBJ1-1*00 | 0.35% |
| CD4+YFP- DKO1 | TGTGCCAGCAGTACGGGACAGGCAAACACAGAAGTCTTCTTT | 1 | TRBV19*00 | TRBD1*00 | TRBJ1-1*00 | 0.35% |
| CD4+YFP- DKO1 | TGTGCCAGCAGCTTGAAATATAACTATGCTGAGCAGTTCTTC | 1 | TRBV3*00 | | TRBJ2-1*00 | 0.35% |
| CD4+YFP- DKO1 | TGTGCCAGCAGCTTAGACTGGGGGGGTGCTGAGCAGTTCTTC | 1 | TRBV3*00 | TRBD2*00 | TRBJ2-1*00 | 0.35% |
| CD4+YFP- DKO1 | TGTGCCAGCAGCTTAGACTGGGGGGAAGAAACGCTGTATTTT | 1 | TRBV3*00 | TRBD2*00 | TRBJ2-3*00 | 0.35% |
| CD4+YFP- DKO1 | TGTGCCAGCAGCTTAGAGGGAAACACCGGGCAGCTCTACTTT | 1 | TRBV3*00 | | TRBJ2-2*00 | 0.35% |
| CD4+YFP- DKO1 | TGTGCCAGCAGCTTAATAGGGGCAAACTCCGACTACACCTTC | 1 | TRBV3*00 | | TRBJ1-2*00 | 0.35% |
| CD4+YFP- DKO1 | TGTGCCAGCAGCTCGACTGGGCACTATGCTGAGCAGTTCTTC | 1 | TRBV3*00 | TRBD2*00 | TRBJ2-1*00 | 0.35% |
| CD4+YFP- DKO1 | TGTGCCAGCAGCCCCCGGGACTGGGGAAACACCTTGTACTTT | 1 | TRBV2*00 | TRBD2*00 | TRBJ2-4*00 | 0.35% |
| CD4+YFP- DKO1 | TGTGCCAGCAGCCCCACTTTTAACCAAGACACCCAGTACTTT | 1 | TRBV2*00 | | TRBJ2-5*00 | 0.35% |
| CD4+YFP- DKO1 | TGTGCCAGCAGCCATAACAGGGGGATCTCCGACTACACCTTC | 1 | TRBV2*00 | TRBD1*00 | TRBJ1-2*00 | 0.35% |
| CD4+YFP- DKO1 | TGTGCCAGCAGCCAAGATTGGGGAAACACAGAAGTCTTCTTT | 1 | TRBV5*00 | TRBD2*00 | TRBJ1-1*00 | 0.35% |
| CD4+YFP- DKO1 | TGTGCCAGCAGCGCACAGGGGACAAACACAGAAGTCTTCTTT | 1 | TRBV4*00 | TRBD1*00 | TRBJ1-1*00 | 0.35% |
| CD4+YFP- DKO1 | TGTGCCAGCAGCGGGACTTCTAGTCAAAACACCTTGTACTTT | 1 | TRBV3*00 | TRBD2*00 | TRBJ2-4*00 | 0.35% |
| CD4+YFP- DKO1 | TGTGCCAGCAGACTGGGGGTCAACTATGCTGAGCAGTTCTTC | 1 | TRBV15*00 | TRBD2*00 | TRBJ2-1*00 | 0.35% |
| CD4+YFP- DKO1 | TGTGCCAGCAGAGACAGGGGTAGTGCAGAAACGCTGTATTTT | 1 | TRBV19*00 | TRBD1*00 | TRBJ2-3*00 | 0.35% |
| CD4+YFP- DKO1 | TGTGCAAGCAGCTTCTTACTGGGGACGGACACCCAGTACTTT | 1 | TRBV16*00 | TRBD2*00 | TRBJ2-5*00 | 0.35% |
| CD4+YFP- DKO1 | TGTGCAAGCAGCTTAGATCGGAGTCAAAACACCTTGTACTTT | 1 | TRBV16*00 | | TRBJ2-4*00 | 0.35% |
| CD4+YFP- DKO1 | TGTGCAAGCAGCTTAGAACAGAGTGCAGAAACGCTGTATTTT | 1 | TRBV16*00 | | TRBJ2-3*00 | 0.35% |
| CD4+YFP- DKO1 | TGTGCAAGCAGCTTAAAATCAAACACCGGGCAGCTCTACTTT | 1 | TRBV16*00 | | TRBJ2-2*00 | 0.35% |
| CD4+YFP- DKO1 | TGTGCAAGCAGCACCCGGGACAGGGGGCGAGAACAGTACTTC | 1 | TRBV16*00 | TRBD1*00 | TRBJ2-7*00 | 0.35% |
| CD4+YFP- DKO1 | TGCACCTGCAGTGCCCGGGACAGGGGAGACACCCAGTACTTT | 1 | TRBV1*00 | TRBD1*00 | TRBJ2-5*00 | 0.35% |
| CD4+YFP- DKO1 | TGCACCTGCAGTGCACGGGCAAACACCGGGCAGCTCTACTTT | 1 | TRBV1*00 | | TRBJ2-2*00 | 0.35% |
| CD4+YFP- DKO1 | TGCACCTGCAGTGCAGCAGGGACTGGAAATACGCTCTATTTT | 1 | TRBV1*00 | TRBD1*00 | TRBJ1-3*00 | 0.35% |
| CD4+YFP- DKO1 | TGCACCTGCAGTGCAAATGGAAACACCGGGCAGCTCTACTTT | 1 | TRBV1*00 | | TRBJ2-2*00 | 0.35% |
| CD4+YFP- DKO1 | TGCACCTGCAGTAACAGGAATAACAACCAGGCTCCGCCTTTT | 1 | TRBV1*00 | TRBD1*00 | TRBJ1-5*00 | 0.35% |
| CD4+YFP- DKO1 | TGTGCCAGCAGTATAACACAGCAAACTCCGACTACACCTTC | 1 | TRBV19*00 | | TRBJ1-2*00 | 0.35% |
| CD4+YFP- DKO1 | TGTGCCAGCTCTCGGGGGGGGAATGAACACCCAGTACTTT | 1 | TRBV12-2*00 | TRBD2*00 | TRBJ2-5*00 | 0.35% |
| CD4+YFP- DKO1 | TGTGCCAGCAGCCCAACCTAGTGCAGAAACGCTGTATTTT | 1 | TRBV24*00 | | TRBJ2-3*00 | 0.35% |
| CD4+YFP- DKO1 | TGTGCTAGCAGTTTTACAGACTATGCTGAGCAGTTCTTC | 1 | TRBV29*00 | | TRBJ2-1*00 | 0.35% |
| CD4+YFP- DKO1 | TGTGCTAGCAGTCGGACAGGCCAAGACACCCAGTACTTT | 1 | TRBV17*00 | TRBD1*00 | TRBJ2-5*00 | 0.35% |
| CD4+YFP- DKO1 | TGTGCTAGCAGTAACCGGGACAACTATGAACAGTACTTC | 1 | TRBV17*00 | TRBD1*00 | TRBJ2-7*00 | 0.35% |
| CD4+YFP- DKO1 | TGTGCTAGCAGCAGGGGTGGCTATGCTGAGCAGTTCTTC | 1 | TRBV17*00 | TRBD1*00 | TRBJ2-1*00 | 0.35% |
| CD4+YFP- DKO1 | TGTGCCTGGAGTCTAGGGCAAAACTCCGACTACACCTTC | 1 | TRBV31*00 | | TRBJ1-2*00 | 0.35% |
| CD4+YFP- DKO1 | TGTGCCTGGAGCCTGGGGGACCAAGACACCCAGTACTTT | 1 | TRBV31*00 | TRBD2*00 | TRBJ2-5*00 | 0.35% |
| CD4+YFP- DKO1 | TGTGCCTGGAGCCGACTGGGTTATGCTGAGCAGTTCTTC | 1 | TRBV31*00 | | TRBJ2-1*00 | 0.35% |
| CD4+YFP- DKO1 | TGTGCCTGGAAACAGCCTTCTGGAAATACGCTCTATTTT | 1 | TRBV31*00 | | TRBJ1-3*00 | 0.35% |
| CD4+YFP- DKO1 | TGTGCCAGCTCTCTTCGGAACCAAGACACCCAGTACTTT | 1 | TRBV12-1*00 | | TRBJ2-5*00 | 0.35% |
| CD4+YFP- DKO1 | TGTGCCAGCTCTCACAGGGTGGGCTCCGACTACACCTTC | 1 | TRBV12-2*00 | TRBD1*00 | TRBJ1-2*00 | 0.35% |
| CD4+YFP- DKO1 | TGTGCCAGCTCCCCAAATAGTGCAGAAACGCTGTATTTT | 1 | TRBV12-1*00 | | TRBJ2-3*00 | 0.35% |
| CD4+YFP- DKO1 | TGTGCCAGCGGTGGGACAGGCTATGCTGAGCAGTTCTTC | 1 | TRBV13-2*00 | TRBD1*00 | TRBJ2-1*00 | 0.35% |
| CD4+YFP- DKO1 | TGTGCCAGCGGTGATGTTGCAAACACAGAAGTCTTCTTT | 1 | TRBV13-2*00 | | TRBJ1-1*00 | 0.35% |
| CD4+YFP- DKO1 | TGTGCCAGCGGTGATGGGGGTGCAGAAACGCTGTATTTT | 1 | TRBV13-2*00 | | TRBJ2-3*00 | 0.35% |
| CD4+YFP- DKO1 | TGTGCCAGCGGTGACAGGGACCAAGACACCCAGTACTTT | 1 | TRBV13-2*00 | TRBD1*00 | TRBJ2-5*00 | 0.35% |
| CD4+YFP- DKO1 | TGTGCCAGCGGTGAGGTGGCAAACTCCGACTACACCTTC | 1 | TRBV13-2*00 | | TRBJ1-2*00 | 0.35% |
| CD4+YFP- DKO1 | TGTGCCAGCGGGGACAGGGGGAATTCGCCCCTCTACTTT | 1 | TRBV13-2*00 | TRBD1*00 | TRBJ1-6*00 | 0.35% |
| CD4+YFP- DKO1 | TGTGCCAGCAGTTTATCAGCAAACTCCGACTACACCTCC | 1 | TRBV15*00 | | TRBJ1-2*00 | 0.35% |
| CD4+YFP- DKO1 | TGTGCCAGCAGTTCACAGGGGACCGGGCAGCTCTACTTT | 1 | TRBV15*00 | TRBD1*00 | TRBJ2-2*00 | 0.35% |
| CD4+YFP- DKO1 | TGTGCCAGCAGTTGGGACAGGGAGACGCCCCTCTACTTT | 1 | TRBV15*00 | TRBD1*00 | TRBJ1-6*00 | 0.35% |
| CD4+YFP- DKO1 | TGTGCCAGCAGTCCCAAGACAGGGGTTGAACAGTACTTC | 1 | TRBV13-1*00 | TRBD1*00 | TRBJ2-7*00 | 0.35% |
| CD4+YFP- DKO1 | TGTGCCAGCAGTCCAGAGAGTCAAAACACCTTGTACTTT | 1 | TRBV3*00 | | TRBJ2-4*00 | 0.35% |
| CD4+YFP- DKO1 | TGTGCCAGCAGTCGGGACACCGCAGAAACGCTGTATTTT | 1 | TRBV15*00 | TRBD1*00 | TRBJ2-3*00 | 0.35% |
| CD4+YFP- DKO1 | TGTGCCAGCAGTCAGGGGGGCAACGAAAGATTATTTTTC | 1 | TRBV21*00 | TRBD2*00 | TRBJ1-4*00 | 0.35% |
| CD4+YFP- DKO1 | TGTGCCAGCAGTGGGGAAAATAATTCGCCCCTCTACTTT | 1 | TRBV13-1*00 | | TRBJ1-6*00 | 0.35% |
| CD4+YFP- DKO1 | TGTGCCAGCAGTGGGACATTCCAAGACACCCAGTACTTT | 1 | TRBV13-3*00 | TRBD1*00 | TRBJ2-5*00 | 0.35% |
| CD4+YFP- DKO1 | TGTGCCAGCAGTGGGACAGCAAACTCCGACTACACCTTC | 1 | TRBV13-1*00 | TRBD1*00 | TRBJ1-2*00 | 0.35% |
| CD4+YFP- DKO1 | TGTGCCAGCAGTGATCCCACTTACTCCGACTACACCTTC | 1 | TRBV13-1*00 | | TRBJ1-2*00 | 0.35% |
| CD4+YFP- DKO1 | TGTGCCAGCAGTGATGGGGGGCAAGACACCCAGTACTTT | 1 | TRBV13-1*00 | TRBD1*00 | TRBJ2-5*00 | 0.35% |
| CD4+YFP- DKO1 | TGTGCCAGCAGTGATACACGTGCAGAAACGCTGTATTTT | 1 | TRBV13-1*00 | | TRBJ2-3*00 | 0.35% |
| CD4+YFP- DKO1 | TGTGCCAGCAGTGAGGGGGGCTCCTATGAACAGTACTTC | 1 | TRBV13-1*00 | TRBD1*00 | TRBJ2-7*00 | 0.35% |

| | | | | | | |
|---|---|---|---|---|---|---|
| CD4+YFP- DKO1 | TGTGCCAGCAGTGAGGGGGAAACCGGGCAGCTCTACTTT | 1 | TRBV13-3*00 | TRBD1*00 | TRBJ2-2*00 | 0.35% |
| CD4+YFP- DKO1 | TGTGCCAGCAGTACGGACAACTCCTATGAACAGTACTTC | 1 | TRBV19*00 | TRBD1*00 | TRBJ2-7*00 | 0.35% |
| CD4+YFP- DKO1 | TGTGCCAGCAGCTTAGACAGGGGGAGCGAACAGTACTTC | 1 | TRBV3*00 | TRBD1*00 | TRBJ2-7*00 | 0.35% |
| CD4+YFP- DKO1 | TGTGCCAGCAGCTTAGAGGACATTGCTGAGCAGTTCTTC | 1 | TRBV25*00 | | TRBJ2-1*00 | 0.35% |
| CD4+YFP- DKO1 | TGTGCCAGCAGCTTAGAACAAAACACAGAAGTCTTCTTT | 1 | TRBV3*00 | | TRBJ1-1*00 | 0.35% |
| CD4+YFP- DKO1 | TGTGCCAGCAGCTCTGCCACGGGCTCCGACTACACCTTC | 1 | TRBV25*00 | | TRBJ1-2*00 | 0.35% |
| CD4+YFP- DKO1 | TGTGCCAGCAGCTCGGACAGGGGGCAGGACTACACCTTC | 1 | TRBV3*00 | TRBD1*00 | TRBJ1-2*00 | 0.35% |
| CD4+YFP- DKO1 | TGTGCCAGCAGCCCTGGGAGCTCCTATGAACAGTACTTC | 1 | TRBV5*00 | | TRBJ2-7*00 | 0.35% |
| CD4+YFP- DKO1 | TGTGCCAGCAGCCCCGGGACAGCCTATGAACAGTACTTC | 1 | TRBV5*00 | TRBD1*00 | TRBJ2-7*00 | 0.35% |
| CD4+YFP- DKO1 | TGTGCCAGCAGCCACGACAACTATGCTGAGCAGTTCTTC | 1 | TRBV5*00 | | TRBJ2-1*00 | 0.35% |
| CD4+YFP- DKO1 | TGTGCCAGCAGCCAAGGAGCAAACTCCGACTACACCTTC | 1 | TRBV5*00 | | TRBJ1-2*00 | 0.35% |
| CD4+YFP- DKO1 | TGTGCCAGCAGCCAAGATAGAGACTCCGACTACACCTTC | 1 | TRBV5*00 | | TRBJ1-2*00 | 0.35% |
| CD4+YFP- DKO1 | TGTGCCAGCAGCGCCAATGCAAACTCCGACTACACCTTC | 1 | TRBV3*00 | | TRBJ1-2*00 | 0.35% |
| CD4+YFP- DKO1 | TGTGCCAGCAGGTCAGGGAGCTCCTATGAACAGTACTTC | 1 | TRBV13-3*00 | | TRBJ2-7*00 | 0.35% |
| CD4+YFP- DKO1 | TGTGCCAGCAGGGACAATAACAACCAGGCTCCGCTTTTT | 1 | TRBV19*00 | TRBD1*00 | TRBJ1-5*00 | 0.35% |
| CD4+YFP- DKO1 | TGTGCCAGCAATTCAGGGAGTCAAAACACCTTGTACTTT | 1 | TRBV13-3*00 | TRBD1*00 | TRBJ2-4*00 | 0.35% |
| CD4+YFP- DKO1 | TGTGCCAGCAACTTCGGGACAGAAGACACCCAGTACTTT | 1 | TRBV14*00 | TRBD1*00 | TRBJ2-5*00 | 0.35% |
| CD4+YFP- DKO1 | TGTGCAAGCAGCTTCTCAGGGAACTCCGACTACACCTTC | 1 | TRBV16*00 | TRBD1*00 | TRBJ1-2*00 | 0.35% |
| CD4+YFP- DKO1 | TGTGCAAGCAGCTTGACAGGGGAAGACACCCAGTACTTT | 1 | TRBV16*00 | TRBD1*00 | TRBJ2-5*00 | 0.35% |
| CD4+YFP- DKO1 | TGTGCAAGCAGCTTGACAGGGACCACAGAAGTCTTCTTT | 1 | TRBV16*00 | TRBD1*00 | TRBJ1-1*00 | 0.35% |
| CD4+YFP- DKO1 | TGTGCAAGCAGCTTAGAGGCAAACTCCGACTACACCTTC | 1 | TRBV16*00 | | TRBJ1-2*00 | 0.35% |
| CD4+YFP- DKO1 | TGTGCAAGCAGCTCGAACTGGGGGTATGAACAGTACTTC | 1 | TRBV16*00 | TRBD2*00 | TRBJ2-7*00 | 0.35% |
| CD4+YFP- DKO1 | TGTGCAAGCAGCTGGGACAGACAAGACACCCAGTACTTT | 1 | TRBV16*00 | TRBD1*00 | TRBJ2-5*00 | 0.35% |
| CD4+YFP- DKO1 | TGTGCAAGCAGCGTGACAGCAAACTCCGACTACACCTTC | 1 | TRBV16*00 | | TRBJ1-2*00 | 0.35% |
| CD4+YFP- DKO1 | TGTGGTGCTGGACAGGGTAACTATGCTGAGCAGTTCTTC | 1 | TRBV20*00 | TRBD1*00 | TRBJ2-1*00 | 0.35% |
| CD4+YFP- DKO1 | TGTGGTGCTAGGGATACAGGGAGCACCGACTACACCTTC | 1 | TRBV20*00 | TRBD1*00 | TRBJ1-2*00 | 0.35% |
| CD4+YFP- DKO1 | TGCTCCAGCAGTCAATCCGGGGACTCCGACTACACCTTC | 1 | TRBV23*00 | | TRBJ1-2*00 | 0.35% |
| CD4+YFP- DKO1 | TGCACCTGCAGTGCCCGGGGAGAAGACACCCAGTACTTT | 1 | TRBV1*00 | TRBD1*00 | TRBJ2-5*00 | 0.35% |
| CD4+YFP- DKO1 | TGCACCTGCAGTGCAGAGAGAAACTCCGACTACACCTTC | 1 | TRBV1*00 | | TRBJ1-2*00 | 0.35% |
| CD4+YFP- DKO1 | TGCACCTGCAGTGCAGAAATGAACGAAAGATTATTTTTC | 1 | TRBV1*00 | | TRBJ1-4*00 | 0.35% |
| CD4+YFP- DKO1 | TGCACCTGCAGTGCAAATTCTGGAAATACGCTCTATTTT | 1 | TRBV1*00 | | TRBJ1-3*00 | 0.35% |
| CD4+YFP- DKO1 | TGCACCTGCAGTGGGACGAACTATGCTGAGCAGTTCTTC | 1 | TRBV1*00 | | TRBJ2-1*00 | 0.35% |
| CD4+YFP- DKO1 | TGTGCCAGCAGTATTACTAGTGCAGAAACGCTGATTTT | 1 | TRBV19*00 | | TRBJ2-3*00 | 0.35% |
| CD4+YFP- DKO1 | TGTGCAAGCAGCTTAGAGTCAAAACACCTTGTACTTT | 1 | TRBV16*00 | | TRBJ2-4*00 | 0.35% |
| CD4+YFP- DKO1 | TGTGCCAGCTCTCGGGGGAACTCCGACTACACCTTC | 1 | TRBV12-1*00 | TRBD1*00 | TRBJ1-2*00 | 0.35% |
| CD4+YFP- DKO1 | TGTGCCAGCCGCCAGGCAAACTCCGACTACACCTTC | 1 | TRBV13-2*00 | | TRBJ1-2*00 | 0.35% |
| CD4+YFP- DKO1 | TGTGCCAGCGGTGATGCAGACTCCGACTACACCTTC | 1 | TRBV13-2*00 | | TRBJ1-2*00 | 0.35% |
| CD4+YFP- DKO1 | TGTGCCAGCGGGGCAAACCAAGACACCCAGTACTTT | 1 | TRBV14*00 | TRBD1*00 | TRBJ2-5*00 | 0.35% |
| CD4+YFP- DKO1 | TGTGCCAGCGGAGGGGACTCCTATGAACAGTACTTC | 1 | TRBV13-2*00 | TRBD1*00 | TRBJ2-7*00 | 0.35% |
| CD4+YFP- DKO1 | TGTGCCAGCACCCTTTCTGGAAATACGCTCTATTTT | 1 | TRBV19*00 | | TRBJ1-3*00 | 0.35% |
| CD4+YFP- DKO1 | TGTGCCAGCACGGACAGAAACACAGAAGTCTTCTTT | 1 | TRBV19*00 | TRBD1*00 | TRBJ1-1*00 | 0.35% |
| CD4+YFP- DKO1 | TGTGCCAGCAGTTTCTCCCAGGGAGTAGTCTTCTTT | 1 | TRBV14*00 | TRBD1*00 | TRBJ1-1*00 | 0.35% |
| CD4+YFP- DKO1 | TGTGCCAGCAGTTTGGGACATACAGAAGTCTTCTTT | 1 | TRBV15*00 | TRBD1*00 | TRBJ1-1*00 | 0.35% |
| CD4+YFP- DKO1 | TGTGCCAGCAGTTTAGTGGGGGGGGAACAGTACTTC | 1 | TRBV15*00 | TRBD2*00 | TRBJ2-7*00 | 0.35% |
| CD4+YFP- DKO1 | TGTGCCAGCAGTCTGTCGTCAGAAACGCTGTATTTT | 1 | TRBV26*00 | | TRBJ2-3*00 | 0.35% |
| CD4+YFP- DKO1 | TGTGCCAGCAGTCGGACAGAAAACACCTTGTACTTT | 1 | TRBV19*00 | TRBD1*00 | TRBJ2-4*00 | 0.35% |
| CD4+YFP- DKO1 | TGTGCCAGCAGTCAGACATCGTATGAACAGTACTTC | 1 | TRBV26*00 | | TRBJ2-7*00 | 0.35% |
| CD4+YFP- DKO1 | TGTGCCAGCAGTGGTGACAATTATGAACAGTACTTC | 1 | TRBV13-1*00 | | TRBJ2-7*00 | 0.35% |
| CD4+YFP- DKO1 | TGTGCCAGCAGTGATTTCGACTCCGACTACACCTTC | 1 | TRBV13-3*00 | | TRBJ1-2*00 | 0.35% |
| CD4+YFP- DKO1 | TGTGCCAGCAGTGATGAGGGGAACACCTTGTACTTT | 1 | TRBV13-1*00 | TRBD1*00 | TRBJ2-4*00 | 0.35% |
| CD4+YFP- DKO1 | TGTGCCAGCAGTGAGGGGAACACAGAAGTCTTCTTT | 1 | TRBV13-1*00 | | TRBJ1-1*00 | 0.35% |
| CD4+YFP- DKO1 | TGTGCCAGCAGTGAGAACTATGCTGAGCAGTTCTTC | 1 | TRBV13-1*00 | | TRBJ2-1*00 | 0.35% |
| CD4+YFP- DKO1 | TGTGCCAGCAGTGAAAGGGGCACAGAAGTCTTCTTT | 1 | TRBV13-1*00 | TRBD1*00 | TRBJ1-1*00 | 0.35% |
| CD4+YFP- DKO1 | TGTGCCAGCAGTATGAGAGCCACAGAAGTCTTCTTT | 1 | TRBV19*00 | | TRBJ1-1*00 | 0.35% |
| CD4+YFP- DKO1 | TGTGCCAGCAGTATATCCCGAGGGGAAGTCTTCTTT | 1 | TRBV19*00 | TRBD1*00 | TRBJ1-1*00 | 0.35% |
| CD4+YFP- DKO1 | TGTGCCAGCAGTACAAACACCGGGCAGCTCTACTTT | 1 | TRBV15*00 | | TRBJ2-2*00 | 0.35% |
| CD4+YFP- DKO1 | TGTGCCAGCAGTAAGGACAACACAGAAGTCTTCTTT | 1 | TRBV15*00 | | TRBJ1-1*00 | 0.35% |
| CD4+YFP- DKO1 | TGTGCCAGCAGCTTTGAGGGGTCCGACTACACCTTC | 1 | TRBV3*00 | TRBD1*00 | TRBJ1-2*00 | 0.35% |
| CD4+YFP- DKO1 | TGTGCCAGCAGCTTGACCCCCTATGAACAGTACTTC | 1 | TRBV3*00 | | TRBJ2-7*00 | 0.35% |
| CD4+YFP- DKO1 | TGTGCCAGCAGCTCGCGGGACAAAGAAGTCTTCTTT | 1 | TRBV25*00 | | TRBJ1-1*00 | 0.35% |
| CD4+YFP- DKO1 | TGTGCCAGCAGCTCGACAGGGTTTGAACAGTACTTC | 1 | TRBV25*00 | TRBD1*00 | TRBJ2-7*00 | 0.35% |
| CD4+YFP- DKO1 | TGTGCCAGCAGCTATTGGTCTGAAACGCTGTATTTT | 1 | TRBV4*00 | | TRBJ2-3*00 | 0.35% |
| CD4+YFP- DKO1 | TGTGCCAGCAGCTACCCGAACTCCGACTACACCTTC | 1 | TRBV4*00 | | TRBJ1-2*00 | 0.35% |
| CD4+YFP- DKO1 | TGTGCCAGCAGCCCTGCAAACACAGAAGTCTTCTTT | 1 | TRBV2*00 | | TRBJ1-1*00 | 0.35% |
| CD4+YFP- DKO1 | TGTGCCAGCAGCCCCAGCTCCTATGAACAGTACTTC | 1 | TRBV3*00 | | TRBJ2-7*00 | 0.35% |
| CD4+YFP- DKO1 | TGTGCCAGCAGCCGACAGGATGCTGAGCAGTTCTTC | 1 | TRBV26*00 | TRBD1*00 | TRBJ2-1*00 | 0.35% |
| CD4+YFP- DKO1 | TGTGCCAGCAGCCACCTAAACTCCGACTACACCTTC | 1 | TRBV2*00 | | TRBJ1-2*00 | 0.35% |
| CD4+YFP- DKO1 | TGTGCCAGCAGCGACAGGGTGGACACCCAGTACTTT | 1 | TRBV2*00 | TRBD1*00 | TRBJ2-5*00 | 0.35% |
| CD4+YFP- DKO1 | TGTGCCAGCAGCACACAAAACTCCGACTACACCTTC | 1 | TRBV3*00 | | TRBJ1-2*00 | 0.35% |
| CD4+YFP- DKO1 | TGTGCCAGCAGCAGGGTCAACGAAAGATTATTTTTC | 1 | TRBV15*00 | TRBD1*00 | TRBJ1-4*00 | 0.35% |
| CD4+YFP- DKO1 | TGTGCCAGGGGGGGGAACTATGCTGAGCAGTTCTTC | 1 | TRBV31*00 | TRBD2*00 | TRBJ2-1*00 | 0.35% |
| CD4+YFP- DKO1 | TGTGCAAGCAGCTTAGGGACATTTGAACAGTACTTC | 1 | TRBV16*00 | TRBD1*00 | TRBJ2-7*00 | 0.35% |
| CD4+YFP- DKO1 | TGTGCAAGCAGCTTAGAGGGGTATGAACAGTACTTC | 1 | TRBV16*00 | | TRBJ2-7*00 | 0.35% |
| CD4+YFP- DKO1 | TGTGCAAGCAGCTTAGAGGACACAGAAGTCTTCTTT | 1 | TRBV16*00 | | TRBJ1-1*00 | 0.35% |

| | | | | | | |
|---|---|---|---|---|---|---|
| CD4+YFP- DKO1 | TGCACCTGTGGGGGGGACCAAGACACCCAGTACTTT | 1 | TRBV1*00 | TRBD2*00 | TRBJ2-5*00 | 0.35% |
| CD4+YFP- DKO1 | TGCACCTGCAGTGCCGACAACTATGAACAGTACTTC | 1 | TRBV1*00 | | TRBJ2-7*00 | 0.35% |
| CD4+YFP- DKO1 | TGTGCCTGGAGTACGGACTGGGGACAGTACTTC | 1 | TRBV31*00 | TRBD2*00 | TRBJ2-7*00 | 0.35% |
| CD4+YFP- DKO1 | TGTGCCATGGGAAGTCAAAACACCTTGTACTTT | 1 | TRBV13-2*00 | | TRBJ2-4*00 | 0.35% |
| CD4+YFP- DKO1 | TGTGCCAGCTCTCTTCTCTATGAACAGTACTTC | 1 | TRBV12-1*00 | | TRBJ2-7*00 | 0.35% |
| CD4+YFP- DKO1 | TGTGCCAGCGGTGGCCGCAGAGAACAGTACTTC | 1 | TRBV13-2*00 | | TRBJ2-7*00 | 0.35% |
| CD4+YFP- DKO1 | TGTGCCAGCGGTAACCAAGACACCCAGTACTTT | 1 | TRBV13-2*00 | | TRBJ2-5*00 | 0.35% |
| CD4+YFP- DKO1 | TGTGCCAGCAGTTTACGCTATGAACAGTACTTC | 1 | TRBV15*00 | | TRBJ2-7*00 | 0.35% |
| CD4+YFP- DKO1 | TGTGCCAGCAGTTCAAACACAGAAGTCTTCTTT | 1 | TRBV15*00 | | TRBJ1-1*00 | 0.35% |
| CD4+YFP- DKO1 | TGTGCCAGCAGTCTTCGGGAAGAAGTCTTCTTT | 1 | TRBV26*00 | TRBD1*00 | TRBJ1-1*00 | 0.35% |
| CD4+YFP- DKO1 | TGTGCCAGCAGCTTACAAGACACCCAGTACTTT | 1 | TRBV25*00 | | TRBJ2-5*00 | 0.35% |
| CD4+YFP- DKO1 | TGTGCCAGCAGCCAAGAAAATACGCTCTATTTT | 1 | TRBV2*00 | | TRBJ1-3*00 | 0.35% |
| CD4+YFP- DKO1 | TGTGCCAGCAGGGACAACTATGAACAGTACTTC | 1 | TRBV19*00 | TRBD1*00 | TRBJ2-7*00 | 0.35% |
| CD4+YFP- DKO1 | TGTGCAAGCAGCCCATCCTATGAACAGTACTTC | 1 | TRBV16*00 | | TRBJ2-7*00 | 0.35% |
| CD4+YFP- DKO1 | TGTGGTGCTTGGGGGGCTGAAACGCTGTATTTT | 1 | TRBV20*00 | | TRBJ2-3*00 | 0.35% |
| CD4+YFP- DKO1 | TGTAGTTCTAGTACTGCAGAAACGCTGTATTTT | 1 | TRBV30*00 | | TRBJ2-3*00 | 0.35% |
| CD4+YFP- DKO1 | TGCACCTGCAGTGCAGCAGACACCCAGTACTTT | 1 | TRBV1*00 | | TRBJ2-5*00 | 0.35% |
| CD4+YFP- DKO1 | TGCACCTGCAGTGCAGGGTATGAACAGTACTTC | 1 | TRBV1*00 | | TRBJ2-7*00 | 0.35% |
| CD4+YFP- DKO1 | TGCACCTGCAGTGCAGATGGCTCCGCTTTTT | 1 | TRBV1*00 | | TRBJ1-5*00 | 0.35% |
| CD4+YFP- DKO1 | TGTGCCAGCAGCCTCTATGAACAGTACTTC | 1 | TRBV4*00 | | TRBJ2-7*00 | 0.35% |
| CD4+YFP- DKO1 | TGTGCCAGCAGCCACCGGGAAGTCTTCTTT | 1 | TRBV4*00 | | TRBJ1-1*00 | 0.35% |
| CD4+YFP- DKO1 | TGTGCCAGCAGCAACACAGAAGTCTTCTTT | 1 | TRBV4*00 | | TRBJ1-1*00 | 0.35% |
| CD4+YFP- DKO1 | TGTGCCTGGAGGACTCCCCAGTACTTC | 1 | TRBV31*00 | | TRBJ2-7*00 | 0.35% |
| CD4+YFP- DKO1 | TGTGCCAGCGGTGATGAACAGTACTTC | 1 | TRBV13-2*00 | | TRBJ2-7*00 | 0.35% |
| CD4+YFP- DKO1 | TGTGCCAGCAGCTATGAACAGTACTTC | 1 | TRBV4*00 | | TRBJ2-7*00 | 0.35% |
| CD4+YFP- DKO1 | TGTGCTAGCAGTAGAAAGGAC | 1 | TRBV17*00 | | TRBJ2-3*00 | 0.35% |

#### CD4+YFP- DKO2

| Sample name | CDR3 Sequence | clone Count | best V Hit | best D Hit | best J Hit | Frequency |
|---|---|---|---|---|---|---|
| CD4+YFP- DKO2 | TGTGCCAGCGGTGATGTGGACTGGGGGAGTGCAGAAACGCTGTATTTT | 408 | TRBV13-2*00 | TRBD2*00 | TRBJ2-3*00 | 89.67% |
| CD4+YFP- DKO2 | TGTGCCAGCATCTCCGGGACTTCAAACACCGGGCAGCTCTACTTT | 2 | TRBV13-3*00 | TRBD2*00 | TRBJ2-2*00 | 0.44% |
| CD4+YFP- DKO2 | TGTGCCAGCTCTCCCTGGGCAGGGAACTCCGACTACACCTTC | 2 | TRBV12-1*00 | TRBD1*00 | TRBJ1-2*00 | 0.44% |
| CD4+YFP- DKO2 | TGTGCCAGCAGTTTTGGGGAAGGGCAGGCTCCGCTTTTT | 2 | TRBV15*00 | TRBD2*00 | TRBJ1-5*00 | 0.44% |
| CD4+YFP- DKO2 | TGTGCCAGCGGTGATGCCCTGGGGGGTCCTAGTGCAGAAACGCTGTATTT | 1 | TRBV13-2*00 | TRBD2*00 | TRBJ2-3*00 | 0.22% |
| CD4+YFP- DKO2 | TGTGCCAGCGGTGATAGGGGAAATTCTGGAAATACGCTCTATTTT | 1 | TRBV13-2*00 | TRBD1*00 | TRBJ1-3*00 | 0.22% |
| CD4+YFP- DKO2 | TGTGCAAGCAGCTTAGCGGCAGGGTCAAACTCCGACTACACCTTC | 1 | TRBV16*00 | TRBD1*00 | TRBJ1-2*00 | 0.22% |
| CD4+YFP- DKO2 | TGCACCTGCAGTGCAGATCCTGGGGACTATGCTGAGCAGTTCTTC | 1 | TRBV1*00 | TRBD2*00 | TRBJ2-1*00 | 0.22% |
| CD4+YFP- DKO2 | TGCACCTGCAGTGCAGAGGGGACTCTCTCCTATGAACAGTACTTC | 1 | TRBV1*00 | TRBD2*00 | TRBJ2-7*00 | 0.22% |
| CD4+YFP- DKO2 | TGCACCTGCAGTGCAACGCCTGGGGGCTCCTATGAACAGTACTTC | 1 | TRBV1*00 | TRBD2*00 | TRBJ2-7*00 | 0.22% |
| CD4+YFP- DKO2 | TGTGCCAGCAGCAACAGGGGGCTAGTCAAAACACCTTGTACTTT | 1 | TRBV13-1*00 | TRBD1*00 | TRBJ2-4*00 | 0.22% |
| CD4+YFP- DKO2 | TGTGCCAGCTCTCTCGAAGGGGTGCACCGGGCAGCTCTACTTT | 1 | TRBV12-1*00 | TRBD1*00 | TRBJ2-2*00 | 0.22% |
| CD4+YFP- DKO2 | TGTGCCTGGAGTCTCTGGGGGGACCAAGACACCCAGTACTTT | 1 | TRBV31*00 | TRBD2*00 | TRBJ2-5*00 | 0.22% |
| CD4+YFP- DKO2 | TGTGCCAGCTCAACCGGGACACCAAACTCCGACTACACCTTC | 1 | TRBV12-1*00 | TRBD1*00 | TRBJ1-2*00 | 0.22% |
| CD4+YFP- DKO2 | TGTGCCAGCGGTGGGGGACAGGGGGCCTCCGACTACACCTTC | 1 | TRBV13-2*00 | TRBD1*00 | TRBJ1-2*00 | 0.22% |
| CD4+YFP- DKO2 | TGTGCCAGCACCCCGGGACAGTCGTATGCTGAGCAGTTCTTC | 1 | TRBV13-3*00 | TRBD1*00 | TRBJ2-1*00 | 0.22% |
| CD4+YFP- DKO2 | TGTGCCAGCAGTGCTACGGACAGGGGAAACACCTTGTACTTT | 1 | TRBV13-3*00 | TRBD1*00 | TRBJ2-4*00 | 0.22% |
| CD4+YFP- DKO2 | TGTGCCAGCAGCTTTTCAATTTCCAACGAAAGATTATTTTTC | 1 | TRBV3*00 | | TRBJ1-4*00 | 0.22% |
| CD4+YFP- DKO2 | TGTGCCAGCAGCTTAGAGGTTACGGTTGCTGAGCAGTTCTTC | 1 | TRBV3*00 | | TRBJ2-1*00 | 0.22% |
| CD4+YFP- DKO2 | TGTGCCAGCAGCTACCGGGACAGCTCCTATGAACAGTACTTC | 1 | TRBV4*00 | TRBD1*00 | TRBJ2-7*00 | 0.22% |
| CD4+YFP- DKO2 | TGTGCTAGCAGGGCGAATAACTATGCTGAGCAGTTCTTC | 1 | TRBV29*00 | | TRBJ2-1*00 | 0.22% |
| CD4+YFP- DKO2 | TGTGCCAGCGGTGATGCTTGGCAAAACACCTTGTACTTT | 1 | TRBV13-2*00 | | TRBJ2-4*00 | 0.22% |
| CD4+YFP- DKO2 | TGTGCCAGCGGTGATGAAAACCAAGACACCCAGTACTTT | 1 | TRBV13-2*00 | | TRBJ2-5*00 | 0.22% |
| CD4+YFP- DKO2 | TGTGCCAGCAGTTTGGGGACAGCCTATGAACAGTACTTC | 1 | TRBV15*00 | TRBD1*00 | TRBJ2-7*00 | 0.22% |
| CD4+YFP- DKO2 | TGTGCCAGCAGTTTAGGGGCAAACTCCGACTACACCTTC | 1 | TRBV15*00 | | TRBJ1-2*00 | 0.22% |
| CD4+YFP- DKO2 | TGTGCCAGCAGTGAGGGACAGGGGGACACCCAGTACTTT | 1 | TRBV13-1*00 | TRBD1*00 | TRBJ2-5*00 | 0.22% |
| CD4+YFP- DKO2 | TGTGCCAGCAGTATGAAGACAGACACAGAAGTCTTCTTT | 1 | TRBV19*00 | TRBD1*00 | TRBJ1-1*00 | 0.22% |
| CD4+YFP- DKO2 | TGTGCCAGCAGCTCGACAGGGGACACAGAAGTCTTCTTT | 1 | TRBV25*00 | TRBD1*00 | TRBJ1-1*00 | 0.22% |
| CD4+YFP- DKO2 | TGTGCCAGCAGCCCCGAGACAGGCTTTGAACAGTACTTC | 1 | TRBV5*00 | TRBD1*00 | TRBJ2-7*00 | 0.22% |
| CD4+YFP- DKO2 | TGTGCCAGCAGGGGGGCGCGGGCAGAAACGCTGTATTTT | 1 | TRBV13-1*00 | TRBD1*00 | TRBJ2-3*00 | 0.22% |
| CD4+YFP- DKO2 | TGTGCCAGCAACCAGGGGGTGTCCTATGAACAGTACCTC | 1 | TRBV15*00 | TRBD1*00 | TRBJ2-7*00 | 0.22% |
| CD4+YFP- DKO2 | TGTGCAAGCAGCTTAGGGGGGAACCAGGCTCCGCTTTTT | 1 | TRBV16*00 | TRBD1*00 | TRBJ1-5*00 | 0.22% |
| CD4+YFP- DKO2 | TGTGCAAGCAGCTTAGACAGTCAAAACACCTTGTACTTT | 1 | TRBV16*00 | | TRBJ2-4*00 | 0.22% |
| CD4+YFP- DKO2 | TGTGCTAGCAGCTACAGTGGAAATACGCTCTATTTT | 1 | TRBV21*00 | | TRBJ1-3*00 | 0.22% |
| CD4+YFP- DKO2 | TGTGCCTGGACAGGGGATGGAAATACGCTCTATTTT | 1 | TRBV31*00 | TRBD1*00 | TRBJ1-3*00 | 0.22% |
| CD4+YFP- DKO2 | TGTGCCAGCAGTTTTTGGGACTCCGACTACACCTTC | 1 | TRBV14*00 | | TRBJ1-2*00 | 0.22% |
| CD4+YFP- DKO2 | TGTGCCAGCAGTTTCAGGGGAGACACCTTGTACTTT | 1 | TRBV15*00 | TRBD1*00 | TRBJ2-4*00 | 0.22% |
| CD4+YFP- DKO2 | TGTGCCAGCAGTTTAGGGGAAAACACCTTGTACTTT | 1 | TRBV15*00 | | TRBJ2-4*00 | 0.22% |
| CD4+YFP- DKO2 | TGTGCCAGCAGCCATAACCAAGACACCCAGTACTTT | 1 | TRBV4*00 | | TRBJ2-5*00 | 0.22% |
| CD4+YFP- DKO2 | TGTGCCAGCAGCCAAGATAAAAACACCTTGTACTTT | 1 | TRBV2*00 | | TRBJ2-4*00 | 0.22% |
| CD4+YFP- DKO2 | TGTGGTGCTAGGGGGGACACCGGGCAGCTCTACTTT | 1 | TRBV20*00 | | TRBJ2-2*00 | 0.22% |
| CD4+YFP- DKO2 | TGTGGTGCTAGGGACCCTGCAGAAACGCTGTATTTT | 1 | TRBV20*00 | | TRBJ2-3*00 | 0.22% |
| CD4+YFP- DKO2 | TGCACCTGCAGTGCGTCTGGAAATACGCTCTATTTT | 1 | TRBV1*00 | | TRBJ1-3*00 | 0.22% |
| CD4+YFP- DKO2 | TGTGCTAGCAGTTTCAACTCCGACTACACCTTC | 1 | TRBV29*00 | | TRBJ1-2*00 | 0.22% |
| CD4+YFP- DKO2 | TGTGCCAGCAGCTTCTCCTATGAACAGTACTTC | 1 | TRBV3*00 | | TRBJ2-7*00 | 0.22% |

#### CD4+YFP- DKO3

| Sample name | CDR3 Sequence | clone Count | best V Hit | best D Hit | best J Hit | Frequency |
|---|---|---|---|---|---|---|
| CD4+YFP- DKO3 | TGTGCCAGCAGCCAACTGGGAGCAAACACCGGGCAGCTCTACTTT | 57 | TRBV5*00 | TRBD2*00 | TRBJ2-2*00 | 29.84% |
| CD4+YFP- DKO3 | TGTGCCAGCAGCCAAGAAGGAATCCAAGACACCCAGTACTTT | 3 | TRBV2*00 | | TRBJ2-5*00 | 1.57% |
| CD4+YFP- DKO3 | TGTGCAAGCAGCTTAGATCGGGGTGAACAGTACTTC | 3 | TRBV16*00 | | TRBJ2-7*00 | 1.57% |
| CD4+YFP- DKO3 | TGTGCCAGCAGCCGGGACTGGGGACCTATGCTGAGCAGTTCTTC | 2 | TRBV13-1*00 | TRBD2*00 | TRBJ2-1*00 | 1.05% |
| CD4+YFP- DKO3 | TGTGCCAGCAGTTTATCGGGGAGTGCAGAAACGCTGTATTTT | 2 | TRBV15*00 | | TRBJ2-3*00 | 1.05% |
| CD4+YFP- DKO3 | TGTGCCAGCAGTCAACTGGGGAACTATGCTGAGCAGTTCTTC | 2 | TRBV26*00 | TRBD2*00 | TRBJ2-1*00 | 1.05% |
| CD4+YFP- DKO3 | TGTGCCAGCAGGCCCCGGGACAGGTACTCCGACTACACCTTC | 2 | TRBV3*00 | TRBD1*00 | TRBJ1-2*00 | 1.05% |
| CD4+YFP- DKO3 | TGTGCAAGCAGCTTAGAAACAAAAGACACCCAGTACTTT | 2 | TRBV16*00 | | TRBJ2-5*00 | 1.05% |
| CD4+YFP- DKO3 | TGTGCTAGCAGCCGGGACAGGGTCGAAGTCTTCTTT | 2 | TRBV17*00 | TRBD1*00 | TRBJ1-1*00 | 1.05% |
| CD4+YFP- DKO3 | TGTGCCAGCAGCTATGGGGGGACAGAAGTCTTCTTT | 2 | TRBV4*00 | TRBD2*00 | TRBJ1-1*00 | 1.05% |
| CD4+YFP- DKO3 | TGTGCCTGGAGTCCAGGGACAGGGAATTCTGGAAATACGCTCTATTTT | 1 | TRBV31*00 | TRBD1*00 | TRBJ1-3*00 | 0.52% |
| CD4+YFP- DKO3 | TGTGCCAGCAGTCTGTCGTTGGACAGGGGCCAAGACACCCAGTACTTT | 1 | TRBV26*00 | TRBD1*00 | TRBJ2-5*00 | 0.52% |
| CD4+YFP- DKO3 | TGCTCCAGCAGACTGGGGGGGGGAACTAGTGCAGAAACGCTGTATTTT | 1 | TRBV23*00 | TRBD2*00 | TRBJ2-3*00 | 0.52% |
| CD4+YFP- DKO3 | TGCACCTGCAGTGCCATGGACAATTATAACAACCAGGCTCCGCTTTTT | 1 | TRBV1*00 | TRBD1*00 | TRBJ1-5*00 | 0.52% |
| CD4+YFP- DKO3 | TGTGCTAGCAGTTTGACAGGGGCAAACACCGGGCAGCTCTACTTT | 1 | TRBV29*00 | TRBD1*00 | TRBJ2-2*00 | 0.52% |
| CD4+YFP- DKO3 | TGTGCCAGCGGTGACCCCGGGACAGTCTATGCTGAGCAGTTCTTC | 1 | TRBV13-2*00 | TRBD1*00 | TRBJ2-1*00 | 0.52% |
| CD4+YFP- DKO3 | TGTGCCAGCAGTGGAGGGACATCTAGTGCAGAAACGCTGTATTTT | 1 | TRBV13-3*00 | TRBD1*00 | TRBJ2-3*00 | 0.52% |
| CD4+YFP- DKO3 | TGTGCCAGCAGTGATGCTGGACTGGGTCAAGACACCCAGTACTTT | 1 | TRBV13-1*00 | TRBD2*00 | TRBJ2-5*00 | 0.52% |
| CD4+YFP- DKO3 | TGTGCCAGCAGTGATGCCGGAGTTTCTGGAAATACGCTCTATTTT | 1 | TRBV13-1*00 | | TRBJ1-3*00 | 0.52% |
| CD4+YFP- DKO3 | TGTGCCAGCAGTGATGGGGGCCCTAACTATGCTGAGCAGTTCTTC | 1 | TRBV13-3*00 | TRBD1*00 | TRBJ2-1*00 | 0.52% |
| CD4+YFP- DKO3 | TGTGCCAGCAGTGACCCTGGGGTTAGTGCAGAAACGCTGTATTTT | 1 | TRBV13-1*00 | TRBD2*00 | TRBJ2-3*00 | 0.52% |
| CD4+YFP- DKO3 | TGTGCCAGCAGTGAGCGGGGGGCTAACTATGCTGAGCAGTTCTTC | 1 | TRBV13-3*00 | TRBD2*00 | TRBJ2-1*00 | 0.52% |
| CD4+YFP- DKO3 | TGTGCCAGCAGTGAACTGGGGGGGCACTATGCTGAGCAGTTCTTC | 1 | TRBV13-3*00 | TRBD2*00 | TRBJ2-1*00 | 0.52% |
| CD4+YFP- DKO3 | TGTGCCAGCAGCTTTGGGGGGGCAAACACCGGGCAGCTCTACTTT | 1 | TRBV3*00 | TRBD2*00 | TRBJ2-2*00 | 0.52% |
| CD4+YFP- DKO3 | TGTGCCAGCAGCTTAGCACAGACTAGTGCAGAAACGCTGTATTTT | 1 | TRBV3*00 | | TRBJ2-3*00 | 0.52% |
| CD4+YFP- DKO3 | TGTGCCAGCAGCTTAAGACTGGGGGGTGCAGAAACGCTGTATTTT | 1 | TRBV3*00 | TRBD2*00 | TRBJ2-3*00 | 0.52% |
| CD4+YFP- DKO3 | TGTGCCAGCAGCCAAGCAGGGGGCTGGTATGCTGAGCAGTTCTTC | 1 | TRBV2*00 | TRBD1*00 | TRBJ2-1*00 | 0.52% |
| CD4+YFP- DKO3 | TGTGCCAGCAGCCAAGACAGGGCTAACCAAGACACCCAGTACTTT | 1 | TRBV5*00 | TRBD1*00 | TRBJ2-5*00 | 0.52% |
| CD4+YFP- DKO3 | TGTGCCAGCAGCCAAGAAGGGGGGAGTGCAGAAACGCTGTATTTT | 1 | TRBV5*00 | TRBD1*00 | TRBJ2-3*00 | 0.52% |
| CD4+YFP- DKO3 | TGTGCCAGCAGTTGACAGGGGGCGAAACCAGGCTCCGCTTTTT | 1 | TRBV22*00 | TRBD1*00 | TRBJ1-5*00 | 0.52% |
| CD4+YFP- DKO3 | TGTGCTAGCAGTAGAGACGGACGAGGAGCTGAGCAGTTCTTC | 1 | TRBV17*00 | | TRBJ2-1*00 | 0.52% |
| CD4+YFP- DKO3 | TGTGCCTGGAGCCTACAGGGAAGTGCAGAAACGCTGTATTTT | 1 | TRBV31*00 | TRBD1*00 | TRBJ2-3*00 | 0.52% |
| CD4+YFP- DKO3 | TGTGCCAGCTCTCTTCCTGGGGGCGAGGACACCTTGTACTTT | 1 | TRBV12-1*00 | TRBD1*00 | TRBJ2-4*00 | 0.52% |
| CD4+YFP- DKO3 | TGTGCCAGCGGTGATCTGGGGCGTCAAAACACCTTGTACTTT | 1 | TRBV13-2*00 | TRBD2*00 | TRBJ2-4*00 | 0.52% |
| CD4+YFP- DKO3 | TGTGCCAGCAGTTTCGGGACAACCACCGGGCAGCTCTACTTT | 1 | TRBV15*00 | TRBD1*00 | TRBJ2-2*00 | 0.52% |
| CD4+YFP- DKO3 | TGTGCCAGCAGTTTACCAGCAAAAAACTCCGACTACACCTTC | 1 | TRBV15*00 | | TRBJ1-2*00 | 0.52% |
| CD4+YFP- DKO3 | TGTGCCAGCAGTTTAGGGGGAAGCTCCTATGAACAGTACTTC | 1 | TRBV15*00 | | TRBJ2-7*00 | 0.52% |
| CD4+YFP- DKO3 | TGTGCCAGCAGTTGGGGACTGGGGGAAAACACCTTGTACTTT | 1 | TRBV15*00 | TRBD2*00 | TRBJ2-4*00 | 0.52% |
| CD4+YFP- DKO3 | TGTGCCAGCAGTCTGGAGGCAAACACCGGGCAGCTCTACTTT | 1 | TRBV24*00 | | TRBJ2-2*00 | 0.52% |
| CD4+YFP- DKO3 | TGTGCCAGCAGTCGTGGACAGTATAATTCGCCCCTCTACTTT | 1 | TRBV22*00 | | TRBJ1-6*00 | 0.52% |
| CD4+YFP- DKO3 | TGTGCCAGCAGTGGACAGGGGGCGGGCACAGAAGTCTTCTTT | 1 | TRBV15*00 | TRBD1*00 | TRBJ1-1*00 | 0.52% |
| CD4+YFP- DKO3 | TGTGCCAGCAGTGATCTGGACAGTGCAGAAACGCTGTATTTT | 1 | TRBV13-1*00 | | TRBJ2-3*00 | 0.52% |
| CD4+YFP- DKO3 | TGTGCCAGCAGTGATGAGGCTTTTCAAAACACCTTGTACTTT | 1 | TRBV13-1*00 | | TRBJ2-4*00 | 0.52% |
| CD4+YFP- DKO3 | TGTGCCAGCAGTGATGAGGGCGGGAACACAGAAGTCTTCTTT | 1 | TRBV13-1*00 | TRBD1*00 | TRBJ1-1*00 | 0.52% |
| CD4+YFP- DKO3 | TGTGCCAGCAGTATTGGGAGGAGTGCAGAAACGCTGTATTTT | 1 | TRBV19*00 | | TRBJ2-3*00 | 0.52% |
| CD4+YFP- DKO3 | TGTGCCAGCAGTATAAACGGGCGTGCAGAAACGCTGTATTTT | 1 | TRBV19*00 | | TRBJ2-3*00 | 0.52% |
| CD4+YFP- DKO3 | TGTGCCAGCAGCTTCGACAGTTCTGGAAATACGCTCTATTTT | 1 | TRBV3*00 | TRBD1*00 | TRBJ1-3*00 | 0.52% |
| CD4+YFP- DKO3 | TGTGCCAGCAGCTTGACAGGGGCCAACCAGGCTCCGCTTTTT | 1 | TRBV4*00 | TRBD1*00 | TRBJ1-5*00 | 0.52% |
| CD4+YFP- DKO3 | TGTGCCAGCAGCTTAGTGATGGGGGGGCAAGAACAGTACTTC | 1 | TRBV3*00 | TRBD2*00 | TRBJ2-7*00 | 0.52% |
| CD4+YFP- DKO3 | TGTGCCAGCAGCTCACGGACAACAAACACAGAAGTCTTCTTT | 1 | TRBV3*00 | TRBD1*00 | TRBJ1-1*00 | 0.52% |
| CD4+YFP- DKO3 | TGTGCCAGCAGCCCGACAACTAGTCAAAACACCTTGTACTTT | 1 | TRBV5*00 | | TRBJ2-4*00 | 0.52% |
| CD4+YFP- DKO3 | TGTGCCAGCAGCCAGGACTGGCTGGATGCTGAGCAGTTCTTC | 1 | TRBV2*00 | TRBD2*00 | TRBJ2-1*00 | 0.52% |
| CD4+YFP- DKO3 | TGTGCCAGCAGCCAAGATTCAGGGGAATATGAACAGTACTTC | 1 | TRBV2*00 | TRBD1*00 | TRBJ2-7*00 | 0.52% |
| CD4+YFP- DKO3 | TGTGCCAGCAGCCAAGAAGGGGGCAACTCCGACTACACCTTC | 1 | TRBV5*00 | TRBD1*00 | TRBJ1-2*00 | 0.52% |
| CD4+YFP- DKO3 | TGTGCCAGCAGGCACCGGGACATCTCCTATGAACAGTACTTC | 1 | TRBV13-1*00 | TRBD1*00 | TRBJ2-7*00 | 0.52% |
| CD4+YFP- DKO3 | TGTGCCAGCAGGGGGACAGCAAACACCGGGCAGCTCTACTTT | 1 | TRBV13-1*00 | TRBD1*00 | TRBJ2-2*00 | 0.52% |
| CD4+YFP- DKO3 | TGTGCCAGCAGACCGGGACAGGGCCAAGACACCCAGTACTTT | 1 | TRBV19*00 | TRBD1*00 | TRBJ2-5*00 | 0.52% |
| CD4+YFP- DKO3 | TGTGCAAGCAGCTTAGAGGGAGCTGGAAATACGCTCTATTTT | 1 | TRBV16*00 | | TRBJ1-3*00 | 0.52% |
| CD4+YFP- DKO3 | TGCACCTGCAGTGCGGGGACAGCAAACTCCGACTACACCTTC | 1 | TRBV1*00 | TRBD1*00 | TRBJ1-2*00 | 0.52% |
| CD4+YFP- DKO3 | TGCACCTGCAGTGCAGGCAGGGGAGGGAACACCTTGTACTTT | 1 | TRBV1*00 | TRBD1*00 | TRBJ2-4*00 | 0.52% |
| CD4+YFP- DKO3 | TGCACCTGCAGTGCAGACAATTCTGGAAATACGCTCTATTTT | 1 | TRBV1*00 | | TRBJ1-3*00 | 0.52% |
| CD4+YFP- DKO3 | TGCACCTGCAGTGGACGGGACAACAACCAGGCTCCGCTTTTT | 1 | TRBV1*00 | TRBD1*00 | TRBJ1-5*00 | 0.52% |
| CD4+YFP- DKO3 | TGTGCTAGCAGTAGAGAATTCTGGAAATACGCTCTATTTT | 1 | TRBV29*00 | | TRBJ1-3*00 | 0.52% |
| CD4+YFP- DKO3 | TGTGCTAGCAGTGCGAAGAACTATGCTGAGCAGTTCTTC | 1 | TRBV17*00 | | TRBJ2-1*00 | 0.52% |
| CD4+YFP- DKO3 | TGTGCCTGGAGTCCAGGACAAAAATATGAACAGTACTTC | 1 | TRBV31*00 | TRBD1*00 | TRBJ2-7*00 | 0.52% |
| CD4+YFP- DKO3 | TGTGCCTGGAGTCACAGGGGGCGCGAGGAACAGTACTTC | 1 | TRBV31*00 | TRBD1*00 | TRBJ2-7*00 | 0.52% |
| CD4+YFP- DKO3 | TGTGCCTGGAGTGGGACAGGGGGCACAGAAGTCTTCTTT | 1 | TRBV31*00 | TRBD1*00 | TRBJ1-1*00 | 0.52% |
| CD4+YFP- DKO3 | TGTGCCAGCCGACAGGGCTCTGGAAATACGCTCTATTTT | 1 | TRBV3*00 | TRBD1*00 | TRBJ1-3*00 | 0.52% |
| CD4+YFP- DKO3 | TGTGCCAGCAGTTTAAGGCAGAACTCCGACTACACCTTC | 1 | TRBV15*00 | | TRBJ1-2*00 | 0.52% |
| CD4+YFP- DKO3 | TGTGCCAGCAGTTTAAGGGAGAACGAAAGATTATTTTTC | 1 | TRBV14*00 | | TRBJ1-4*00 | 0.52% |
| CD4+YFP- DKO3 | TGTGCCAGCAGTCTGTCGCGGGGGTTTGAACAGTACTTC | 1 | TRBV26*00 | TRBD1*00 | TRBJ2-7*00 | 0.52% |

| | | | | | | |
|---|---|---|---|---|---|---|
| CD4+YFP- DKO3 | TGTGCCAGCAGTCTGTCGCGGAATTCGCCCCTCTACTTT | 1 | TRBV26*00 | | TRBJ1-6*00 | 0.52% |
| CD4+YFP- DKO3 | TGTGCCAGCAGTCTGCCAGGGGAAAACACCTTGTACTTT | 1 | TRBV24*00 | TRBD1*00 | TRBJ2-4*00 | 0.52% |
| CD4+YFP- DKO3 | TGTGCCAGCAGTCGACTGGGTTATGCTGAGCAGTTCTTC | 1 | TRBV14*00 | TRBD2*00 | TRBJ2-1*00 | 0.52% |
| CD4+YFP- DKO3 | TGTGCCAGCAGTGATGCGGGAACCGGGCAGCTCTACTTT | 1 | TRBV13-1*00 | | TRBJ2-2*00 | 0.52% |
| CD4+YFP- DKO3 | TGTGCCAGCAGTGATGGCCAGAAAAACACCTTGTACTTT | 1 | TRBV13-1*00 | | TRBJ2-4*00 | 0.52% |
| CD4+YFP- DKO3 | TGTGCCAGCAGCTTTCGGGGGGACTCCGACTACACCTTC | 1 | TRBV3*00 | TRBD2*00 | TRBJ1-2*00 | 0.52% |
| CD4+YFP- DKO3 | TGTGCCAGCAGCTCCGGGGGCAACCAGGCTCCGCTTTTT | 1 | TRBV25*00 | TRBD1*00 | TRBJ1-5*00 | 0.52% |
| CD4+YFP- DKO3 | TGTGCCAGCAGCCCTCCGGGTGCAGAAACGCTGTATTTT | 1 | TRBV3*00 | | TRBJ2-3*00 | 0.52% |
| CD4+YFP- DKO3 | TGTGCCAGCAGCCCGGGGAGGGATGCTGAGCAGTTCTTC | 1 | TRBV13-3*00 | TRBD1*00 | TRBJ2-1*00 | 0.52% |
| CD4+YFP- DKO3 | TGTGCCAGCAGCCCGGACACCTCCTATGAACAGTACTTC | 1 | TRBV5*00 | TRBD1*00 | TRBJ2-7*00 | 0.52% |
| CD4+YFP- DKO3 | TGTGCCAGCAGCCGCGGGGACTCCTATGAACAGTACTTC | 1 | TRBV2*00 | | TRBJ2-7*00 | 0.52% |
| CD4+YFP- DKO3 | TGTGCCAGCAGCCGGGACTGGGGAAGCACCCAGTACTTT | 1 | TRBV4*00 | TRBD2*00 | TRBJ2-5*00 | 0.52% |
| CD4+YFP- DKO3 | TGTGCCAGCAGCCAGGGACAAAACACAGAAGTCTTCTTT | 1 | TRBV2*00 | TRBD1*00 | TRBJ1-1*00 | 0.52% |
| CD4+YFP- DKO3 | TGTGCCAGCAGACCGGGACTGGATGCTGAGCAGTTCTTC | 1 | TRBV14*00 | TRBD2*00 | TRBJ2-1*00 | 0.52% |
| CD4+YFP- DKO3 | TGTGCAAGCAGCTTAGTTGGGGGGTATGAACAGTACTTC | 1 | TRBV16*00 | TRBD2*00 | TRBJ2-7*00 | 0.52% |
| CD4+YFP- DKO3 | TGTGCAAGCAGCTTAGACAGACAAAACACCTTGTACTTT | 1 | TRBV16*00 | | TRBJ2-4*00 | 0.52% |
| CD4+YFP- DKO3 | TGTGCAAGCAGCTCTGACAGGAGCACAGAAGTCTTCTTT | 1 | TRBV16*00 | TRBD1*00 | TRBJ1-1*00 | 0.52% |
| CD4+YFP- DKO3 | TGTGCAAGCAGCAGACTGGGGGAAGACACCCAGTACTTT | 1 | TRBV16*00 | TRBD2*00 | TRBJ2-5*00 | 0.52% |
| CD4+YFP- DKO3 | TGTGGTGCTAGTAGGGGAGGTGCAGAAACGCTGTATTTT | 1 | TRBV20*00 | TRBD1*00 | TRBJ2-3*00 | 0.52% |
| CD4+YFP- DKO3 | TGTGGTGCTAGGCAGGACAGAAACTCCGACTACACCTTC | 1 | TRBV20*00 | TRBD1*00 | TRBJ1-2*00 | 0.52% |
| CD4+YFP- DKO3 | TGTAGTTCTAGACAGGGAAACTATGCTGAGCAGTTCTTC | 1 | TRBV30*00 | TRBD1*00 | TRBJ2-1*00 | 0.52% |
| CD4+YFP- DKO3 | TGCTCCAGCAGTCAATCTTCAGTAAACACCTTGTACTTT | 1 | TRBV23*00 | | TRBJ2-4*00 | 0.52% |
| CD4+YFP- DKO3 | TGCACCTGCAGTGCGGGACAGGGGACTGAGCAGTTCTTC | 1 | TRBV1*00 | TRBD1*00 | TRBJ2-1*00 | 0.52% |
| CD4+YFP- DKO3 | TGCACCTGCAGTGCGACTAGTGCAGAAACGCTGTATTTT | 1 | TRBV1*00 | | TRBJ2-3*00 | 0.52% |
| CD4+YFP- DKO3 | TGCACCTGCAGTGGGACTGCCTCCTATGAACAGTACTTC | 1 | TRBV1*00 | TRBD2*00 | TRBJ2-7*00 | 0.52% |
| CD4+YFP- DKO3 | TGTGCCAGCAGTTTCCCAGAAGAGACACCCAGTACTTT | 1 | TRBV15*00 | TRBD2*00 | TRBJ2-5*00 | 0.52% |
| CD4+YFP- DKO3 | TGTGCCTGGAGTCAACTGGGGCTATGAACAGTACTTC | 1 | TRBV31*00 | | TRBJ2-7*00 | 0.52% |
| CD4+YFP- DKO3 | TGTGCTAGCAGTCAATGGCATTATGAACAGTACTTC | 1 | TRBV21*00 | | TRBJ2-7*00 | 0.52% |
| CD4+YFP- DKO3 | TGTGCCTGGAGCCCCCCCTATGCTGAGCAGTTCTTC | 1 | TRBV31*00 | TRBD2*00 | TRBJ2-1*00 | 0.52% |
| CD4+YFP- DKO3 | TGTGCCAGCTCCCCTAGTGCAGAAACGCTGTATTTT | 1 | TRBV18*00 | | TRBJ2-3*00 | 0.52% |
| CD4+YFP- DKO3 | TGTGCCAGCGGGGGCAAAAACTCCGACTACACCTTC | 1 | TRBV13-2*00 | | TRBJ1-2*00 | 0.52% |
| CD4+YFP- DKO3 | TGTGCCAGCAGTTTGGGGGGAAACACCTTGTACTTT | 1 | TRBV15*00 | TRBD2*00 | TRBJ2-4*00 | 0.52% |
| CD4+YFP- DKO3 | TGTGCCAGCAGTGTCAGAGACGAAAGATTATTTTTC | 1 | TRBV13-3*00 | | TRBJ1-4*00 | 0.52% |
| CD4+YFP- DKO3 | TGTGCCAGCAGTATAGAACAGTATGAACAGTACTTC | 1 | TRBV19*00 | | TRBJ2-7*00 | 0.52% |
| CD4+YFP- DKO3 | TGTGCCAGCAGCTATAACCAAGACACCCAGTACTTT | 1 | TRBV4*00 | | TRBJ2-5*00 | 0.52% |
| CD4+YFP- DKO3 | TGTGCCAGCAGCCCGAGCTCCTATGAACAGTACTTC | 1 | TRBV3*00 | | TRBJ2-7*00 | 0.52% |
| CD4+YFP- DKO3 | TGTGCCAGCAGCCCATCCCAAGACACCCAGTACTTT | 1 | TRBV3*00 | | TRBJ2-5*00 | 0.52% |
| CD4+YFP- DKO3 | TGTGCCAGCAGCCAAGGGGATTCGCCCCTCTACTTT | 1 | TRBV5*00 | | TRBJ1-6*00 | 0.52% |
| CD4+YFP- DKO3 | TGTGCCAGCAGCAGAAACCAAGACACCCAGTACTTT | 1 | TRBV2*00 | | TRBJ2-5*00 | 0.52% |
| CD4+YFP- DKO3 | TGTGCAAGCAGCTTCACAAACCAGGCTCCGCTTTTT | 1 | TRBV16*00 | | TRBJ1-5*00 | 0.52% |
| CD4+YFP- DKO3 | TGTGCAAGCAGCTTACTGGGGGGGCGCAAGTACTTC | 1 | TRBV16*00 | TRBD2*00 | TRBJ2-7*00 | 0.52% |
| CD4+YFP- DKO3 | TGTGCAAGCAGCTTAGGAAAATATGAACAGTACTTC | 1 | TRBV16*00 | | TRBJ2-7*00 | 0.52% |
| CD4+YFP- DKO3 | TGTGGTGCTACGACAGGGGGCTCCGACTACACCTTC | 1 | TRBV20*00 | TRBD1*00 | TRBJ1-2*00 | 0.52% |
| CD4+YFP- DKO3 | TGTGGTGCTAGGGAGACCAACGAAAGATTATTTTTC | 1 | TRBV20*00 | | TRBJ1-4*00 | 0.52% |
| CD4+YFP- DKO3 | TGCACCTGCAGTGTCAACACCGGGCAGCTCTACTTT | 1 | TRBV1*00 | | TRBJ2-2*00 | 0.52% |
| CD4+YFP- DKO3 | TGTGCCAGCGGGAACCAAGACACCCAGTACTTT | 1 | TRBV13-2*00 | | TRBJ2-5*00 | 0.52% |
| CD4+YFP- DKO3 | TGTGCCAGCAGTTTTCAAAACACCTTGTACTTT | 1 | TRBV15*00 | | TRBJ2-4*00 | 0.52% |
| CD4+YFP- DKO3 | TGTGCCAGCAGCCGACAATATGAACAGTACTTC | 1 | TRBV3*00 | | TRBJ2-7*00 | 0.52% |
| CD4+YFP- DKO3 | TGTGCCAGCAGGAGGGGGTATGAACAGTACTTC | 1 | TRBV13-1*00 | TRBD1*00 | TRBJ2-7*00 | 0.52% |
| CD4+YFP- DKO3 | TGTGCCAGGACAGCAAACTCCGACTACACCTTC | 1 | TRBV31*00 | | TRBJ1-2*00 | 0.52% |
| CD4+YFP- DKO3 | TGTGGTGCTAGGGGTGACACAGAAGTCTTCTTT | 1 | TRBV20*00 | | TRBJ1-1*00 | 0.52% |
| CD4+YFP- DKO3 | TGTGCCAGCAGCAAATATGAACAGTACTTC | 1 | TRBV4*00 | | TRBJ2-7*00 | 0.52% |
| CD4+YFP- DKO3 | TGTGCCAGCCTTTGAACAGTACTTC | 1 | TRBV12-2*00 | | TRBJ2-7*00 | 0.52% |
